# From the Roundabout of Molecular Events to Nanomaterial-Induced Chronic Inflammation Prediction

**DOI:** 10.1101/2020.02.27.966036

**Authors:** Hana Majaron, Boštjan Kokot, Aleksandar Sebastijanović, Carola Voss, Rok Podlipec, Patrycja Zawilska, Trine Berthing, Carolina Ballester López, Pernille Høgh Danielsen, Claudia Contini, Mikhail Ivanov, Ana Krišelj, Petra Čotar, Qiaoxia Zhou, Jessica Ponti, Vadim Zhernovkov, Matthew Schneemilch, Zahra Doumandji, Mojca Pušnik, Polona Umek, Stane Pajk, Olivier Joubert, Otmar Schmid, Iztok Urbančič, Martin Irmler, Johannes Beckers, Vladimir Lobaskin, Sabina Halappanavar, Nick Quirke, Alexander P. Lyubartsev, Ulla Vogel, Tilen Koklič, Tobias Stoeger, Janez Štrancar

## Abstract

Nanomaterial-induced diseases cannot be reliably predicted because of the lack of clearly identified causal relationships, in particular between acute exposures and chronic symptoms. By applying advanced microscopies and omics to *in vitro* and *in vivo* systems, together with *in silico* molecular modelling, we have here determined that the long-lasting response to a single exposure originates in the counteracting of a newly discovered nanomaterial quarantining and nanomaterial cycling among different lung cell types. This allows us to predict the nanomaterial-induced spectrum of lung inflammation using only *in vitro* measurements and *in silico* modelling. Besides its profound implications for cost-efficient animal-free predictive toxicology, our work also paves the way to a better mechanistic understanding of nanomaterial-induced cancer, fibrosis, and other chronic diseases.

## Main Text

Chronic diseases such as asthma, lung cancer, heart disease, and brain damage with accelerated cognitive decline, are considered to be major causes of death, ^[1–3]^ and are known to be associated with air pollution and the inhalation of particulate matter and nanoparticles. ^[4]^

According to the OECD and WHO, they kill four million people globally every year. ^[5,6]^ Ever-increasing exposure to nanomaterials, a consequence of the rapidly developing nanotechnology industry, is causing concern because of the threat to people’s health. Despite advances in targeted test assays ^[7]^ and QSAR ^[8,9]^ models for nanotoxicology, neither *in vitro* nor *in silico* tools can reliably predict *in vivo* adverse outcomes. ^[10,11]^ This is particularly the case with the systemic and chronic adverse effects that are associated with the pathological changes that evolve in organs and tissues over long periods.

Because we do not understand how these problems evolve, decision-makers around the world (OECD, US EPA, NIH, EC, JRC, etc.) have highlighted the need to explain the molecular mechanisms involved, in particular using adverse-outcome pathways (AOPs). ^[12]^ These are now seen as the most promising construct to predict the apical endpoints, based on detecting the key events in the toxicity pathways. To this end, the community is actively searching for *in vitro* systems as an inexpensive, predictive ^[13,14]^, high-throughput alternative to conventional testing strategies ^[15]^ that would be capable of reproducing the nanomaterial’s *in vivo* mechanism of action, especially the single-exposure-initiated chronic inflammation ^[16–21]^ with the co-observed dysregulated lipid metabolism. ^[22–27]^

Here we demonstrate, for the first time, the prediction of nanomaterial-induced acute and chronic inflammation for selected nanomaterials, such as quartz and titanium dioxide. We employed strategically selected *in vitro* and *in silico* tests based on here-discovered molecular events and their causal relationships: 1) nanomaterial surface quarantining in lipid- nanomaterial composites, enabled by an upregulated lipid metabolism, and 2) nanomaterial cycling between different lung cell types, fueled by a lipid-metabolism-associated influx of new immune cells.

### Quarantining of Nanomaterials

To uncover the causal relationships between events leading from pulmonary nanomaterial exposure to chronic inflammation, we applied a complex set of complementary *in vivo, in vitro* and *in silico* experiments employing state-of-the-art microscopy, spectroscopy, omics and modelling approaches. TiO_2_ nanotubes were selected as the model material because they induce very high and long-lasting chronic inflammatory responses *in vivo* (Supplementary Information (SI) section S2d). With cells *in vitro* remaining viable for longer period after exposure, this nanomaterial allows us to study in detail the mechanism of the inflammatory response. Importantly, this nanomaterial also induces similar bio-nano composites on the surface of the epithelial lung cells both *in vitro* (Figure 1C) and *in vivo* (Figure 1B, violet structures). Note that similar structures were also observed after exposure to crystalline quartz (DQ12) (Figure 1C, SI section S1c), a well-known occupational hazard causing chronic inflammation. ^[16]^

**Figure 1.**
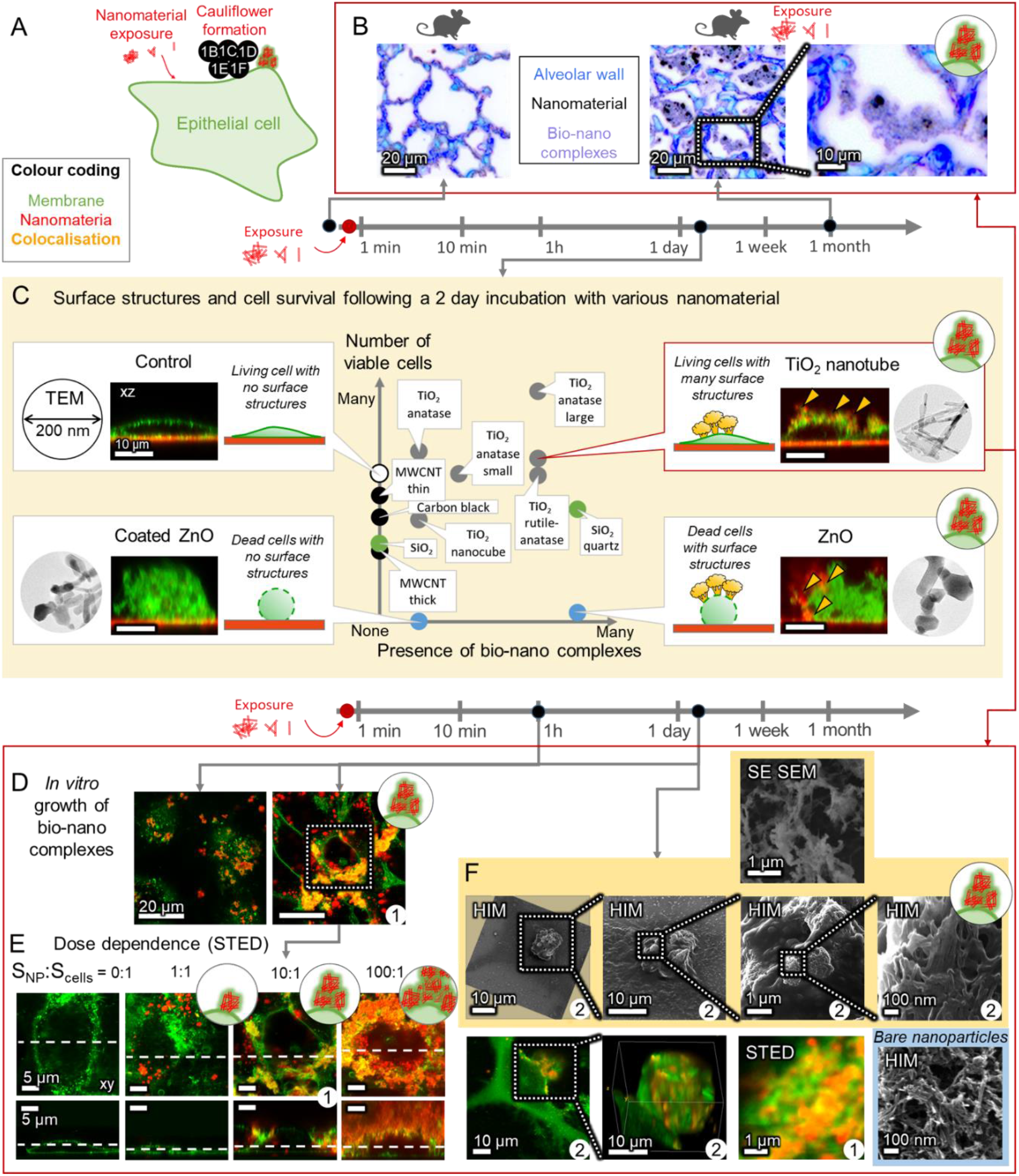
Formation of bio-nano composites on epithelial cell surface, referred to as cauliflowers. (**A**) A general scheme of events shown in this figure. (**B**) Hyperspectral-color-inverted dark-field micrographs of TiO_2_ nanotubes (black) in bio-nano composites (violet) observed in alveoli (blue) 1 month after instillation of the nanomaterial in mice. In fluorescence micrographs of *in vitro* alveolar epithelial (LA-4) cells (**C**-**F**) membranes are shown in green and nanoparticles in red. Images with the same number in the lower-right corner are images of the same cell. (**C**) Presence of cauliflowers, cell survival and xz cross-sections after a 2-day exposure to several nanomaterials at a nanomaterial-to-cell surface ratio (S_NP_:S_cells_) of 10:1 (nanoparticles observed in backscatter). Insets show 200-nm-large TEM micrographs of nanoparticles used. (**D**) Time-dependent cauliflower formation by LA-4 exposed to TiO_2_ nanotubes at S_NP_:S_cells_ = 10:1. (**E**) Super-resolved STED xy and xz cross-sections of dose-dependent cauliflower growth reveal that cauliflowers are located on the outer surface of cells after 2 days. S_NP_:S_cells_ are 0:1, 1:1, 10:1 and 100:1. (**F**) link to 3D: 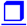 High-resolution correlativeSTED, SE SEM and HIM images reveal the detailed structures of cauliflowers at a S_NP_:S_cells_ = 10:1. For associated data see section S1in Supplementary Information (SI).

Based on previously observed strong interactions between TiO_2_ nanotubes and epithelial plasma membranes ^[28]^ we would expect that at higher surface doses (surface-of-nanomaterial- to-cell-surface dose 10:1 or more) these nanoparticles to completely disrupt the epithelial cell membranes. Surprisingly, our experiments show that the epithelial cells survive exposure to surface doses as high as 100:1 (Figure 1E, SI sections S0e and S0f). A few days after the exposure, the majority of the nanoparticles are found in large bio-nano composites on the epithelial surface, consisting of at least nanoparticles and lipids, which we term cauliflowers because of their shapes in our fluorescence micrographs (Figure 1D, Figure 1E, yellow color, Figure 1F).

Because the cauliflowers need several days to form and contain excessive amount of lipids, we next explore changes to the membrane structures and the lipid metabolism in an active biological response to the nanomaterial exposure.

### The Role of Lipids

Coinciding with the formation of the lipid-rich bio-nano composites (Figure 2B), i.e., two days after the nanomaterial exposure, a strong upregulation of lipid metabolism-related genes is observed (Figure 2D). Moreover, further modulation of the lipid synthesis pathway by blocking its key enzyme, fatty acid synthase (FAS), with resveratrol precludes the formation of large cauliflowers (Figure 2I), confirming that the epithelial cells actively respond to the nanomaterial exposure with increased lipid synthesis. The additionally synthesized lipids are used to immobilize and quarantine the nanoparticle surface, rendering such composites more loosely packed compared to the agglomerates of pure nanoparticles, as demonstrated by an increased fluorescence lifetime (Figure 2C). This process enables the layered growth of the cauliflowers over time, hindering any further interaction and lowering the active dose of the nanomaterial.

**Figure 2.**
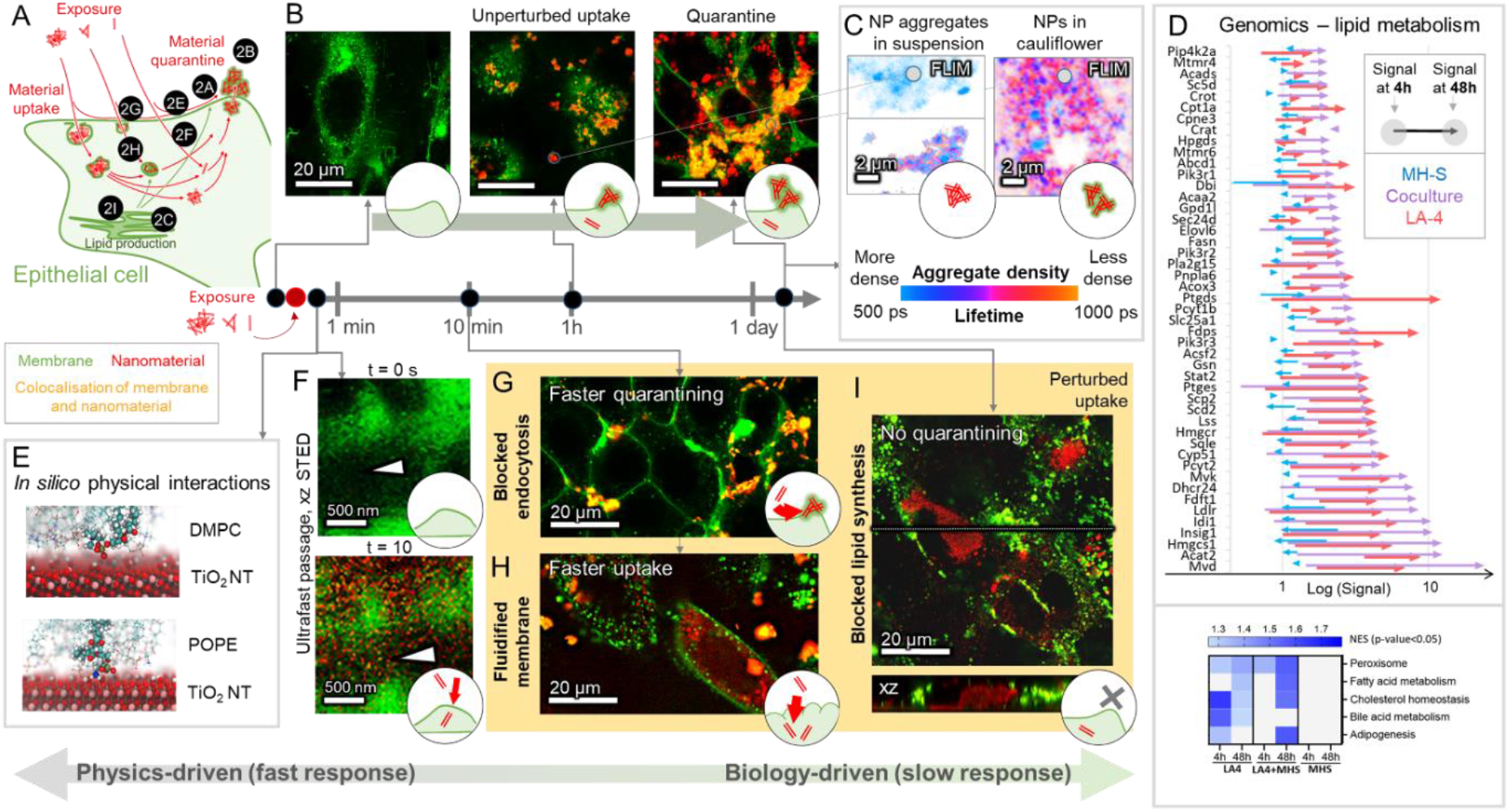
Role of lipids in cauliflower formation. (**A**) General scheme of events. In fluorescence micrographs *in vitro*, cell membranes are displayed in green and TiO_2_ nanotubes in red, surface dose was 10:1 (except **F**). (**B**) Unperturbed uptake of TiO_2_ nanotubes after 0, 1 h and 2 days by epithelial LA-4 lung cells, same as Figure 1D. (**C**) Increased fluorescence lifetime (FLIM) of fluorophore on TiO_2_ nanotubes in cauliflowers (right) compared to passively formed nanomaterial agglomerates in suspension (left) corresponds to increased distance between fluorophores on the nanotubes (e.g., separation due to lipid interspacing). (**D**) Transcriptional signature of lipid metabolism genes (top) and hallmark gene sets (bottom) for MH-S macrophages (blue), LA-4 epithelial cells (red) and their co-culture (purple) after 4 hours (beginning of arrow) and 48 hours (end of arrow) of nanomaterial exposure (NES). (**E**) Final state of full-atom *in silico* simulation confirms strong interaction between disordered lipids and the TiO_2_ nanotubes (DMPC links to movie and 3D: 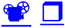, POPE 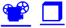). (**F**) 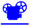 xz cross-sections immediately before (above) and 10 s after (below) instant delivery of TiO_2_ nanotubes onto cells by nebulisation (1:1 surface dose) show ultrafast membrane passage of the nanotubes through the cell plasma membrane into the cell (arrowhead). Drug-perturbed uptakes (to compare with **B**): (**G**) 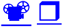 chlorpromazine-blocked clathrin-mediated endocytosis, (**H**) 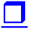 fluidified cell plasma membrane induced by cholesterol depletion (beta-methyl- cyclodextrin) (**I**) 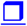 inhibited fatty acid synthesis (resveratrol-blocked fatty-acid synthase). For associated data see SI section S2.

As the internalization of the nanoparticles typically precedes the cauliflower formation (SI section S2b), we investigate the causal relationship between the two phenomena by blocking an important route of nanoparticle uptake, i.e., clathrin-mediated endocytosis (SI section S0d), using chlorpromazine. Interestingly, small “proto” cauliflowers are formed soon after exposure (15 min time scale) (Figure 2G), indicating an additional mechanism of formation that requires no intracellular processing. In this case the formation of cauliflowers presumably relies on the strong physical affinity between the nanoparticles and the lipids, which is supported by the *in silico* simulations (Figure 2E) and the *in vitro* experiments on model lipid membranes (SI section S0c). However, these “proto” cauliflowers are rarely seen under normal conditions, which leads us to conclude that this additional mechanism of formation is usually less likely, possibly due to the efficient particle uptake that displaces the nanomaterials away from the plasma membrane, preventing their further interaction.

Interestingly, the depletion of cholesterol as the major membrane constituent by beta-methyl- cyclodextrin, which increases the fluidity of the plasma membrane, leads to the strong suppression of rapid (membrane-lipid-drain only) cauliflower formation (Figure 2H). This indicates an important interaction between the nanoparticles and the cholesterol, which is reflected in the strongly upregulated cholesterol and lipid synthesis pathways in the epithelial cells *in vitro* (Figure 2D heatmap, SI section S2d), as well as in mouse lungs *in vivo* (SI section S2d). In the case of cholesterol-depleted plasma membranes, the majority of the nanoparticles cross the plasma membranes on a timescale of minutes, resulting in a fine distribution of the particles inside the cell. The dominant role of such a passage can also be observed when the nanoparticles are delivered in a highly dispersed form through an aerosol, directly to the epithelial plasma cell membranes and pass through them ^[29]^ in a matter of seconds (Figure 2F, link to movie: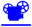).

For the alveolar barrier of the lung in particular, the lipid-synthesis-driven formation of bio- nano composites appears to be an important part of the active response of alveolar epithelial cells, enabling their survival after exposure to the nanomaterial, even at higher doses (SI sections S0e and S0g). As we consistently observed the quarantining of the nanomaterials on the cell surface that follows the nanomaterial internalization, we further explore the cellular mechanisms that ^[29]^facilitate the export of the internalized material.

### The Role of Actin

As exocytosis involves cytoskeletal actin remodeling, we examined the role of actin in the process. Almost simultaneously with the nanoparticle uptake and well before the cauliflowers form, many nanoparticles interact with actin fibers (Figure 3D), forming nanoparticle-actin 3D composites resembling Faberge eggs (Figure 3B). Hours after exposure, the same interaction causes actin network transformations from linear aligned to branched fibers (Figure 3E), which is associated with an increased cell motility ^[30]^ as well as with internal vesicular trafficking ^[31,32]^ and exocytosis. ^[33,34]^

**Figure 3.**
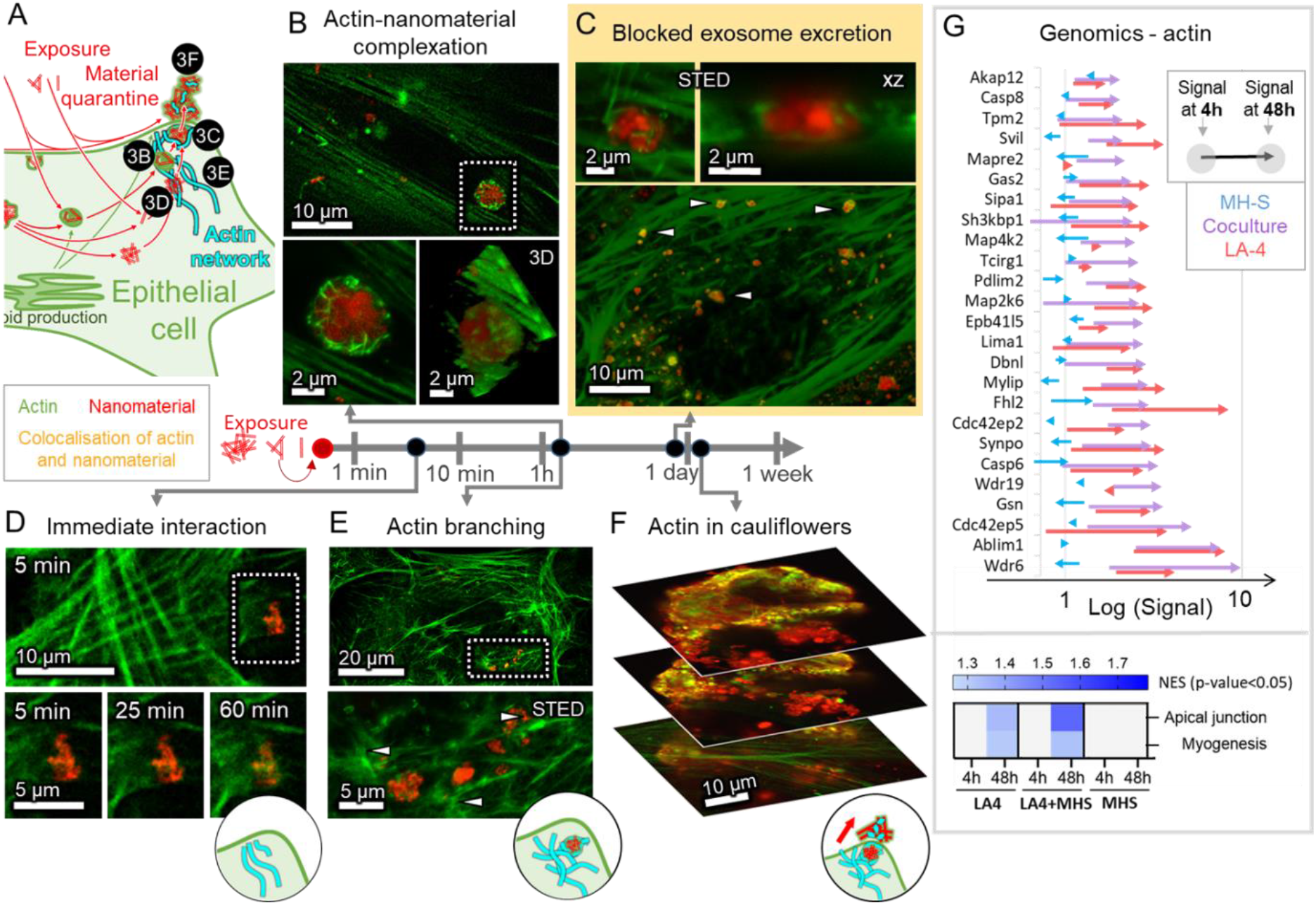
Role of actin in cauliflower formation. (**A**) General scheme of events. Fluorescence micrographs of the actin network of LA-4 cells (green) after exposure to TiO_2_ nanotubes (red) at a 10:1 surface dose. (**D**) 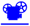 Soon after exposure, actin interacts with internalized nanoparticles, (**B**) 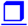 leading to formation of actin-nanoparticle composites after a few hours. (**E**) Synchronously, the actin network branches (arrowheads), indicating changes in internal processes and reshaping of the cell. (**C**) 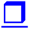 Blocking the final stage of exocytosis with jasplakinolide traps nanoparticles in actin rings, prepared for exocytosis (arrowheads and zoom-ins). (**F**) 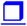 After a few days, actin fragments are observed in cauliflowers (arrowheads). (**G**) Transcriptional signature of actin-network-related genes (top) and hallmark gene sets (bottom) for LA-4 (red), macrophages (blue), and their co-cultures (purple) after 4 hours (beginning of arrow) and 48 hours (end of arrow) of nanomaterial exposure. For associated data see SI section S3.

By blocking the actin fiber dynamics (polymerization and depolymerization) with jasplakinolide, the excretion of exocytotic vesicles can be stopped, thereby enabling the simultaneous visualization and identification of the nanoparticles trapped in the exocytotic vesicles (actin rings) (Figure 3C). As actin can be identified extracellularly within the cauliflowers (Figure 3F, link to 3D: 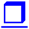), the excretion of nanoparticles is apparently more destructive to the actin network than normal homeostatic exocytosis, where actin is retained inside the cells. Actin adherence to the nanomaterial is also reflected in the coronome analysis of the mobile fraction of nanoparticles after the exposure, in which we have previously detected an abundant fraction of actin proteins, but were unable to explain it. ^[28]^ The loss and destruction of actin also triggers the upregulation of the pathways related to actin synthesis (Figure 3G).

The creation of cauliflowers on the cell surface thus involves both membrane lipids and actin (heatmaps in Figure 2D and Figure 3C) that directly interact with the nanoparticle surface. Due to the strong binding of the amines and phosphates identified by *in silico* simulations (Figure 2E), it is reasonable to expect that various biomolecules, including lipids, proteins and nucleic acids, strongly bind to the same particle surface. Moreover, multiple binding sites on the nanomaterial and biomolecules or their supramolecular structures lead directly to crosslinking and the formation of large bio-nano composites, such as the observed cauliflowers. This implies that a lack of strong interaction identified within the *in silico* modelling of biomolecule- nanomaterial surface pairs, is predictive of the absence of bio-nano composite formation, and can be used in safety prediction.

The ability of the alveolar epithelium to supply enough biomolecules to crosslink and thereby quarantine the received dose of nanomaterial explains the cell survival, even for relatively large local doses of nanomaterials, which can also be observed *in vivo* (Figure 1). The process of nanomaterial quarantining, however, seems to contradict the observation of simultaneous chronic pulmonary inflammation, raising the question of the role of the neighboring cells, especially the alveolar macrophages, which are responsible for the alveolar immune defense and thereby the alveolar integrity. To address this, we expose a co-culture of LA-4 epithelial cells and MH-S alveolar macrophages in the same way as we did with the epithelial monoculture.

### Macrophage Action Against Epithelial Defense

With a co-culture of MH-S alveolar macrophages on top of nearly confluent LA-4 epithelial cells we aimed to mimic the cell populations of the alveolar surface, where alveolar macrophages represent approximately 3–5% of all the alveolar cells. ^[35]^ Upon exposure of the co-culture to TiO_2_ nanotubes, part of the material becomes internalized by the phagocytes, which cannot entirely prevent the nanomaterial from reaching the epithelial cells (SI section S0h 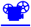), which is in line with previous *in vivo* observations. ^[36]^ Aside from that, the macrophages unexpectedly slow down considerably, after having taken up large amounts of nanoparticles (graph in SI section S0h), making their clearance function even less efficient. This explains why the exposed epithelium also produces cauliflowers in our co-culture (SI section S0h), reproducing the bio-nano composites observed *in vivo* in the alveolar region of the lungs of particle-exposed mice (Figure 1B).

Although the nanoparticles are quarantined in cauliflowers on the surface of the LA-4 cells, enabling their survival, the same structures trigger the attack of macrophages, as seen in the experiment when unexposed macrophages were added to pre-exposed and therefore cauliflower-rich epithelium (Figure 4A). After internalization of the cauliflowers, macrophages are able to degrade only their organic part, as revealed by the decreased lifetime of the fluorescent probes bound to the nanoparticles, indicating denser packing of the nanoparticles in macrophages compared to the cauliflowers (FLIM maps in Figure 4A insets). Freeing the nanoparticles exposes the macrophage interior to the bare nanoparticle surfaces, leading to macrophage death and subsequent disintegration, as observed in monoculture (Figure 4D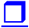), possibly caused by the lack of exocytosis and suppressed normally elevated lipid synthesis signature (Figure 2C). A similar macrophage fate is also observed after they have attacked the epithelial cells (Figure 4E 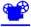) or other macrophages with internalized nanomaterials (Figure 4F 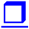). When nanomaterial-exposed macrophages die, they release bare nanomaterial, which is later taken up again by the epithelial cells. This can be observed experimentally: after nanomaterial-laden macrophages were added to the unexposed epithelial layer, nanoparticles are localized inside the epithelial cells (Figure 4B).

**Figure 4.**
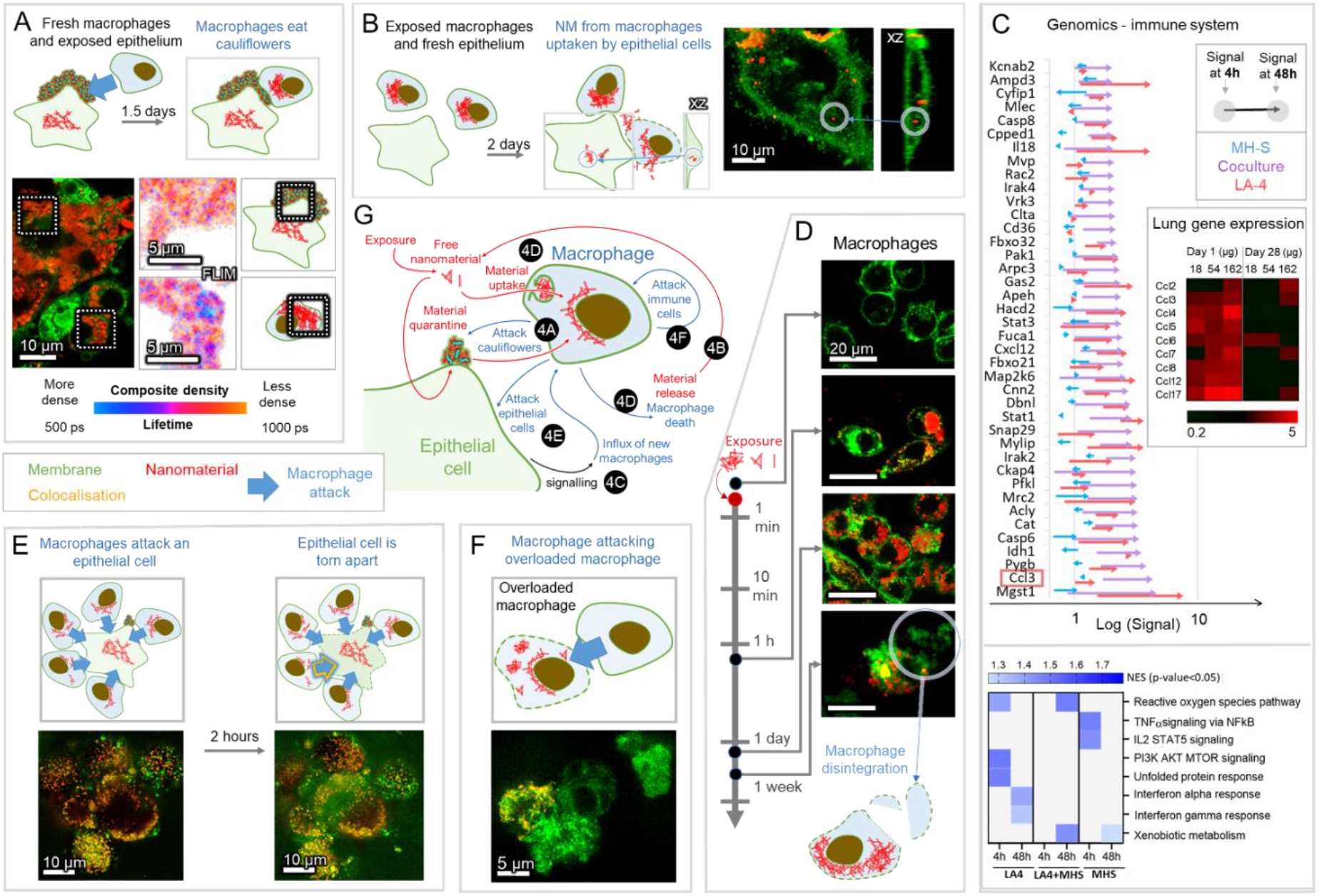
The cycle of uptake, quarantine and release in nanomaterial-exposed co-culture. In all fluorescence micrographs, cell membranes are displayed in green and TiO_2_ nanotubes in red, and the surface dose of nanoparticles is 10:1. (**A**) Unexposed macrophages (MH-S) were added to washed LA-4 cells with cauliflowers. Within 1.5 days, MH-S phagocyte the cauliflowers from the LA-4 cell surface, and degrade their organic (lipid) part, thereby compacting the nanoparticles (fluorescence-lifetime-maps FLIM, right). (**B**) Washed nanomaterial-laden MH-S were added to unexposed LA-4. After 2 days, the nanomaterial is found in LA-4 cells (encircled). (**C**) Transcriptional signature of genes related to the immune response (top) and hallmark gene sets (bottom) for LA-4 (red), MH-S (blue) and their co-culture (purple) after 4 hours (beginning of arrow) and 48 hours (end of arrow) of nanomaterial exposure, with lung gene expression of some CCL monocyte attractants after 1 and 28 days. **(D)** Nanoparticle uptake by MH-S followed by their disintegration after a few days (encircled): 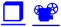 (control) 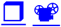 (2 h) 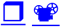 (2 days) 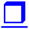 (4 days, MH-S disintegration) (**E**) 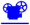 Time lapse of MH-S attacking and tearing apart a nanomaterial-laden LA-4 cell. (**F**) 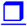 Observation of MH- S attacking another nanomaterial-laden macrophage. (**G**) A general scheme of events shown in this figure. For associated data see SI section S4.

This reuptake in turn, again leads to quarantined nanomaterial on the self-protected epithelial cells. *In vivo*, however, dead macrophages are replaced through an influx of new monocyte-derived macrophages, attracted to the site by respective macrophage/monocyte chemo- attractants such as the C-C motif ligand 3 (Ccl3)) from the epithelial cells, or Ccl2-17 for the lungs of nanomaterial-exposed mice (SI section S2d). This macrophage replenishment brings the entire system to conditions very similar to the initial exposure, while the reuptake of the nanomaterial by the epithelium finally closes the chain of events, together forming a vicious cycle, generating a never-before-seen loop of persistent inflammation (Figure 4G, Figure 5A). Strikingly, the same chemokine expressions can be detected both *in vivo* (Figure 4C inset) and *in vitro* in the co-culture of LA-4 and MH-S cells (Figure 4C, purple arrows), but not in the monocultures of LA-4 (Figure 4C, red arrows) nor of MH-S (Figure 4C, blue arrows). This pro-inflammatory signaling represents the last missing piece of evidence that the *in vitro* co-culture can reproduce the entire cycle of the chronic-inflammation-initiating mechanism (black arrow in Figure 4G). Can we thus predict such an *in vivo* chronic inflammation response by measuring the specific states of simple *in vitro* tests?

**Figure 5.**
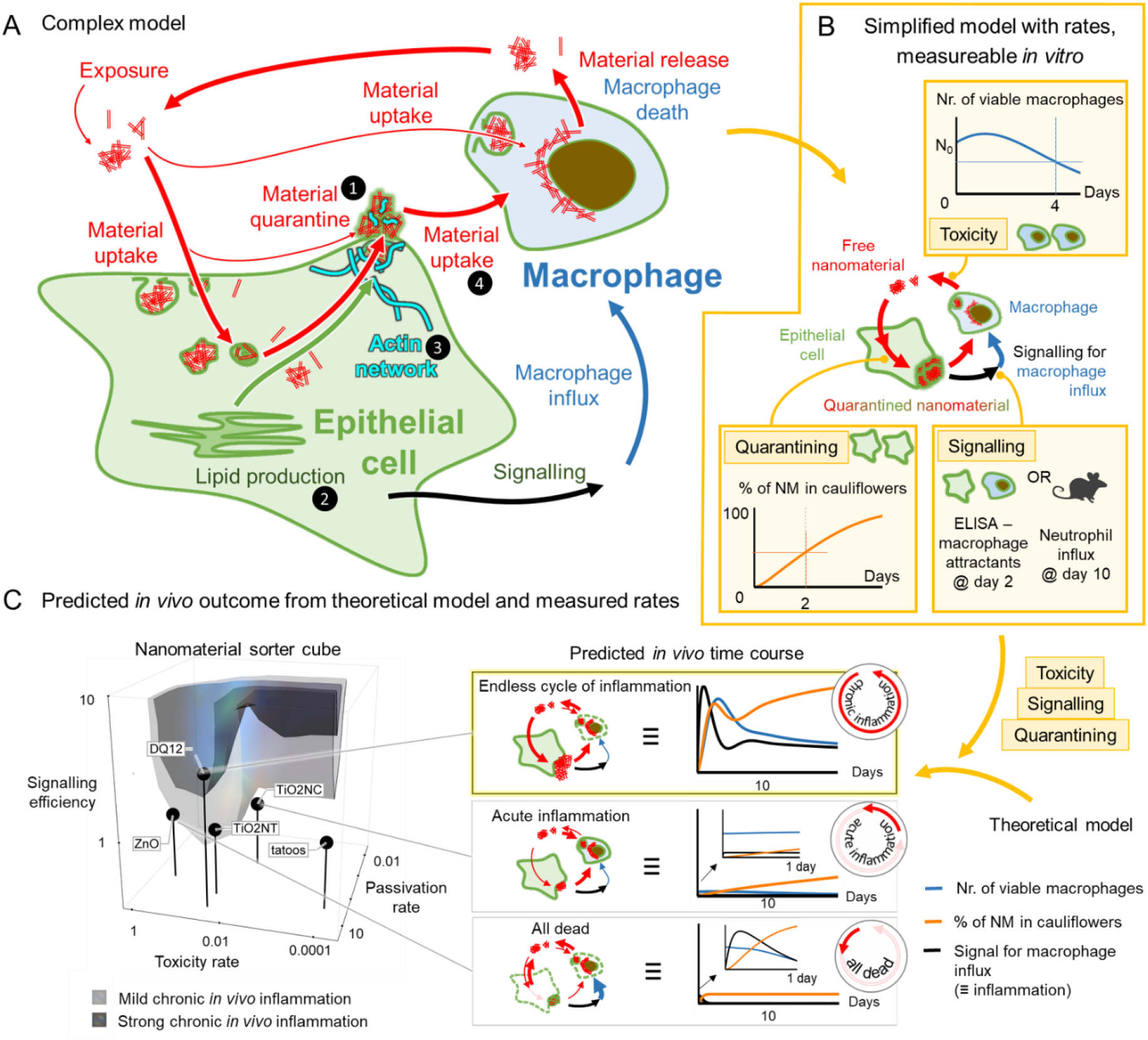
Cycle of uptake, quarantine and release of nanomaterial between epithelial cells and macrophages in co-cultures. (A) The grand scheme connecting all inter- and intracellular events from Figures 1–4, simplified to (B) theoretical model, defined by the rates of cauliflower formation, nanomaterial toxicity and signaling efficiency. These nanomaterial descriptors can be determined from single time-point measurements *in vitro* and/or *in vivo*. (C) The model is evaluated using these determined parameters, producing *in vivo* time courses (right) of the relative amount of nanomaterial in cauliflowers (orange), the number of viable macrophages (blue) and the signal for their influx (black). The value of the latter at day 10, when the acute phase is expected to subside, is contoured in the 3D space of the aforementioned rates (cube on the left). Nanomaterials are placed in the same 3D cube according to their measured descriptors, enabling the prediction of the degree of nanoparticle-induced chronic inflammation. For associated data see SI section S5.

### Acute or Chronic? The Birth of Predictive Tools

The proposed pathway (Figure 5A), connecting an acute nanoparticle exposure to a chronic inflammation via a chain of causally related events, allows us to construct a simplified cyclical theoretical model that describes the flow of the nanomaterial between four separate compartments (outside the cells, inside the epithelial cells, quarantined in cauliflowers, and in macrophages). This model is defined with three key descriptors (SI section S5b, depicted in Figure 5B), measurable in appropriate *in vitro* assays for any nanomaterial of interest (yellow shaded boxes in Figure 5B):

- The rate of toxicity of the nanomaterials to individual cells *(tox)* is determined by the measured number of viable macrophages in the MH-S monoculture after 4 days (Figure 5B, toxicity);
- The rate of nanomaterial quarantine by epithelial cells *(cff)* is calculated from the rate *tox* and the measured fraction of nanomaterial in the cauliflowers in the LA-4 monoculture after 2 days (Figure 5B, quarantining);
- The efficiency of the signaling and the monocyte influx replacing the dying macrophages *(signalEff)* is calculated from the rates *tox, cff* and either via the measured macrophage attractants in the *in vitro* co-culture of LA-4 and MH-S after 2 days or via the measured influx of inflammatory cells (polymorphonuclear leukocyte) *in vivo* after at least 10 days (Figure 5B, signalling), a time point where the chronification of the response is secured.

Whether the cycle stops or continues indefinitely depends heavily on the rates of the associated processes, calculated from the measured descriptors as described in SI section S5b. Using these rates, the model can predict the *in vivo* time course of the amount of quarantined nanomaterial in the cauliflowers, signaling for macrophage influx, as well as of the total macrophage number, and accordingly predict the nanomaterial-specific acute-to-chronic inflammation outcome (Figure 5C - time traces). For example, for a very toxic nanomaterial such as ZnO, the model yields a rapid decline in the number of cells, preventing quarantine of the nanomaterial and resulting in the destruction of the alveolar layer, which is consistent with the *in vivo* observations. ^[37]^ For a material with an intermediate toxicity and quarantine rate, e.g., TiO_2_ nanocubes, the model predicts weak transient inflammation, with all the nanomaterial ending up in cells, as observed *in vivo*. ^[16]^ Finally, for a material such as TiO_2_ nanotubes or DQ12 with intermediate toxicity and a high quarantine rate, persistently high inflammation and large cauliflowers (Figure 5C – time traces) are predicted, reproducing the *in vivo* observations (Figure 1B). In this 3D space of nanomaterial descriptors (Figure 5C – 3D plot) we can now delineate regions eliciting a similar outcome, thus sorting nanomaterials according to their mode of action.

### Conclusions and Perspectives

The here described lipid-mediated nanomaterial quarantine, a previously unknown epithelial defense mechanism, prevents an acute epithelium damage upon nanomaterial exposure and delays the immune response. The resulting continuous cycling of nanomaterial between different cell types leads to a vicious cycle of molecular events, clarifying the associated adverse outcome pathway and explaining chronic inflammation. Such nonlinear initiation of an adverse outcome pathway could inspire future research towards a mechanistic understanding and possible treatment strategies of the endless adverse cycles in nanomaterial-induced cancer, fibrosis, and other chronic diseases.

The unraveled pathway allowed us to predict the *in vivo* outcome, being either acute inflammation or a long-term chronic one, by using only strategically selected *in vitro* measurements and *in silico* modeling, applicable to any nanomaterial, regardless of its specific properties, such as shape, size, charge, surface functionalization, etc. Based on this, we contend that the game-changing screening strategy in nanotoxicology should be based on understanding the response of the organism to nanomaterial exposure from the initial contact with the nanomaterial to the potential adverse outcome. Although this requires the use of advanced imaging, omics, particle labelling and tracking techniques at the stage of analyzing the *in vivo* and *in vitro* data, it could lead to novel, cost-efficient, high-throughput, alternative-to-animal testing strategies.

## Experimental Section

This is a condensed description of the methods. Details are available in the Supplementary Information in the general section “S0a – general materials and methods” for general methods as well as for each experiment separately.

### Materials

Alexa Fluor 647 NHS ester (Termo Fisher), Star 520 SXP NHS ester (Abberior), ATTO 594 NHS ester (Atto-tec), CellMask Orange (Invitrogen), SiR Actin (Cytoskeleton), Star Red-DPPE (Abberior), 4-(8,9-Dimethyl-6,8-dinonyl-2-oxo-8,9-dihydro-2H-pyrano[3,2- g]quinolin-3-yl)-1-(3-(trimethylammonio) propyl)pyridin-1-ium dibromide(SHE-2N), 3- (Benzo[d]thiazol-2-yl)-6,8,8,9-tetramethyl-2-oxo-8,9-dihydro-2H-pyrano[3,2-g]quinoline-4- carbonitrile (SAG-38), LCIS-Live Cell Imaging Solution (Invitrogen), PBS-phosphate buffer saline (Gibco), 100x dcb: 100-times diluted bicarbonate buffer (pH 10, osmolarity 5 miliosmolar, mixed in-house), F-12K cell culture medium (Gibco), RPMI 1640 cell culture medium (Gibco), Trypsin (Sigma), Penicillin-Streptomycin (Sigma), Non-essential amino acids (Gibco), Beta mercaptoethanol (Gibco), glucose (Kemika), BSA-bovine serum albumin (Sigma), Hydrogen peroxide (Merck), Chlorpromazine (Alfa Aesar), MBCD-Metyl-Beta-Cyclodextran (Acros organics), Resveratrol (Sigma), #1.5H µ-dishes (Ibidi,) #1.5H µ-Slide 8- well (Ibidi), Limulus Amebocyte Lysate Assay (Lonza, Walkersville, MD, USA), 10% neutral buffered formalin (CellPath Ltd, UK), haematoxylin and eosin (H&E), Pelcotec™ SFG12 Finder Grid Substrate- Si wafers (Ted Pella), Aeroneb®Pro nebulizer (from VITROCELL® Cloud 6 system), GeneChip® WT PLUS Reagent Kit (Thermo Fisher/Affymetrix), RNeasy Plus Mini Kit (Qiagen), WT PLUS Reagent Kit (Thermo Fisher Scientific Inc., Waltham, USA), Mouse Clariom S arrays (Thermo Fisher Scientific)

Nanomaterials used in this study

Synthesized in-house by P. Umek:

TiO_2_ nanotubes (PU-nTOX-01-03) and TiO_2_ nanocubes (PU-nTOX-01-21);

Kind gift from U. Vogel:

carbon black (Printex 90), TiO_2_ MKNA015 (MKN- TiO_2_ -A015), TiO_2_ MKNA100 (MKN- TiO_2_ -A100) and quartz silica (SiO_2_ DQ12);

Kind gift from JRC Nanomaterial Repository:

NM-101 TiO2 anatase (TiO2-NM101-JRCNM01001a), NM-105 TiO2 rutil-anatase (TiO2- NM105-JRCNM01005a), NM-110 ZnO (ZnO-NM110-JRCNM62101a), and NM 111 ZnO (ZnO-NM111-JRCNM01101a), NM-200 SiO2 (SiO2-NM200-JRCNM02000a), NM-401 MWCNT (MWCNTs-NM401-JRCNM04001a), NM-402 MWCNT (MWCNTs-NM402- JRCNM04002a).

### Software

Imspector (version 16.2.8282-metadata-win64-BASE) software provided by Abberior SPCImage 7.3 (Becker & Hickl)

Fiji, ImageJ 1.52p (NIH)

syGlass (http://www.syglass.io/, RRID:SCR_017961)

Mathematica 12.0, licence L5063-5112 (Wolfram)

genomics software: GSEA by Broad Institute

modelling: GROMACS 2018.3 (calculation), VMD (visualisation)

TiO_2_ nanotubes synthesis and labelling

The TiO_2_ anatase nanotubes used in this paper were synthesized, functionalized with AEAPMS, and labelled with STED-compatible fluorescent probes via a covalent reaction between the AEAPMS and ester functional group on the probe. All this was done in-house as described in reference. ^27^ Labelled TiO_2_ was then stored suspended in 100x diluted bicarbonate buffer. For the multi-nanomaterial exposure experiments we used other NMs as well. In this case, the nanomaterials were suspended in PBS and sonicated in ice bath using a tip sonicator (Sonicator 4000, Misonix, with 419 Microtip probe) for 15 min with 5s ON/ 5s OFF steps.

### Cell culture

Murine epithelial lung tissue cell line (LA- 4; cat. no. ATCC CCL-196) and murine alveolar lung macrophage (MH-S; cat. No. CRL2019) cell line were purchased from and cultured according to American Type Culture Collection (ATCC) instructions. Cells were cultured in TPP cell culture flasks at 37 °C in a 5% CO_2_ humidified atmosphere until monolayers reached desired confluency. All experiments were performed with cells before the twentieth passage. For long–term live cell experiments we used a homemade stage-top incubator which maintains a humidified atmosphere with a 5% CO_2_ and is heated to 37 °C.

Medium used for culturing of the epithelial LA-4 cells is Ham’s F-12K medium (Gibco) supplemented with 15% FCS (ATCC), 1% P/S (Sigma), 1% NEAA (Gibco), 2 mM L-gln.

For alveolar macrophages, MH-S, cell line we used RPMI 1640 (Gibco) medium supplemented with 10% FCS (ATCC), 1% P/S (Sigma), 2 mM L-gln, and 0.05 mM beta mercapthoethanol (Gibco).

### *In vitro* sample preparation and exposure

LA-4 and MH-S cells were seeded in Ibidi 1.5H dishes of various surface area, depending on the experiment. After 24 h, nanomaterial (c=1mg/mL) was added at an appropriate surface dose (S_NP_:S_cells_), according to the experiment needs. Before exposure, nanomaterial suspension was sonicated for 10 s in an ultrasonic bath (Bransonic ultrasonic cleaner, Branson 2510EMT). Cells were then incubated at 37 °C and 5% CO_2_ atmosphere with the nanomaterial for the desired time in order to observe the cells at the post-exposure time points of interest. If the experiment required monoculture of either cell line, sample were prepared as described above, if however, we experimented with the co-cultures, sample preparation differed slightly. For co- cultures, we grew LA-4 and MH-S in separate dishes up to desired confluency (lower than for monocultures) and then mixed them together by adding MH-S in the LA-4 dish at a ratio of 1: 40. Co-cultures were then incubated for 24 h more, exposed to nanomaterial as described above and incubated for additional desired amount of time. Growth medium for co-cultures was mixture of equal volumes of F12K and RPMI 1640. Cells were then labelled with fluorescent dyes according to the manufacturer’s recommendations. Right before observing the live cells, unbound fluorescent label was washed and medium was exchanged for LCIS.

In some experiments we used different chemicals for modulation of the cell metabolism. For blocking the Clathrin-mediated endocytosis, cells were treated with 100 μM Chlorpromazine for 15 min. Membrane cholesterol was extracted with a 24 h incubation with 0.5 - 1 mM MBCD. FAS was inhibited with overnight 100 μM Resveratrol incubation. Finally, for actin stabilization, we used higher concentration (≥1mM) of Sir-Actin Label based on Jasplankinolide. All the chemical modulators were added before exposure to nanomaterial and continued to be incubated with the cells even after during incubation with the nanomaterial for abovementioned time periods.

For the reuptake experiments different cell lines were grown separately, and washed with PBS before adding MH-S to LA-4.

#### HIM, SEM

Samples were prepared as usual but we grew them on Si-wafers. After reaching desired confluency samples were freeze-dried with metal mirror freezing technique.

Imaging *in vitro*

#### STED

Super-resolution and confocal fluorescence micrographs were acquired using custom build STED microscope from Abberior with an Olympus IX83 microscope and two avalanche photodiodes as detectors (APDs). The microscope is equipped with two 120 picosecond pulsed laser sources (Abberior) with excitation wavelengths 561 and 640 nm and maximal power of 50 µW in the sample plane. Pulse repetition frequency for experiments was 40 - 80 MHz, depending on the experiment. STED depletion laser wavelength is 775 nm with same repetition frequency as excitation lasers, pulse length of 1.2 ns and maximal power of 170 mW in the sample plane. Filter sets used for detection were either 585–625 nm (green channel) or 650– 720 nm (red channel). Images were acquired using Imspector (version 16.2.8282-win64) software also provided by Abberior. All microscope settings were tuned separately for maximal resolution during each of the experiments and are listed with alongside the recorded images in Supplementary Information.

#### FLIM

Fluorescence lifetime images (FLIM) were obtained on the same custom-built STED microscope (Abberior instruments) as confocal and STED fluorescence images in this study. This time, the emitted fluorescence was detected using PMT detectors and TCSPC technology developed by Becker & Hickl. 16-channel GaASP PMT detectors attached to a spectrograph with diffraction grating 600 l mm^−1^ were used to measure fluorescence lifetime of emitted photons with wavelengths ranging from 560 to 760 nm. Spectral information was discarded and the lifetimes were gathered in Imspector 16.2 (Abberior Instruments).

The fluorescence lifetime data was analyzed with SPCImage 7.3 software (Becker & Hickl), where the Decay matrix was calculated from the brightest pixel in the image (monoexponential fitting), binning was set to 3 and threshold to 5. The rainbow LUT was rescaled to range from 500 ps to 1000 ps for all images and both intensity and contrast of the lifetime-coded image were adjusted for easier comparison of lifetimes between samples.

### Imaging of nanomaterial in backscatter mode

In Figure 1c, simultaneously with measuring fluorescence from CellMask Orange in the cell membrane (as described in STED section), backscattered light was detected as well to locate the nanomaterial in the sample. A tunable Chameleon Discovery laser (Coherent) with 100 fs long pulses, pulse repetition frequency 80 MHz, and maximal average power of 1.7 W at 850 nm was used as the scattering light. The pre-attenuated laser light with a wavelength of 750 nm first passed through a 785 nm built-in dichroic where a fraction of the power was directed onto the sample through the same 60x WI objective (NA 1.2) as the excitation light for fluorescence imaging. The light scattered off the nanomaterial and passed back through the same objective and dichroic, now mostly passing through the dichroic towards the detectors. After passing through a pinhole (0.63 A.U.), the backscattered light was spectrally separated from the fluorescence by a short-pass 725 nm dichroic, afterwards being detected on the same PMT, as described in the FLIM section, this time set to collect light with wavelengths above 725nm.

Due to the large coherence of the laser, the backscattered light exhibited a strong speckle pattern, which was diminished by a 100-nm-wide Gaussian blur on the scattering image, thus decreasing false negative colocalisation of nanomaterial on account of spatial resolution.

#### SEM

SEM imaging has been performed on MIRA3 Flexible FE-SEM produced by TESCAN, by detection of secondary electrons. Beam powers used have been between 5.0 kV and 15 kV with variable field of view 1.8 μm to 180 μm. All samples have been measured under high pressure vacuum (HiVac). All analysis has been performed in Tescan developed software.

#### HIM

Super-resolution imaging on the nanoscale was carried out using Helium Ion Microscope (Orion NanoFab, Zeiss) available at IBC at the Helmholtz-Zentrum Dresden - Rossendorf e. V., a member of the Helmholtz Association. Microscope equipped with GFIS injection system and additional in-situ backscatter spectrometry and secondary ion mass spectrometry can achieve 0.5 nm lateral resolution imaging using 10-35 keV He ion beams. Measurements of secondary electrons (Se) emitted from the first few nm of the sample were done by He ion acceleration of 30 keV, current of 1.7 pA and were acquired under high vacuum inside the sample chamber (3×10-7 mBar). Field-of-view was varied from 60 μm x 60 μm down to 1 μm x 1 μm, with pixel steps small as 2 nm. Imaging was performed on non-tilted and tilted sample stage (45 degrees) for better 3-D visualization.

#### TEM

ZnO and coated ZnO: Of each material 1 mg was dispersed in 1 mL MilliQ water, except CNTs in 1 mL tannic acid solution 300 mg L^−1^, using a vial tweeter for 15 min. Each suspension was diluted 1/10 and 3 µL drop deposited on Formvar Carbon coated 200 mesh copper grids (Agar Scientific, USA) and dehydrated overnight in a desiccator before analysis. Images were collected by JEOL JEM-2100 HR-transmission electron microscope at 120kV (JEOL, Italy) at JRC. ^[38]^

TiO_2_ nanotubes: The nanoparticles were dispersed in water and the dispersion sonicated in water bath for ∼3h before use. Of each sample 5 µl was deposed onto glow-discharged copper grid (Agar scientific Ltd, UK) for one minute and the excess of sample was removed blotting with filter paper. After shortly washing with one drop of water, the grid was therefore immersed into a 2% uranyl acetate (UA) solution for 20 s and blotted again with filter paper. The grids were imaged using a JEOL JEM-2100F fitted with a Gatan Orius SC 1000 camera (2×4k).

### Transcriptomics *in vitro*

Cells were grown in 6-well plates and exposed to TiO_2_ nanotubes for 4 h and 48 h, control samples were taken at 0 h and 48 h. Samples were prepared as described above. Briefly, growth medium was removed and the 6-well plates containing cells only were frozen at −70°C. Total RNA was isolated employing the RNeasy Plus Mini Kit (Qiagen). The Agilent 2100 Bioanalyzer was used to assess RNA quality and RNA with RIN>7 was used for microarray analysis.

Total RNA (120 ng) was amplified using the WT PLUS Reagent Kit (Thermo Fisher Scientific Inc., Waltham, USA). Amplified cDNA was hybridized on Mouse Clariom S arrays (Thermo Fisher Scientific). Staining and scanning (GeneChip Scanner 3000 7G) was done according to manufacturer’s instructions.

Statistical analysis for all probe sets included limma t-test and Benjamini-Hochberg multiple testing correction. Raw p-values of the limma t-test were used to define sets of regulated genes (p<0.01). Detection Above Background (dabg) p-values were used to exclude background signals: significant genes were filtered for p<0.05 in more than half of the samples in at least one group. Array data has been submitted to the GEO database at NCBI (GSE146036).

In the arrow graphs, only genes which were up- or down-regulated more than two times compared to non-exposed cells are shown. The signal (x axis) is drawn in logarithmic scale. Expression is normalized to expression of control samples.

### *In vivo* data

#### Preparation and characterization of TiO_2_ nanotube suspensions

TiO_2_ nanotubes were suspended in nanopure water with 2 % v/v mouse serum (prepared in-house) to a final concentration of 3.24 mg ml^−1^. The suspension was probe sonicated on ice for 16 min with 10 % amplitude. 3.24 mg ml^−1^ corresponds to a dose of 162 µg TiO_2_ nanotubes per 50 µl instillation volume per mice. The vehicle of nanopure water with 2 % v/v mouse serum was probe sonicated using the same protocol. The dose of 162 µg per mouse corresponds to an average surface dose of 3:1 S_nanomaterials_:S_cells_ and is equivalent to 15 working days at the 8-h time-weighted average occupational exposure limit for TiO_2_ by Danish Regulations (6.0 mg m^−3^ TiO_2_).

The average hydrodynamic particle size of the TiO_2_ nanotube in suspension (3.24 mg ml^−1^) was determined by Dynamic Light Scattering (DLS). The TiO_2_ nanotube suspension had a bimodal size distribution with a major peak at 60 nm and a narrow peak at 21 nm. ^[16]^ The intensity-based average size was 168.7 nm and the polydispersity index (PI) was 0.586, indicating some polydispersity in the suspensions. Endotoxin levels were measured using the Limulus Amebocyte Lysate Assay. The level of endotoxins was low in TiO_2_ tube suspensions (0.095 endotoxin units (EU) ml^−1^), and in nanopure water with 2 % mouse serum (0.112 EU ml^−1^).

#### Animal handling and exposure

Seven-week-old female C57BL/6jBomtac mice (Taconic, Ejby, Denmark) were randomized in groups for TiO_2_ nanotube exposure (*N*=5 mice per group for histology) and vehicle controls (*N* = 2-4 mice per group). At 8 weeks of age the mice were anaesthetized and exposed to 0 µg or 162 µg TiO_2_ nanotube in 50 µl vehicle by single intratracheal instillation. In brief, the mice were intubated in the trachea using a catheter. The 50 μl suspension was instilled followed by 200 µL air. The mouse was transferred to a vertical hanging position with the head up. This ensures that the administered material is maintained in the lung. Animal experiments were performed according to EC Directive 2010/63/UE in compliance with the handling guidelines established by the Danish government and permits from the Experimental Animal Inspectorate (no. 2015-15-0201-00465). Prior to the study, the experimental protocols were approved by the local Animal Ethics Council.

More details regarding the animal study can be found in Danielsen et al.. ^[16]^

#### Histology and enhanced dark-field imaging

At 28, 90 or 180 days post-exposure mice were weighed and anesthetized. Lungs were filled slowly with 4% formalin under 30 cm water column pressure. A knot was made on the trachea to secure formaldehyde in lungs to fixate tissue in “inflated state”. Lungs were then removed and placed in 4% neutral buffered formaldehyde for 24 hours. After fixation the samples were trimmed, dehydrated and embedded in paraffin. 3 µm thin sections were cut and stained with haematoxylin and eosin (H&E). Cytoviva enhanced dark-field hyperspectral system (Auburn, AL, USA) was used to image particles and organic debris in the histological sections of mouse lungs. Enhanced dark-field images were acquired at 100x on an Olympus BX 43 microscope with a Qimaging Retiga4000R camera.

#### Transcriptomics *in vivo*

Microarray mRNA analysis was performed using Agilent 8 × 60 K oligonucleotide microarrays (Agilent Technologies Inc., Mississauga, ON, Canada) as described previously ^[39]^ with six replicas for each condition. Bioinformatics analysis of the row data: signal intensities were Loess normalized using the limma package in R/Bioconductor. ^[40]^ Analysis of differentially expressed genes (DEGs) was performed using the limma package. The genes were considered as significantly differentially expressed if the BH-adjusted p-values were less than or equal to 0.1. Statistical analysis is same as for the *in vitro* transcriptomics above.

#### Comparison of transcriptomics *in vitro* and *in vivo*

Mice were exposed to 18, 54 or 162 µg of TiO_2_ nanotubes per mouse and lungs were harvested on 1^st^ and 28^th^ day post exposure for transcriptomic analysis to evaluate overlapping sets of genes differentially expressed in the *in vivo* and *in vitro* experimental data. The goal of the analysis is to determine and compare alterations in lipid metabolism, immune response in terms of proinflammatory signaling and cholesterol metabolism between two experimental systems. For the assessment of the monocyte influx, all genes encoding monocyte chemoattractive (C-C motif) chemokines were selected and their expression evaluated.

### Modelling

Atomistic molecular dynamics simulation

#### System composition

Atomistic molecular dynamics simulations have been carried out for DMPC and POPE lipids near anatase (101) TiO_2_ surface in water environment. Anatase slab (71.8 x 68.2 x 30.5 Å) with (101) surface normal to the z axis is used as a model of a nanoparticle surface. The slab contains 4536 Ti atoms of which 504 are five-fold coordinated atoms on the surface. (101) anatase surface was chosen as a surface of the lowest energy. At neutral pH TiO_2_ surface is covered by hydroxyl groups and is negatively charged. In our model we bind hydroxyl groups to 5- coordinated surface Ti atoms so that the surface charge density is close to the experimental value at neutral pH. Thus we add 151 hydroxyl groups to randomly picked Ti surface atoms (which constitutes 30% of their total amount) which results in a surface charge density of −0.62 electrons nm^−2^, which is in line with the experimental results. ^[41]^

The TiO_2_ slab is then placed in the middle of the simulation box with 3D periodic boundary conditions. The box size in X and Y directions is defined by the slab length and width so that the slab is periodic in those directions. The height of the box is set to 130 Å to accommodate the TiO_2_ slab (thickness of 30.5 Å), eventual formed lipid bilayer on the both sides (2 x 40 Å) as well as their hydration layers (2 x 10 Å). 82 lipid molecules (POPE or DMPC) are inserted at random unoccupied positions in the box in random orientations, after that the box is filled with water molecules (about 12000). Then, a small number of water molecules are picked at random and are substituted with Na^+^ and Cl^-^ ions to balance the negative surface charge of the slab and provide NaCl concentration of 0.15 M in the water phase of the simulated system.

### Simulation protocol

First, energy minimization of the simulated systems using the steepest gradient descent method is performed, followed by a short 100 ps pre-equilibration run at constant volume and temperature. After that, the pressure in the system is equilibrated to 1 bar using anisotropic Berendsen barostat ^[42]^ with relaxation time of 5 ps during 10 ns, which is finally followed by 1 μs production run in the NVT ensemble. Leap-frog algorithm with time step 1 fs is used to integrate the equations of motion. Center-of-mass motion is removed every 100 steps. Verlet cut-off scheme ^[43]^ with the buffer tolerance of 0.005 kJ mol^-1^ ps^-1^ per atom is used to generate the pair lists. Minimum cut-off of 1.4 nm is used for both short ranged electrostatic and VdW interactions. Long range electrostatics are calculated using PME ^[44]^ with the grid spacing of 0.12 nm and cubic interpolation. Long range dispersion corrections are applied to both energy and pressure. Velocity rescaling thermostat ^[45]^ is used to control the temperature, which is set to 303 K with the relaxation time of 1 ps. All bonds with hydrogen atoms are constrained using the LINCS algorithm. ^[46]^ Atom coordinates and energies are saved every 5 ps. All simulations were performed by the Gromacs 2019 software package. ^[47]^ Visualization of the simulations is done by VMD. ^[48]^

### Models used

Lipids are described by the Slipids force field^[49]^. For TiO_2_, we use parameters optimized to fit results on charge density distributions and water-TiO_2_ surface coordination obtained in *ab-initio* simulations of TiO_2_-water interface. ^[50]^ These parameters are listed in tables in SI section S2e. Water molecules are represented by the TIP3P model ^[51]^, and for Na^+^ and Cl^-^ ions Yoo and Aksimentiev ion parameters is used. ^[52]^ Lorentz-Berthelot rules are applied to determine Lennard-Jones parameters for cross-interactions.

### Model of chronic inflammation following nanomaterial exposure

The theoretical model of chronic inflammation following nanomaterial exposure is described by a series of differential equations (see S5b), describing the events observed in *in vitro* and *in vivo* experiments in this work. This minimal-complexity *in vivo* model consists of 6 variables (surface of nanomaterial in epithelial cells, in cauliflowers, in macrophages and freely-floating nanomaterial, surface of macrophages and surface of epithelial cells), 4 fixed parameters which are calibrated for each model system and later locked (endocytosis rate, rate of cauliflower endocytosis, delay between cauliflower production and signaling for macrophage influx, and epithelial cell replication rate) and 3 NM-associated parameters (cauliflower formation rate *cff*, signaling efficiency *signEff*, and toxicity *tox*). Separate *in vitro* models were obtained from the *in vivo* model by swapping the macrophage influx with macrophage replication and leaving out non-existent cells for monocultures.

The system of equations was solved numerically using Wolfram Mathematica 12.0, license L5063-5112 to obtain the time evolution and final state of the model. The same software was also used for visualization of the results.

The phase space was scanned by calculating the time evolution of the appropriate system of equations from chapter S5b for a set of nanomaterials with appropriately interspaced parameters: toxicity (*tox*), cauliflower formation (*cff*) and signaling efficiency (*signalEff*). For each parameter, 30 logarithmically-equally-spaced values in a sensible range were chosen – the total amount of values in the grid was thus 30 x 30 x 30 = 27.000.

More detailed information can be found in Supplementary Information.

## Supporting Information

All supporting information below is available free of charge *via* the Internet at adjacent links.

Source data is publicly available online at http://lbfnanobiodatabase.ijs.si/file/data/cauliflowerpaper/ with all 3Ds, movies and raw tiffs as a part of a database develop for H2020 Smart Nano Tox project.

Source data for Figures 1-5 and for figures in the Supplementary Information is available at the following links:

Source Raw Data Fig.1 (.rar)

Source Raw Data Fig.2 (.rar)

Source Raw Data Fig.3 (.rar)

Source Raw Data Fig.4 (.rar)

Source Raw Data Fig.5 (.rar)

Source Data for Supplementary Information (.rar)

Source data for *in vitro* genomics was deposited in the GEO database under the number GSE146036 and is accessible via the link https://www.ncbi.nlm.nih.gov/geo/query/acc.cgi?acc=GSE146036 using the token listed in the Cover letter.

## Author Contributions

The manuscript was written through contributions of all authors. All authors have given approval to the final version of the manuscript.

‡These authors contributed equally as first authors: Hana Majaron, BoŠtjan Kokot and Aleksandar Sebastijanović.

‡‡These authors contributed equally as corresponding authors: Janez Štrancar, Tobias Stoeger, Tilen Koklič.

HM, BK, AS, CV, RP, PZ, TB, CBL, PHD, CC, VZ, MS, OJ, MIr, JB, VL, SH, NQ, AL, UV, TK, TS, JS designed the study and analysis.

HM, BK, AS, CV, RP, PZ, TB, PHD, CC, AK, PC, QZ, JP, ZMD, MP, PU, SP, MIr, SH prepared the samples.

HM, BK, AS, CV, RP, PZ, TB, PHD, CC, AK, PC, QZ, JP, ZMD, MP, MIr, SH performed the experiments.

HM, BK, AS, CV, RP, PZ, TB, CBL, CC, AK, QZ, JP, VZ, ZMD, MP, MIr, JB, SH, TK, TS, JS analyzed the data.

HM, MIv and MS performed the modelling.

OJ, JB, VL, SH, NQ, AL, UV, TK, TS, JS supervised the study.

HM, BK and AS prepared the manuscript with input from all other authors: CV, RP, PZ, TB, CBL, PHD, CC, MIv, AK, PC, QZ, JP, VZ, MS, ZMD, MP, PU, SP, OJ, OS, IU, MIr, JB, VL, SH, NQ, AL, UV, TK, TS, JS.

## Acknowledgments

We are grateful to the team at TeScan for FE-SEM measurements and would like to thank G. Hlawacek and N. Klingner for assistance on HIM. We thank K. Richter for excellent technical assistance for the transcriptomics analysis and J. Birkelund Sørli with *in vivo* experiments. We kindly thank JRC Nanomaterials Repository for providing us with various nanomaterials and the team from Syglass for their support.

Corresponding Authors

Janez Štrancar

Tobias Stoeger

Tilen Koklič

