## Supplementary Information for "From the Roundabout of Molecular Events to Nanomaterial-Induced Chronic Inflammation Prediction"

#### Detailed table of contents

A word on supplement arrangement ........ 10

--- Materials, methods and nanomaterial characterisation --- 11
S0a – General materials and methods 11
Materials 11
Animals and cells 11
Chemicals 11
Materials 11
Nanomaterials used in this study 12
Software 12
Chemicals for imaging 12
Non-commercial fluorescent probes 13
Main message 13
*Figure S1 - Figure S3* 13
Characterisation 13
Cell culture 15
Nanomaterial synthesis and labelling 15
General in vitro sample preparation and exposure 15
Imaging in vitro 16
STED 16
FLIM 16
Imaging of nanomaterial in backscatter mode 16
SEM 17
HIM 17
FMS 17
TEM 17
Transcriptomics in vitro 17
Transcriptomics in vivo 18
Comparison of transcriptomics in vitro and in vivo 18
In vivo experiments 18
Modelling 18
S0b – Nanomaterial characterisation 19
Zeta potential 19
*Figure S4* 19
Materials and methods 19
Results 20
DLS, TEM of $TiO_2$ nanotubes 20
TEM of $TiO_2$ nanotubes and other nanomaterials, used in this study 20

Cell viability ... 53
Metabolic activity ... 53
Experiment names ... 54
Controls and statistics ... 54
S0g – Cell viability LA-4 – various nanomaterials, same surface dose ... 55
LA-4 murine alveolar cells, PI and Hoechst assay, various nanomaterial ... 55
*Figure S33 ... 55*
Materials and methods ... 55
Experiment names ... 56
Results ... 56
S0h – In vitro exposure of cocultures of MH-S&LA-4 to $TiO_2$ ... 57
Materials and methods ... 57
Experiment names ... 58
Controls and statistics ... 59
*Figure S34 ... 59*
Link to time-lapse ... 60
--- Supplementary information for experiments in Figures 1 – 5 in the main text --- ... 61
S1 – Quarantine of nanomaterials ... 62
S1b – In vivo data ... 63
Main message ... 63
*Figure S35 ... 63*
Materials and methods ... 63
Preparation and characterization of $TiO_2$ tube suspensions ... 63
Animal handling and exposure ... 63
Histology and enhanced darkfield imaging ... 64
Controls and statistics ... 64
S1c – Cauliflowers with various nanomaterial ... 66
Main message ... 66
Materials and methods ... 66
Experiment names ... 68
Control and statistics ... 72
*Figures S 36-49 ... 72*
S1d – Time evolution of cauliflower growth ... 88
*Figure S50-Figure S53 ... 88*
*Figure S54-Figure S56 ... 88*
Materials and methods ... 88

Comparison of transcriptome response to TiO2 perturbation for in vivo and in vitro conditions163

Experiment names ... 163
S2e – In silico data – atomistic molecular dynamics simulation ... 164
Pool of lipids ... 164
Main message ... 164
*Figure S104-Figure S105 ... 164*
*Table S 9-10 ... 164*
Link to time-lapse ... 165
Link to 3D ... 165
Materials and methods ... 166
*System composition ... 166*
*Simulation protocol ... 167*
*Models used ... 167*
Modelling of bilayer adhesion to $TiO_2$ surface ... 167
Main message ... 167
*Figure S106 ... 167*
Materials and methods ... 168
S2f – Ultrafast passage of nanomaterial of cell membrane ... 169
*Figure S107-Figure S111 ... 169*
Link to time-lapse ... 169
Materials and methods ... 169
Experiment names ... 170
Controls and statistics ... 170
S2g – Blocking clathrin-mediated endocytosis ... 175
*Figure S112-Figure S121 ... 175*
Materials and methods ... 175
Link to time-lapse ... 176
Link to 3D ... 176
Experiment names ... 177
Controls and statistics ... 178
S2h – Membrane cholesterol extraction with Methyl-Beta-Cyclodextran ... 186
*Figure S122-Figure S125 ... 186*
Materials and methods ... 186
Experiment names ... 187
Controls and statistics ... 187
Link to 3D ... 192
Link to time-lapse ... 192

S2i – Block of lipid synthesis by FAS inhibition ... 193
*Figure S126-Figure S131 ... 193*
Materials and methods ... 193
Experimental names ... 193
Controls and statistics ... 194
Link to 3D ... 200
Link to time-lapse ... 201
S3 – The role of actin ... 202
S3b – Actin – nanomaterial aggregation ... 203
*Figure S132-Figure S134 ... 203*
Materials and methods ... 203
Experiment names ... 203
Controls and statistics ... 204
Link to 3D ... 207
S3c – Inhibition of exosome excretion by high concentrations of Jaspakinolide ... 209
*Figure S135-Figure S140 ... 209*
Materials and methods ... 209
Experiment names ... 209
Controls and statistics ... 210
Link to 3D ... 216
S3d – Nanoparticle actin interaction ... 218
*Figure S141-Figure S144 ... 218*
Materials and methods ... 218
Experiment names ... 218
Controls and statistics ... 218
Link to time-lapse ... 223
S3e – Actin branching ... 224
*Figure S145-Figure S147 ... 224*
Materials and methods ... 224
Experiment names ... 224
Controls and statistics ... 224
S3f – Actin in cauliflowers ... 228
*Figure S148-Figure S151 ... 228*
Materials and methods ... 228
Experiment names ... 228
Controls and statistics ... 228

#### A word on supplement arrangement

The first part of the supplement (S0a to S0h, pages 11 - 62) contains general supporting material for the main paper (“Materials, methods and nanomaterial characterisation”) as well as complementary data used in the main text but not shown in Figures in the main text (“Complementary experiments, not included in figures in the main text”).
From S1 (page 64) onwards, detailed supplementary material for each image from the main text is shown (“Supplementary information for experiments in Figures 1 – 5 in the main text”). It contains details on experimental design, controls and repetitions (fluorescence images shown both in separate channels and overlayed), and names (cyphers) of experiments to ease locating them in the depository/database.
We first supply the main text image duplicates for easier orientation. The supplement section correspond to the panels in the main text. For example, section S2c in the supplement corresponds to the experiment in Figure 2c in the main text.

### --- Materials, methods and nanomaterial characterisation ---

#### S0a – General materials and methods

##### Materials

###### Animals and cells

- LA-4 murine alveolar epithelial cells (ATCC)
- MH-S murine alveolar macrophages (ATCC)
- NR8383 rat alveolar macrophages (ATCC)
- C57BL/6jBomtac mice (Taconic, Ejby, Denmark)

###### Chemicals

- LCIS: Live Cell Imaging Solution (Invitrogen)
- PBS: phosphate buffer saline (Gibco)
- 100x dcb: 100-times diluted bicarbonate buffer, pH 10, osmolarity 5 miliosmolar, mixed in-house
- F-12K: cell culture medium for LA-4 (Gibco)
- RPMI 1640: cell culture medium for MH-S (Gibco)
- Trypsin (Sigma)
- Penicillin-Streptomycin (Sigma)
- Non-essential amino acids (Gibco)
- Beta mercaptoethanol (Gibco)
- dH2O: deionised water
- glucose (Kemika)
- BSA: bovine serum albumin (Sigma)
- Hydrogen peroxide (Merck)
- Chlorpromazine (Alfa Aesar)
- MBCD: Metyl-Beta-Cyclodextran (Acros organics)
- Resveratrol (Sigma)
- KCl (Kemika)
- HCl (Merck)
- KOH (Carlo Erba)
- Limulus Amebocyte Lysate Assay (Lonza, Walkersville, MD, USA)
- 10% neutral buffered formalin (CellPath Ltd, UK)
- DMEM (Sigma-Aldrich, France, Saint-Quentin-Fallavier)
- heat-inactivated FBS (Sigma-Aldrich, France, Saint-Quentin-Fallavier)
- 4 mM L-glutamine (SIGMA-G7513)
- antibiotic/antimycotic composed of 100 U/mL of penicillin, 100 μg/mL of streptomycin (SIGMA-P0781) and 0.25 μg/mL of amphotericin B (SIGMA-A2942)
- LDH assay (Roche-4744934001, Germany)
- Triton (Sigma-Aldrich, France, Saint-Quentin-Fallavier)
- WST-1 assay (MV Berridge, AS Tan, KD McCoy, 1996) (Roche, 11644807001, USA)
- WST-1 kit (Roche, Germany)
- GeneChip® WT PLUS Reagent Kit (Thermo Fisher/Affymetrix)
- RNeasy Plus Mini Kit (Qiagen)
- WT PLUS Reagent Kit (Thermo Fisher Scientific Inc., Waltham, USA)

##### Materials

- #1.5H μ-Dish (Ibidi)

- #1.5H µ -Slide 8-well (Ibidi)
- Pelcotec™ SFG12 Finder Grid Substrate - Si wafers for HIM (Ted Pella)
- Aeroneb®Pro nebulizer (from VITROCELL® Cloud 6 system)
- Mouse Clariom S arrays (Thermo Fisher Scientific)

#### Nanomaterials used in this study

Synthesized in-house by P. Umek:
$TiO_2$ nanotubes (PU-nTOX-01-03) and $TiO_2$ nanocubes (PU-nTOX-01-21);

Kind gift from U. Vogel:
carbon black (Printex 90), $TiO_2$ MKNA015 (MKN- $TiO_2$ -A015), $TiO_2$ MKNA100 (MKN-$TiO_2$ -A100) and quartz silica ($SiO_2$ DQ12);

Kind gift from JRC Nanomaterial Repository:
NM-101 TiO2 anatase (TiO2-NM101-JRCNM01001a), NM-105 TiO2 rutil-anatase (TiO2-NM105-JRCNM01005a), NM-110 ZnO (ZnO-NM110-JRCNM62101a), and NM 111 ZnO (ZnO-NM111-JRCNM01101a), NM-200 SiO2 (SiO2-NM200-JRCNM02000a), NM-401 MWCNT (MWCNTs-NM401-JRCNM04001a), NM-402 MWCNT (MWCNTs-NM402-JRCNM04002a).

#### Software

- Imspector (version 16.2.8282-metadata-win64-BASE), provided by Abberior
- SPCImage 7.3 (Becker & Hickl)
- Fiji, ImageJ 1.52p (NIH)
- syGlass (http://www.syglass.io/, RRID:SCR_017961)
- Mathematica 12.0, licence L5063-5112 (Wolfram)
- genomics software: GSEA by Broad Institute
- modelling: GROMACS 2018.3 (calculation), VMD (visualisation)

#### Chemicals for imaging

Staining for *ex vivo* imaging:

- haematoxylin and eosin (H&E) (Trine)

Fluorescent probes used for $TiO_2$ labelling:

- Alexa Fluor 647 NHS ester (Termo Fisher), $\lambda_{Ex/Em}$: 651/672 nm
- Star 520 SXP NHS ester (Abberior), $\lambda_{Ex/Em}$: 520/640
- ATTO 594 NHS ester (Atto-tec), $\lambda_{Ex/Em}$: 603/626 nm

Fluorescent probes used for cell labelling:

- CellMask Orange (Invitrogen)
- SiR Actin (Cytoskeleton)
- Priopidium Iodide, PI (Sigma)
- Hoechst 33342 (Sigma)
- pHrodo Red Transferrin conjugate (Invitrogen)
- Lysotracker Red (Invitrogen)
- MitoTracker™ Orange CMXRos (Invitrogen)
- Star Red-DPPE (Abberior)

#### Non-commercial fluorescent probes

##### *Main message*

Characterisation of used home-made fluorescent probes SHE-2N and SAG-38. SHE-2N labels plasma membrane, SAG-28 labels lipids droplets.
Supporting raw and analysed data:

Figure S1 - Figure S3

##### *Characterisation*

Spectra were recorded in toluene at $10^{-7}$ $M$ concentration on Perkin Elmer LS-55 spectrofluorometer. For information regarding the probe synthesis contact the authors.

- 4-(8,9-Dimethyl-6,8-dinonyl-2-oxo-8,9-dihydro-2H-pyrano[3,2-g]quinolin-3-yl)-1-(3-(trimethylammonio)propyl)pyridin-1-ium dibromide (**SHE-2N**)

**SHE-2N** is an amphiphilc fluorescent probe designed for labelling of plasma membrane. Lipophilic tails and coumarin moiety are incorporated into lipid bilayer, while two positive charges at the headgroup reduce distribution to other membranes of the cell and thus provide stable labelling of plasma membrane. Additionally, labelling of plasma membrane is quick and with low photobleaching.

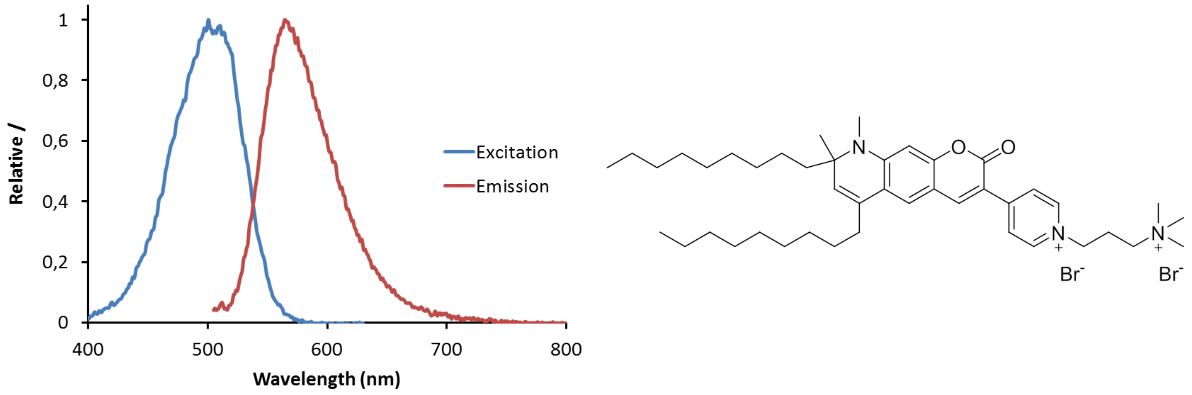

*Figure S1: Left: normalized fluorescence excitation (blue line) and emission (red line) spectra of **SHE-2N** recorded in toluene at $10^{-7}$ M concentration. Right: structure of membrane probe **SHE-2N**.*

- 3-(Benzo[d]thiazol-2-yl)-6,8,8,9-tetramethyl-2-oxo-8,9-dihydro-2H-pyrano[3,2-g]quinoline-4-carbonitrile (**SAG-38**)

**SAG-38** is a lipophilic coumarin-based probe designed as an alternative to nile red for labelling of lipid droplets. Emission of **SAG-38** is red-shifted in comparison to emission of nile red and ideal for recording of emission within 580–625 nm.

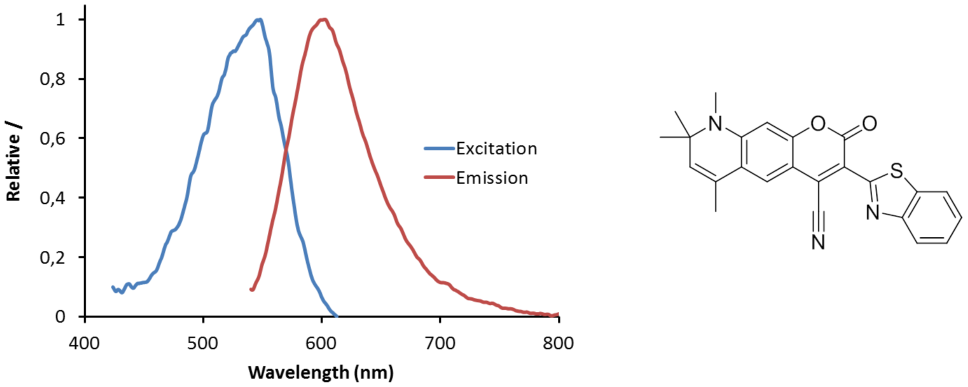

*Figure S2: Left: normalized fluorescence excitation (blue line) and emission (red line) spectra of* ***SAG-38*** *recorded in toluene at* $10^{-7}$ *M concentration. Right: structure of probe* ***SAG-38****.*

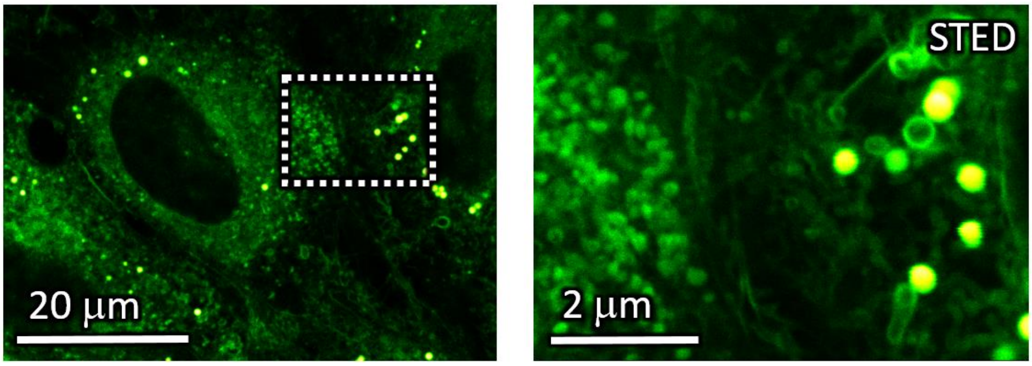

*Figure S*3: A confocal (left) and STED (right) image of a non-exposed LA-4 cell labelled with SAG-38 – the probe labels various vesicles, including lamellar bodies (bright, full vesicles).

- experiment 20190607/e03_s02_t05_SAG-38_STED.msr :
  - LA-4 cells were seeded @30% confluence in an Ibidi #1.5H µ-Slide 8-well
  - after 2 days, 10 µM probe slightly lipophilic probe SAG-38 was freshly diluted to 10 µM in LCIS, added to cells and incubated for 10 minutes, after which the image was taken
- analysis:
  - rescaling of green channel

| | |
|---|---|
| Cell line | LA-4 (membranes and lipid droplets, SAG-38) |
| NPs | |
| exposure | |
| imaging | xy confocal |

| | |
|---|---|
| pixelsize (x,y) | 50 nm |
| FOV (x,y) | 60 µm |
| pixelsize (z) | |
| FOV (z) | |
| imaging time | |
| number of frames | |

| | |
|---|---|
| 561nm | 2% |
| 640nm | |
| STED | |
| filter sets | 605 nm – 625 nm |
| dwell time | 10 µs |
| objective | 60x wi (NA1.2) |

| | |
|---|---|
| Cell line | LA-4 (membranes and lipid droplets, SAG-38) |
| NPs | |
| exposure | |
| imaging | xy STED |

| | |
|---|---|
| pixelsize (x,y) | 20 nm |
| FOV (x,y) | 15x11 µm |
| pixelsize (z) | |
| FOV (z) | |
| imaging time | |
| number of frames | |

| | |
|---|---|
| 561nm | 20% |
| 640nm | |
| STED | 25% |
| filter sets | 605 nm – 625 nm |
| dwell time | 10 µs |
| objective | 60x wi (NA1.2) |

#### Cell culture

Murine epithelial lung tissue cell line (LA – 4; cat. no. ATCC CCL-196) and murine alveolar lung macrophage (MH-S; cat. No. CRL2019) cell line were purchased from and cultured according to American Type Culture Collection (ATCC) instructions. Cells were cultured in TPP cell culture flasks at 37 °C in a 5% $CO_2$ humidified atmosphere until monolayers were 80% confluent, at which time they were subcultured or seeded for experimentation. All experiments were performed with cells before the twentieth subculture. For long–term live cell experiments, a homemade incubator which maintains a humidified atmosphere with a 5% $CO_2$ and is heated to 37 °C.

Medium used for culturing of the epithelial LA-4 cells is Ham's F-12K (Kaihn's) medium, produced by Gibco, supplemented with 15% FCS (ATCC), 1% P/S (Sigma), and 1% NEAA (Gibco), 2 mM L-gln. For the specific composition of Ham's F-12K medium, look for details on the TermoFisher webpage under the catalogue number 21127022.

For Alveolar macrophages, MH-S, cell line we used RPMI 1640 (Gibco, ATCC modification) medium, supplemented with 10% FCS (ATCC), 1% P/S (Sigma), 2 mM L-gln, and 0.05 mM beta mercapthoethanol (Gibco). For the specific composition of this medium, look for details on the TermoFisher webpage under the catalogue number A1049101.

#### Nanomaterial synthesis and labelling

The $TiO_2$ anatase nanotubes used in this paper were synthesized, functionalized with AEAPMS, and labelled with STED-compatible fluorescent probes via a covalent reaction between the AEAPMS and ester functional group on the probe. All this was done in-house [1]. Labelled $TiO_2$ was then stored suspended in 100x diluted bicarbonate buffer. For the multi-nanomaterial exposure experiments we used other nanomaterials as well. In this case, the nanomaterials were suspended in PBS and sonicated in ice bath using a tip sonicator (Sonicator 4000, Misonix, with 419 Microtip probe) for 15 min with 5s ON/ 5s OFF steps.

#### General *in vitro* sample preparation and exposure

LA-4 and MH-S cells were seeded in Ibidi 1.5H dishes of various surface area, depending on the experiment. After 24 h, nanomaterial (c=1mg/mL) was added at an appropriate surface dose (SNP:Scells), according to the experiment needs (listed for each experiment in sections S1 – S5). Before exposure, nanomaterial suspension was sonicated for 10 s in an ultrasonic bath (Bransonic ultrasonic cleaner, Branson 2510EMT). Cells were then incubated at 37 °C and 5% CO2 atmosphere with the nanomaterial for the desired time in order to observe the cells at the post-exposure time points of interest. If the experiment required monoculture of either cell line, sample were prepared as described above, if however, we experimented with the co-cultures, sample preparation differed slightly. For co-cultures, we grew LA-4 and MH-S in separate dishes up to desired confluency (lower than for monocultures) and then mixed them together by adding MH-S in the LA-4 dish at a ratio of 1 : 40. Co-cultures were then incubated for 24 h more, exposed to nanomaterial as described above and incubated for additional desired amount of time. Growth medium for co-cultures was mixture of equal volumes of F12K and RPMI 1640. Cells were then labelled with fluorescent dyes according to the manufacturer's recommendations. Right before observing the live cells, unbound fluorescent label was washed and medium was exchanged for LCIS.

In some experiments we used different chemicals for modulation of the cell metabolism. For blocking the Clathrin-mediated endocytosis, cells were treated with 100 μm Chlorpromazine for 15 min. Membrane cholesterol was extracted with a 24 h incubation with 0.5 - 1 mM MBCD. FAS was inhibited with overnight 100 µM Resveratrol incubation. Finally, for actin stabilization, we used higher concentration (≥1mM) of Sir-Actin Label based on Jasplankinolide. All the chemical modulators were added before exposure to nanomaterial and continued to be incubated with the cells even after during incubation with the nanomaterial for abovementioned time periods.

For the reuptake experiments different cell lines were grown separately, and washed with PBS before adding MH-S to LA-4.

HIM, SEM
Samples were prepared as usual but we grew them on Si-wafers. After reaching desired confluency samples were freeze-dried with metal mirror freezing technique.

#### Imaging *in vitro*

##### STED

Super-resolution and confocal fluorescence micrographs were acquired using custom build STED microscope from Abberior with an Olympus IX83 microscope and two avalanche photodiodes as detectors (APDs). The microscope is equipped with two 120 picosecond pulsed laser sources (Abberior) with excitation wavelengths 561 and 640 nm and maximal power of 50 µW in the sample plane. Pulse repetition frequency for experiments was 40 - 80 MHz, depending on the experiment. STED depletion laser wavelength is 775 nm with same repetition frequency as excitation lasers, pulse length of 1.2 ns and maximal power of 170 mW in the sample plane. Filter sets used for detection were either 605–625 nm (green channel) or  650–720 nm (red channel). Images were acquired using Imspector (version 16.2.8282-metadata-win64-BASE) software also provided by Abberior. All microscope settings were tuned separately for maximal resolution during each of the experiments and are listed with alongside the recorded images in Supplement.

##### FLIM

Fluorescence lifetime images (FLIM) were obtained on the same custom-built STED microscope (Abberior instruments) as confocal and STED fluorescence images in this study. This time, the emitted fluorescence was detected using PMT detectors and TCSPC technology developed by Becker & Hickl. 16-channel GaASP PMT detectors attached to a spectrograph with diffraction grating 600 l/mm were used to measure fluorescence lifetime of emitted photons with wavelengths ranging from 560 to 760 nm. Spectral information was discarded and the lifetimes were gathered in Imspector 16.2 (Abberior Instruments).

The fluorescence lifetime data was analysed with SPCImage 7.3 software (Becker & Hickl), where the Decay matrix was calculated from the brightest pixel in the image (monoexponential fitting), binning was set to 3 and threshold to 5. The rainbow LUT was rescaled to range from 500 ps to 1000 ps for all images and both intensity and contrast of the lifetime-coded image were adjusted for easier comparison of lifetimes between samples.

##### Imaging of nanomaterial in backscatter mode

In Figure 1c, simultaneously with measuring fluorescence from CellMask Orange in the cell membrane (as described in STED section), backscattered light was detected as well to locate the nanomaterial in the sample. A tuneable Chameleon Discovery laser (Coherent) with 100 fs

long pulses, pulse repetition frequency 80 MHz, and maximal average power of 1.7 W at 850 nm was used as the scattering light. The pre-attenuated laser light with a wavelength of 750 nm first passed through a 785 nm built-in dichroic where a fraction of the power was directed onto the sample through the same 60x WI objective (NA 1.2) as the excitation light for fluorescence imaging. The light scattered off the nanomaterial and passed back through the same objective and dichroic, now mostly passing through the dichroic towards the detectors. After passing through a pinhole (0.63 A.U.), the backscattered light was spectrally separated from the fluorescence by a short-pass 725 nm dichroic, afterwards being detected on the same PMT, as described in the FLIM section, this time set to collect light with wavelengths above 725nm.
Due to the large coherence of the laser, the backscattered light exhibited a strong speckle pattern, which was diminished by a 100-nm-wide Gaussian blur on the scattering image, thus decreasing false negative colocalisation of nanomaterial on account of spatial resolution.

##### SEM

SEM imaging has been performed on MIRA3 Flexible FE-SEM produced by TESCAN, by detection of secondary electrons. Beam powers used have been between 5.0 kV and 15 kV with variable field of view 1.8 μm to 180 μm. All samples have been measured under high pressure vacuum (HiVac). All analysis has been performed in Tescan developed software.

##### HIM

Super-resolution imaging on the nanoscale was carried out using Helium Ion Microscope (Orion NanoFab, Zeiss) available at IBC at the Helmholtz-Zentrum Dresden - Rossendorf e. V., a member of the Helmholtz Association. Microscope equipped with GFIS injection system and additional in-situ backscatter spectrometry and secondary ion mass spectrometry can achieve 0.5 nm lateral resolution imaging using 10-35 keV He ion beams. Measurements of secondary electrons (Se) emitted from the first few nm of the sample were done by He ion acceleration of 30 keV, current of 1.7 pA and were acquired under high vacuum inside the sample chamber (3x10-7 mBar). Field-of-view was varied from 60 μm x 60 μm down to 1 μm x 1 μm, with pixel steps small as 2nm. Imaging was performed on non-tilted and tilted sample stage (45 degrees) for better 3-D visualization.

##### FMS

Fluorescence micrographs of cell viability have been acquired on inverted fluorescence Nikon Eclipse TE 2000-E microscope with Xe-Hg source (Sutter Lambda LS, Novato, CA). Filter sets which have have been used were manufactured by BrightLine from Semrock, Rochester, NY. Images were taken with EMCCD camera (iXon3 897 from Andor, Belfast, UK).

##### TEM

See S0b – Nanomaterial characterisation.

#### Transcriptomics *in vitro*

Cells were grown in 6-well plates and exposed to $TiO_2$ nanotubes for 4 h and 48 h, control samples were taken at 0 h and 48 h. Samples were prepared as described above. Briefly, growth medium was removed and the 6-well plates containing cells only were frozen at -70°C. Total RNA was isolated employing the RNeasy Plus Mini Kit (Qiagen). The Agilent

2100 Bioanalyzer was used to assess RNA quality and RNA with RIN>7 was used for microarray analysis.
Total RNA (120 ng) was amplified using the WT PLUS Reagent Kit (Thermo Fisher Scientific Inc., Waltham, USA). Amplified cDNA was hybridized on Mouse Clariom S arrays (Thermo Fisher Scientific). Staining and scanning (GeneChip Scanner 3000 7G) was done according to manufacturer`s instructions.
Statistical analysis for all probe sets included limma t-test and Benjamini-Hochberg multiple testing correction. Raw p-values of the limma t-test were used to define sets of regulated genes ($p<0.01$). Detection Above Background (dabg) p-values were used to exclude background signals: significant genes were filtered for $p<0.05$ in more than half of the samples in at least one group. Array data has been submitted to the GEO database at NCBI (GSE146036).
In the arrow graphs, only genes which were up- or down-regulated more than two times compared to non-exposed cells are shown. The signal (x axis) is drawn in logarithmic scale. Expression is normalized to expression of control samples.

#### Transcriptomics *in vivo*

Microarray mRNA analysis was performed using Agilent 8 × 60 K oligonucleotide microarrays (Agilent Technologies Inc., Mississauga, ON, Canada) as described previously [2] with six replicas for each condition. Bioinformatics analysis of the row data: signal intensities were Loess normalized using the limma package in R/Bioconductor [3]. Analysis of differentially expressed genes (DEGs) was performed using the limma package. The genes were considered as significantly differentially expressed if the BH-adjusted p-values were less than or equal to 0.1. Statistical analysis is same as for the *in vitro* transcriptomics above.

#### Comparison of transcriptomics *in vitro* and *in vivo*

Mice were exposed to 18, 54 or 162 µg of $TiO_2$ nanotubes per mouse and lungs were harvested on 1st and 28th day post exposure for transcriptomic analysis to evaluate overlapping sets of genes differentially expressed in the *in vivo* and *in vitro* experimental data. The goal of the analysis is to determine and compare alterations in lipid metabolism, immune response in terms of proinflammatory signalling and cholesterol metabolism between two experimental systems. For the assessment of the monocyte influx, all genes encoding monocyte chemoattractive (C-C motif) chemokines were selected and their expression evaluated.

#### *In vivo* experiments

See S1b – *In vivo* data.

#### Modelling

See S2e – *In silico* data – atomistic molecular dynamics simulation, S5b – Model of chronic inflammation following nanomaterial exposure and determination of its parameters and S5c – Phase space of chronic inflammation

### S0b – Nanomaterial characterisation

#### Zeta potential

##### Main message

Surface charge characterisation by measurement of ζ-potential on all our nanomaterials used for exposure in experiments presented in this paper.

Supporting raw and analysed data:

Figure S4

##### Materials and methods

All ζ-potential measurements were carried out on the NanoBrook ZetaPALS Potential Analyzer (Brookhaven) with Biolab ζ-potential electrode for non-organic solvents (AQ-1203, Brookhaven) (Holtsville, NY). Samples were prepared and measured as follows:
Samples for measurements were prepared as 10x dilutions of TiO2 nanotubes (1 mg/mL) in dH2O with 10 mM KCl (Kemika). Desired initial pH values were adjusted with HCl (Merck) or KOH (Carlo Erba) and measured with Seven Multi, Mettler Toledo pH meter equipped with 1 mm thick pH electrode (Inlab ExpertPro, Mettler Toledo). ζ-potential has been measured across the range of 8 or more pH values. Analyser was used according to the manufacturers recommendations under the following parameters:

- Size: 10-100 nm
- Concentration: 0.1 mg/mL
- Cycles: 10
- Runs: 10
- Mode of measurement: Smoluchowski
- Temperature: 20°C - 23°C
- Media: Aqueous

Samples for ζ-potential measurement are prepared from batches synthesised under different conditions (look under S0a – General materials and methods, Nanomaterial synthesis and labelling):

- $TiO_2$-17-non-labeled
- $TiO_2$-17-Alexa 647
- $TiO_2$-17-Star 520S
- $TiO_2$-40-non-labeled
- $TiO_2$-40-Alexa 647
- $TiO_2$-03-non-labeled

#### Results

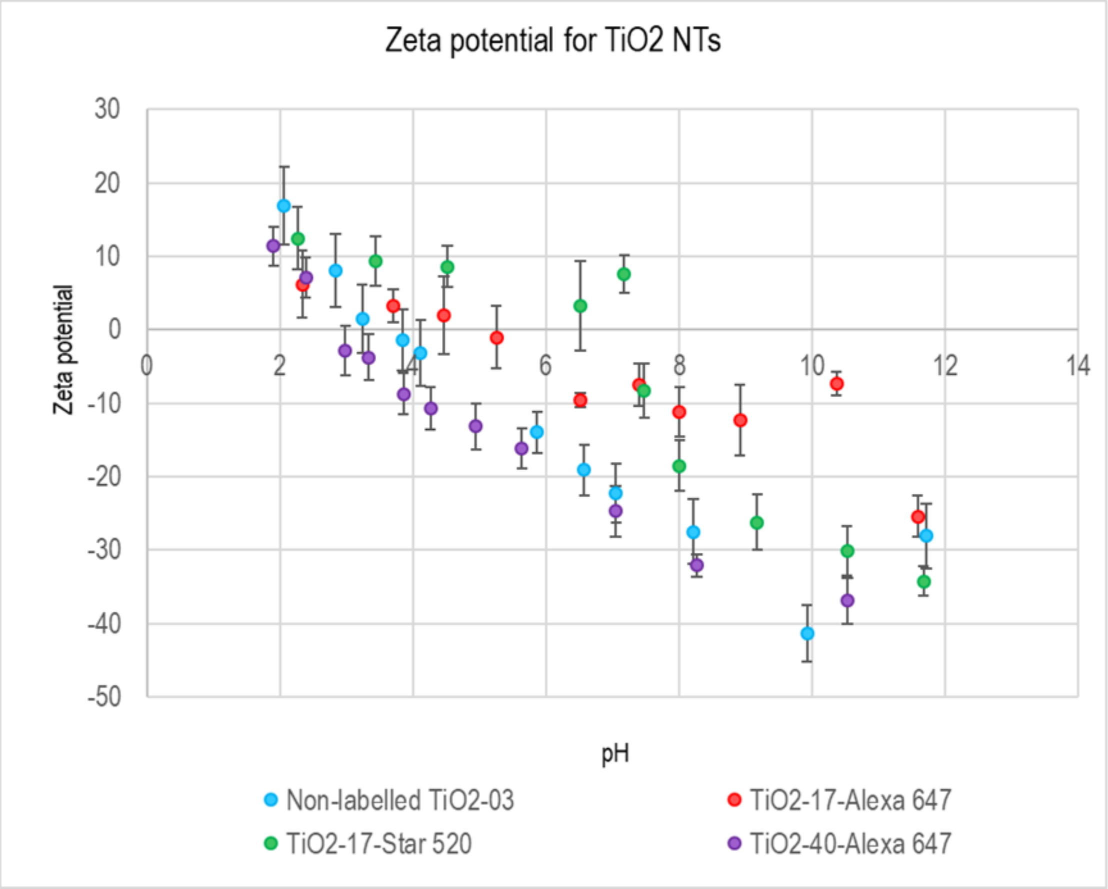

*Figure S4 Measured ζ-potential of AEAPMS functionalized $TiO_2$ nanotubes from the batch 40. Error bars represent the standard error.*

#### DLS, TEM of $TiO_2$ nanotubes

Detailed nanomaterial characterisation of $TiO_2$ nanotubes can be found in the recently submitted paper "Effects of physicochemical properties of $TiO_2$ nanomaterials for pulmonary inflammation, acute phase response and alveolar proteinosis in intratracheally exposed mice [4].The length of $TiO_2$ nanotubes was found to be between 40-500 nm and diameter 6-11 nm, based on TEM images. The DLS showed a bimodal distribution with a minor peak at 21 nm and a major peak at 60 nm.

#### TEM of $TiO_2$ nanotubes and other nanomaterials, used in this study

##### Main message

Nanomaterial structure is determined with transmission electron microscope with nm spatial resolution.

##### Materials and methods

Samples for TEM images of nanomaterials NM101, NM105, NM200, NM402 were prepared using the following protocol.

Of each material 1 mg was dispersed in 1 mL MilliQ water, except CNTs in 1 mL tannic acid solution 300mg/L, using a vial tweeter for 15 min. Each suspension was diluted 1/10 and 3 µL drop deposited on Formvar Carbon coated 200 mesh copper grids (Agar Scientific, USA) and dehydrated overnight in a desiccator before analysis. Images were collected by JEOL JEM-2100 HR-transmission electron microscope at 120kV (JEOL, Italy).

Samples for TEM images of nanomaterials DQ-12, $TiO_2$ A015, $TiO_2$ A100, $TiO_2$ nanotubes, $TiO_2$ nanocubes were prepared using the following protocol.
The nanoparticles were dispersed in water and the dispersion sonicated in water bath for ~3h before use.
Of each sample 5 µl was deposed onto glow-discharged copper grid (Agar scientific Ltd, UK) for one minute and the excess of sample was removed blotting with filter paper. After shortly washing with one drop of water, the grid was therefore immersed into a 2% uranyl acetate (UA) solution for 20 s and blotted again with filter paper. The grids were imaged using a JEOL JEM-2100F fitted with a Gatan Orius SC 1000 camera (2x4k).

TEM images of carbon black nanoparticles (Printex90) were reprinted with permission from reference [5].

#### Images and statistics

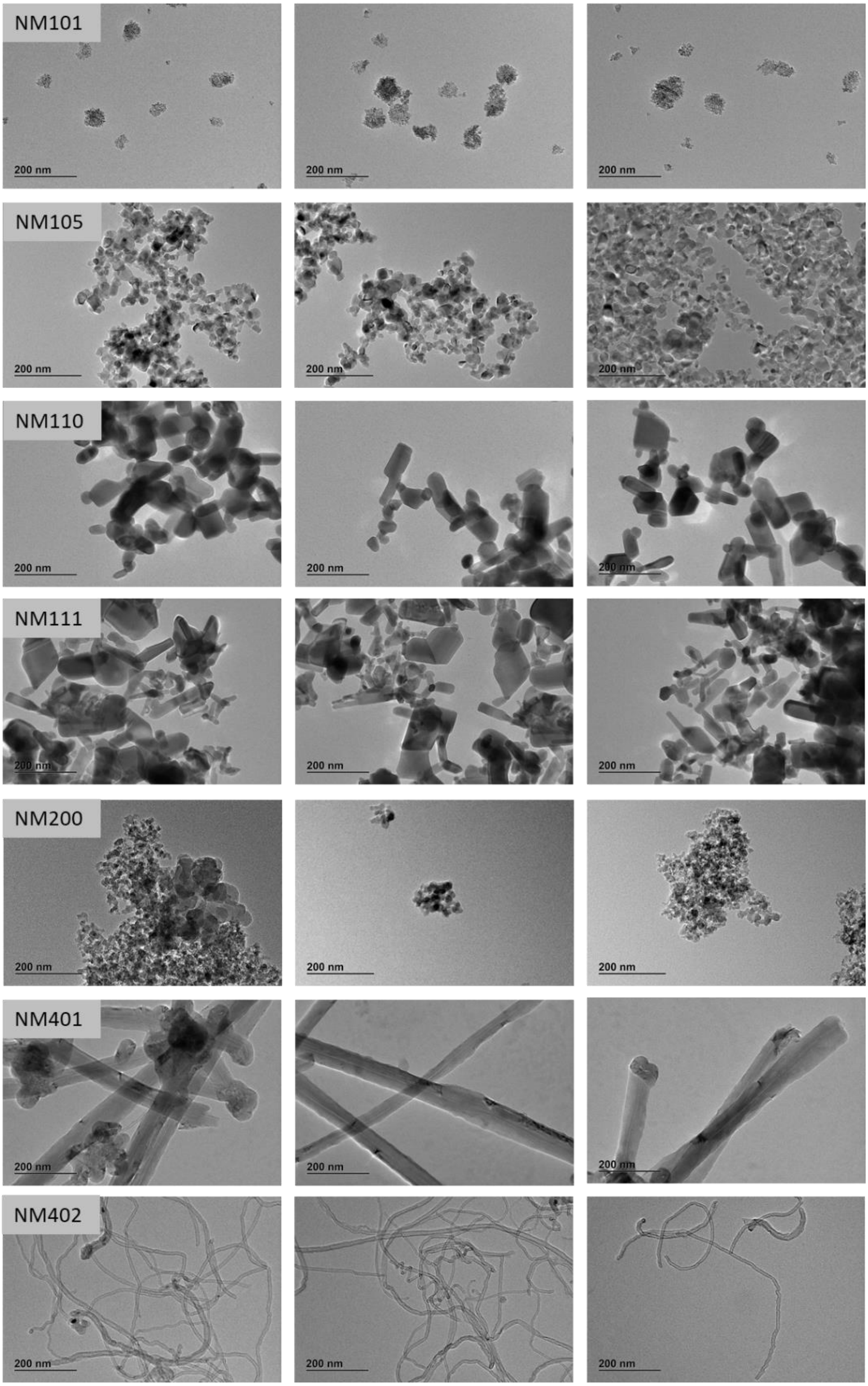

*Figure S5: TEM images of NM101, NM105, NM110, NM11, NM200, NM401 and NM402.*

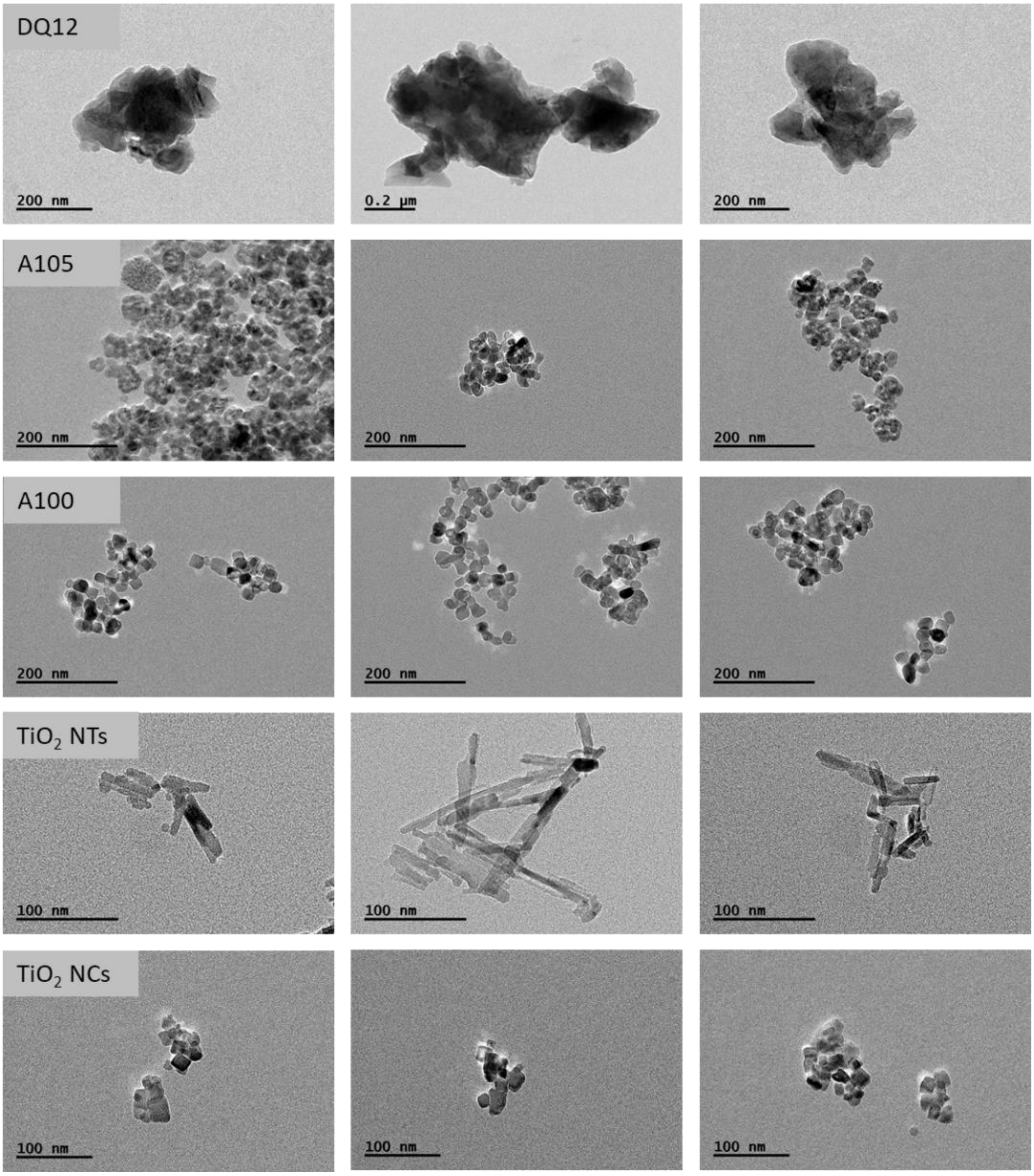

*Figure S6: TEM images of DQ12, A105, A100, $TiO_2$ nanotubes and $TiO_2$ NCs.*

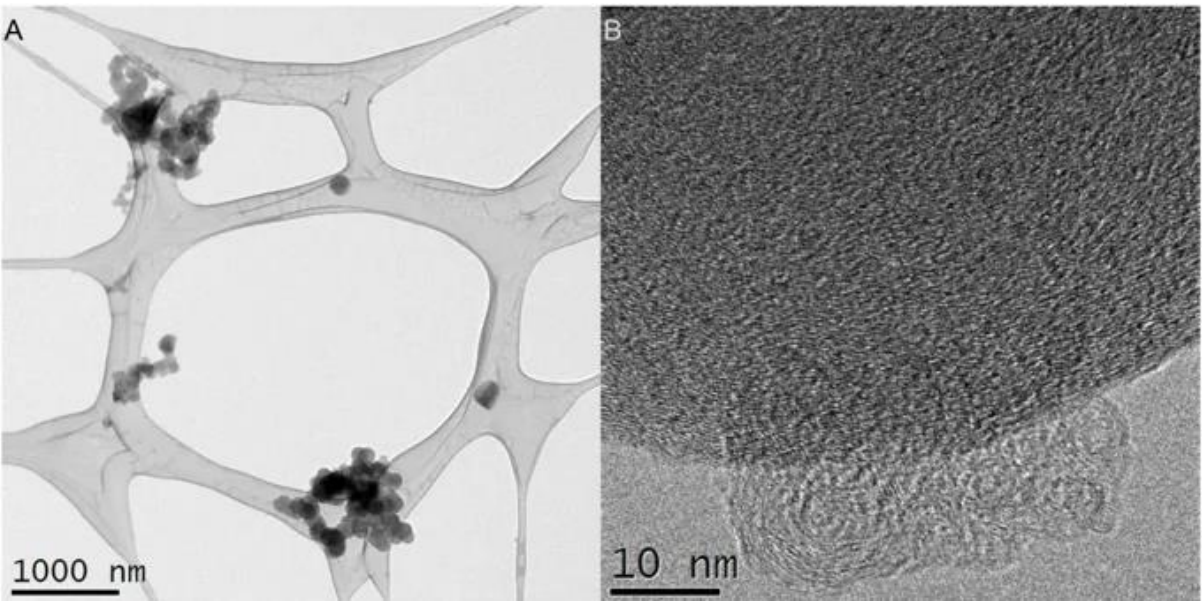

*Figure S7: TEM images of carbon black Printex90, reprinted with permission from reference [5].*

#### --- Complementary experiments, not included in figures in the main text ---

#### S0c – Interaction between $TiO_2$ nanotubes and lipid model membranes

##### Main message

Non-specific interaction between $TiO_2$ nanotubes and liposomes as lipid model membrane.

##### Short description

The interaction between $TiO_2$ nanotubes and lipid membranes has been evaluated employing liposomes as lipid model system. The TEM characterisation shown a clear interaction between the nanotubes and liposomes.

##### Materials and methods

All materials were purchase from Sigma-Aldrich unless stated otherwise. DOPC lipids were purchased from Avanti Polar Lipids Inc. (USA) in frozen powder form. They are manufactured to 99.9% purity and dissolved in chloroform (CHCl3) for easy stock dilution and use. Extrusion kit including polycarbonate filters were also purchased from Avanti Polar Lipids Inc. (USA).

Liposomes were fabricated using Freeze-Thawing cycles and extrusion. A lipid film (~ 5 mg) has been deposited into a glass vial and dried in $N_2$ stream and placed in a vacuum chamber overnight for complete CHCl3 removal. The film was hydrated the following day with DI water to a final concentration of 2 mg $ml^{-1}$. The lipid dispersion underwent freezing-thawing cycles for 5 times and then extruded 21 times using a 200 nm polycarbonate filter.

The $TiO_2$ nanotubes were dispersed in water and the dispersion sonicated in water bath for ~3h before use.

5 µl of sample was deposed onto glow-discharged copper grid (Agar scientific Ltd, UK) for one minute and the excess of sample was removed blotting with filter paper. After shortly washing with one drop of water, the grid was therefore immersed into a 2% uranyl acetate (UA) solution for 20 s and blotted again with filter paper. The grids were imaged using a JEOL JEM-2100F fitted with a Gatan Orius SC 1000 camera (2x4k).

##### Results

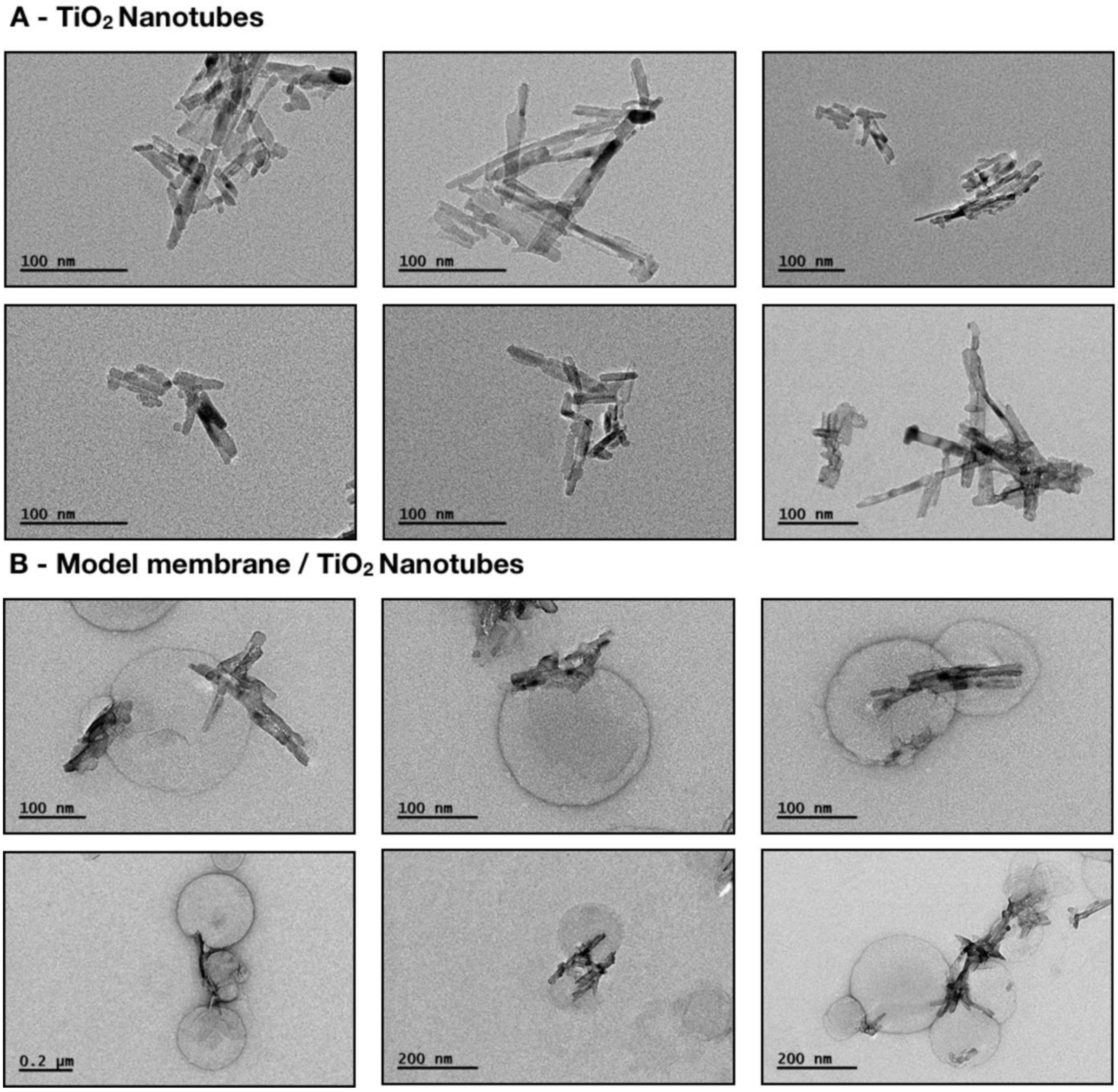

*Figure S8: TEM micrographs of (A) $TiO_2$ nanotubes and (B) lipid model membranes interacting with $TiO_2$ nanotubes.*

### S0d – Uptake of nanomaterial

#### Main message

After exposure, the $TiO_2$ nanotubes are taken up into the LA-4 cells and have been colocalised with endosomes, lysosomes and other vesicles.

Supporting raw and analysed data:

main images:

Figure S9-Figure S10

controls and statistics:

Figure S11-Figure S14

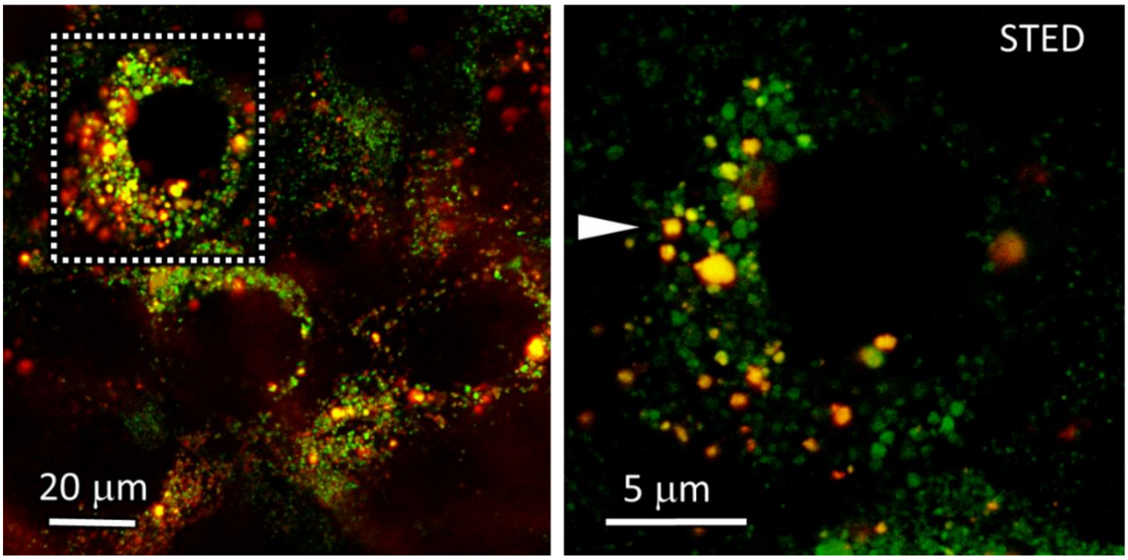

*Figure S9: Confocal (left) and STED (right) colocalisation of $TiO_2$ nanotubes (Alexa Fluor 647, red) and endosomes (green, pHrodo Red Transferrin conjugate) after 30 minutes incubation*

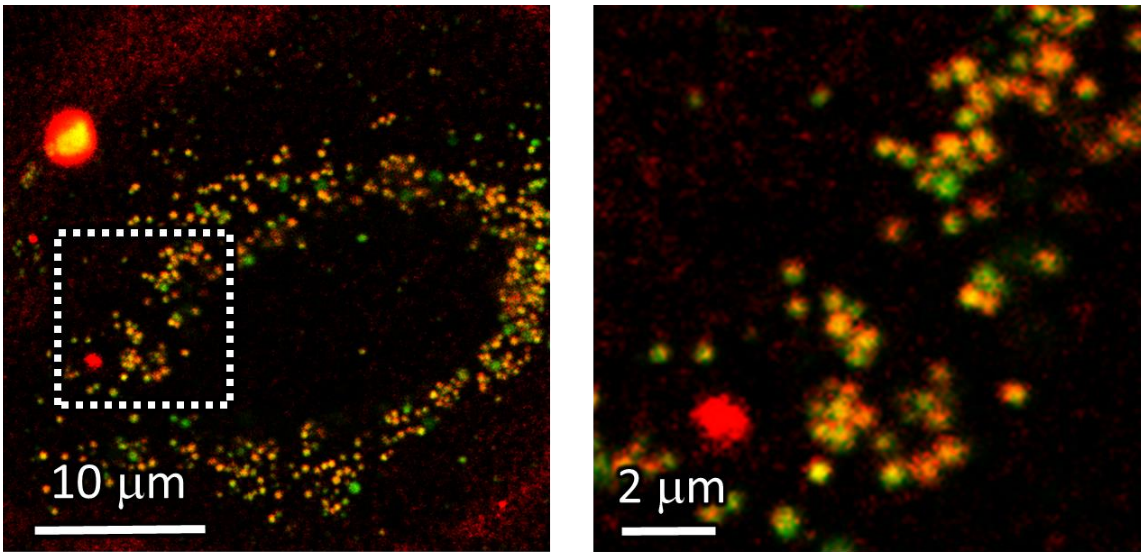

*Figure S10: Colocalisation of $TiO_2$ nanotubes (Alexa Fluor 647, red) and lysosomes (Lysotracker Red, green) after 70 minutes of incubation.*

#### Co-localisation of endosomes and $TiO_2$

controls and statistics for endosome colocalisation:

Figure S11-Figure S13

##### *Materials and methods*

- Experiment:
  - LA-4 cells were seeded @30% confluence in an Ibidi 1.5H µ-Dish.
  - After 48 hours LA-4 cells were washed with warm F12-K medium and placed on ice for 10 minutes.
  - Next, cells were washed 3 times with cold Live Cell Imaging Solution (LCIS) containing 20 mM glucose and 1% BSA.
  - 400 µL pHrodo Red Transferrin conjugate at 50 µg/mL in LCIS containing 20 mM glucose and 1% BSA was added.
  - After 30 minutes incubation at 37°C, 35 µL freshly filtered 1 mg/mL $TiO_2$-Alexa647 in 100x dcb was added directly to the cells and mixed to achieve a 10:1 surface dose

- Analysis:
  - Confocal: logarithmic scale in red channel, cut-off at 5 counts; set maximum to 100 counts on green channel
  - STED: logarithmic scale in both channels, in red channel cut-off at 2 counts, set maximum to 50 counts; on green channel cut-off at 2 counts, set maximum to 75 counts;

| | |
|---|---|
| pixelsize (x,y) | 100 nm |
| FOV (x,y) | 80 µm |

| | |
|---|---|
| 561nm | 30% |
| 640nm | 30% |
| STED | - |
| filter sets | 605 nm – 625 nm, 650 nm – 720 nm |
| dwell time | 10 µs |
| objective | wi 60x (NA0.3) |

| | |
|---|---|
| Cell line | LA-4 (endosomes, pHrodo Red Transferrin Conjugate) |
| NPs | TiO2 (Alexa647) |
| exposure | 10:1, 0min-60min |
| imaging | 0min-30min<br>xyz confocal, 30min |

| | |
|---|---|
| pixelsize (x,y) | 30 nm |
| FOV (x,y) | 37 µm |
| pixelsize (z) | |
| FOV (z) | |
| imaging time | |
| number of frames | |

| | |
|---|---|
| 561nm | 40% |
| 640nm | 40% |
| STED | 20% |
| filter sets | 605 nm – 625 nm, 650 nm – 720 nm |
| dwell time | 10 µs |
| objective | wi 60x (NA1.1) |

##### *Experiment names*

- Main experiment name:
  - 2018_12_06/e01_s02_t03_POSTIVE_CONTROL_LA-4_50ug_ml_pHrodo_30min_NP-Alexa647_1to10.msr
- Supplement names:
  - 2018_12_06/e01_s02_t03_POSTIVE_CONTROL_LA-4_50ug_ml_pHrodo_30min_NP-Alexa647_1to10.msr

    - 2018_12_06/e01_s02_t03_POSITIVE_CONTROL_LA-4_50ug_ml_pHrodo_30min_washed_NP-Alexa647
    - 2018_12_06/e01_s03_t04_POSITIVE_CONTROL_LA-4_50ug_ml_pHrodo_30min_washed_NP-Alexa647
    - 2018_12_06/e01_s05_t05_POSITIVE_CONTROL_LA-4_50ug_ml_pHrodo_30min_washed_NP-Alexa647
    - 2018_12_06/e04_s01_t01_POSTIVE_CONTROL_LA-4_50ug_ml_pHrodo_30min_washed_NP-Alexa647_1to10
- *Control:*
    - 2019_11_22/e03_s02_t03_LA-4_pHrodo_unwashed_25min

##### *Controls and statistics*

Colocalisation of nanomaterial with endosomes after 30 minutes incubation

Figure S11-Figure S12

Control for crosstalk of the pHrodo probe into the nanomaterial channel

Figure S13

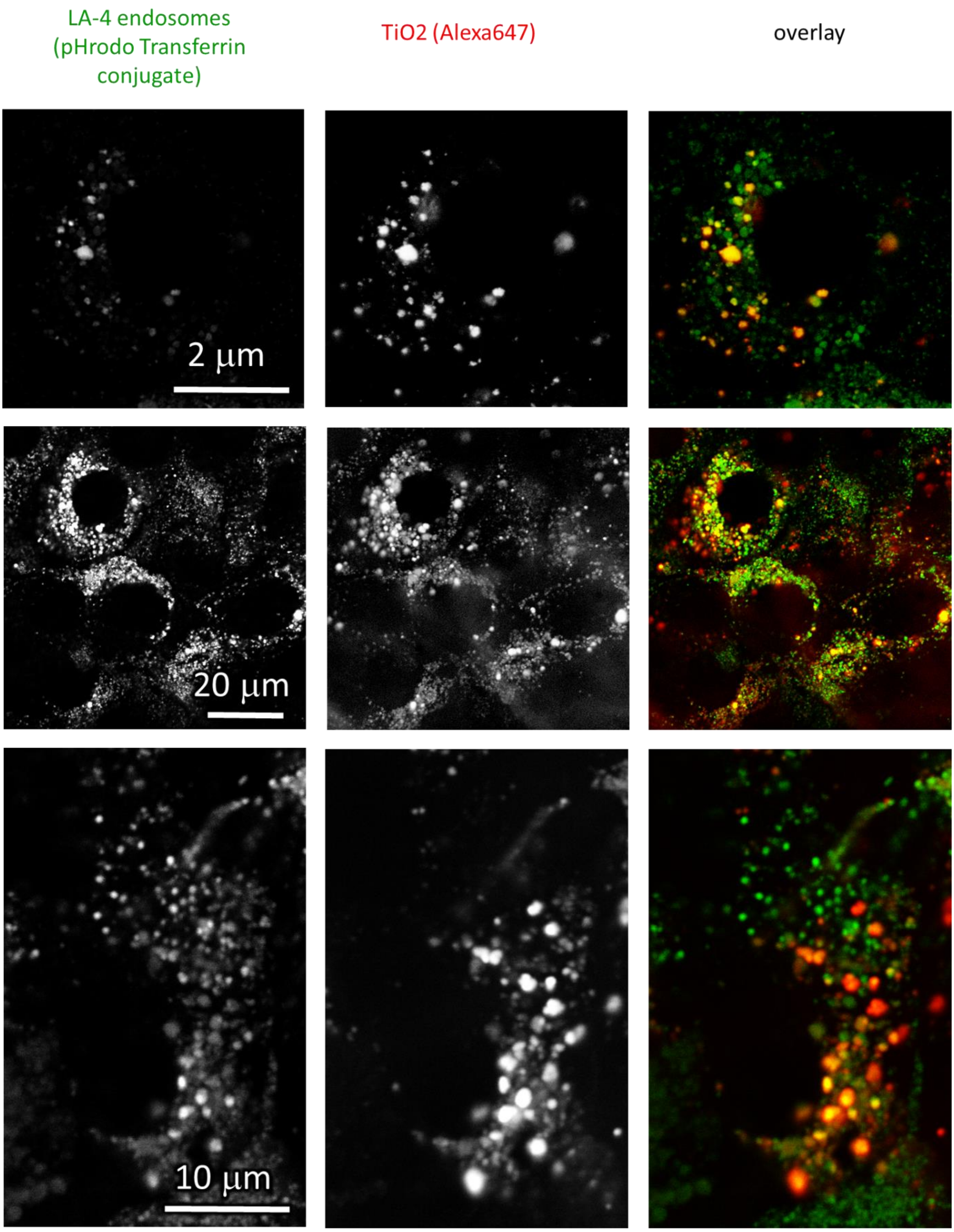

*Figure S11: Statistics for colocalisation of nanomaterial with endosomes after 30 minutes incubation.*

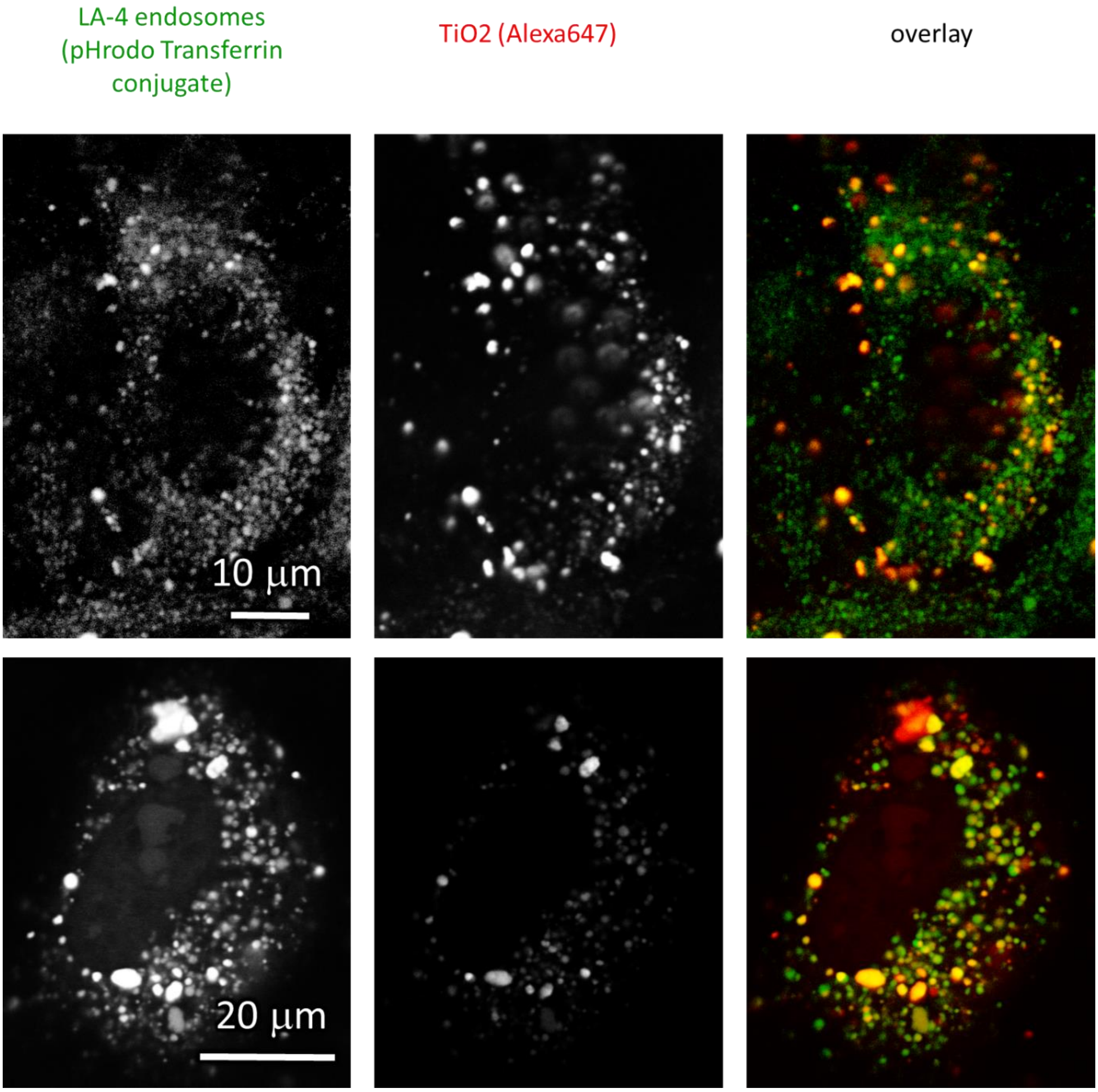

*Figure S12: Statistics for colocalisation of nanomaterial with endosomes after 30 minutes incubation.*

**CONTROL: LA-4 + pHrodo Transferrin**

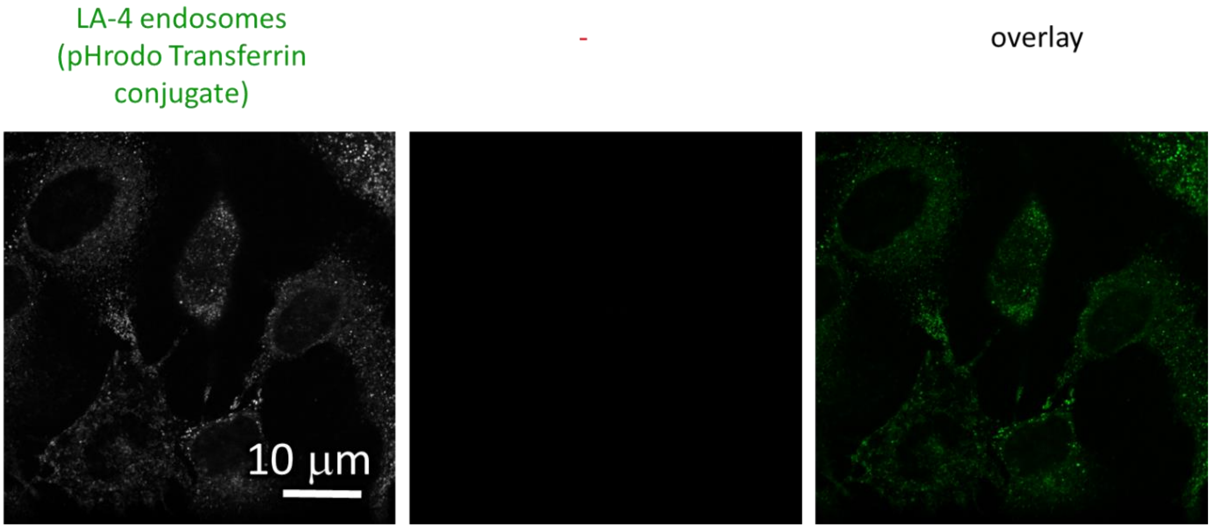

*Figure S13: Control for possible false colocalisation of nanomaterial with endosomes after 30 minutes incubation due to crosstalk of the pHrodo probe into the nanomaterial channel.*

#### Colocalisation of lysosomes and $TiO_2$ nanotubes

Lysosome colocalisation:

Figure S14

##### *Materials and methods*

- experiment 20181115/e03_s01_t04_LA-4 lysotracker TiO2Alexa647.msr:
  - LA-4 cells were seeded @30% confluence in an Ibidi #1.5H µ-Slide 8-well
  - after 3 days the cells were incubated with 150 µL 50nM Lysotracker in LCIS for 2h 20minutes. Afterwards, the medium was exchanged with 200 µL LCIS
  - 1 µL freshly filtered 1mg/ml $TiO_2$-40-Alexa647 in 100x dcb was added to cells to achieve a 1:1 surface dose of nanomaterial. The cells were continuously filmed on a heated insert at 32C for 1h 30 minutes
- analysis:
  - rescaling of green channel (max G = 200 counts)
  - rescaling of red channel (max R = 12 counts)

| | |
|---|---|
| Cell line | LA-4 (lysosomes, Lysotracker) |
| NPs | TiO2 (Alexa647) |
| exposure | 1:1, 0h-1h |
| imaging | xyt confocal, 0h-1h |

| | |
|---|---|
| pixelsize (x,y) | 100 nm |
| FOV (x,y) | 45 µm |
| pixelsize (z) | |
| FOV (z) | |
| imaging time | 1h |
| number of frames | 932 |

| | |
|---|---|
| 561nm | 20% |
| 640nm | 20% |
| STED | |
| filter sets | 605 nm – 625 nm, 650 nm – 720 nm |
| dwell time | 10 µs |
| objective | 60x wi (NA1.2) |

##### *Experiment names*

- Main experiment name:

• 20181115/e03_s01_t04_LA-4 lysotracker TiO2Alexa647.msr

##### *Controls and statistics*

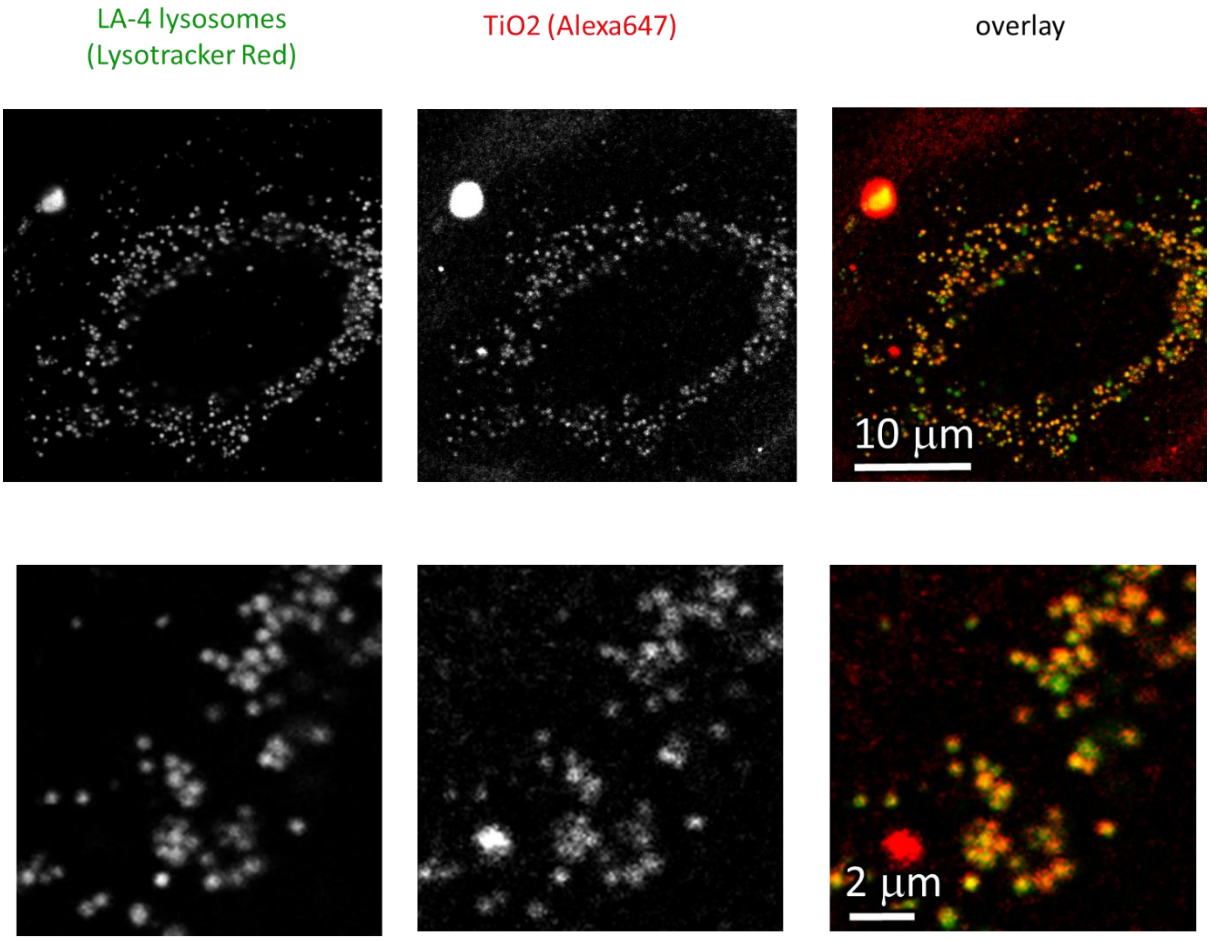

*Figure S14: Colocalisation of lysosomes and nanomaterial after 1 hour of  incubation.*

### S0e – Effect of $TiO_2$ nanotubes on the mitochondrial morphology of the epithelial lung cells

#### Main message

Mitochondria fragment in 3h after exposure to $TiO_2$ nanotubes. After two days the fragmentation is reverted to a state of normal, pre-exposure morphology.
Supporting raw and analysed data:

Figure S15-Figure S29

#### Short description

We evaluated the effects of titanium dioxide nanotubes on fluorescent labelled mitochondrial network of LA-4 murine lung epithelial cells, with fluorescent confocal microscopy and mitochondrial network analysis. LA-4 cells were treated with nanoparticles in 1:1/ 10:1/ 100:1 nanoparticle surface to cell surface ratio and then incubated for 3h, 1 and 2 days. 3h after incubation LA-4 cells with different concentrations of TiO2, the mitochondria were fragmented and closely resembled to positive control (H2O2), in comparison to negative control, where the length of mitochondria were significantly longer. 24 hours after of incubation TiO2NT with LA-4 cells we noticed slightly longer and less fragmented mitochondria in comparison to 3h incubation, which indicates that the mitochondrial morphology started to recover. That was seen especially at low concentration (1:1), while at higher concentration still prevailed fragmented mitochondria. 48 hours after incubation of LA-4 with TiO2NT, mitochondrial morphology became even more similar to negative control, indicating that low concentrations of TiO2NT altered the mitochondrial morphology only transiently. Only when treated with the highest concentration (100:1), the cells still exhibited fragmented mitochondria, but they were longer than those incubated 24h.

#### Materials and methods: Confocal imaging of LA-4 cells

The LA-4 murine lung epithelial cells were seeded into a 35 mm Ibidi μ-Dish and incubated in the complete culturing medium for a day (F12K medium, 15% FCS, 1% P/S (antibiotics), 1% NEAA (nonessential amino acids)). The powder of $TiO_2$ nanotubes was resuspended in distilled water, with addition of 2% of fetal calf serum and 0.5 % absolute ethanol, to the final concentration of 3,24 mg/mL, on ice and in sterile conditions (MISONIX Ultrasound Liquid Processor with 419 MicrotipTM probe for 16 minutes in an ice-bath, setting to the 10 % of the power). Just before treating the cells with nanoparticles in 1:1/ 10:1/ 100:1 nanoparticle surface to cell surface ratio, suspension of nanoparticles was diluted in the cell medium to desire concentration. After 3 hours/ 1 day/ 2 days of incubation, the samples were washed, incubated with 50 nM MitoTracker™ Red CMXRos (ThermoFisher Scientific) for 30 min, followed by observation in the Live cell imaging solution. For positive control 16,3 mM hydrogen peroxide (Merck) was used with incubation time 10 min. For imaging, an Abberior Instruments STED microscope equipped with a 60× water immersion objective was used. Images were acquired at 50 nm pixel size and 0.63 to 1.1 pinhole, depending on the sample. Mitrotracker was excited with the pulsed laser at 561 and fluorescence recorded with an avalanche photodiode within 580−625 nm (filters by Semrock).

#### Experiment names

- Main experiment:
  - 20180925/e01_s01_t01_LA-4_50nM_Mitotracker_3h2D.msr
  - 20180925/e01_s01_t01_LA-4_50nM_Mitotracker_1day_2D.msr
  - 20180925/e01_s01_t01_LA-4_50nM_Mitotracker_2days_2D.msr
- *Controls:*
  - 20180925/e01_s01_t01_LA-4_50nM_Mitotracker_16,3mM_H2O2_10 min_2D.msr

#### Controls and statistics

Representative image for each dose and time point:

Figure S15-Figure S16: Analysis of mitohondria fragmentation through mitochondria size and distribution estimation.

3h incubation:

Figure S17-Figure S21

12h incubation:

Figure S22-Figure S25

48h incubation:

Figure S26-Figure S29

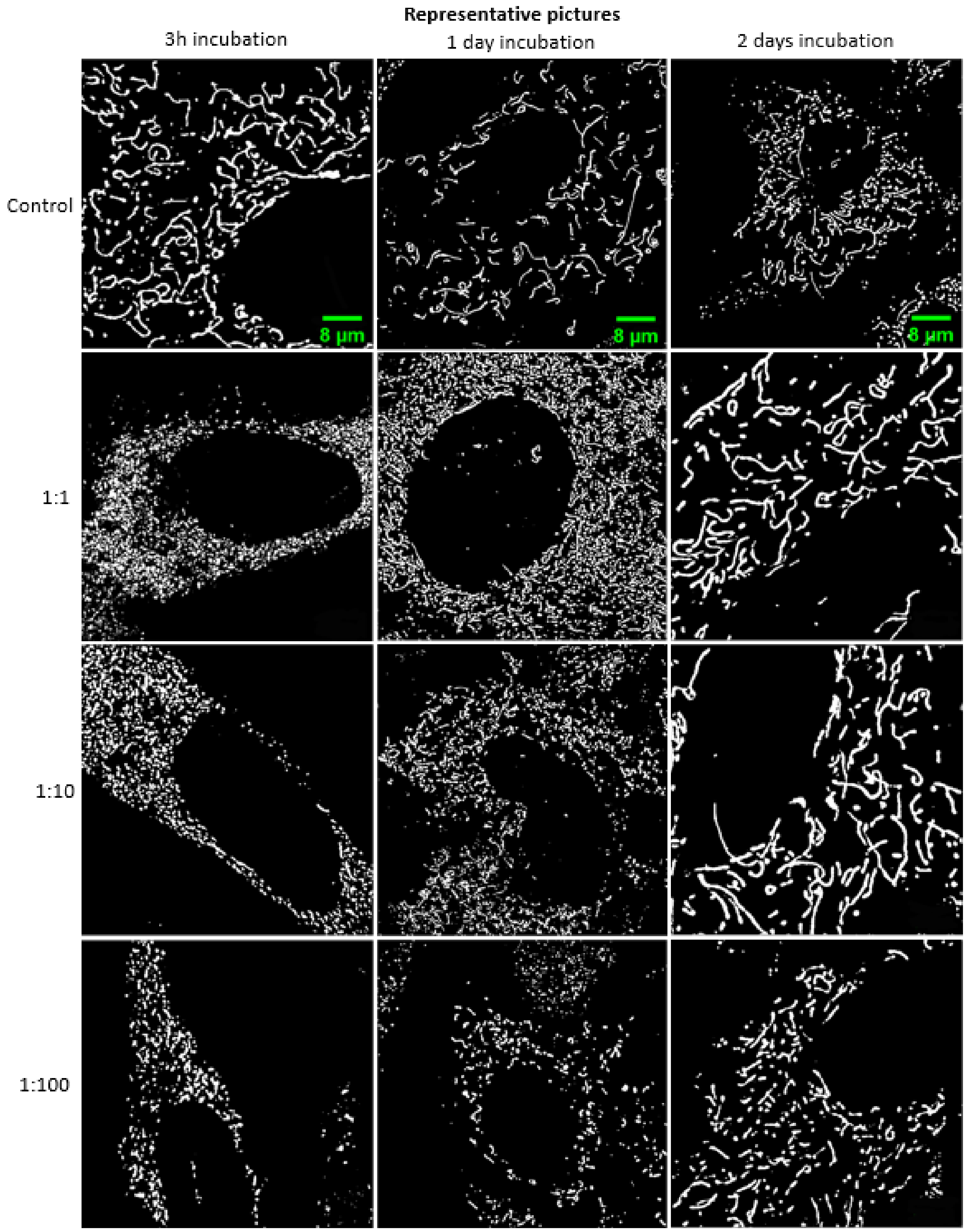

*Figure S15: Representative image for each dose and time point together with control.*

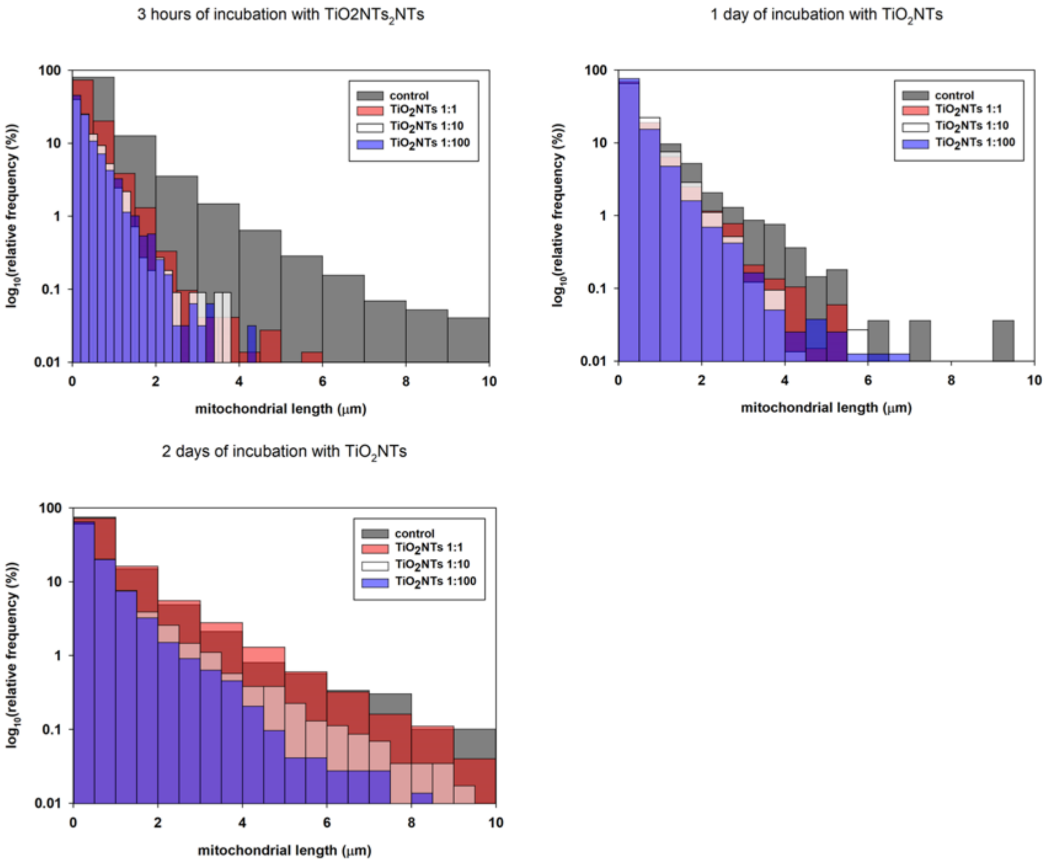

*Figure S16: Analysis of mitohondria fragmentation through mitochondria size and distribution estimation.*

*3 hour incubation of LA-4 cells with $TiO_2$NT*

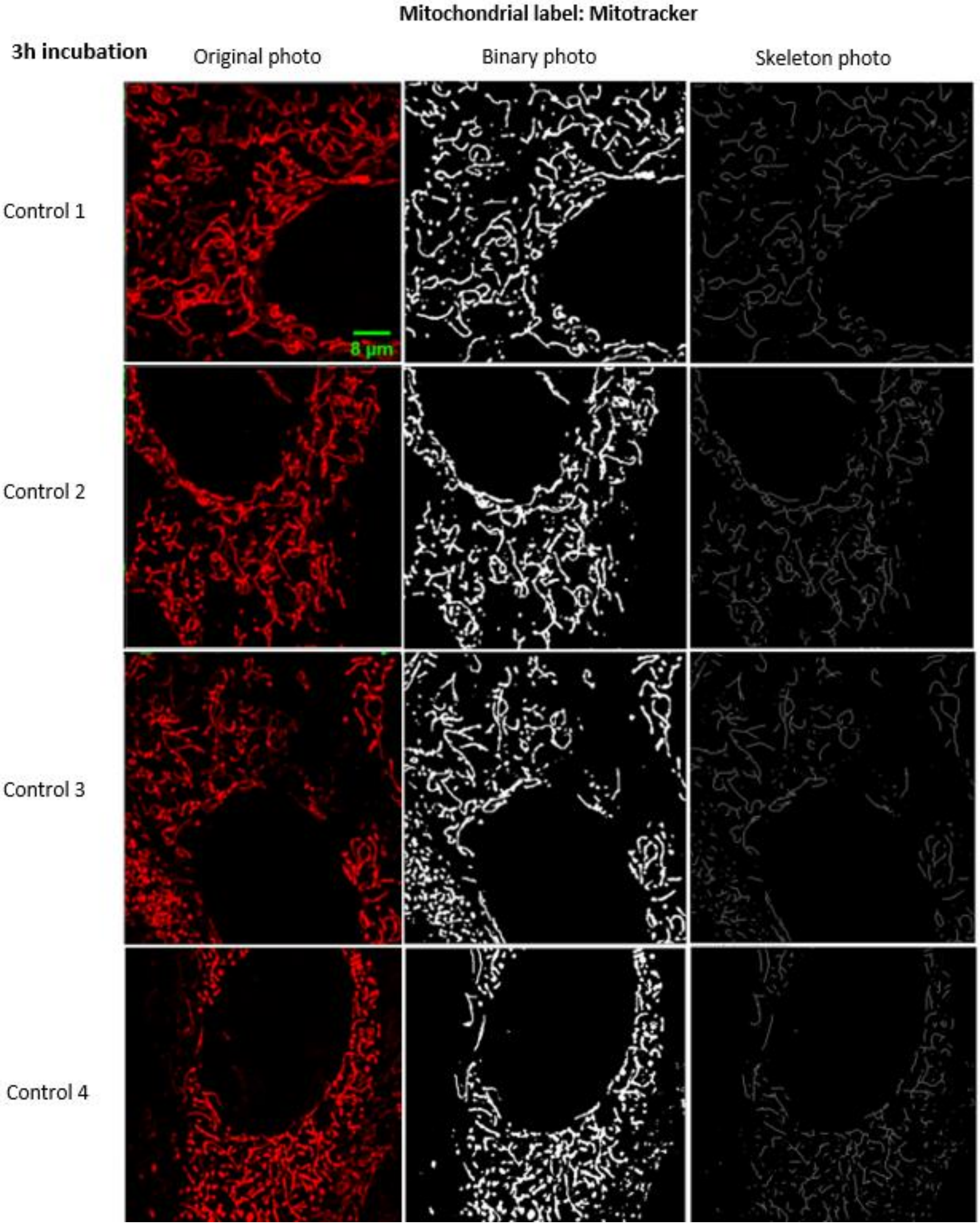

*Figure S17: Statistics for 3h incubation control (non exposed sample) with underlying analysis. Left column is one-channel raw data.*

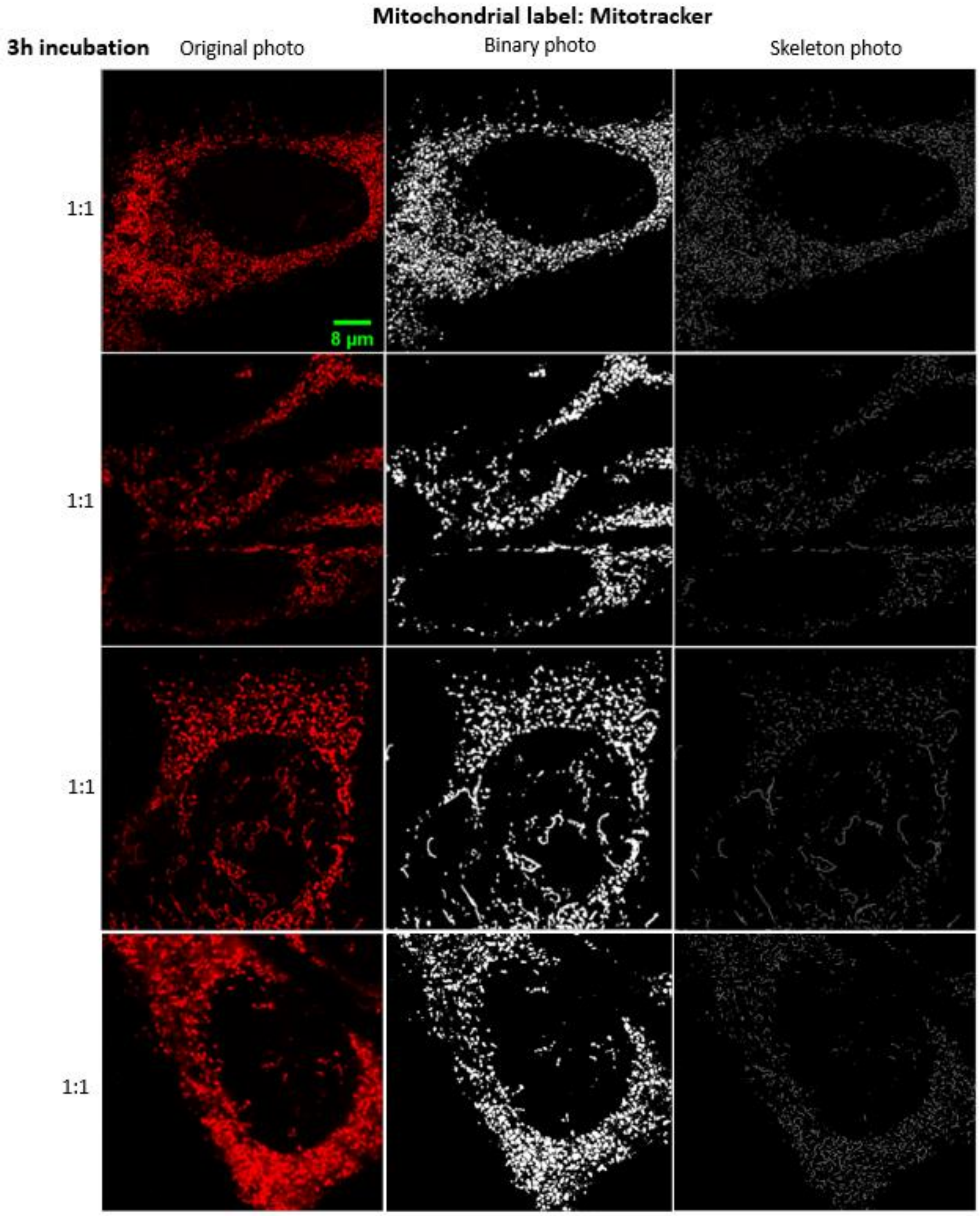

*Figure S18: Statistics  for 3h incubation 1:1 (surface of nanoparticles: surface of cells) dose with underlying analysis. Left column is one-channel raw data.*

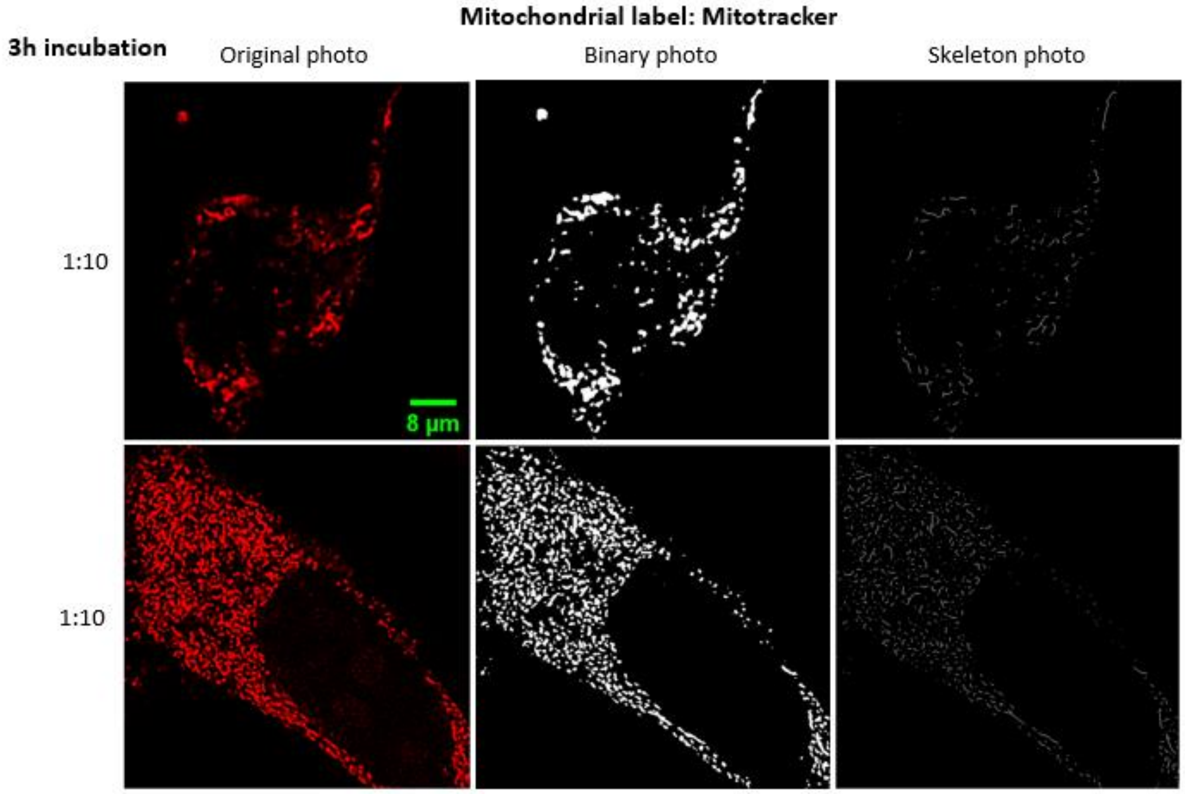

*Figure S19: Statistics  for 3h incubation 10:1 (surface of nanoparticles: surface of cells) dose with underlying analysis. Left column is one-channel raw data.*

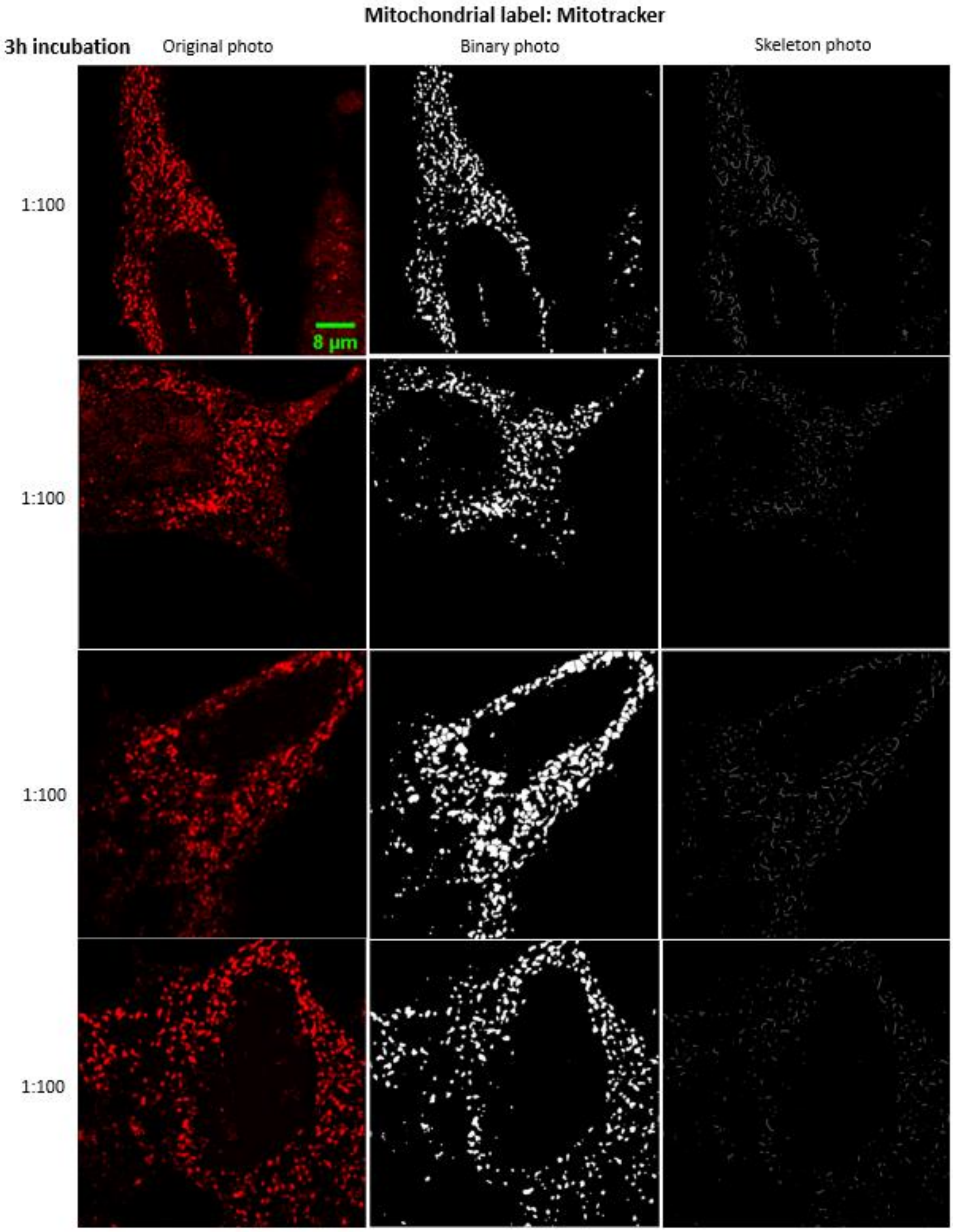

*Figure S20: Statistics for 3h incubation 100:1 (surface of nanoparticles: surface of cells) dose with underlying analysis. Left column is one-channel raw data.*

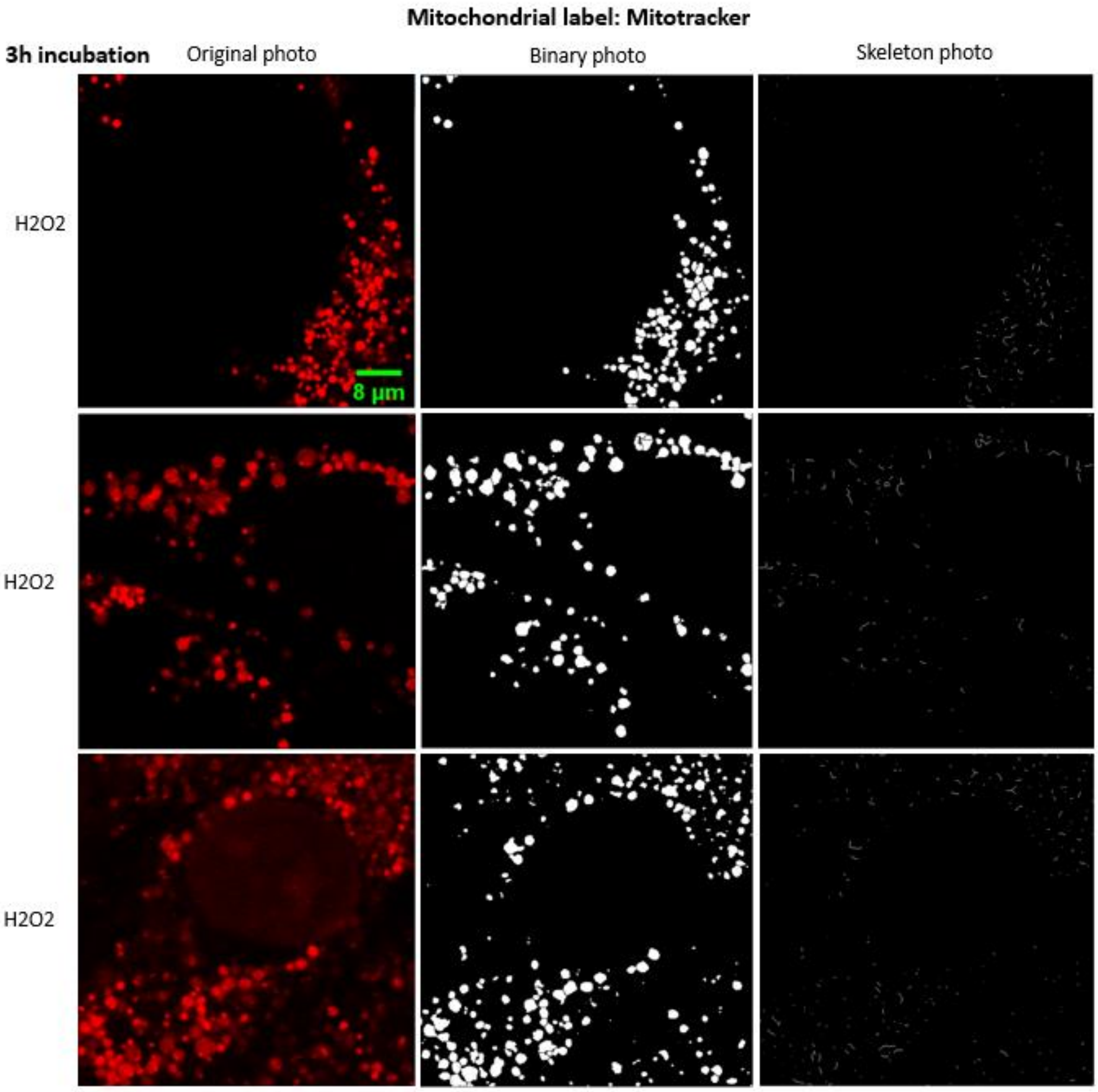

*Figure S21: Statistics for 3h incubation control with $H_2O_2$ with underlying analysis. Left column is one-channel raw data.*

*1 day incubation of LA-4 cells with $TiO_2NT$*

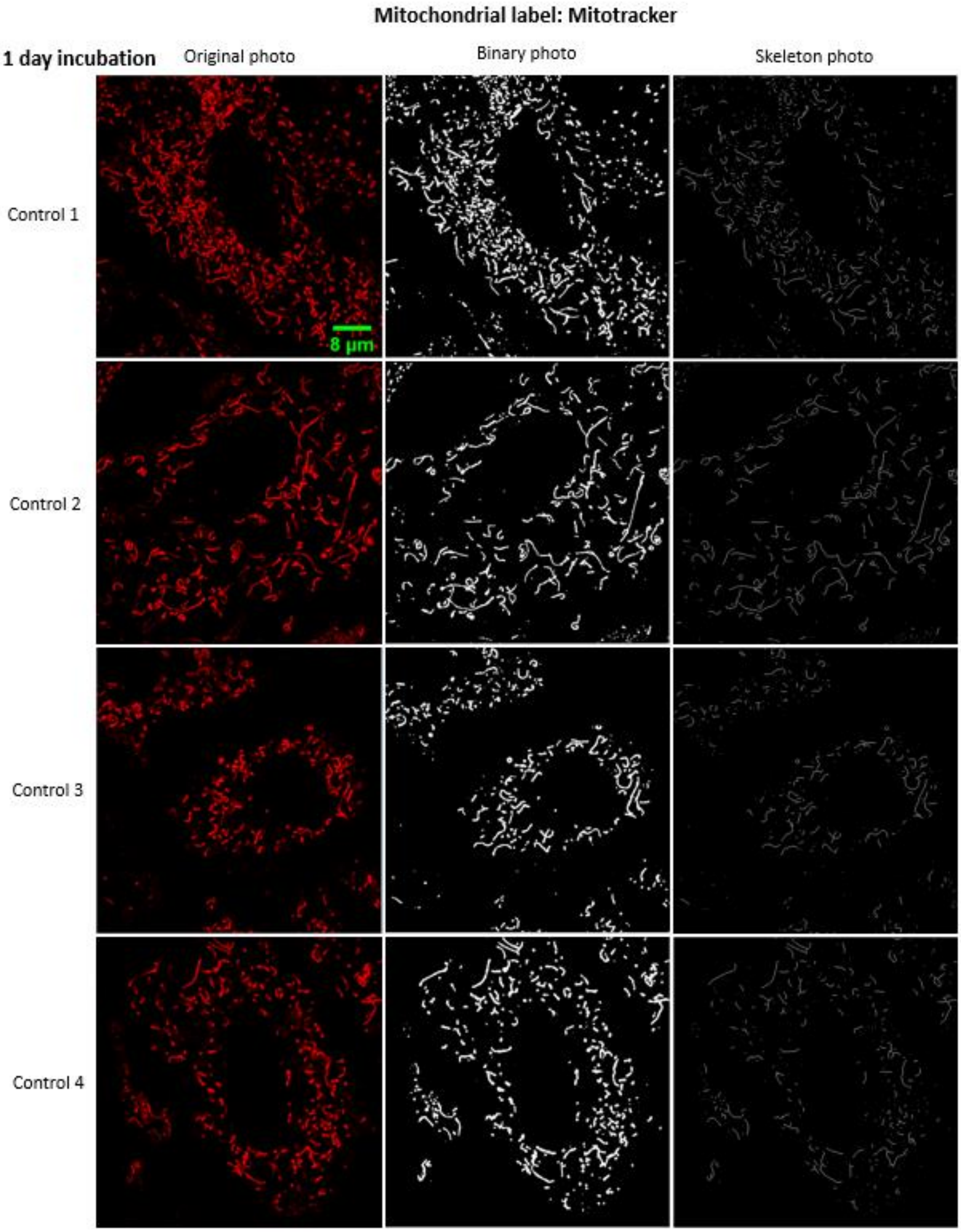

*Figure S22: Statistics for 24h incubation control (non exposed sample) with underlying analysis. Left column is one-channel raw data.*

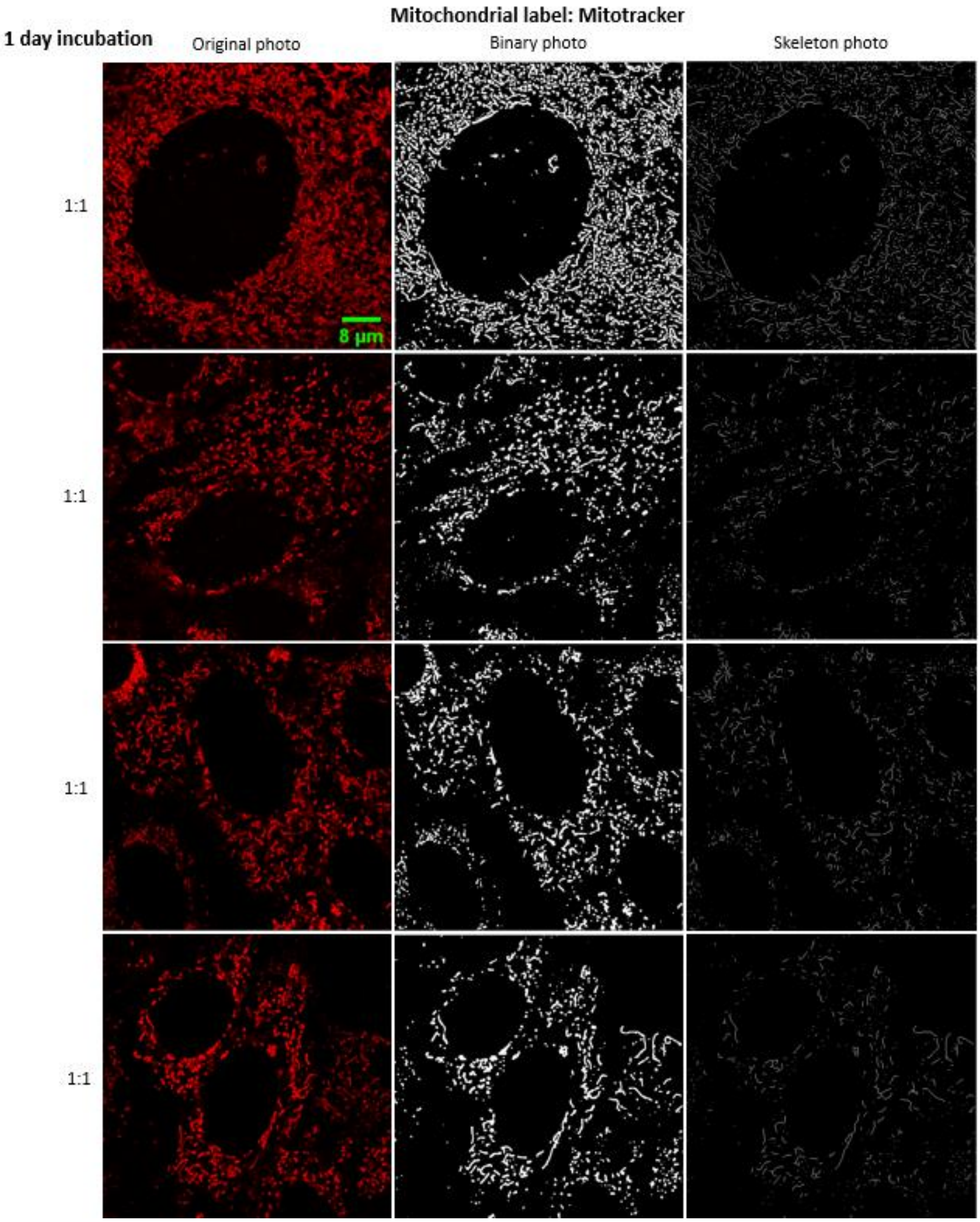

*Figure S23: Statistics  for 24h incubation 1:1 (surface of nanoparticles: surface of cells) dose with underlying analysis. Left column is one-channel raw data.*

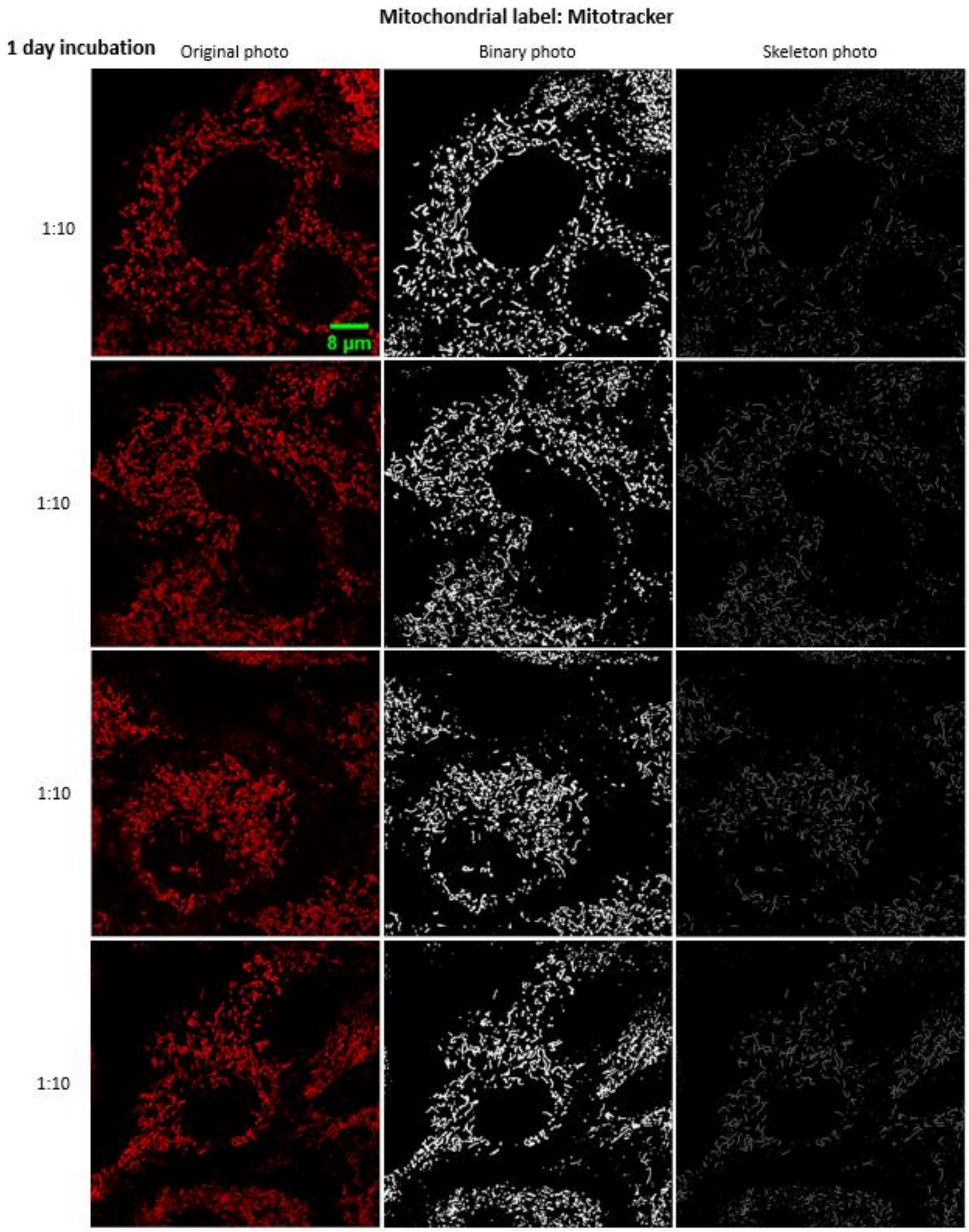

*Figure S24: Statistics for 24h incubation 10:1 (surface of nanoparticles: surface of cells) dose with underlying analysis. Left column is one-channel raw data.*

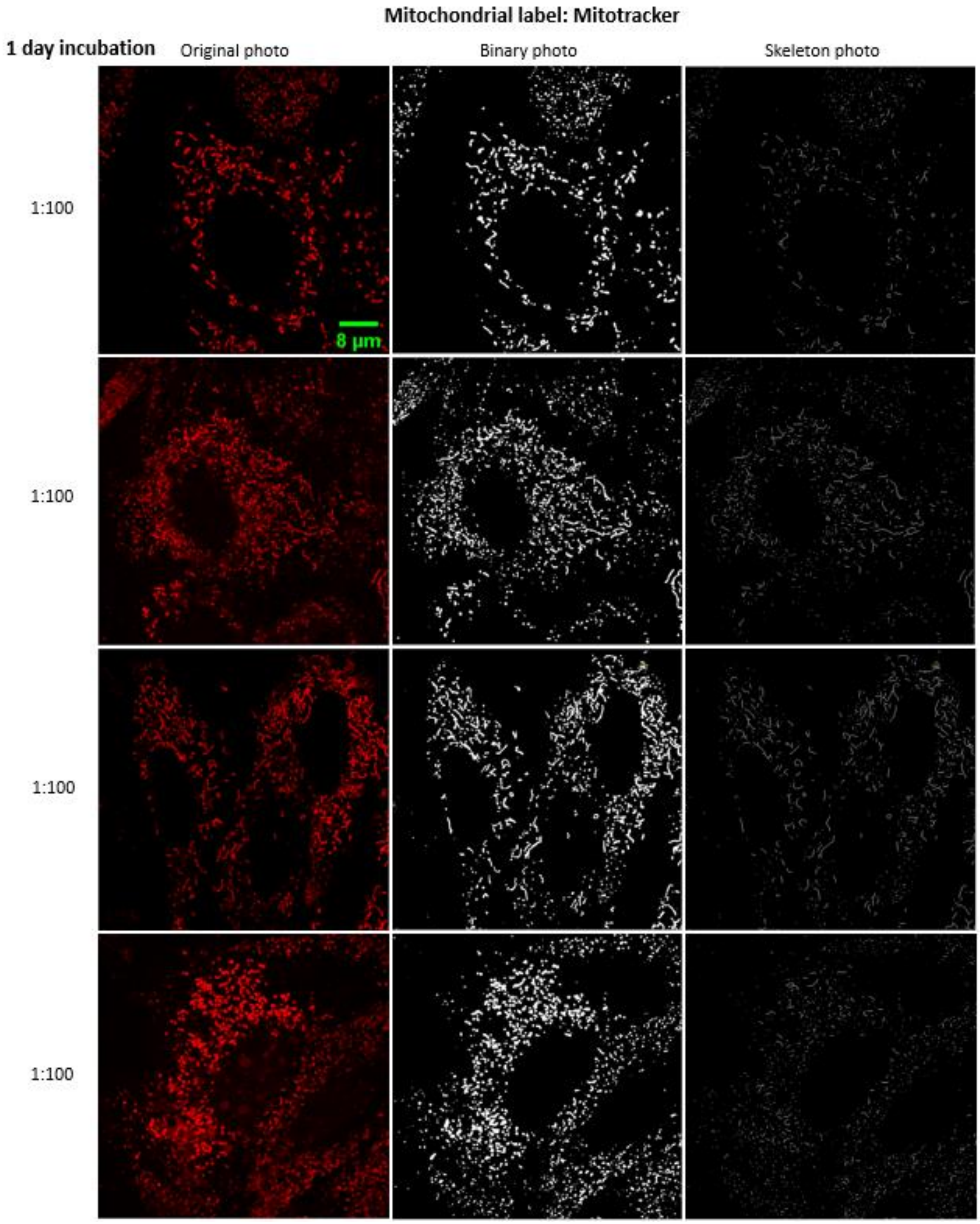

*Figure S25: Statistics  for 24h incubation 100:1 (surface of nanoparticles: surface of cells) dose with underlying analysis. Left column is one-channel raw data.*

#### *2 days incubation of LA-4 cells with $TiO_2$NT*

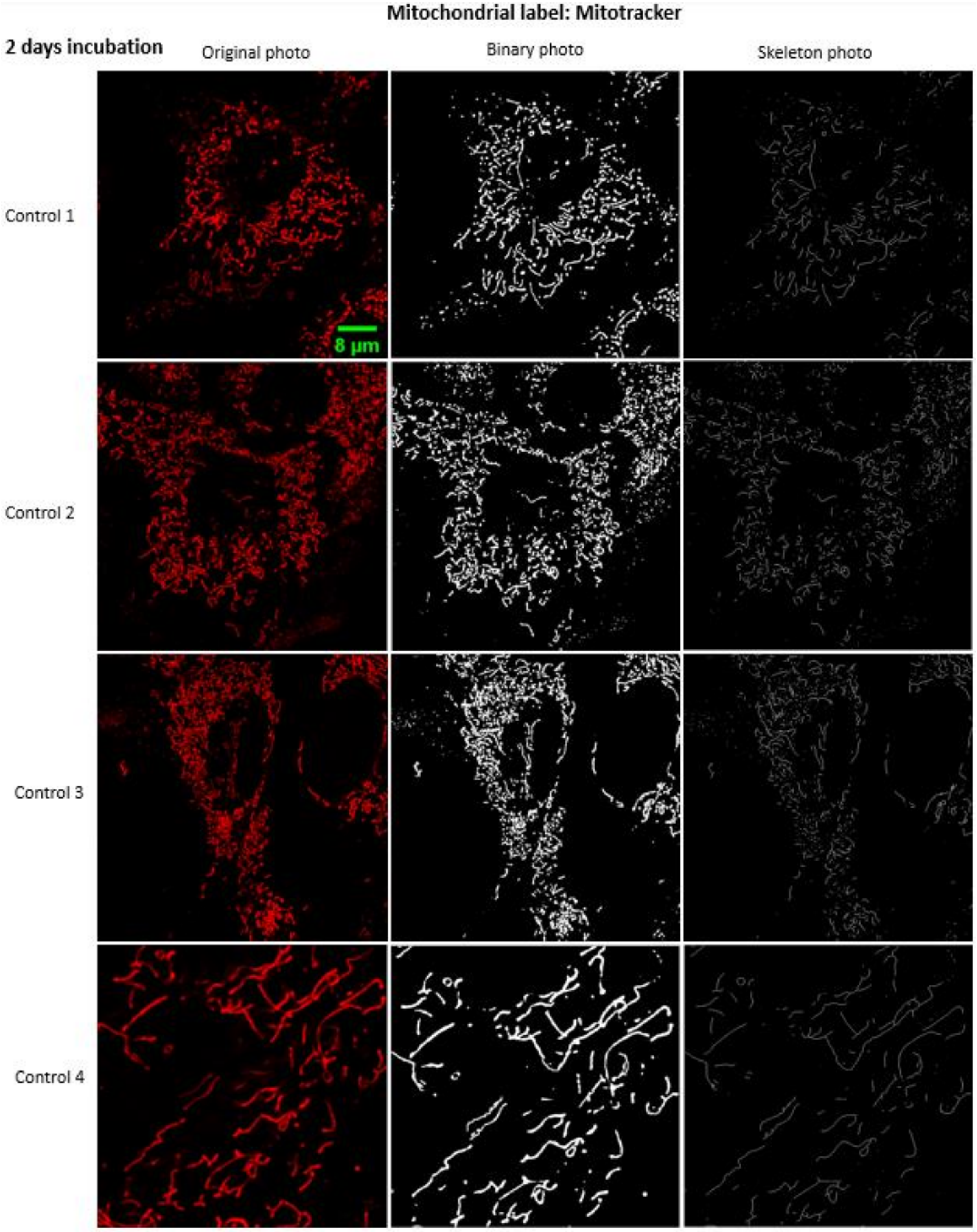

*Figure S26: Statistics for 48h incubation control (non exposed sample) with underlying analysis. Left column is one-channel raw data.*

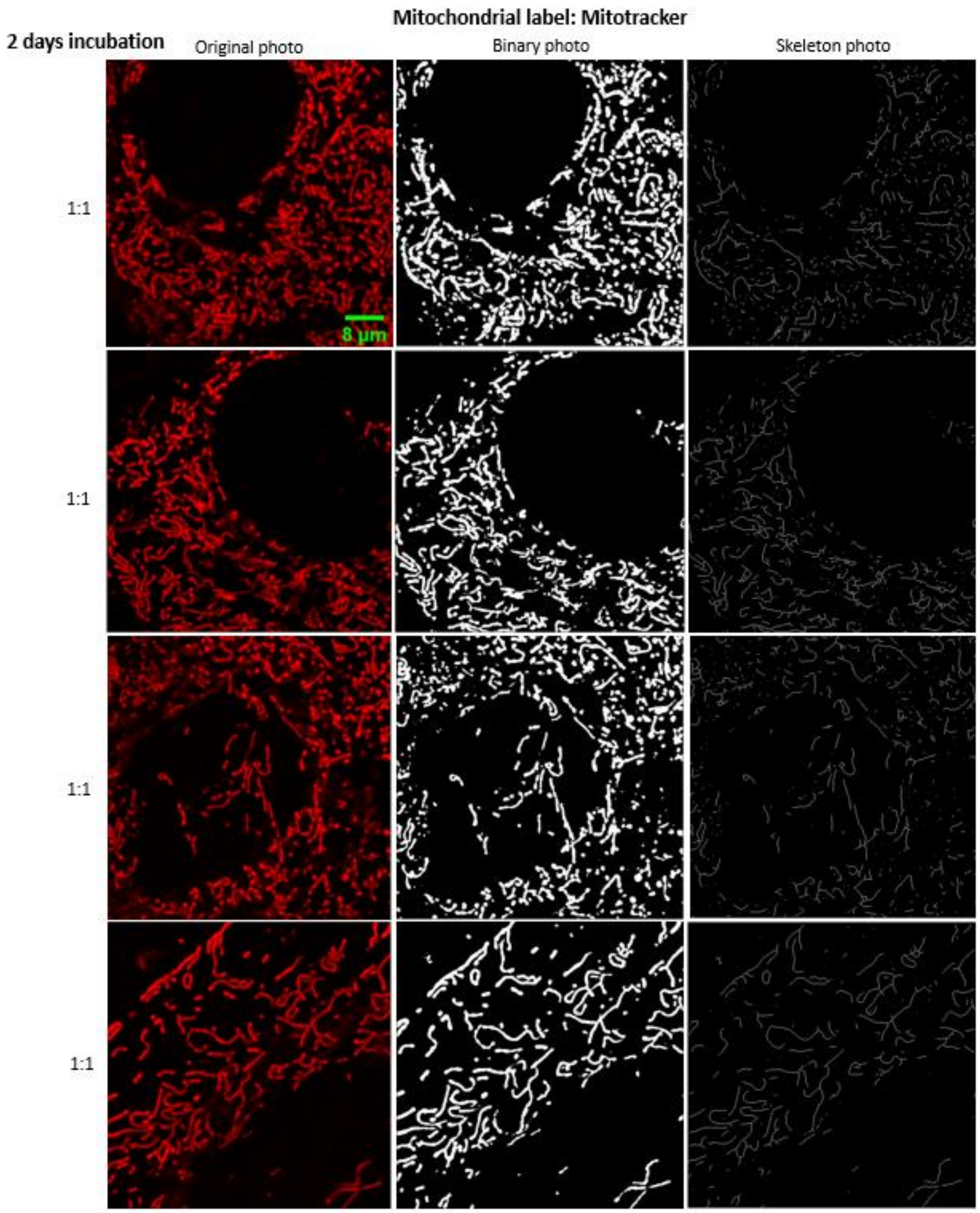

*Figure S27: Statistics for 48h incubation 1:1 (surface of nanoparticles: surface of cells) dose with underlying analysis. Left column is one-channel raw data.*

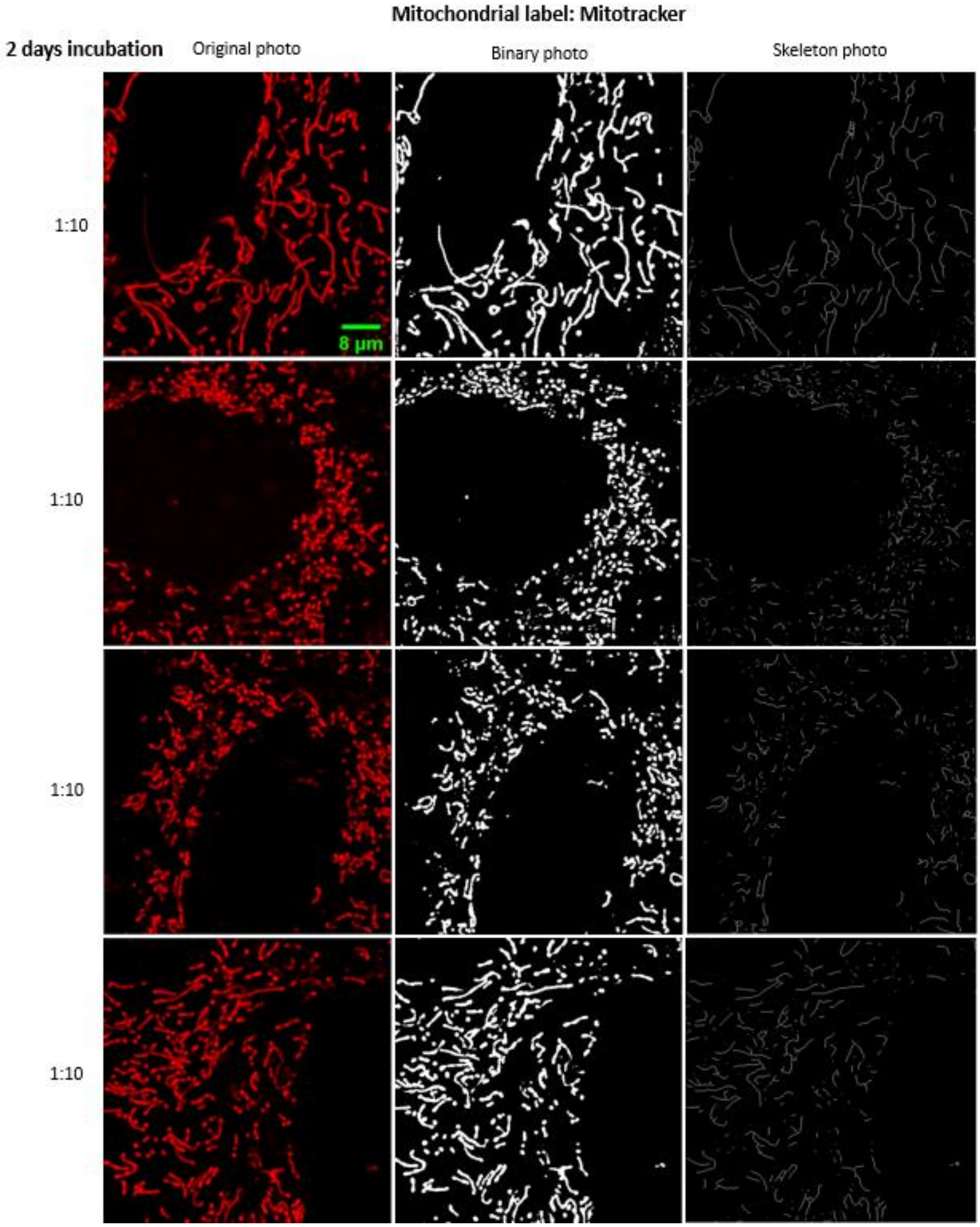

*Figure S28: Statistics for 48h incubation 10:1 (surface of nanoparticles: surface of cells) dose with underlying analysis. Left column is one-channel raw data.*

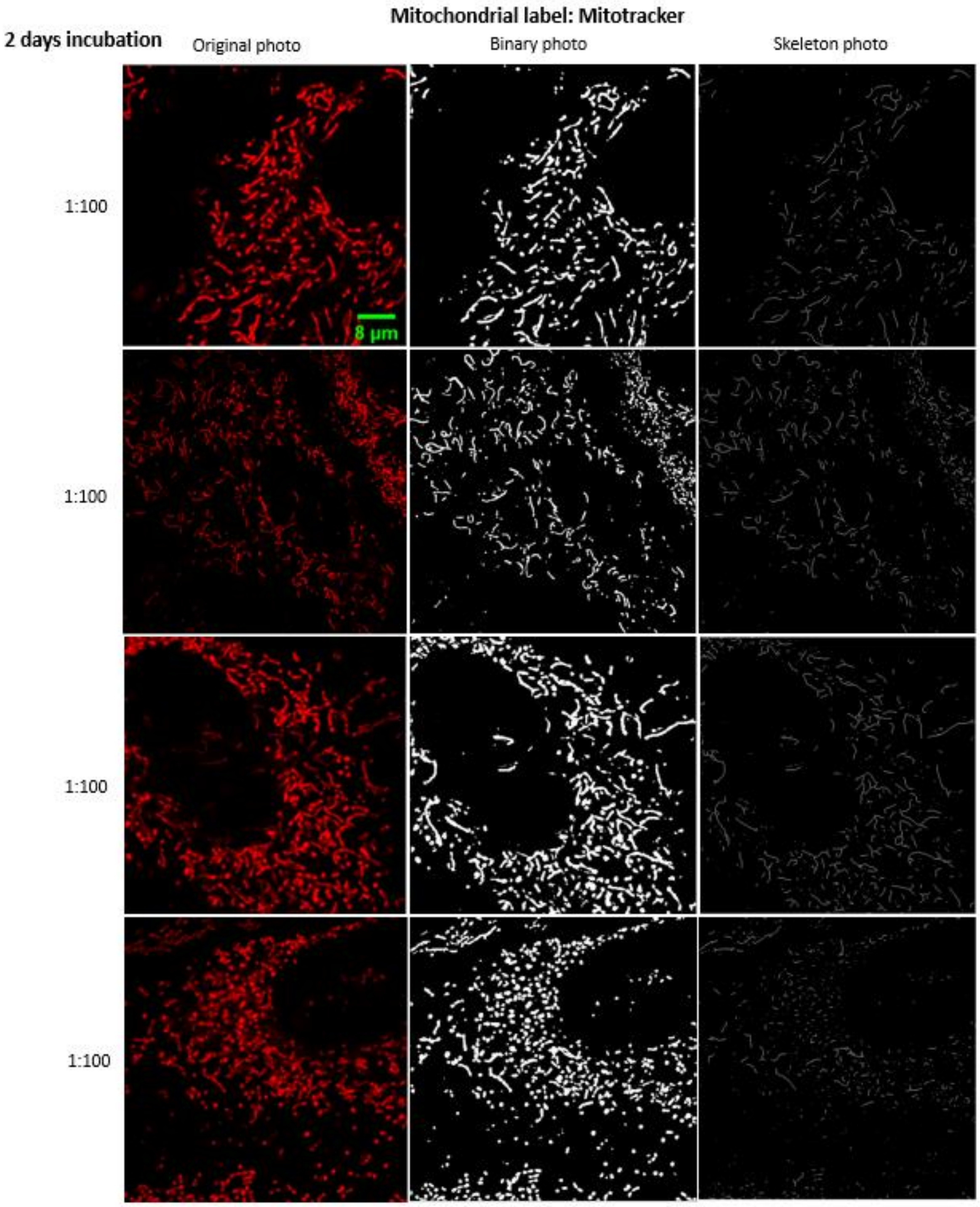

*Figure S29: Statistics  for 48h incubation 100:1 (surface of nanoparticles: surface of cells) dose with underlying analysis. Left column is one-channel raw data.*

#### S0f – Cell viability – $TiO_2$ nanotubes, dose dependence

The cell viability after exposure to increasing doses of $TiO_2$ nanotubes was tested on LA-4 murine epithelial cells and MH-S murine alveolar macrophages using the WST-1 assay as well as on NR8383 alveolar rat macrophages using the LDH and WST-1 assay.

##### Effect of $TiO_2$ nanotubes on the cell viability of lung epithelial cells and alveolar macrophages

###### Main message

Dose response for cell viability was observed in LA-4 and MH-S cells 24h after exposure to $TiO_2$ nanotubes. For higher concentrations of $TiO_2$ nanotubes, for LA-4 cells the cell viability exhibited a plateau effect while MH-S show a continuous decline of metabolic activity. This indicates that epithelial LA-4 cells cope with higher doses compared to the macrophages. The same effect was observed also with crystalline silica (DQ12), and multiwall carbon nanotubes NM-401.

###### Short description

The effects of titanium dioxide nanotubes on viability of LA-4 lung epithelial cells and MH-S alveolar macrophages were compared using WST-1 assay. LA-4 cells and MH-S cells were exposed to titanium dioxide nanotubes in submerged conditions and incubated for 24h with the nanoparticle surface to cell surface ratio of 1:0.39, 1:3.85, 1:19.25, 1:38.5, and 1:385, respectively. 24h after incubation cells with different concentrations of $TiO_2$, the cell viability was measured by WST-1 assay. A significant dose-response was observed with the increased concentration of particles, and there was a plateau effect in the metabolic activity of LA-4 with the nanoparticle surface to cell surface ratio higher than 1:19.25 while MH-S shows a continuous decline of metabolic activity.

###### Materials and methods: Cell viability test of LA-4 and MH-S cells

LA-4 (ATCC# CCL-196) murine lung epithelial cells were cultured in HAMs F12 cell culture medium with stable Glutamin, 1% NEAA (nonessential amino acids), 15% FCS, and 1% Penicillin/Streptomycin. MH-S (ATCC# CRL-2019) murine alveolar macrophages were cultured in RPMI 1640 cell culture medium with stable Glutamin, 0.05mM 2ME (2-Mercaptoethanol), 10% FBS, and 1% Penicillin/Streptomycin. Cells were seeded at a density of $1x10^5$ cells/well into a 24-well plate and incubated in complete culture medium overnight. $TiO_2$ nanotubes were dispersed in distilled water, to the final concentration of 5 mg/mL. The suspension was sonicated on ice with a probe sonifier for 2 min 40 sec at 30% power and 100% cycle time. The nanoparticle suspension was diluted in cell culture medium to concentration of 1, 10, 50, 100, and 1000 ug/mL, which refer to the nanoparticle surface to cell surface ratio of 1:0.39, 1:3.85; 1:19.25; 1:38.5; and 1:385, respectively. Experiments using crystalline silica (DQ12), ZnO and Printex 90 were performed in a similar manner with varying dose ranges. Multiwall carbon nanotubes NM-401 were dispersed in distilled water with 0.2mg/ml real lung surfactant from pig, to the final concentration of 1.5mg/ml. The suspension was sonicated in ultrasonic water bath for 5 min and then sonicated on ice with a probe sonifier for 30 sec at 30% power and 100% cycle time. The nanoparticle suspension was diluted in cell culture medium to desire concentrations just before treating the cells. WST-1 assay was carried out 24h after the particle exposure using cell proliferation reagent WST-1 kit (Roche, Germany) for cell viability measurement. WST-1 solution was either 1:15 (LA-4 cells) or 1:10 (MH-S cells) diluted in complete cell culture medium and incubated with the cells at 37°C for 40 min (LA-4 cells) or 15 min (MH-S cells). 200ul of the culture medium supernatant were removed and centrifuged at 14000 rpm for 10 min after the incubation, and the absorbance (OD value) of the solution was determined at 450nm by microplate reader (Infinite®F200, Tecan). Cell viability of the samples were calculated as followed: cell viability = (sample OD – blank OD)/(control OD – blank OD)*100%.

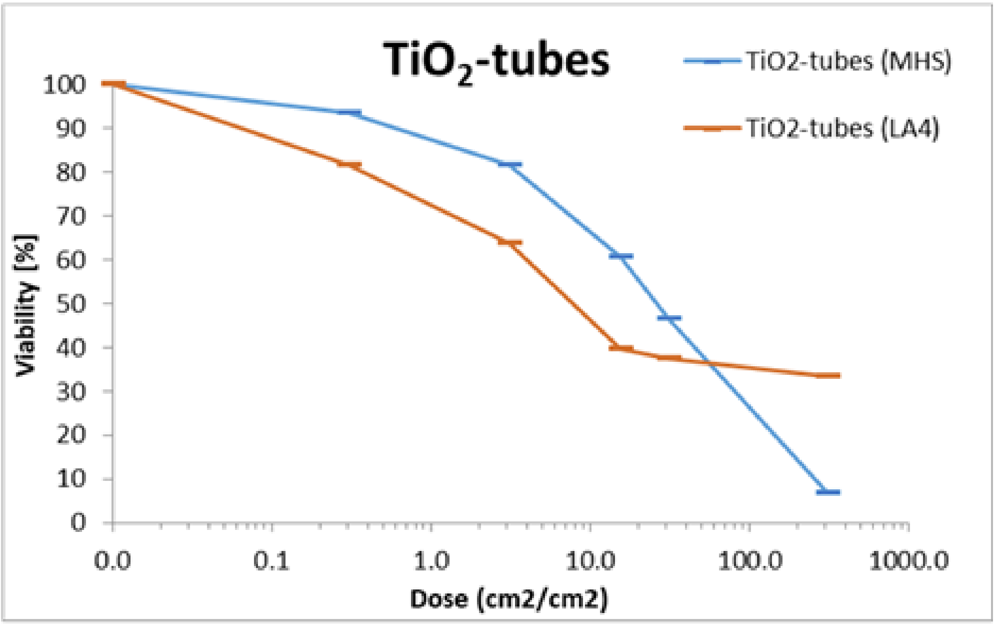

*Figure 30: Dose dependant viability data of exposed MH-S alveolar macrophages and LA-4 alveolar epithelial cells to $TiO_2$ nanotubes in monoculture. Error bars represent the standard error.*

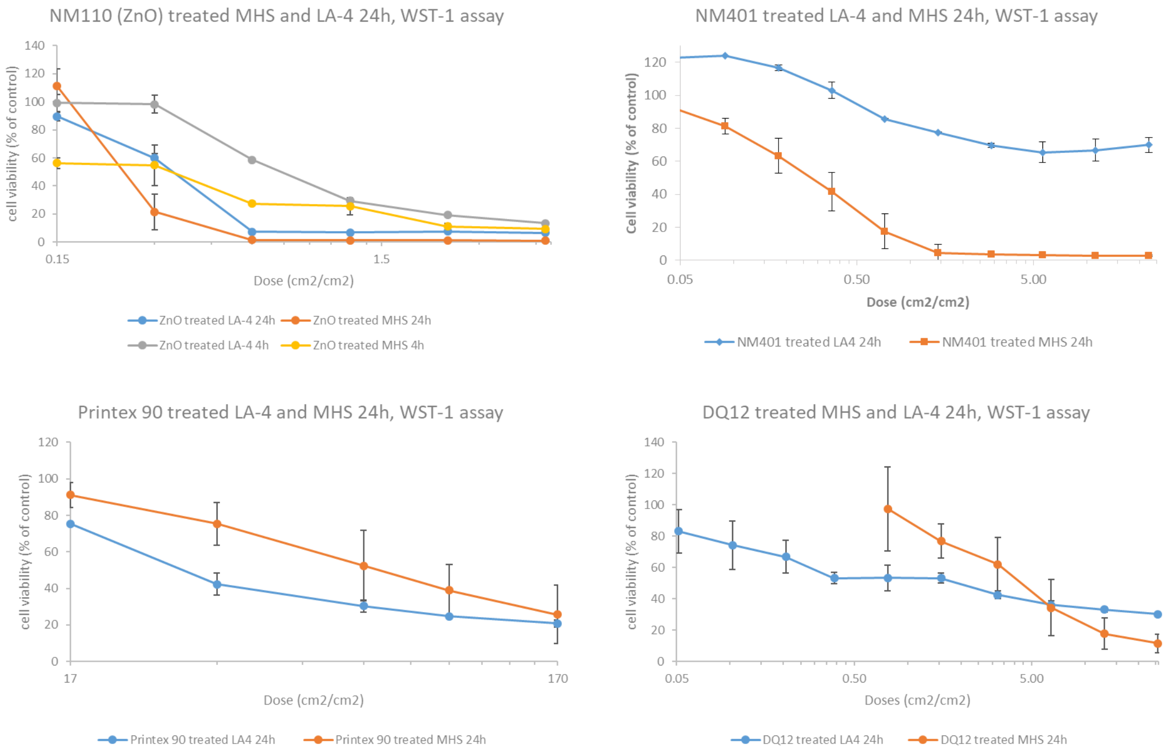

*Figure 31: Dose dependant viability data of exposed MH-S alveolar macrophages and LA-4 alveolar epithelial cells to ZnO (NM 110), multi wall carbon nanotubes (NM401), carbon black (Printex90) and silica (DQ12) in monoculture. Error bars represent the standard error.*

#### NR8383 alveolar rat macrophages, LDH and WST-1 assay, $TiO_2$ nanotubes

##### Main message

$TiO_2$ nanotubes don't show toxicity, even at the highest concentration tested. Cellular metabolism is slightly affected for concentrations until 100 µg/mL.

Supporting raw and analysed data:

Figure S32

Table S 1

##### Materials and methods

###### *Cell culture and exposure*

NR8383 alveolar rat macrophages cell line was obtained from the American Type Culture Collection (ATCC, USA) and were grown in DMEM supplemented with 15 % heat-inactivated FBS, 4 mM L-glutamine (SIGMA-G7513) and a mixture of antibiotic/antimycotic composed of 100 U/mL of penicillin, 100 µg/mL of streptomycin (SIGMA-P0781) and 0.25 µg/mL of amphotericin B (SIGMA-A2942), at 37°C in a humidified mixture of air (95%) and CO2 (5%).

For all experiments, cells were seeded 24 h before exposure to nanoparticles at a density of 5 x $10^4$ cells/mL. Cells were exposed to nanoparticles in cell media without FBS. The different concentrations of nanoparticles were mixed at room temperature to ensure homogeneity of the samples before exposure to cells. Cells not exposed to nanoparticles were served as controls in each experiment. Each experiment was conducted on 4 independent replicates.

###### *Cell viability*

Lactate dehydrogenase (LDH) leakage was analyzed using the LDH assay (Roche-4744934001, Germany) following the manufacturer's instruction. Briefly, NR8383 cells were seeded at 5 x $10^4$ cells/mL in 96-well plates and exposed to different concentrations ranging 2.87 to 91.84 $cm^2$ of $TiO_2$ nanotubes per $cm^2$ of cells ($cm^2/cm^2$). These specific surface concentrations are equivalent of mass concentration of 6.25 and 200 µg/mL. After 24 h of exposure, plates were centrifuged at 800 xg for 10 min and 100 µl of each supernatant were transferred to a new 96-well plate with black bottom that already prefilled with 100 µL of the LDH reaction mixture. Extracellular media was incubated for 30 min at room temperature, then, 50 µL of a stop solution was added and the absorbance was measured at 490 nm on a microplate reader. NR8383 cells treated with 5 % Triton was considered as positive control. Unexposed NR8383 cells were considered as negative control. Nanoparticles cytotoxicity was expressed as the percent of LDH leakage measured in positive control cells. Dose-effect relationships were assessed by ANOVA and Dunett's test. p values < 0.05 were considered significant.

###### *Metabolic activity*

Metabolic activity was assessed using the WST-1 assay [6] (Roche, 11644807001, USA), according to manufacturer's protocol. NR8383 cells were seeded at 5 x $10^4$ cells/mL in 96-well plates and exposed to different concentrations (6,25 to 200 µg/mL, corresponding to 2.87 to 91.84 $cm^2/cm^2$) of $TiO_2$ nanotubes. After 24 h of exposure, WST-1 reagent was added to each well. Cells were incubated at 37 °C for 2 h. The absorbance of the solution was determined at 480 nm on microreader (BioRad-iMARK). IC50 was measured for each nanoparticle according to Reed-Muench method [7] from WST-1 results.

#### Experiment names

- Main experiment name: 2019 09 30 - Oliver - Copie de NanoTube TiO2

#### Controls and statistics

Characterisation of $TiO_2$ nanotubes:

Table S 1

Viability of rat macrophages in dependence of dose:

Figure S32

| Hydrodynamic Radius | PDI | Specific area BET ($m^2/g$) (2) |
|---|---|---|
| 371± 3 | 0.623 ± 0.2 | 152 $m^2/g$ |

*Table S 1: Main characteristics of NanoTube $TiO_2$*

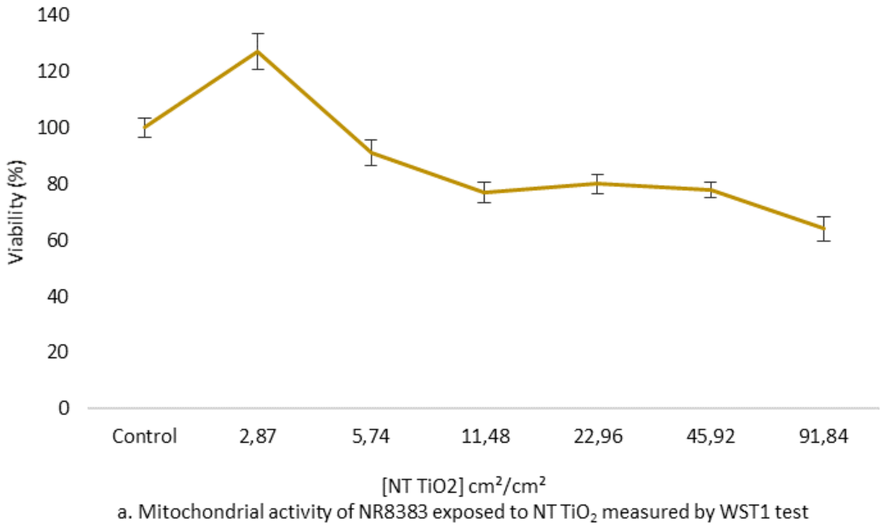

a. Mitochondrial activity of NR8383 exposed to NT $TiO_2$ measured by WST1 test

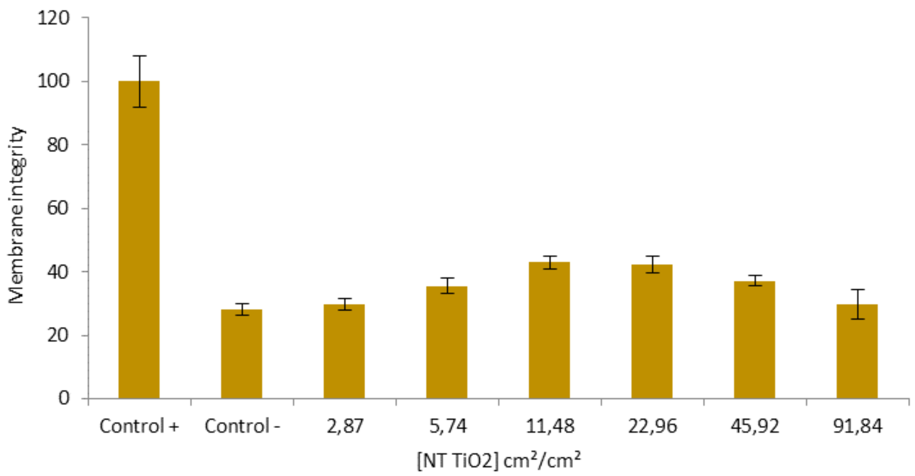

b. Membrane integrity of NR8383 exposed to NT $TiO_2$ measured by LDH test

*Figure S32: Viability of NR8383 exposed to nanoparticles of NanoTube $TiO_2$ in both tests of WST1 (top) and LDH (bottom) Cells exposed to $TiO_2$ nanotubes showed a significant decrease of viability from 5.74 $cm^2/cm^2$. IC50 > 91.84 $cm^2/cm^2$ ( 200 µg/mL).*

### S0g – Cell viability LA-4 – various nanomaterials, same surface dose

#### LA-4 murine alveolar cells, PI and Hoechst assay, various nanomaterial

##### Main message

We tested the toxicity of various nanomaterials to the LA-4 alveolar epithelial cells using the PI/Hoechst assay. ZnO exhibits the highest toxicity on LA-4 cell line. Cell viability after exposure to other nanomaterials remains high, but morphological changes are evident in comparison with the control samples.

The cauliflower-forming potential of the same nanomaterials is examined in "S1c – Cauliflowers with various nanomaterial" and the TEM micrographs are in section "S0b – Nanomaterial characterisation".

Supporting raw and analysed data:

Figure S33

##### Materials and methods

- cell preparation and exposure was performed simultaneously and using the same nanomaterials and protocol as in S1c.
- preparation for imaging:
    - Propidium Iodide (PI) (Sigma, final concentration 0.3 µg/mL) was used to label the nuclei of dead cells and Hoechst 33342 (Sigma, final concentration 10 µg/mL) to label the nuclei of all cells. Cells were imaged in LCIS without washing
- Cells have been imaged with the FMS setup: inverted fluorescence Nikon Eclipse TE 2000-E microscope with Xe-Hg source (Sutter Lambda LS, Novato, CA). Excitation 352-402, dichroic

409 and emission 417-477 broad-band pass filter set have been used to image Hoechst 33342 labelled cells and excitation 503-538, dichroic 560 and emission 596-664 filter set have been used to image PI labelled cells (BrightLine from Semrock, Rochester, NY). Images were taken with EMCCD camera (iXon3 897 from Andor, Belfast, UK)

- Analysis:
  - Each well of the 8-well Ibidi, was imaged on three random locations to get the unbiased overview of the sample. Absolute number of cells in each frame was then determined manually as Hoechst 33342 + and number of dead cells as PI+ and Hoechst 33342 +
  - For the analysis median number of cells per well was used

#### Experiment names

- Main experiment name:
  - 2019_11_29 LA-4 Multiple NP exposure Viability

#### Results

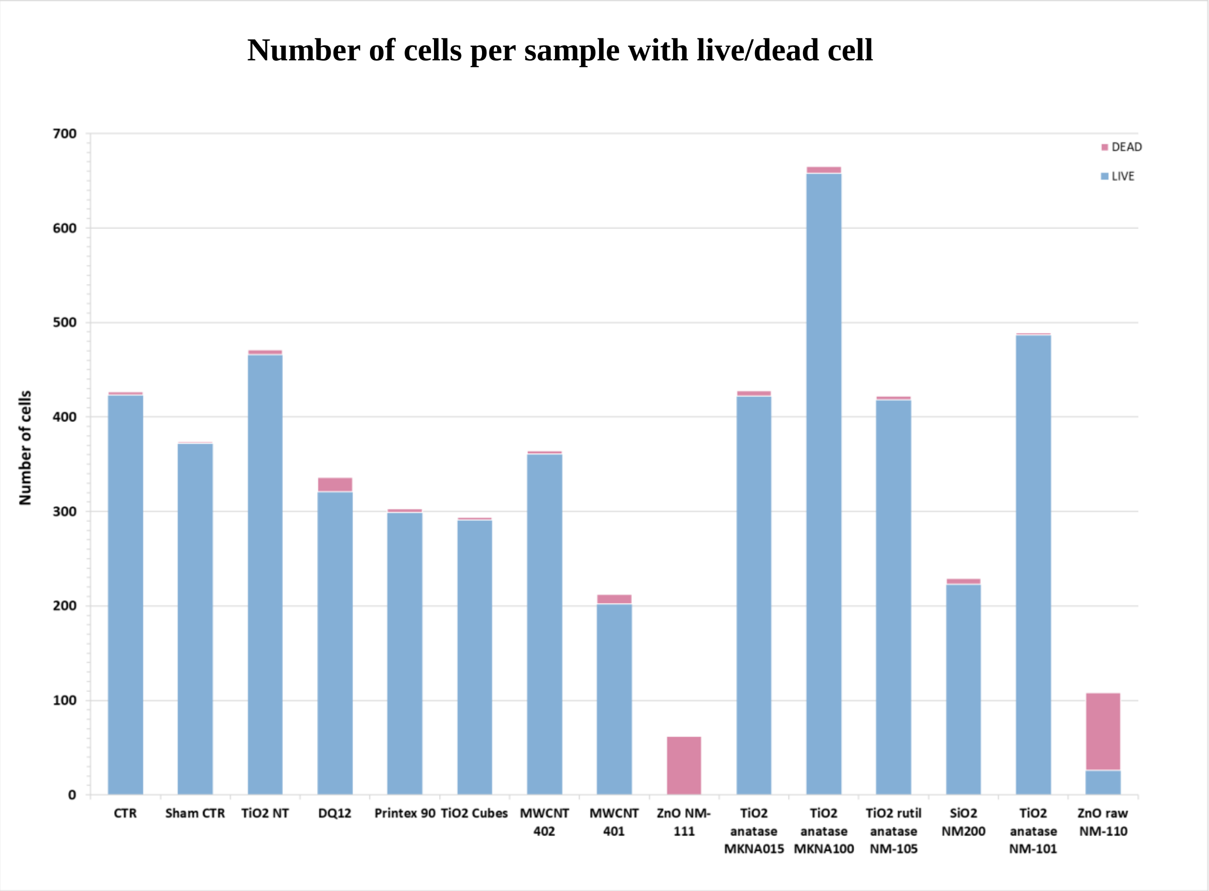

*Figure S33: Median number of LA-4 per field of view in the sample with viability after a 2-day exposure to 10:1 surface dose of nanomaterials, as determined by PI-Hoechst 33342 staining.*

#### S0h – *In vitro* exposure of cocultures of MH-S&LA-4 to $TiO_2$

##### Main message

Tracking of macrophages grown on top of epithelial cell layer through the first 3 days after exposure to see the relative surface cleaned by the macrophages as well as their slow down.

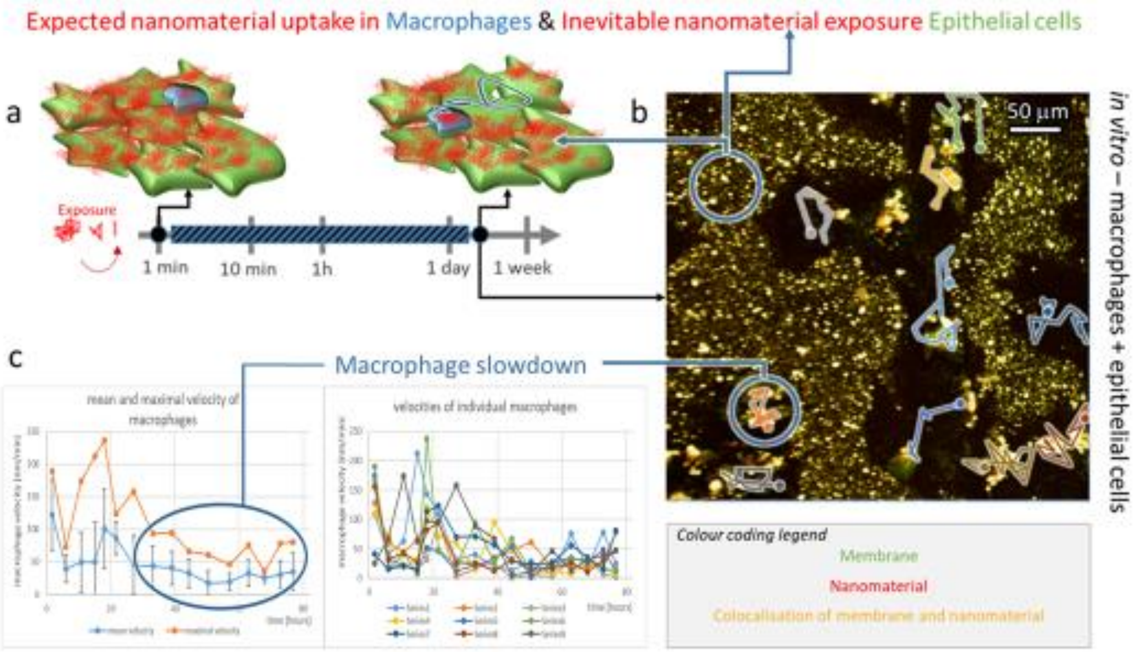

##### Materials and methods

- experiment:
    - LA-4 cells were seeded @30% confluence in an Ibidi #1.5H µ-Dish
    - MH-S cells were seeded @30% confluence in a separate Ibidi #1.5H µ-Dish
    - After 48 hours of separate incubation, LA-4 and MH-S were mixed together. Growth media for cocultures was mixture of F12K and RPMI-1640 in 1:1 ratio
    - after 72 hours 35 µl freshly filtered 1 mg/ml $TiO_2$-17-Alexa647 in 100x dcb was added directly to the cocultures of LA-4 and MH-S cells (in 400 µL mixed medium) and mixed to achieve 10:1 surface dose
    - After approximately 96 hours exposed cocultures of LA-4 and MH-S was incubated with 1 um SHE-2N membrane label for 5 minutes in incubator at 37°C and 5% CO2, afterwards they were flushed with 1x400 ml LCIS and observed in 400 µL mixture of F12K and RPMI-1640 in 1:1 ratio (200 µL) and Live cell imaging solution (LCIS – 200 µL) in the home-made incubator at 37°C for following 72 h

- analysis:
    - The colour intensity scale is changed from linear to logarithmic scale in both green and red channel to get better contrast. Images were analysed in Photo Shop.

    - For the dynamics of movement analysis seventeen images with peak intensity and approximately five hour increments have been chosen for the analysis. They have been aligned in order to perform single cell tracking experiment with ImageJ' Manual tracking software. All data have been exported and analysed in excel (Fig 1.). Manual tracking plugin enables users to overlay lines over chosen dots (MH-S in our case) and thus renders cell trajectories.

#### Experiment names

- 20190517_ e01 m01 s01 LA-4 & MH-S_SHE 2N exposed to TiO2 Alexa 647_72h_FRI-SAT_24h
- 20190517_ e01 m02 s01 LA-4 & MH-S_SHE 2N exposed to TiO2 Alexa 647_72h_SAT-SUN_48h
- 20190517_ e01 m03 s01 LA-4 & MH-S_SHE 2N exposed to TiO2 Alexa 647_SUN_48h_before crash
- 20190517_ e01 m03 s01 LA-4 & MH-S_SHE 2N exposed to TiO2 Alexa 647_MON_72h_ after the crash
- Seventeen representative images, which can be found in 20190517_e01 m01 s01 LA-4 & MH-S_SHE 2N exposed to 10 to 1 $TiO_2$ Alexa 647_72h_Raw folder are: 0, 43, 103, 156, 206, 226, 293, 359, 435, 500, 566, 643, 725, 784, 846, 902, 940

#### Controls and statistics

##### Figure S34

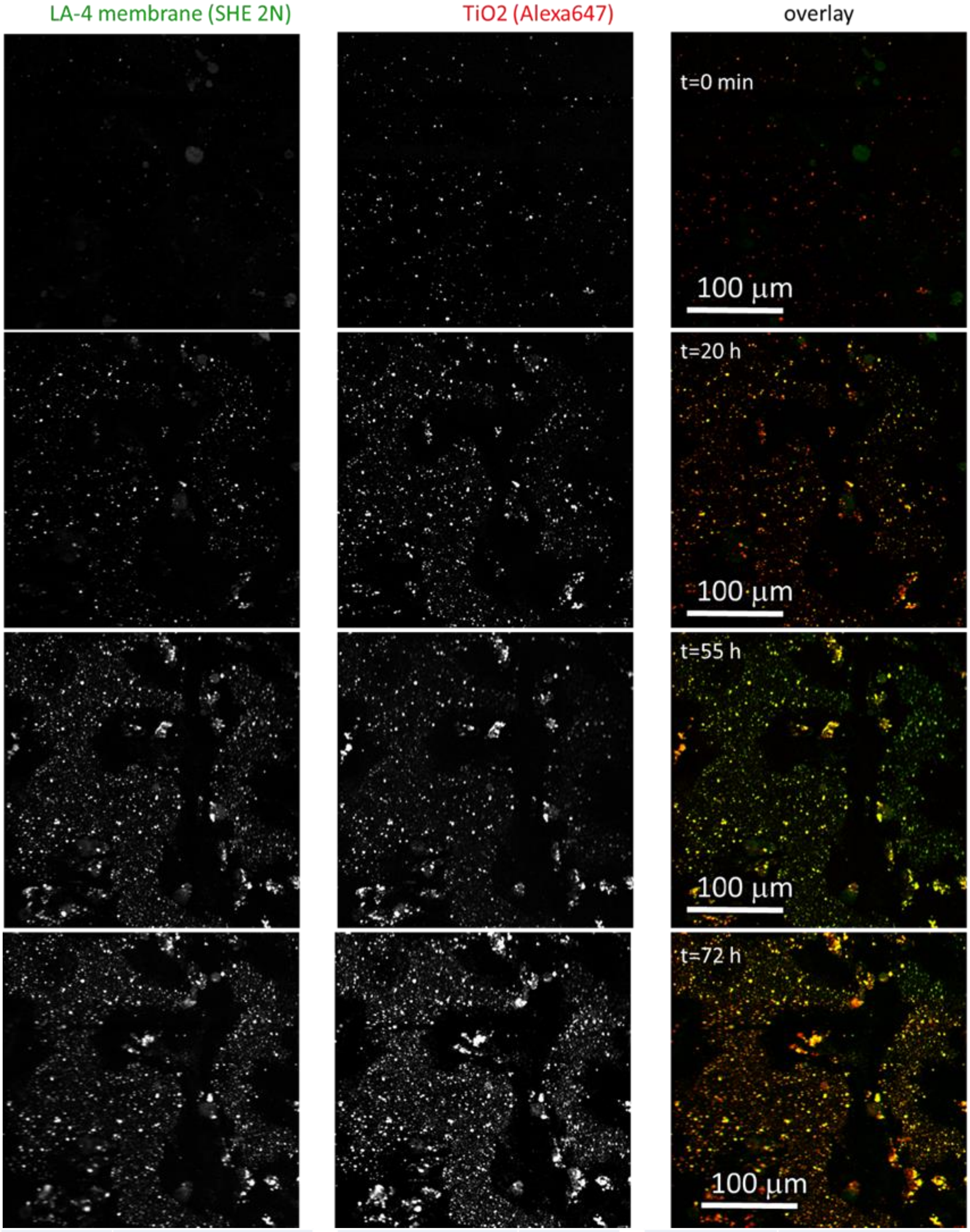

| | | | | | |
|---|---|---|---|---|---|
| Cell line | LA-4 & MHS (membrane, SHE 2N) | pixelsize (x,y) | 300 nm | 561nm | 15% |
| NPs | TiO2 (Alexa647) | FOV (x,y) | 400 µm | 640nm | 15% |
| exposure | 10:1, 0h-72h | pixelsize (z) | / nm | STED | % |
| imaging | xyz confocal, 72h | FOV (z) | / µm | filter sets | 605 nm – 625 nm, 650 nm – 720 nm |
| | | imaging time | 72 hours | dwell time | 10 µs |
| | | number of frames | 1015 | objective | 10x (NA0.3) |

*Figure S34: Co-culture of LA-4 and MH-S exposed to $TiO_2$ nanotubes for 72 h. MH-S are unable to phagocyte and remove all nanotubes from the epithelial layer after 72 h. See also Movie S1.*

#### Link to time-lapse

- Macrophage activity in a coculture with epithelial cells over the course of 72h:
    - *http://lbfnanobiodatabase.ijs.si/file/data/cauliflowerpaper/20190517__e01_m01_s01_LA-4_&_MH-S_SHE_2N_exposed_to_TiO2_Alexa_647_72h_MH-S_tracking.gif*

*Movie S1: Co-culture of LA-4 and MH-S exposed to $TiO_2$ nanotubes for 72 h. MH-S are unable to phagocyte and remove all nanotubes from the epithelial layer after 72 h. See also Figure S34.*

#### --- Supplementary information for experiments in Figures 1 – 5 in the main text ---

From here on, detailed supplementary material for each image from the main text is shown. It contains details on experimental design, controls and repetitions (fluorescence images shown both in separate channels and overlayed), and names (cyphers) of experiments to ease locating them in the depository/database.

We first supply the main text image duplicates for easier orientation. The supplement sections correspond to the panels in the main text. For example, section S2c in the supplement corresponds to the experiment depicted in Figure 2c in the main text.

### S1 – Quarantining of nanomaterials

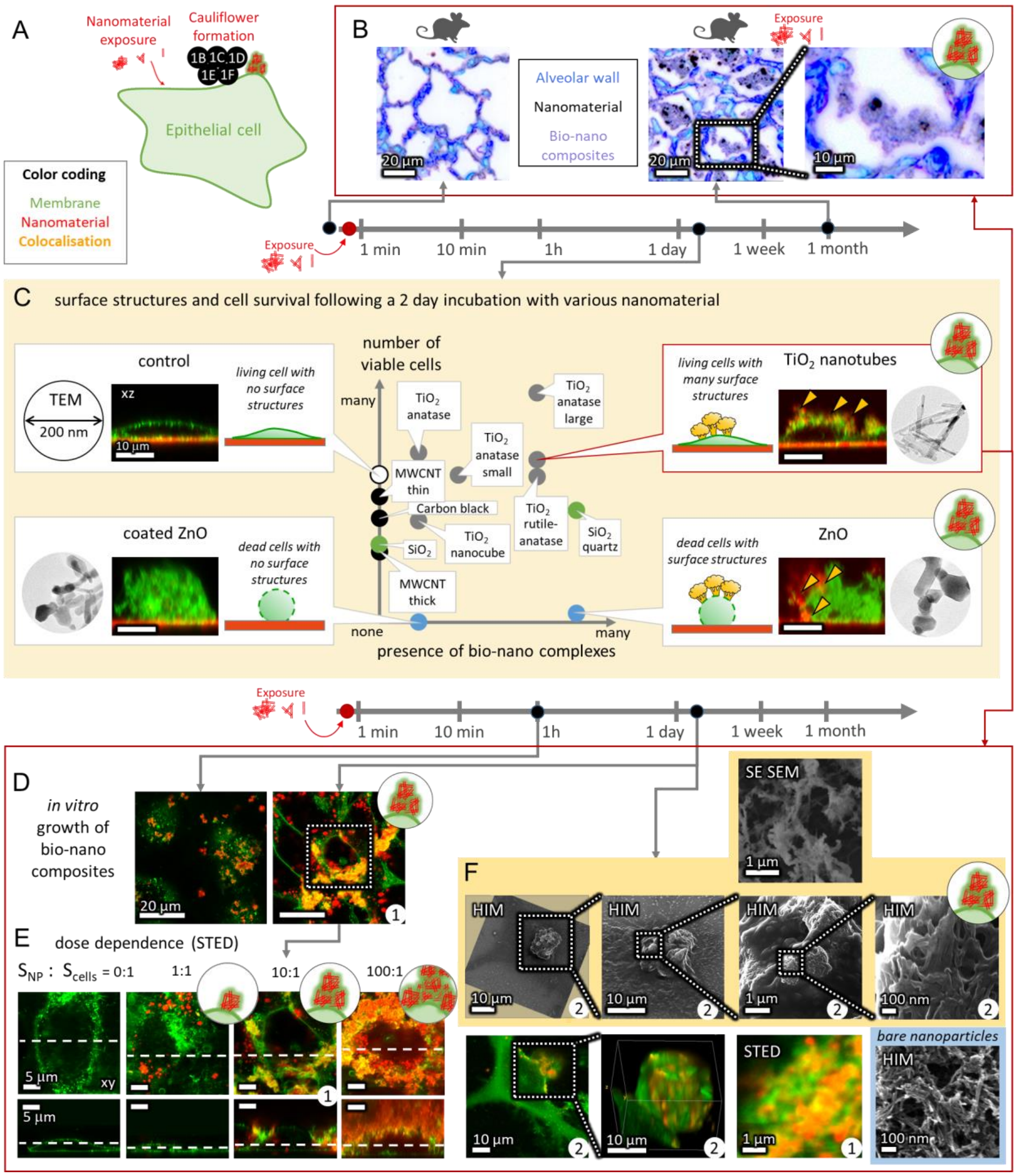

#### S1b – *In vivo* data

##### Main message

One month after intratracheal instillation in mice of $TiO_2$ tube, nanomaterial was mainly seen in bio-nano composites in alveolar spaces and in macrophages.

Supporting raw and analysed data:
For 28 days:

Figure S35

##### Materials and methods

The materials and methods used for intratracheal instillation of mice with $TiO_2$ tube are described in detail by Danielsen et. al[4] and included here in an abbreviated version.

###### *Preparation and characterization of $TiO_2$ tube suspensions*

For $TiO_2$ tube characterization see nanomaterial characterization section in S0b – Nanomaterial characterisation and [1].
The $TiO_2$ tube was suspended in nanopure water with 2 % v/v mouse serum (prepared in-house) to a final concentration of 3.24 mg/ml. The suspension was probe sonicated on ice for 16 min with 10 % amplitude. 3.24 mg/ml corresponds to a dose of 162 µg $TiO_2$ tube per 50 µl instillation volume per mice. The vehicle of nanopure water with 2 % v/v mouse serum was probe sonicated using the same protocol. The dose of 162 µg/mouse corresponds to an average surface dose of 3:1 $S_{nanomaterials}$:$S_{cells}$ and is equivalent to 15 working days at the 8-h time-weighted average occupational exposure limit for $TiO_2$ by Danish Regulations (6.0 $mg/m^3$ $TiO_2$).
The average hydrodynamic particle size of the $TiO_2$ tube in suspension (3.24 mg/ ml) was determined by Dynamic Light Scattering (DLS). The $TiO_2$ tube suspension had a bimodal size distribution with a major peak at 60 nm and a narrow peak at 21 nm[4]. The intensity-based z-average size was 168.7 nm and the polydispersity index (PI) was 0.586, indicating some polydispersity in the suspensions. Endotoxin levels were measured using the Limulus Amebocyte Lysate Assay. The level of endotoxins was low in $TiO_2$ tube suspensions (0.095 endotoxin units (EU)/mL), and in nanopure water with 2 % mouse serum (0.112 EU/ml).

###### *Animal handling and exposure*

Seven-week-old female C57BL/6jBomtac mice (Taconic, Ejby, Denmark) were randomized in groups for $TiO_2$ tube exposure (N=5 mice/group for histology) and vehicle controls (N = 2-4 mice/group). At 8 weeks of age the mice were anaesthetized and exposed to 0 µg or 162 µg $TiO_2$ tube in 50 µl vehicle by single intratracheal instillation. In brief, the mice were intubated in the trachea using a catheter. The 50 µl suspension was instilled followed by 200 µL air. The mouse was transferred to a vertical hanging position with the head up. This ensures that the administered material is maintained in the lung. Animal experiments were performed according to EC Directive 2010/63/UE in compliance with the handling guidelines established by the Danish government and permits from the Experimental Animal Inspectorate (no. 2015-15-0201-00465). Prior to the study, the experimental protocols were approved by the local Animal Ethics Council.
More details regarding the animal study can be found in Danielsen et al.[4].

##### *Histology and enhanced darkfield imaging*

At 28, 90 or 180 days post-exposure mice were weighed and anesthetized. Lungs were filled slowly with 4% formalin under 30 cm water column pressure. A knot was made on the trachea to secure formaldehyde in lungs to fixate tissue in “inflated state”. Lungs were then removed and placed in 4% neutral buffered formaldehyde for 24 hours. After fixation the samples were trimmed, dehydrated and embedded in paraffin. 3 µm thin sections were cut and stained with haematoxylin and eosin (H&E). Cytoviva enhanced darkfield hyperspectral system (Auburn, AL, USA) was used to image particles and organic debris in the histological sections of mouse lungs. Enhanced darkfield images were acquired at 100x on an Olympus BX 43 microscope with a Qimaging Retiga4000R camera.

##### Controls and statistics

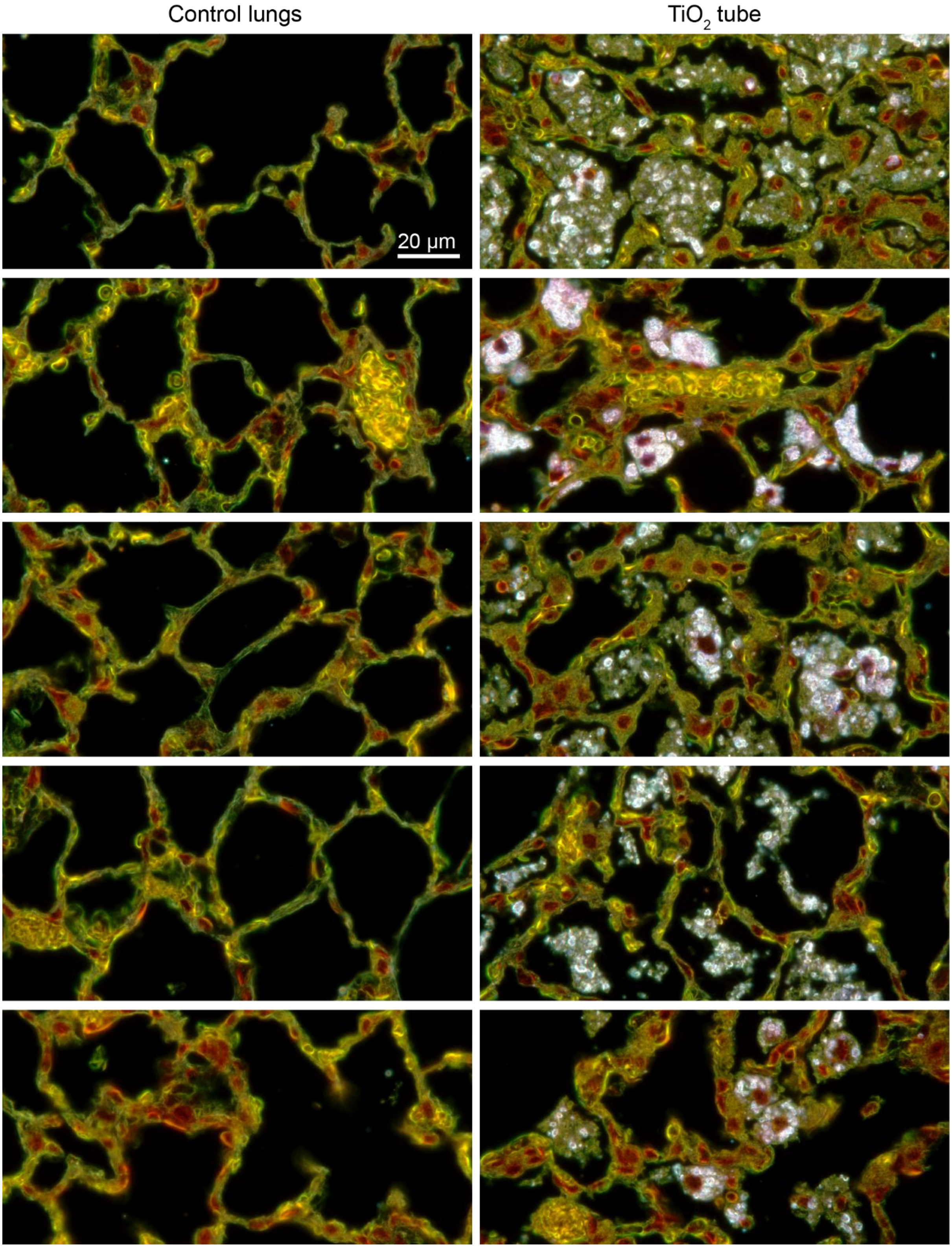

*Figure S35: Alveoli of murine lungs one month after intratracheal instillation of control vehicle or $TiO_2$ tube nanomaterial (white). $TiO_2$ tube nanomaterial was mainly seen in macrophages and in bio-nano composites in alveolar spaces and often close to epithelial cells of the alveolar wall. Enhanced darkfield microscopy of H&E stained histological tissue sections.*

#### S1c – Cauliflowers with various nanomaterial

##### Main message

Different nanomaterials exhibit different surface structures and viability of LA-4 cells – from large surface structures to none. Toxicity of the same nanomaterials is examined in "S0g – Cell viability LA-4 – various nanomaterials, same surface dose" and the TEM images of the nanomaterials are shown in section "S0b – Nanomaterial characterisation".

##### Materials and methods

- Experiment
    - Cells were grown up to 80% confluency in Ibidi #1.5H µ-Slide 8-well chambers for 24h which was followed by exposure to various nanomaterials at a 10 : 1 ($S_{nano}$:$S_{cells}$) surface dose
    - nanomaterials used in experiment are in the Table S2

*Table S2: List of nanomaterials exposed on LA-4 cells in this study with their specific β-BET surface [m2/g]*

| Name | official ID - name | Nanomaterial code | β-BET surface [m2/g] |
|---|---|---|---|
| **$TiO_2$ nanotubes** | $TiO_2$ nanotubes | PU-nTOX-01-03 | 152 |
| **$SiO_2$ DQ12** | Quartz DQ12 | / | 10,1 |
| **Printex 90** | Printex 90, carbon black | / | 310 |
| **$TiO_2$ nanocubes** | $TiO_2$ nanocubes | PU-nTOX-01-21 | 86 |
| **$TiO_2$ rut-an NM-105** | NM105 TiO2 rut-anat | TiO2-NM105-JRCNM01005a | 46 |
| **$TiO_2$ anat MKNA015** | MKNA015 | MKN-TiO2-A015 | 250 |
| **$TiO_2$ anat MKNA100** | MKNA100 | MKN-TiO2-A100 | 16,8 |
| **$TiO_2$ an NM-101** | NM101 TiO2 anatase | TiO2-NM101-JRCNM01001a | 230 |
| **$SiO_2$ NM-200** | NM200 Silica | SiO2-NM200-JRCNM02000a | 189 |
| **MWCNT 402** | NM402 MWCNT | MWCNTs-NM402-JRCNM04002a | 226 |
| **MWCNT 401** | NM401 SWCNT | MWCNTs-NM401-JRCNM04001a | 18 |
| **ZnO NM-111** | NM111 ZnO | ZnO-NM111-JRCNM01101a | 12 |
| **ZnO raw NM-110** | noncoated ZnO | ZnO-NM110-JRCNM62101a | 15 |

    - cells were incubated with nanomaterials for additional 48h
- preparation for imaging:
    - after incubation cell media was exchanged for 150 µL of fresh medium with 5 µg $ml^{-1}$ of Cell Mask Orange and incubated for additional 10 min
    - cells were then washed 1x200 µL LCIS and mixture of 50%LCIS and 50%F12-K medium was added for imaging
    - samples were held on the microscope stage heated at 37°C for 2h

- Imaging:
    - To get an unbiased overview of the sample we used a home-made script which semi-randomly selected all the imaging places. On each well, we'd select three regions of interest. In these regions, the first image was of our choosing remaining five images were randomly selected by our script. Each well is thus, represented with 18 images.
    - The extent of the surface structures was determined according to the cauliflower forming potential scale (CFP) devised in our lab (Table S3). Samples were evaluated during the imaging.

*Table S3: Cauliflower forming potential scale used for sample description*

| CFP | Description |
|---|---|
| 0 | no cauliflowers in 1 cm^2 |
| 1 | few cauliflowers per 1 cm^2 |
| 2 | up to 1 % cells have small cauliflowers |
| 3 | up to 10% cells have small cauliflowers |
| 4 | most cells have small cauliflowers OR up to 10% cells have large cauliflowers |
| 5 | most cells have large cauliflowers |

According to the CFP scale all evaluated nanomaterial have been given a CFP score presented in the Table S4

*Table S4: CFP score of exposed LA-4 cells for all the nanomaterials used in this study*

| Name | sample name | β-BET surface [m2/g] | min CFP [ /5] | Max CFP [ /5] | total CFP |
|---|---|---|---|---|---|
| **SiO2 DQ12** | PC-nTOX-02-39 | 10,1 | 2 | 5 | 5 |
| **ZnO raw NM-110** | PC-nTOX-02-50 | 12 | x | x | x |
| **ZnO NM-111** | PC-nTOX-02-44 | 15 | x | x | x |
| **TiO2 anatase MKNA100** | PC-nTOX-02-46 | 16,8 | 4 | 4 | 4 |
| **MWCNT 401** | PC-nTOX-02-43 | 18 | 0 | 0 | 0 |
| **TiO2 rut-anat NM-105** | PC-nTOX-02-47 | 46 | 4 | 4 | 4 |
| **TiO2 NQs** | PC-nTOX-02-41 | 86 | 1 | 1 | 1 |
| **TiO2 NTs** | PC-nTOX-02-38 | 152 | 2 | 4 | 4 |
| **SiO2 NM-200** | PC-nTOX-02-48 | 189 | 0 | 0 | 0 |
| **MWCNT 402** | PC-nTOX-02-42 | 226 | 0 | 0 | 0 |
| **TiO2 anatase NM-101** | PC-nTOX-02-49 | 230 | 1 | 1 | 1 |
| **TiO2 anatase MKNA015** | PC-nTOX-02-45 | 250 | 2 | 2 | 2 |
| **Printex 90** | PC-nTOX-02-40 | 300 | 0 | 0 | 0 |
| **Sham Control** | PC-nTOX-02-51 | - | - | - | - |
| **Control** | - | - | - | - | - |

##### Experiment names

CTRL >

20191129_exp08_LA-4-CMO_ctrl-48h-3-pos_0_xy_MC-PMT_AC_8bit_ovrly1
20191129_exp08_LA-4-CMO_ctrl-48h-3-pos_0_xzy_MC-PMT_AC_8bit_ovrly1
20191129_exp08_LA-4-CMO_ctrl-48h-2-pos_0_xy_MC-PMT_AC_8bit_ovrly1
20191129_exp08_LA-4-CMO_ctrl-48h-2-pos_0_xzy_MC-PMT_AC_8bit_ovrly1
20191129_exp08_LA-4-CMO_ctrl-48h-2-pos_5_xy_MC-PMT_AC_8bit_ovrly1
20191129_exp08_LA-4-CMO_ctrl-48h-2-pos_5_xzy_MC-PMT_AC_8bit_ovrly1

Sham >

20191129_exp15_LA-4-CMO_sham-48h-1-pos_4_xy_MC-PMT_AC_8bit_ovrly1
20191129_exp15_LA-4-CMO_sham-48h-1-pos_4_xzy_MC-PMT_AC_8bit_ovrly1

20191129_exp15_LA-4-CMO_sham-48h-1-pos_5_xzy_MC-PMT_AC_8bit_ovrly1
20191129_exp15_LA-4-CMO_sham-48h-3-pos_0_xy_MC-PMT_AC_8bit_ovrly1
20191129_exp15_LA-4-CMO_sham-48h-3-pos_0_xzy_MC-PMT_AC_8bit_ovrly1

TiO2 NTs >
20191129_exp01_LA-4-CMO_TiO2-NT-N-48h-4-pos_4_xy_MC-PMT_AC_8bit_ovrly1
20191129_exp01_LA-4-CMO_TiO2-NT-N-48h-4-pos_4_yzx_MC-PMT_AC_8bit_ovrly1
20191129_exp01_LA-4-CMO_TiO2-NT-N-48h-4-pos_3_xy_MC-PMT_AC_8bit_ovrly1
20191129_exp01_LA-4-CMO_TiO2-NT-N-48h-4-pos_3_xzy_MC-PMT_AC_8bit_ovrly1
20191129_exp01_LA-4-CMO_TiO2-NT-N-48h-6-pos_0_xy_MC-PMT_AC_8bit_ovrly1
20191129_exp01_LA-4-CMO_TiO2-NT-N-48h-6-pos_0_xzy_MC-PMT_AC_8bit_ovrly1

DQ12 >
20191129_exp02_LA-4-CMO_DQ12-N-48h-1-pos_0_xy_MC-PMT_AC_8bit_ovrly1
20191129_exp02_LA-4-CMO_DQ12-N-48h-1-pos_0_xzy_MC-PMT_AC_8bit_ovrly1
20191129_exp02_LA-4-CMO_DQ12-N-48h-3-pos_5_xy_MC-PMT_AC_8bit_ovrly1
20191129_exp02_LA-4-CMO_DQ12-N-48h-3-pos_5_xzy_MC-PMT_AC_8bit_ovrly1
20191129_exp02_LA-4-CMO_DQ12-N-48h-1-pos_1_xy_MC-PMT_AC_8bit_ovrly1

20191129_exp02_LA-4-CMO_DQ12-N-48h-3-pos_5_yzx_MC-PMT_AC_8bit_ovrly1

Printex 90 >
20191129_exp03_LA-4-CMO_Printex90-N-48h-3-pos_0_xy_MC-PMT_AC_8bit_ovrly1
20191129_exp03_LA-4-CMO_Printex90-N-48h-3-pos_0_xzy_MC-PMT_AC_8bit_ovrly1
20191129_exp03_LA-4-CMO_Printex90-N-48h-1-pos_4_xy_MC-PMT_AC_8bit_ovrly1
20191129_exp03_LA-4-CMO_Printex90-N-48h-1-pos_4_xzy_MC-PMT_AC_8bit_ovrly1
20191129_exp03_LA-4-CMO_Printex90-N-48h-2-pos_1_xy_MC-PMT_AC_8bit_ovrly1
20191129_exp03_LA-4-CMO_Printex90-N-48h-2-pos_1_xzy_MC-PMT_AC_8bit_ovrly1

TiO2 NQs >
20191129_exp04_LA-4-CMO_TiO2-NQs-N48h-3-pos_3_xy_MC-PMT_AC_8bit_ovrly1
20191129_exp04_LA-4-CMO_TiO2-NQs-N48h-3-pos_3_xzy_MC-PMT_AC_8bit_ovrly1

20191129_exp04_LA-4-CMO_TiO2-NQs-N48h-2-pos_3_xy_MC-PMT_AC_8bit_ovrly1
20191129_exp04_LA-4-CMO_TiO2-NQs-N48h-2-pos_3_yzx_MC-PMT_AC_8bit_ovrly1
20191129_exp04_LA-4-CMO_TiO2-NQs-N48h-4-pos_3_xy_MC-PMT_AC_8bit_ovrly1
20191129_exp04_LA-4-CMO_TiO2-NQs-N48h-4-pos_3_xzy_MC-PMT_AC_8bit_ovrly1

MWCNT-401 >
20191129_exp06_LA-4-CMO_MWCNT-NM401-N-48h-1-pos_0_xy_MC-PMT_AC_8bit_ovrly1
20191129_exp06_LA-4-CMO_MWCNT-NM401-N-48h-1-pos_0_xzy_MC-PMT_AC_8bit_ovrly1
20191129_exp06_LA-4-CMO_MWCNT-NM401-N-48h-2-pos_5_xy_MC-PMT_AC_8bit_ovrly1
20191129_exp06_LA-4-CMO_MWCNT-NM401-N-48h-2-pos_5_yzx_MC-PMT_AC_8bit_ovrly1
20191129_exp06_LA-4-CMO_MWCNT-NM401-N-48h-3-pos_4_xy_MC-PMT_AC_8bit_ovrly1
20191129_exp06_LA-4-CMO_MWCNT-NM401-N-48h-3-pos_4_yzx_MC-PMT_AC_8bit_ovrly1

MWCNT-402 >
20191129_exp05_LA-4-CMO_MWCNT-NM402-N-48h-3-pos_1_xy_MC-PMT_AC_8bit_ovrly1
20191129_exp05_LA-4-CMO_MWCNT-NM402-N-48h-3-pos_1_xzy_MC-PMT_AC_8bit_ovrly1
20191129_exp05_LA-4-CMO_MWCNT-NM402-N-48h-2-pos_0_xy_MC-PMT_AC_8bit_ovrly1
20191129_exp05_LA-4-CMO_MWCNT-NM402-N-48h-2-pos_0_xzy_MC-PMT_AC_8bit_ovrly1
20191129_exp05_LA-4-CMO_MWCNT-NM402-N-48h-1-pos_1_xy_MC-PMT_AC_8bit_ovrly1
20191129_exp05_LA-4-CMO_MWCNT-NM402-N-48h-1-pos_1_yzx_MC-PMT_AC_8bit_ovrly1

ZnO-NM111 >
20191129_exp07_LA-4-CMO_ZnO-coat-NM111-N-48h-1-pos_0_xy_MC-PMT_AC_8bit_ovrly1
20191129_exp07_LA-4-CMO_ZnO-coat-NM111-N-48h-1-pos_0_xzy_MC-PMT_AC_8bit_ovrly1
20191129_exp07_LA-4-CMO_ZnO-coat-NM111-N-48h-2-pos_0_xy_MC-PMT_AC_8bit_ovrly1
20191129_exp07_LA-4-CMO_ZnO-coat-NM111-N-48h-2-pos_0_xzy_MC-PMT_AC_8bit_ovrly1
20191129_exp07_LA-4-CMO_ZnO-coat-NM111-N-48h-2-pos_3_xy_MC-PMT_AC_8bit_ovrly1
20191129_exp07_LA-4-CMO_ZnO-coat-NM111-N-48h-2-pos_3_xzy_MC-PMT_AC_8bit_ovrly1

ZnO-Raw >
20191129_exp14_LA-4-CMO_ZnOraw-NM110-N-48h-3-pos_2_xy_MC-PMT_AC_8bit_ovrly1
20191129_exp14_LA-4-CMO_ZnOraw-NM110-N-48h-3-pos_2_xzy_MC-PMT_AC_8bit_ovrly1
20191129_exp14_LA-4-CMO_ZnOraw-NM110-N-48h-3-pos_0_xy_MC-PMT_AC_8bit_ovrly1
20191129_exp14_LA-4-CMO_ZnOraw-NM110-N-48h-3-pos_0_xzy_MC-PMT_AC_8bit_ovrly1
20191129_exp14_LA-4-CMO_ZnOraw-NM110-N-48h-1-pos_2_xy_MC-PMT_AC_8bit_ovrly1
20191129_exp14_LA-4-CMO_ZnOraw-NM110-N-48h-1-pos_2_yzx_MC-PMT_AC_8bit_ovrly1

TiO2-anatase L MKNA 100 >
20191129_exp10_LA-4-CMO_TiO2anatL-MKNA100-N-48h-3-pos_2_xy_MC-PMT_AC_8bit_ovrly1
20191129_exp10_LA-4-CMO_TiO2anatL-MKNA100-N-48h-3-pos_2_xzy_MC-PMT_AC_8bit_ovrly1
20191129_exp10_LA-4-CMO_TiO2anatL-MKNA100-N-48h-3-pos_1_xy_MC-PMT_AC_8bit_ovrly1
20191129_exp10_LA-4-CMO_TiO2anatL-MKNA100-N-48h-3-pos_1_xzy_MC-PMT_AC_8bit_ovrly1
20191129_exp10_LA-4-CMO_TiO2anatL-MKNA100-N-48h-3-pos_4_xy_MC-PMT_AC_8bit_ovrly1
20191129_exp10_LA-4-CMO_TiO2anatL-MKNA100-N-48h-3-pos_4_yzx_MC-PMT_AC_8bit_ovrly1

TiO2-tutile-anatase-MKNA 105 >
20191129_exp11_LA-4-CMO_TiO2rut-anat-NM105-N-48h-2-pos_3_xy_MC-PMT_AC_8bit_ovrly1
20191129_exp11_LA-4-CMO_TiO2rut-anat-NM105-N-48h-2-pos_3_xzy_MC-PMT_AC_8bit_ovrly1
20191129_exp11_LA-4-CMO_TiO2rut-anat-NM105-N-48h-1-pos_2_xy_MC-PMT_AC_8bit_ovrly1
20191129_exp11_LA-4-CMO_TiO2rut-anat-NM105-N-48h-1-pos_2_xzy_MC-PMT_AC_8bit_ovrly1
20191129_exp11_LA-4-CMO_TiO2rut-anat-NM105-N-48h-1-pos_0_xy_MC-PMT_AC_8bit_ovrly1
20191129_exp11_LA-4-CMO_TiO2rut-anat-NM105-N-48h-1-pos_0_xzy_MC-PMT_AC_8bit_ovrly1

TiO2-anatase-NM101 >
20191129_exp13_LA-4-CMO_TiO2-anat-NM101-N-48h-3-pos_4_xy_MC-PMT_AC_8bit_ovrly1
20191129_exp13_LA-4-CMO_TiO2-anat-NM101-N-48h-3-pos_4_xzy_MC-PMT_AC_8bit_ovrly1
20191129_exp13_LA-4-CMO_TiO2-anat-NM101-N-48h-1-pos_4_xy_MC-PMT_AC_8bit_ovrly1

20191129_exp13_LA-4-CMO_TiO2-anat-NM101-N-48h-1-pos_3_yzx_MC-PMT_AC_8bit_ovrly1

20191129_exp13_LA-4-CMO_TiO2-anat-NM101-N-48h-1-pos_0_xy_MC-PMT_AC_8bit_ovrly1

20191129_exp13_LA-4-CMO_TiO2-anat-NM101-N-48h-1-pos_0_yzx_MC-PMT_AC_8bit_ovrly1

SiO2 NM200 >

20191129_exp12_LA-4-CMO_SiO2-NM200-N-48h-3-pos_0_xy_MC-PMT_AC_8bit_ovrly1

20191129_exp12_LA-4-CMO_SiO2-NM200-N-48h-3-pos_0_yzx_MC-PMT_AC_8bit_ovrly1

20191129_exp12_LA-4-CMO_SiO2-NM200-N-48h-2-pos_0_xy_MC-PMT_AC_8bit_ovrly1

20191129_exp12_LA-4-CMO_SiO2-NM200-N-48h-2-pos_0_xzy_MC-PMT_AC_8bit_ovrly1

20191129_exp12_LA-4-CMO_SiO2-NM200-N-48h-1-pos_4_xy_MC-PMT_AC_8bit_ovrly1

20191129_exp12_LA-4-CMO_SiO2-NM200-N-48h-1-pos_4_xy_MC-PMT_AC_8bit_ovrly1

#### Control and statistics

LA-4 cells exposed to 10 to 1 surface dose of various nanomaterials.

Figures S 36-49

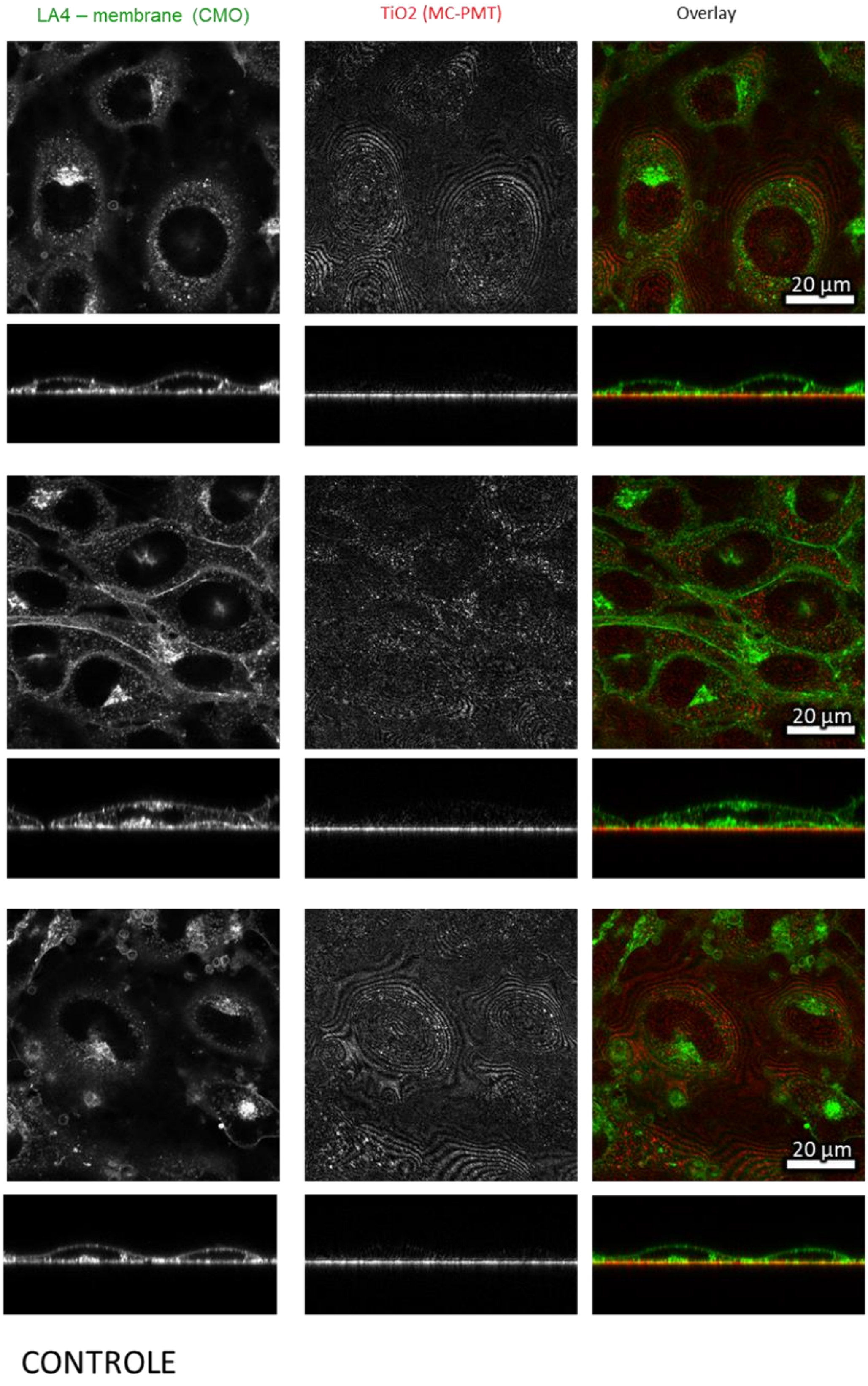

*Figure S36: Confocal images of unexposed LA-4 cells after 48 h of incubation in xy plane (upper windows) and xz plane (lower windows)*

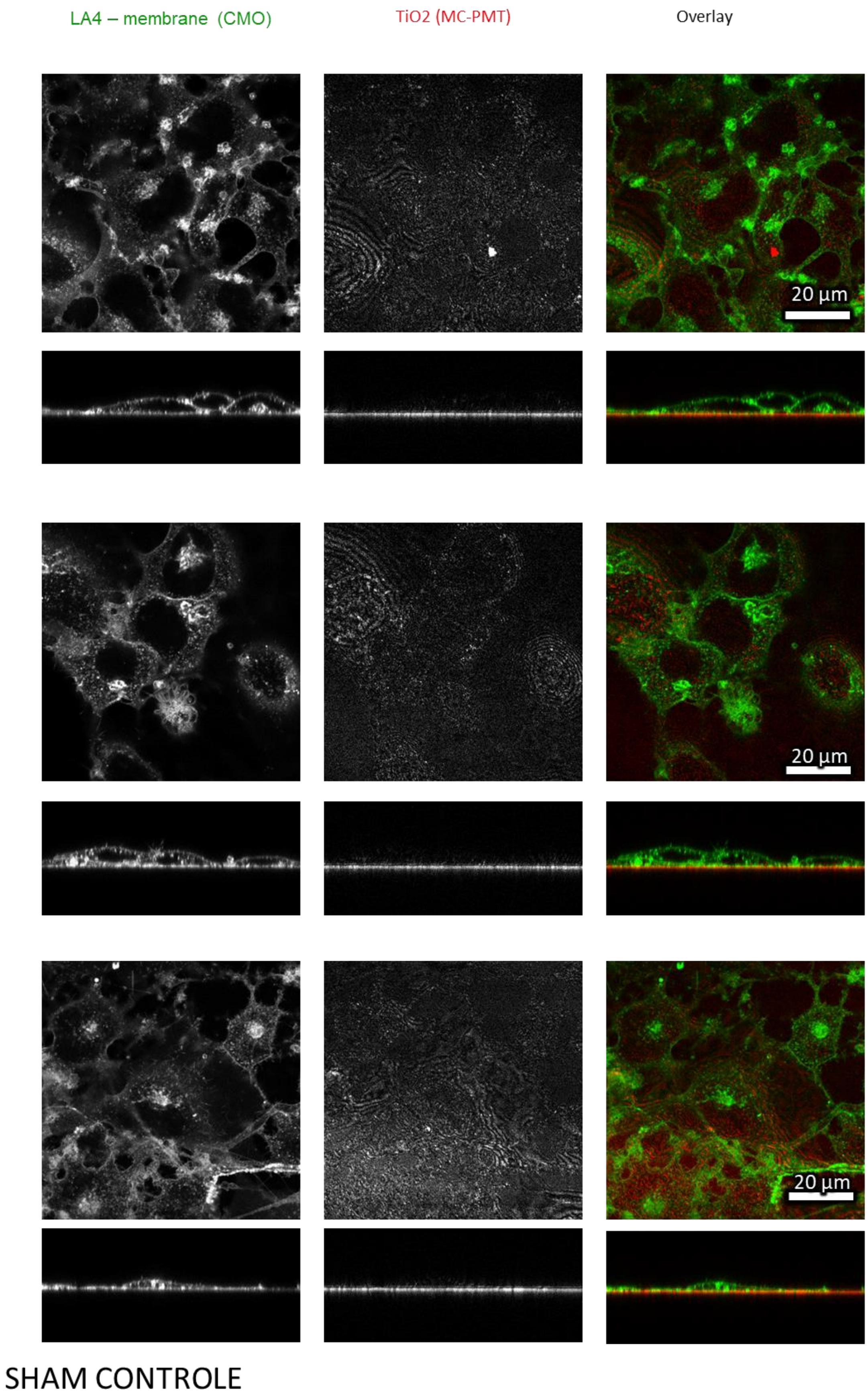

*Figure S37: Confocal images of unexposed LA-4 cells after 48 h of incubation in tip sonicated medium in xy plane (upper windows) and xz plane (lower windows)*

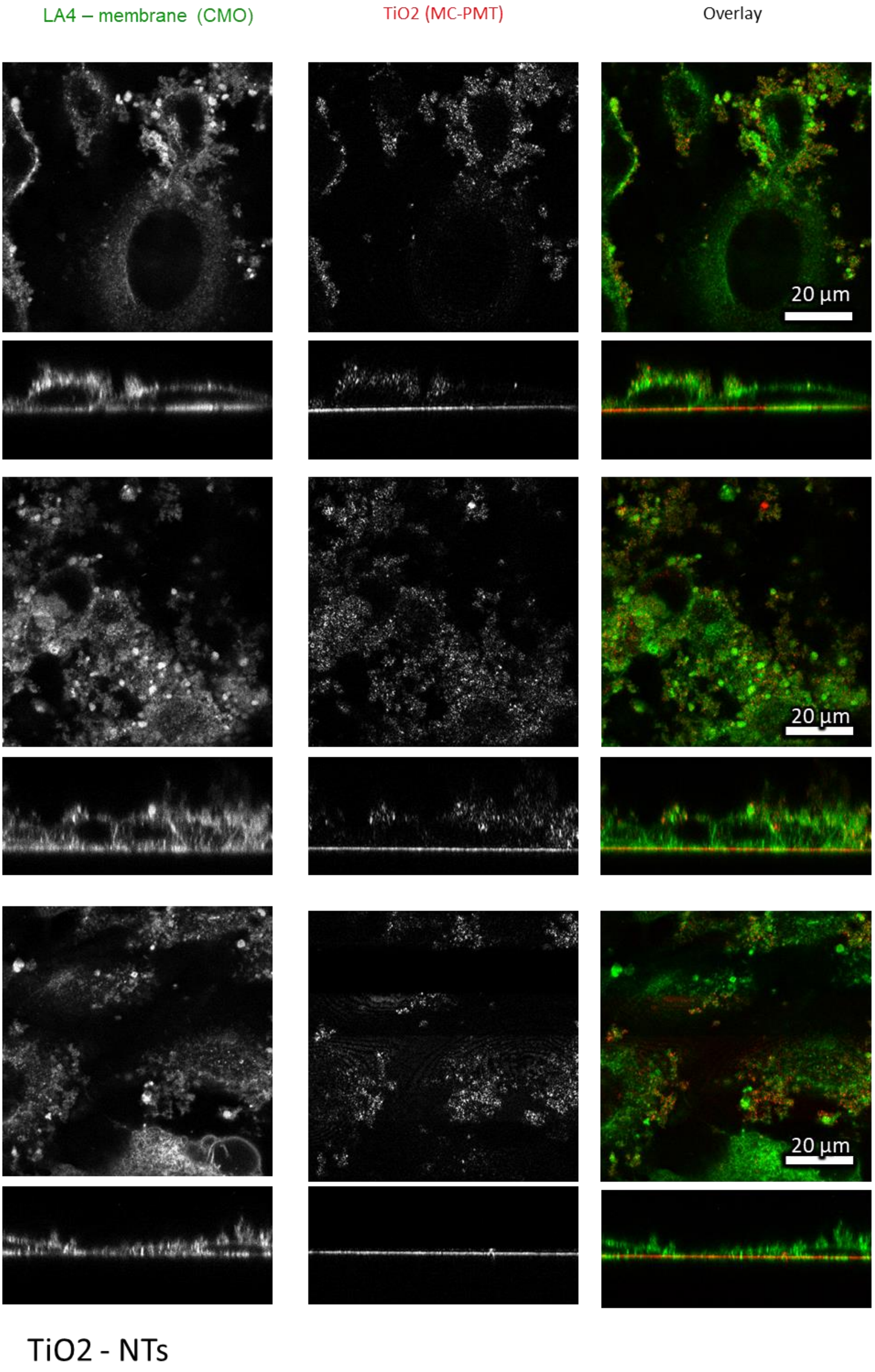

*Figure S38: Confocal images of LA-4 cells exposed to 10 to 1 surface dose of $TiO_2$ nanotubes for 48 h in xy plane (upper windows) and xz plane (lower windows)*

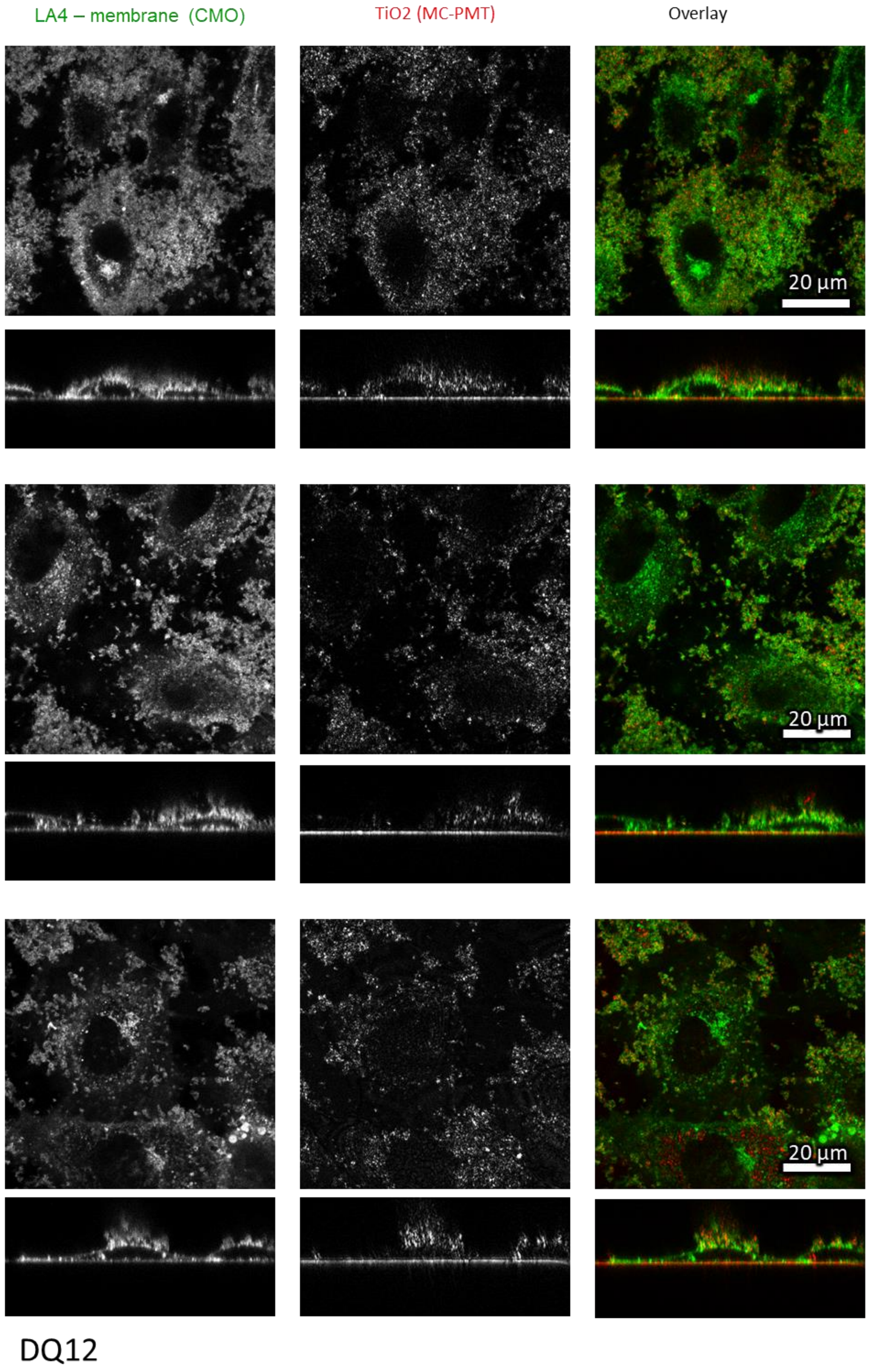

*Figure S39: Confocal images of LA-4 cells exposed to 10 to 1 surface dose of DQ12 for 48 h in xy plane (upper windows) and xz plane (lower windows)*

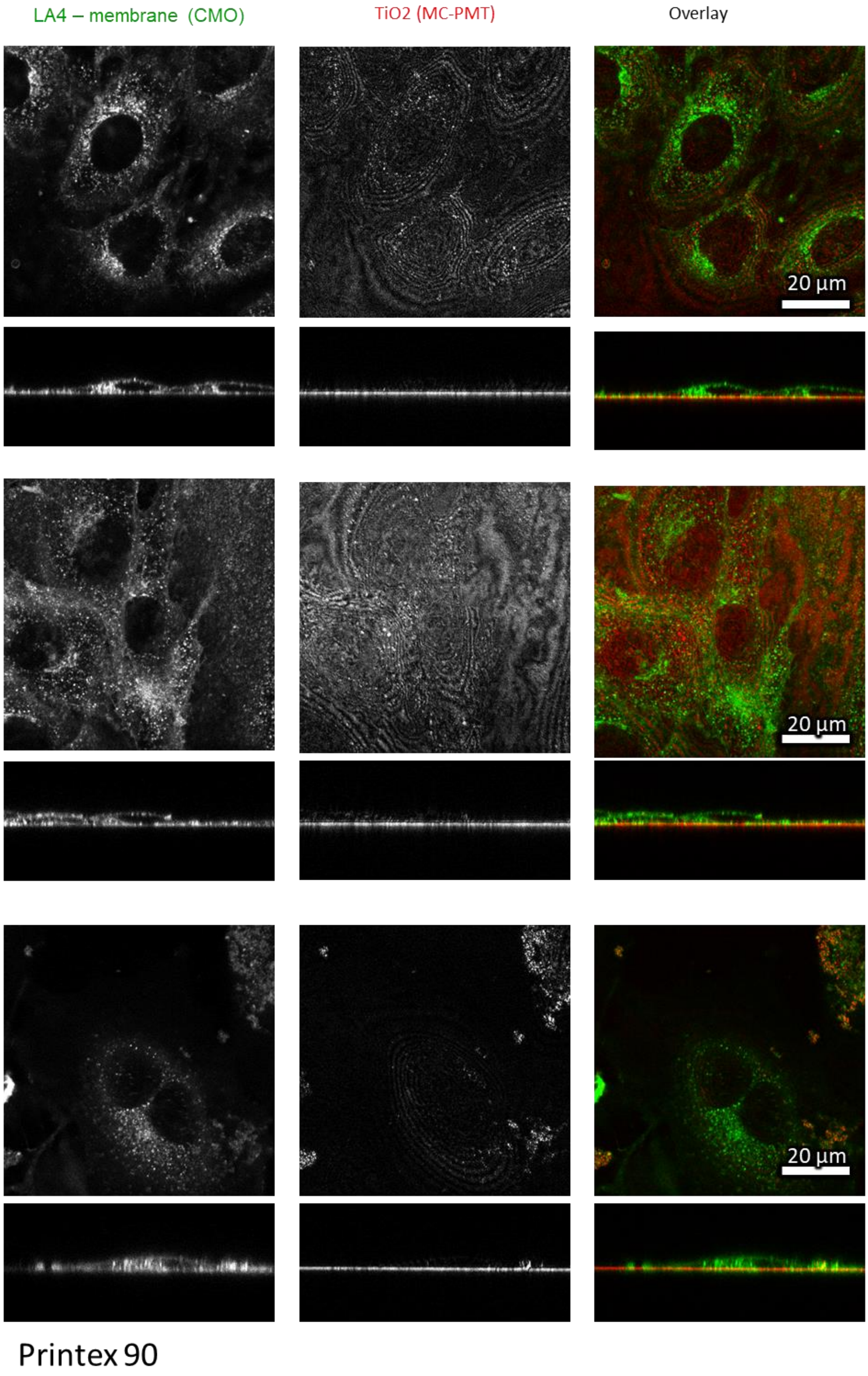

*Figure S40: Confocal images of LA-4 cells exposed to 10 to 1 surface dose of Printex 90 for 48 h in xy plane (upper windows) and xz plane (lower windows)*

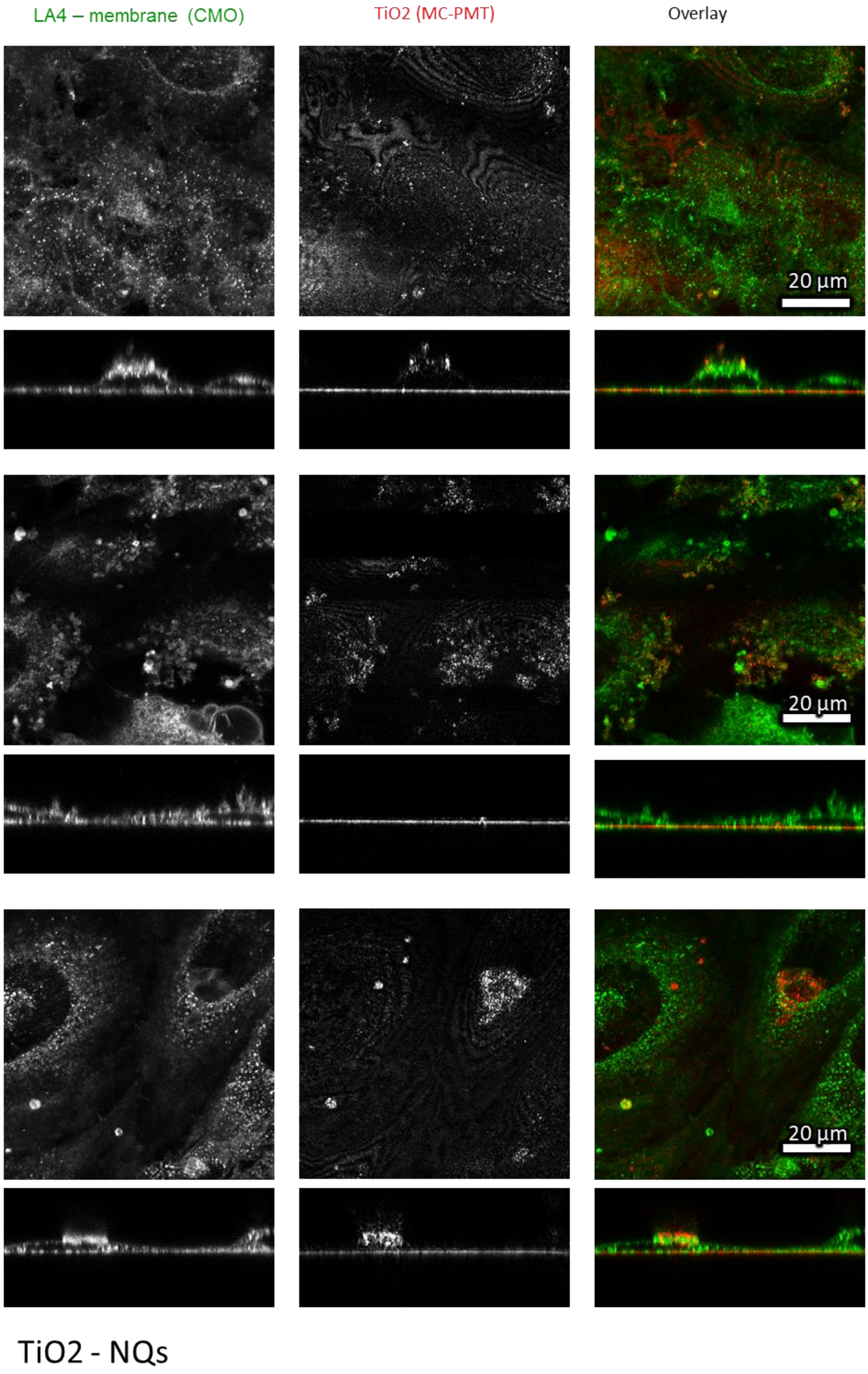

*Figure S41: Confocal images of LA-4 cells exposed to 10 to 1 surface dose of $TiO_2$ nanoqubes for 48 h in xy plane (upper windows) and xz plane (lower windows)*

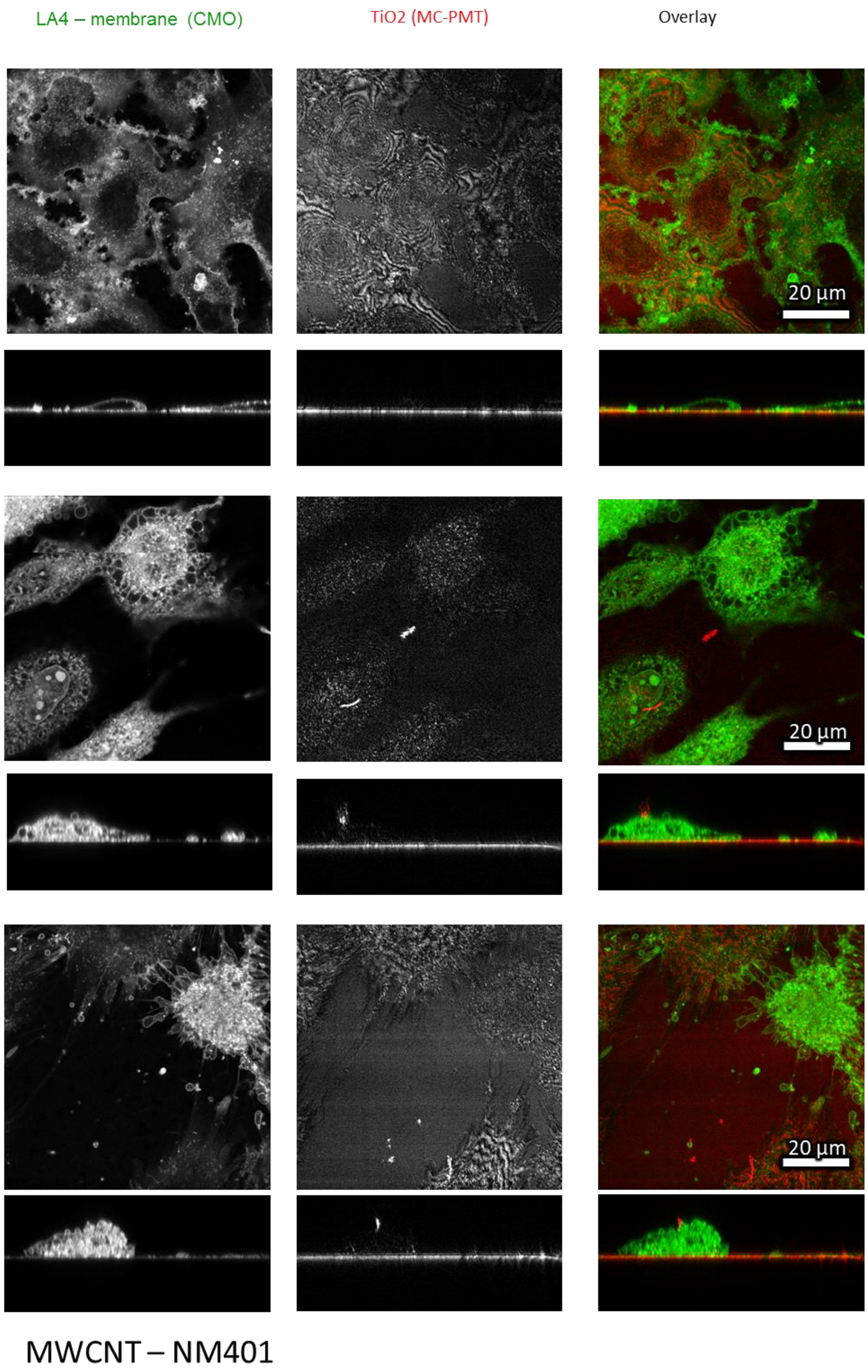

*Figure S42: Confocal images of LA-4 cells exposed to 10 to 1 surface dose of MWCNT-NM401 for 48 h in xy plane (upper windows) and xz plane (lower windows)*

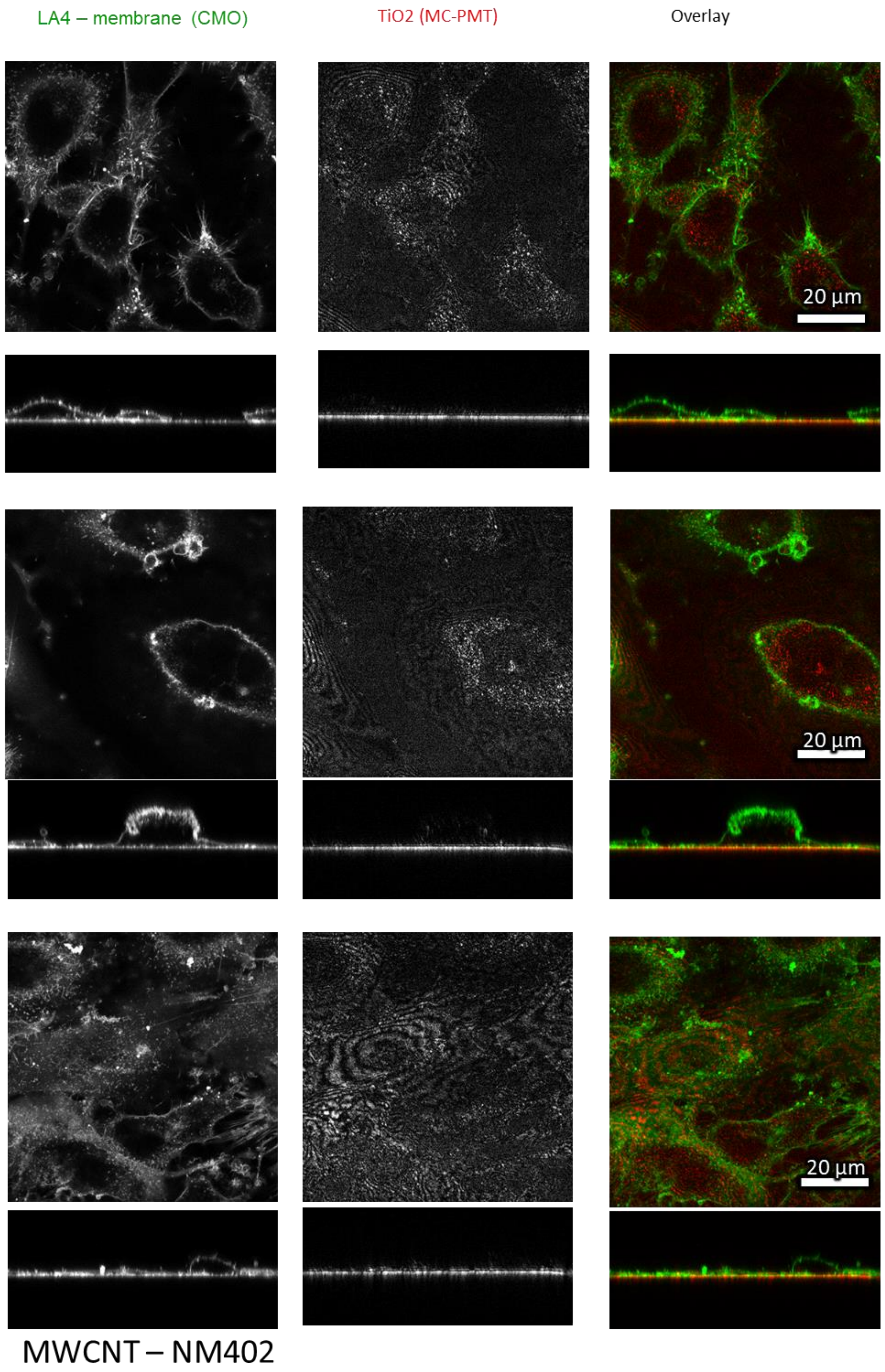

*Figure S43: Confocal images of LA-4 cells exposed to 10 to 1 surface dose of MWCNT-NM402 for 48 h in xy plane (upper windows) and xz plane (lower windows)*

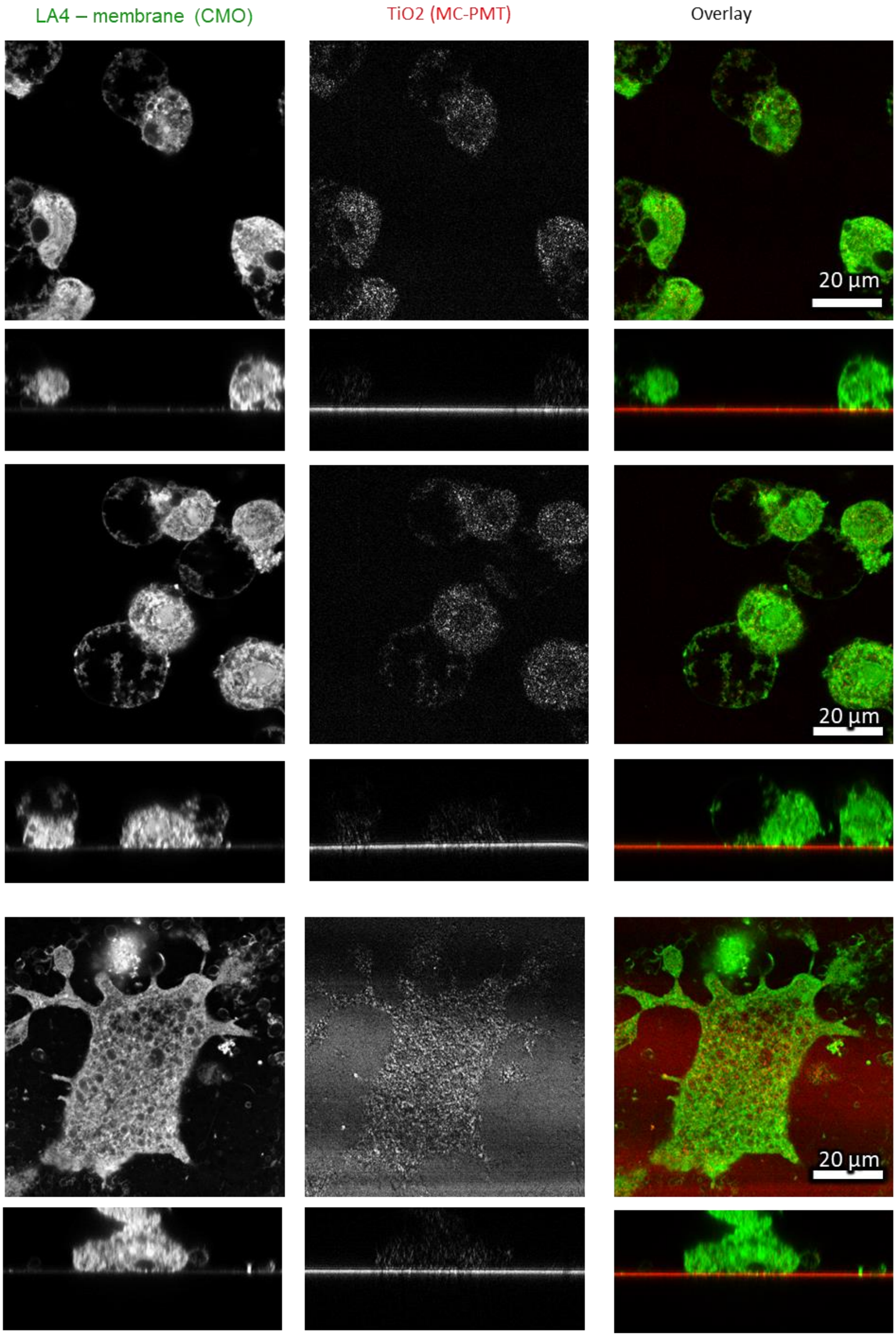

#### ZnO – NM111

*Figure S44: Confocal images of LA-4 cells exposed to 10 to 1 surface dose of ZnO-NM111 for 48 h in xy plane (upper windows) and xz plane (lower windows)*

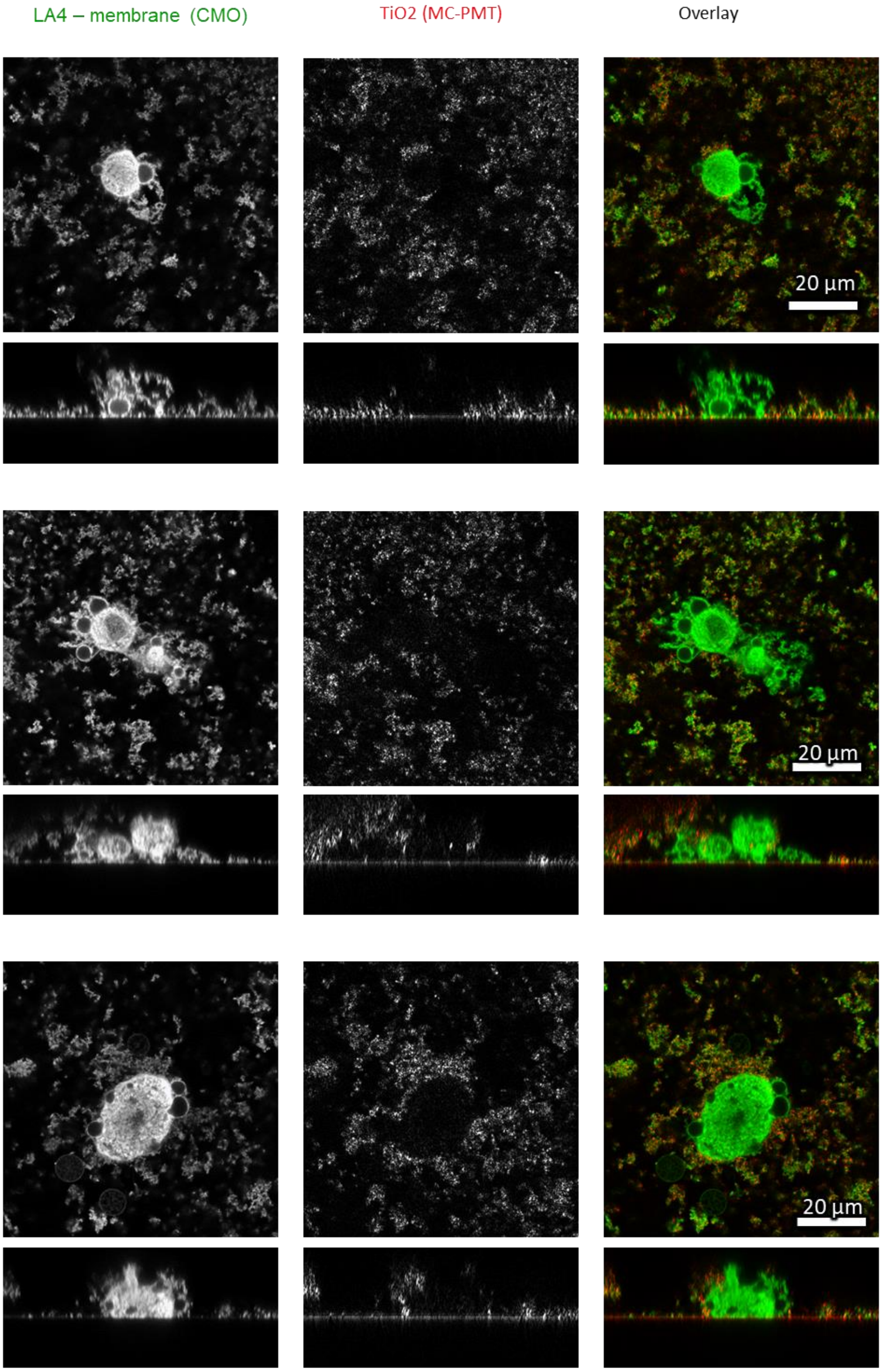

#### ZnO - Raw

*Figure S45: Confocal images of LA-4 cells exposed to 10 to 1 surface dose of ZnO-Raw for 48 h in xy plane (upper windows) and xz plane (lower windows)*

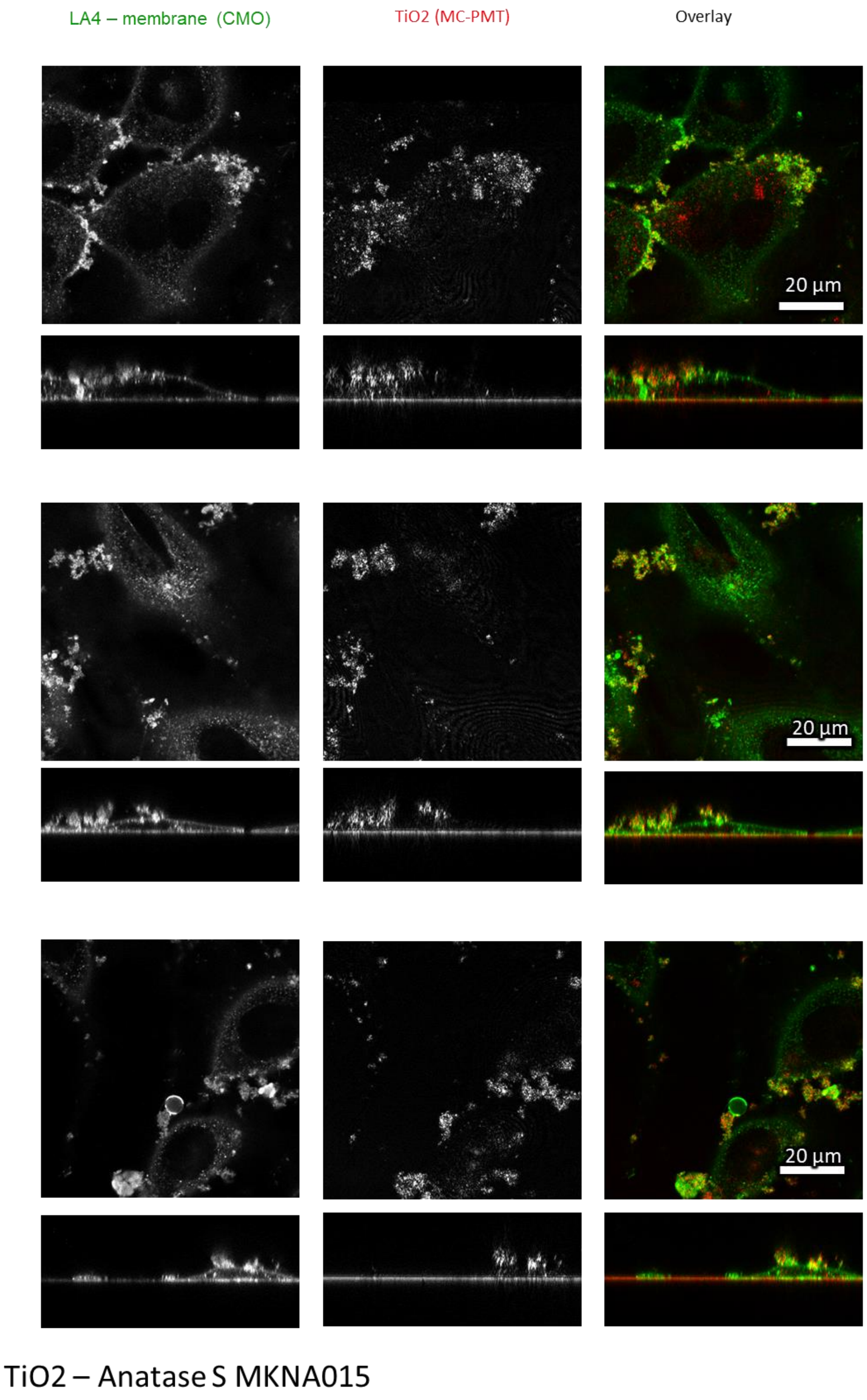

*Figure S46: Confocal images of LA-4 cells exposed to 10 to 1 surface dose of $TiO_2$ anatase MKNA015 for 48 h in xy plane (upper windows) and xz plane (lower windows)*

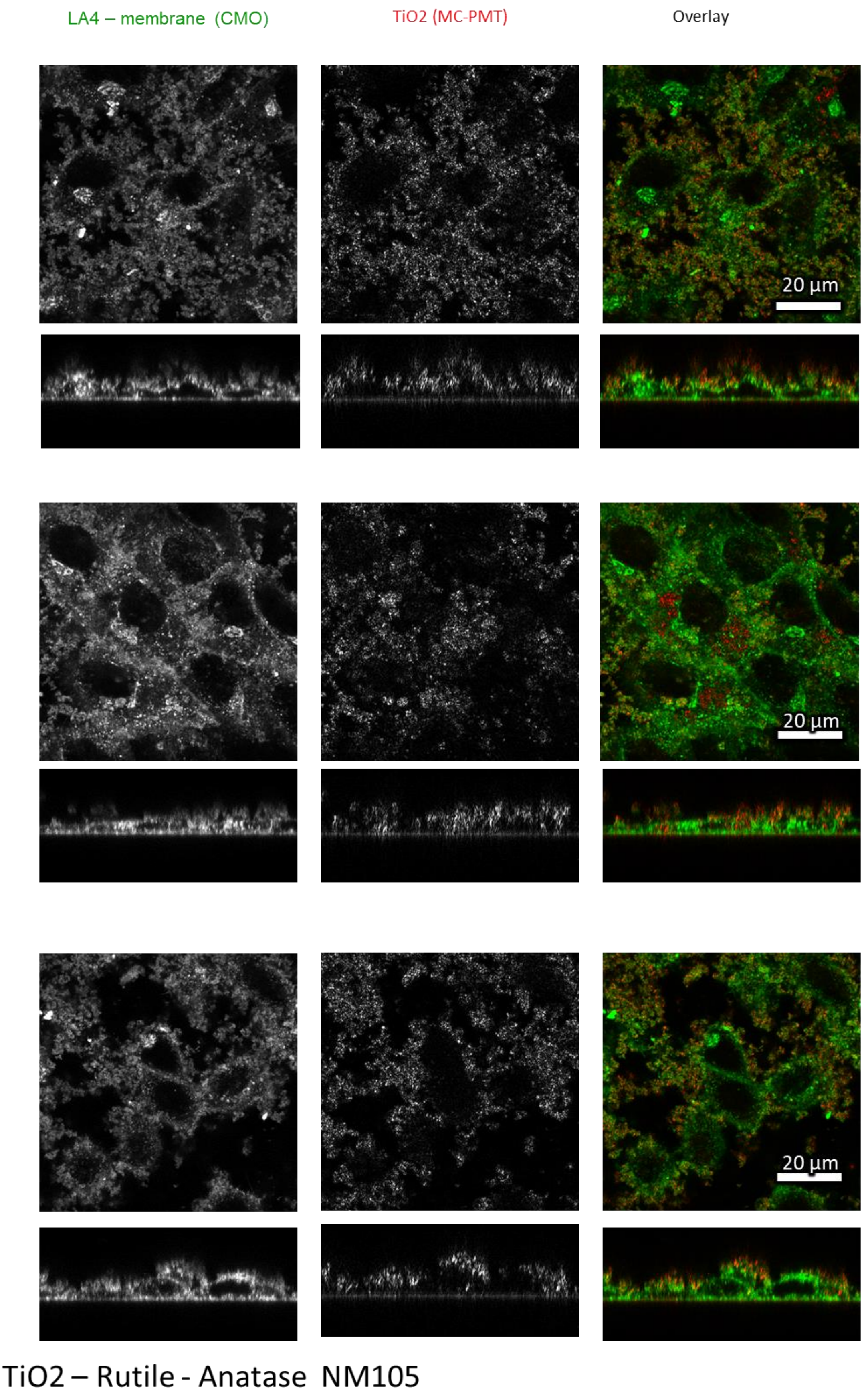

*Figure S47: Confocal images of LA-4 cells exposed to 10 to 1 surface dose of $TiO_2$ rutile-anatase for 48 h in xy plane (upper windows) and xz plane (lower windows)*

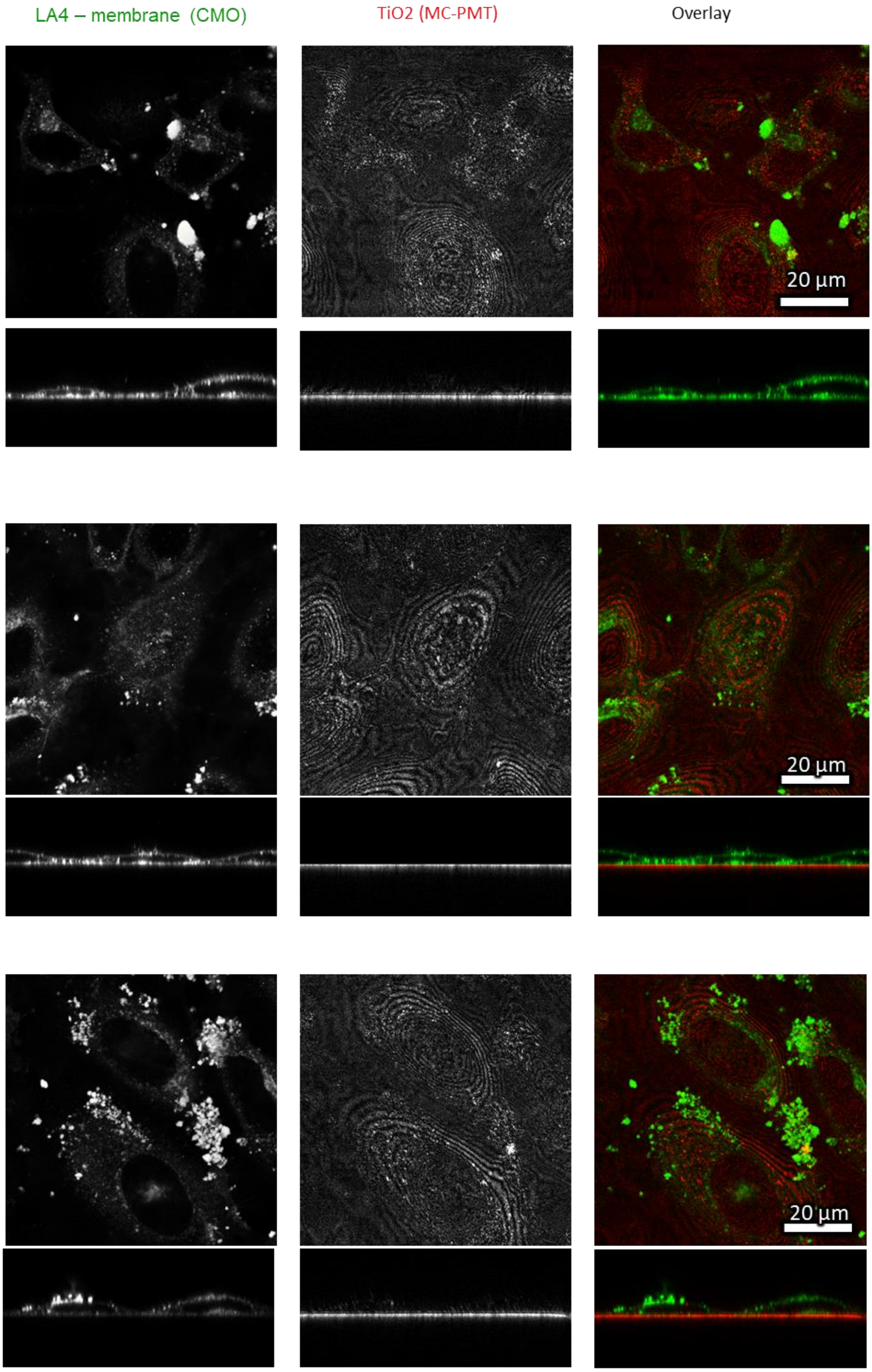

### SiO2 – NM200

*Figure S48: Confocal images of LA-4 cells exposed to 10 to 1 surface dose of $SiO_2$ for 48 h in xy plane (upper windows) and xz plane (lower windows)*

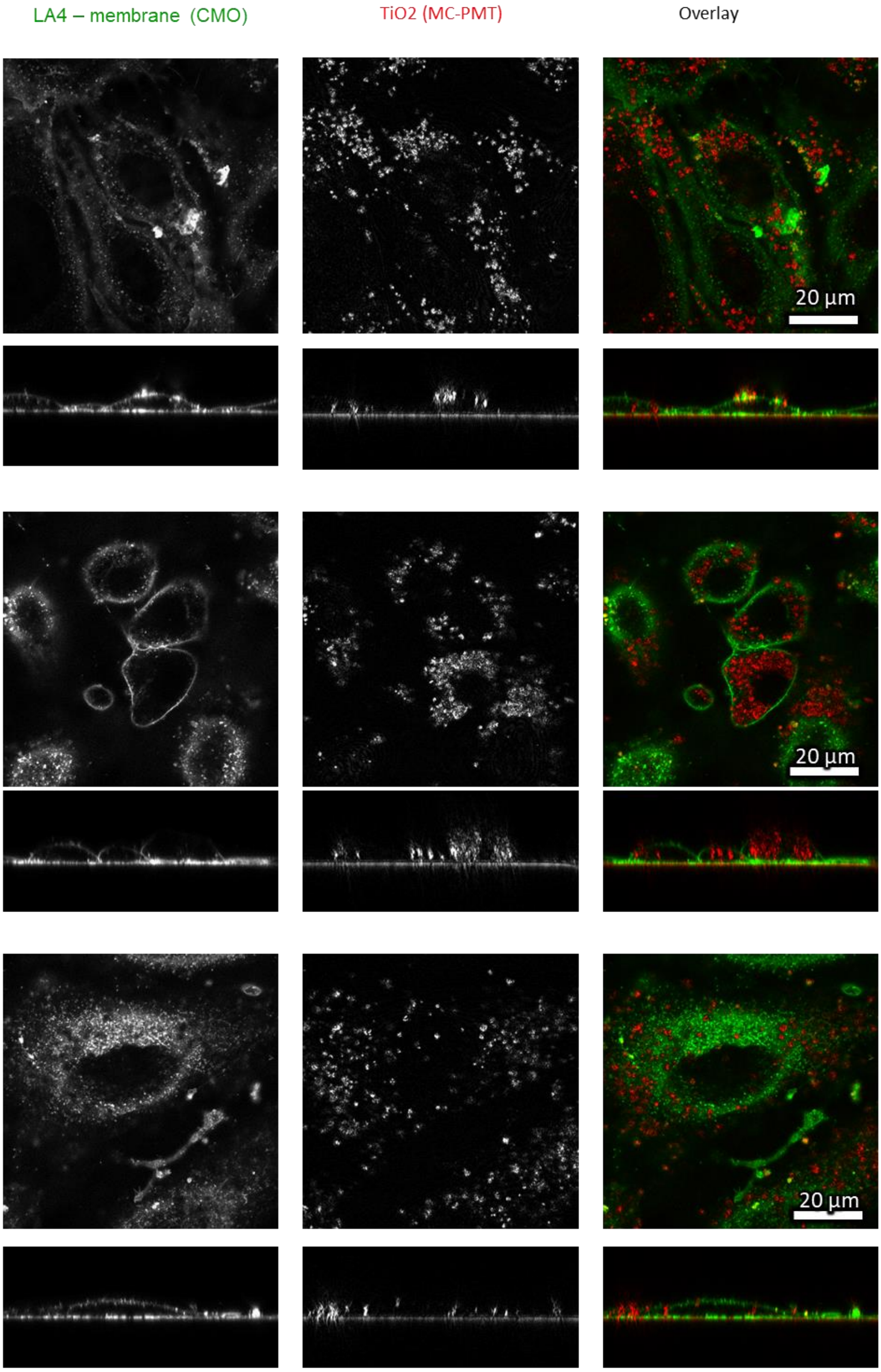

#### TiO2 – Anatase NM101

*Figure S49: Confocal images of LA-4 cells exposed to 10 to 1 surface dose of $TiO_2$ anatase NM101 for 48 h in xy plane (upper windows) and xz plane (lower windows)*

| | |
|---|---|
| Cell line | LA-4 (membrane, CMO) |
| NPs | TiO2 (Star 520) |
| exposure | 10:1, 48 h |
| imaging | xz confocal, 48 h |

| | |
|---|---|
| pixelsize (x,y) | 100 nm |
| FOV (x,y) | 80um |
| pixelsize (z) | 100nm |
| FOV (z) | 35um |
| imaging time | /min |
| number z-stacks | / |

| | |
|---|---|
| 561nm | 100% |
| 640nm | 0% |
| Discovery 2PE 999nm | ON |
| filter sets | 605 nm – 625 nm, 650 nm – 720 nm |
| dwell time | 10 ms |
| objective | wi60x (NA1.2) |

| | |
|---|---|
| Cell line | LA-4 (membrane, CMO) |
| NPs | TiO2 (Star 520) |
| exposure | 10:1, 48 h |
| imaging | xy confocal, 48 h |

| | |
|---|---|
| pixelsize (x,y) | 100 nm |
| FOV (x,y) | 80x80um |
| pixelsize (z) | /nm |
| FOV (z) | /um |
| imaging time | /min |
| number z-stacks | / |

| | |
|---|---|
| 561nm | 100% |
| 640nm | 0% |
| Discovery 2PE 999nm | ON |
| filter sets | 605 nm – 625 nm, 650 nm – 720 nm |
| dwell time | 10 ms |
| objective | wi60x (NA1.2) |

#### S1d – Time evolution of cauliflower growth

##### Main message

Proto-cauliflowers start forming after 1h, after 48h big cauliflower-like structures can be observed at dose 10:1 (surface of $TiO_2$-NTs: surface of the cells).

Supporting raw and analysed data:

For 1h, 10:1 :

Figure S50-Figure S53

For 25h, 10:1 :

Figure S54-Figure S56

##### Materials and methods

See supplement S1e – Dose-dependent exposure of LA-4 to $TiO_2$ nanotubes.

##### Experiment names

See supplement S1e – Dose-dependent exposure of LA-4 to $TiO_2$ nanotubes.

##### Controls and statistics

For 1h, 10:1 :

Figure S50-Figure S53

For 25h, 10:1 :

Figure S54-Figure S56

control: 10:1, 1h exposure confocal xy

| | |
|---|---|
| Cell line | LA-4 (membrane, CellMaskOrange) |
| NPs | $TiO_2$ (Alexa647) |
| exposure | 10:1,1h incubaton |
| imaging | xy confocal, @1h |

| | |
|---|---|
| pixelsize (x,y) | 100 nm |
| FOV (x,y) | 80 µm |
| pixelsize (z) | / |
| FOV (z) | / |
| imaging time | / |
| number of frames | / |

| | |
|---|---|
| 561nm | 30% |
| 640nm | 30% |
| STED | 20% |
| filter sets | 605 nm – 625 nm, 650 nm – 720 nm |
| dwell time | 20 µs |
| objective | wi60x (NA1.2) |

control: 10:1, 1h exposure confocal xy

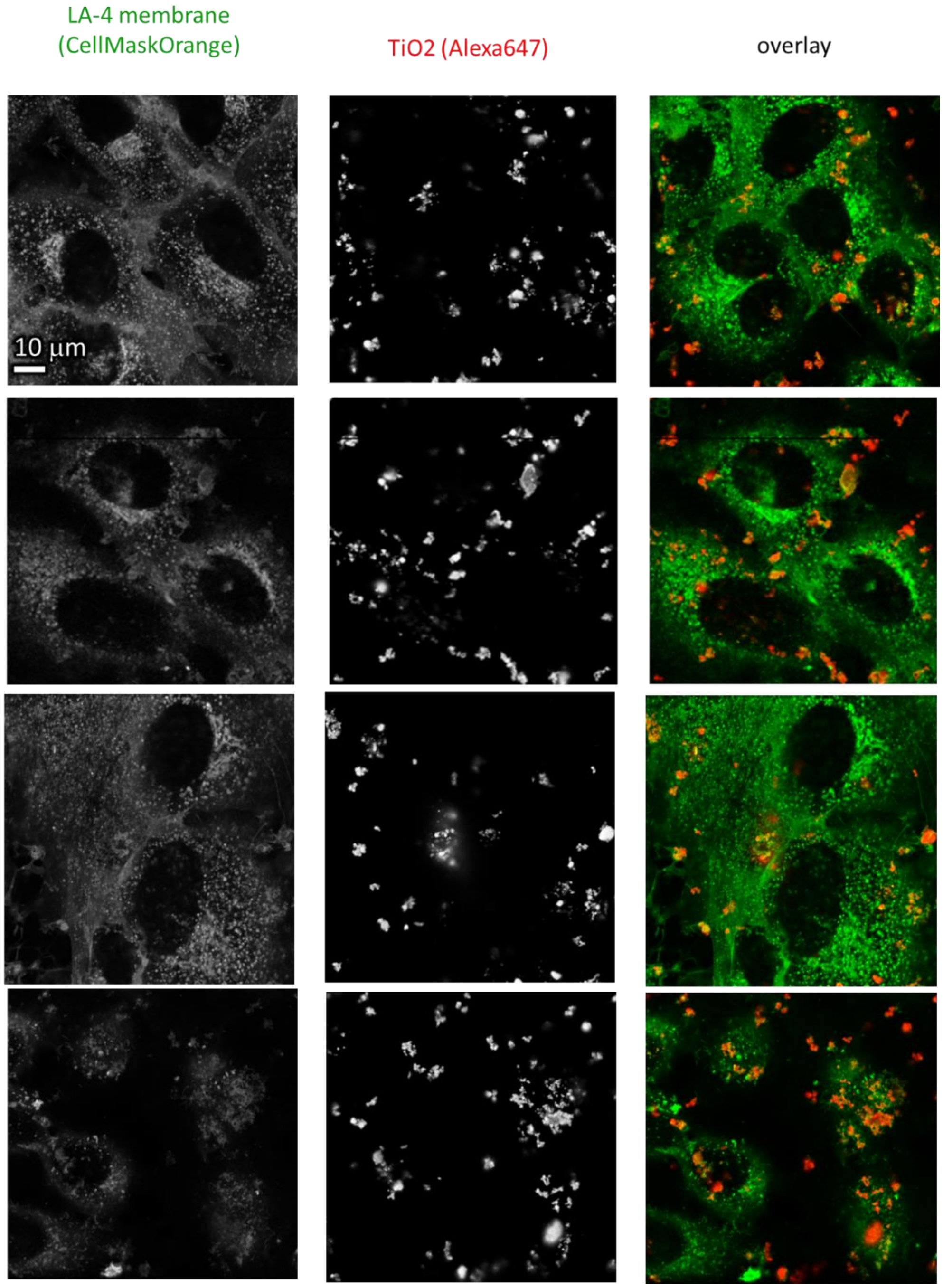

*Figure S50: Statistics for time point 1h – confocal xy recordings.*

control: 10:1, 1h exposure STED xy

| | |
|---|---|
| Cell line | LA-4 (membrane, CellMaskOrange) |
| NPs | $TiO_2$ (Alexa647) |
| exposure | 10:1,1h incubaton |
| imaging | xy confocal, @1h |

| | |
|---|---|
| pixelsize (x,y) | 30 nm |
| FOV (x,y) | 25 µm |
| pixelsize (z) | 62 nm |
| FOV (z) | 12 µm |
| imaging time | / |
| number of frames | / |

| | |
|---|---|
| 561nm | 30% |
| 640nm | 30% |
| STED | 20% |
| filter sets | 605 nm – 625 nm, 650 nm – 720 nm |
| dwell time | 20 µs |
| objective | wi60x (NA1.2) |

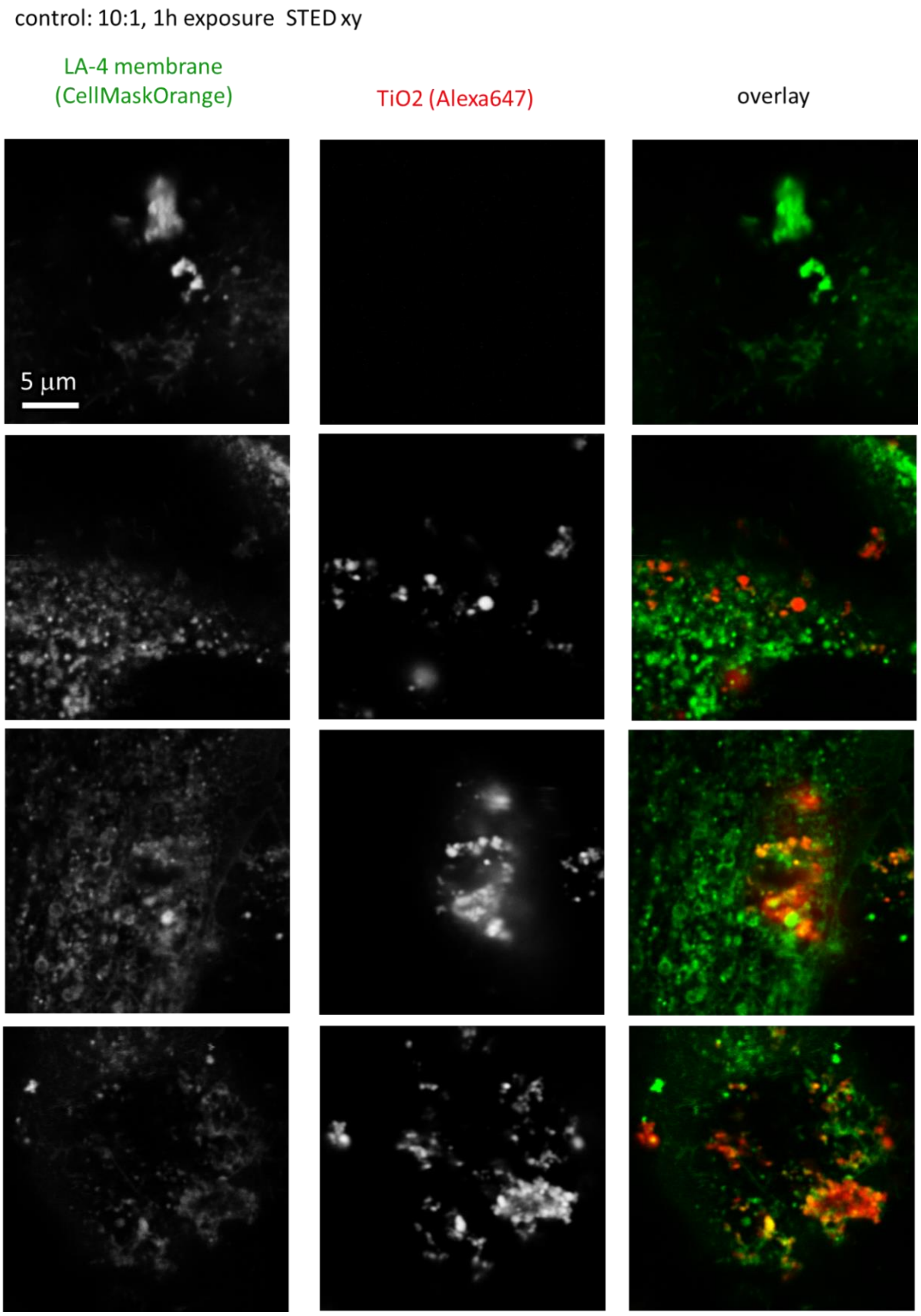

*Figure S51: Statistics for time point 1h – STED xy recordings.*

control: 10:1, 1h exposure  confocal xz

| | |
|---|---|
| Cell line | LA-4 (membrane, CellMaskOrange) |
| NPs | $TiO_2$ (Alexa647) |
| exposure | 10:1,1h incubaton |
| imaging | xy confocal, @1h |

| | |
|---|---|
| pixelsize (x,y) | 100 nm |
| FOV (x,y) | 70 µm |
| pixelsize (z) | 131 nm |
| FOV (z) | 25 µm |
| imaging time | / |
| number of frames | / |

| | |
|---|---|
| 561nm | 30% |
| 640nm | 30% |
| STED | 0 |
| filter sets | 605 nm – 625 nm, 650 nm – 720 nm |
| dwell time | 20 µs |
| objective | wi60x (NA1.2) |

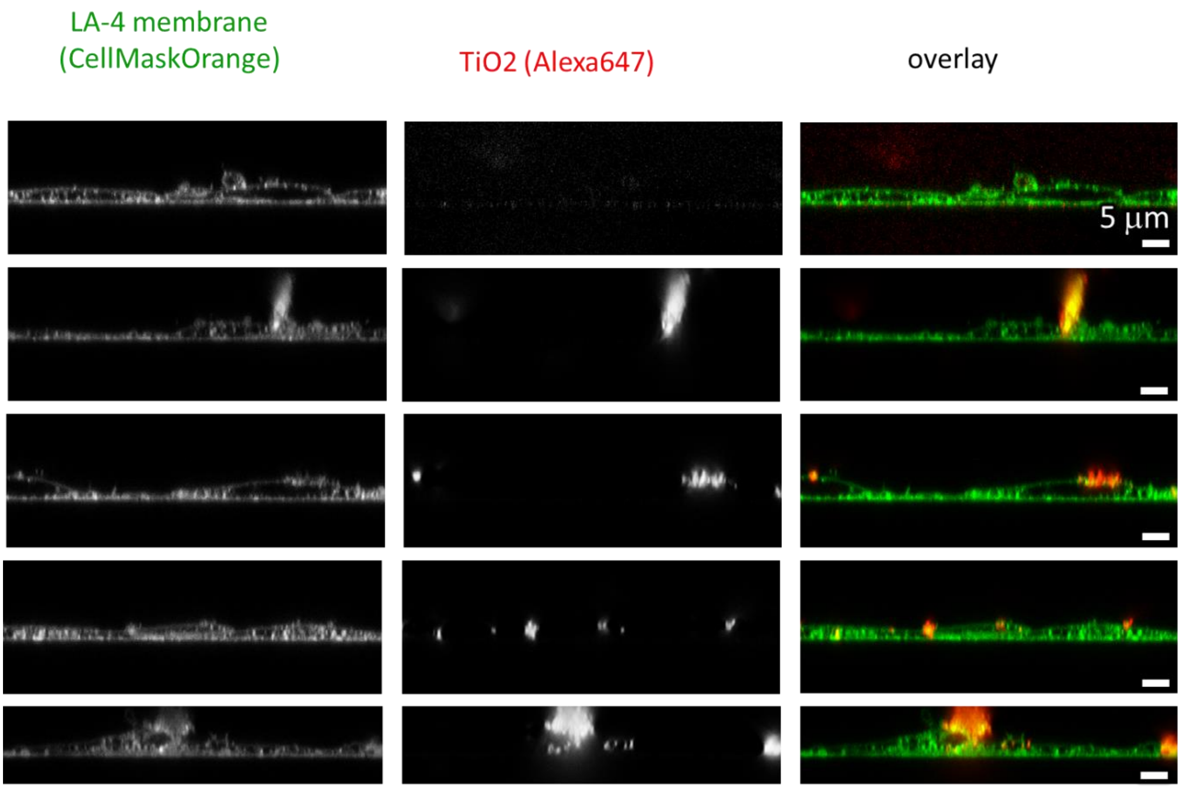

*Figure S52: Statistics for time point 1h – confocal xz recordings.*

control: 10:1, 1h exposure STED xz

| | |
|---|---|
| Cell line | LA-4 (membrane, CellMaskOrange) |
| NPs | $TiO_2$ (Alexa647) |
| exposure | 10:1,1h incubaton |
| imaging | xy confocal, @1h |

| | |
|---|---|
| pixelsize (x,y) | 100 nm |
| FOV (x,y) | 70 µm |
| pixelsize (z) | 131 nm |
| FOV (z) | 25 µm |
| imaging time | / |
| number of frames | / |

| | |
|---|---|
| 561nm | 30% |
| 640nm | 30% |
| STED | 0 |
| filter sets | 605 nm – 625 nm, 650 nm – 720 nm |
| dwell time | 20 µs |
| objective | wi60x (NA1.2) |

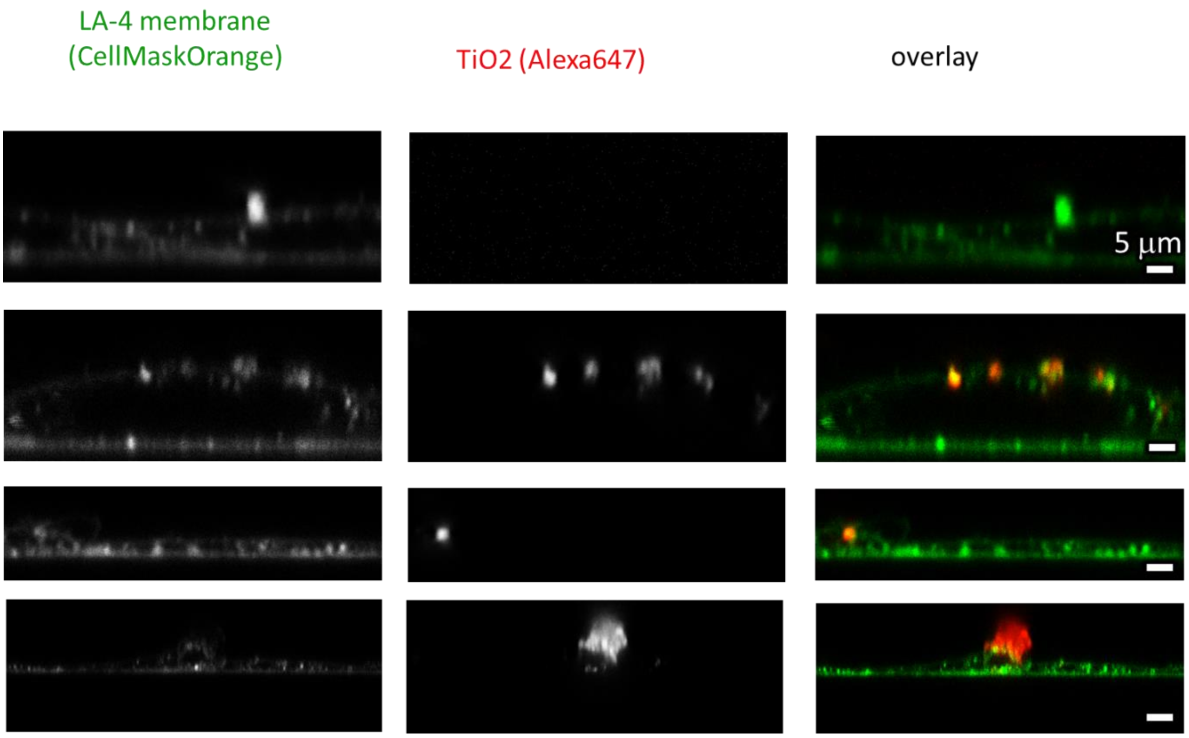

*Figure S53: Statistics for time point 1h – STED xz recordings.*

confocal xy – statistics over sample

| | |
|---|---|
| Cell line | LA-4 (membrane, CellMaskOrange) |
| NPs | $TiO_2$ (Alexa647) |
| exposure | 10:1, 25h incubaton |
| imaging | xy confocal, @25h |

| | |
|---|---|
| pixelsize (x,y) | 100 nm |
| FOV (x,y) | 70 µm |
| pixelsize (z) | / |
| FOV (z) | / |
| imaging time | / |
| number of frames | / |

| | |
|---|---|
| 561nm | 20% |
| 640nm | 20% |
| STED | / |
| filter sets | 605 nm – 625 nm, 650 nm – 720 nm |
| dwell time | 20 µs |
| objective | wi60x (NA1.2) |

| | |
|---|---|
| Cell line | LA-4 (membrane, CellMaskOrange) |
| NPs | $TiO_2$ (Alexa647) |
| exposure | 10:1, 25h incubaton |
| imaging | xy confocal, @25h |

| | |
|---|---|
| pixelsize (x,y) | 300 nm |
| FOV (x,y) | 70 µm |
| pixelsize (z) | / |
| FOV (z) | / |
| imaging time | / |
| number of frames | / |

| | |
|---|---|
| 561nm | 20% |
| 640nm | 20% |
| STED | / |
| filter sets | 605 nm – 625 nm, 650 nm – 720 nm |
| dwell time | 20 µs |
| objective | wi60x (NA1.2) |

control: 10:1, 25h exposure confocal xy

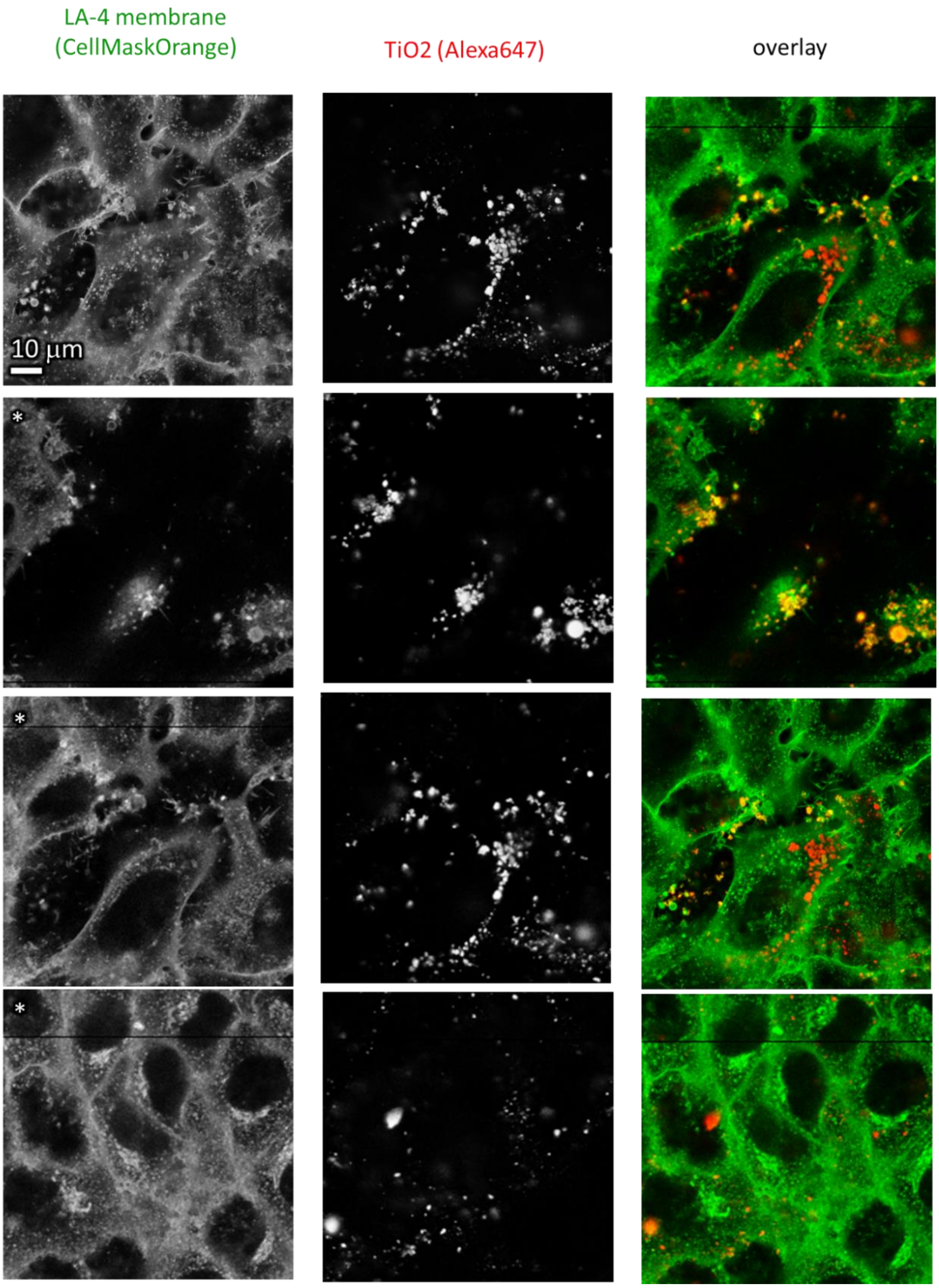

*Figure S54: Statistics for time point 25h – confocal xy recordings.*

control: 10:1, 25h exposure  confocal xz

| | |
|---|---|
| Cell line | LA-4 (membrane, CellMaskOrange) |
| NPs | $TiO_2$ (Alexa647) |
| exposure | 10:1,25h incubaton |
| imaging | xy confocal, @1h |

| | |
|---|---|
| pixelsize (x,y) | 100 nm |
| FOV (x,y) | 70 µm |
| pixelsize (z) | 131 nm |
| FOV (z) | 25 µm |
| imaging time | / |
| number of frames | / |

| | |
|---|---|
| 561nm | 30% |
| 640nm | 30% |
| STED | 0 |
| filter sets | 605 nm – 625 nm, 650 nm – 720 nm |
| dwell time | 20 µs |
| objective | wi60x (NA1.2) |

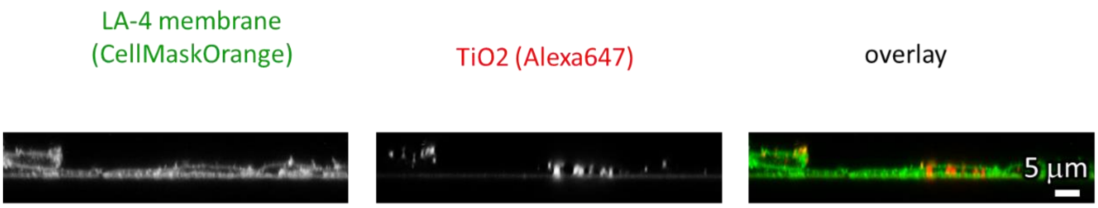

*Figure S55: Statistics for time point 25h – confocal xz recordings.*

control: 10:1, 25h exposure  STED xz

| | |
|---|---|
| Cell line | LA-4 (membrane, CellMaskOrange) |
| NPs | $TiO_2$ (Alexa647) |
| exposure | 10:1,25h incubaton |
| imaging | xy confocal, @25h |

| | |
|---|---|
| pixelsize (x,y) | 100 nm |
| FOV (x,y) | 70 µm |
| pixelsize (z) | 131 nm |
| FOV (z) | 25 µm |
| imaging time | / |
| number of frames | / |

| | |
|---|---|
| 561nm | 30% |
| 640nm | 30% |
| STED | 0 |
| filter sets | 605 nm – 625 nm, 650 nm – 720 nm |
| dwell time | 20 µs |
| objective | wi60x (NA1.2) |

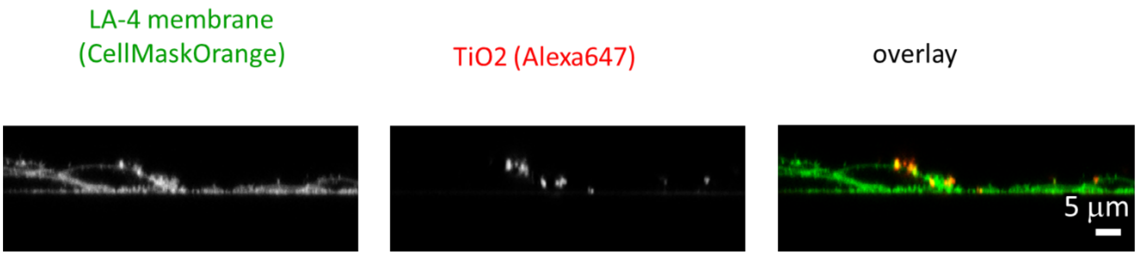

*Figure S56: Statistics for time point 25h – STED xz recordings.*

#### S1e – Dose-dependent exposure of LA-4 to $TiO_2$ nanotubes

##### Main message

With increasing dose, the amount of nanomaterial in cauliflower-like structures increases. At low doses the amount of nanomaterial in the cells also increases but remains constant from the dose 10:1 upwards (surface of $TiO_2$-NTs: surface of the cells).

Supporting raw and analysed data:

Figure S57-Figure S76

##### Materials and methods

- experiment (dose dependence):
    - LA-4 cells were seeded @30% confluence in an Ibidi Ibidi #1.5H µ-Dish
    - after 48 hours freshly filtered 1 mg/ml $TiO_2$-17-Alexa647 in PBS was mixed into 315 µl fresh media, PBS was added to the final volume of 350 µl and added to the cells. The volume of added nanoparticles for each sample was 3.5 µl for 1:1 surface dose ($S_{NPs}$:$S_{cellsurface}$) and 35 µL for 10:1 surface dose. For surface dose 100:1, the nanoparticles were filtered directly into full cell medium.
    - after exposure for 53 hours, media with nanoparticles was slowly removed and the cells were incubated with 5 µg/ml CellMaskOrange for 6 minutes at 37C, afterwards media was carefully exchanged with 100 µl LCIS, cells have been observed at room temperature.
- experiment (1 h incubation):
    - LA-4 cells were seeded @60% confluence in an Ibidi #1.5H µ-Dish
    - after 48 hours, the cells were incubated with 1.6 µg/ml CellMaskOrange in PBS for 6 minutes at 37C and 5% CO2, afterwards they were flushed with LCIS and observed in 315 µl LCIS on a heated insert on the microscope (28C)
    - on the microscpe, 35 µl freshly filtered 1 mg/ml $TiO_2$-40-Alexa647 in PBS were added to cells to achieve a 10:1 surface dose. Cells were observed the first 3 hours after incubation
    - experiment (25 h incubation):
    - LA-4 cells were seeded @60% confluence in an Ibidi #1.5H µ-Dish
    - after 23 hours 35 µl freshly filtered 1 mg/ml $TiO_2$-40-Alexa647 in 100x dcb was mixed into 350 µl fresh media on cells to achieve 10:1 surface dose
    - after exposure for 25 hours, the cells were incubated with 1.6 µg/ml CellMaskOrange in PBS for 6 minutes at 37C and 5% CO2, afterwards they were flushed with LCIS and observed in 315 µl LCIS on a heated insert on the microscope (28C)
- Analysis:
    - threshold of both channels in all STED images has been set to 1 count, since this is the noise of our detector
    - maximum in all STED images in both channels has been adjusted for maximal visibility
    - all STED images in the supplement have been analyzed in same manner
    - brightness of all confocal overlays has also been adjusted for maximal visibility (in supplement)
    - all images in supplement are represented with brightness/threshold adjusted overlay and non-contrasted raw (.png) of both channel separately

#### Experiment names

- Main experiment name:
    - 20191104/e03_s01_t01_LA-4 CellMaskOrange_TiO2 Alexa647_STED.msr
    - 20180202/e03_s04_t04_LA-4 CellMask TiO2 Alexa647-wellLabeled - 1 to 1 _ SNPs to Scells - cellsB.msr
    - 20180202/e04_s02_t03_LA-4 CellMask TiO2 Alexa647-wellLabeled - 10 to 1 _SNPs to Scells - cellsG.msr
    - 20180202/e05_s06_t07_LA-4 CellMask TiO2 Alexa647-wellLabeled - 100to 1 _SNPs to Scells - cellsH-biggerZ.msr
    - 20180202/e06_s02_t02_LA-4 CellMask - control - cellsA.msr
    - 20180202/e04_s03_t04_LA-4 CellMask TiO2 Alexa647-wellLabeled - 10 to 1 _SNPs to Scells - cellsG.msr
    - 20190308/e02_t08_LA-4 CellMask TiO2Alexa647 10 to 1.msr
    - 20190308/e03_t01_LA-4 CellMask TiO2Alexa647 10 to 1_25h incubation.msr

- Supplement experiment names:
    - dose dependence - 20180202
        - STED and confocals:
            - e06_s05_t05_LA-4 CellMask - control - cellsA.msr
            - e06_s04_t04_LA-4 CellMask - control - cellsA.msr
            - e06_s03_t03_LA-4 CellMask - control - cellsA.msr
            - e05_s06_t06_LA-4 CellMask TiO2 Alexa647-wellLabeled - 100to 1 _SNPs to Scells - cellsH.msr
            - e05_s05_t05_LA-4 CellMask TiO2 Alexa647-wellLabeled - 100to 1 _SNPs to Scells - cellsH.msr
            - e05_s04_t04_LA-4 CellMask TiO2 Alexa647-wellLabeled - 100to 1 _SNPs to Scells - cellsH.msr
            - e05_s03_t03_LA-4 CellMask TiO2 Alexa647-wellLabeled - 100to 1 _SNPs to Scells - cellsH.msr
            - e04_s04_t05_LA-4 CellMask TiO2 Alexa647-wellLabeled - 10 to 1 _SNPs to Scells - cellsG.msr
            - e03_s04_t05_LA-4 CellMask TiO2 Alexa647-wellLabeled - 1 to 1 _ SNPs to Scells - cellsB.msr
            - e03_s03_t03_LA-4 CellMask TiO2 Alexa647-wellLabeled - 1 to 1 _ SNPs to Scells - cellsB.msr
            - e03_s02_t02_LA-4 CellMask TiO2 Alexa647-wellLabeled - 1 to 1 _ SNPs to Scells - cellsB.msr
            - e03_s01_t01_LA-4 CellMask TiO2 Alexa647-wellLabeled - 1 to 1 _ SNPs to Scells - cellsB.msr
            - e03_s01_t01_LA-4 CellMask TiO2 Alexa647-wellLabeled - 1 to 1 _ SNPs to Scells - cellsB - time.msr
        - confocals:
            - e06_s01_t01_LA-4 CellMask - control - cellsA - time.msr
            - e03_s01_t01_LA-4 CellMask TiO2 Alexa647-wellLabeled - 1 to 1 _ SNPs to Scells - cellsB - time.msr
            - e04_s01_t01_LA-4 CellMask TiO2 Alexa647-wellLabeled - 10 to 1 _SNPs to Scells - cellsG - time.msr
            - e05_s01_t01_LA-4 CellMask TiO2 Alexa647-wellLabeled - 100to 1 _SNPs to Scells - cellsH - time.msr
            - e05_s02_t02_LA-4 CellMask TiO2 Alexa647-wellLabeled - 100to 1 _SNPs to Scells - cellsH - time.msr

#### Controls and statistics

control – no exposure to nanomaterial:

Figure S57-Figure S59

Figure S71-Figure S72

1:1 surface to surface dose:

Figure S58-Figure S62

Figure S71-Figure S72

10:1 surface to surface dose:

Figure S63-Figure S65

Figure S73-Figure S74

100:1 surface to surface dose:

Figure S66-Figure S68

Figure S75-Figure S76

control: 0:1 exposure STED xz

| | |
|---|---|
| Cell line | LA-4 (membrane, CellMaskOrange) |
| NPs | / |
| exposure | / |
| imaging | xz STED, @48h |

| | |
|---|---|
| pixelsize (x,y) | 30 nm |
| FOV (x,y) | 30 µm |
| pixelsize (z) | 85 nm |
| FOV (z) | 20 µm |
| imaging time | / |
| number of frames | / |

| | |
|---|---|
| 561nm | 30% |
| 640nm | 3% |
| STED | 20% |
| filter sets | 605 nm – 625 nm, 650 nm – 720 nm |
| dwell time | 80 µs |
| objective | wi60x (NA1.2) |

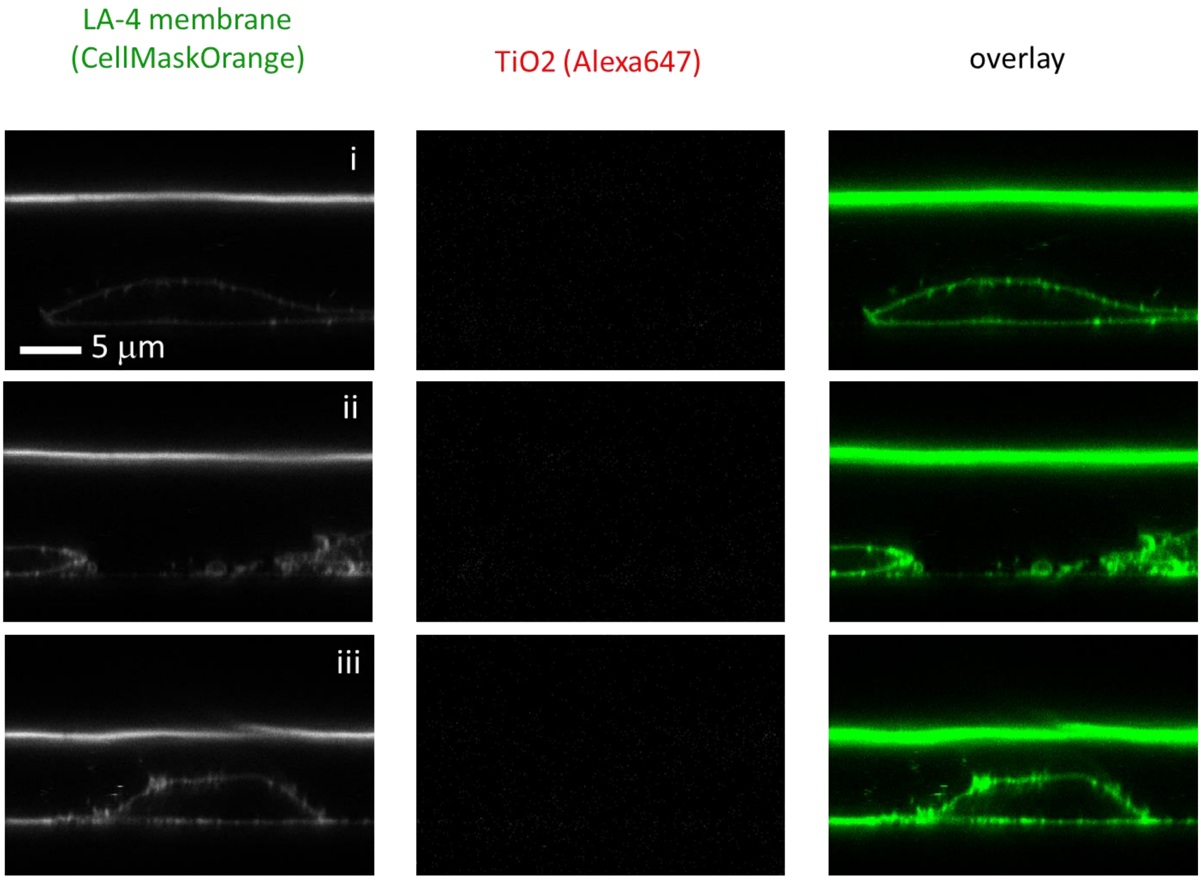

*Figure S57: Statistics for control with no exposure to nanomaterial – STED xz recordings.*

control: 0:1 exposure STED xy

| | |
|---|---|
| Cell line | LA-4 (membrane, CellMaskOrange) |
| NPs | / |
| exposure | / |
| imaging | xy STED, @48h |

| | |
|---|---|
| pixelsize (x,y) | 30 nm |
| FOV (x,y) | 30 µm |
| pixelsize (z) | / |
| FOV (z) | / |
| imaging time | / |
| number of frames | / |

| | |
|---|---|
| 561nm | 30% |
| 640nm | 3% |
| STED | 20% |
| filter sets | 605 nm – 625 nm, 650 nm – 720 nm |
| dwell time | 80 µs |
| objective | wi60x (NA1.2) |

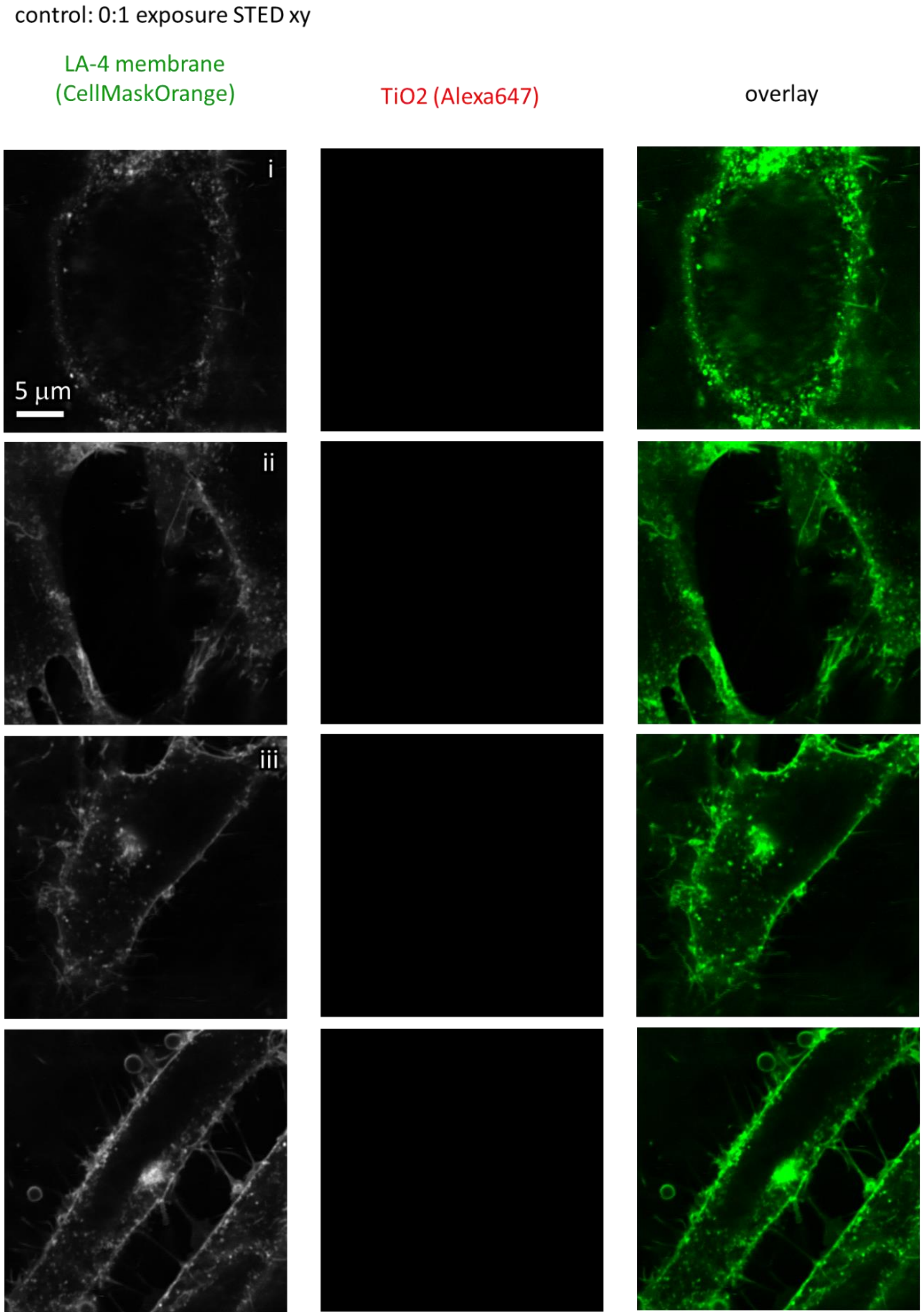

*Figure S58: Statistics for control with no exposure to nanomaterial – STED xy recordings.*

control: 0:1 exposure confocal xy (STED is a zoom)

| | |
|---|---|
| Cell line | LA-4 (membrane, CellMaskOrange) |
| NPs | / |
| exposure | / |
| imaging | xy confocal, @48h |

| | |
|---|---|
| pixelsize (x,y) | 100 nm |
| FOV (x,y) | 80 µm |
| pixelsize (z) | / |
| FOV (z) | / |
| imaging time | / |
| number of frames | / |

| | |
|---|---|
| 561nm | 30% |
| 640nm | 3% |
| STED | 20% |
| filter sets | 605 nm – 625 nm, 650 nm – 720 nm |
| dwell time | 80 µs |
| objective | wi60x (NA1.2) |

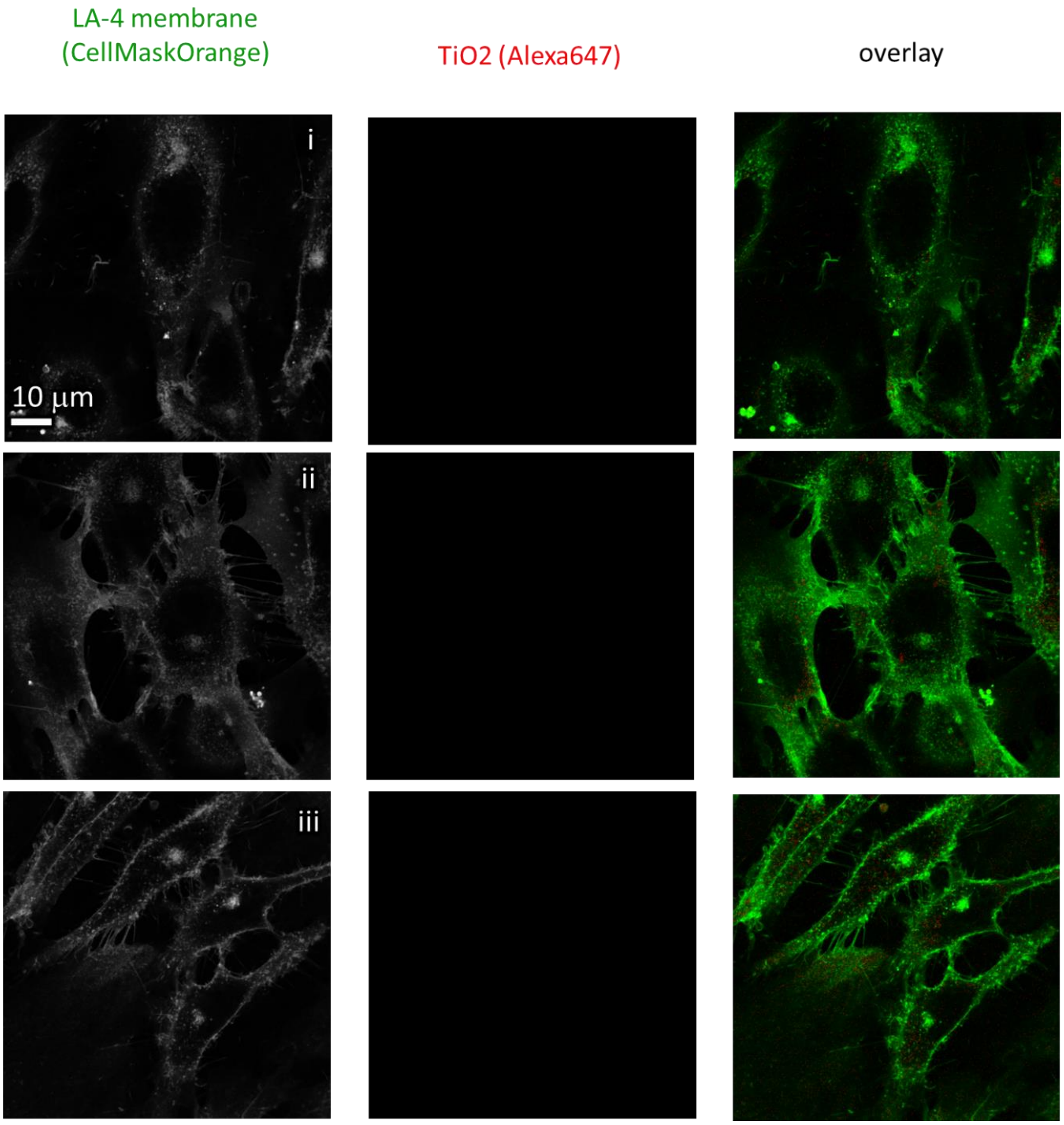

*Figure S59: Statistics for control with no exposure to nanomaterial – confocal xy recordings.*

exposure: 1:1 exposure STED xz

| | |
|---|---|
| Cell line | LA-4 (membrane, CellMaskOrange) |
| NPs | $TiO_2$ (Alexa647) |
| exposure | 1:1, 48h incubaton |
| imaging | xz STED, @48h |

| | |
|---|---|
| pixelsize (x,y) | 30 nm |
| FOV (x,y) | 30 µm |
| pixelsize (z) | 85 nm |
| FOV (z) | 20 µm |
| imaging time | / |
| number of frames | / |

| | |
|---|---|
| 561nm | 30% |
| 640nm | 3% |
| STED | 20% |
| filter sets | 605 nm – 625 nm, 650 nm – 720 nm |
| dwell time | 80 µs |
| objective | wi60x (NA1.2) |

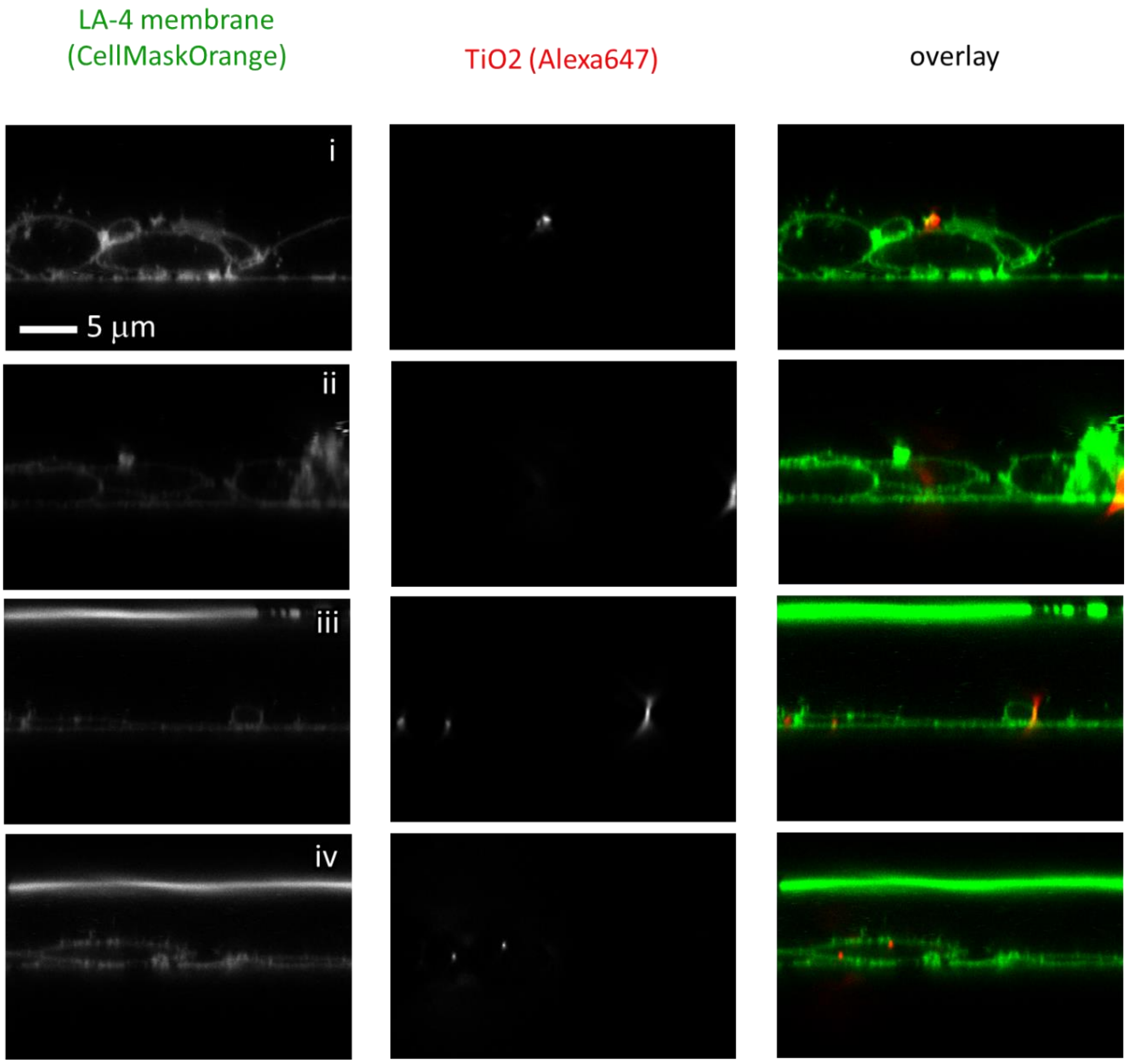

*Figure S60: Statistics at surface nanoparticles to surface cells dose 1:1 – STED xz recordings.*

control: 1:1 exposure STED xy

| | |
|---|---|
| Cell line | LA-4 (membrane, CellMaskOrange) |
| NPs | $TiO_2$ (Alexa647) |
| exposure | 1:1, 48h incubaton |
| imaging | xy STED, @48h |

| | |
|---|---|
| pixelsize (x,y) | 30 nm |
| FOV (x,y) | 30 µm |
| pixelsize (z) | / |
| FOV (z) | / |
| imaging time | / |
| number of frames | / |

| | |
|---|---|
| 561nm | 30% |
| 640nm | 3% |
| STED | 20% |
| filter sets | 605 nm – 625 nm, 650 nm – 720 nm |
| dwell time | 80 µs |
| objective | wi60x (NA1.2) |

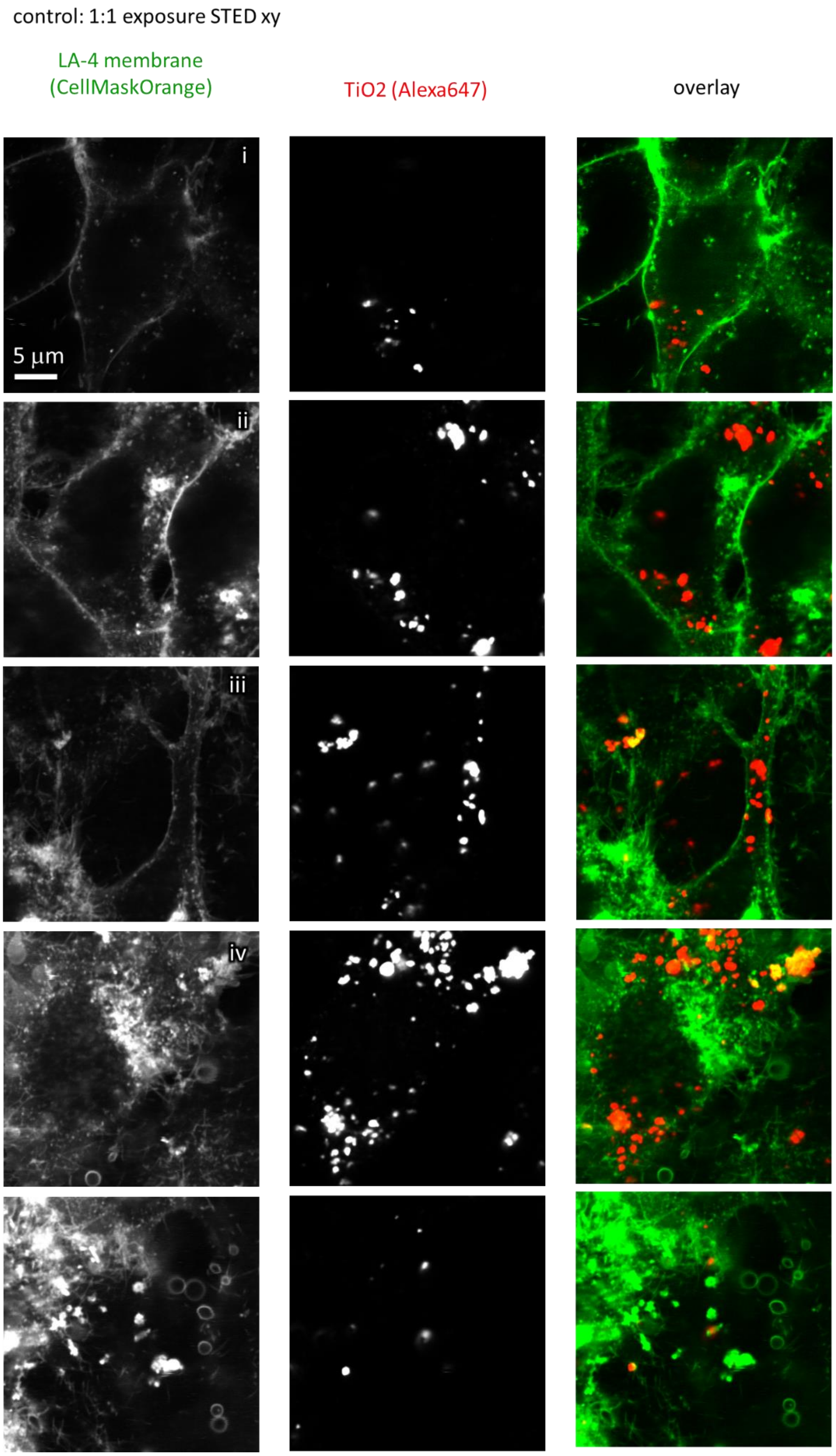

*Figure S61: Statistics at surface nanoparticles to surface cells dose 1:1 – STED xy recordings.*

control: 1:1 confocal xy

| | |
|---|---|
| Cell line | LA-4 (membrane, CellMaskOrange) |
| NPs | $TiO_2$ (Alexa647) |
| exposure | 1:1, 48h incubaton |
| imaging | xy confocal, 48h |

| | |
|---|---|
| pixelsize (x,y) | 100 nm |
| FOV (x,y) | 80 µm |
| pixelsize (z) | / |
| FOV (z) | / |
| imaging time | / |
| number of frames | / |

| | |
|---|---|
| 561nm | 30% |
| 640nm | 3% |
| STED | / |
| filter sets | 605 nm – 625 nm, 650 nm – 720 nm |
| dwell time | 80 µs |
| objective | wi60x (NA1.2) |

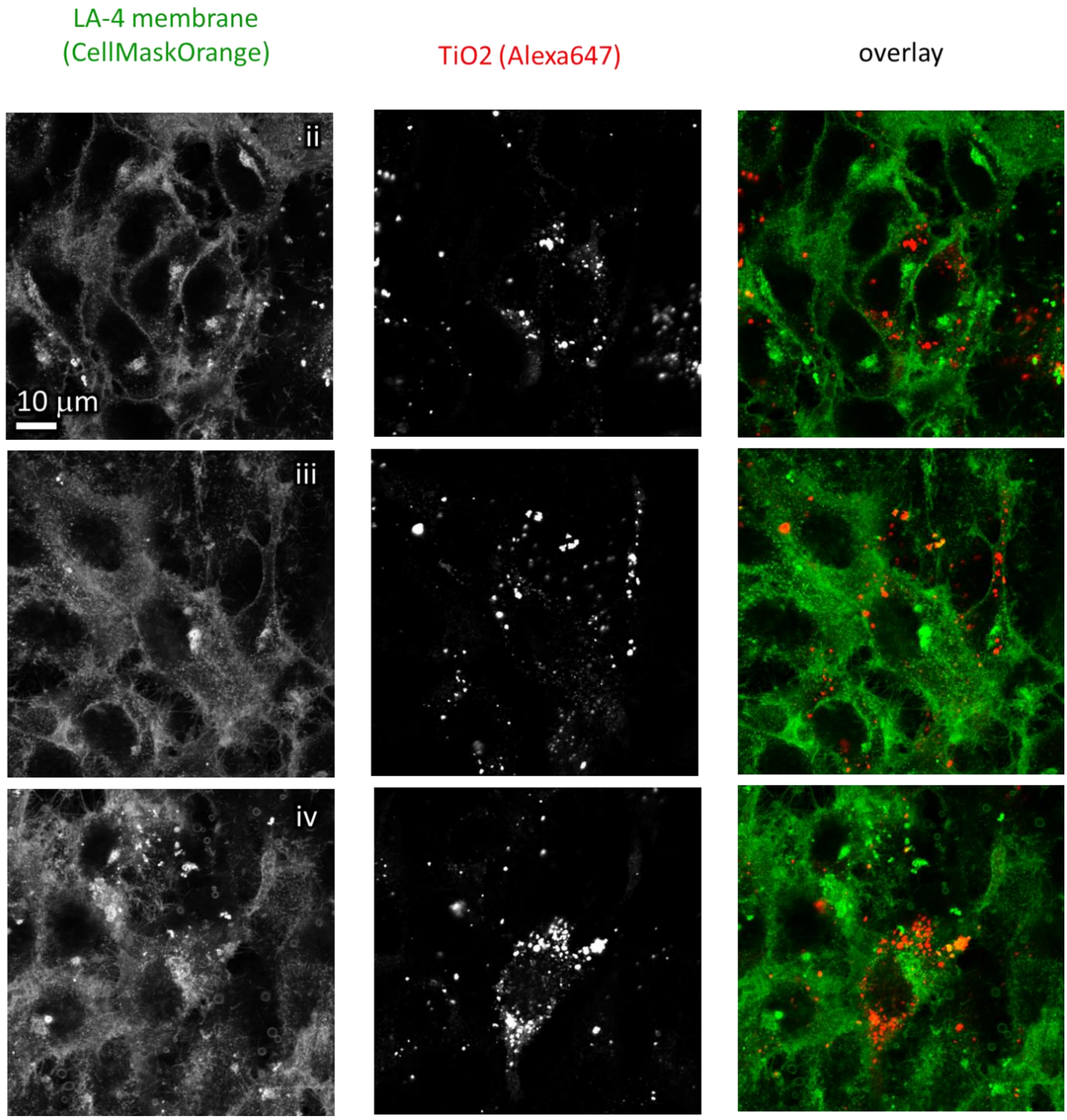

*Figure S62: Statistics at surface nanoparticles to surface cells dose 1:1 – confocal xy recordings.*

exposure: 10:1 exposure STED xz

| | |
|---|---|
| Cell line | LA-4 (membrane, CellMaskOrange) |
| NPs | $TiO_2$ (Alexa647) |
| exposure | 10:1, 48h incubaton |
| imaging | xz STED, @48h |

| | |
|---|---|
| pixelsize (x,y) | 30 nm |
| FOV (x,y) | 30 µm |
| pixelsize (z) | 85 nm |
| FOV (z) | 20 µm |
| imaging time | / |
| number of frames | / |

| | |
|---|---|
| 561nm | 30% |
| 640nm | 3% |
| STED | 20% |
| filter sets | 605 nm – 625 nm, 650 nm – 720 nm |
| dwell time | 80 µs |
| objective | wi60x (NA1.2) |

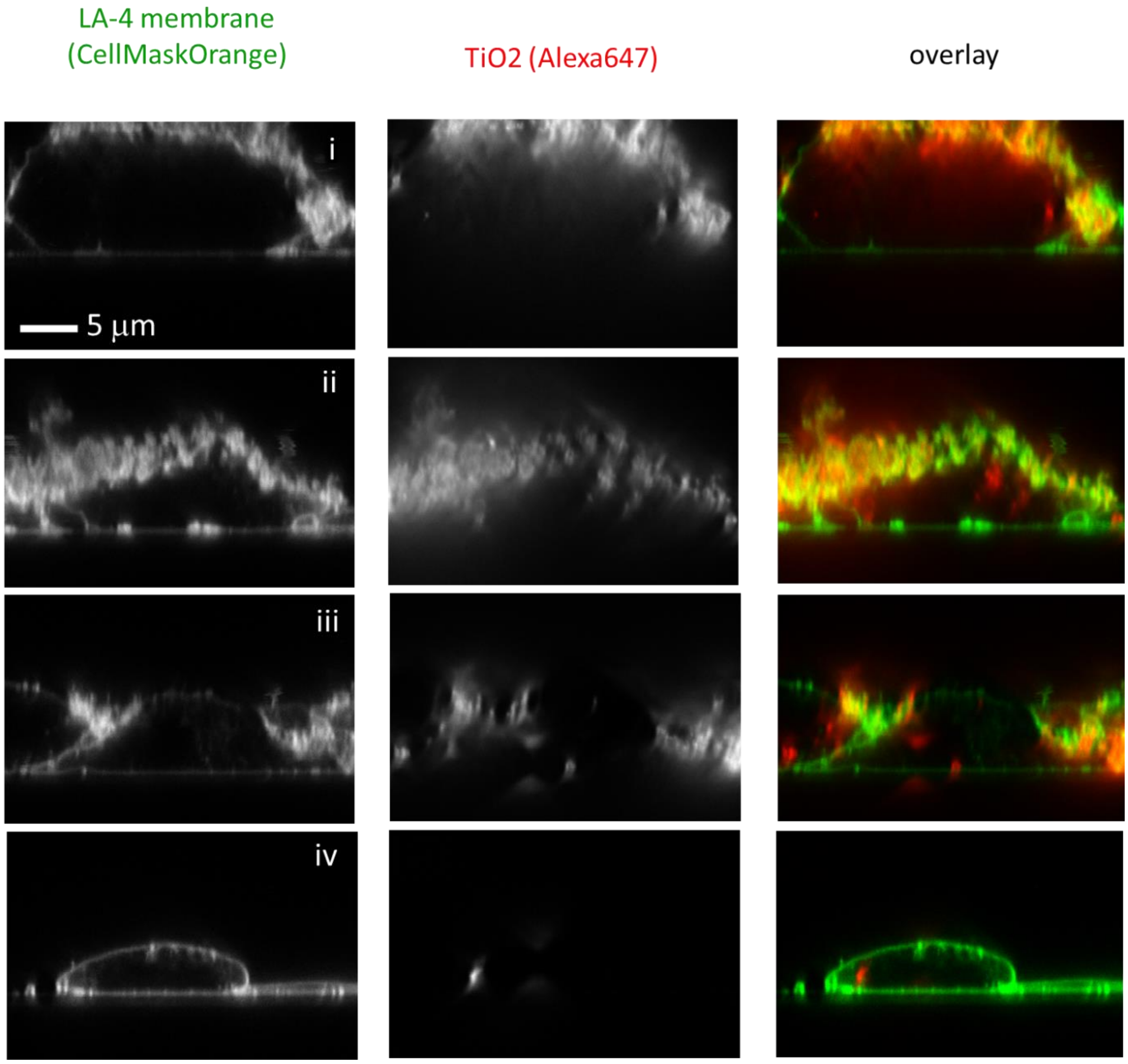

*Figure S63: Statistics at surface nanoparticles to surface cells dose 10:1 – STED xz recordings.*

| | |
|---|---|
| Cell line | LA-4 (membrane, CellMaskOrange) |
| NPs | $TiO_2$ (Alexa647) |
| exposure | 10:1, 48h incubaton |
| imaging | xy STED, 48h |

| | |
|---|---|
| pixelsize (x,y) | 30 nm |
| FOV (x,y) | 30 µm |
| pixelsize (z) | / |
| FOV (z) | / |
| imaging time | / |
| number of frames | / |

| | |
|---|---|
| 561nm | 30% |
| 640nm | 3% |
| STED | 20% |
| filter sets | 605 nm – 625 nm, 650 nm – 720 nm |
| dwell time | 80 µs |
| objective | wi60x (NA1.2) |

control: 10:1 exposure STED xy

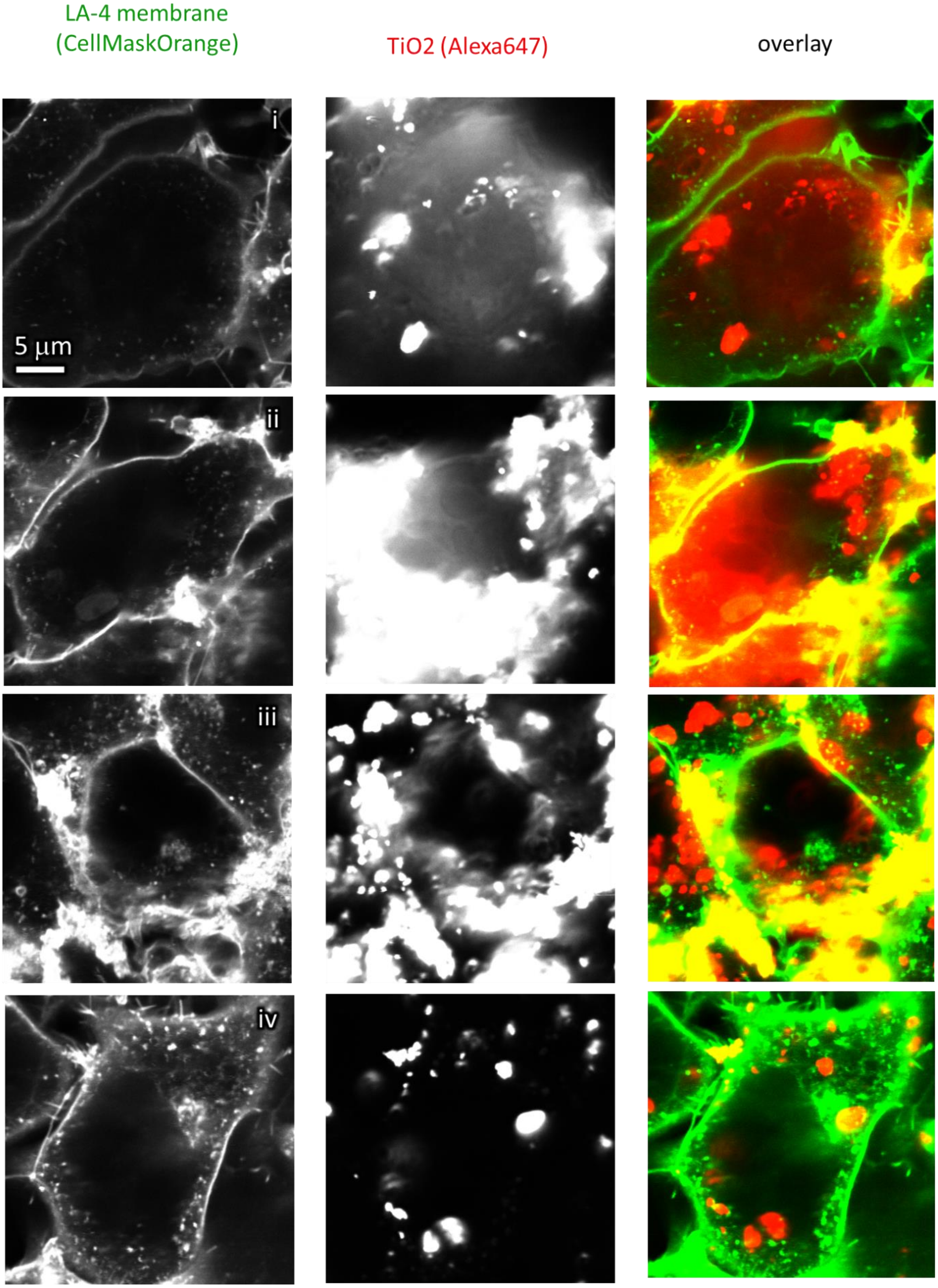

*Figure S64: Statistics at surface nanoparticles to surface cells dose 100:1 – STED xy recordings.*

control: 10:1 exposure confocal xy (STED is a zoom)

| | |
|---|---|
| Cell line | LA-4 (membrane, CellMaskOrange) |
| NPs | $TiO_2$ (Alexa647) |
| exposure | 10:1, 48h incubaton |
| imaging | xy confocal, 48h |

| | |
|---|---|
| pixelsize (x,y) | 100 nm |
| FOV (x,y) | 80 µm |
| pixelsize (z) | / |
| FOV (z) | / |
| imaging time | / |
| number of frames | / |

| | |
|---|---|
| 561nm | 30% |
| 640nm | 3% |
| STED | 20% |
| filter sets | 605 nm – 625 nm, 650 nm – 720 nm |
| dwell time | 80 µs |
| objective | wi60x (NA1.2) |

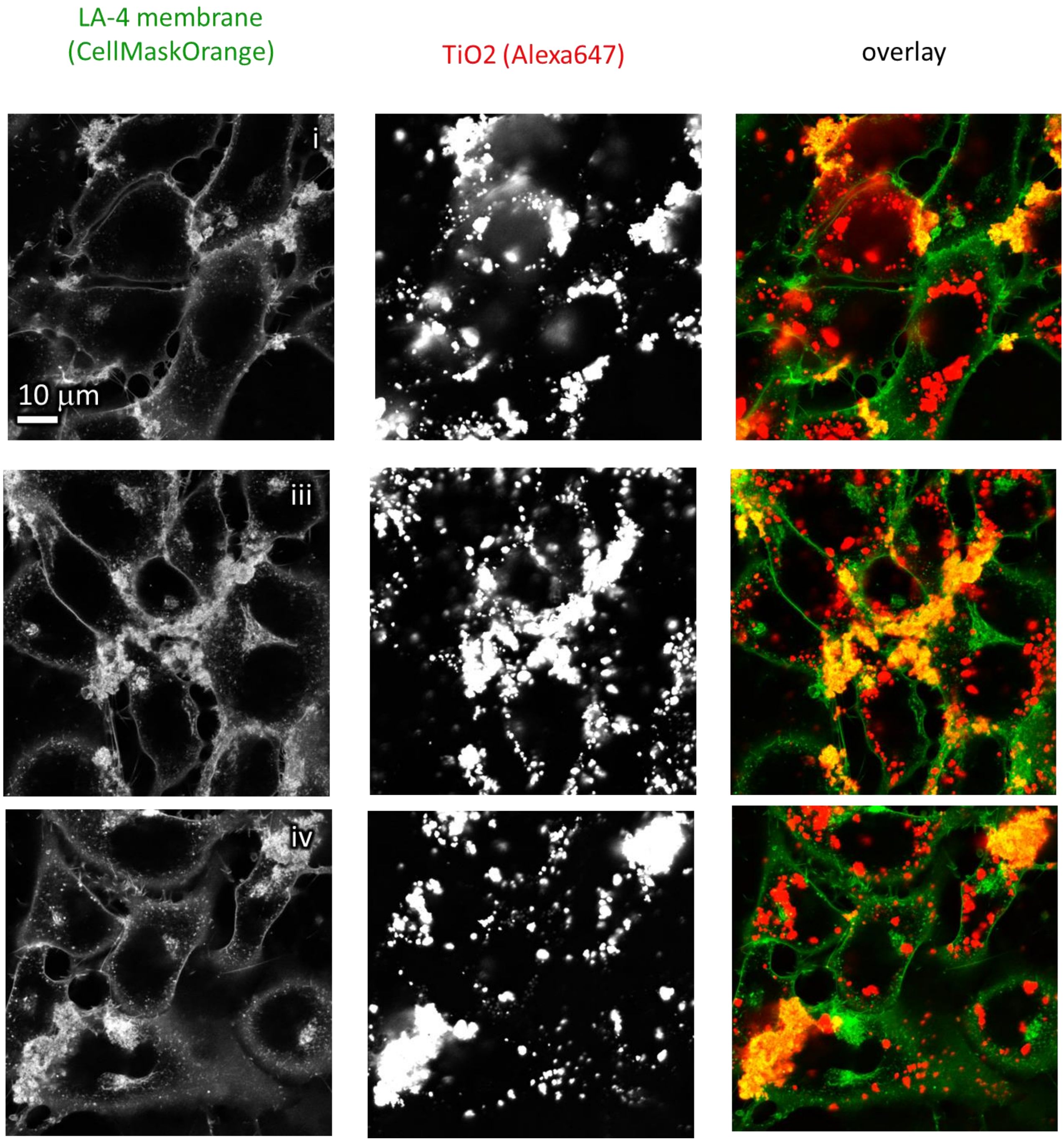

*Figure S65: Statistics at surface nanoparticles to surface cells dose 10:1 – confocal xy recordings.*

exposure: 100:1 exposure STED

| | |
|---|---|
| Cell line | LA-4 (membrane, CellMaskOrange) |
| NPs | $TiO_2$ (Alexa647) |
| exposure | 100:1, 48h incubaton |
| imaging | xz STED, @48h |

| | |
|---|---|
| pixelsize (x,y) | 30 nm |
| FOV (x,y) | 30 µm |
| pixelsize (z) | 85 nm |
| FOV (z) | 20 µm |
| imaging time | / |
| number of frames | / |

| | |
|---|---|
| 561nm | 30% |
| 640nm | 3% |
| STED | 20% |
| filter sets | 605 nm – 625 nm, 650 nm – 720 nm |
| dwell time | 80 µs |
| objective | wi60x (NA1.2) |

| | |
|---|---|
| iv pixelsize (x,y) | 30 nm |
| FOV (x,y) | 30 µm |
| pixelsize (z) | 104 nm |
| FOV (z) | 50 µm |
| imaging time | / |
| number of frames | / |

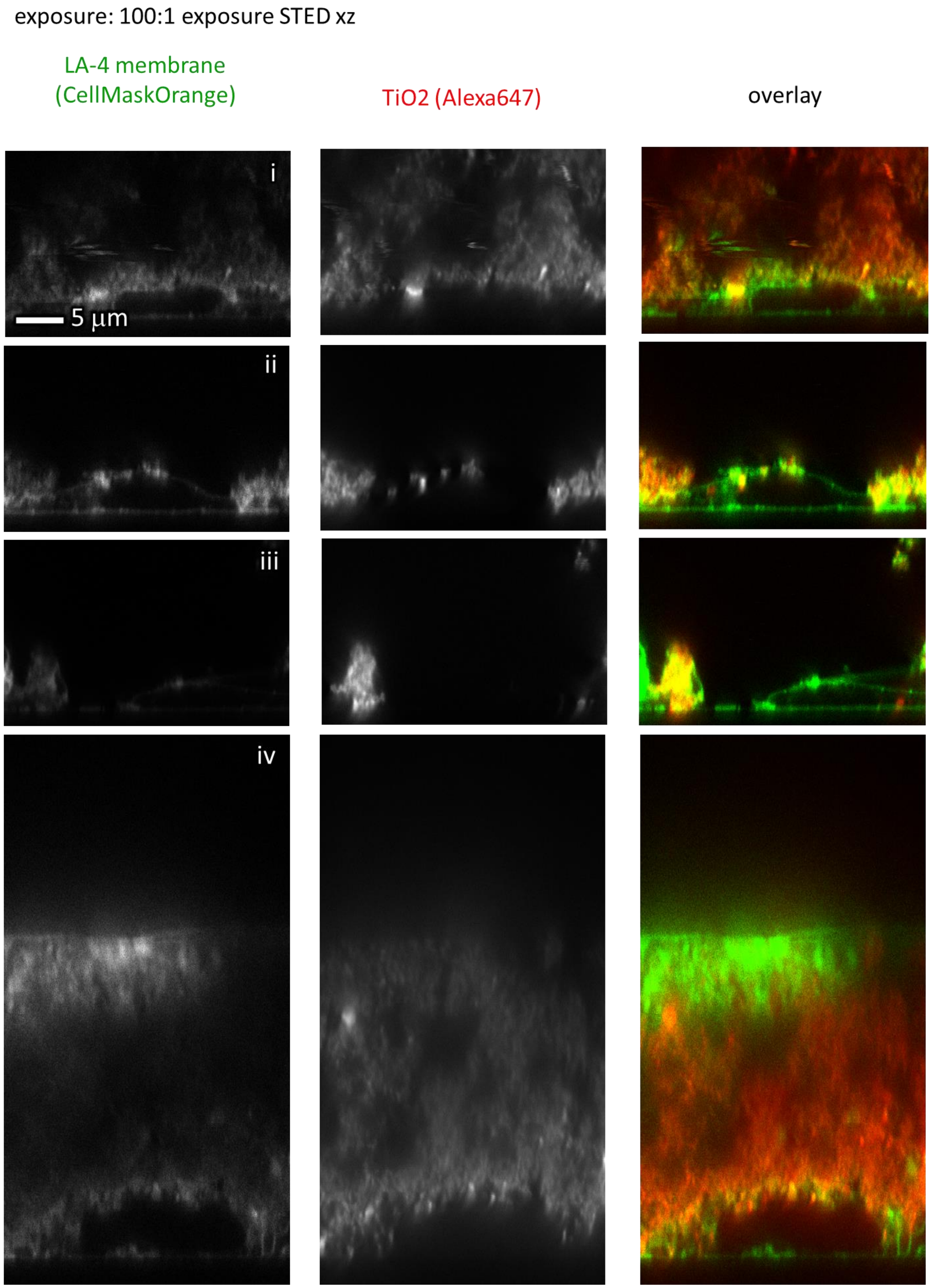

*Figure S66: Statistics at surface nanoparticles to surface cells dose 100:1 – STED xz recordings.*

control: 100:1 exposure STED xy

| | |
|---|---|
| Cell line | LA-4 (membrane, CellMaskOrange) |
| NPs | $TiO_2$ (Alexa647) |
| exposure | 100:1, 48h incubaton |
| imaging | xy STED, @48h |

| | |
|---|---|
| pixelsize (x,y) | 30 nm |
| FOV (x,y) | 30 µm |
| pixelsize (z) | / |
| FOV (z) | / |
| imaging time | / |
| number of frames | / |

| | |
|---|---|
| 561nm | 30% |
| 640nm | 3% |
| STED | 20% |
| filter sets | 605 nm – 625 nm, 650 nm – 720 nm |
| dwell time | 80 µs |
| objective | wi60x (NA1.2) |

iv

| | |
|---|---|
| pixelsize (x,y) | 30 nm |
| FOV (x,y) | 30 µm |
| pixelsize (z) | / |
| FOV (z) | / |
| imaging time | / |
| number of frames | / |

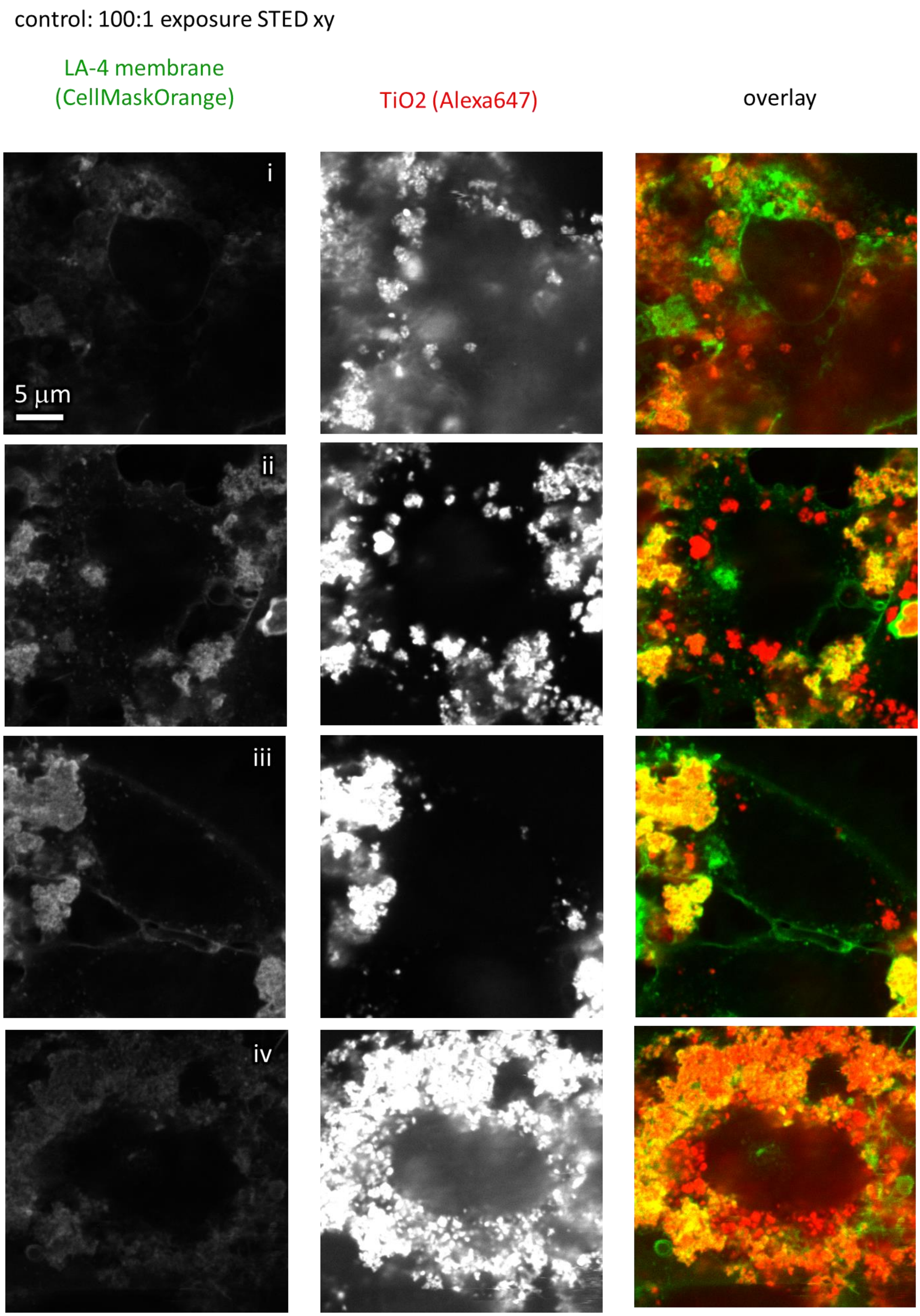

*Figure S67: Statistics at surface nanoparticles to surface cells dose 100:1 – confocal xy recordings.*

control: 100:1 exposure confocal xy

| | |
|---|---|
| Cell line | LA-4 (membrane, CellMaskOrange) |
| NPs | $TiO_2$ (Alexa647) |
| exposure | 100:1, 48h incubaton |
| imaging | xy confocal, @48h |

| | |
|---|---|
| pixelsize (x,y) | 100 nm |
| FOV (x,y) | 80 µm |
| pixelsize (z) | / |
| FOV (z) | / |
| imaging time | / |
| number of frames | / |

| | |
|---|---|
| 561nm | 30% |
| 640nm | 3% |
| STED | 20% |
| filter sets | 605 nm – 625 nm, 650 nm – 720 nm |
| dwell time | 80 µs |
| objective | wi60x (NA1.2) |

LA-4 membrane (CellMaskOrange)

TiO2 (Alexa647)

overlay

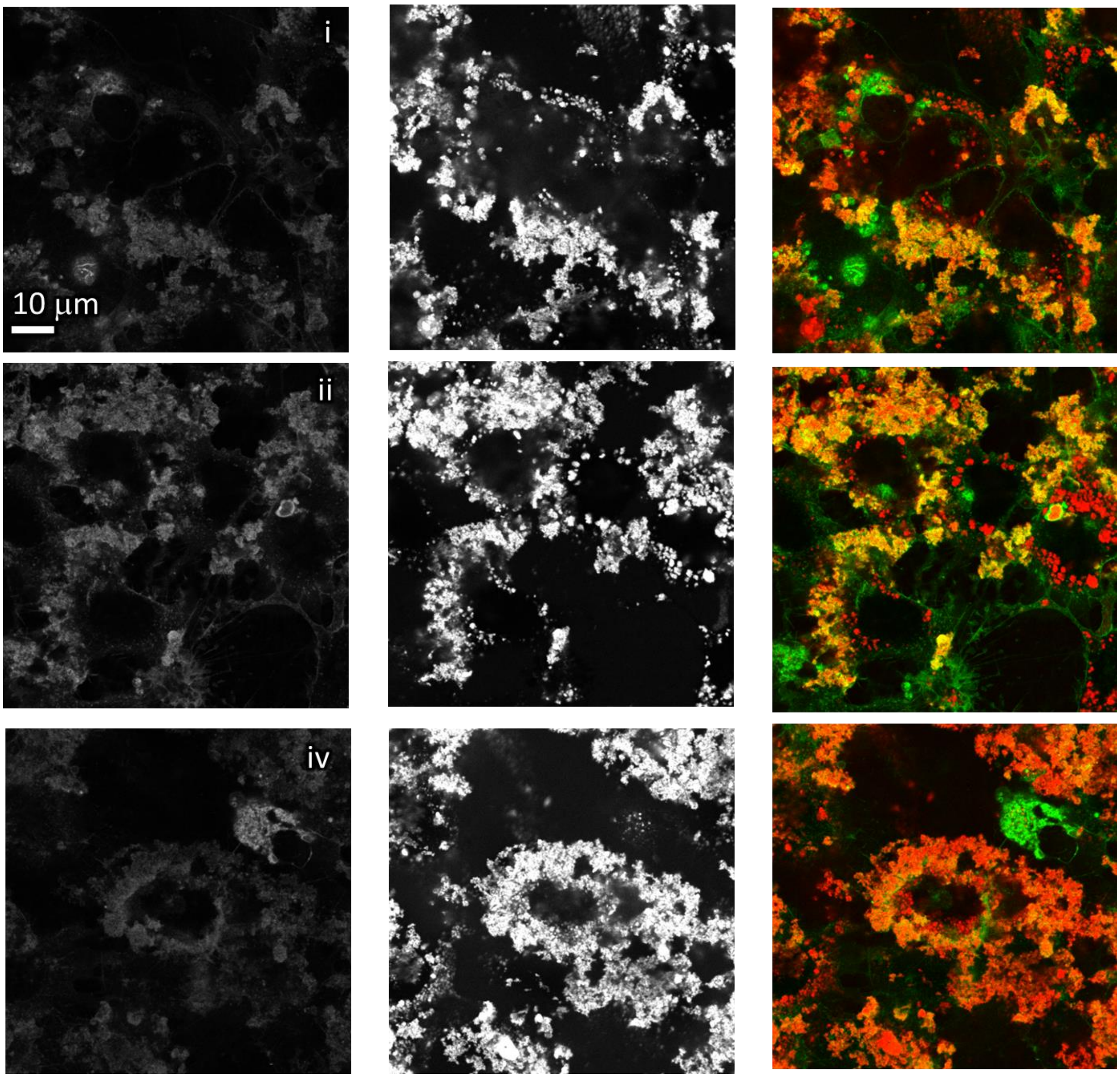

*Figure S68: Statistics at surface nanoparticles to surface cells dose 100:1 – confocal xy recordings.*

confocal xy – statistics over sample

LA-4 membrane (CellMaskOrange)

TiO2 (Alexa647)

overlay

| | |
|---|---|
| Cell line | LA-4 (membrane, CellMaskOrange) |
| NPs | / |
| exposure | / |
| imaging | xy confocal, @48h |

| | |
|---|---|
| pixelsize (x,y) | 100 nm |
| FOV (x,y) | 80 μm |
| pixelsize (z) | / |
| FOV (z) | / |
| imaging time | / |
| number of frames | / |

| | |
|---|---|
| 561nm | 30% |
| 640nm | 30% |
| STED | 20% |
| filter sets | 605 nm – 625 nm, 650 nm – 720 nm |
| dwell time | 20 μs |
| objective | wi60x (NA1.2) |

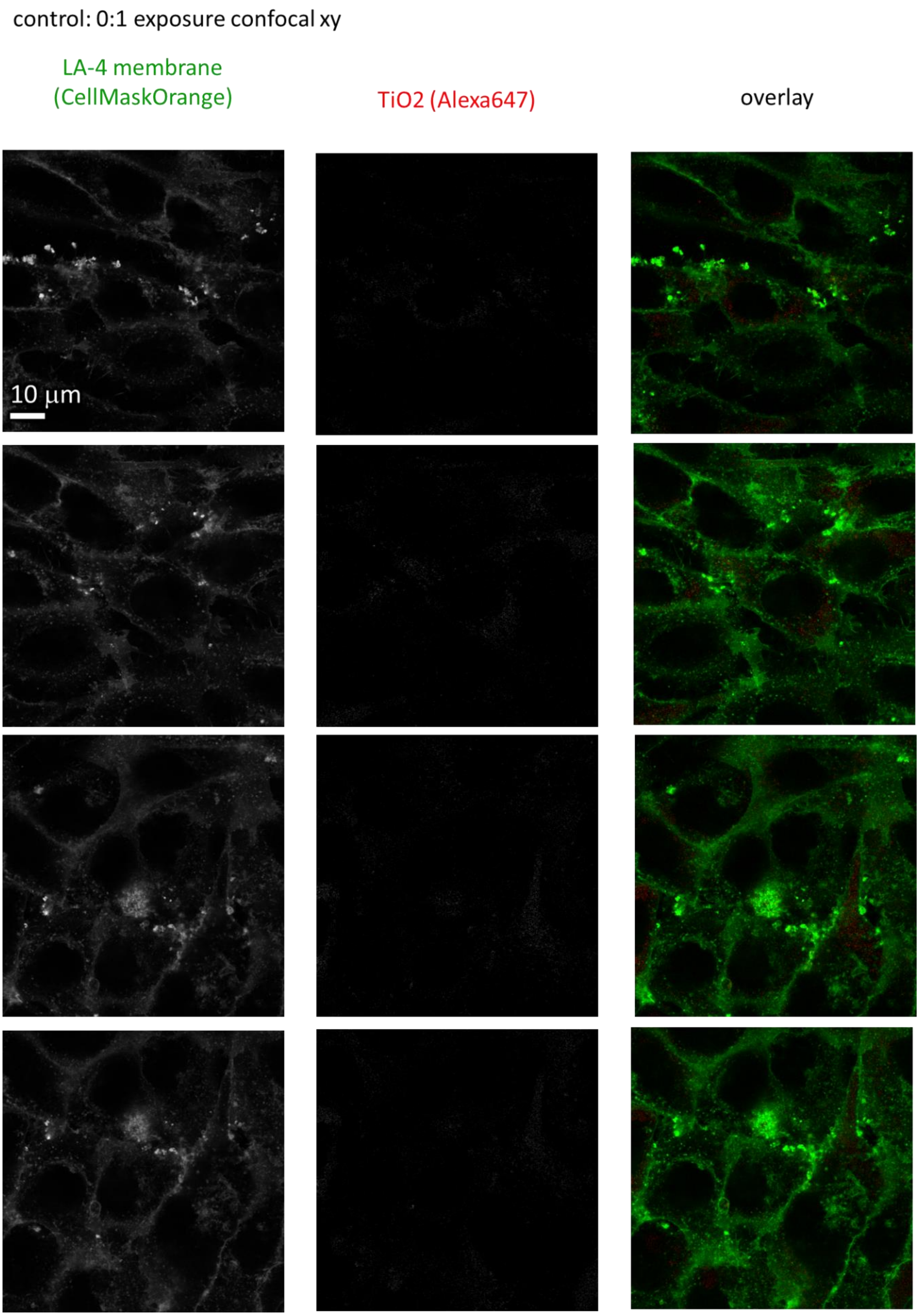

*Figure S69: Statistics for control with no exposure to nanomaterial – confocal xy recordings.*

control: 0:1 exposure confocal xy

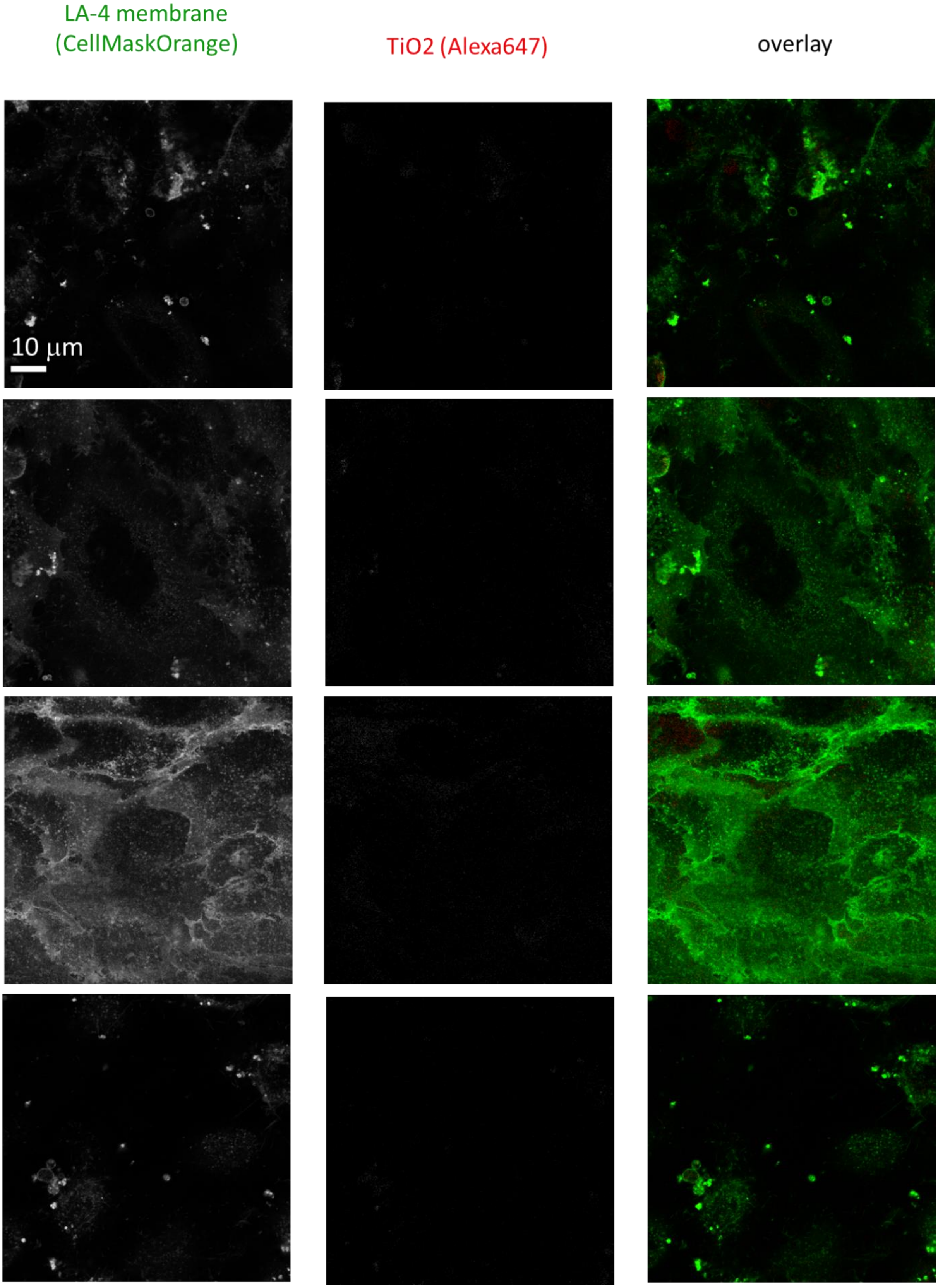

control: 0:1 exposure confocal xy

LA-4 membrane (CellMaskOrange) | TiO2 (Alexa647) | overlay

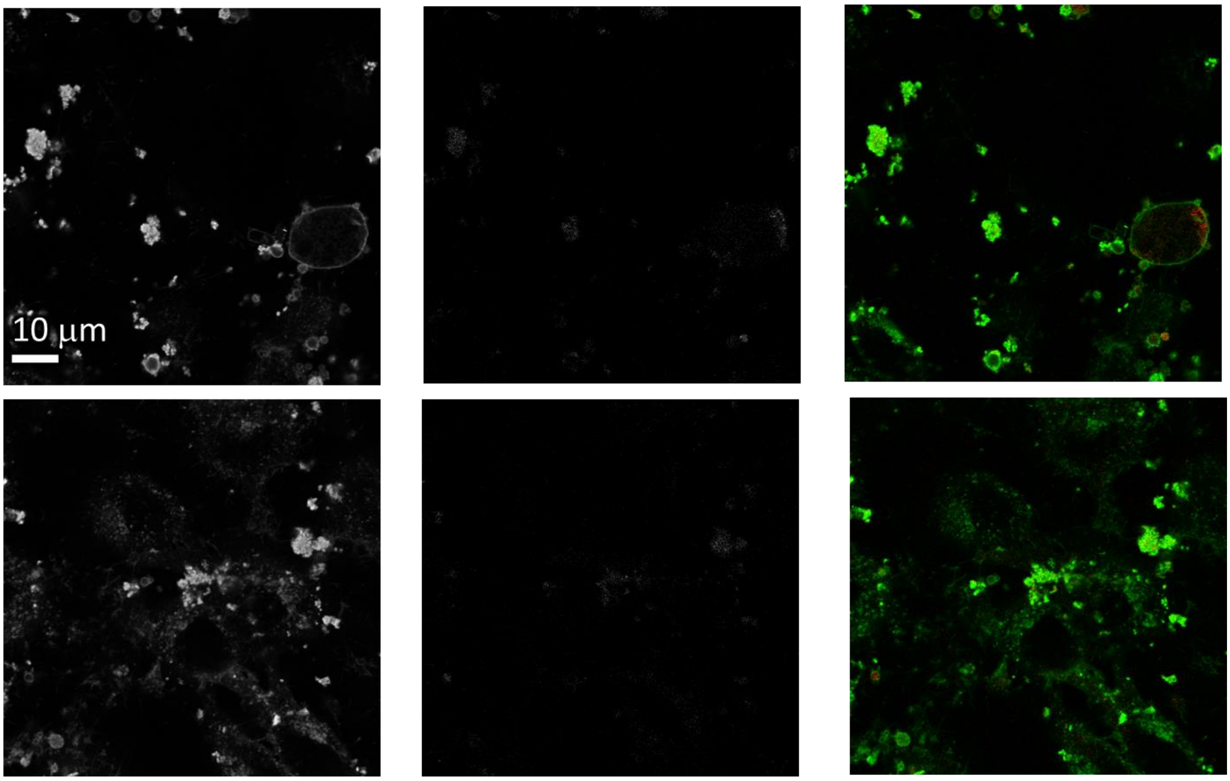

*Figure S70: Statistics for control with no exposure to nanomaterial – confocal xy recordings.*

control: 1:1 exposure confocal xy

| | |
|---|---|
| Cell line | LA-4 (membrane, CellMaskOrange) |
| NPs | $TiO_2$ (Alexa647) |
| exposure | 1:1, 48h incubation |
| imaging | xy confocal, @48h |

| | |
|---|---|
| pixelsize (x,y) | 100 nm |
| FOV (x,y) | 80 µm |
| pixelsize (z) | / |
| FOV (z) | / |
| imaging time | / |
| number of frames | / |

| | |
|---|---|
| 561nm | 30% |
| 640nm | 30% |
| STED | 20% |
| filter sets | 605 nm – 625 nm, 650 nm – 720 nm |
| dwell time | 20 µs |
| objective | wi60x (NA1.2) |

control: 1:1 exposure confocal xy

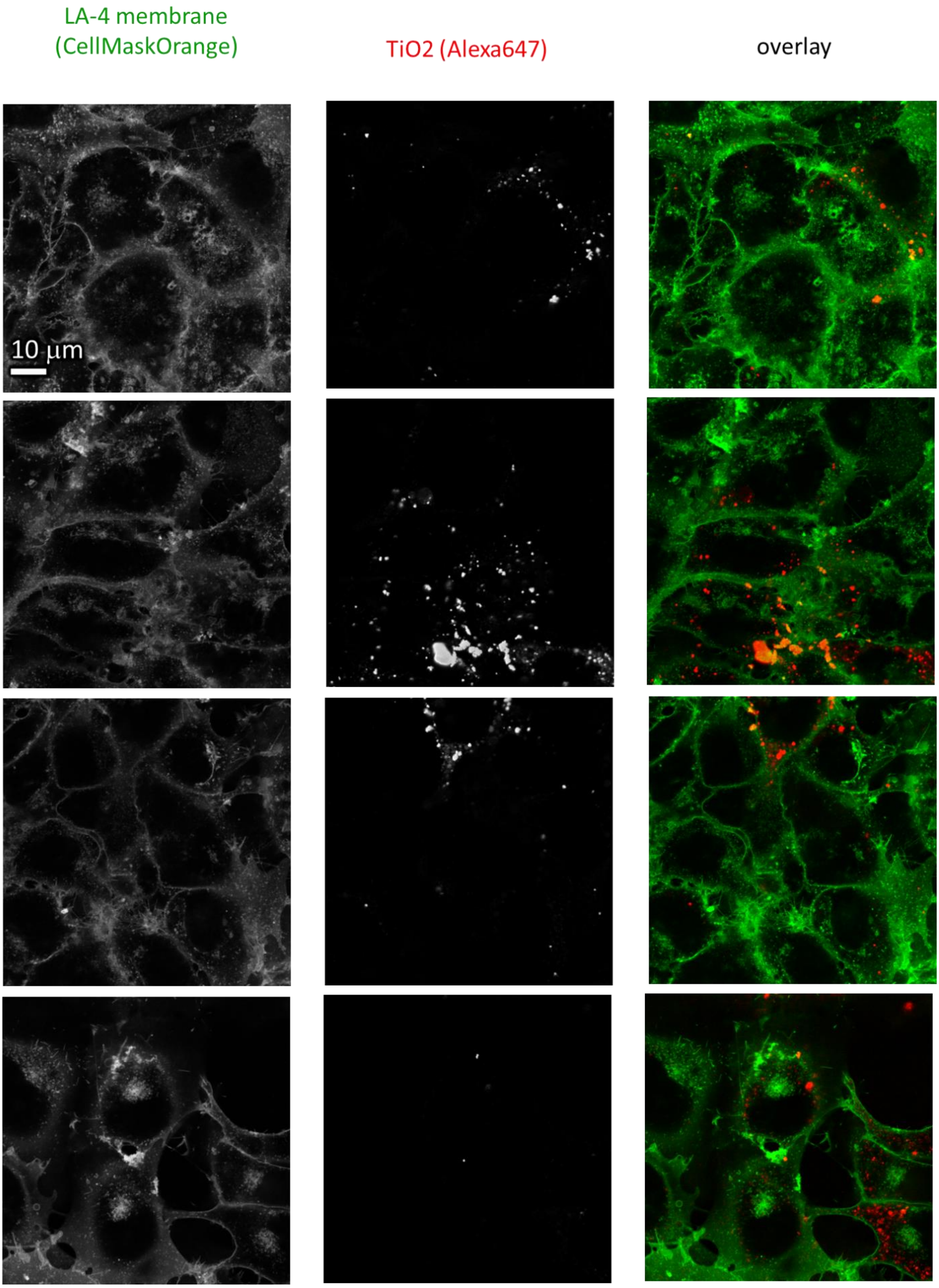

*Figure S71: Statistics at surface nanoparticles to surface cells dose 1:1 – confocal xy recordings.*

control: 1:1 exposure confocal xy

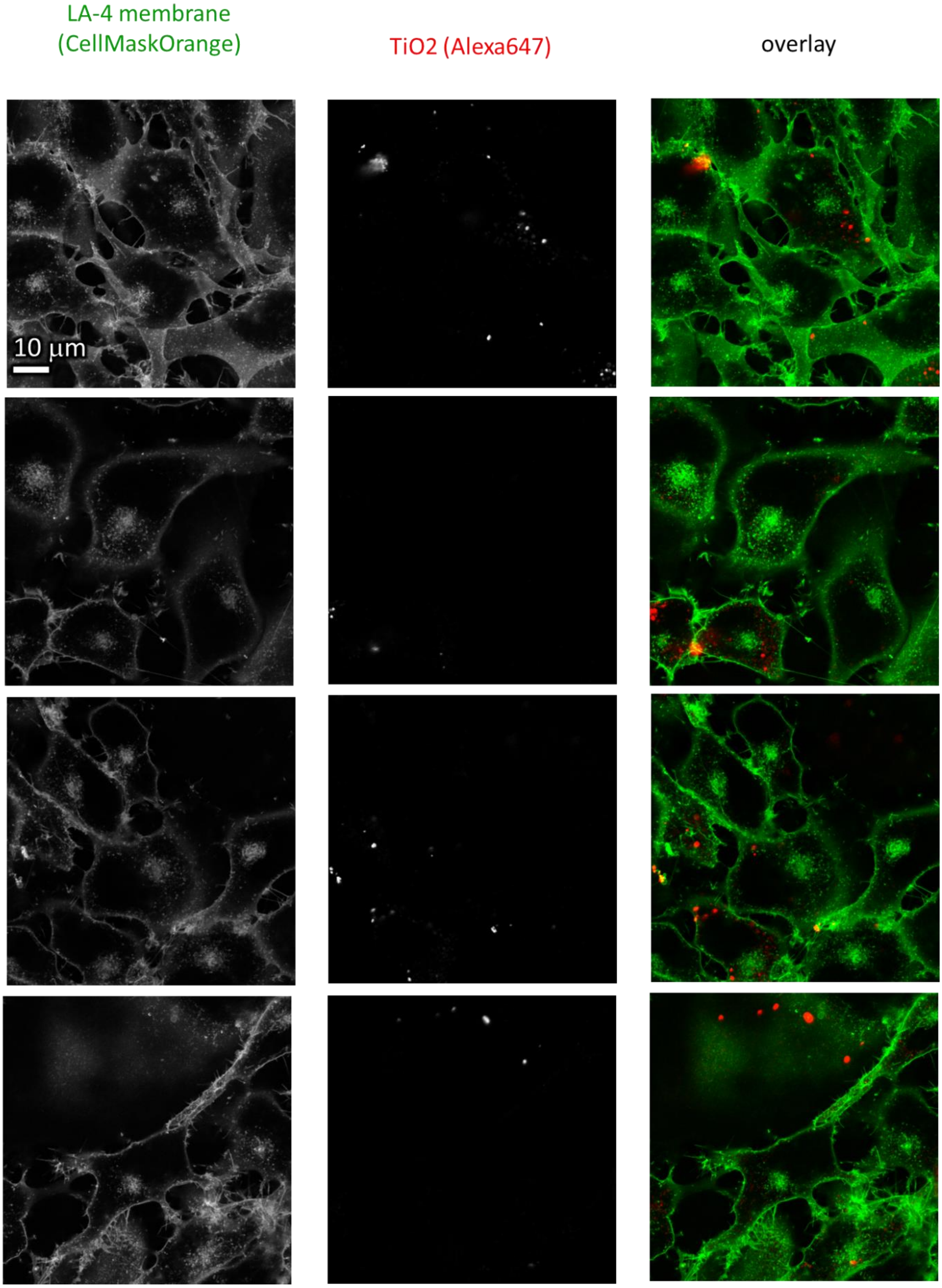

control: 1:1 exposure confocal xy

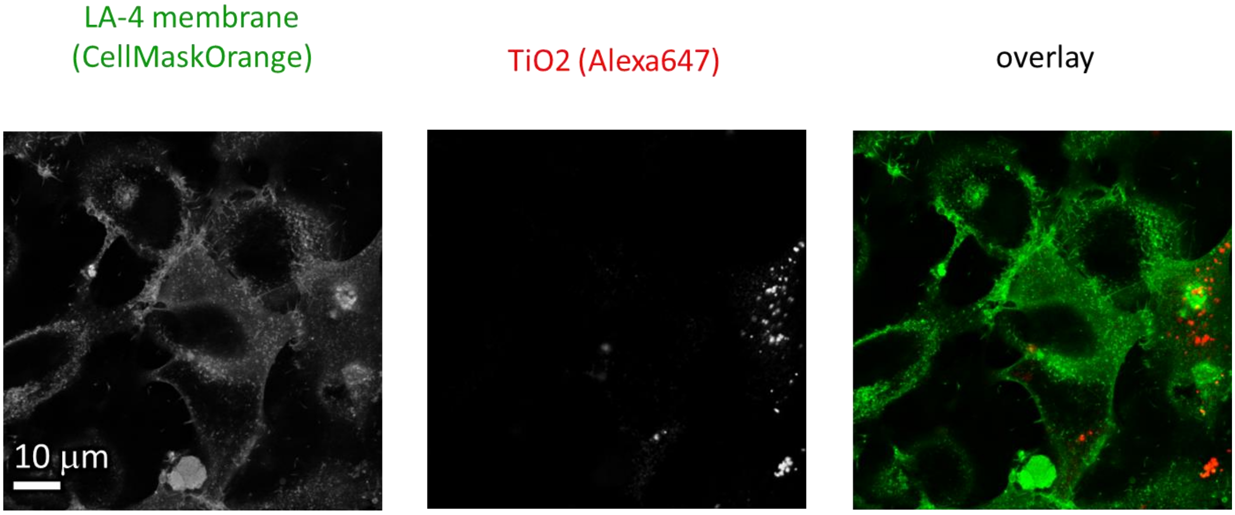

*Figure S72: Statistics at surface nanoparticles to surface cells dose 1:1 – confocal xy recordings.*

control: 10:1 exposure confocal xy

| | |
|---|---|
| Cell line | LA-4 (membrane, CellMaskOrange) |
| NPs | $TiO_2$ (Alexa647) |
| exposure | 10:1, 48h incubation |
| imaging | xy confocal, @48h |

| | |
|---|---|
| pixelsize (x,y) | 100 nm |
| FOV (x,y) | 80 μm |
| pixelsize (z) | / |
| FOV (z) | / |
| imaging time | / |
| number of frames | / |

| | |
|---|---|
| 561nm | 30% |
| 640nm | 30% |
| STED | 20% |
| filter sets | 605 nm – 625 nm, 650 nm – 720 nm |
| dwell time | 20 μs |
| objective | wi60x (NA1.2) |

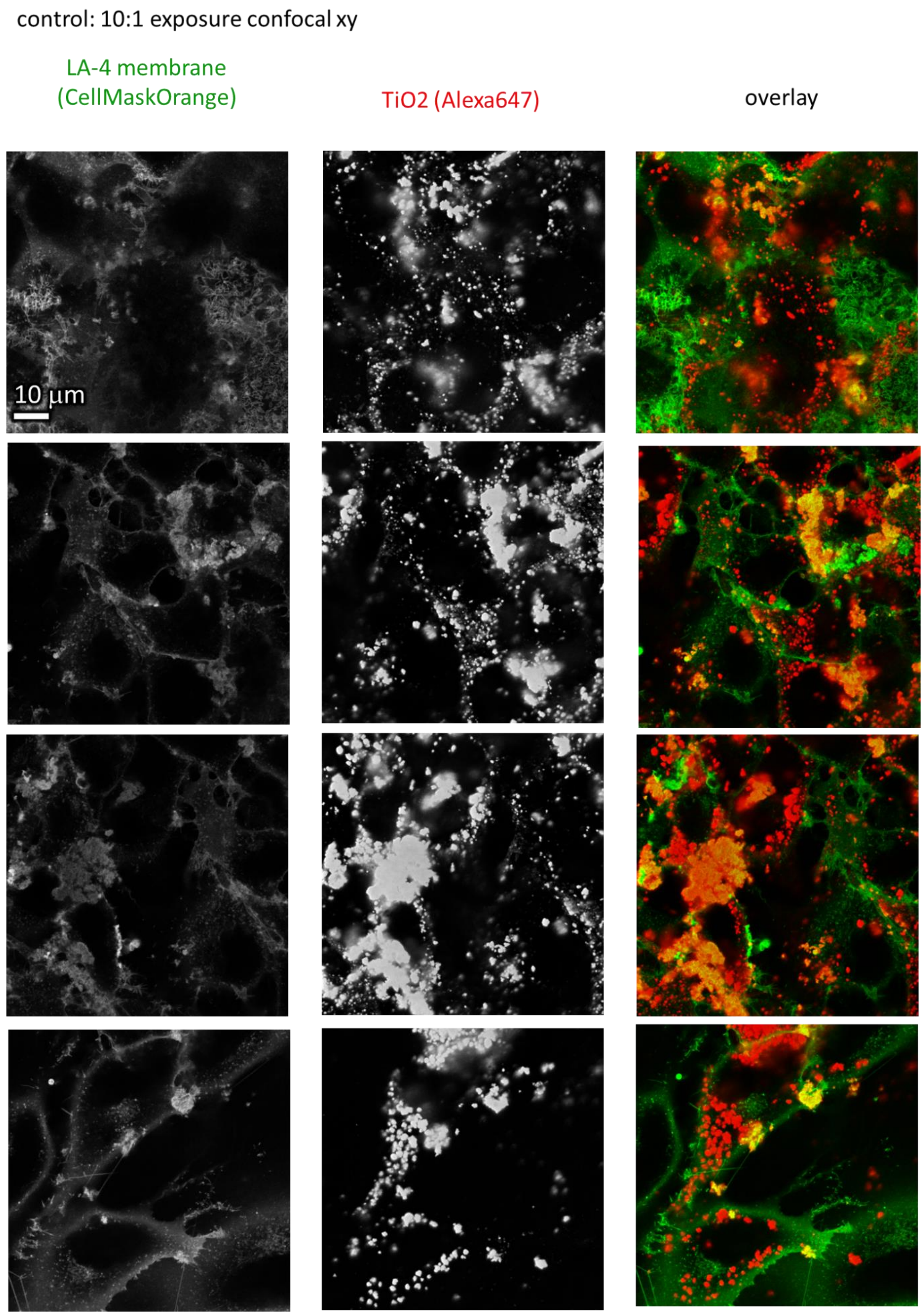

*Figure S73: Statistics at surface nanoparticles to surface cells dose 10:1 – confocal xy recordings.*

control: 10:1 exposure confocal xy

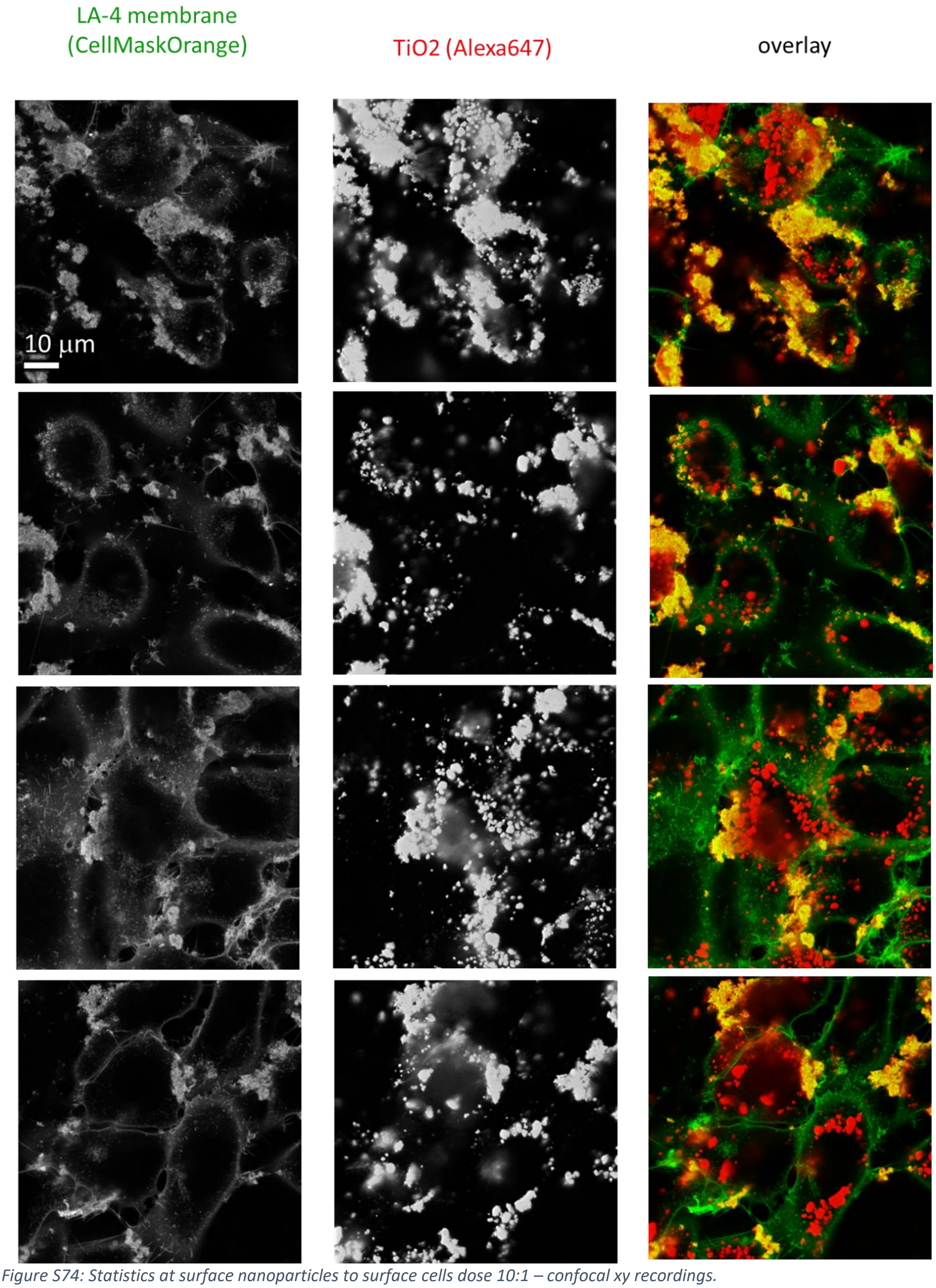

*Figure S74: Statistics at surface nanoparticles to surface cells dose 10:1 – confocal xy recordings.*

control: 100:1 exposure confocal xy

| | |
|---|---|
| Cell line | LA-4 (membrane, CellMaskOrange) |
| NPs | $TiO_2$ (Alexa647) |
| exposure | 100:1, 48h incubation |
| imaging | xy confocal, @48h |

| | |
|---|---|
| pixelsize (x,y) | 100 nm |
| FOV (x,y) | 80 µm |
| pixelsize (z) | / |
| FOV (z) | / |
| imaging time | / |
| number of frames | / |

| | |
|---|---|
| 561nm | 30% |
| 640nm | 30% |
| STED | 20% |
| filter sets | 605 nm – 625 nm, 650 nm – 720 nm |
| dwell time | 20 µs |
| objective | wi60x (NA1.2) |

control: 100:1 exposure confocal xy

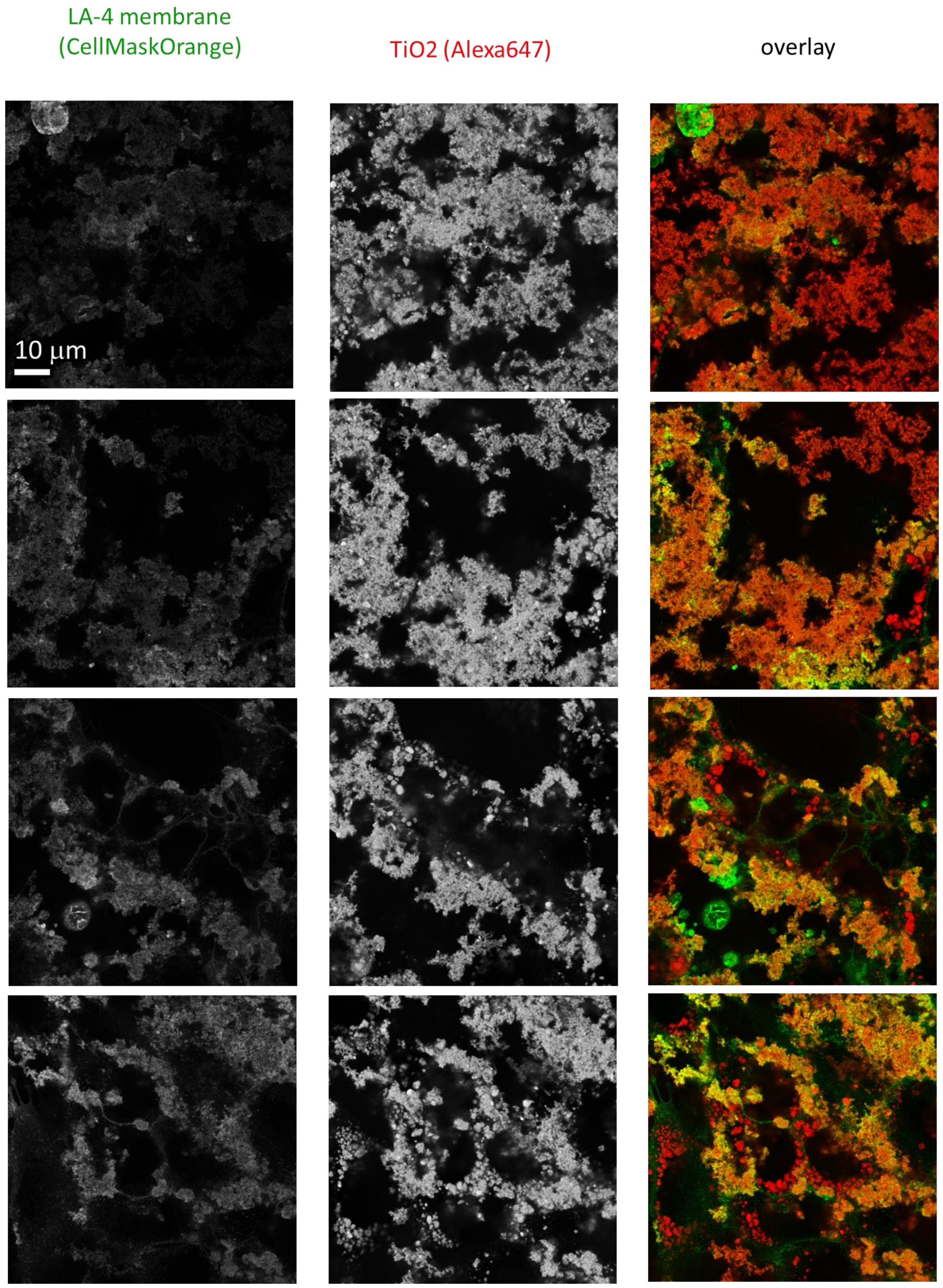

*Figure S75: Statistics at surface nanoparticles to surface cells dose 100:1 – confocal xy recordings.*

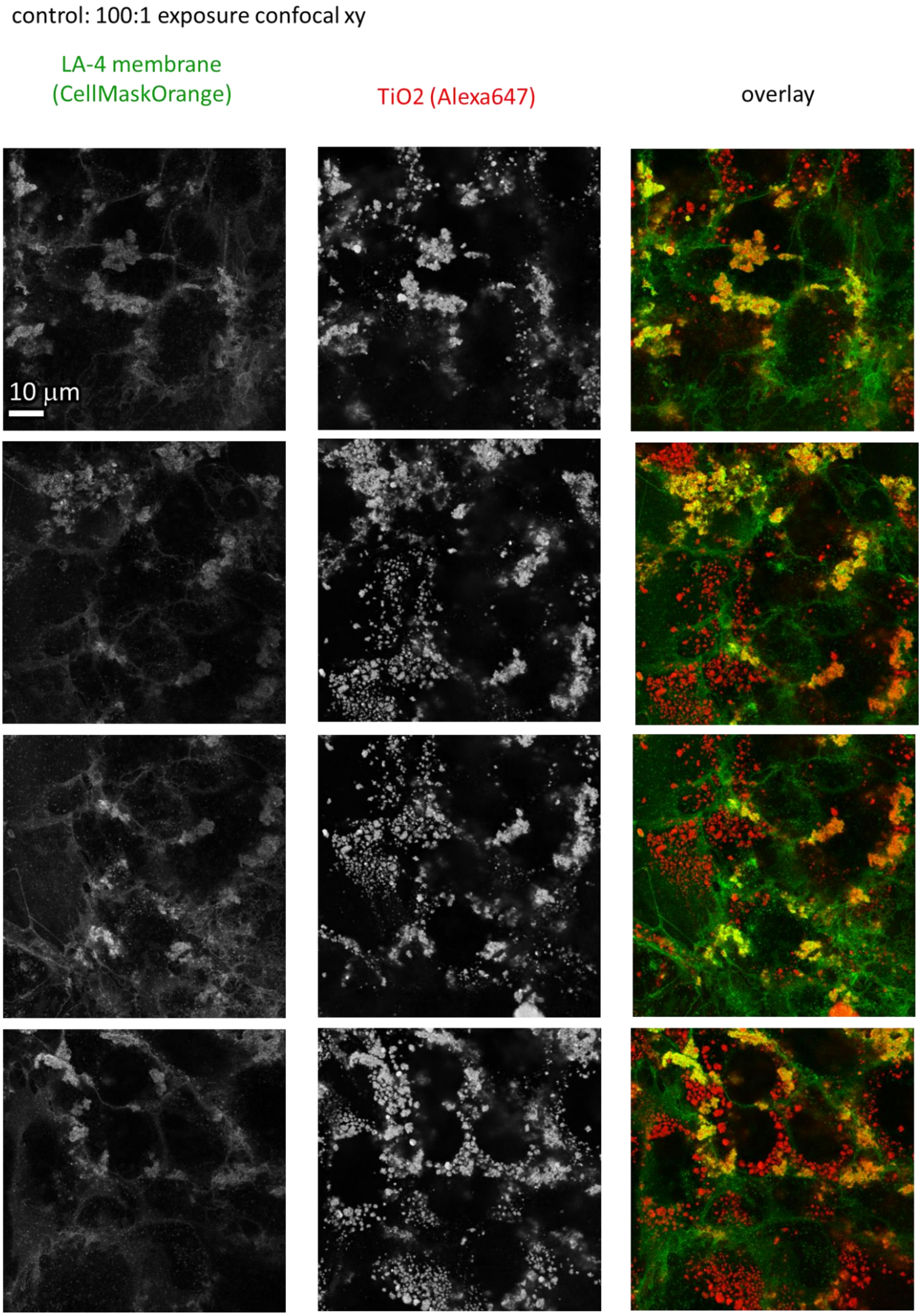

*Figure S76: Statistics at surface nanoparticles to surface cells dose 100:1 – confocal xy recordings.*

#### S1f – STED, HIM, SE SEM resolution comparison

For STED supplement see

S1d – Time evolution of cauliflower growth and S1e – Dose-dependent exposure of LA-4 to $TiO_2$ nanotubes.

##### Main message

Large magnification high-resolution HIM and SE SEM show that cauliflower-like bio-nano composites really are composed of small tubes and lipids as we claim based on STED high-resolution images.

Supporting raw and analysed data:

Figure S77-Figure S84

##### Materials and methods

In both HIM and SEM experiments samples were prepared in the same way. LA-4 cells were seeded in flasks with Si wafers (Pelcotec™ SFG12 Finder Grid Substrate, Ted Pella) placed on the bottom. After 48h Si wafers with cells on the surface were transferred to the 8-well holders. In this stage, Si wafers were approximately 70% confluent. Fresh medium was added together with 10:1(surface:surface) $TiO_2$ nanotubes. Samples where incubated for following 24 h before imaging or freezing. Freezing has been performed by freeze drying.

##### Controls and statistics – HIM

Several positions in several samples:

Figure S77-Figure S83

Acceleration voltage 30 keV for all experiments

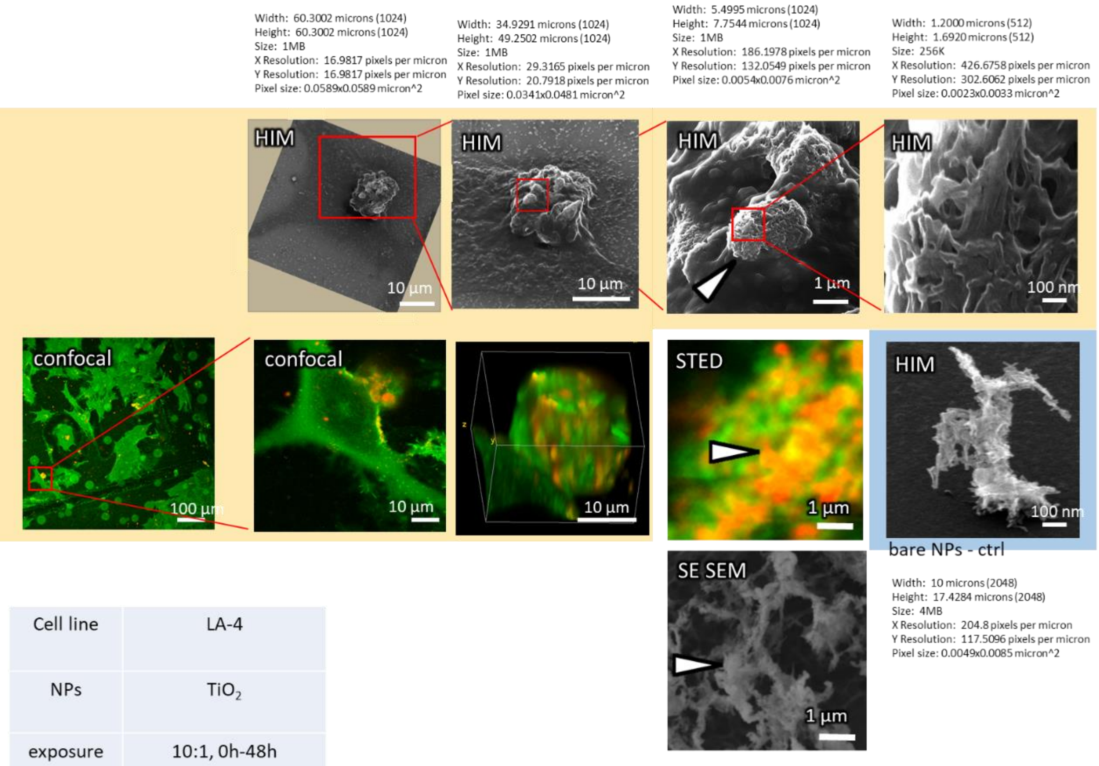

| | |
|---|---|
| Cell line | LA-4 |
| NPs | $TiO_2$ |
| exposure | 10:1, 0h-48h |
| imaging | HIM, @ 24h |

| | |
|---|---|
| Cell line | LA-4 (membrane, CellMaskOrange) |
| NPs | $TiO_2$-Alexa647 |
| exposure | / |
| imaging | xy confocal wide, @48h<br>xy confocal, @48h<br>xy STED, @48h<br>xy 3D confocal, @48h |

xy

| | |
|---|---|
| image | xy conf wide/ conf/ STED |
| pixelsize (x,y) | 100 nm/100 nm/30 nm |
| FOV (x,y) | 80 µm/40 µm/10 µm |

xyz – 3D

| | |
|---|---|
| pixelsize (x,y) | 100 nm |
| FOV (x,y) | 30 µm |
| pixelsize (z) | 500 nm |
| FOV (z) | 30 µm |

| | |
|---|---|
| 561nm | 30% |
| 640nm | 3% |
| STED | 20% |
| filter sets | 605 nm – 625 nm,<br>650 nm – 720 nm |
| dwell time | 80 µs |
| objective | wi60x (NA1.2) |

*Figure S77: Fluorescence and HIM correlation micorscopy with 3D scan as well as higher magnifications.*

#### Cauliflover formation (LA4)

Title: A3-cell-2.czi
Width:  85.0000 microns (2048)
Height:  85.0001 microns (2048)
Size:  4MB
X Resolution:  24.0941 pixels per micron
Y Resolution:  24.0941 pixels per micron
Pixel size: 0.0415x0.0415 micron^2

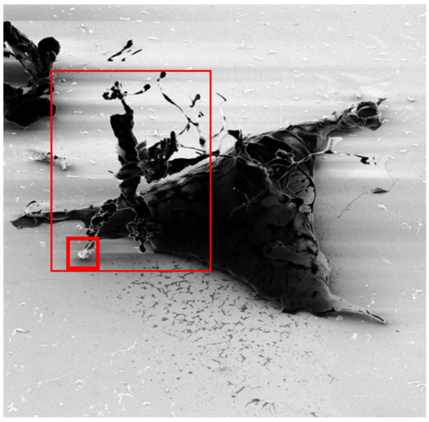

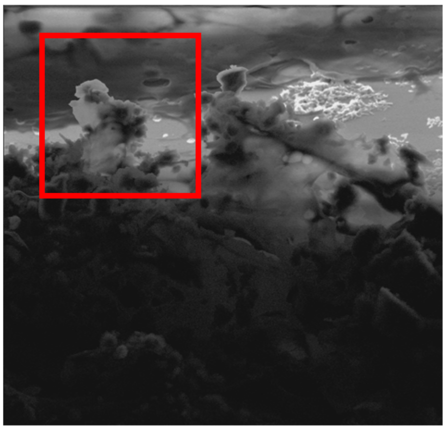

LA4 cell with destroyed part of a membrane which is ful of TiO2 nanoparticles

| Cell line | LA-4 |
|---|---|
| NPs | $TiO_2$ |
| exposure | 10:1, 0h-48h |
| imaging | HIM, @ 24h |

*Figure S78: Different magnifications of samples exposed to 10:1 (surface of nanomaterial: surface of cell) dose using HIM.*

### Cauliflover formation (LA4)

Title: C3-5.czi
Width:  75 microns (1024)
Height:  130.7127 microns (1024)
Size:  1MB
X Resolution:  13.6533 pixels per micron
Y Resolution:  7.8340 pixels per micron
Pixel size: 0.0732x0.1276 micron^2

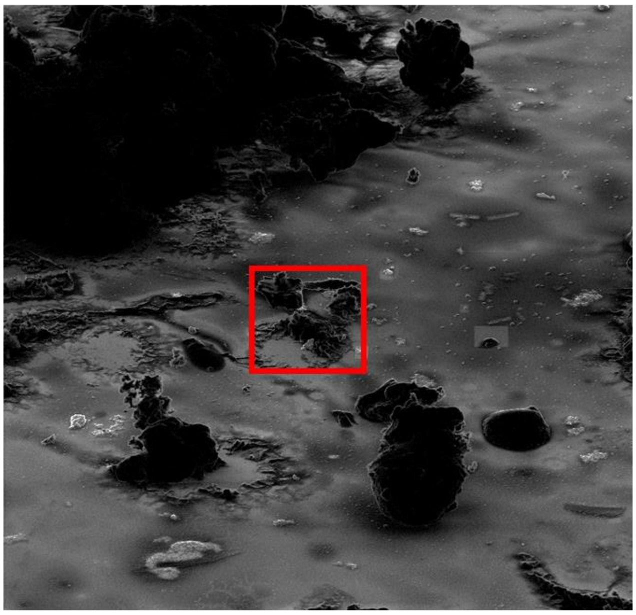

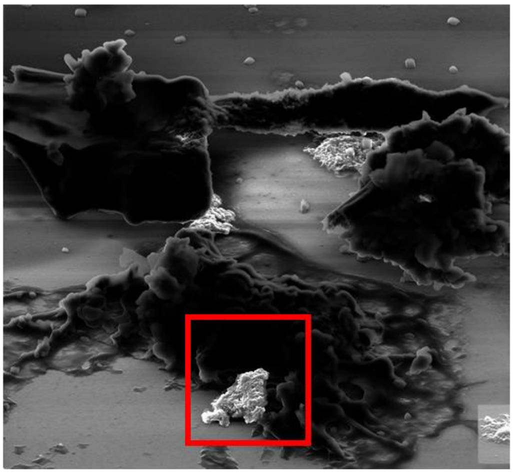

Title: C3-9.czi
Width:  2.5 microns (2048)
Height:  4.3571 microns (2048)
Size:  4MB
X Resolution:  819.2 pixels per micron
Y Resolution:  470.0384 pixels per micron
Pixel size: 0.0012x0.0021 micron^2

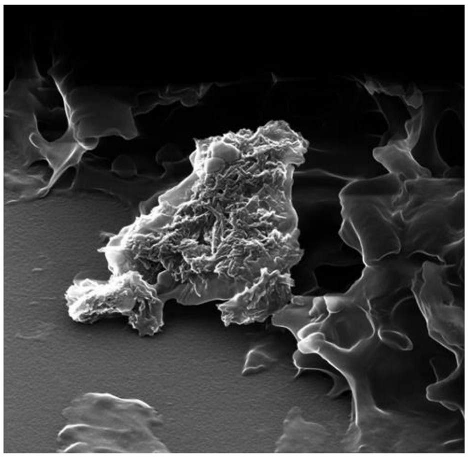

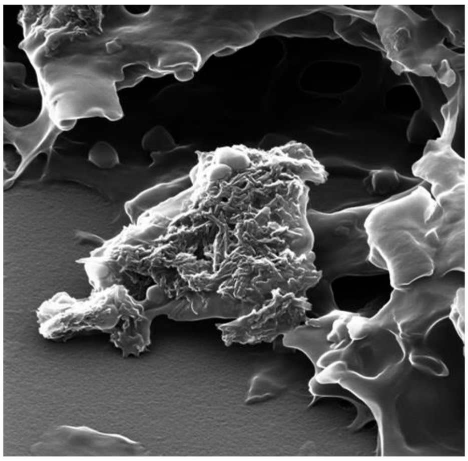

| Cell line | LA-4 |
|---|---|
| NPs | $TiO_2$ |
| exposure | 10:1, 0h-48h |
| imaging | HIM, @ 24h |

*Figure S79: Different magnifications of samples exposed to 10:1 (surface of nanomaterial: surface of cell) dose using HIM.*

### Destroyed MHS, debris of membranes with NPs

Title: A8-4.czi
Width: 350 microns (2048)
Height: 609.9927 microns (2048)
Size: 4MB
X Resolution: 5.8514 pixels per micron
Y Resolution: 3.3574 pixels per micron
Pixel size: 0.1709x0.2978 micron^2

Title: A8-6.czi
Width: 70 microns (2048)
Height: 121.9985 microns (2048)
Size: 4MB
X Resolution: 29.2571 pixels per micron
Y Resolution: 16.7871 pixels per micron
Pixel size: 0.0342x0.0596 micron^2

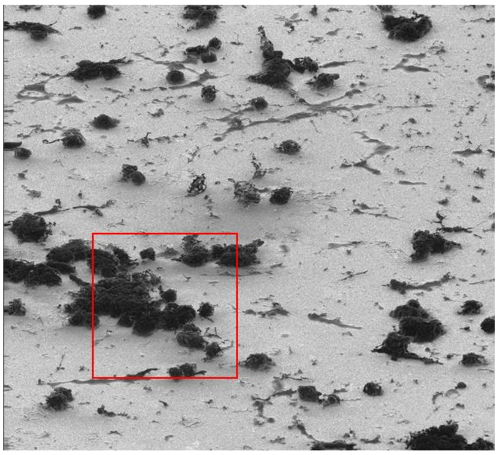

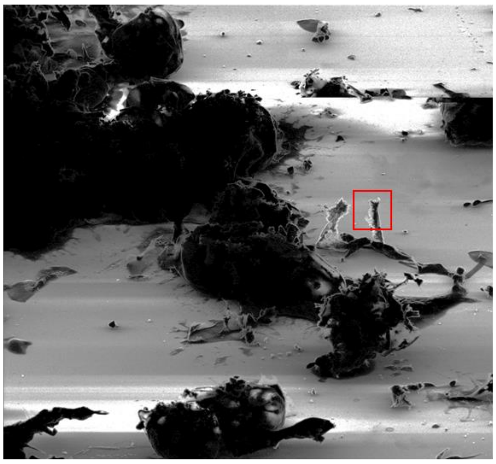

Title: A8-13.czi
Width: 1.9986 microns (1024)
Height: 3.4833 microns (1024)
Size: 1MB
X Resolution: 512.3530 pixels per micron
Y Resolution: 293.9765 pixels per micron
Pixel size: 0.0020x0.0034 micron^2

Title: A8-11.czi
Width: 4.5 microns (1024)
Height: 7.8428 microns (1024)
Size: 1MB
X Resolution: 227.5556 pixels per micron
Y Resolution: 130.5662 pixels per micron
Pixel size: 0.0044x0.0077 micron^2

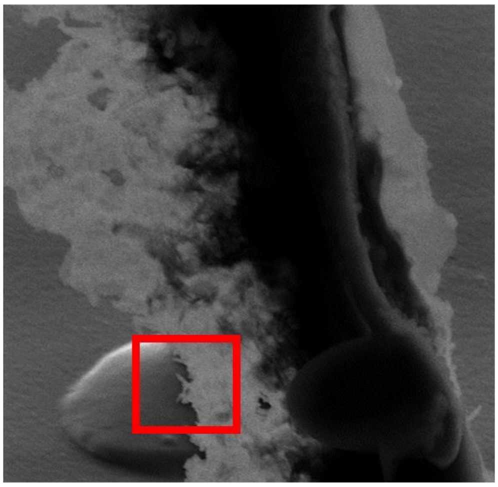

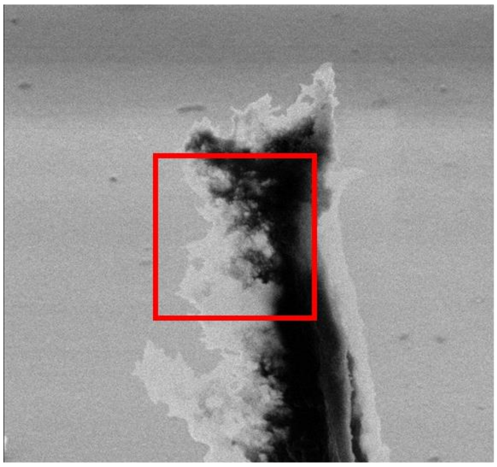

Single TiO2 nanotubes nicely observed at the surface
Whole image is just 2µm * 2µm

| | |
|---|---|
| Cell line | MHS |
| NPs | $TiO_2$ |
| exposure | 10:1, 0h-48h |
| imaging | HIM, @ 24h<br>SE SEM, @ 24h |

*Figure S80: Different magnifications of samples exposed to 10:1 (surface of nanomaterial: surface of cell) dose using HIM.*

### MHS, short term exposure with NPs

Title: C3-7.czi
Width: 12 microns (2048)
Height: 20.9140 microns (2048)
Size: 4MB
X Resolution: 170.6667 pixels per micron
Y Resolution: 97.9247 pixels per micron
Pixel size: 0.0059x0.0102 micron^2

Title: C3-11.czi
Width: 15.0000 microns (2048)
Height: 26.1425 microns (2048)
Size: 4MB
X Resolution: 136.5333 pixels per micron
Y Resolution: 78.3397 pixels per micron
Pixel size: 0.0073x0.0128 micron^2

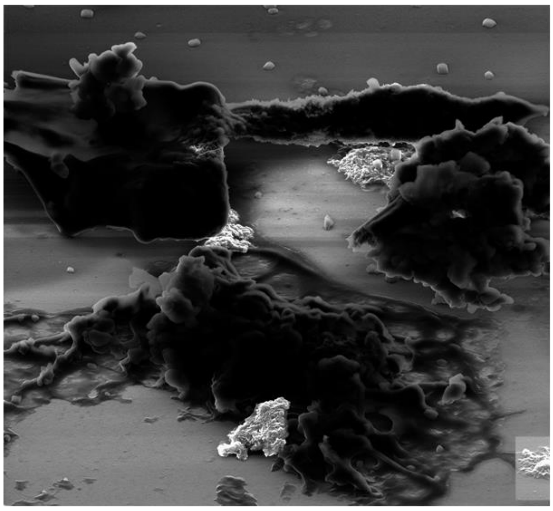

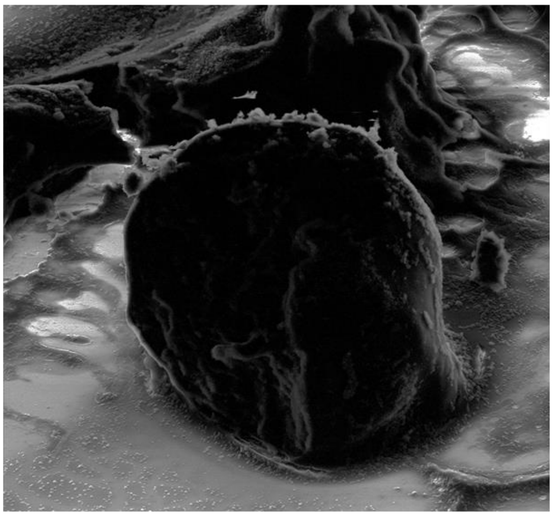

| | |
|---|---|
| Cell line | MHS |
| NPs | $TiO_2$ |
| exposure | 10:1, 0h-48h |
| imaging | HIM, @ 1h |

*Figure S81: Different magnifications of samples exposed to 10:1 (surface of nanomaterial: surface of cell) dose using HIM.*

### MHS, long term exposure with NPs

Title: B3-cell-20-tilted.czi
Width: 125 microns (2048)
Height: 193.2184 microns (2048)
Size: 4MB
X Resolution: 16.384 pixels per micron
Y Resolution: 10.5994 pixels per micron
Pixel size: 0.0610x0.0943 micron^2

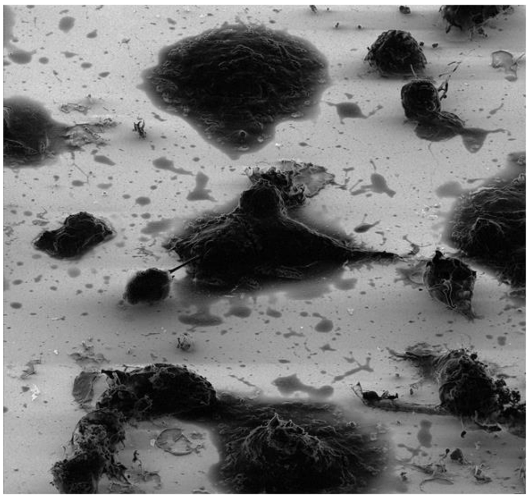

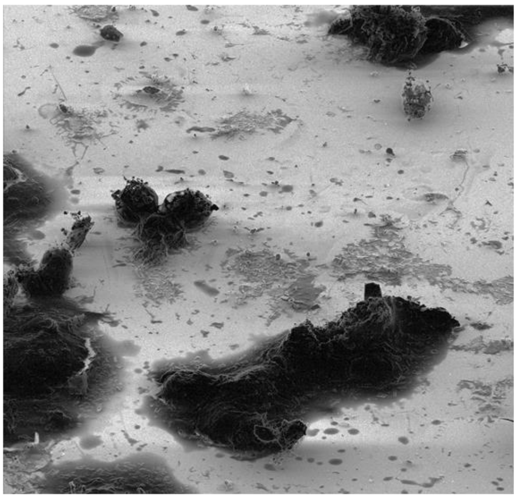

Title: B3-cell-24-tilted.czi
Width: 40 microns (2048)
Height: 61.8299 microns (2048)
Size: 4MB
X Resolution: 51.2 pixels per micron
Y Resolution: 33.1231 pixels per micron
Pixel size: 0.0195x0.0302 micron^2

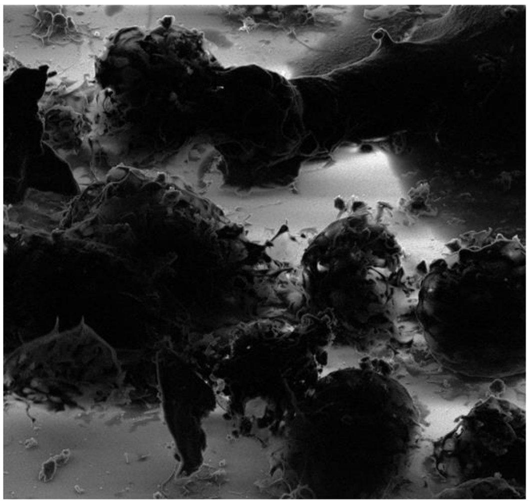

| | |
|---|---|
| Cell line | LA-4 |
| NPs | $TiO_2$ |
| exposure | 10:1, 0h-48h |
| imaging | HIM, @ 24h |

*Figure S82: Different magnifications of samples exposed to 10:1 (surface of nanomaterial: surface of cell) dose using HIM.*

### MHS, long term exposure with NPs

Title: A7-cell-14-tilted.czi
Width: 425 microns (2048)
Height: 656.9427 microns (2048)
Size: 4MB
X Resolution: 4.8188 pixels per micron
Y Resolution: 3.1175 pixels per micron
Pixel size: 0.2075x0.3208 micron^2

Title: A7-cell-15-tilted.czi
Width: 90.0000 microns (2048)
Height: 139.1173 microns (2048)
Size: 4MB
X Resolution: 22.7556 pixels per micron
Y Resolution: 14.7214 pixels per micron
Pixel size: 0.0439x0.0679 micron^2

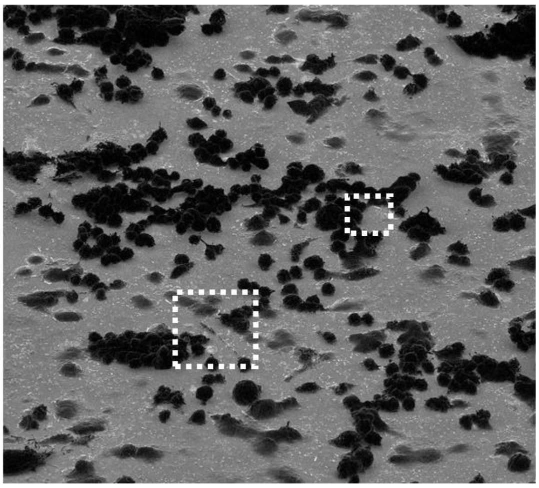

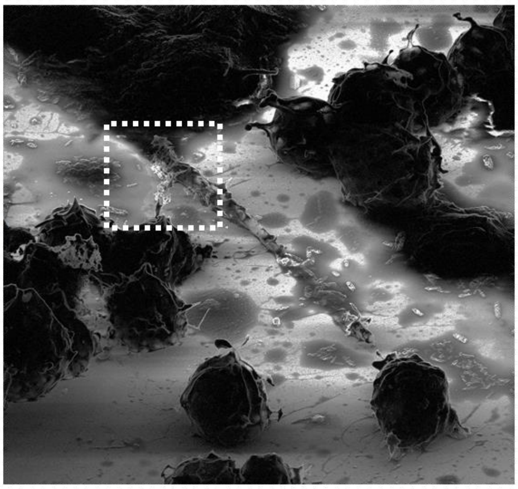

Title: A7-cell-17-tilted.czi
Width: 15.0000 microns (2048)
Height: 23.1862 microns (2048)
Size: 4MB
X Resolution: 136.5333 pixels per micron
Y Resolution: 88.3284 pixels per micron
Pixel size: 0.0073x0.0113 micron^2

Title: A7-cell-19-tilted.czi
Width: 1.4999 microns (1024)
Height: 2.3185 microns (1024)
Size: 1MB
X Resolution: 682.6956 pixels per micron
Y Resolution: 441.6605 pixels per micron
Pixel size: 0.0015x0.0023 micron^2

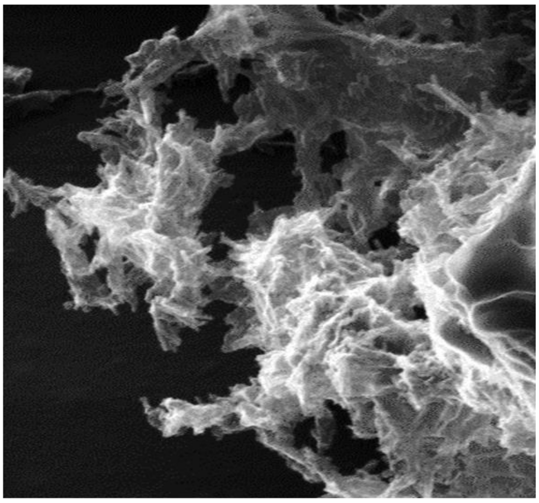

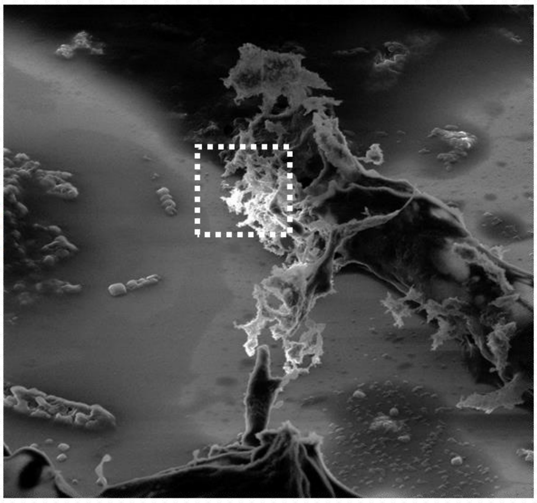

| | |
|---|---|
| Cell line | LA-4 |
| NPs | $TiO_2$ |
| exposure | 10:1, 0h-48h |
| imaging | HIM, @ 24h |

*Figure S83: Different magnifications of samples exposed to 10:1 (surface of nanomaterial: surface of cell) dose using HIM.*

#### Controls and statistics – SE SEM

Larger field of view on the sample with several zoom-ins:

Figure S84

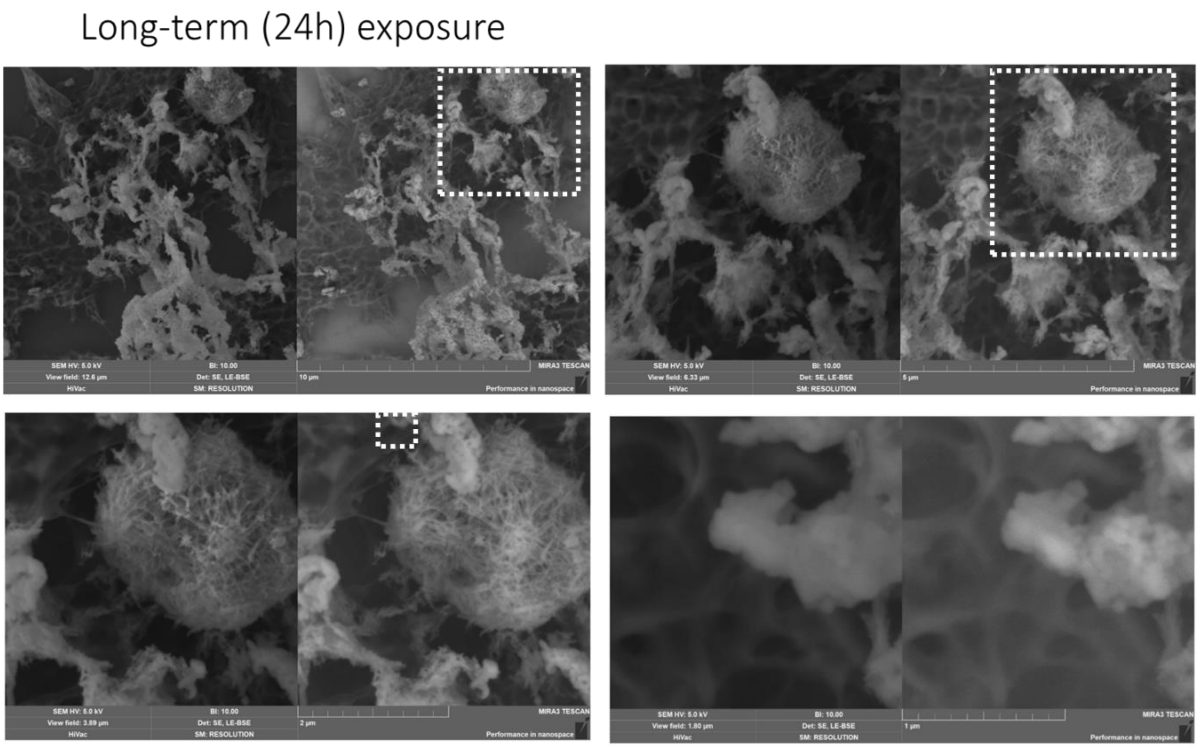

| | |
|---|---|
| Cell line | LA-4 |
| NPs | $TiO_2$ |
| exposure | 10:1, 0h-24h |
| imaging | SE SEM, @ 24h |

*Figure S84: Different magnifications of sample exposed to 10:1 (surface of nanomaterial: surface of cell) dose using SE SEM.*

### S2 – The role of lipids

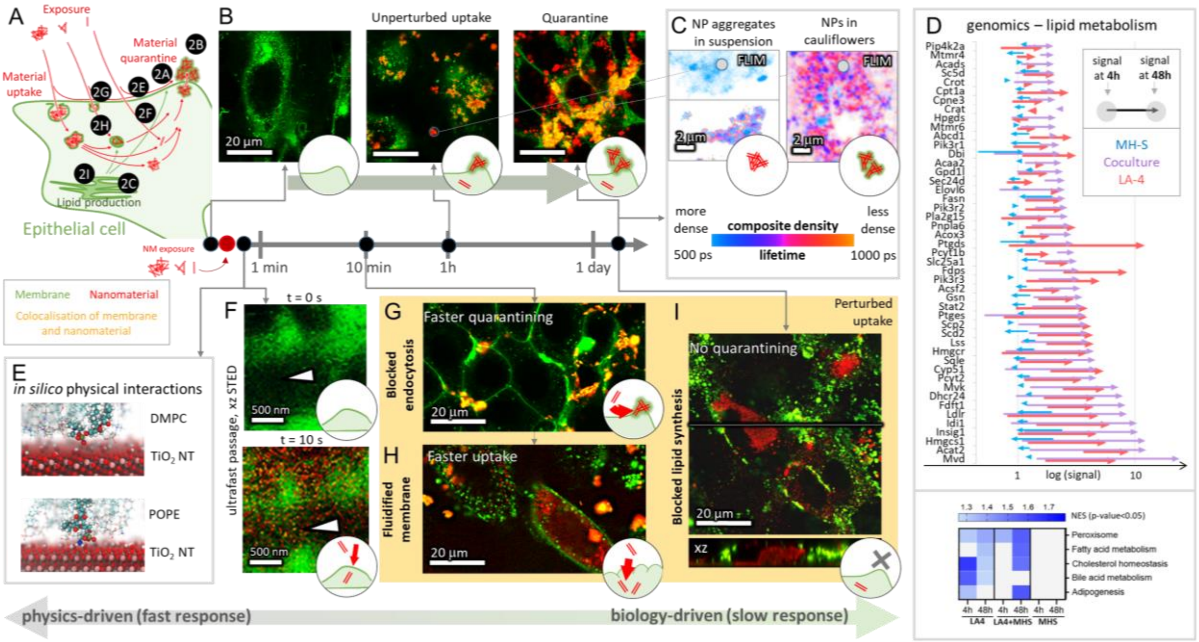

#### S2b – Time evolution of cauliflower growth

See supplement section

S1d – Time evolution of cauliflower growth.

#### S2c – FLIM of cauliflowers

##### Main message

The fluorescence lifetime of labelled $TiO_2$ nanotubes in cauliflowers is longer than their lifetime in dense aggregates and shorter than the lifetime of free-floating labelled $TiO_2$ nanotubes, indicating that the density of nanomaterial in cauliflowers is somewhere in between dense aggregates and free-floating nanomaterial. Also, the bleed-through from the membrane labels into the FLIM channel shown in this paper is negligible.

Supporting raw and analysed data:

Figure S85-Figure S94

##### Materials and methods

- experiment 20190419_e06_s02_t02 (upper left image - giant aggregate FLIM):
    - 11 µl labelled $TiO_2$-40-Alexa647 in 100x dcb was added to 200 µl LCIS in an Ibidi #1.5H µ-Slide 8-well to achieve an effective surface dose 10:1 (in regard to the bottom surface of the well)
    - after 4 hours, the sample was observed on the heated stage at 32C

| | |
|---|---|
| Cell line | |
| NPs | TiO2 (Alexa647) |
| exposure | 10:1, 4h |
| imaging | xy confocal FLIM |

| | |
|---|---|
| pixelsize (x,y) | 100 nm |
| FOV (x,y) | 51,2 µm |
| TCSPC pixelsize | 122 ps |
| TCSPC FOV | 19.5 ns |

| | |
|---|---|
| 561nm | |
| 640nm | 10% |
| STED | |
| diffraction grating | 523 – 722 nm |
| dwell time | 50 µs |
| objective | 60x wi (NA1.2) |

- experiment 20190628_fc04_s01_t01 (lower left image - small aggregate FLIM)
    - 10 µl labelled $TiO_2$-17-Alexa647 in 100x dcb was added to 200 µl LCIS in an Ibidi #1.5H µ-Slide 8-well to achieve an effective surface dose 10:1 (in regard to the bottom surface of the well)
    - after 1 hour, the sample was observed on the heated stage at 32C

| | |
|---|---|
| Cell line | |
| NPs | TiO2 (Alexa647) |
| exposure | 10:1, 1h |
| imaging | xy confocal FLIM |

| | |
|---|---|
| pixelsize (x,y) | 100 nm |
| FOV (x,y) | 51,2 µm |
| TCSPC pixelsize | 122 ps |
| TCSPC FOV | 19.5 ns |

| | |
|---|---|
| 561nm | |
| 640nm | 30% |
| STED | |
| diffraction grating | 523 – 722 nm |
| dwell time | 200 µs |
| objective | 60x wi (NA1.2) |

- experiment 20190621_e01.100_s02_t04 (right image - cauliflower FLIM):
    - LA-4 cells were seeded @50% confluence in an Ibidi #1.5H µ-Dish with a 4-Well Culture-Insert (Ibidi)
    - after 36 hours, 35 µl freshly filtered 1 mg/ml $TiO_2$-17-Alexa647 in 100x dcb was mixed into 130 µl fresh F12K medium and added to cells in one of the inserts to achieve 100:1 surface dose

- 29 hours later, the cells in the well were first incubated with 1.6 µg/ml CellMaskOrange in LCIS for 5 minutes, then incubated 15 minutes with freshly diluted 5 µM SAG-38 in LCIS at room temperature, and finally flushed with LCIS and observed in 150 µL LCIS in the home-made incubator at 37C

| | |
|---|---|
| Cell line | LA-4 (membrane, CellMaskOrange and lipid bodies, SAG-38) |
| NPs | TiO2 (Alexa647) |
| exposure | 100:1, 29h |
| imaging | xy confocal FLIM |

| | |
|---|---|
| pixelsize (x,y) | 100 nm |
| FOV (x,y) | 51,2 µm |
| TCSPC pixelsize | 122 ps |
| TCSPC FOV | 19.5 ns |

| | |
|---|---|
| 561nm | |
| 640nm | 10% |
| STED | |
| diffraction grating | 523 – 722 nm |
| dwell time | 100 µs |
| objective | 60x wi (NA1.2) |

- analysis:
  - the fluorescence lifetime data was sent from Imspector 16.2 (Abberior Instruments) to SPCImage 7.3 (Becker & Hickl), where the Decay matrix was calculated from the brightest pixel in the image (monoexponential fitting), binning was set to 1 and threshold to 5. In the supplement, wherever the lifetime of non-aggregated nanomaterial was determined, the binning was set to 10 (also noted in the image itself)
  - the rainbow LUT was rescaled to range from 500 ps to 1000 ps and intensity and contrast of the lifetime-coded image were adjusted for easier comparison between experiments
  - the lifetime-coded image and color legend were exported as a TIF and imported into IrfanView, where the negative was obtained for more clear presentation in the paper (green color was avoided in the FLIM image to prevent confusing it with the green-coded membranes in ordinary fluorescence images)

#### Experiment names

- Main experiment name:
  - 20190419_e06_s02_t02_TiO2-40-Alexa647 in LCIS 10.1_aggregate.img
  - 20190628_fc04_s01_t01_TiO2-17-Alexa647 in LCIS_FLIM.img
  - 20190621_e01.100_s02_t04_LA-4CellMask TiO2-17-Alexa647 100.1_cauliflower.img

- Supplement Experiment names:
  - 20190419/e06_s01_t01_TiO2Alexa647 before autoclaving in LCIS 10.1_bottom_FLIM_noSpectralFlim.msr
  - 20190419/e12_s01_t01_Alexa647 before autoclaving in LCIS_middle_noSpectralFlim.msr
  - 20190607/e03_s02_t02_SAG-38_conf.msr
  - 20190628/fc04_s01_t01_TiO2-17-Alexa647 in LCIS_FLIM.msr
  - 20190628/fc01_s01_t03_LA-4 free Alexa647 1h_FLIM.msr
  - 20190808/e01_s01_LA-4 CMO C75 c0.msr

#### Controls and statistics

Fluorescence lifetime mapping of $TiO_2$ nanotubes, labelled with Alexa Fluor 647 – in aggregates and in suspension

Figure S85-Figure S90

Fluorescence lifetime of $TiO_2$ nanotubes, labelled with Alexa Fluor 647, in cauliflowers

Figure S91

Fluorescence lifetime of free Alexa Fluor 647 in suspension

Figure S92

Fluorescence lifetime of free Alexa Fluor 647 in cells

Figure S93

Crosstalk reference for CellMaskOrange and SAG-38

Figure S94

experiment 20190419_e06_s01_t01 (smaller aggregates of TiO2-Alexa647 in LCIS):

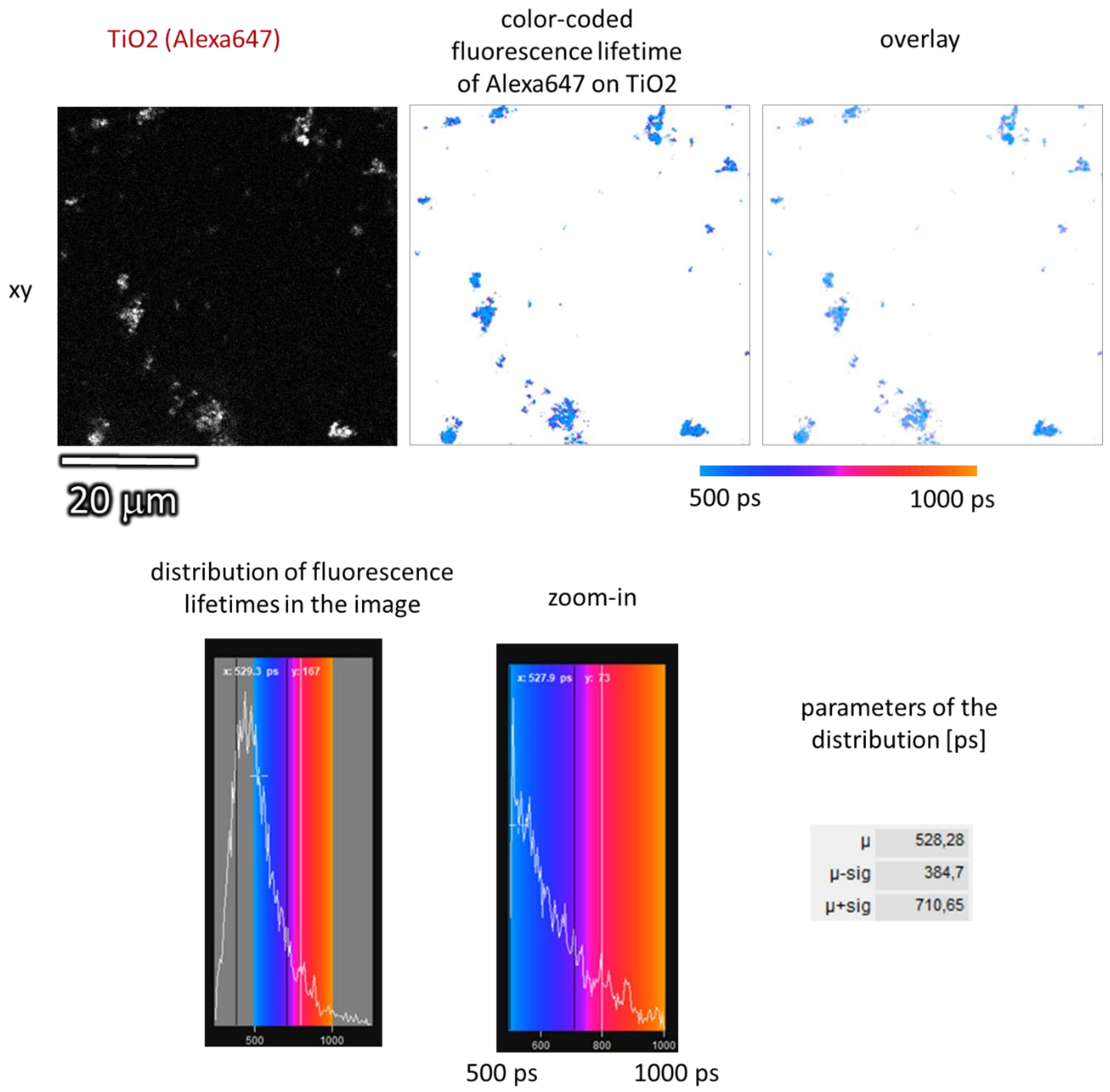

| | |
|---|---|
| Cell line | |
| NPs | TiO2 (Alexa647) |
| exposure | 10:1, 1h |
| imaging | xy confocal FLIM |

| | |
|---|---|
| pixelsize (x,y) | 100 nm |
| FOV (x,y) | 51,2 µm |
| TCSPC pixelsize | 122 ps |
| TCSPC FOV | 19.5 ns |

| | |
|---|---|
| 561nm | |
| 640nm | 10% |
| STED | |
| diffraction grating | 523 – 722 nm |
| dwell time | 200 µs |
| objective | 60x wi (NA1.2) |

*Figure S85: Analysis of fluorescence lifetimes of aggregated labelled nanomaterial – fluorescence image, fluorescence-lifetime-color-coded image and distribution of fluorescence lifetimes in the image.*

experiment 20190419_e06_s01_t01 (smaller aggregates of TiO2-Alexa647 in LCIS):

use binning of 10 pixels to measure the free-floating nanomaterial:

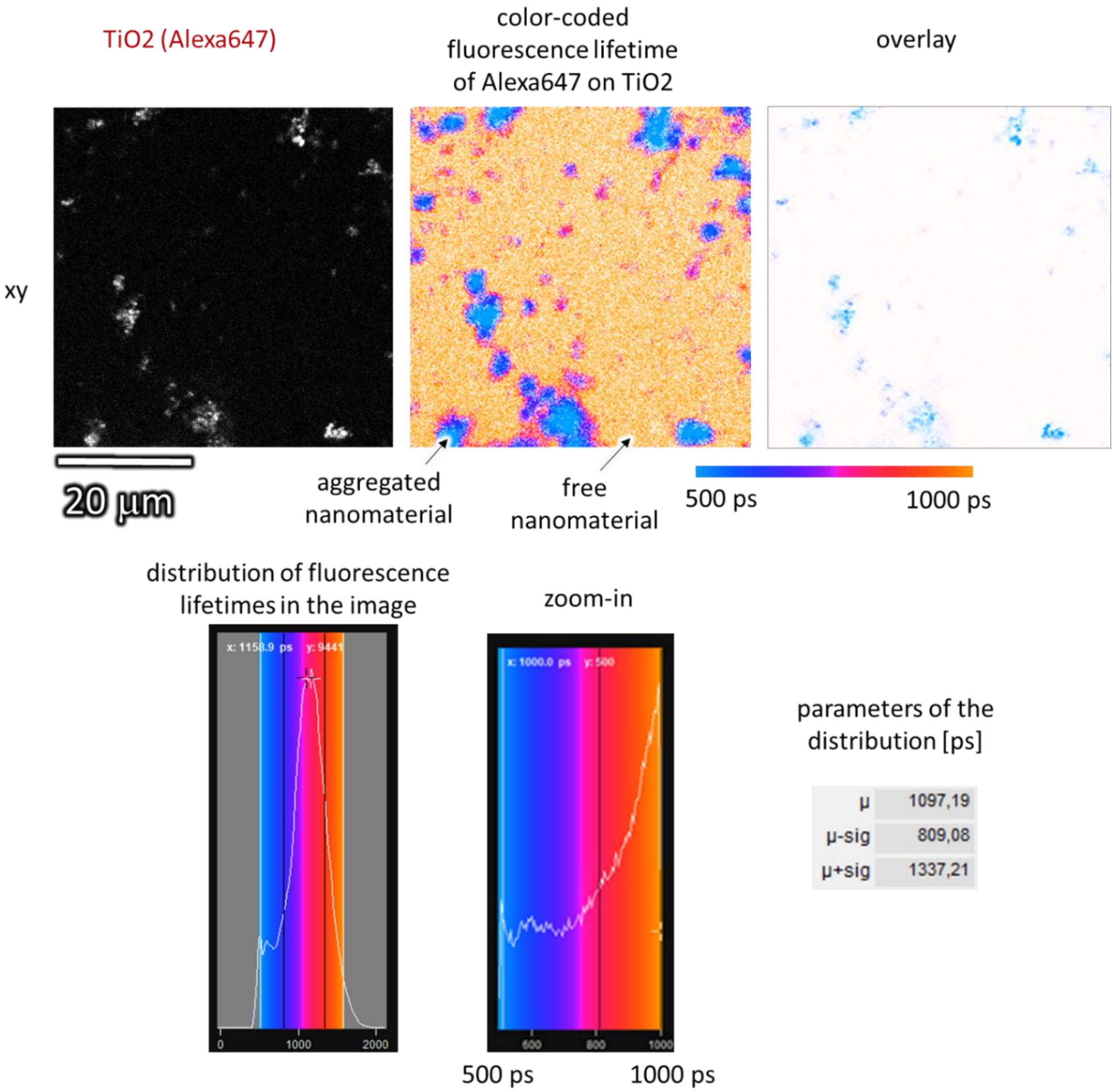

| | |
|---|---|
| Cell line | |
| NPs | TiO2 (Alexa647) |
| exposure | 10:1, 1h |
| imaging | xy confocal FLIM |

| | |
|---|---|
| pixelsize (x,y) | 100 nm |
| FOV (x,y) | 51,2 µm |
| TCSPC pixelsize | 122 ps |
| TCSPC FOV | 19.5 ns |

| | |
|---|---|
| 561nm | |
| 640nm | 10% |
| STED | |
| diffraction grating | 523 – 722 nm |
| dwell time | 200 µs |
| objective | 60x wi (NA1.2) |

*Figure S86: Analysis of fluorescence lifetimes of free-floating and aggregated labelled nanomaterial – fluorescence image, fluorescence-lifetime-color-coded image and distribution of fluorescence lifetimes in the image.*

experiment 20190419_e06_s02_t02 (giant aggregate of $TiO_2$-Alexa647 in LCIS):

TiO2 (Alexa647) | color-coded fluorescence lifetime of Alexa647 on TiO2 | overlay

xy

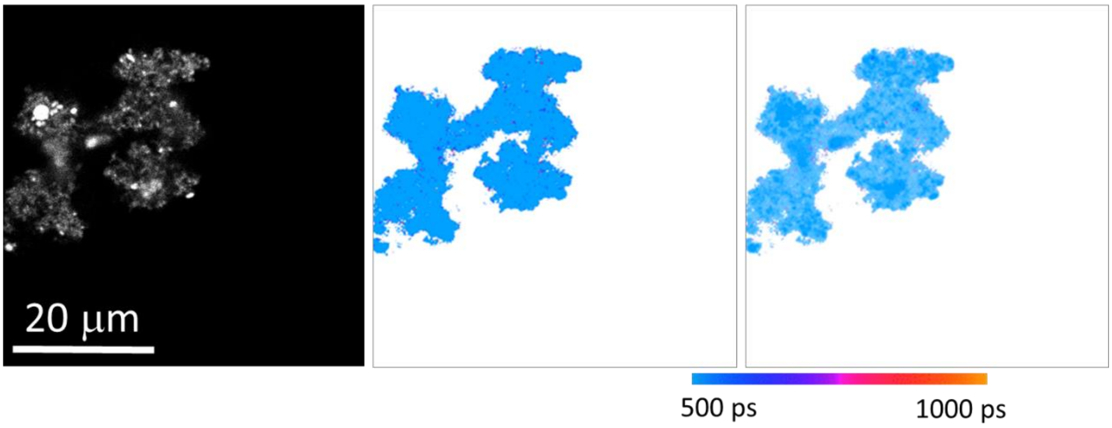

distribution of fluorescence lifetimes in the image | zoom-in

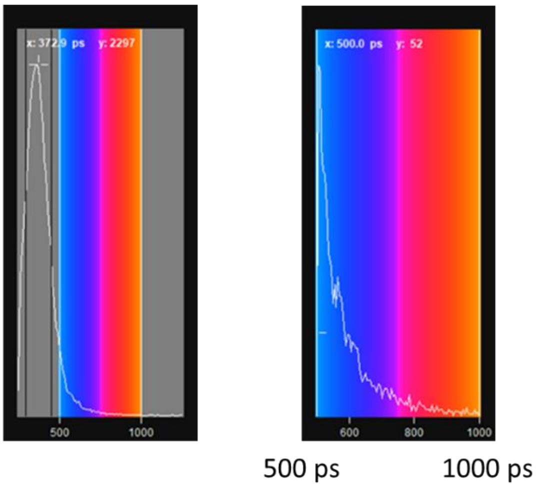

parameters of the distribution [ps]

| | |
|---|---|
| µ | 374,11 |
| µ-sig | 295,27 |
| µ+sig | 454,31 |

| | |
|---|---|
| Cell line | |
| NPs | TiO2 (Alexa647) |
| exposure | 10:1, 4h |
| imaging | xy confocal FLIM |

| | |
|---|---|
| pixelsize (x,y) | 100 nm |
| FOV (x,y) | 51,2 µm |
| TCSPC pixelsize | 122 ps |
| TCSPC FOV | 19.5 ns |

| | |
|---|---|
| 561nm | |
| 640nm | 10% |
| STED | |
| diffraction grating | 523 – 722 nm |
| dwell time | 50 µs |
| objective | 60x wi (NA1.2) |

*Figure S87: Analysis of fluorescence lifetimes of aggregated labelled nanomaterial – fluorescence image, fluorescence-lifetime-color-coded image and distribution of fluorescence lifetimes in the image.*

experiment 20190419_e06_s02_t02 (giant aggregate of TiO2-Alexa647 in LCIS):

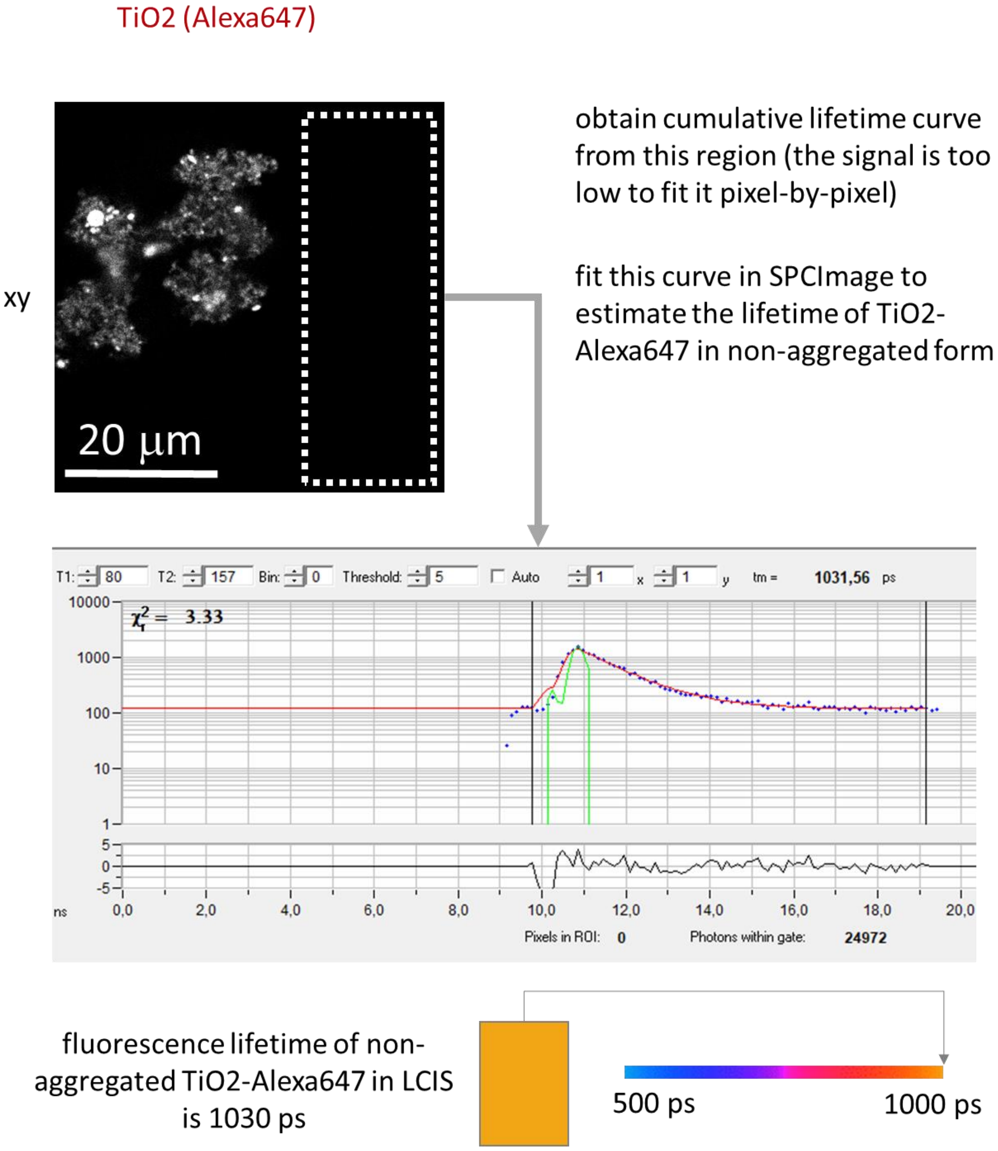

| | |
|---|---|
| Cell line | |
| NPs | TiO2 (Alexa647) |
| exposure | 10:1, 4h |
| imaging | xy confocal FLIM |

| | |
|---|---|
| pixelsize (x,y) | 100 nm |
| FOV (x,y) | 51,2 µm |
| TCSPC pixelsize | 122 ps |
| TCSPC FOV | 19.5 ns |

| | |
|---|---|
| 561nm | |
| 640nm | 10% |
| STED | |
| diffraction grating | 523 – 722 nm |
| dwell time | 50 µs |
| objective | 60x wi (NA1.2) |

*Figure S88: Analysis of fluorescence lifetime of free-floating labelled nanomaterial – fluorescence image and mean fluorescence lifetime in the marked area.*

experiment 20190628_fc04_s01_t01 (smaller aggregates of TiO2-Alexa647 in LCIS):

TiO2 (Alexa647) | color-coded fluorescence lifetime of Alexa647 on TiO2 | overlay

xy

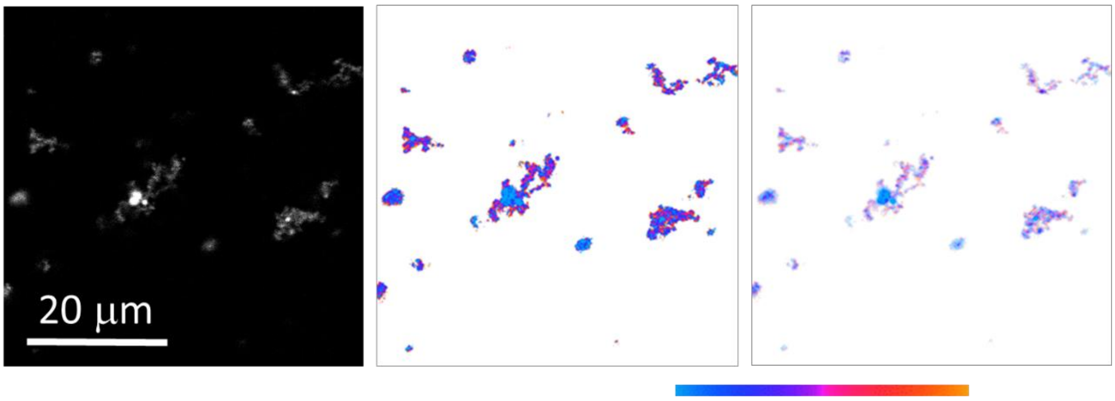

500 ps 1000 ps

distribution of fluorescence lifetimes in the image | zoom-in

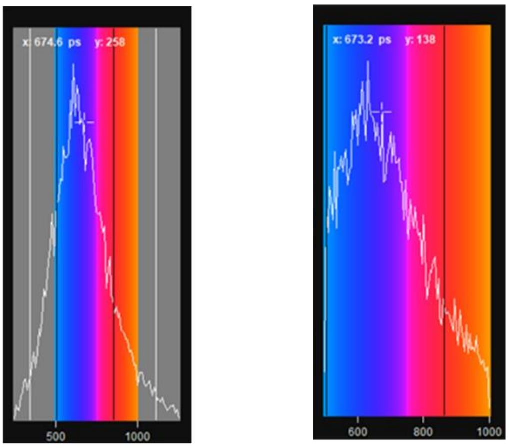

500 ps 1000 ps

parameters of the distribution [ps]

| | |
|---|---|
| μ | 673,64 |
| μ-sig | 506,83 |
| μ+sig | 862,69 |

| | |
|---|---|
| Cell line | |
| NPs | TiO2 (Alexa647) |
| exposure | 10:1, 1h |
| imaging | xy confocal FLIM |

| | |
|---|---|
| pixelsize (x,y) | 100 nm |
| FOV (x,y) | 51,2 μm |
| TCSPC pixelsize | 122 ps |
| TCSPC FOV | 19.5 ns |

| | |
|---|---|
| 561nm | |
| 640nm | 30% |
| STED | |
| diffraction grating | 523 – 722 nm |
| dwell time | 200 μs |
| objective | 60x wi (NA1.2) |

*Figure S89: Analysis of fluorescence lifetimes of aggregated labelled nanomaterial – fluorescence image, fluorescence-lifetime-color-coded image and distribution of fluorescence lifetimes in the image.*

experiment 20190628_fc04_s01_t01 (smaller aggregates of TiO2-Alexa647 in LCIS):

use binning of 10 pixels to measure the free-floating nanomaterial:

TiO2 (Alexa647) | color-coded fluorescence lifetime of Alexa647 on TiO2 | overlay

xy

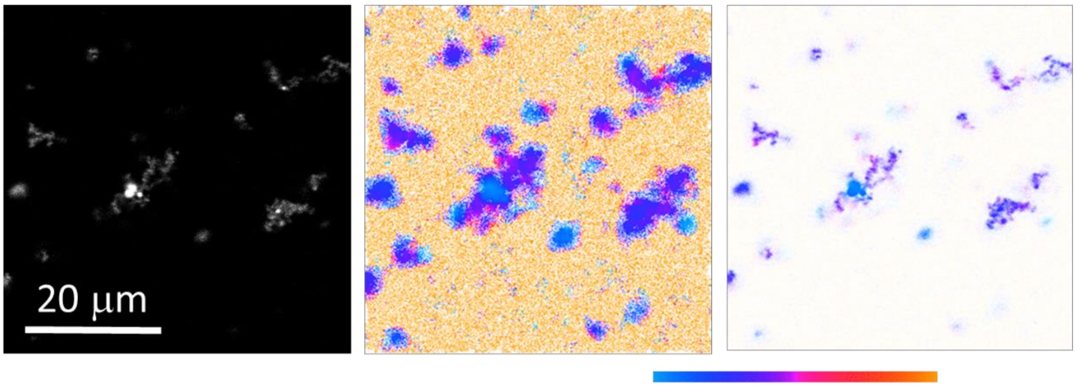

500 ps 1000 ps

distribution of fluorescence lifetimes in the image | zoom-in

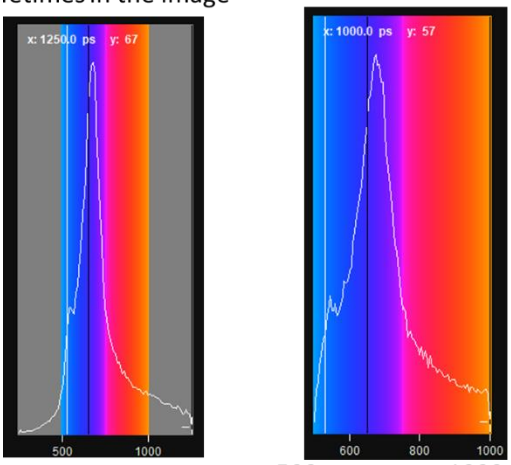

500 ps 1000 ps

parameters of the distribution

| | | |
|---|---|---|
| µ | 2099,66 | 57 |
| µ-sig | 651 | 1559 |
| µ+sig | 2723 | 57 |

| | |
|---|---|
| Cell line | |
| NPs | TiO2 (Alexa647) |
| exposure | 10:1, 1h |
| imaging | xy confocal FLIM |

| | |
|---|---|
| pixelsize (x,y) | 100 nm |
| FOV (x,y) | 51,2 µm |
| TCSPC pixelsize | 122 ps |
| TCSPC FOV | 19.5 ns |

| | |
|---|---|
| 561nm | |
| 640nm | 30% |
| STED | |
| diffraction grating | 523 – 722 nm |
| dwell time | 200 µs |
| objective | 60x wi (NA1.2) |

*Figure S90: Analysis of fluorescence lifetimes of aggregated and free-floating labelled nanomaterial – fluorescence image, fluorescence-lifetime-color-coded image and distribution of fluorescence lifetimes in the image.*

experiment 20190621_e01.100_s02_t04 (cauliflower FLIM):

| | TiO2 (Alexa647) | color-coded fluorescence lifetime of Alexa647 on TiO2 | overlay |
|---|---|---|---|

xy

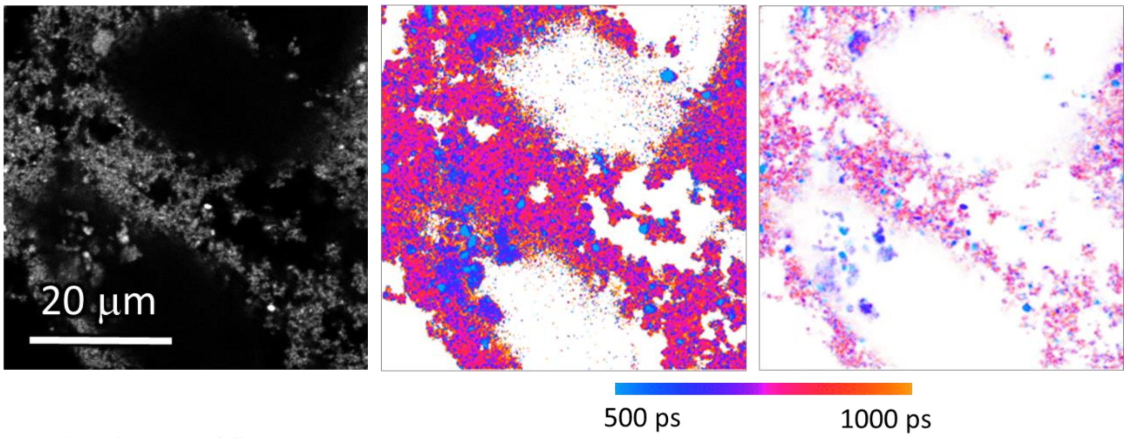

distribution of fluorescence lifetimes in the image

zoom-in

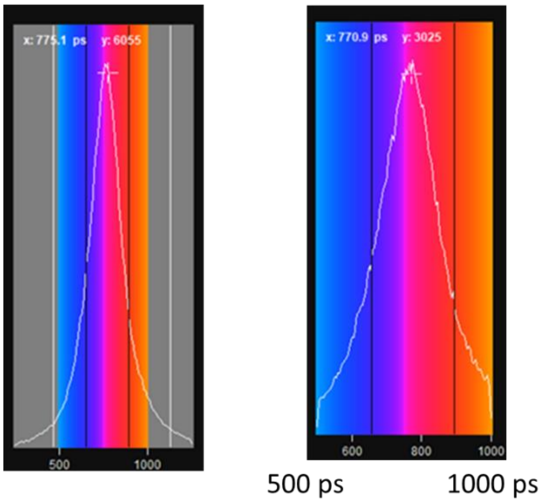

parameters of the distribution

| | |
|---|---|
| µ | 773,76 |
| µ-sig | 655,77 |
| µ+sig | 895,16 |

| | |
|---|---|
| Cell line | LA-4 (membrane, CellMaskOrange and lipid bodies, SAG-38) |
| NPs | TiO2 (Alexa647) |
| exposure | 100:1, 29h |
| imaging | xy confocal FLIM |

| | |
|---|---|
| pixelsize (x,y) | 100 nm |
| FOV (x,y) | 51,2 µm |

| | |
|---|---|
| TCSPC pixelsize | 122 ps |
| TCSPC FOV | 19.5 ns |

| | |
|---|---|
| 561nm | |
| 640nm | 10% |
| STED | |
| diffraction grating | 523 – 722 nm |
| dwell time | 100 µs |
| objective | 60x wi (NA1.2) |

*Figure S91: Analysis of fluorescence lifetimes of nanomaterial in cauliflowers – fluorescence image, fluorescence-lifetime-color-coded image and distribution of fluorescence lifetimes in the image.*

experiment 20190419_ee12_s01_t01 (free Alexa647 in LCIS):

Alexa647
color scale 0 – 3 counts

xy

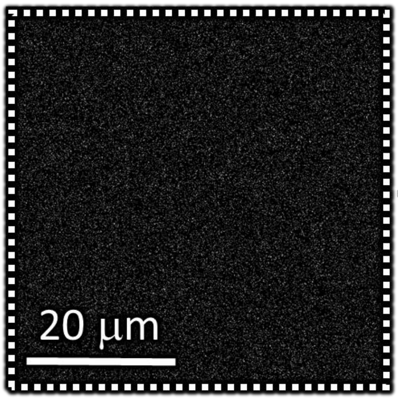

obtain cumulative lifetime curve from this region (the signal is too low to fit it pixel-by-pixel)

fit this curve in SPCImage to estimate the lifetime of TiO2-Alexa647 in non-aggregated form

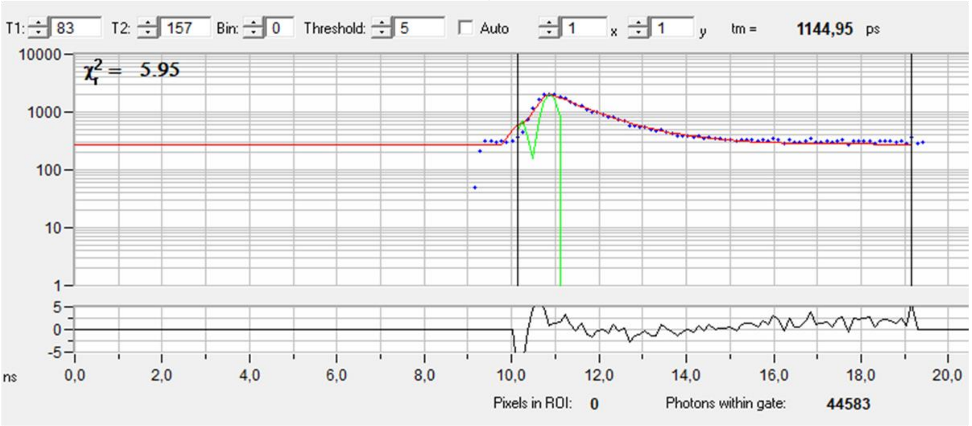

fluorescence lifetime of Alexa647 in LCIS is 1140 ps

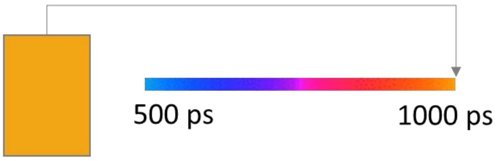

| | |
|---|---|
| Cell line | |
| NPs | Free Alexa647 |
| exposure | |
| imaging | xy confocal FLIM |

| | |
|---|---|
| pixelsize (x,y) | 100 nm |
| FOV (x,y) | 51,2 µm |
| TCSPC pixelsize | 122 ps |
| TCSPC FOV | 19.5 ns |

| | |
|---|---|
| 561nm | |
| 640nm | 10% |
| STED | |
| diffraction grating | 523 – 722 nm |
| dwell time | 50 µs |
| objective | 60x wi (NA1.2) |

*Figure S92: Determination of fluorescence lifetime of free-floating labelled nanomaterial – fluorescence image and mean fluorescence lifetime in the marked area.*

experiment 20190628_fc02_s02 (FLIM of free Alexa647 on LA4 cells):

| TiO2 (Alexa647) | color-coded fluorescence lifetime of Alexa647 on TiO2 | overlay |
|---|---|---|

xy

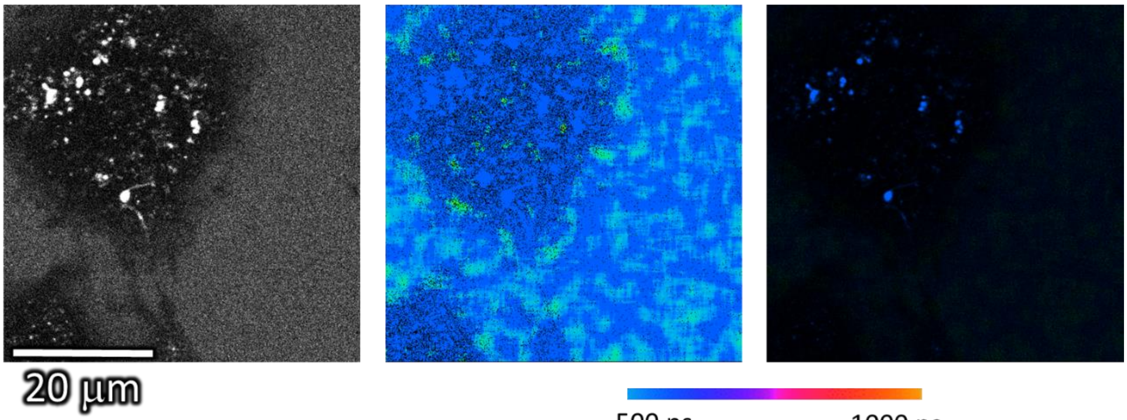

| distribution of fluorescence lifetimes in the image | zoom-in |
|---|---|

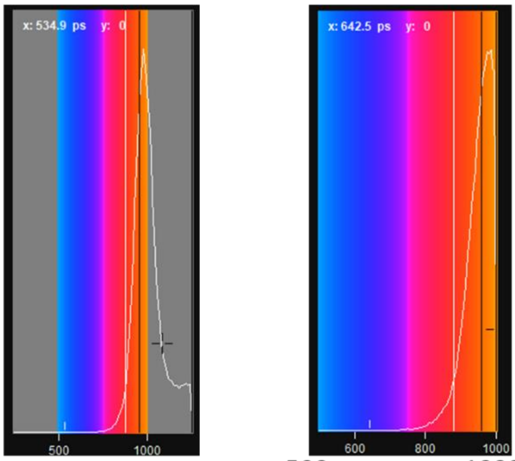

parameters of the distribution [ps]

| | | |
|---|---|---|
| µ | 1085,53 | 1940 |
| µ-sig | 956,91 | 6669 |
| µ+sig | 1386,39 | 1940 |

| | |
|---|---|
| Cell line | LA-4 (Alexa647, 2uM, 1h incubation) |
| NPs | |
| exposure | |
| imaging | xy confocal FLIM |

| | |
|---|---|
| pixelsize (x,y) | 100 nm |
| FOV (x,y) | 51,2 µm |
| TCSPC pixelsize | 122 ps |
| TCSPC FOV | 19.5 ns |

| | |
|---|---|
| 561nm | |
| 640nm | 30% |
| STED | |
| diffraction grating | 523 – 722 nm |
| dwell time | 600 µs |
| objective | 60x wi (NA1.2) |

1hour 2uM Alexa647 >> c(Alexa) on 1:1 NPs

*Figure S93: Analysis of fluorescence lifetimes of free Alexa Fluor 647 if exposed to LA-4 cells for 1 hour at 2uM concentration – fluorescence image, fluorescence-lifetime-color-coded image and distribution of fluorescence lifetimes in the image.*

experiment 20190808_e01_s01 (CellMask Orange)

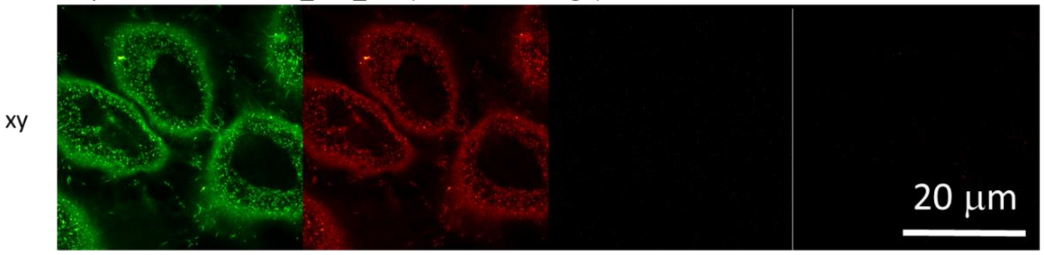

experiment 20190607_e03_s02_t02 (SAG-38)

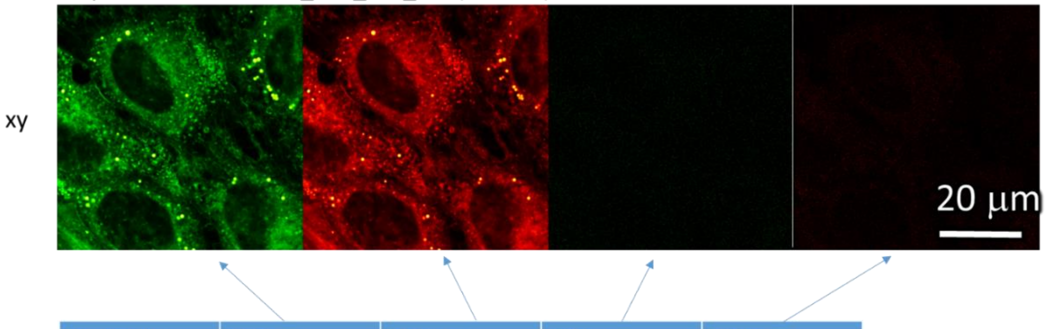

| excitation | 561nm | 561nm | 640nm | 640nm | |
|---|---|---|---|---|---|
| detector | 605-625 nm | 650-720 nm | 605-625 nm | 650-720 nm | |
| CellMask Orange | 204 | 69 | 2 | 3 | # of counts on the detector |
| SAG-38 | 180 | 151 | 3 | 3 | |

640 nm laser does not excite CellmaskOrange or SAG-38
-> the signal of these probes is not seen in FLIM images when the sample is excited with 640 nm laser
-> all FLIM histograms, excited with 640nm laser can be attributed to Alexa647

| | |
|---|---|
| Cell line | LA-4 (membrane, CellMaskOrange) |
| NPs | TiO2 (Alexa647) |
| exposure | 10:1, 2 days |
| imaging | xy confocal |

| | |
|---|---|
| pixelsize (x,y) | 100 nm |
| FOV (x,y) | 80.0 µm |
| pixelsize (z) | |
| FOV (z) | |
| imaging time | |
| number of frames | |

| | |
|---|---|
| 561nm | 5% |
| 640nm | 5% |
| STED | |
| filter sets | 605 nm – 625 nm, 650 nm – 720 nm |
| dwell time | 10 µs |
| objective | 60x wi (NA 1.2) |

| | |
|---|---|
| Cell line | LA-4 (lipid bodies, SAG-38) |
| NPs | TiO2 (Alexa647) |
| exposure | 10:1, 2 days |
| imaging | xy confocal |

| | |
|---|---|
| pixelsize (x,y) | 50 nm |
| FOV (x,y) | 60.0 µm |
| pixelsize (z) | |
| FOV (z) | |
| imaging time | |
| number of frames | |

| | |
|---|---|
| 561nm | 2% |
| 640nm | 2% |
| STED | |
| filter sets | 605 nm – 625 nm, 650 nm – 720 nm |
| dwell time | 10 µs |
| objective | 60x wi (NA 1.2) |

*Figure S94: Crosstalk reference for CellMaskOrange and SAG-38 – these two probes are not excited by 640 nm laser. Since all FLIM measurements in this paper are done by 640 nm excitation, they represent the fluorescence lifetimes of solely the nanomaterial with no artefacts from the membrane label signal.*

#### S2d – Transcriptomics *in vitro* and *in vivo* after exposure to $TiO_2$ and comparison of both

##### *In vitro*

##### Main Message

The transcriptome profile of LA-4 cells exposed to $TiO_2$ nanotubes is analysed in terms of gene sets and pathways, which are mechanistically involved in cauliflower formation: lipid metabolism, immune system, vesicular trafficking and actin cytoskeleton organization.

##### Results

###### *Gene analysis*

Lipid Metabolism

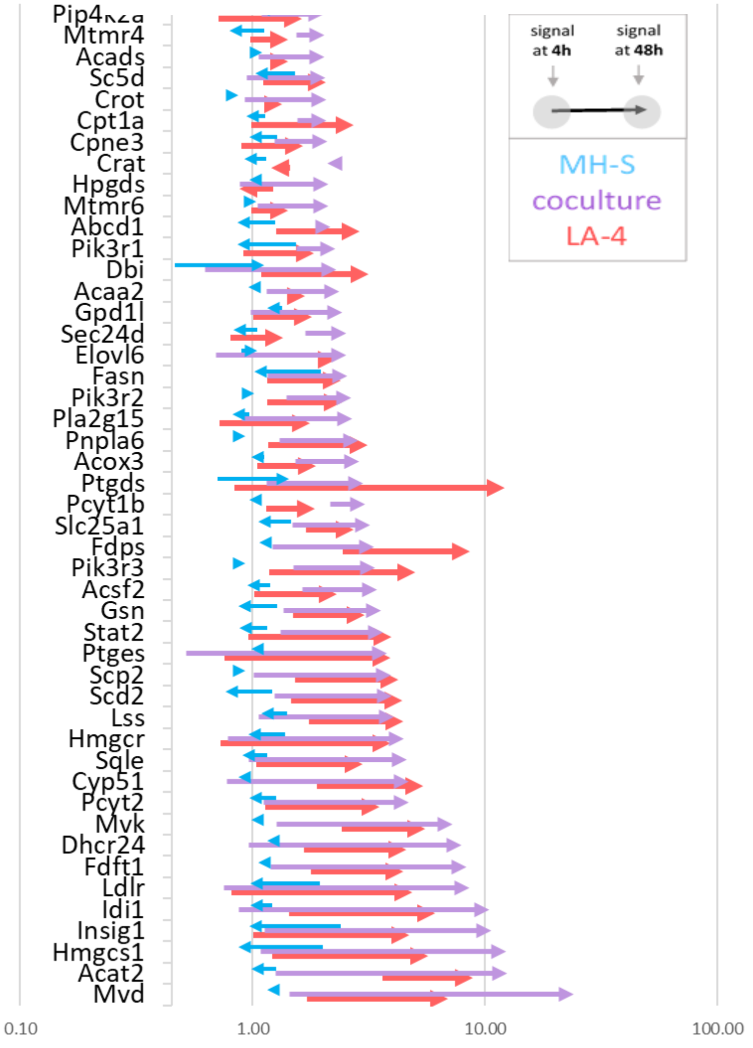

*Figure S95: Lipid metabolism related genes with more than two fold increased expressions. Fold increase in expression is determined according to the non-exposed control sample at the same time point. Arrows represent the time evolution of gene expression so that the beginning of the arrow shows expression at 4 h and an arrowhead shows the expression at the 48 h time point*

Actin Related Genes

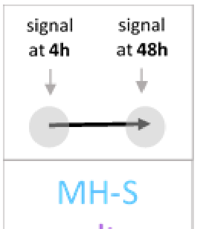

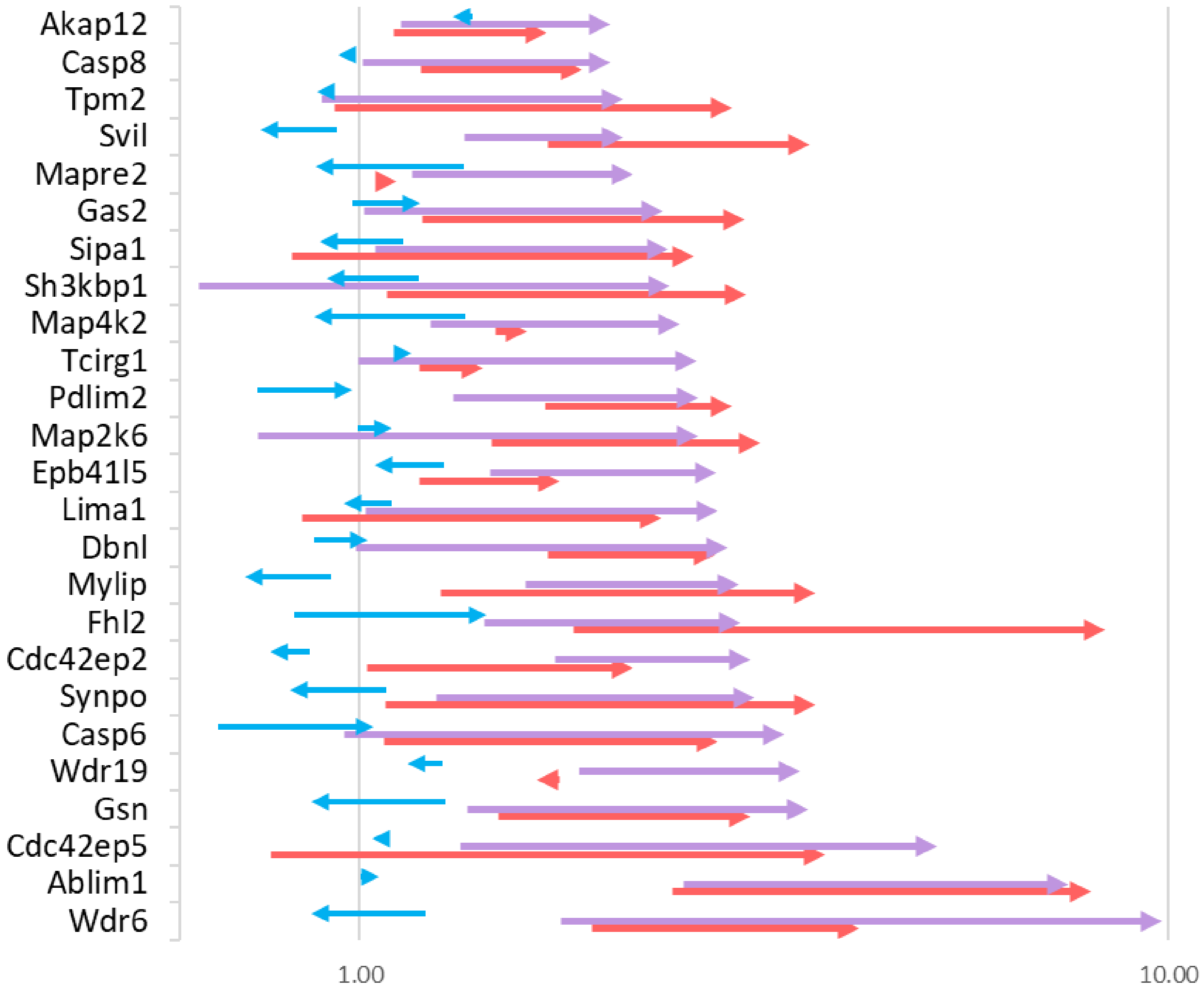

*Figure S96: Actin cytoskeleton related genes with more than two fold increased expressions. Fold increase in expression is determined according to the non-exposed control sample at the same time point. Arrows represent the time evolution of gene expression so that the beginning of the arrow shows expression at 4 h and an arrowhead shows the expression at the 48 h time point*

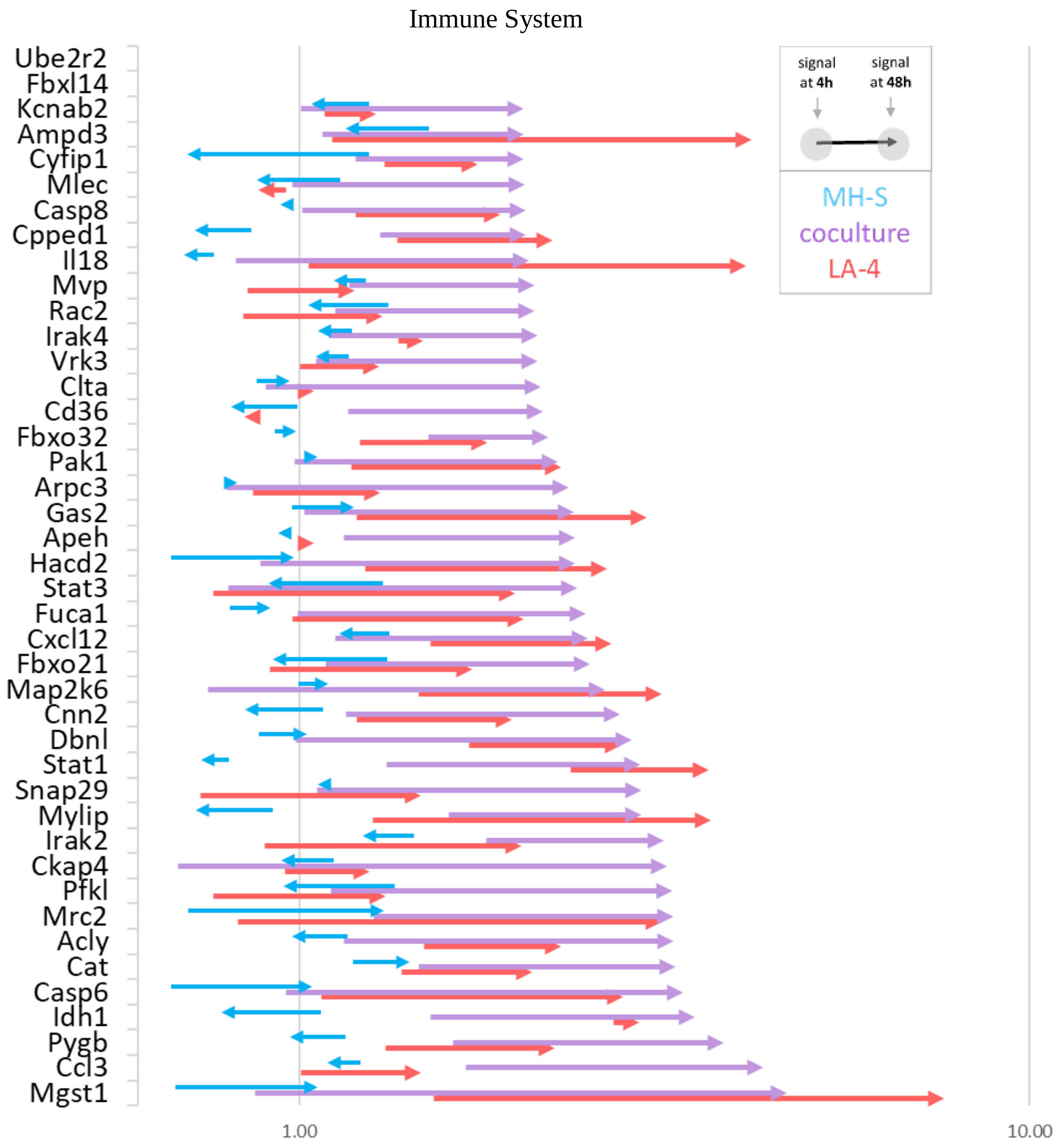

*Figure S97: Immune response related genes with more than two fold increased expressions. Fold increase in expression is determined according to the non-exposed control sample at the same time point. Arrows represent the time evolution of gene expression so that the beginning of the arrow shows expression at 4 h and an arrowhead shows the expression at the 48 h time point.*

#### Pathway analysis

To further analyse the cellular response to the $TiO_2$ exposure, gene expression analysis was performed and related pathways were assessed by gene set enrichment analysis of mono and co-cultures of epithelial cells (LA-4) and macrophages (MH-S) at different time points. As described for cauliflower formation, also lipid metabolism and actin-dependent pathways were significantly enriched in epithelial cells and co-cultures with macrophages, but not in single cultures of macrophages. This supports the idea that upon uptake, nanoparticle excretion and cauliflowers formation is specific for epithelial cells but for professional phagocytes. Most of the pathways are abundantly enriched during the early phase 4h after nanomaterial exposure, such as cholesterol homeostasis, bile acid metabolism, from the lipid pathways, while others like peroxisome regulation show a stronger enrichment over time. This time dependence can be taken from the colour coded NES values (Figure S98 left) or the respective enrichment plots (Figure S98 right). Interestingly, only macrophages monocultures showed an inflammatory signature evidenced by the enrichment of TNFα/NFkB and IL-2/STAT5 signalling pathways, as well as for alveolar macrophages typical oxidative phosphorylation. None of these showed enrichment in epithelial cells single cultures (Figure S98 left).

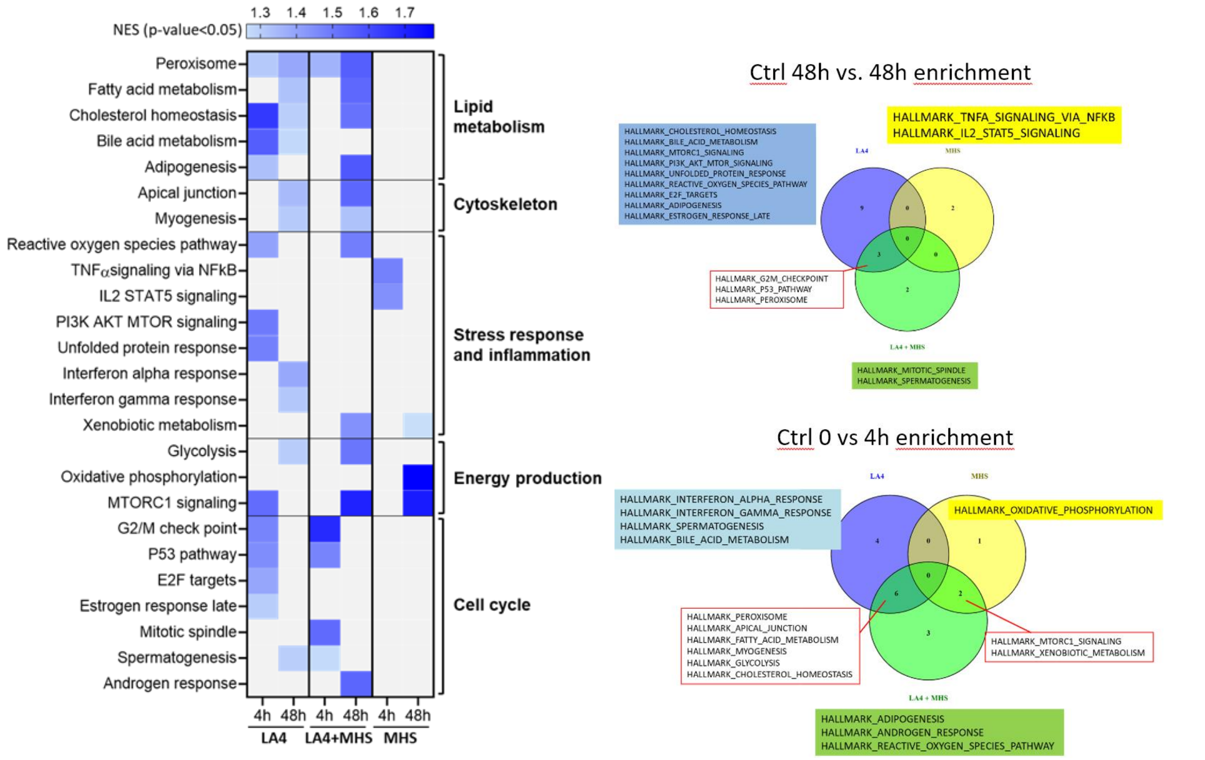

*Figure S98: Left: Heat maps for enrichment of hallmark pathways: lipid metabolism, cytoskeleton stress response and inflammation, energy production and cell cycle, (NES: normalized enrichment score) gene expression on a pathway level for for genes increased in two-fold. Right: Venn diagrams.*

4h 48h

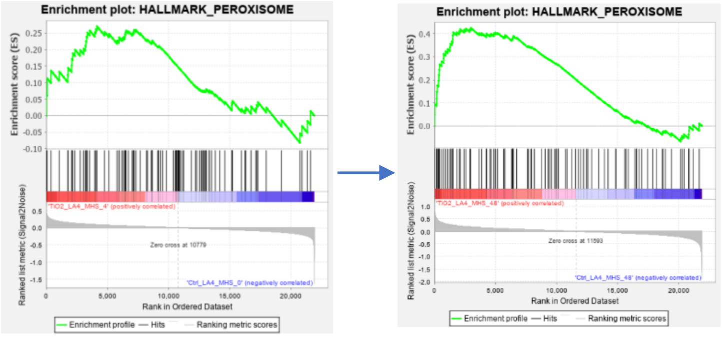

*Figure S99: Enrichment plots of time progression of the peroxisome pathway (co-cultures)*

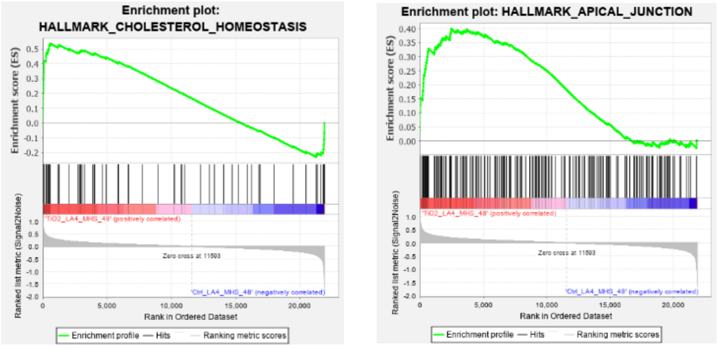

*Figure S100: Enrichment plots of lipid metabolism and cytoskeleton pathways (co-cultures, 48h)*

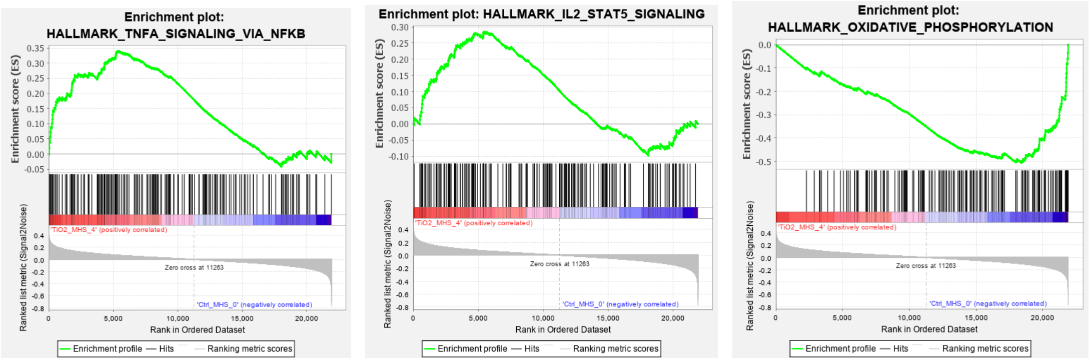

*Figure S101: Enrichment plots of pro-inflammatory and oxidative phosphorylation pathways (MH-S mono-culture, 4h)*

#### Materials and Methods

##### *Sample preparation*

Samples for the experiments, LA-4, MH-S, and cocultures were grown in 6-well plates until desired confluency. Cocultures were seeded so that the ratio of LA-4:MH-S was approximately 40:1 which reflects the physiological conditions of human alveoli. Cells were exposed to nanoparticles ($TiO_2$ or MWCNT) at a 10:1 surface dose ($NP_{surface}$ to $Cell_{surface}$ ratio) when they reached 90 % confluency. Cells were exposed to $TiO_2$ nanotubes and MWCNT for 4 h and 48 h and control samples were taken at 0 h and 48 h. Samples were prepared as described above. Briefly, growth medium was removed and 6-well plates containing cells only were frozen at -70°C. Detailed preparation of samples is given in the following tables:

*Table S5: Sample preparation for the 4 h exposure of LA-4, MH-S and their co-culture to $TiO_2$ nanotubes*

| 4h exposure | LA-4 | Coculture | MH-S |
|---|---|---|---|
| 1st day | 25 % LA-4 | 25 % LA-4 | / |
| 3rd day | Δ medium | Δ medium + 2 % MH-S | MH-S 50 % |
| 5th day | Δ medium + $TiO_2$ | | |
| 5th day | HARVEST DAY | | |

*Table S6: Sample preparation for the 48 h exposure of LA-4, MH-S and their co-culture to $TiO_2$ nanotubes*

| 48h exposure | LA-4 | Coculture | MH-S |
|---|---|---|---|
| 1st day | 15 % LA-4 | 15 % LA-4 | / |
| 3rd day | Δ medium | Δ medium + 1 % MH-S | MH-S 20 % |
| 5th day | Δ medium + $TiO_2$ | | |
| 7th day | HARVEST DAY | | |

Control samples were prepared in the same manner without exposure to nanoparticles.

##### *RNA isolation*

Total RNA was isolated employing the RNeasy Plus Mini Kit (Qiagen). The Agilent 2100 Bioanalyzer was used to assess RNA quality and RNA with RIN>7 was used for microarray analysis.
Total RNA (120 ng) was amplified using the WT PLUS Reagent Kit (Thermo Fisher Scientific Inc., Waltham, USA). Amplified cDNA was hybridized on Mouse Clariom S arrays (Thermo Fisher Scientific). Staining and scanning (GeneChip Scanner 3000 7G) was done according to manufacturer`s instructions.

##### *Statistical analysis*

Statistical analysis for all probe sets included limma t-test and Benjamini-Hochberg multiple testing correction. Raw p-values of the limma t-test wereused to define sets of regulated genes ($p<0.01$). Detection Above Background (dabg) p-values were used to exclude background signals: significant genes were filtered for $p<0.05$ in more than half of the samples in at least one group. Array data has been submitted to the GEO database at NCBI (GSE146036).

##### *Gene Set Enrichment Analysis*

GSEA software from the Broad Institute (*http://www.gsea-msigdb.org/gsea/index.jsp*) (*Gene set enrichment analysis: a knowledge-based approach for interpreting genome-wide expression profiles.Proc Natl Acad Sci U S A. 2005; 102: 15545-15550*) was used to identify enrichment of defined gene sets in our microarray data (GSE146036). Among the molecular signature databases offered by the GSEA collection, The Hallmark (H) database was selected due its consistent representation of specific biological processes and lack of redundancy (*Liberzon, Arthur, et al. "The molecular signatures database hallmark gene set collection." Cell systems 1.6 (2015): 417-425.*). Analysis was run under default settings of the GSEA software.

##### *Arrow graphs*

In the arrow graphs, only genes which were up- or down-regulated more than two-fold compared to non-exposed cells are shown. The signal (x axis) is drawn in logarithmic scale. Expression is normalized to expression of control samples. Genes in arrow charts are ordered according to their fold increase at 48 hours (fold increase was calculated in regard to non-exposed cells at the same time-point). Gene sub-sets were selected manually.

#### Experiment names

20200117_genomics_TiO2_results.xlsx

##### *In vivo*

#### Main Message

The transcriptome profile of female C57BL/6 mice at 1 and 28 days post-exposure to $TiO_2$ nanotubes shows enriched gene sets involved in fatty acid metabolism after 28 days thus supporting our findings in *in vitro* system. Furthermore, it shows a strong immune response, in terms of chemokine signaling, additionally strengthening our hypothesis.

*Table S7: Lipid-related pathways from the KEGG pathway analysis. The analysis was performed using the GSEA method and the KEGG signalling pathways database. Pathways were selected if they were identified with FDR corrected p-value < 0.05. The values in the table indicate -log10(FDR corrected p-value).*

| | *in vivo* | | | | | |
|---|---|---|---|---|---|---|
| KEGG term name | 18 µg d1 | 18 µg d28 | 54 µg d1 | 54 µg d28 | 162 µg d1 | 162 µg d28 |
| Glycosphingolipid biosynthesis - ganglio series | 1.18 | 1.19 | 1.01 | 1.15 | - | - |
| Sphingolipid signaling pathway | 2.13 | - | 4.54 | - | 1.71 | - |
| Glycerophospholipid metabolism | - | - | - | - | - | - |
| Ether lipid metabolism | 1.56 | - | - | 1.73 | - | - |
| Fatty acid degradation | - | 5.63 | 1.10 | 2.40 | - | - |
| Fatty acid metabolism | - | 4.31 | - | 1.95 | - | - |
| Fatty acid elongation | - | 1.52 | - | 1.46 | - | - |
| Biosynthesis of unsaturated fatty acids | - | - | - | - | - | - |
| Glycosaminoglycan biosynthesis - keratan sulfate | 1.40 | - | - | 1.46 | - | - |
| Non-alcoholic fatty liver disease (NAFLD) | - | 10.52 | 1.80 | 2.55 | - | - |

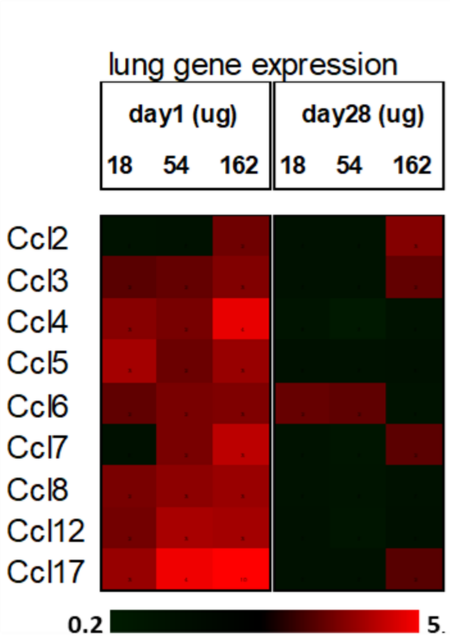

*Figure S102: Genes encoding monocyte chemoattractive (C-C motif) chemokines upregulated in mice exposed to $TiO_2$ nanotubes after 1 and 28 days.*

#### Cell composition in bronchoalveolar lavage fluid (BAL)

Danielsen et al. [4] assessed inflammatory cells recruitment in BAL fluid at 1, 3, 28, 90 and 180 days post-exposure as a pulmonary inflammatory response marker. For the $TiO_2$ tube, the number of neutrophils and macrophages was statistically significant increased at day 28 post-exposure. As a response to acute inflammation number of neutrophils was, in the dose dependent matter, highest after the first day of exposure and kept decreasing towards 28$^{th}$ day, but remained elevated nevertheless. Number of macrophages was elevated on the 1$^{st}$ day post-exposure and kept increasing until the 28$^{th}$ day in a dose dependent matter, indicative of chronic inflammation. The cell dynamics reflects the transcriptomics data both in vitro and in vivo, where we can see acute response transitioning into a chronic one.

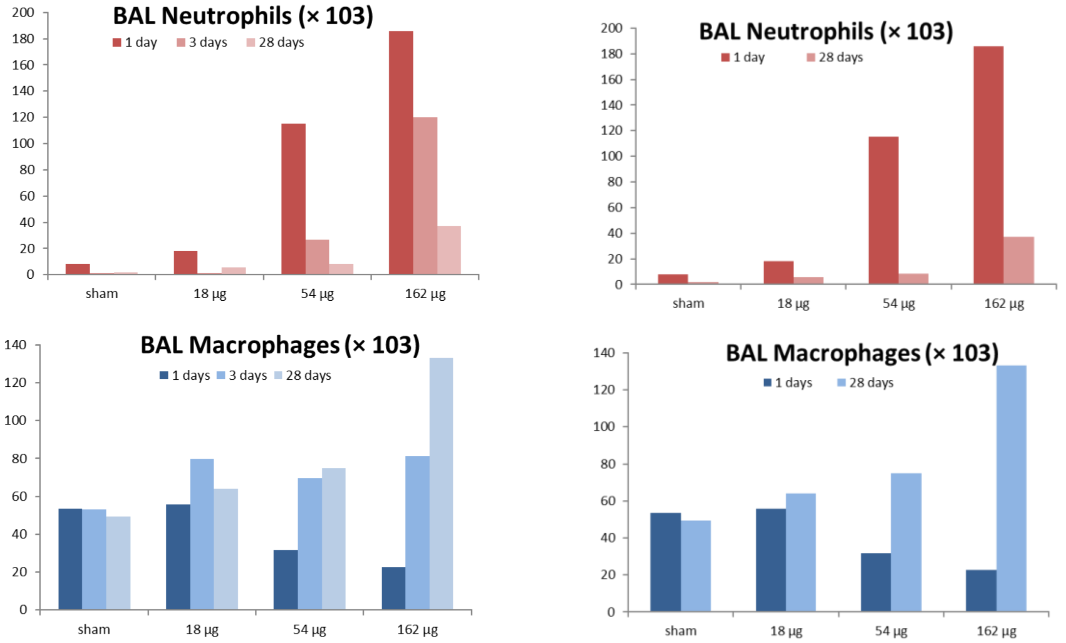

*Figure S103: Neutrophil and macrophage recruitment in BAL fluid at 1, 3, 28, 90 and 180 days post-exposure*

#### Materials and Methods

##### *Sample preparation*

The materials and methods used for intratracheal instillation of mice with TiO2 tube are described in detail by Danielsen et. al [4] and included in this document (S1b – *In vivo* data) in a short version. Female C57BL/6 mice were exposed by single intratracheal instillation to 18, 54 or 162 µg/mouse of a TiO2 tube. Lung tissues were harvested on day 1 and 28 after exposure. Microarray mRNA analysis was performed using Agilent 8 × 60 K oligonucleotide microarrays (Agilent Technologies Inc., Mississauga, ON, Canada) as described previously [2] with 6 replicas for each condition. Bioinformatics analysis of the row data: signal intensities were Loess normalized using the limma package in R/Bioconductor [3]. Analysis of differentially expressed genes (DEGs) was performed using the limma package. The genes were considered as significantly differentially expressed if the BH-adjusted p-values were less than or equal to 0.1.

##### *RNA isolation*

RNA was isolated and treated in the same manner as in the *in vitro* experiment described above.

##### *Statistical analysis*

Statistical analysis for all probe sets included limma t-test and Benjamini-Hochberg multiple testing correction. Raw p-values of the limma t-test were used to define sets of regulated genes ($p<0.01$). Detection Above Background (dabg) p-values were used to exclude background signals: significant genes were filtered for $p<0.05$ in more than half of the samples in at least one group. Array data has been submitted to the GEO database at NCBI (GSE146036).

##### *Gene Set Enrichment Analysis*

The KEGG gene set enrichment analysis was performed using camera function from the limma R package [3]. Pathways were selected if they were identified with FDR corrected p-value < 0.05. For the assessment of the monocyte influx, all genes encoding monocyte chemoattractive (C-C motif) chemokines were selected and their expression evaluated (Figure S102).

##### *Experiment names*

KEGG_pathway_analysis.xlsx

#### Comparison *in vivo* and *in vitro*

*Table S8: Lipid-related pathways from the KEGG pathway analysis. The analysis was performed using the GSEA method and the KEGG signalling pathways database. Pathways were selected if they were identified with FDR corrected p-value < 0.05. The values in the table indicate -log10(FDR corrected p-value).*

| | *in vitro* | | | | | | *in vivo* | | | | | |
|---|---|---|---|---|---|---|---|---|---|---|---|---|
| KEGG term name | LA-4 4h | LA-4 48h | LA-4 / MH-S | LA-4 / MH-S | MH-S 4h | MH-S 48h | 18 µg d1 | 18 µg d28 | 54 µg d1 | 54 µg d28 | 162 µg d1 | 162 µg d28 |

| | | | 4h | 48h | | | | | | | | |
|---|---|---|---|---|---|---|---|---|---|---|---|---|
| Glycosphingolipid biosynthesis - ganglio series | - | 1.35 | - | - | - | - | 1.18 | 1.19 | 1.01 | 1.15 | - | - |
| Sphingolipid signaling pathway | - | - | 1.57 | - | 1.81 | - | 2.13 | - | 4.54 | - | 1.71 | - |
| Glycerophospholipid metabolism | - | - | - | - | 1.76 | - | - | - | - | - | - | - |
| Ether lipid metabolism | - | - | - | - | - | - | 1.56 | - | - | 1.73 | - | - |
| Fatty acid degradation | - | 1.17 | - | 1.57 | - | - | - | 5.63 | 1.10 | 2.40 | - | - |
| Fatty acid metabolism | 1.14 | 2.49 | - | 3.75 | - | - | - | 4.31 | - | 1.95 | - | - |
| Fatty acid elongation | 1.04 | - | - | - | - | - | - | 1.52 | - | 1.46 | - | - |
| Biosynthesis of unsaturated fatty acids | - | - | - | 2.89 | - | - | - | - | - | - | - | - |
| Glycosaminoglycan biosynthesis - keratan sulfate | - | - | 1.17 | - | - | - | 1.40 | - | - | 1.46 | - | - |
| Non-alcoholic fatty liver disease (NAFLD) | - | - | 1.29 | - | 10.64 | - | - | 10.52 | 1.80 | 2.55 | - | - |
| Glycosaminoglycan biosynthesis - chondroitin sulfate / dermatan sulfate | - | - | - | - | 1.31 | - | 1.65 | - | 2.33 | 1.70 | - | - |
| Fatty acid biosynthesis | - | - | - | - | - | - | - | 2.03 | - | - | - | - |

##### Comparison of transcriptome response to TiO2 perturbation for *in vivo* and *in vitro* conditions

Mice were exposed to 18, 54 or 162 µg of $TiO_2$ nanotubes per mouse and lungs were harvested on 1$^{st}$ and 28$^{th}$ day post exposure for transcriptomic analysis to evaluate overlapping sets of genes differentially expressed in the *in vivo* and *in vitro* experimental data. The goal of the analysis is to determine and compare alterations in lipid metabolism, immune response in terms of proinflammatory signalling and cholesterol metabolism between two experimental systems.

###### *Experiment names*

KEGG_pathway_analysis.xlsx

#### S2e – *In silico* data – atomistic molecular dynamics simulation

##### Pool of lipids

*Main message*

From a pool of lipids (non-bilayer) the POPE and DMPC lipids attach to $TiO_2$ surface by lipid head-group. Due to less dense packing on the surface than in a bilayer a interleaved bi-layer is formed (with thickness between 2 and 3 nm (instead of usual 4 nm for bilayer).

Supporting raw and analysed data:

Figure S104-Figure S105

Table S 9-10

After a few tens of nanosecond from the simulations start, the lipids formed bilayer-like patches on the $TiO_2$ surface (Figure S105), binding to the surface by the headgroups, see Figure 2e (main text) and Figure S104. We found that PC and PE lipids bind to $TiO_2$ differently: while PE lipids bind by amino-group directly to the $TiO_2$ surface, PC lipids bind by the phosphate group through the intermediate layer of water molecules. These binding modes are shown in Figure S105 (A,B), and density profiles of N and P atoms of lipids are shown in Figure S105 (C,D). One can see that in the case of PE lipids the density maxima are sharper, and density profile of N-atoms is on a shorter distance from the bilayer surface compared to PC lipids, which is indication that PE lipids bind to anatase (101) surface stronger than PE lipids. In the both cases, one can see formation of the second lipid layer with lipids having hydrophobic tails contacting lipid tails of the first layer, and with lipid headgroups exposed to solvent. It would be naturally to suggest that headgroups of lipids of the second layer can bind to the surface of another $TiO_2$ nanoparticle, thus building lipid-nanoparticles complexes.

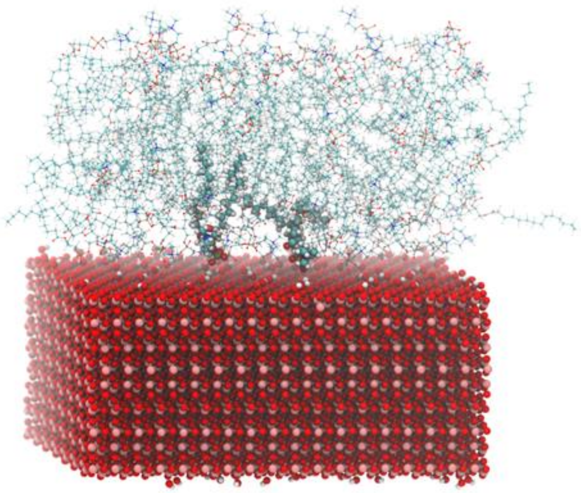

*Figure S104: DMPC lipid bilayer formed on $TiO_2$ anatase(101) surface. Two lipid molecules are highlighted; water molecules are not shown. In the beginning of simulations the lipids were dispersed in the solution*

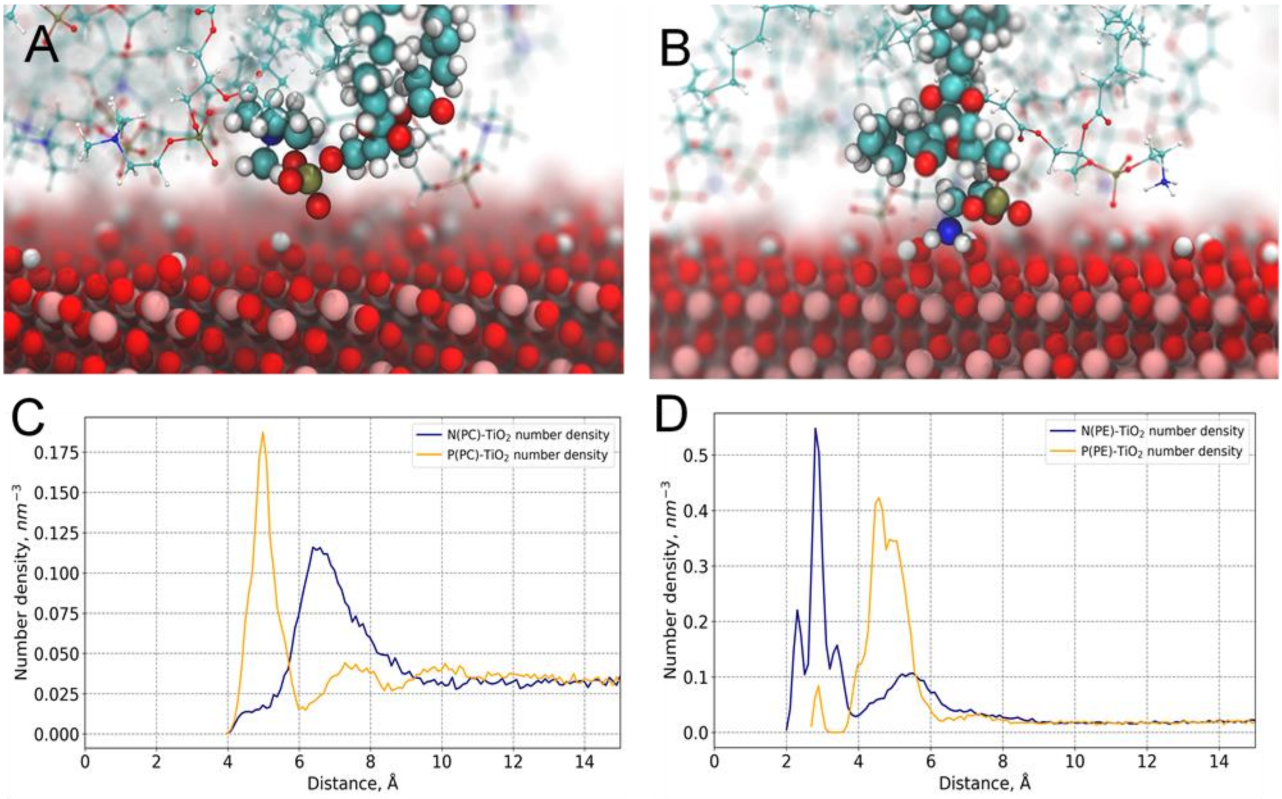

*Figure S105: Binding modes of DMPC (A) and POPE (B) lipids to anatase (101) $TiO_2$ surface, density profiles (in terms of number density) of DMPC choline and phosphate groups (C), and of POPE amino- and phosphate groups (D) near anatase (101) $TiO_2$ surface. The density profiles were computer after 50 ns of equilibration simulations. The distance is counted from the outmost layer of Ti atoms in the slab.*

#### Link to time-lapse

- [http://lbfnanobiodatabase.ijs.si/file/data/cauliflowerpaper/Fig2_atomistic_molecular_dynamics_binding_anatase-101-DMPC(40ns).mp4](http://lbfnanobiodatabase.ijs.si/file/data/cauliflowerpaper/Fig2_atomistic_molecular_dynamics_binding_anatase-101-DMPC(40ns).mp4)

*Movie S2: Time-lapse of the first 40 ns of DMPC lipids binding to anatase (101) $TiO_2$ surface*

- [http://lbfnanobiodatabase.ijs.si/file/data/cauliflowerpaper/Fig2_atomistic_molecular_dynamics_binding_anatase-101-POPE(17.5ns).mp4](http://lbfnanobiodatabase.ijs.si/file/data/cauliflowerpaper/Fig2_atomistic_molecular_dynamics_binding_anatase-101-POPE(17.5ns).mp4)

*Movie S3: Time-lapse of the first 17.5 ns of POPE lipids binding to anatase (101) $TiO_2$ surface*

#### Link to 3D

- [http://lbfnanobiodatabase.ijs.si/file/data/cauliflowerpaper/Fig2_atomistic_molecular_dynamics_binding_anatase-101-DMPC-rotation.mp4](http://lbfnanobiodatabase.ijs.si/file/data/cauliflowerpaper/Fig2_atomistic_molecular_dynamics_binding_anatase-101-DMPC-rotation.mp4)

*Movie S4:Final state of the atomistic molecular dynamics simulation of DMPC lipids binding to anatase (101) $TiO_2$ surface*

- [http://lbfnanobiodatabase.ijs.si/file/data/cauliflowerpaper/Fig2_atomistic_molecular_dynamics_binding_anatase-101-POPE-rotation.mp4](http://lbfnanobiodatabase.ijs.si/file/data/cauliflowerpaper/Fig2_atomistic_molecular_dynamics_binding_anatase-101-POPE-rotation.mp4)

*Movie S5: Final state of the atomistic molecular dynamics simulation of POPE lipids binding to anatase (101) $TiO_2$ surface*

| Atom type | Comment | q (e) | σ (Å) | ε (kJ/mol) |
|---|---|---|---|---|

| | | | | |
|---|---|---|---|---|
| Ti(O6) | Bulk Ti | 2.248 | 1.9 | 13.79 |
| Ti(O5) | Surface Ti | 2.159 | 1.9 | 13.79 |
| O(Ti3) | Oxygen in $TiO_2$ bulk | -1.124 | 3.51 | 0.409 |
| O(Ti2) | Bridge oxygen on $TiO_2$ | -1.035 | 3.42 | 0.401 |
| O(Ti,H) | Hydroxyl oxygen | -0.913 | 3.29 | 0.389 |
| H (O) | Hydrogen | 0.417 | 0 | 0 |

*Table S 9: Non-bonded force field parameters for $TiO_2$. For each atom type, coordinated atoms are given in parenthesis*

| Bond type | $b_o$ (Å) | $k_b$ (kJ/mol Å$^2$) |
|---|---|---|
| Ti-O(Ti3) bulk | 1.9 | 8000. |
| Ti-O(Ti2) bridge | 1.9 | 8000. |
| Ti-O(H) hydroxyl | 1.9 | 8000. |
| O-H hydroxyl | 1.0 | 3267. |
| Angle type | $\theta_0$ (deg) | $k_\Theta$ (kJ/mol deg$^2$) |
| Ti-O-H hydroxyl | 114.85 | 5433. |

*Table S 10: Bonded parameters for $TiO_2$*

#### *Materials and methods*

##### System composition

Atomistic molecular dynamics simulations have been carried out for DMPC and POPE lipids near anatase (101) $TiO_2$ surface in water environment. Anatase slab (71.8 x 68.2 x 30.5 Å) with (101) surface normal to the z axis is used as a model of a nanoparticle surface. The slab contains 4536 Ti atoms of which 504 are five-fold coordinated atoms on the surface. (101) anatase surface was chosen as a surface of the lowest energy. At neutral pH $TiO_2$ surface is covered by hydroxyl groups and is negatively charged. In our model we bind hydroxyl groups to 5-coordinated surface Ti atoms so that the surface charge density is close to the experimental value at neutral pH. Thus we add 151 hydroxyl groups to randomly picked Ti surface atoms (which constitutes 30% of their total amount) which results in a surface charge density of -0.62 electrons/nm$^2$, which is in line with the experimental results[8].

The $TiO_2$ slab is then placed in the middle of the simulation box with 3D periodic boundary conditions. The box size in X and Y directions is defined by the slab length and width so that the slab is periodic in those directions. The height of the box is set to 130 Å to accommodate the $TiO_2$ slab (thickness of 30.5 Å), eventual formed lipid bilayer on the both sides (2 x 40 Å) as well as their hydration layers (2 x 10 Å). 82 lipid molecules (POPE or DMPC) are inserted at random unoccupied positions in the box in random orientations, after that the box is filled with water molecules (about 12000). Then, a small number of water molecules are picked at random and are substituted with $Na^+$ and $Cl^-$ ions to balance the negative surface charge of the slab and provide NaCl concentration of 0.15 M in the water phase of the simulated system.

###### Simulation protocol

First, energy minimization of the simulated systems using the steepest gradient descent method is performed, followed by a short 100 ps pre-equilibration run at constant volume and temperature. After that, the pressure in the system is equilibrated to 1 bar using anisotropic Berendsen barostat[9] with relaxation time of 5 ps during 10 ns, which is finally followed by 1 µs production run in the NVT ensemble. Leap-frog algorithm with time step 1 fs is used to integrate the equations of motion. Center-of-mass motion is removed every 100 steps. Verlet cut-off scheme[10] with the buffer tolerance of 0.005 kJ x $mol^{-1}$ x $ps^{-1}$ per atom is used to generate the pair lists. Minimum cut-off of 1.4 nm is used for both short ranged electrostatic and VdW interactions. Long range electrostatics are calculated using PME[11] with the grid spacing of 0.12 nm and cubic interpolation. Long range dispersion corrections are applied to both energy and pressure. Velocity rescaling thermostat[12] is used to control the temperature, which is set to 303 K with the relaxation time of 1 ps. All bonds with hydrogen atoms are constrained using the LINCS algorithm[13]. Atom coordinates and energies are saved every 5 ps. All simulations were performed by the Gromacs 2019 software package[14]. Visualization of the simulations is done by VMD[15].

###### Models used

Lipids are described by the Slipids force field[16]. For $TiO_2$, we use parameters optimized to fit results on charge density distributions and water-$TiO_2$ surface coordination obtained in *ab-initio* simulations of $TiO_2$-water interface[17]. These parameters are listed in tables in supplement S5b, S5c and S5d. Water molecules are represented by the TIP3P model[18], and for $Na^+$ and $Cl^-$ ions Yoo and Aksimentiev ion parameters is used[19]. Lorentz-Berthelot rules are applied to determine Lennard-Jones parameters for cross-interactions.

#### Modelling of bilayer adhesion to $TiO_2$ surface

##### *Main message*

Anatase $TiO_2$ tube (cylinder) with radius of 10 nm cannot wrap into a non-perturbed bilayer.

Supporting raw and analysed data:

###### Figure S106

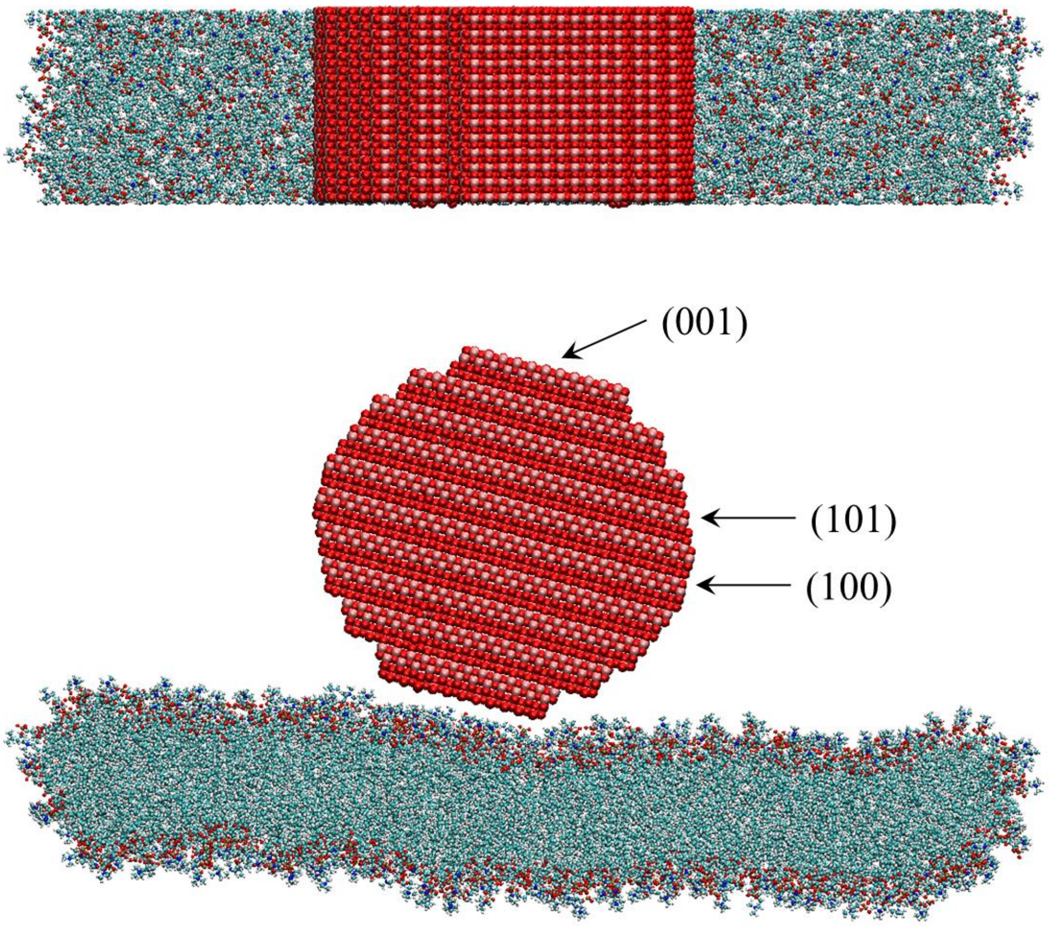

*Figure S106: Top and side views of a 10 nm cylinder undergoing wrapping by a CG bilayer in the ribbon geometry. Red arrows show the direction of restraining potentials.*

##### Materials and methods

The simulation was performed using GROMACS 2018.3 software with a 2 fs time step. Temperatures were maintained at 310 K using the Nose-Hoover thermostat with 5 ps time constant. Pressure was maintained at 1 atm using an anisotropic Parrinello-Rahman barostat with 10 ps time constant. Electrostatic interactions were calculated with a Particle Mesh Ewald (PME) summation. Lennard-Jones and real space electrostatic interaction potentials were truncated at 1.4 nm. The DOPC lipids were modeled using the fully atomistic 118 site Slipids force field which was developed for use in conjunction with TIP3P water[16,20,21]. This system was shown to reproduce many experimentally observed properties of bilayers including the lipid specific area and volume, bilayer thickness, isothermal area compressibility, and nuclear magnetic resonance (NMR) order parameters and scattering form factors. The titania force field was derived from electron densities generated by DFT simulation and was able to reproduce the DFT water density profiles at six low-energy titania cleavage planes. The axis of the cylinder lies along the [010] crystallographic direction, which is perpendicular to three of the four anatase cleavage planes with least surface energy, as determined by DFT simulation[22]. The surface thus comprises cleavage planes from the {001}, {101} and {100} families, as indicated in the diagram.

#### S2f – Ultrafast passage of nanomaterial of cell membrane

##### Main message

Nanoparticles passively (by physical interaction driven) pass exposed LA-4 membrane with only thin layer of water on top, with a 1s time-scale.

Supporting raw and analysed data:

Figure S107-Figure S111

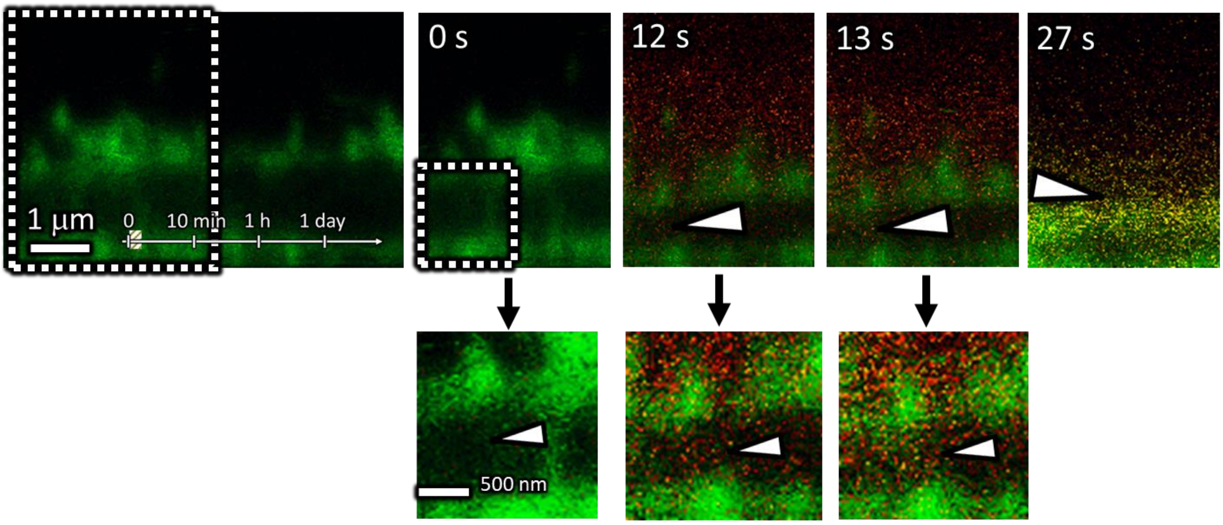

*Figure S107: Composition of three times points alongside time point 0s wth additional zoomin to region of interest. Note that the nanomaterial passes through the membrane on 1s time scale (between frame 12s and 13s). At the end of exposure a lot nanomaterial is stuck to the top of the membrane and some nanomaterial can be seen inside. See also Movie S6.*

##### Link to time-lapse

- [http://lbfnanobiodatabase.ijs.si/file/data/cauliflowerpaper/movie-NanoRain-TiO2_NT_nebulization_obto_LA-4_-_40s_duration_filtered_scaled_fliped-gif_40s.gif](http://lbfnanobiodatabase.ijs.si/file/data/cauliflowerpaper/movie-NanoRain-TiO2_NT_nebulization_obto_LA-4_-_40s_duration_filtered_scaled_fliped-gif_40s.gif)

*Movie S6: Nanoparticles (in red $TiO_2$ – Alexa 647) are nebulized directly to LA-4 epithelial cell membrane (in green – Cellmask Orange) and imaged in time with 1s time scale between frames. Nanoparticle signal was thresholded to 2 counts (there is one count noise on the detector) and membrane signal has been adjusted between frames for visibility due to bleaching of the signal. Whole movie is 40s long. See also Figure S107.*

##### Materials and methods

- experiment:
  - LA-4 cells were seeded @60% confluence in an Ibidi #1.5H µ-Dish
  - after 24 hours, cells were checked to be @ approx. 100% confluency (bright field microscope), cells were incubated with 1.6 µg/ml CellMaskOrange for 5 minutes at room 37C in the incubator, afterwards they were flushed with 3x400 µl LCIS and in 300 µL LCIS at room temperature
  - right prior exposure LCIS has been almost completely removed leaving just thin water layer on top
  - Ibidi #1.5H µ-Dish has been kept @ 37C during measurement, but not the tubing

- about 30 s after LCIS removal cells have been exposed to nebulized (sprayed on top) nanomaterial
  - 3 µl of 33 mg/ml $TiO_2$ (Alexa Fluor 647 labeled) has been added to Aeroneb®Pro nebulizer taken from VITROCELL® Cloud 6 system and mounted to standard plastic 50 ml centrifuge with a diameter of 2.75 cm. Nebulizer has been 10 cm above cell surface.
- cells have been observed 50s in total after the start of experiment

- analysis:
  - images were analyzed to present co-localization on each image separately
    - contrast on each image has been adjusted for maximal visibility due to bleaching in green channel or appearance of more signal due to nanoparticle signal rise in time in red channel – bleaching rate of CellMask™ Orange can be seen on raw images of green signal
    - contrast has been re-scaled to maximum value for each image seprately
    - contrast was adjusted for yellow color to appear where both colors have been present in one pixel above noise (noise = 1 count) – using standard gamma correction
    - bottom threshold has been set to 1 count or above in both channels to reduce noise enough for the desired features to be seen
  - top of the images has been cropped for maximal visibility of desired phenomena on paper

#### Experiment names

- Main experiment name:
  - 20180206 /e02_s01_t01_LA-4 CellMask TiO2Alexa647_after nebulization_xztSTED.msr (date and name of .msr file so it can be easily found)

#### Controls and statistics

All separate channels and overlays for all time points:

Figure S108-Figure S111

| | |
|---|---|
| Cell line | LA-4 (membrane, CellMaskOrange) |
| NPs | $TiO_2$ (Alexa647) |
| exposure | 1:1, 3s-50s (live nebulization) |
| imaging | xzt STED, 0-50s |

| | |
|---|---|
| pixelsize (x,y) | 30 nm |
| FOV (x,y) | 7.95 µm |
| pixelsize (z) | 33 nm |
| FOV (z) | 6.58 µm |
| imaging time | 50 s |
| number of frames | 50 |

| | |
|---|---|
| 561nm | 20% |
| 640nm | 20% |
| STED | 13% |
| filter sets | 605 nm – 625 nm, 650 nm – 720 nm |
| dwell time | 60 µs |
| objective | wi60x (NA1.2) |

TiO2 (Alexa647)

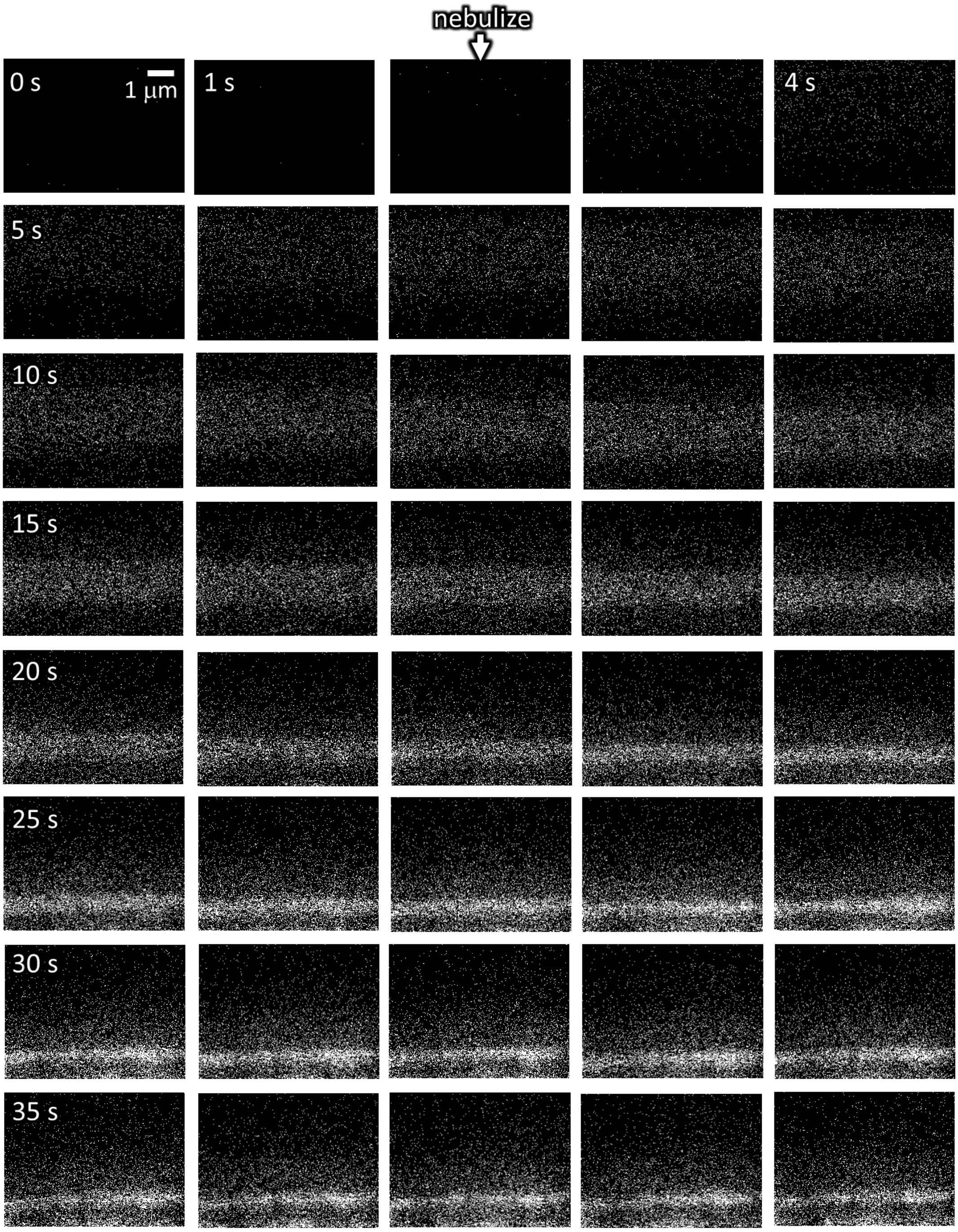

*Figure S108: Red channel-raw of 40 s time series with times step of 1s.*

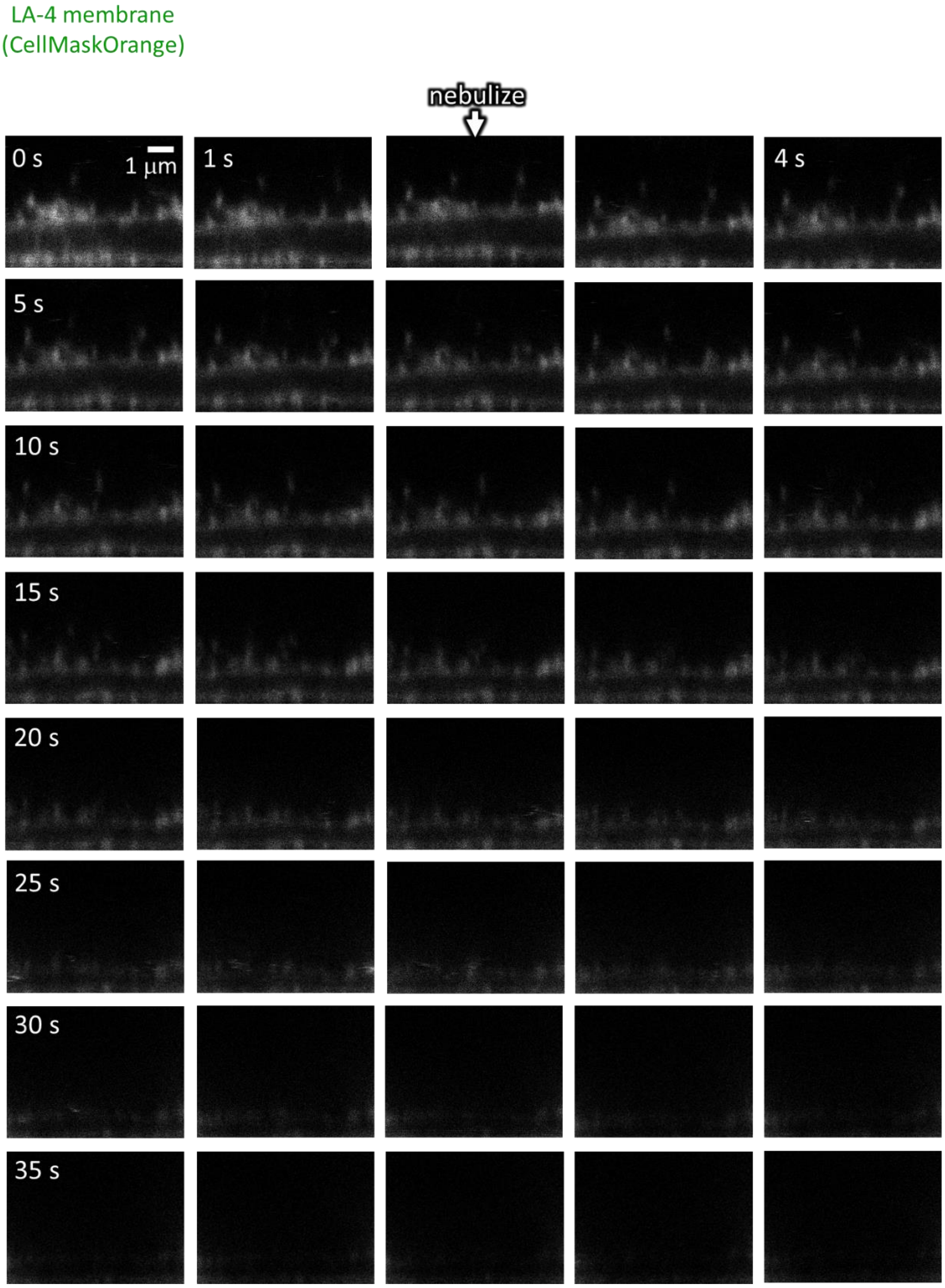

*Figure S109: Green channel-raw of 40 s time series with times step of 1s.*

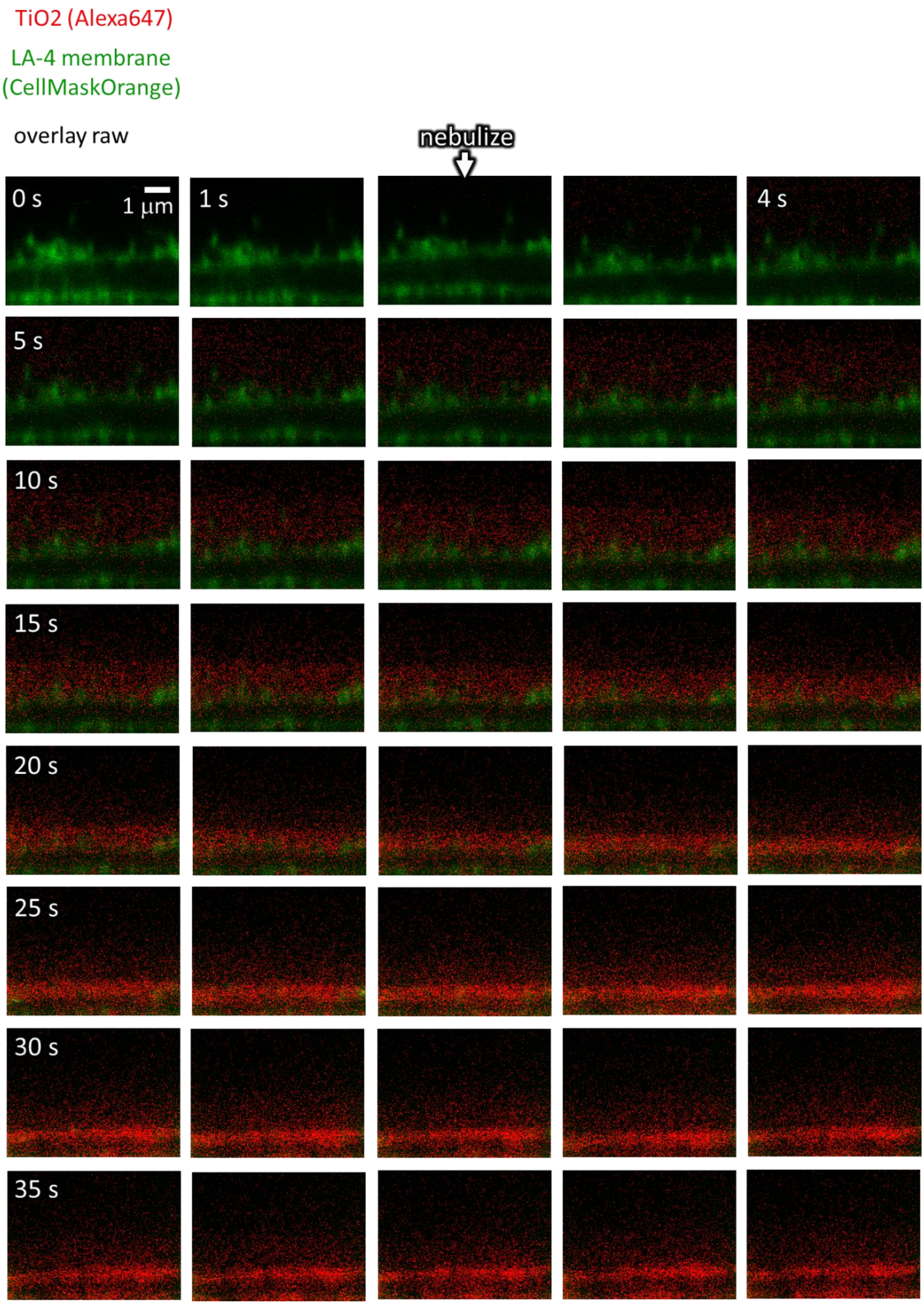

*Figure S110: Overlay-raw of 40 s time series with times step of 1s.*

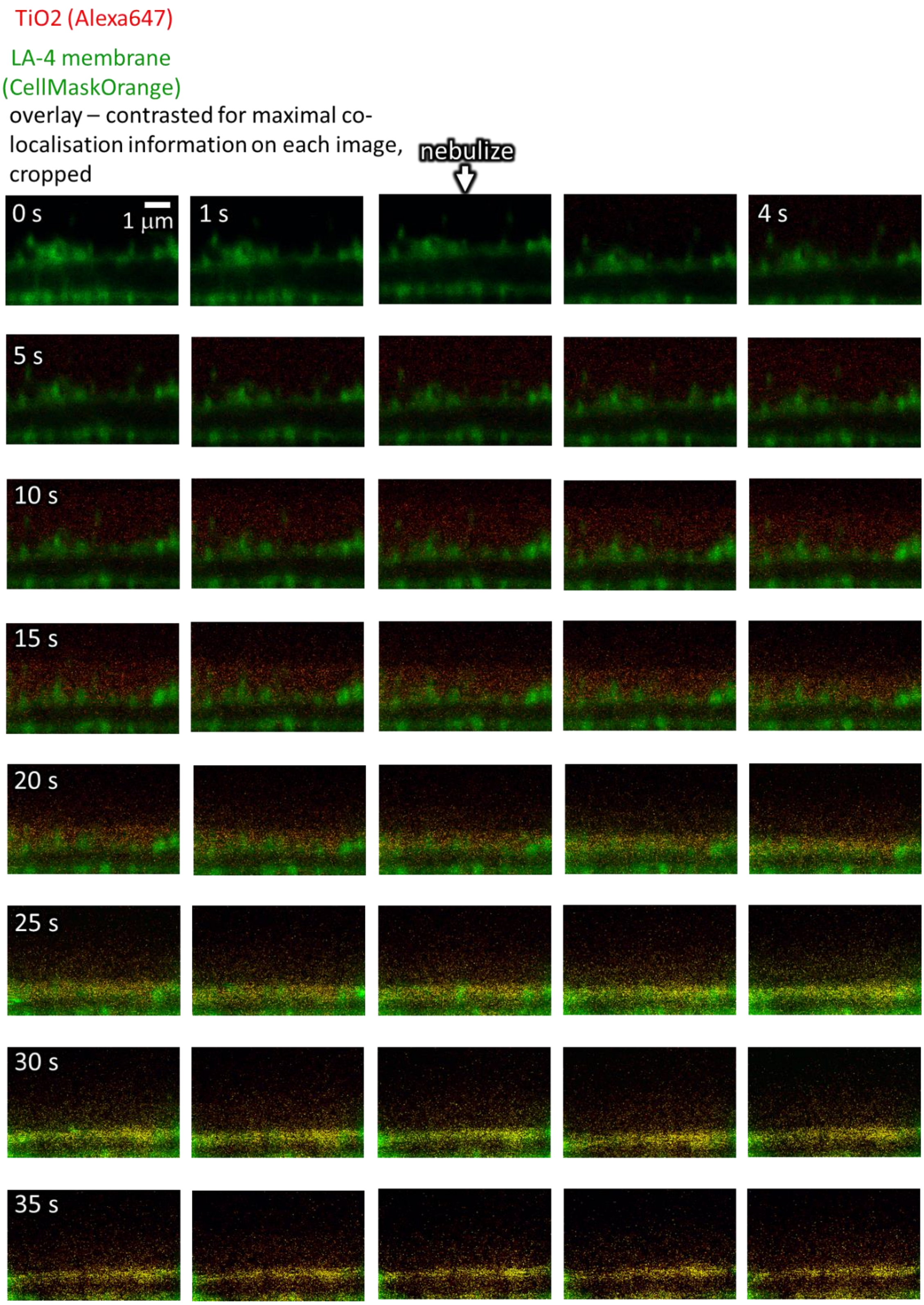

*Figure S111: Contrasted overlay of 40 s time series with times step of 1s.*

#### S2g – Blocking clathrin-mediated endocytosis

##### Main message

When blocking clathrin-mediated endocytosis using chlorpromazine, no nanomaterial enters the cell in the first 4 hours. However, at this timepoint, cauliflowers can be observed on the surface of the cell. They are smaller than fully-grown cauliflowers at day 2, but noticeably larger than the cauliflowers in control cells without blocked endocytosis.

Supporting raw and analysed data:

Figure S112-Figure S121

##### Materials and methods

- experiment **LA-4 + CellMask + $TiO_2$-Alexa 647 + 100 µm Chlorpromazine**
  - LA-4 cells were seeded @30% confluence in an Ibidi #1.5H µ-Dish.
  - After 48 hours LA-4 cells were washed with warm F12-K medium and placed on ice for 10 minutes.
  - Next, cells were washed 3 times with cold Live Cell Imaging Solution (LCIS) containing 20 mM glucose and 1% BSA .
  - 400 µL of 100 µm chlorpromazine in LCIS containing 20 mM glucose and 1% BSA was added and incubated at 37°C for 15 minutes.
  - Then 1.5 µg/mL of CellMask was added to medium.
  - After 15 minutes incubation at 37°C, 35 µL freshly filtered 1 mg/ml $TiO_2$-Alexa647 in 100x dcb was added directly to the cells and mixed to achieve 10:1 surface dose
- analysis:
  - Confocal: logarithmic scale on red channel, cut-off at 2 counts; set maximum to 350 counts on red channel; set maximum to 50 counts on green channel
  - STED: logarithmic scale on red channel, cut-off at 2 counts, set maximum to 500 counts on red channel; set maximum to 50 counts on green channel

| | |
|---|---|
| pixelsize (x,y) | 50 nm |
| FOV (x,y) | 100 µm |
| pixelsize (z) | 50 nm |
| FOV (z) | 30 µm |

| | |
|---|---|
| 561nm | 30% |
| 640nm | 20% |
| STED | - |
| filter sets | 605 nm – 625 nm, 650 nm – 720 nm |
| dwell time | 10 µs |
| objective | wi 60x (NA1.1) |

| | |
|---|---|
| Cell line | LA-4 (membrane, CellMask) |
| NPs | TiO2 (Alexa647) |
| exposure | 10:1, 0h-4h |
| imaging | 0min-5h xyz confocal, h |

| | |
|---|---|
| pixelsize (x,y) | 30 nm |
| FOV (x,y) | 26x33 µm |
| pixelsize (z) | |
| FOV (z) | |
| imaging time | - |
| number of frames | - |

| | |
|---|---|
| 561nm | 40% |
| 640nm | 20% |
| STED | 15% |
| filter sets | 605 nm – 625 nm, 650 nm – 720 nm |
| dwell time | 10 µs |
| objective | wi 60x (NA1.1) |

- experiment **LA-4 + DPPE Star580 + $TiO_2$-Alexa 647 + 200 µm Chlorpromazine**:
  - LA-4 cells were seeded @30% confluence in an Ibidi #1.5H µ-Dish

- After 48 hours LA-4 cells were washed with warm F12-K medium and placed on ice for 10 minutes.
    - Next cells were washed 3 times with cold Live Cell Imaging Solution (LCIS) containing 20 mM glucose and 1% BSA .
    - 400 µL of 200 um chlorpromazine in LCIS containing 20 mM glucose and 1% BSA was added and incubated at 37°C for 15 minutes.
    - Then 4 µL of 1mM DPPE-Star580 was added to medium
    - After 15 minutes incubation at 37°C, 35 µL freshly filtered 1 mg/ml $TiO_2$-Alexa647 in 100x dcb was added directly to the cells and mixed to achieve a 10:1 surface dose
- analysis:
    - Confocal: logarithmic scale on red channel, cut-off at 3 counts; set maximum to 300 counts on both channels
    - STED: logarithmic scale on red channel, cut-off at 3 counts; set maximum to 175 counts on both channels

#### Link to time-lapse

- http://lbfnanobiodatabase.ijs.si/file/data/cauliflowerpaper/20190613_e01_s02_t01_LA-4_1uM_DPPEstar580_200uM_chlorpromazine_TiO2Alexa_1to10_1s_is_20min.gif

*Movie S7: First 4 hours following exposure of LA-4 (membranes labelled with DPPEstar580, green) with chlorpromazine-blocked clathrin-mediated endocytosis to 10:1 TiO2 (Alexa 647, red), first 4 hours following exposure. 1 second in the movie corresponds to 20 minutes in real time. See also Figure S113.*

- http://lbfnanobiodatabase.ijs.si/file/data/cauliflowerpaper/20190613_e02_s01_t01_LA-4_1uM_DPPEstar580_200uM_chlorpromazine_TiO2Alexa_1to10_1s_is_20min.gif

*Movie S8: First 3.5 hours following exposure of LA-4 (membranes labelled with DPPEstar580, green) with chlorpromazine-blocked clathrin-mediated endocytosis to 10:1 TiO2 (Alexa 647, red). 1 second in the movie corresponds to 20 minutes in real time. See also Figure S118.*

#### Link to 3D

- http://lbfnanobiodatabase.ijs.si/file/data/cauliflowerpaper/20190613_e01_s02_t04_LA-4_1uM_DPPEstar580_200uM_chlorpromazine_TiO2Alexa_10to1_4h_xyz_FOV_100_x_100_x_30_um.gif

*Movie S9: Final state after 4 hours of exposure of LA-4 (membranes labelled with Star580 DPPE, green) with chlorpromazine-blocked clathrin-mediated endocytosis to 10:1 TiO2 (Alexa 647, red). Field-of-view is 100 x 100 x 30 µm. See also Figure S113.*

- http://lbfnanobiodatabase.ijs.si/file/data/cauliflowerpaper/20190613_e02_s01_t03_LA-4_1uM_DPPEstar580_200uM_chlorpromazine_TiO2Alexa_10to1_3.5h_xyz_FOV_100_x_100_x_30_um.mp4

*Movie S10: Final state after 3.5 hours of exposure of LA-4 (membranes labelled with Star580 DPPE, green) with chlorpromazine-blocked clathrin-mediated endocytosis to 10:1 TiO2 (Alexa 647, red). Field-of-view is 100 x 100 x 30 µm. See also Figure S118.*

| | |
|---|---|
| pixelsize (x,y) | 50 nm |
| FOV (x,y) | 100 µm |
| pixelsize (z) | 50 nm |
| FOV (z) | 30 µm |

| | |
|---|---|
| 561nm | 30% |
| 640nm | 30% |
| STED | - |
| filter sets | 605 nm – 625 nm,<br>650 nm – 720 nm |
| dwell time | 10 µs |
| objective | wi 60x (NA0.3) |

| | |
|---|---|
| Cell line | LA-4 (membrane, DPPE-Star580) |
| NPs | TiO2 (Alexa647) |
| exposure | 10:1, 0h-4h |
| imaging | 0min-4h<br>xyz confocal, 4h |

| | |
|---|---|
| pixelsize (x,y) | 20 nm |
| FOV (x,y) | 50 µm |
| pixelsize (z) | |
| FOV (z) | |
| imaging time | 3.5h |
| number of frames | 150 |

| | |
|---|---|
| 561nm | 20% |
| 640nm | 30% |
| STED | 15% |
| filter sets | 605 nm – 625 nm,<br>650 nm – 720 nm |
| dwell time | 10 µs |
| objective | wi 60x (NA1.1) |

#### Experiment names

- Main experiment name:
  - e02_s03_t01_LA-4_1,5ug_ml_CellMask_100um_chlorpromazine_TiO2Alexa_1to10_4h

- Experiment names:

  *CellMask:*
  - e02_s01_t01_LA-4_1,5ug_ml_CellMask_100um_chlorpromazine_TiO2Alexa_1to10_3D
  - e02_s02_t01_LA-4_1,5ug_ml_CellMask_100um_chlorpromazine_TiO2Alexa_1to10_3h
  - e02_s03_t01_LA-4_1,5ug_ml_CellMask_100um_chlorpromazine_TiO2Alexa_1to10_4h
  - e02_s04_t01_LA-4_1,5ug_ml_CellMask_100um_chlorpromazine_TiO2Alexa_1to10_5h
  - e02_s01_t02_LA-4_1,5ug_ml_CellMask_100um_chlorpromazine_TiO2Alexa_1to10_timeLapse
  - 20190611/e03_s01_t01_LA-4_1,5ug_ml_CellMask_200uM_chlorpromazine_TiO2Alexa_1to10

  *DPPE-Star580:*
  - 20190613/e01_s02_t04_LA-4_1um_DPPEstar580_200um_chlorpromazine_TiO2Alexa_1to10_3D_4h.msr
  - 20190613/e01_s02_t02_LA-4_1um_DPPEstar580_200um_chlorpromazine_TiO2Alexa_1to10_1.5h.msr
  - 20190613/e01_s02_t03_LA-4_1um_DPPEstar580_200um_chlorpromazine_TiO2Alexa_1to10_3h.msr
  - 20190613/e01_s02_t01_LA-4_1um_DPPEstar580_200um_chlorpromazine_TiO2Alexa_1to10_0.5h.msr
  - 20190613/e02_s01_t01_LA-4_1um_DPPEstar580_200um_chlorpromazine_TiO2Alexa_1to10_0.5h
  - 20190613/e02_s01_t02_LA-4_1um_DPPEstar580_200um_chlorpromazine_TiO2Alexa_1to10_3.5h_3D

- 20190613/e02_s01_t03_LA-4_1um_DPPEstar580_200um_chlorpromazine_TiO2Alexa_1to10_3.5h_3D

*Controls:*

- 20190618/e01_s01_t01_LA-4_1um_DPPEstar580_200um_chlorpromazine_0.5h_3D
- 20190121/e01_s01_t01_LA-4_50ug_ml_pHrodo_200um_chlorpromazine_TiO2-Alexa647_steps_30minInk
- 20190121/e01_s02_t02_LA-4_50ug_ml_pHrodo_200um_chlorpromazine_TiO2-Alexa647_steps_30minInk
- 20190611/e01_s01_t02_LA-4_1ug_ml_CellMask_100uM_chlorpromazine_test3hInk3D

#### Controls and statistics

Time-course of cauliflower growth in cells with inhibited endocytosis

Figure S112-Figure S118

control – cells with blocked endocytosis, not exposed to nanomaterial

Figure S119 - Figure S120

control – colocalisation of $TiO_2$ with endosome probe and chlorpromazine

Figure S121

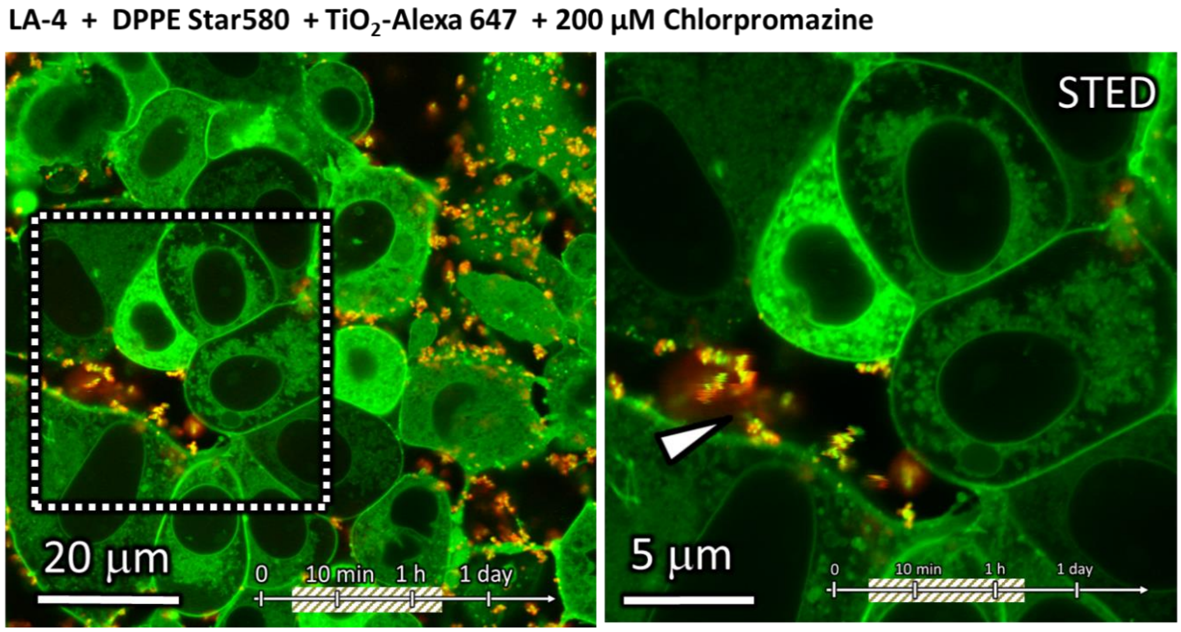

*Figure S112: Cauliflower growth in cells with inhibited endocytosis (time-point 2.5 hours)*

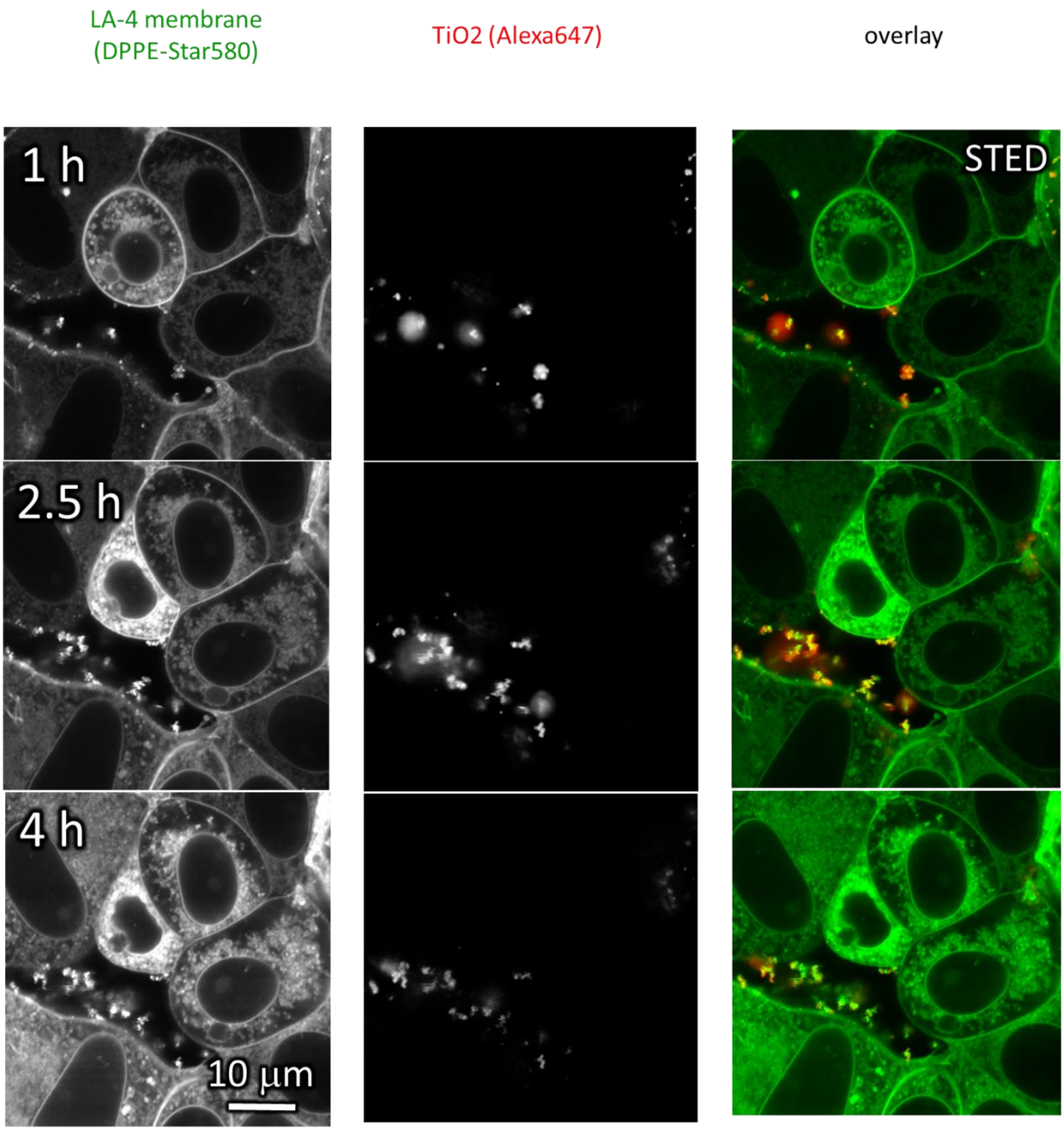

*Figure S113: Time-course of cauliflower growth in cells with inhibited endocytosis – xy cross-sections. See also Movie S7 and Movie S9*

LA-4 + CellMask + $TiO_2$-Alexa 647 + 100 µM Chlorpromazine

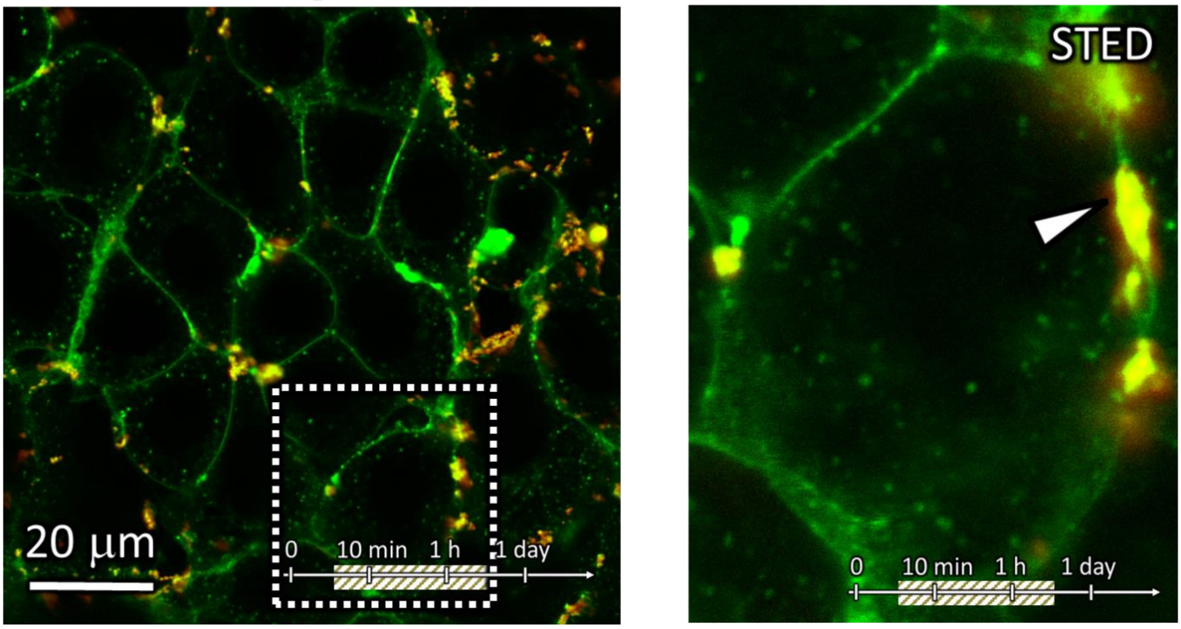

*Figure S114: Cauliflower growth in cells with inhibited endocytosis (time-point 5 hours)*

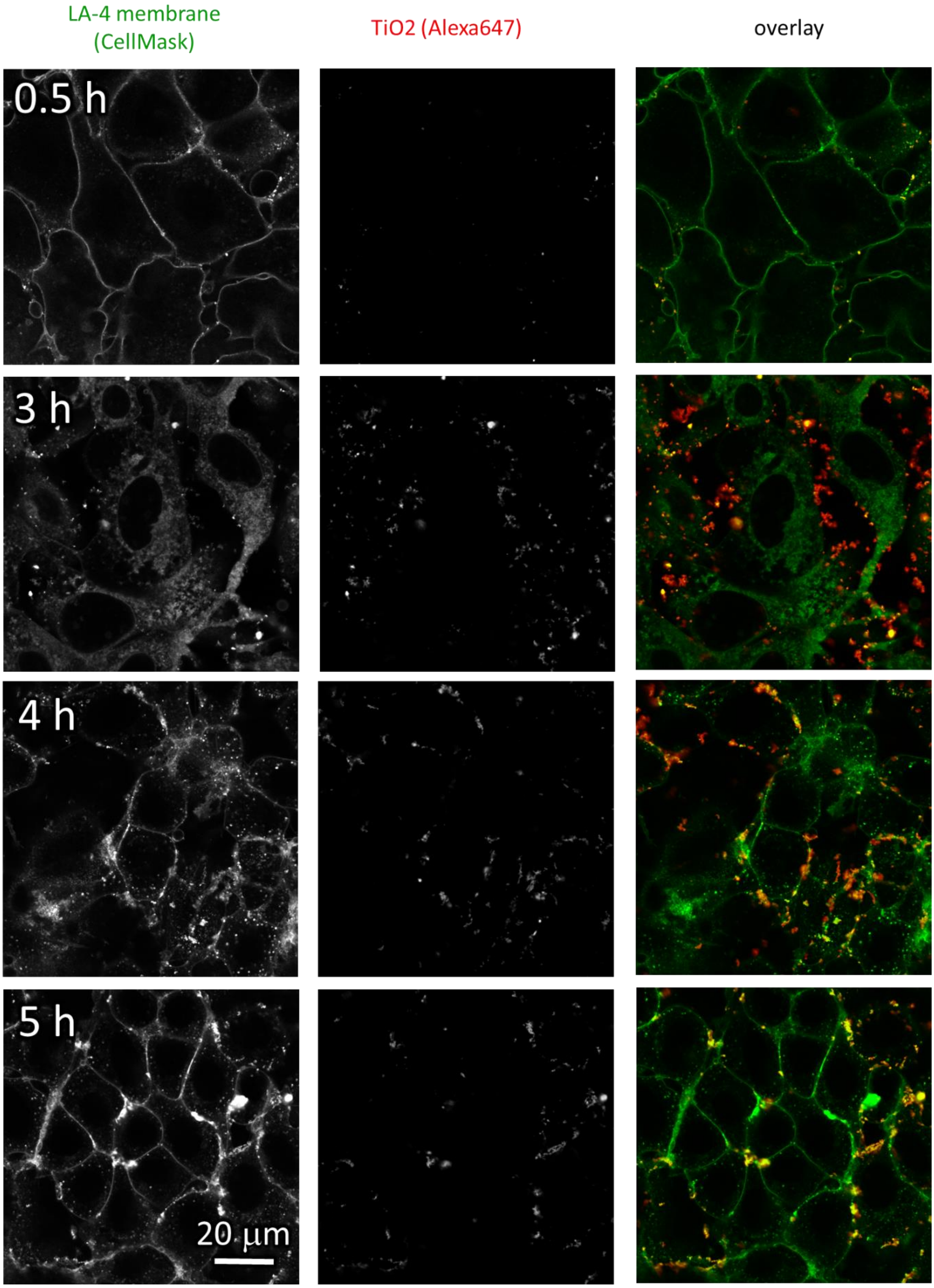

*Figure S115: Time-course of cauliflower growth in cells with inhibited endocytosis – xy cross-sections.*

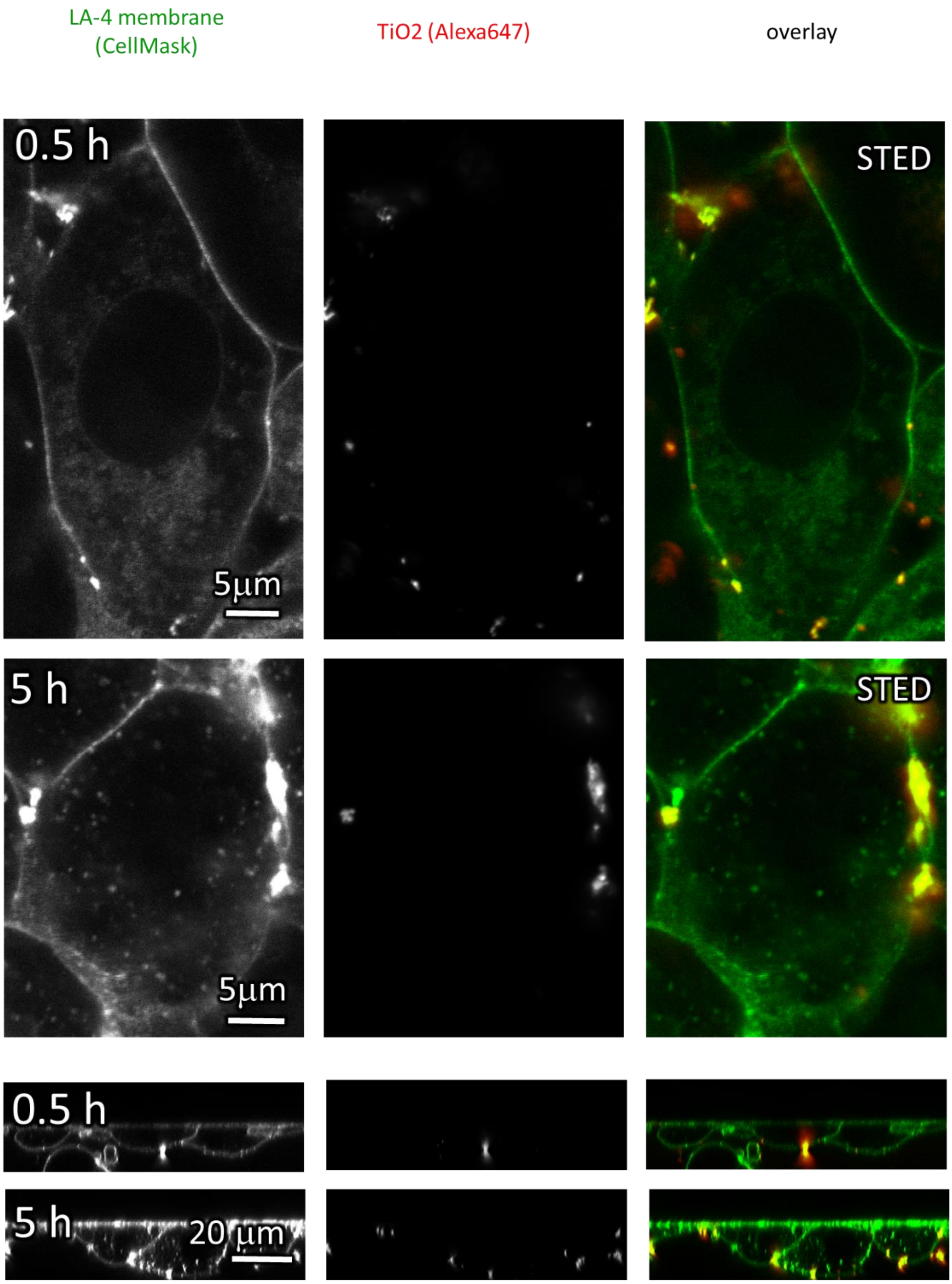

*Figure S116: Time-course of cauliflower growth in cells with inhibited endocytosis – xy and xz cross-sections.*

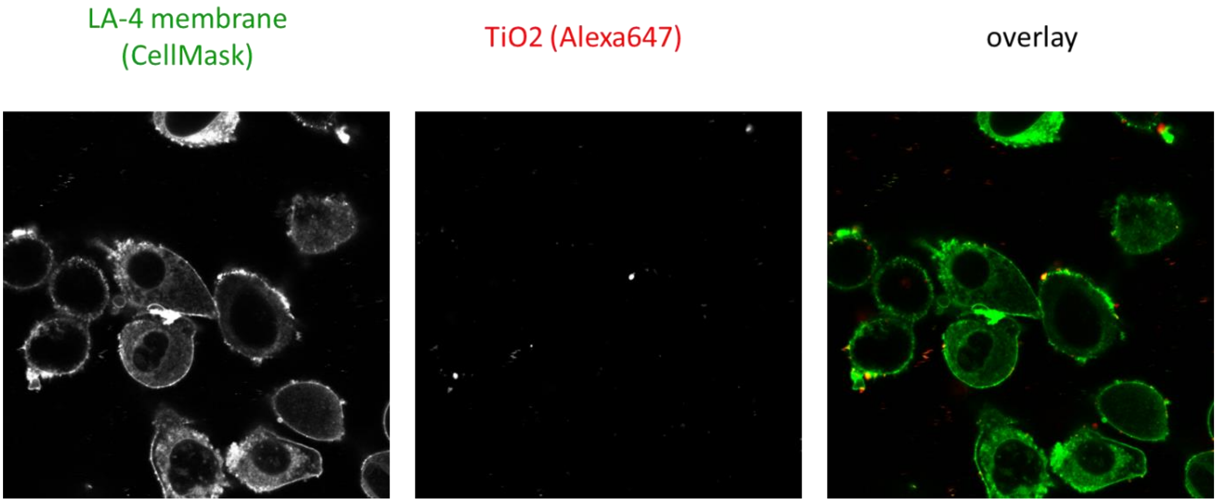

*Figure S117: Cauliflower growth in cells with inhibited endocytosis (time-point 0.5 hours).*

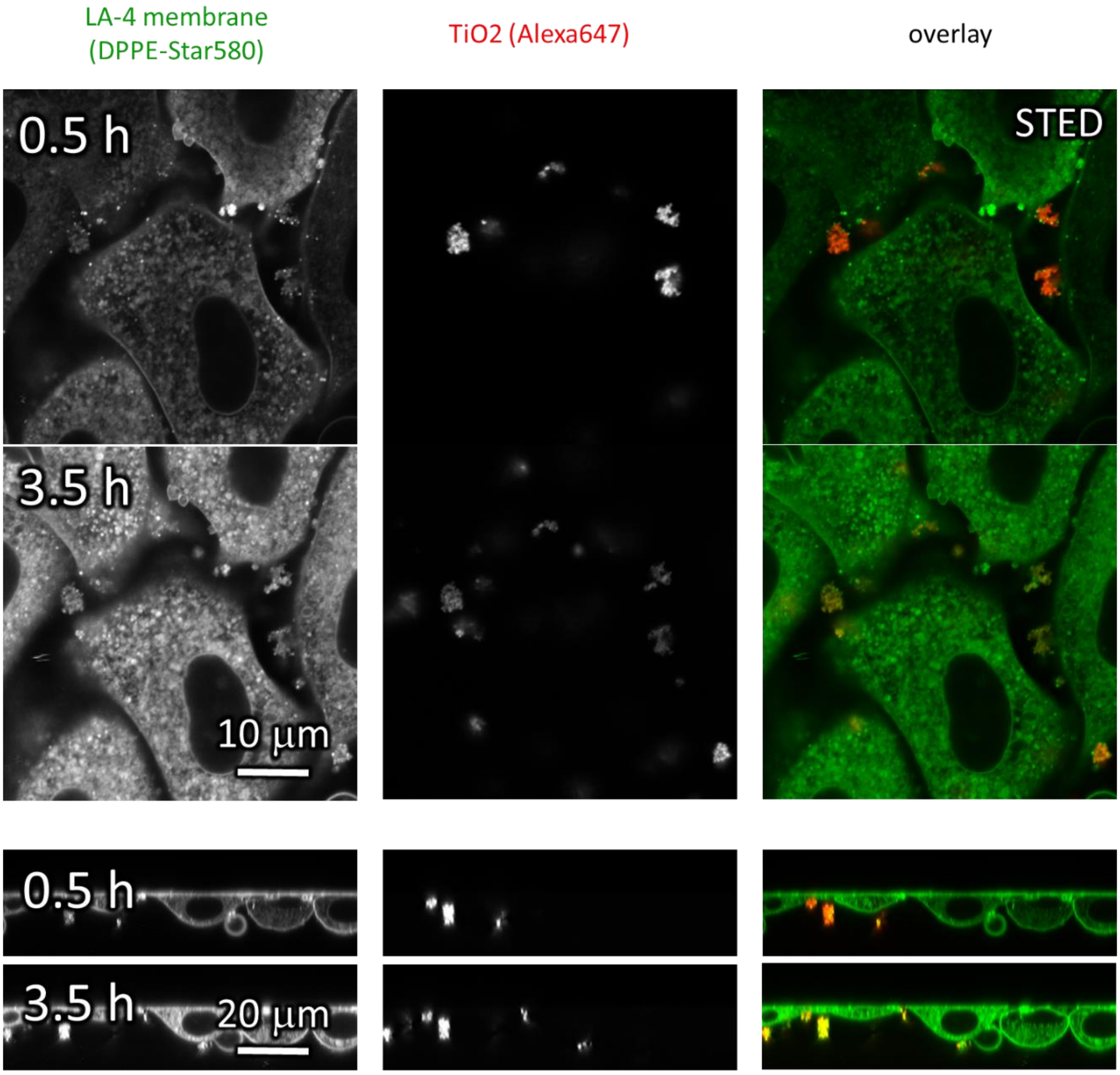

*Figure S118: Time-course of cauliflower growth in cells with inhibited endocytosis – xy and xz cross-sections. See also Movie S8 and Movie S10.*

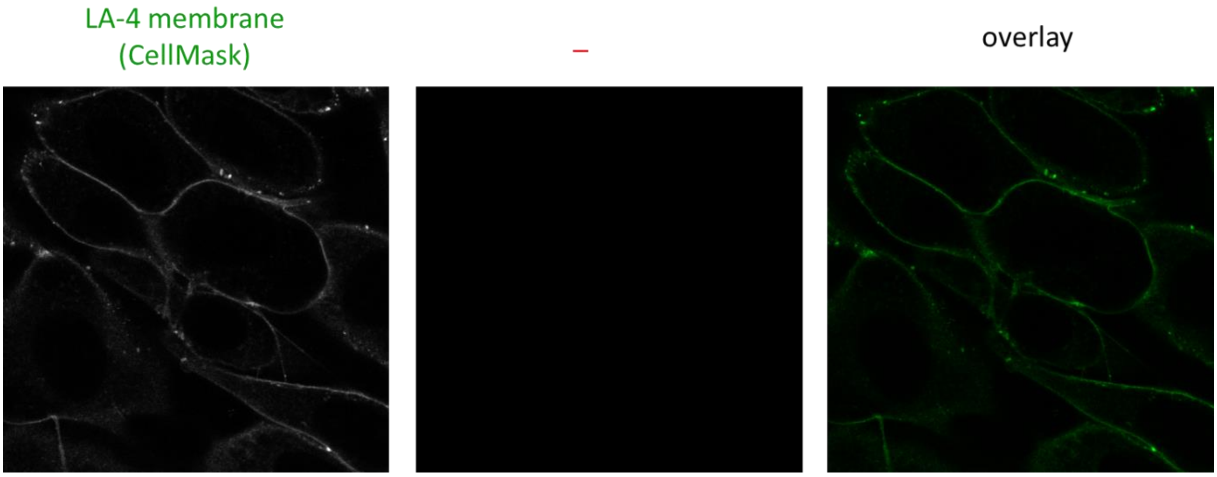

*Figure S119: Control –labelled cells with blocked endocytosis, not exposed to nanomaterial.*

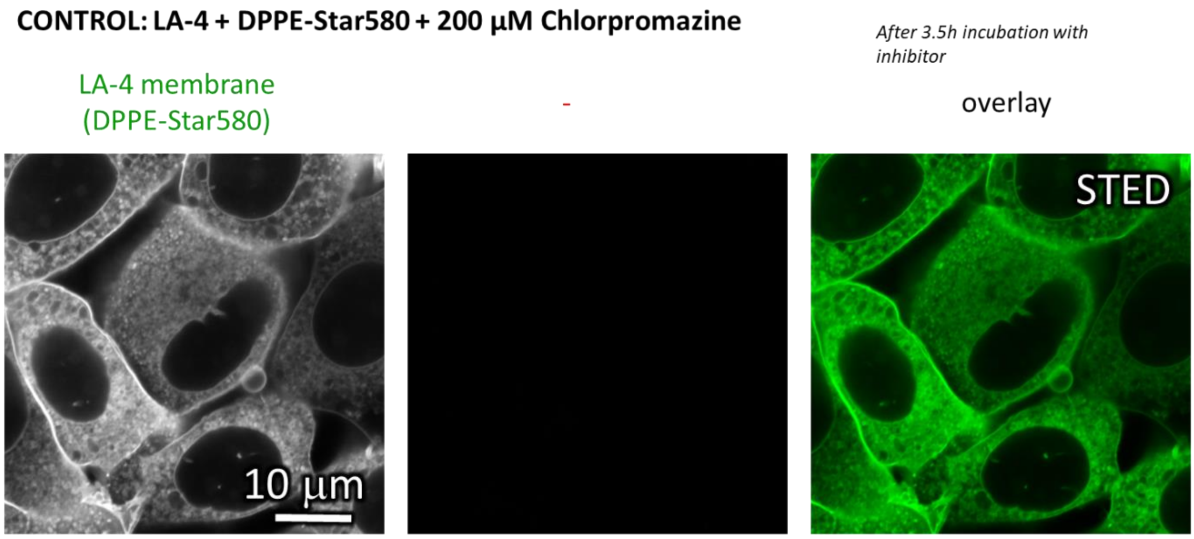

*Figure S120: Control –labelled cells with blocked endocytosis, not exposed to nanomaterial.*

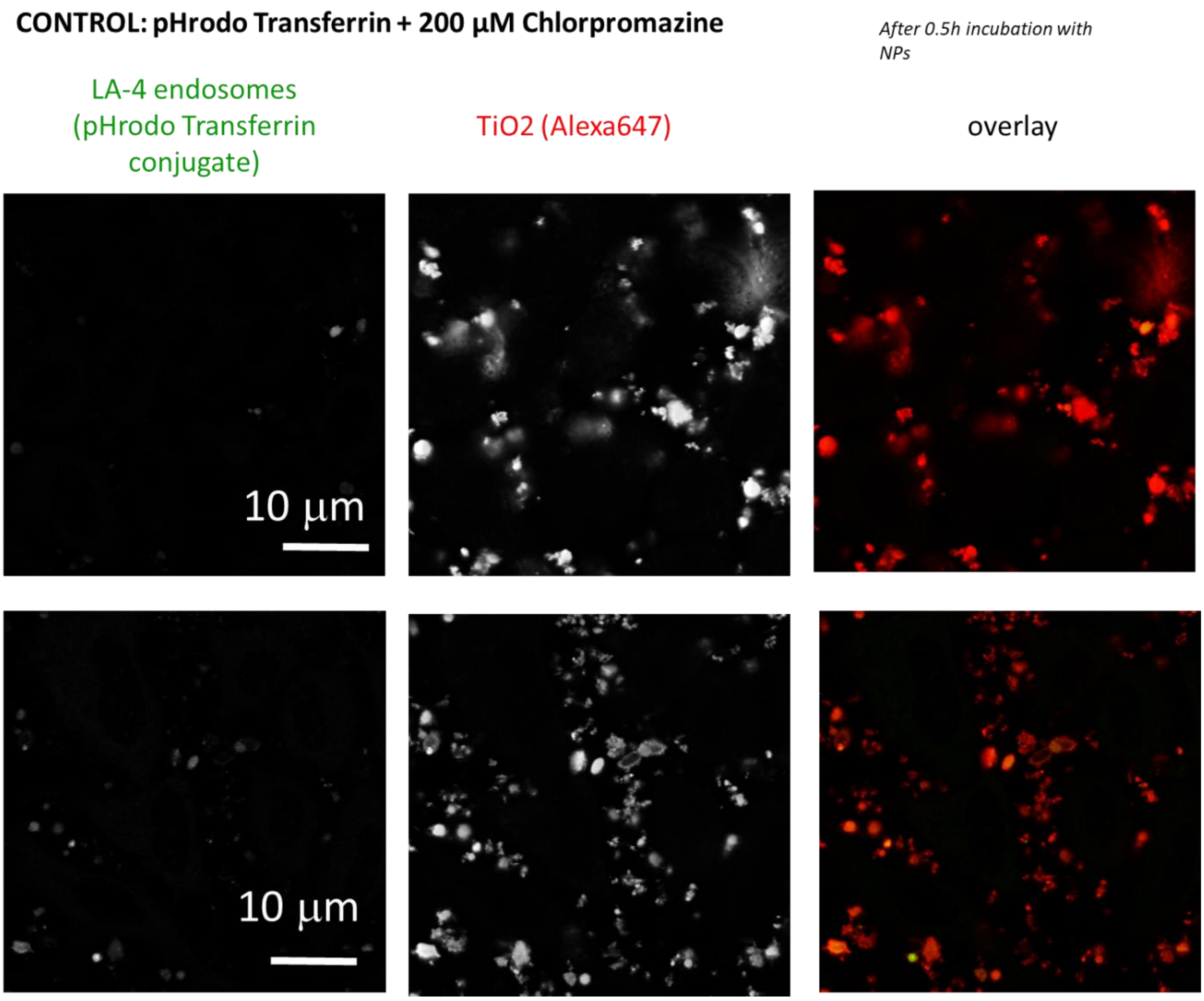

*Figure S121: Control – colocalisation of labelled $TiO_2$ with endosome probe (pHrodo Transferrin) when incubated with chlorpromazine.*

#### S2h – Membrane cholesterol extraction with Methyl-Beta-Cyclodextran

##### Main message

Observing formation of cauliflower-like structures after the extraction of cholesterol from plasma membrane with Metyl – Beta – Cyclodextrin.

Supporting raw and analysed data:

Figure S122-Figure S125

##### Materials and methods

- experiment:
    - LA-4 cells were seeded @60% confluence in an Ibidi #1.5H µ-Dish
    - after 24 hours medium was exchanged for fresh mixture of LCIS and medium with 1 mM Metyl-Beta-Cyclodextran (MBCD). Immediately after 35 µL of freshly filtered 1 mg/mL $TiO_2$-40-ATTO 594 in 100x dcb was added directly to the cells (in 400 µL medium with 1 mM MBCD) and mixed to achieve 10:1 surface dose. Cells were incubated for additional 24h
    - After 48 hours cells were incubated with 1 um Star Red DPPE for 5 min in incubator at 37°C, 5% CO2. Cells were not washed with LCIS in order to not wash away MBCD. Cells were imaged on the microscope stage heated on 37°C.
- analysis:
    - Sharpening algorithm (σ = 2, c = 0,3; described below) was applied in red channel and 1 pixel Gaussian blur in green
    - Contrast is adjusted to achieve better visibility

    To reduce the noise and accentuate fine details on some images we used a sharpening algorithm for subtraction of a blurred image (Gaussian low-past filter) from the original image. This can be considered as a convolution operation on an image with a kernel mask that is a two-dimensional Gaussian function:

$$g(x,y) = \frac{1}{\sigma\sqrt{2\pi}} e^{-(x^2+y^2)/2\sigma^2}$$

    Where σ represents the size of the Gaussian kernel mask which determines the range of frequencies removed by the Gaussian filter. This blurred image is then subtracted from the original image as:

$$F(x,y) = \frac{c}{2c-1} I(x,y) - \frac{(1-c)}{2c-1} U(x,y)$$

    where the F(x, y) represents the brightness value of a pixel at the coordinate (x, y) in the filtered image, and I(x, y) and U(x, y) represent the brightness values of the corresponding pixels in the original and blurred images, respectively. The constant c controls the relative weightings of the original and blurred images in the difference equation.
    Such a subtraction enhances high-frequency spatial detail at the expense of low-frequency spatial information in the image.

#### Experiment names

- Main experiment name:
  - 20190612_e01 m02 s01_LA-4_SR DPPE_TiO2 Star520S_1 mM MBCD_overnight incubation_NM in the cell

- Supplement Experiment names:
  - 20190612_e03 m03 s01 _LA-4_SR DPPE_TiO2 Star520S_1 mM MBCD_overnight incubation_uptake of TiO2
  - 20190612_e01 m04 s02 LA-4_SR DPPE TiO2 Star 520 0.5mM MBCD overnight much NM in the cell
  - 20190612_e01 m02 s01_LA-4_SR DPPE_TiO2 Star520S_1 mM MBCD_overnight incubation_NM in the cell_STED
  - 20190612_e01 m03 s01 LA-4_SR DPPE TiO2 Star 520 0.5 mM MBCD overnight much NM in the cell_3D.msr

#### Controls and statistics

Shortly after treatment with 1 mM MBCD cells show increased uptake of single and small aggregates of nanomaterial. We see less large cauliflower structures on the surface of the cells:

Figure S122-Figure S124

Four time points from a time lapse movies of the cell rapidly uptaking $TiO_2$ nanotubes:

Figure S125

Main Experiment

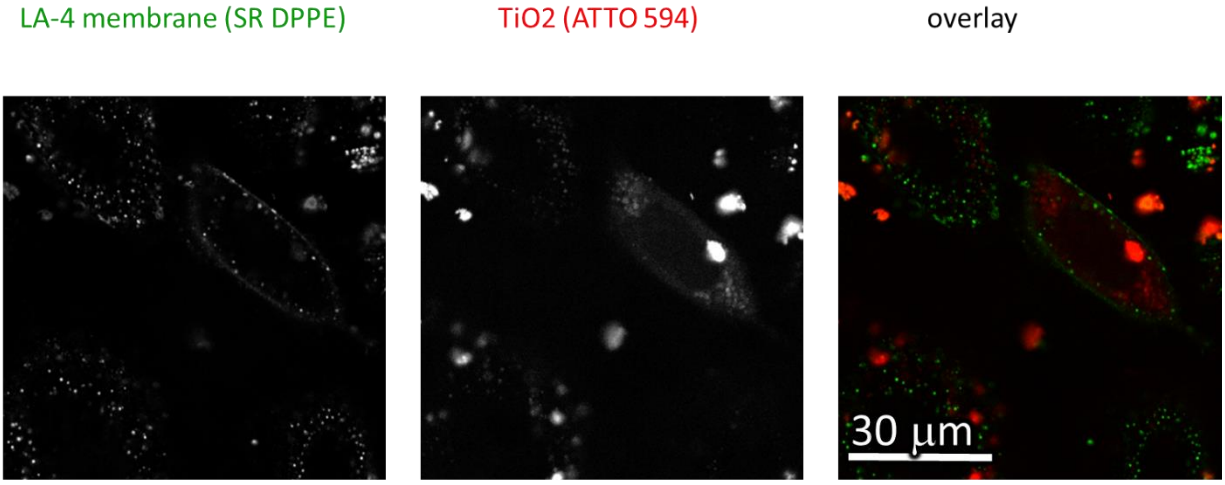

| | | | | | |
|---|---|---|---|---|---|
| Cell line | LA-4 (membrane, StarREd-DPPE) | pixelsize (x,y) | 200 | 561nm | 30 % |
| NPs | TiO2 (ATTO 594) | FOV (x,y) | 80 µm | 640nm | 40 % |
| exposure | 10:1, 24h-48h 1 mM MBCD | pixelsize (z) | / nm | STED | / % |
| imaging | xy confocal, 48h | FOV (z) | / µm | filter sets | 605 nm – 625 nm, 650 nm – 720 nm |
| | | imaging time | / | dwell time | 10 µs |
| | | number of frames | / | objective | 6wi0x (NA1.2) |

*Figure S122: Confocal images of LA-4 in green and $TiO_2$ nanotubes in red channel. Cells treated with 1 mM MBCD for 48 hours. No collocalization and no cauliflower-like formations. Cells have been stained with Star Red-DPPE dye which tends to redistribute inside the cell into membranes of different vesicles. For that reason we can't see clear outlines of all the cells. Note how cells are full of individual nanotubes.*

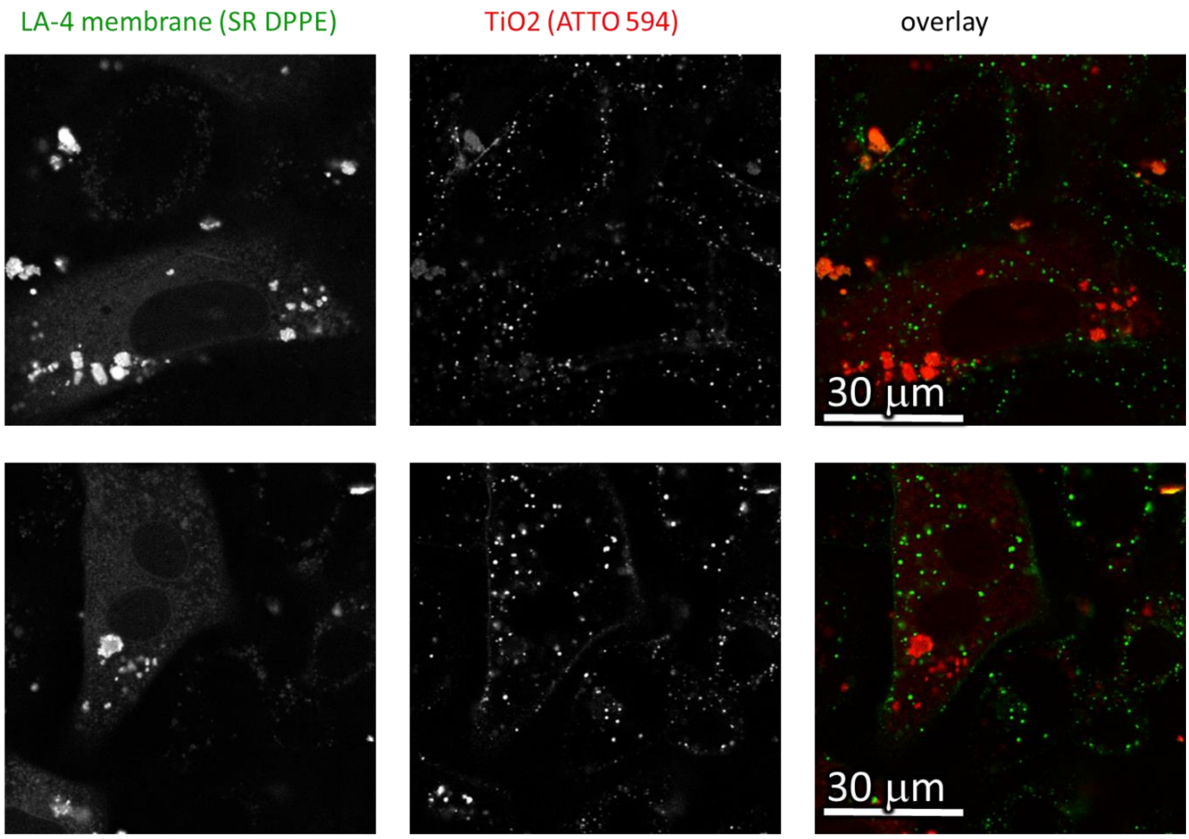

| | |
|---|---|
| Cell line | LA-4 (membrane, StarREd-DPPE) |
| NPs | TiO2 (ATTO 594) |
| exposure | 10:1, 24h-48h<br>1 mM MBCD |
| imaging | xyt STED /,<br>xy confocal, 48h |

| | |
|---|---|
| pixelsize (x,y) | 200 |
| FOV (x,y) | 80 µm |
| pixelsize (z) | / nm |
| FOV (z) | / µm |
| imaging time | / |
| number of frames | / |

| | |
|---|---|
| 561nm | 30 % |
| 640nm | 40 % |
| STED | / % |
| filter sets | 605 nm – 625 nm,<br>650 nm – 720 nm |
| dwell time | 10 µs |
| objective | 6wi0x (NA1.2) |

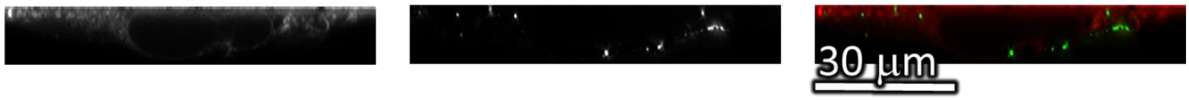

| | |
|---|---|
| Cell line | LA-4 (membrane, StarREd-DPPE) |
| NPs | TiO2 (ATTO 594) |
| exposure | 10:1, 24h-48h<br>1 mM MBCD |
| imaging | xz STED 48h,<br>xz confocal / |

| | |
|---|---|
| pixelsize (x,y) | 40 |
| FOV (x,y) | 80 µm |
| pixelsize (z) | 40 nm |
| FOV (z) | 12 µm |
| imaging time | / |
| number of frames | / |

| | |
|---|---|
| 561nm | 30 % |
| 640nm | 30 % |
| STED | 20 % |
| filter sets | 605 nm – 625 nm,<br>650 nm – 720 nm |
| dwell time | 10 µs |
| objective | 6wi0x (NA1.2) |

*Figure S123: Confocal images of LA-4 in green and $TiO_2$ nanotubes in red channel. Cells treated with 1 mM MBCD for 48 hours. No collocalization and no cauliflower-like formations. Cells in first two windows are XY shots of different spots on the sample. In 3rd window we see STED of the XZ plane of the large cell in the first window. Note how cells are full of individual nanotubes.*

LA-4 membrane (SR DPPE) TiO2 (ATTO 594) overlay

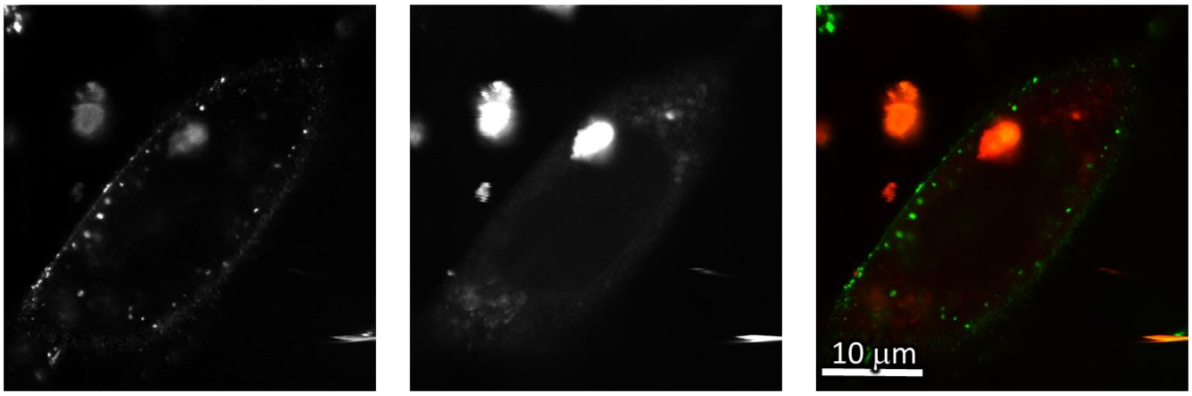

| | |
|---|---|
| Cell line | LA-4 (membrane, StarRed-DPPE) |
| NPs | TiO2 (ATTO 594) |
| exposure | 10:1, 24h-48h |
| imaging | xy STED, 48h |

| | |
|---|---|
| pixelsize (x,y) | 30 |
| FOV (x,y) | 39 x 40,4 µm |
| pixelsize (z) | / nm |
| FOV (z) | / µm |
| imaging time | / |
| number of frames | / |

| | |
|---|---|
| 561nm | 30 % |
| 640nm | 30 % |
| STED | 20 % |
| filter sets | 605 nm – 625 nm, 650 nm – 720 nm |
| dwell time | 10 µs |
| objective | 6wi0x (NA1.2) |

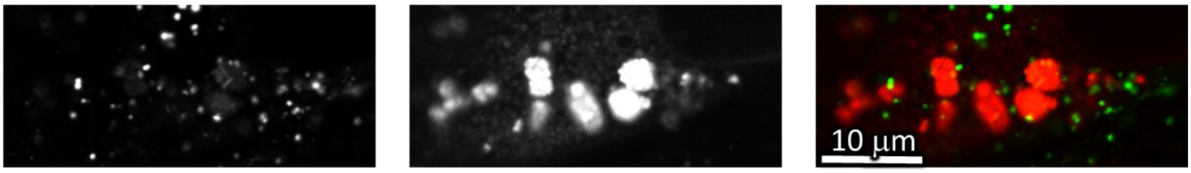

| | |
|---|---|
| Cell line | LA-4 (membrane, StarRed-DPPE) |
| NPs | TiO2 (ATTO 594) |
| exposure | 10:1, 24h-48h |
| imaging | xy confocal, 48h |

| | |
|---|---|
| pixelsize (x,y) | 40 |
| FOV (x,y) | 37 x 16 µm |
| pixelsize (z) | / nm |
| FOV (z) | / µm |
| imaging time | / |
| number of frames | / |

| | |
|---|---|
| 561nm | 30 % |
| 640nm | 30 % |
| STED | / % |
| filter sets | 605 nm – 625 nm, 650 nm – 720 nm |
| dwell time | 10 µs |
| objective | 6wi0x (NA1.2) |

*Figure S124: Confocal images of LA-4 in green and $TiO_2$ nanotubes in red channel. Cells treated with 1 mM MBCD for 48 hours. No collocalization and no cauliflower-like formations. Confocal images of a XY plane in the 1st window and XZ plane in the 2nd rst window. Note how cells are full of individual nanotubes. See also Movie S11.*

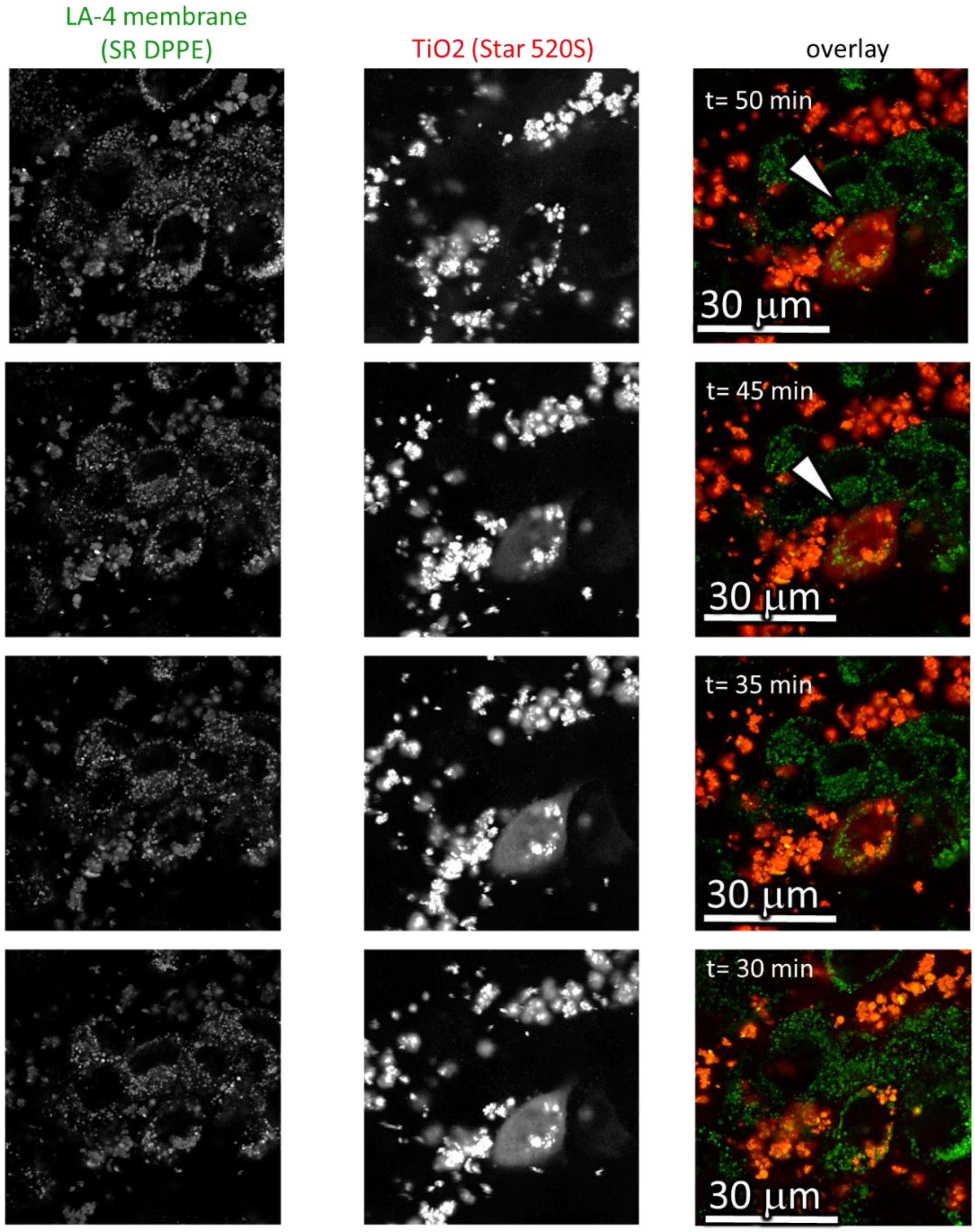

| | |
|---|---|
| Cell line | LA-4 (membrane, StarREd-DPPE) |
| NPs | TiO2 (ATTO 594) |
| exposure | 10:1, 24h-48h |
| imaging | xy confocal, 48h |

| | |
|---|---|
| pixelsize (x,y) | 80 |
| FOV (x,y) | 80 µm |
| pixelsize (z) | / nm |
| FOV (z) | / µm |
| imaging time | 1h |
| number of frames | 61 |

| | |
|---|---|
| 561nm | 30 % |
| 640nm | 40 % |
| STED | / % |
| filter sets | 605 nm – 625 nm, 650 nm – 720 nm |
| dwell time | 10 µs |
| objective | 6wi0x (NA1.2) |

*Figure S125 Confocal images of LA-4 in green and $TiO_2$ nanotubes in red channel. Cells treated with 1 mM MBCD for 48 hours. No collocalization and no cauliflower-like formations. Confocal images of XY planes of different time points in a time lapse video. Filling of the cell with nanotubes in course of 20 min. See also Movie S12.*

Link to 3D

- http://lbfnanobiodatabase.ijs.si/file/data/cauliflowerpaper/20190612_e01_m03_s01__LA-4_SR_DPPE_TiO2_Star_520_0.5mM_MBCD_overnight_much_NM_in_the_cell__3D.mp4

*Movie S11: 3D representation of the LA-4 cells (StarRed DPPE, green) filled with $TiO_2$ nanotubes (Star 520 SXP, red). Cells were treated with 1 mM MBCD for 48 hours beforehand to decrease cholesterol concentration in membrane. See also Figure S124*

Link to time-lapse

- http://lbfnanobiodatabase.ijs.si/file/data/cauliflowerpaper/e03_m03_s01_LA-4_SR_DPPE_TiO2_Star_520_added_1_mMMBCD_NM_uptake.gif

*Movie S12: Time-lapse of filling of the LA-4 cells (StarRed DPPE, green) with $TiO_2$ nanotubes (Star 520 SXP, red) in course of 20 min. Cells were treated with 1 mM MBCD for 48 hours beforehand to decrease cholesterol concentration in membrane. See also Figure S125.*

#### S2i – Block of lipid synthesis by FAS inhibition

##### Main message

Formation of cauliflower-like structures after the chemical inhibition of fatty acid synthase (FAS) enzyme complex by Resveratrol after 2 days of incubation with $TiO_2$ nanotubes.

Supporting raw and analysed data:

Figure S126-Figure S131

##### Materials and methods

- Main experiment:
  - LA-4 cells were seeded @60% confluence in an Ibidi #1.5H µ-Dish in cell medium
  - After 24 hours medium was exchanged for fresh medium with 100 µM Resveratrol and 35 ml of freshly filtered 1 mg/ml $TiO_2$-17-Alexa 647 in 100x dcb was added directly to the cells (in 400 µL medium/100 µM Resveratrol) and mixed to achieve 10:1 surface dose. Cells were incubated for additional 24h.
  - After incubation with $TiO_2$ and Resveratrol cells were incubated with 1 µM SHE-2N for 5 min in incubator at 37°C, 5% CO2. Cells were not washed with LCIS in order not to wash away Resveratrol. Cells were imaged on the microscope stage heated on 37°C.
- Supplement experiments:
  - Sample preparation was the same for supplement experiments. Only difference was labelling of the cells where we stained one sample with 1 µm CellMask and another with 1 µm SirActin in the incubator at 37°C, 5% CO2 for 5 min and 2 hours respectively. Cells were not washed with LCIS in order not to wash away Resveratrol. Cells were imaged on the microscope stage heated on 37°C.

Analysis:

- Images analysed with ImageJ
  - Signal count was multiplied 1.5 times to gain better visibility
  - Sharpening algorithm ($\sigma$ = 2, c = 0,6) was applied on a red channel and 1 pix Gaussian Blur on a green channel

##### Experimental names

- Main experiment name:
  - 20190614_e02 m05 s03 CM 100 µM Resveratrol LA-4 SHE 2N and 10 - 1 S520 TiO2_larger FOV

- Supplement Experiment names:
  - 20190619_e06 m01 s01_LA-4_CellMask_ NEG CNOTROLE for 100 µM Resveratrol and 10 - 1 S520 TiO2_3D
  - 20190619_e05 m05 s04 LA-4_CM 100 µM Resveratrol and 10 - 1 S520 TiO2_Alexa 647_larger feed of view
  - 20190619_e04 m08 s07 LA-4_1 um SA_10 to 1_ TiO2_Star520S_100 µM Resveratrol 24h incubation

- 20190619_e05 m05 s05 LA-4_CM 100 µM Resveratrol and 10 - 1 S520 TiO2_Alexa 647_larger feed of view
- 20190619_e05 m09 s07 LA-4_CM 100 µM Resveratrol and 10 - 1 S520 TiO2_Alexa 647_larger feed of view
- 20190614_e01 m05 s05_ LA-4 SHE 2N and 10 - 1 S520 TiO2_controle
- 20190614_e02 m05 s03 CM 100 µM Resveratrol LA-4 SHE 2N and 10 - 1 S520 TiO2_larger FOV
- 20190614_e02 m04 s03 LA-4 treated with 100 µM Resveratrol and 10-1 TiO2 3D

#### Controls and statistics

Absence of large cauliflower-like structures on the surface of the cells treated with 100 µM Resveratrol after 2 days of incubation with $TiO_2$ nanotubes:

Figure S126

Control experiment where the cells were not treated with the Resveratrol and were exposed to $TiO_2$ under the same conditions. We see formation of cauliflower-like structures on the surface on the cells:

Figure S127-Figure S129

Cells treated with 100 µM Resveratrol and exposed to $TiO_2$ nanotubes for 2 days. Different labels were used, as described in boxes underlying the images:

Figure S130-Figure S131

Main Experiment

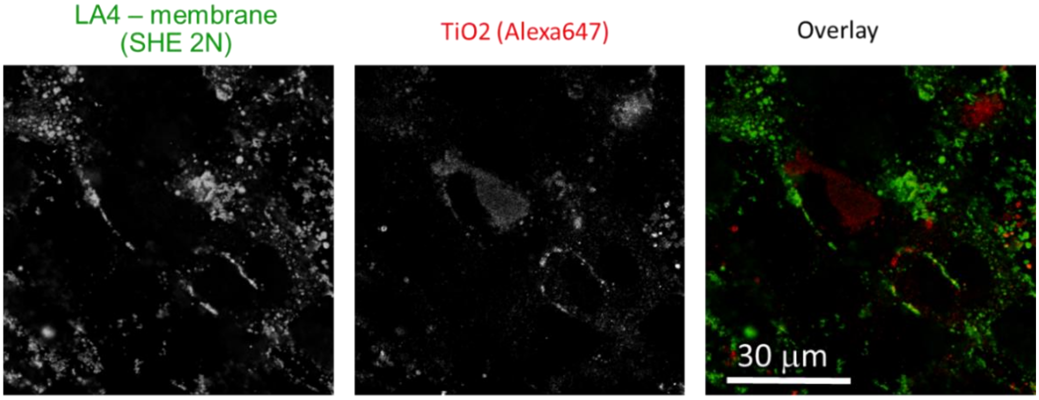

| | |
|---|---|
| Cell line | LA-4 (membrane, SHE 2N) |
| NPs | TiO2 (Alexa 647) |
| exposure | 10:1, 24h-48h |
| imaging | xyz confocal, 48h Resveratrol 100 uM |

| | |
|---|---|
| pixelsize (x,y) | 200 nm |
| FOV (x,y) | 80 um |
| pixelsize (z) | / nm |
| FOV (z) | / um |
| imaging time | / min |
| number z-stacks | |

| | |
|---|---|
| 561nm | 30% |
| 640nm | 30% |
| STED | 0% |
| filter sets | 605 nm – 625 nm, 650 nm – 720 nm |
| dwell time | 10 ms |
| objective | wi60x (NA1.2) |

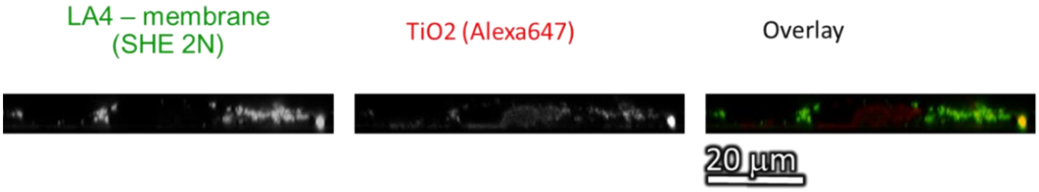

| | |
|---|---|
| Cell line | LA-4 (membrane, SHE 2N) |
| NPs | TiO2 (Alexa 647) 100 uM Resveratrol |
| exposure | 10:1, 24h-48h |
| imaging | xz confocal, 48h Resveratrol 100 uM |

| | |
|---|---|
| pixelsize (x,y) | 50 nm |
| FOV (x,y) | 70 um |
| pixelsize (z) | 50 nm |
| FOV (z) | 9 um |
| imaging time | / min |
| number z-stacks | |

| | |
|---|---|
| 561nm | 30% |
| 640nm | 30% |
| STED | 0% |
| filter sets | 605 nm – 625 nm, 650 nm – 720 nm |
| dwell time | 10 ms |
| objective | wi60x (NA1.2) |

*Figure S126: Confocal images of LA-4 in green and $TiO_2$ nanotubes in red channel. Cells treated with 100 µM Resveratrol for 48 hours. No collocalization and no cauliflower-like formations. XY on upper window and XZ on lower.*

Supplement Experiments

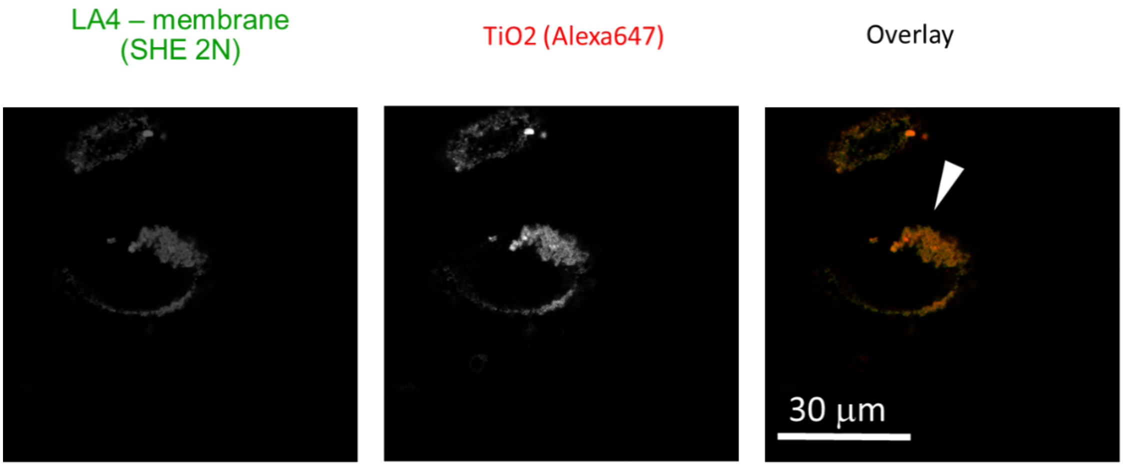

| | |
|---|---|
| Cell line | LA-4 (membrane, SHE 2N) |
| NPs | TiO2 (Alexa 647) |
| exposure | 10:1, 24h-48h |
| imaging | xz confocal, 48h CONTROLE |

| | |
|---|---|
| pixelsize (x,y) | 200 nm |
| FOV (x,y) | 80 x 80 um |
| pixelsize (z) | / |
| FOV (z) | / |
| imaging time | / |
| number z-stacks | / |

| | |
|---|---|
| 561nm | 30% |
| 640nm | 30% |
| STED | 0% |
| filter sets | 605 nm – 625 nm, 650 nm – 720 nm |
| dwell time | 10 ms |
| objective | wi60x (NA1.2) |

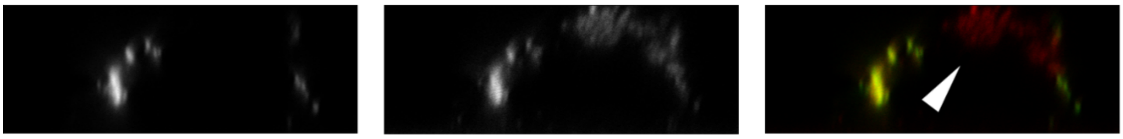

| | |
|---|---|
| Cell line | LA-4 (membrane, SHE 2N) |
| NPs | TiO2 (Alexa 647) |
| exposure | 10:1, 24h-48h |
| imaging | xz confocal, 48h CONTROLE |

| | |
|---|---|
| pixelsize (x,y) | 50 nm |
| FOV (x,y) | 36 um |
| pixelsize (z) | 50 nm |
| FOV (z) | 13 um |
| imaging time | / |
| number z-stacks | |

| | |
|---|---|
| 561nm | 30% |
| 640nm | 30% |
| STED | 0% |
| filter sets | 605 nm – 625 nm, 650 nm – 720 nm |
| dwell time | 10 ms |
| objective | wi60x (NA1.2) |

*Figure S127: Confocal images of LA-4 in green and $TiO_2$ nanotubes in red channel. Cells not treated with Resveratrol. We can see collocalization and cauliflower-like formations on the topp of the cell. XY on upper window and XZ on lower.*

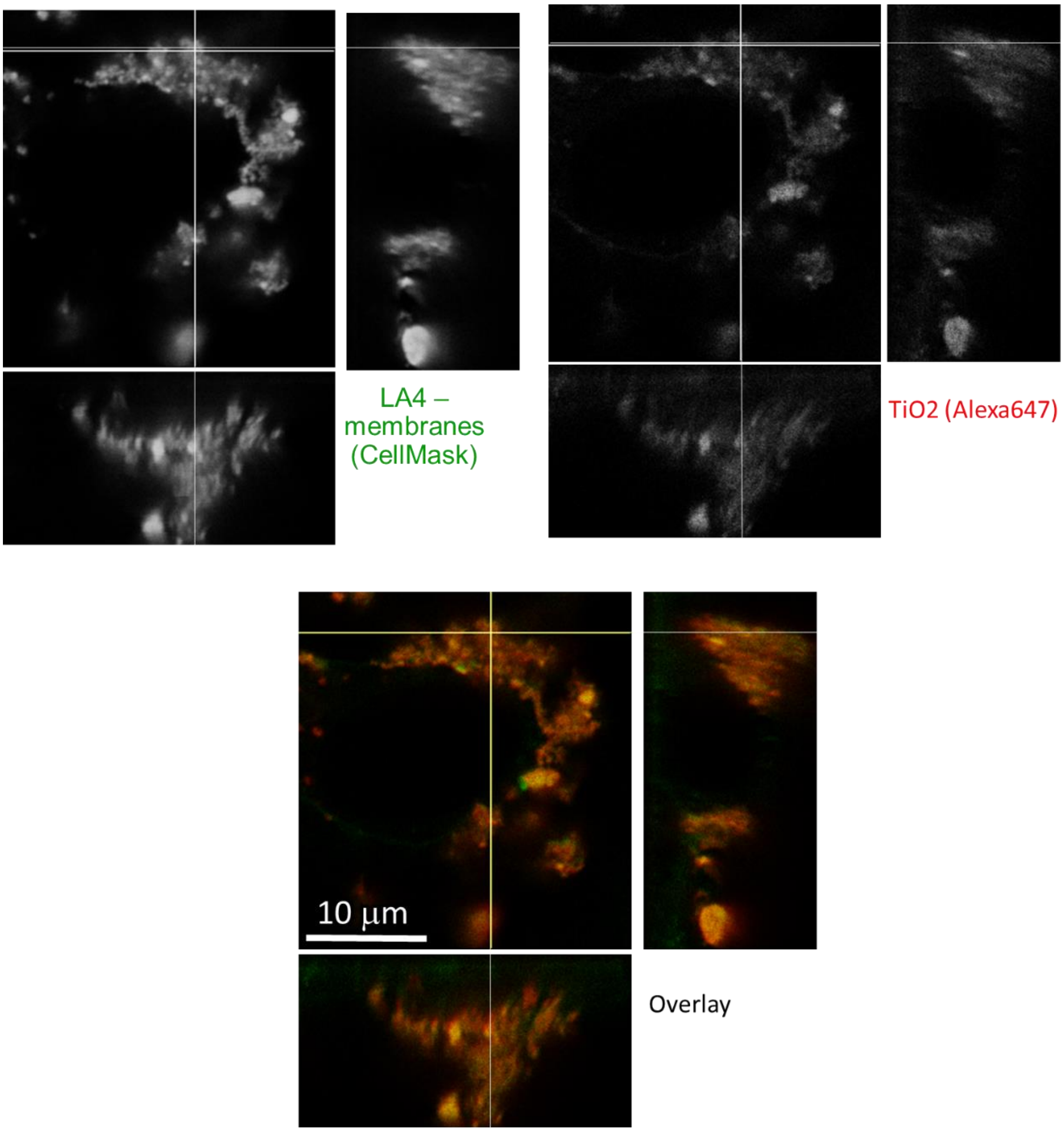

| | |
|---|---|
| Cell line | LA-4 (membrane, CellMask) |
| NPs | TiO2 (Alexa 647) CONTROLE |
| exposure | 10:1, 24h-48h |
| imaging | xyz STED, 48h CONTROLE |

| | |
|---|---|
| pixelsize (x,y) | 100 nm |
| FOV (x,y) | 28 x 31 um |
| pixelsize (z) | 100 nm |
| FOV (z) | 15 um |
| imaging time | 10 min |
| number z-stacks | 150 |

| | |
|---|---|
| 561nm | 50% |
| 640nm | 50% |
| STED | 10% |
| filter sets | 605 nm – 625 nm, 650 nm – 720 nm |
| dwell time | 10 ms |
| objective | wi60x (NA1.2) |

*Figure S128: Confocal images of LA-4 in green and $TiO_2$ nanotubes in red channel. Image is just one z-slice from a whole z-stack. Cells not treated with Resveratrol. We can see collocalization and cauliflower-like formations on the topp of the cell. XY on upper window and XZ on lower.*

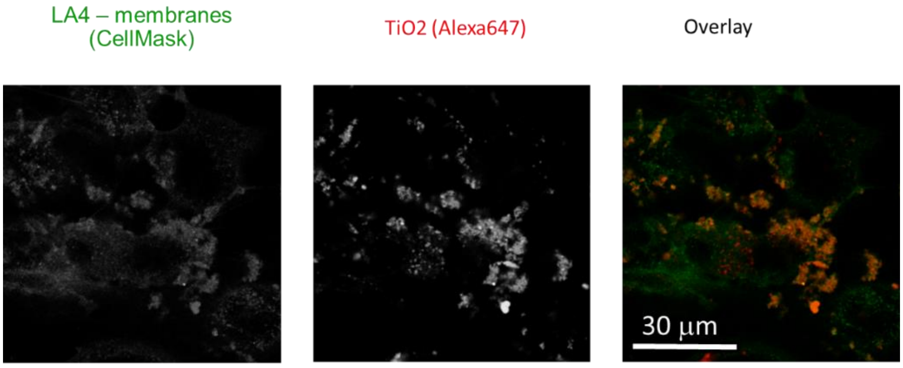

| | |
|---|---|
| Cell line | LA-4 (membrane, CellMask) |
| NPs | TiO2 (Alexa 647) |
| exposure | 10:1, 24h-48h CONTROLE |
| imaging | xy confocal, 48h CONTROLE |

| | |
|---|---|
| pixelsize (x,y) | 200 nm |
| FOV (x,y) | 80x 80 um |
| pixelsize (z) | / |
| FOV (z) | / |
| imaging time | / |
| number z-stacks | / |

| | |
|---|---|
| 561nm | 30% |
| 640nm | 30% |
| STED | 0% |
| filter sets | 605 nm – 625 nm, 650 nm – 720 nm |
| dwell time | 10 ms |
| objective | wi60x (NA1.2) |

*Figure S129: Confocal images of LA-4 in green and $TiO_2$ nanotubes in red channel. Cells not treated with Resveratrol. We can see collocalization and cauliflower-like formations on the topp of the cell. XY on upper window and XZ on lower.*

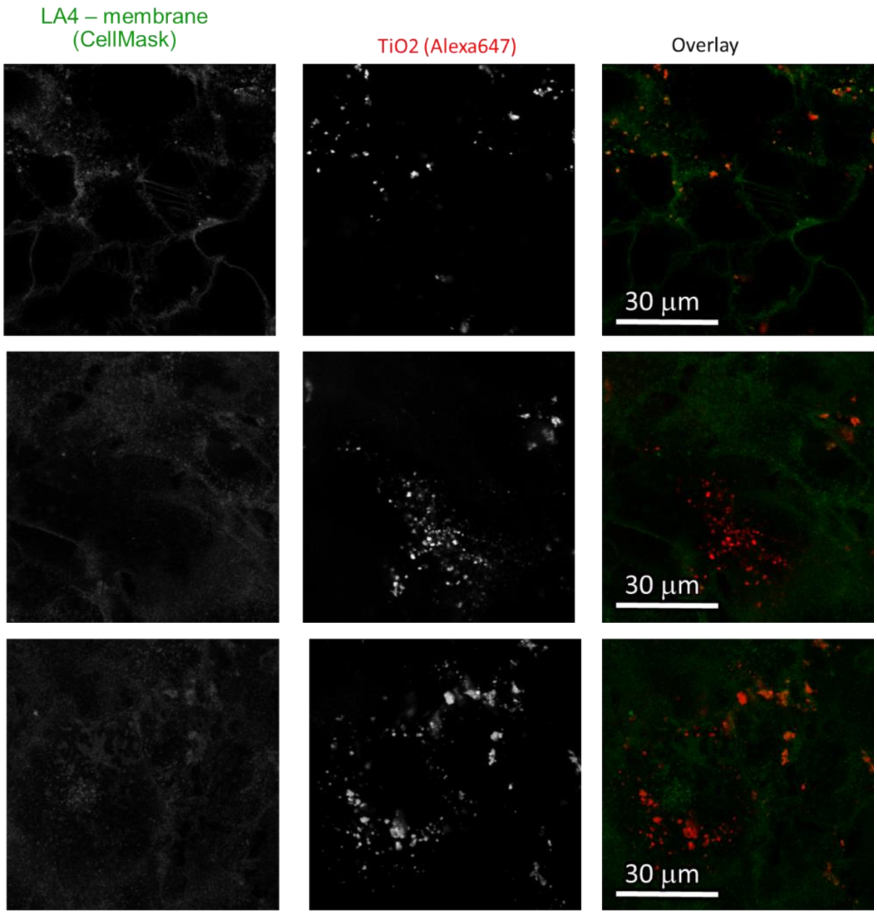

| | |
|---|---|
| Cell line | LA-4 (membrane, CellMask) |
| NPs | TiO2 (Alexa 647) |
| exposure | 10:1, 24h-48h |
| imaging | xy confocal, 48h Resveratrol 100 uM |

| | |
|---|---|
| pixelsize (x,y) | 200 nm |
| FOV (x,y) | 80x 80 um |
| pixelsize (z) | / |
| FOV (z) | / |
| imaging time | / |
| number z-stacks | / |

| | |
|---|---|
| 561nm | 30% |
| 640nm | 30% |
| STED | 0% |
| filter sets | 605 nm – 625 nm, 650 nm – 720 nm |
| dwell time | 10 ms |
| objective | wi60x (NA1.2) |

*Figure S130: Confocal images of LA-4 in green and $TiO_2$ nanotubes in red channel. Cells treated with 100 µM Resveratrol for 48 hours. No collocalization and no cauliflower-like formations. XY shots of three different FoVs.*

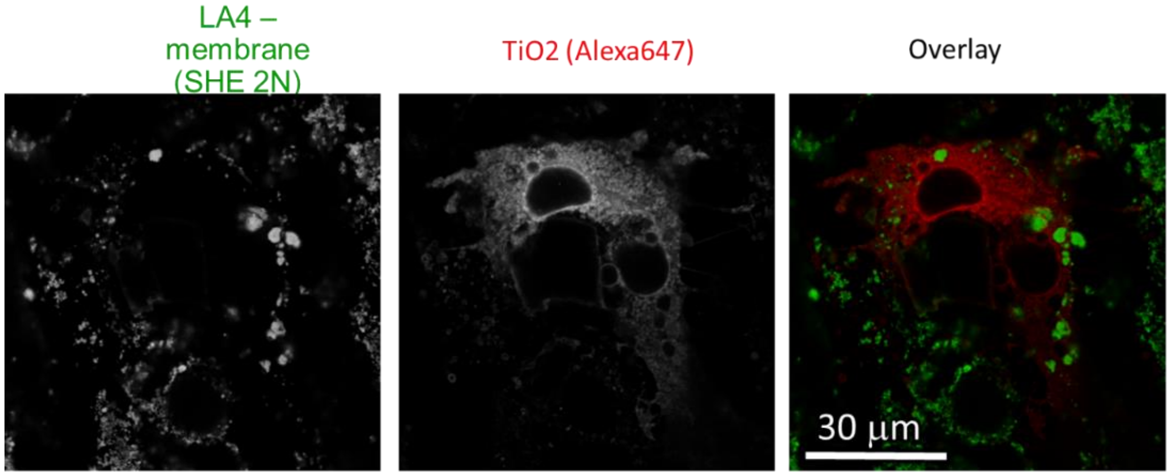

| | |
|---|---|
| Cell line | LA-4 (membrane, CellMask) |
| NPs | TiO2 (Alexa 647) |
| exposure | 10:1, 24h-48h |
| imaging | xy confocal, 48h Resveratrol 100 uM |

| | |
|---|---|
| pixelsize (x,y) | 200 nm |
| FOV (x,y) | 80x 80 um |
| pixelsize (z) | / |
| FOV (z) | / |
| imaging time | / |
| number z-stacks | / |

| | |
|---|---|
| 561nm | 30% |
| 640nm | 30% |
| STED | 0% |
| filter sets | 605 nm – 625 nm, 650 nm – 720 nm |
| dwell time | 10 ms |
| objective | wi60x (NA1.2) |

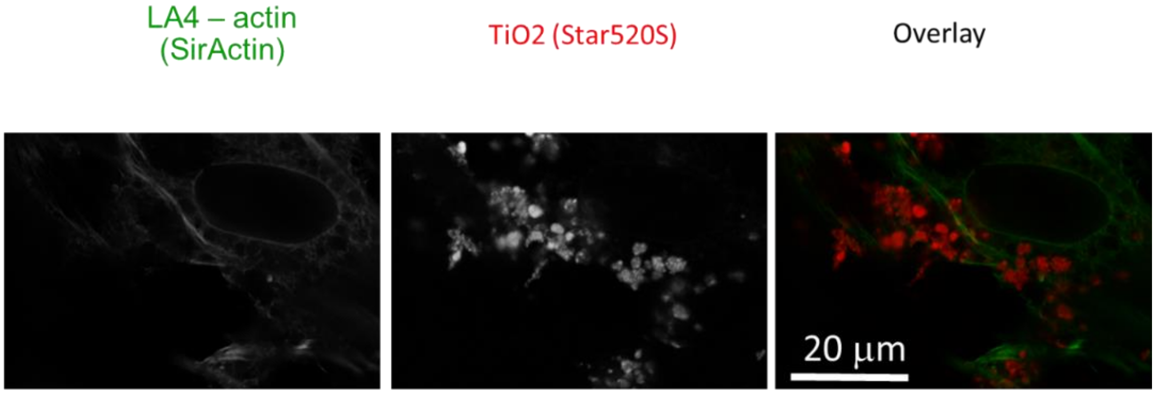

| | |
|---|---|
| Cell line | LA-4 (actin, SirActin) |
| NPs | TiO2 (Star 520S) |
| exposure | 10:1, 24h-48h |
| imaging | xy confocal, 48h Resveratrol 100 uM |

| | |
|---|---|
| pixelsize (x,y) | 60 nm |
| FOV (x,y) | 65 x 45 um |
| pixelsize (z) | / |
| FOV (z) | / |
| imaging time | / |
| number z-stacks | / |

| | |
|---|---|
| 561nm | 50% |
| 640nm | 50% |
| STED | 20% |
| filter sets | 605 nm – 625 nm, 650 nm – 720 nm |
| dwell time | 10 ms |
| objective | wi60x (NA1.2) |

*Figure S131: Confocal images of LA-4 in green and $TiO_2$ nanotubes in red channel. Cells treated with 100 µM Resveratrol for 48 hours. No collocalization and no cauliflower-like formations. In upper window we can see LA-4 cells stained with plasma membrane dye Cell Mask Orange. In the lower window LA-4 cells have been stained with Sir Actin dye which stains F-actin.*

Link to 3D

- http://lbfnanobiodatabase.ijs.si/file/data/cauliflowerpaper/20190619_Treated_with_100_mM_Resveratrol_overnight_overview_NM_in_the_cell_XYZ.mp4

*Movie S13: Cells treated with 100 µM Resveratrol, show reduced cells cauliflower formation and more nanomaterial in the cytosolic side of the cell.*

Link to time-lapse

- http://lbfnanobiodatabase.ijs.si/file/data/cauliflowerpaper/20190619_e06_m01_s01_LA-4_CellMask__NEG_CNOTROLE__for_100mM_Resveratrol_and_10_-_1_S520_TiO2_3D.mp4

*Movie S14: Cells not treated with Resveratrol grow huge cauliflowers on the surface of membranes outside of the cell*

### S3 – The role of actin

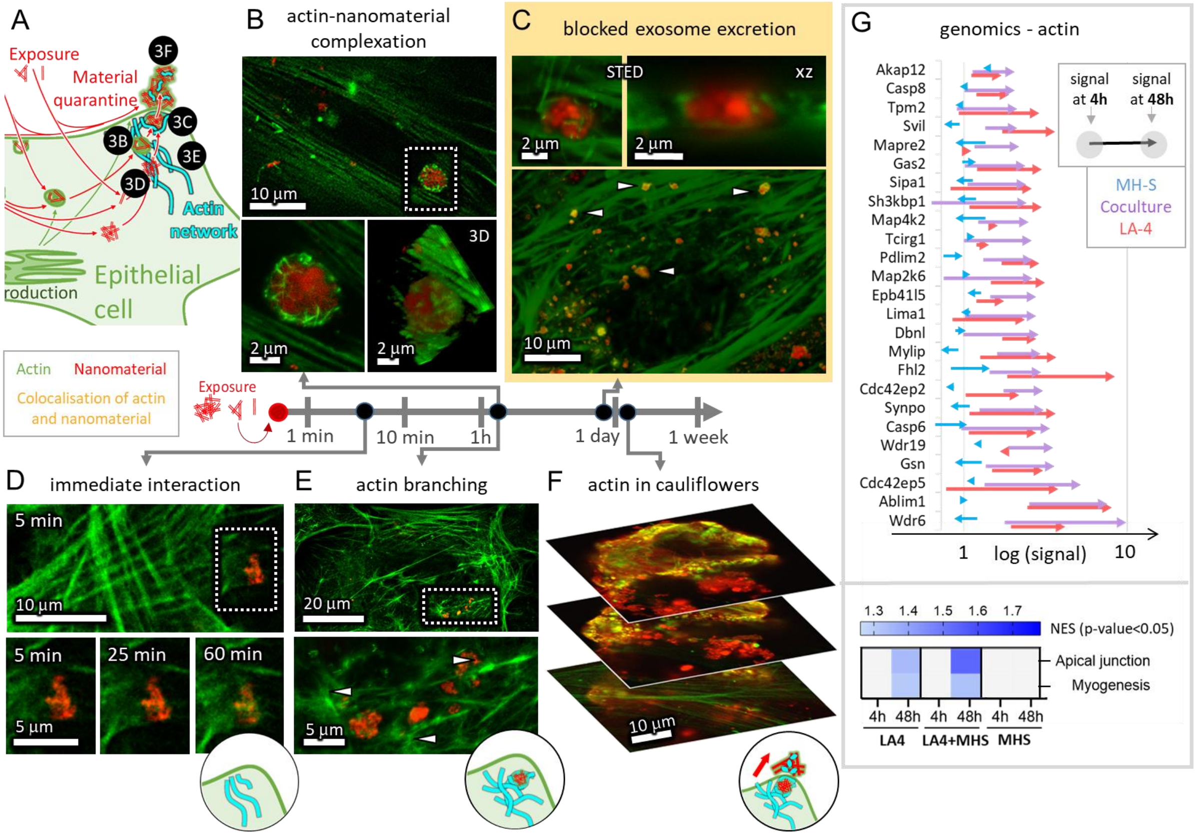

#### S3b – Actin – nanomaterial aggregation

##### Main message

Fluorescent micrographs of cytoskeleton after few hours of incubation with the $TiO_2$ nanotubes.

Supporting raw and analysed data:

Figure S132-Figure S134

##### Materials and methods

- Main experiment:
  - LA-4 cells were seeded @70% confluence in an Ibidi #1.5H µ-Dish and incubated at 37°C and 5% CO2
  - after 24 hours fresh 400 µL medium with 1 µM of SirActin (Spirochrome Probes for Bioimaging) was added and incubated for additional 2 hours
  - After incubation cells were washed 3x400 µL LCIS and exposed to 35 µl freshly filtered 1 mg/ml $TiO_2$-40-ATTO 594 in 100x dcb (in 400 µL LCIS), mixed to achieve 10:1 surface dose
  - Cells were observed in 400 µL LCIS with label on the microscope stage heated at 37°C for 2h

- Supplement experiments:
  - 20190122_e01_m05_LA-4_SA_actin interaction with TiO2_ATTO 594 has the same sample preparation procedure
  - 20190711_e01 m16 s10_LA-4_1 um SA & TiO2_S520_1 - 10_24h incubation _3D_faberges sample was seeded @40% confluency in an Ibidi #1.5H µ-Dish and incubated at 37°C and 5% CO2
  - After 24 hours after 24 hours fresh 400 µL medium with 1 µM of Sir Actin was added and exposed to 35 µl freshly filtered 1 mg/ml $TiO_2$-17-Star 520S and incubated for further 24h
  - After incubation cells were washed 3x400 µL LCIS and observed in 400 µL LCIS on the microscope stage heated at 37°C for 2h

- Analysis (Image J):
  - 3D STED: in red channel signal was multiplied to gain maximal visibility and noise was reduced by Gaussian Blur (1 pix); in green channel only Gaussian Blur (1 pix) was applied
  - Confocal: red channel has logarithmic scale, and Gaussian filter (1 pix) was used
  - For STED image in red channel Gaussian Blur was applied; in green to better visualize fine actin strands we sharpened the image ($\sigma = 2$, $c = 0,4$) using following:

##### Experiment names

- Main experiment name:
  - 20190122_e02_m05_Aleksandar_LA-4_SA and TiO2_ATTO 594_ Faberge_exp09_01

- 20190122_e02_m05_Aleksandar_LA-4_SA and TiO2_ATTO 594_ Faberge_exp09_02
- 20190122_e02_m05_Aleksandar_LA-4_SA and TiO2_ATTO 594_ Faberge_exp09_03

- Supplement Experiment names:
  - 20190122_e01_m05_LA-4_SA and TiO2_ATTO594_interaction_ after 120_min.msr
  - 20190122_e01_m06_LA-4_SA and TiO2_ATTO594_interaction_ after 120_min.msr
  - 20190711_e01 m16 s10_LA-4_1 um SA and TiO2_S520_1 - 10_24h incubation _3D_faberges

#### Controls and statistics

Four vertical (z) sections of a 3D stack of images showing one large $TiO_2$ nanotubes -actin structure:

Figure S132

STED zoom in of the 3D stack from previous figure and large field of view of the same structure:

Figure S133

STED time lapse of an actin engulfing a $TiO_2$ aggregate:

Figure S134

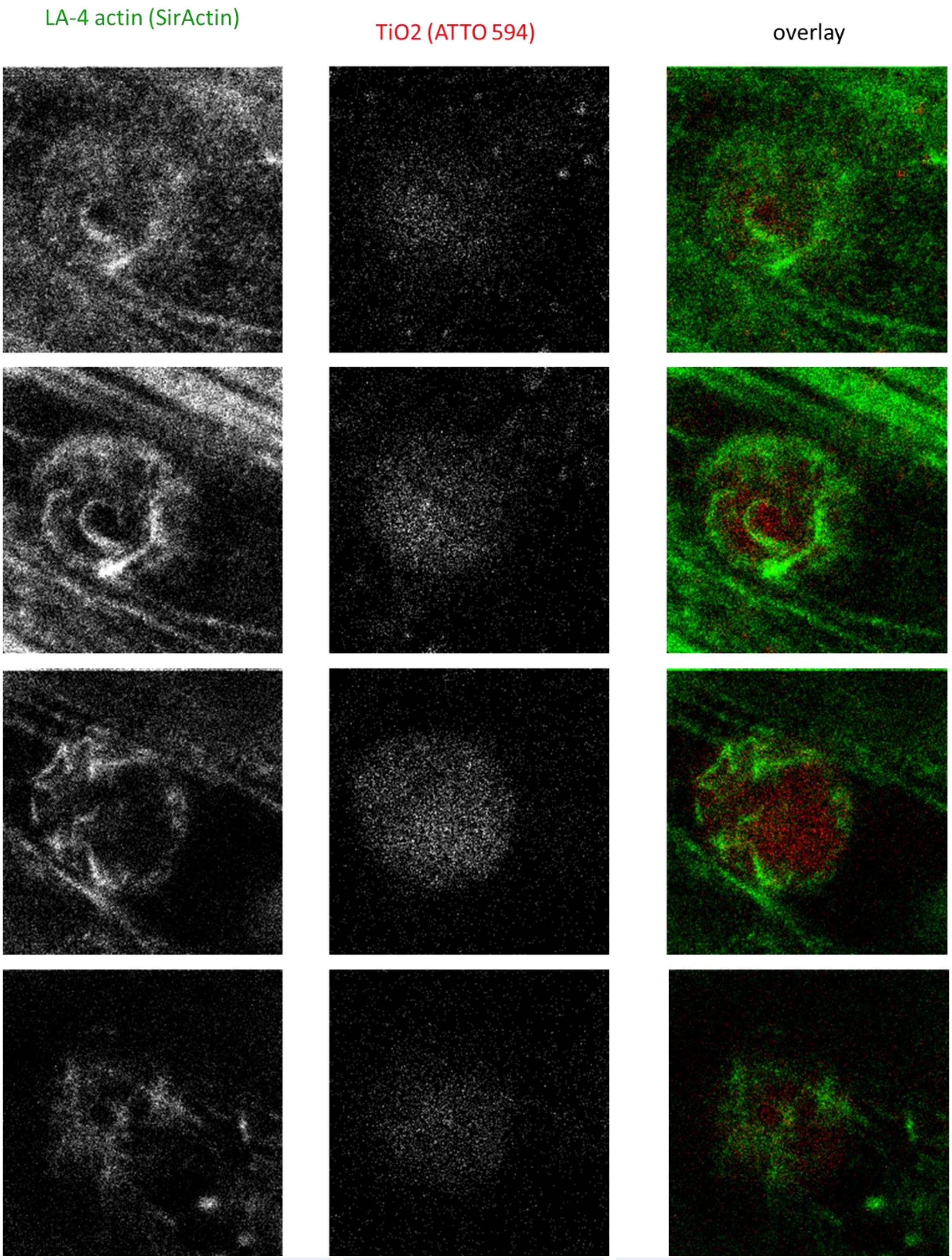

| | | | | | |
|---|---|---|---|---|---|
| Cell line | LA-4 (actin, SirActin) | pixelsize (x,y) | 40 nm | 561nm | 26% |
| NPs | TiO2 (ATTO 594) | FOV (x,y) | 9 µm | 640nm | 15% |
| exposure | 10:1, 26h-28h | pixelsize (z) | 60 nm | STED | 10% |
| imaging | XYZ STED, 26h | FOV (z) | 4 µm | filter sets | 605 nm – 625 nm, 650 nm – 720 nm |
| | | imaging time | / | dwell time | 10 µs |
| | | number of frames | 68 | objective | wi60x (NA1.2) |

*Figure S132: Confocal images of LA-4 in green and $TiO_2$ nanotubes in red channel. Cells are stained with Sir Actin label for F-actin. STED images of XY planes of different slices of a whole Z-stack.*

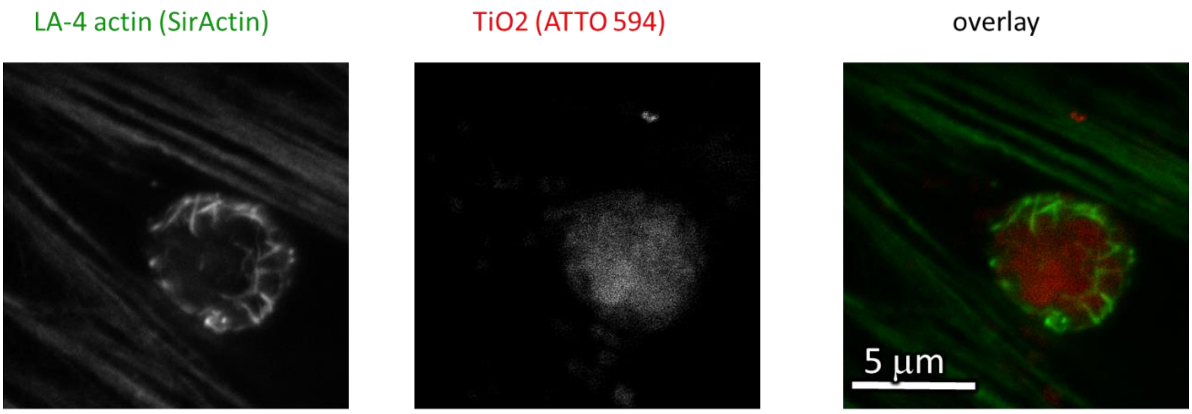

| | |
|---|---|
| Cell line | LA-4 (actin, SirActin) |
| NPs | TiO2 (ATTO 594) |
| exposure | 10:1, 26h-28h |
| imaging | xy STED, 26h<br>xy confocal, 26h |

| | |
|---|---|
| pixelsize (x,y)<br>(STED) | 15nm |
| FOV (x,y) | 14 µm |
| pixelsize (z) | / nm |
| FOV (z) | / µm |
| imaging time | / |
| number of frames | / |

| | |
|---|---|
| 561nm | 30% |
| 640nm | 15% |
| STED | 10% |
| filter sets | 605 nm – 625 nm,<br>650 nm – 720 nm |
| dwell time | 10 µs |
| objective | wi60x (NA 1.2) |

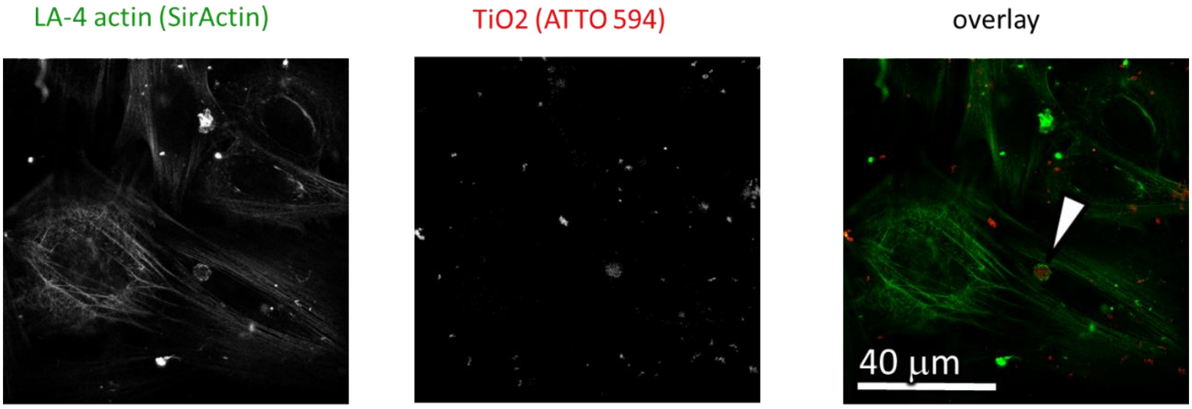

| | |
|---|---|
| Cell line | LA-4 (actin, SirActin) |
| NPs | TiO2 (Alexa647) |
| exposure | 10:1, 26h-28h |
| imaging | xy STED, 26h<br>xy confocal, 26h |

| | |
|---|---|
| pixelsize (x,y)<br>(confocal) | 200 nm |
| FOV (x,y) | 100 µm |
| pixelsize (z) | / nm |
| FOV (z) | / µm |
| imaging time | / |
| number of frames | / |

| | |
|---|---|
| 561nm | 5% |
| 640nm | 10% |
| STED | / |
| filter sets | 605 nm – 625 nm,<br>650 nm – 720 nm |
| dwell time | 10 µs |
| objective | wi60x (NA 1.2) |

*Figure S133: STED and confocal images of LA-4 in green and $TiO_2$ nanotubes in red channel. Cells are stained with Sir Actin label for F-actin. STED zoom – in (XY plane) the upper window shows the same actin-NTs agregat from the previous figure. In lower window is the confocal image (XY plane) where the specific location of the actin-NTs agregate in the cell is clearly visible.*

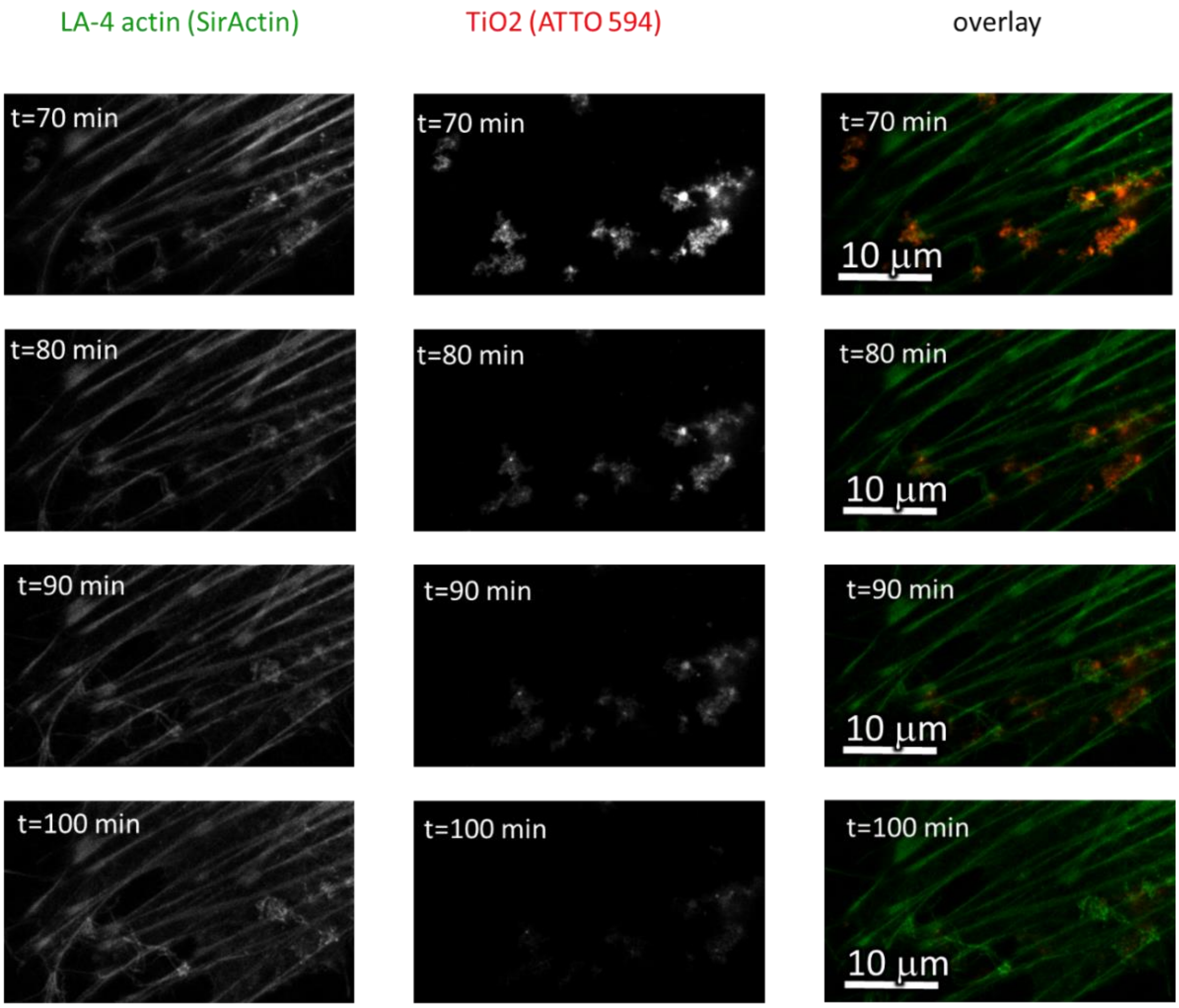

| | |
|---|---|
| Cell line | LA-4 (actin, SirActin) |
| NPs | TiO2 (ATTO 594) |
| exposure | 10:1, 26h-27h |
| imaging | xyt STED, 27h-27h |

| | |
|---|---|
| pixelsize (x,y) | 70 nm |
| FOV (x,y) | 37 μm |
| pixelsize (z) | / |
| FOV (z) | / |
| imaging time | 1 hour |
| number of frames | 611 |

| | |
|---|---|
| 561nm | 40% |
| 640nm | 25% |
| STED | 15% |
| filter sets | 605 nm – 625 nm, 650 nm – 720 nm |
| dwell time | 10 μs |
| objective | wi60x (NA1.2) |

*Figure S134: STED time lapse of LA-4 in green and $TiO_2$ nanotubes in red channel. Shown are time points from 70 – 100 min with 10 min increments. Actin fibers are completely engulfing $TiO_2$ nanotubes.*

#### Link to 3D

- http://lbfnanobiodatabase.ijs.si/file/data/cauliflowerpaper/20190122_e02_m05_Aleksandar_LA-4_Sir-Actin_Fabrege_exp09_03_.gif

*Movie S15: Interaction of actin fibres with nanoaggregates in the cell after few hours - Faberge egg*

- http://lbfnanobiodatabase.ijs.si/file/data/cauliflowerpaper/20190122_e02_m05_Aleksandar_LA-4_Sir-Actin_Fabrege_exp09_03_.mp4

*Movie S16: Interaction of actin fibres with nanoaggregates in the cell after few hours - Faberge egg*

- http://lbfnanobiodatabase.ijs.si/file/data/cauliflowerpaper/20190122_e02_m05_Aleksandar_LA-4_Sir-Actin_Fabrege_exp09_03_1.mp4

*Movie S17: Interaction of actin fibres with nanoaggregates in the cell after few hours - Faberge egg*

- http://lbfnanobiodatabase.ijs.si/file/data/cauliflowerpaper/20190711_e01_m16_s10_LA-4_1_uM_SA_&_TiO2_S520_1_-_10_24h_incubation__3D_faberges.gif

*Movie S18: Interaction of actin with uptaken nanomaterial after 24h - Faberge egg pattern*

#### S3c – Inhibition of exosome excretion by high concentrations of Jaspakinolide

##### Main message

Upon stabilisation of actin fibres, cell is incapable to compress formed actin rings around to-be-exported cargo, which we see as red globes of nanomaterial, unable to be extruded, wrapped by green actin filaments close to the surface of the cell.

Supporting raw and analysed data:

Figure S135-Figure S140

##### Materials and methods

- Main experiment:
  - LA-4 cells were seeded @30% confluence in an Ibidi #1.5H μ-Dish
  - A) 24 hours later, the cells were incubated with 100 nM SirActin and Jaspakinolide overnight at the incubator at 37°C and 5% CO2. On the 3rd day the cells were washed with 1x400 μl LCIS and observed in 400 μl LCIS at microscope stage heated on 37°C
  - B) 24 hours later, the cells were incubated with 1 μM SirActin and Jaspakinolide overnight at the incubator at 37°C and 5% CO2. On the 3rd day the cells were washed with 1x400 μl LCIS and observed in 400 μl LCIS at microscope stage heated on 37°C
  - C) After 24 hours 35 ml freshly filtered 1 mg/ml $TiO_2$-17-Alexa647 in 100x dcb was added directly to the cells (in 400 μL medium with 1 um SirActin) and mixed to achieve 10:1 surface dose
  - With A and B setup we allowed the cauliflowers to grow and observe what role actin has with already formed cauliflowers as opposed with C setup, where stabilisation of actin would weaken cauliflower forming potential

- Main experiment (Fig 4c) has been performed as described in A, in supplement we can see same effect of C
  - Supplement experiments have been prepared in the same way

- Analysis:
  - Confocal images are just exported in raw format
  - 0,5 pixel Gaussian blur was applied on STED images

##### Experiment names

- Main experiment name:
  - 20190627_e08 m13 s03 LA-4_1um SA & 10 - 1 TiO2_S520 3DI_omega vesicles
  - 20190627_e08 m13 s12 LA-4_1um SA & 10 - 1 TiO2_S520 3DI_omega vesicles_xy STED

- Supplement Experiment names:
  - 20190627_e01 m04 s03 LA-4_1um SA & 10 - 1 TiO2_S520 3DI_confocal
  - 20190627_e01 m01 s01 LA-4_1um SA & 10 - 1 TiO2_S520 3DI_confocal

- 20190627_e01 m03 s02 LA-4_1um SA & 10 - 1 $TiO_2$_S520 3DI_confocal
- 20190627_e01 m06 s05 LA-4_1um SA & 10 - 1 $TiO_2$_S520 3DI_confocal
- 20190711_e01m09s05_LA-4 1um SA and 10 to 1 $TiO_2$ Star520S_3D
- 20190711_e01 m16 s10_LA-4_ 1um SA and 10 to 1 $TiO_2$_Star520s_Omega vesicles_3D

#### Controls and statistics

Confocal and STED images of actin rings after Jaspankinolide treatment:

Figure S135-Figure S139

LA-4 actin (SirActin) | TiO2 (Star 520S) | overlay

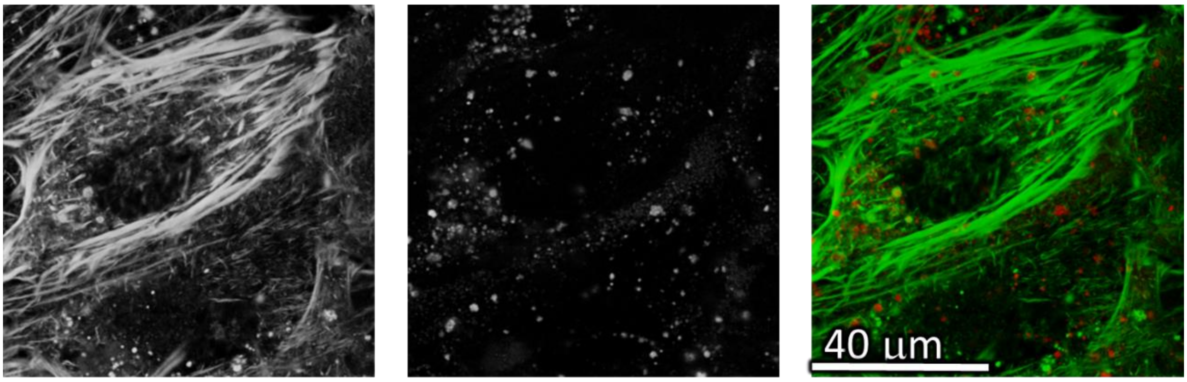

| | |
|---|---|
| Cell line | LA-4 (actin, SirActin) |
| NPs | TiO2 (Star 520S) |
| exposure | 10:1, 24h-60h C |
| imaging | xy confocal, 60h |

| | |
|---|---|
| pixelsize (x,y) | 200 nm |
| FOV (x,y) | 80 µm |
| pixelsize (z) | / nm |
| FOV (z) | / µm |
| imaging time | / |
| number of frames | / |

| | |
|---|---|
| 561nm | 30% |
| 640nm | 50% |
| STED | 20% |
| filter sets | 605 nm – 625 nm, 650 nm – 720 nm |
| dwell time | 10 µs |
| objective | 60wi0x (NA1.2) |

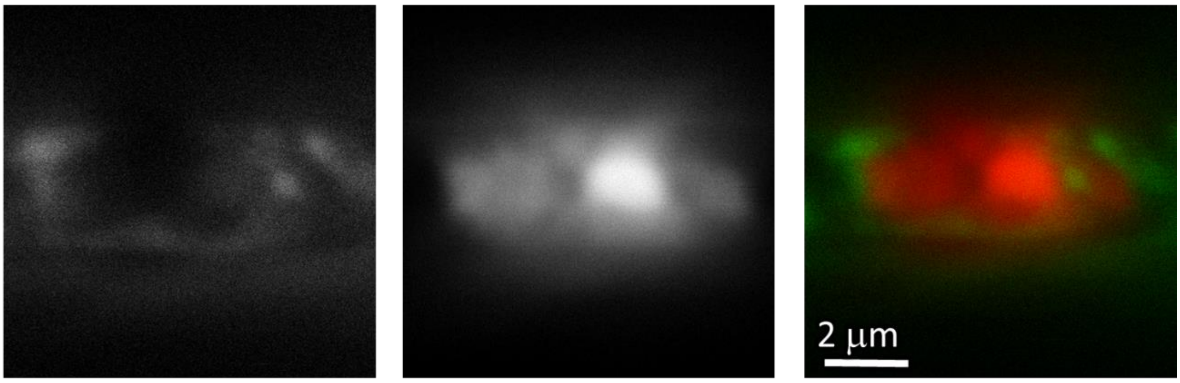

| | |
|---|---|
| Cell line | LA-4 (actin, SirActin) |
| NPs | TiO2 (Star 520S) |
| exposure | 10:1, 24h-60h C |
| imaging | xz STED, 60h |

| | |
|---|---|
| pixelsize (x,y) | 10 nm |
| FOV (x,y) | 8,5 µm |
| pixelsize (z) | 18 nm |
| FOV (z) | 20 µm |
| imaging time | / |
| number of frames | / |

| | |
|---|---|
| 561nm | 30 % |
| 640nm | 35 % |
| STED | 15 % |
| filter sets | 605 nm – 625 nm, 650 nm – 720 nm |
| dwell time | 10 µs |
| objective | 60wi0x (NA1.2) |

*Figure S135: Confocal image and STED images of LA-4 in green and $TiO_2$ nanotubes in red channel. Cells are stained with Sir Actin label for F-actin. Confocal image of large FoV in upper window and STED zoom – in (XZ plane) in the lower window showing representative actine rings, unable to compress and excrete nanomaterial outside the cell due to Jaspankinolide treatmant.*

LA-4 actin (SirActin) | TiO2 (Star 520S) | overlay

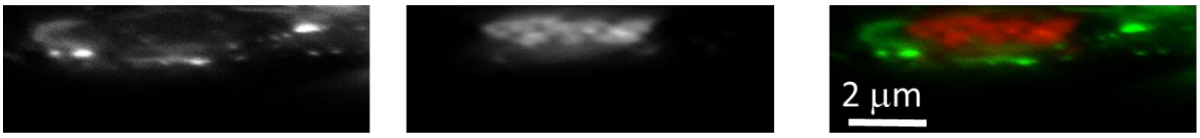

| pixelsize (x,y) | 10 nm |
| --- | --- |
| FOV (x,y) | 9 µm |
| pixelsize (z) | 15,6 nm |
| FOV (z) | 5 µm |

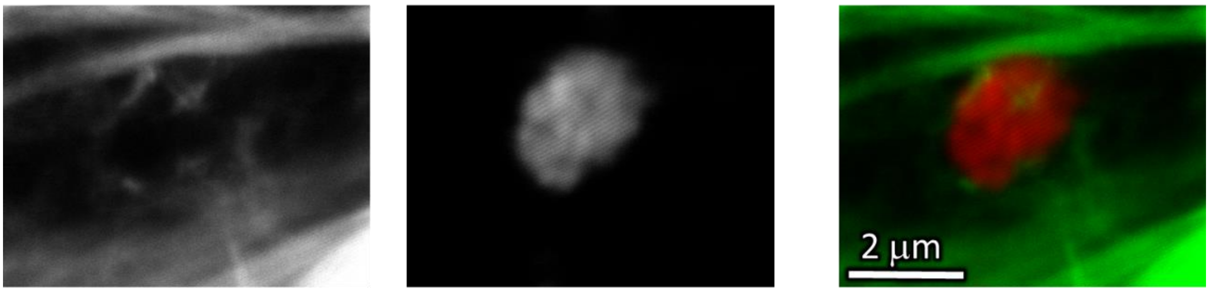

| pixelsize (x,y) | 5nm |
| --- | --- |
| FOV (x,y) | 6*4,6 µm |
| pixelsize (z) | / |
| FOV (z) | / |

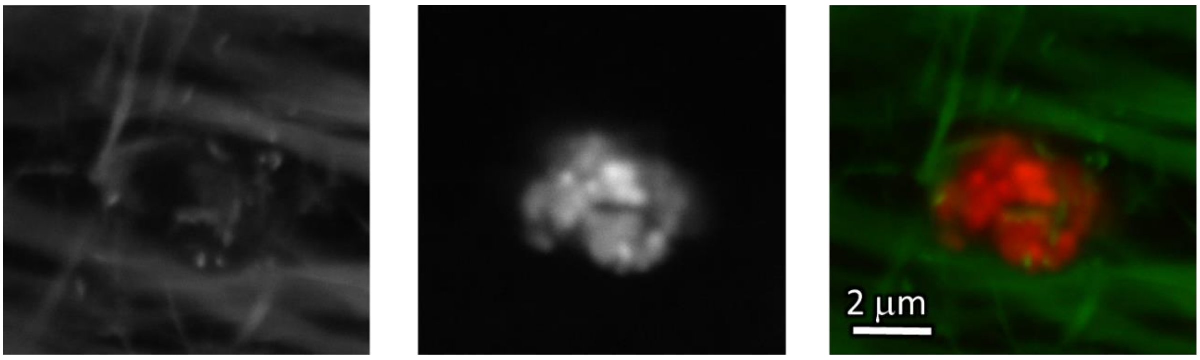

| pixelsize (x,y) | 8 nm |
| --- | --- |
| FOV (x,y) | 8,6 µm |
| pixelsize (z) | 8 nm |
| FOV (z) | 8,2 µm |

*Figure S136: STED images of LA-4 in green and $TiO_2$ nanotubes in red channel. Cells are stained with Sir Actin label for F-actin. STED zoom – ins (XZ plane in upper window and two XY planes in middle and lower window) showing representative actine rings, unable to compress and excrete nanomaterial outside the cell due to Jaspankinolide treatmant. All parameters which are left out from boxes are the same as for the above images.*

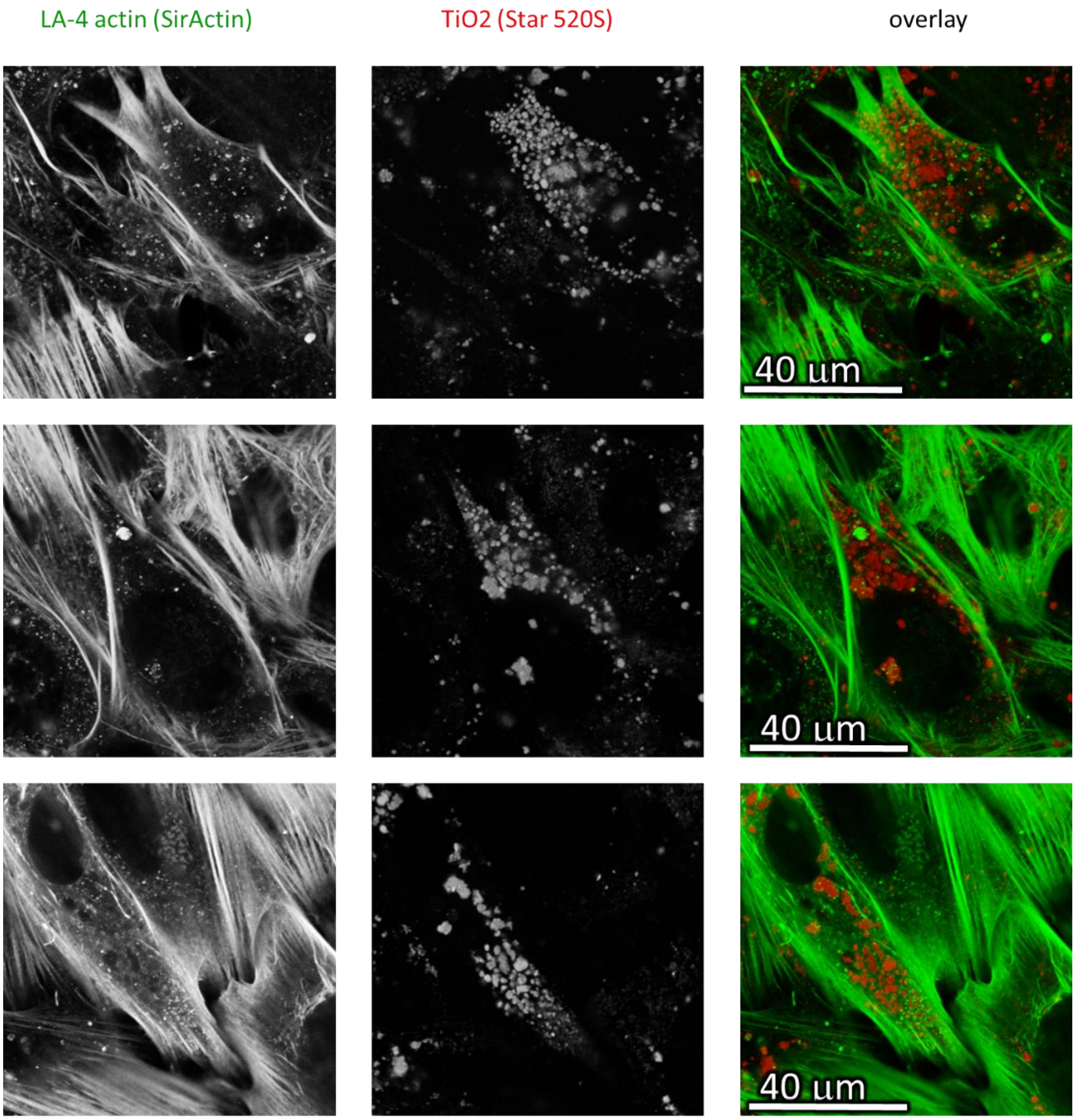

| | |
|---|---|
| Cell line | LA-4 (actin, SirActin) |
| NPs | TiO2 (Star 520S) |
| exposure | 10:1, 24h-60h |
| imaging | xy confocal, 60h |

| | |
|---|---|
| pixelsize (x,y) | 200 nm |
| FOV (x,y) | 80 µm |
| pixelsize (z) | / nm |
| FOV (z) | / µm |
| imaging time | / |
| number of frames | / |

| | |
|---|---|
| 561nm | 30% |
| 640nm | 50% |
| STED | / |
| filter sets | 605 nm – 625 nm,<br>650 nm – 720 nm |
| dwell time | 10 µs |
| objective | 60wi0x (NA1.2) |

*Figure S137: Confocal images of LA-4 in green and $TiO_2$ nanotubes in red channel. Cells are stained with Sir Actin label for F-actin. Confocal image of large FoVs in all window showing representative actine rings, unable to compress and excrete nanomaterial outside the cell due to Jaspankinolide treatmant.*

LA-4 actin (SirActin) | TiO2 (Star 520S) | overlay

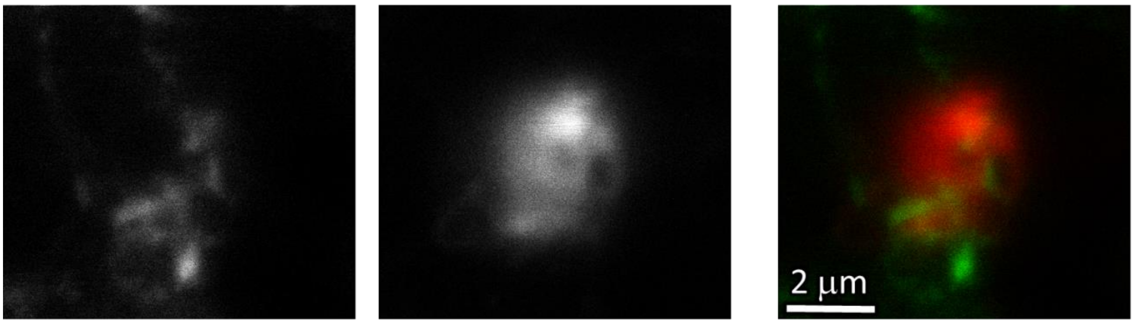

| | |
|---|---|
| Cell line | LA-4 (actin, SirActin) |
| NPs | TiO2 (Star 520S) |
| exposure | 10:1, 24h-60h |
| imaging | xy STED, 60h |

| | |
|---|---|
| pixelsize (x,y) | 10 nm |
| FOV (x,y) | 7,4 x 6,6 µm |
| pixelsize (z) | / nm |
| FOV (z) | / µm |
| imaging time | / |
| number of frames | / |

| | |
|---|---|
| 561nm | 50% |
| 640nm | 65% |
| STED | 30% |
| filter sets | 605 nm – 625 nm, 650 nm – 720 nm |
| dwell time | 10 µs |
| objective | 60wi0x (NA1.2) |

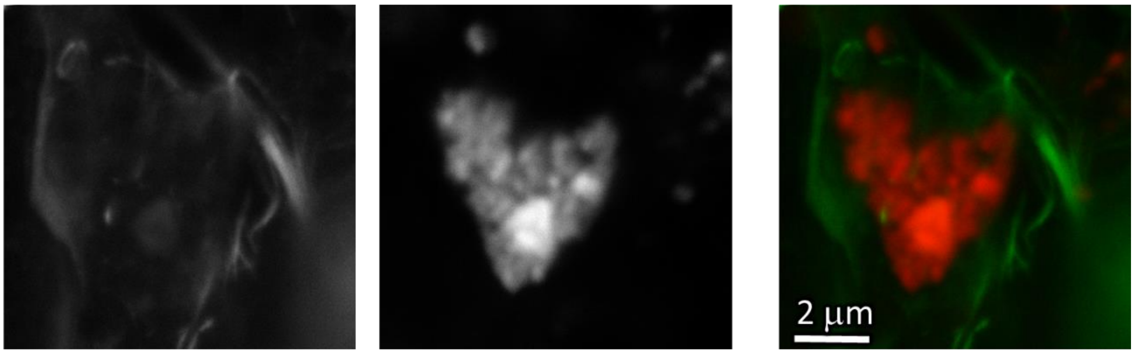

| | |
|---|---|
| Cell line | LA-4 (actin, SirActin) |
| NPs | TiO2 (Star 520S) |
| exposure | 10:1, 24h-60h A |
| imaging | xy STED, 60h |

| | |
|---|---|
| pixelsize (x,y) | 10 nm |
| FOV (x,y) | 9 x 9 µm |
| pixelsize (z) | / nm |
| FOV (z) | / µm |
| imaging time | / |
| number of frames | / |

| | |
|---|---|
| 561nm | 30% |
| 640nm | 50% |
| STED | 15% |
| filter sets | 605 nm – 625 nm, 650 nm – 720 nm |
| dwell time | 10 µs |
| objective | 60wi0x (NA1.2) |

*Figure S138: STED images of LA-4 in green and $TiO_2$ nanotubes in red channel. Cells are stained with Sir Actin label for F-actin. STED zoom – ins (XY planes in both windows) showing representative actine rings, unable to compress and excrete nanomaterial outside the cell due to Jaspankinolide treatmant.*

LA-4 actin (SirActin) | TiO2 (Star 520S) | overlay

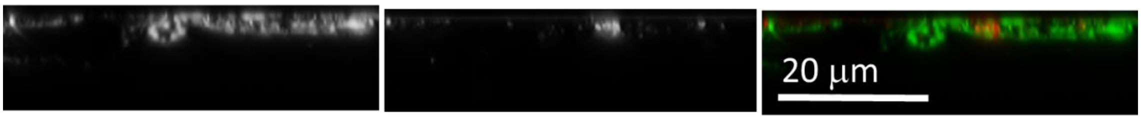

| | |
|---|---|
| Cell line | LA-4 (actin, SirActin) |
| NPs | TiO2 (Star 520S) |
| exposure | 10:1, 24h-60h |
| imaging | xz STED, 60h |

| | |
|---|---|
| pixelsize (x,y) | 40 nm |
| FOV (x,y) | 50 µm |
| pixelsize (z) | 40 nm |
| FOV (z) | 15 µm |
| imaging time | / |
| number of frames | / |

| | |
|---|---|
| 561nm | 30% |
| 640nm | 50% |
| STED | 15% |
| filter sets | 605 nm – 625 nm, 650 nm – 720 nm |
| dwell time | 10 µs |
| objective | 60wi0x (NA1.2) |

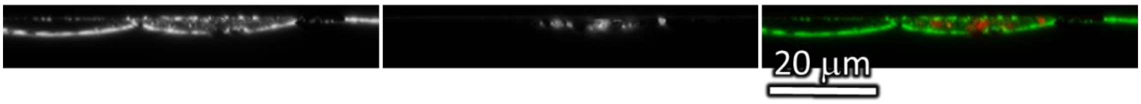

| | |
|---|---|
| Cell line | LA-4 (actin, SirActin) |
| NPs | TiO2 (Star 520S) |
| exposure | 10:1, 24h-60h |
| imaging | yz STED, 60h |

| | |
|---|---|
| pixelsize (x,y) | 30 nm |
| FOV (x,y) | 70 µm |
| pixelsize (z) | 50nm |
| FOV (z) | 30 µm |
| imaging time | / |
| number of frames | / |

| | |
|---|---|
| 561nm | 30% |
| 640nm | 50% |
| STED | 15% |
| filter sets | 605 nm – 625 nm, 650 nm – 720 nm |
| dwell time | 10 µs |
| objective | 60wi0x (NA1.2) |

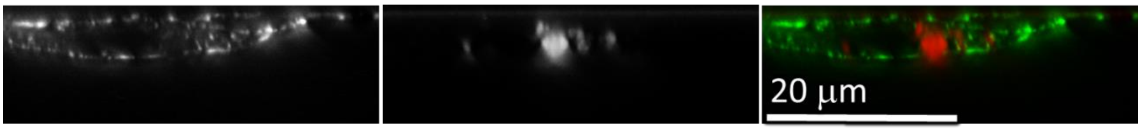

| | |
|---|---|
| Cell line | LA-4 (actin, SirActin) |
| NPs | TiO2 (Star 520S) |
| exposure | 10:1, 24h-60h |
| imaging | yz STED, 60h |

| | |
|---|---|
| pixelsize (x,y) | 40 nm |
| FOV (x,y) | 40 µm |
| pixelsize (z) | 40 nm |
| FOV (z) | 13 µm |
| imaging time | / |
| number of frames | / |

| | |
|---|---|
| 561nm | 30% |
| 640nm | 50% |
| STED | 15% |
| filter sets | 605 nm – 625 nm, 650 nm – 720 nm |
| dwell time | 10 µs |
| objective | 60wi0x (NA1.2) |

*Figure S139: STED images of LA-4 in green and $TiO_2$ nanotubes in red channel. Cells are stained with Sir Actin label for F-actin. STED images (XZ plane in upper window and two YZ planes in middle and lower window) showing representative actine rings, unable to compress and excrete nanomaterial outside the cell due to Jaspankinolide treatmant.*

LA-4 actin (SirActin) | TiO2 (Star 520S) | overlay

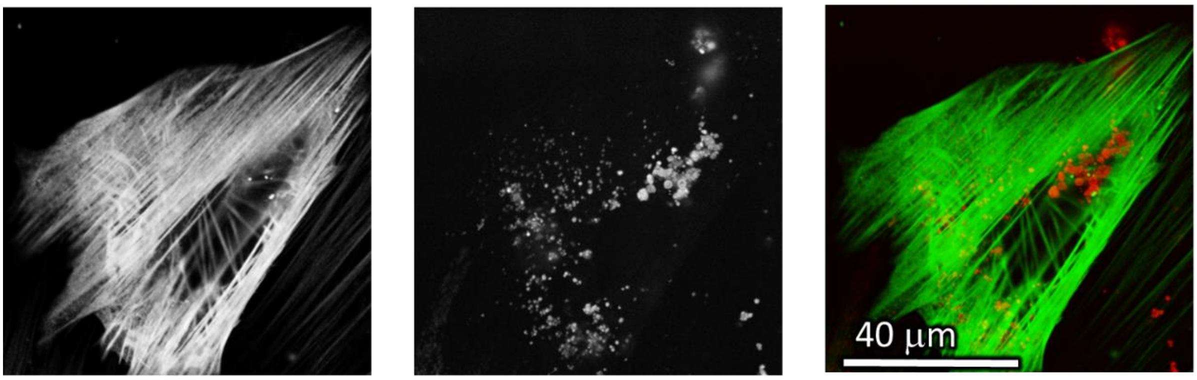

| | |
|---|---|
| Cell line | LA-4 (actin, SirActin) |
| NPs | TiO2 (Star 520S) |
| exposure | 10:1, 24h-48h<br>A |
| imaging | XY confocal, 48h |

| | |
|---|---|
| pixelsize (x,y) | 150 nm |
| FOV (x,y) | 80 µm |
| pixelsize (z) | / nm |
| FOV (z) | / µm |
| imaging time | / |
| number of frames | / |

| | |
|---|---|
| 561nm | 30% |
| 640nm | 30% |
| STED | /% |
| filter sets | 605 nm – 625 nm,<br>650 nm – 720 nm |
| dwell time | 10 µs |
| objective | 60wi0x (NA1.2) |

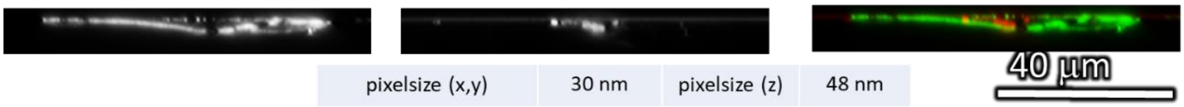

| | | | |
|---|---|---|---|
| pixelsize (x,y) | 30 nm | pixelsize (z) | 48 nm |
| FOV (x,y) | 60 µm | FOV (z) | 12 µm |

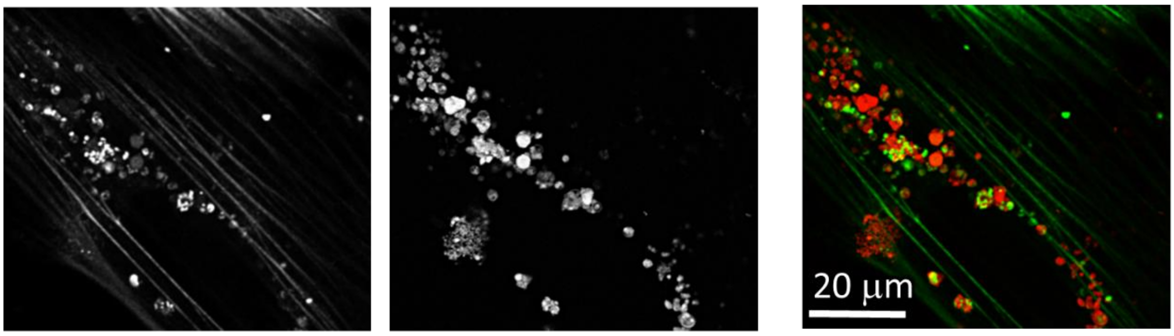

| | |
|---|---|
| Cell line | LA-4 (actin, SirActin) |
| NPs | TiO2 (Star 520S) |
| exposure | 10:1, 24h-48h<br>A |
| imaging | XYZ confocal, 48h |

| | |
|---|---|
| pixelsize (x,y) | 210 nm |
| FOV (x,y) | 70 x 62<br>µm |
| pixelsize (z) | 30 nm |
| FOV (z) | 7 µm |
| imaging time | / |
| number of frames | / |

| | |
|---|---|
| 561nm | 30% |
| 640nm | 30% |
| STED | /% |
| filter sets | 605 nm – 625 nm,<br>650 nm – 720 nm |
| dwell time | 10 µs |
| objective | 60wi0x (NA1.2) |

*Figure S140: STED images of LA-4 in green and $TiO_2$ nanotubes in red channel. Cells are stained with Sir Actin label for F-actin. Confocal image of large FoV in upper window, STED zoom – in (XZ plane) in the middle window and one sXY plane of a whole Z-stack (3D)in the lower window showing representative actine rings, unable to compress and excrete nanomaterial outside the cell due to Jaspankinolide treatmant*

Link to 3D

- http://lbfnanobiodatabase.ijs.si/file/data/cauliflowerpaper/20190711_e01_m16_s10_LA-4__1um_SA_and_10_to_1_TiO2_Star520s_Omega_vesicles_3D.mp4

*Movie S19: Representative actine rings, unable to compress and excrete nanomaterial outside the cell due to Jaspankinolide treatmant. Cells are stained with Sir Actin label for F-actin (green) and were exposed to 10:1 surface dose of $TiO_2$ nanotubes (Star 520 SXP, red).*

#### S3d – Nanoparticle actin interaction

##### Main message

Uptake of a bigger nanomaterial aggregate by the cell within one hour of exposure.

Supporting raw and analysed data:

Figure S141-Figure S144

##### Materials and methods

- Main experiment:
    - LA-4 cells were seeded @60% confluence in an Ibidi #1.5H µ-Dish
    - after 24 hours cells were incubated with 1 µm Sir Actin for 2 hours in the incubator at 37°C, 5% CO2 afterwards they were washed with 3x400 ml LCIS
    - After 2h of incubation 35 µl freshly filtered 1 mg/ml $TiO_2$-40-ATTO 594 in 100x dcb was added directly to the cells (in 400 µL medium) and mixed to achieve 10:1 surface dose and observed in 400 µL LCIS at microscope stage heated on 37°C

- Analysis:
    - threshold has been set to 1 count and brightness has been adjusted for maximal visibility
    - 0,5 pixel Gaussian blur and sharpening algorithm were applied as well

##### Experiment names

- Main experiment name:
    - 20190122_e01_m02_LA-4_SirActin and 10 to 1_ TiO2_ATTO 594_uptake of NM

- Supplement Experiment names:
    - 20190127_e01 m02_LA-4_SA and 10 to 1 TiO2_ATTO 594_uptake
    - 20190129_e01m01s01_Native LA-4_SA and 10 to 1 TiO2_ATTO 594_large FOV
    - 20190129_e01m01s04_Native LA-4_SA_static and parallel filaments
    - 20190129_e01m01s05_Native LA-4_SA_filaments_STED

##### Controls and statistics

Uptake of $TiO_2$ nanotubes immediately after exposure of the cells:

Figure S141

Control, unexposed cells labelled with Sir-Actin:

Figure S142-Figure S143

Uptake of $TiO_2$ nanotubes immediately after exposure of the cells:

Figure S144

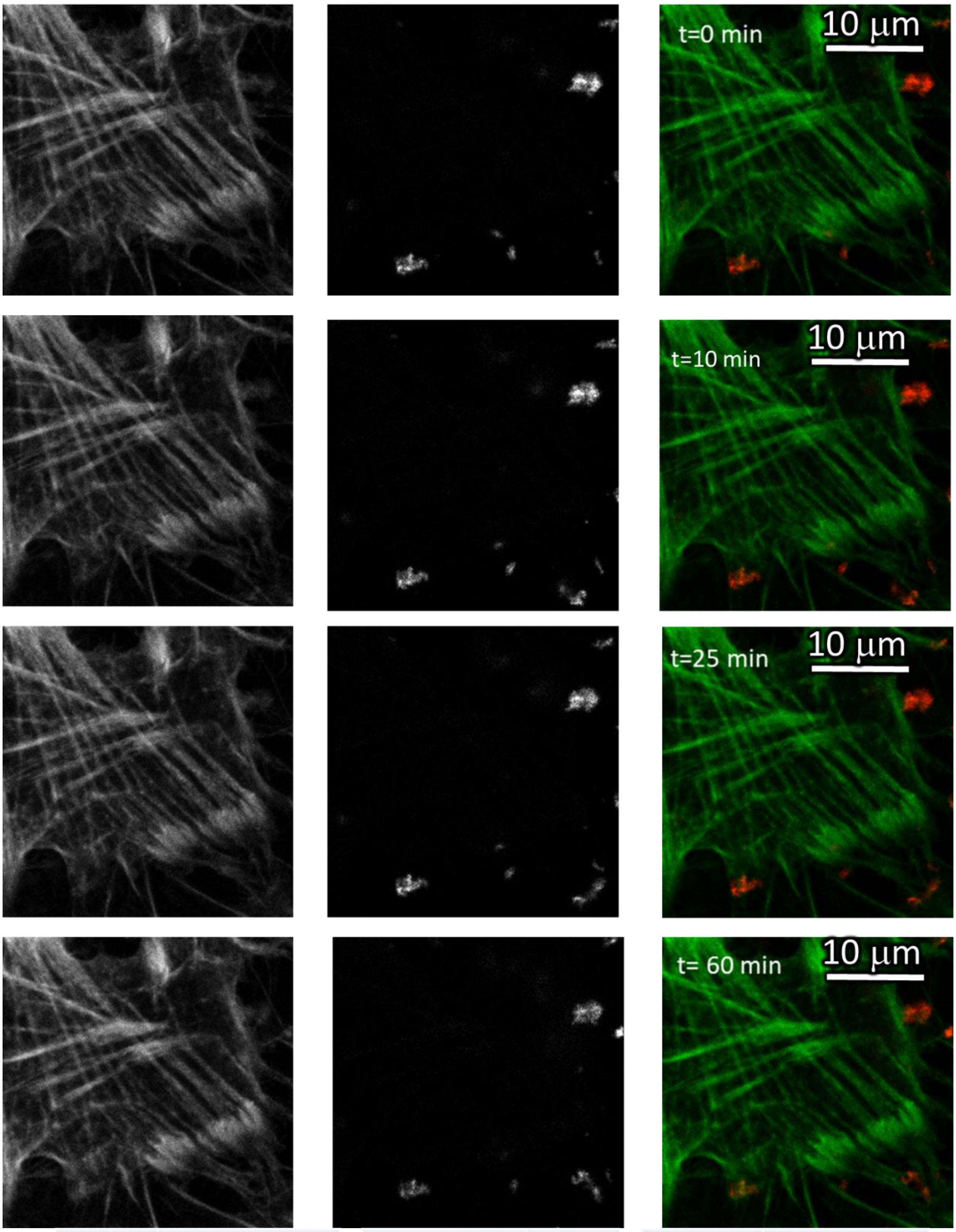

| | |
|---|---|
| Cell line | LA-4 (actin, Sir Actin) |
| NPs | TiO2 (ATTO 594) |
| exposure | 10:1, 0h-1h |
| imaging | xyt STED, 0h-1h |

| | |
|---|---|
| pixelsize (x,y) | 100 nm |
| FOV (x,y) | 30 mm |
| pixelsize (z) | / nm |
| FOV (z) | / mm |
| imaging time | 20 min |
| number of frames | 511 |

| | |
|---|---|
| 561nm | 25% |
| 640nm | 25% |
| STED | 14% |
| filter sets | 605 nm – 625 nm, 650 nm – 720 nm |
| dwell time | 10 ms |
| objective | wi60x (NA1.2) |

*Figure S141: STED images of LA-4 in green and $TiO_2$ nanotubes in red channel. Cells are stained with Sir Actin label for F-actin. Images are four different time points with 10, 15 and 25 min increments, respectively. Actin actively engulfs $TiO_2$ nanotubes and pulls them towards cell interior.*

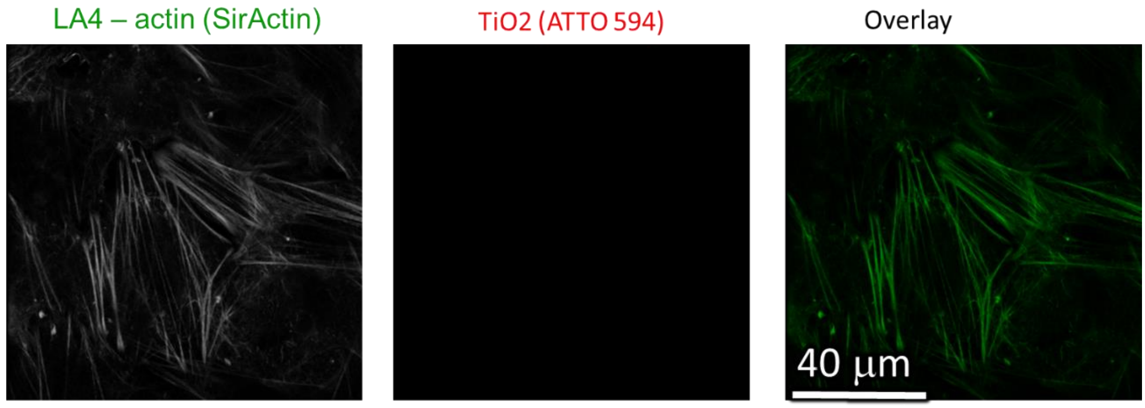

| | |
|---|---|
| Cell line | LA-4 (actin, Sir Actin) |
| NPs | TiO2 (ATTO 594) |
| exposure | CONTROLE |
| imaging | Xy confocal, 0h-1h |

| | |
|---|---|
| pixelsize (x,y) | 200 nm |
| FOV (x,y) | 100 mm |
| pixelsize (z) | / nm |
| FOV (z) | / mm |
| imaging time | / min |
| number of frames | / |

| | |
|---|---|
| 561nm | /% |
| 640nm | 13% |
| STED | /% |
| filter sets | 605 nm – 625 nm, 650 nm – 720 nm |
| dwell time | 10 ms |
| objective | wi60x (NA1.2) |

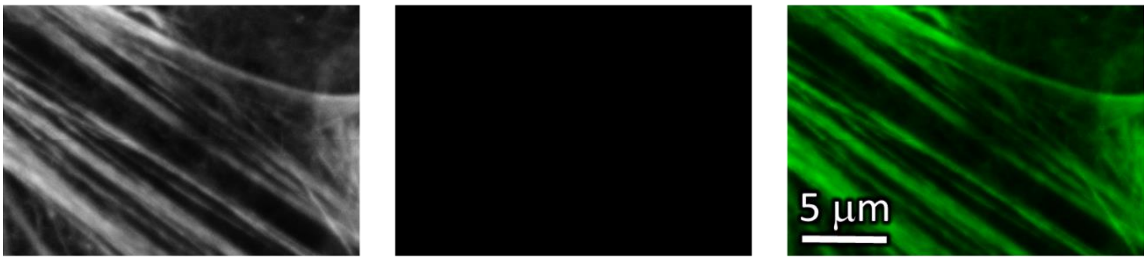

| | |
|---|---|
| Cell line | LA-4 (actin, Sir Actin) |
| NPs | TiO2 (ATTO 594) |
| exposure | CONTROLE |
| imaging | xy STED |

| | |
|---|---|
| pixelsize (x,y) | 15 nm |
| FOV (x,y) | 20 x 14 mm |
| pixelsize (z) | / nm |
| FOV (z) | / mm |
| imaging time | / min |
| number of frames | / |

| | |
|---|---|
| 561nm | /% |
| 640nm | 25% |
| STED | 15% |
| filter sets | 605 nm – 625 nm, 650 nm – 720 nm |
| dwell time | 10 ms |
| objective | wi60x (NA1.2) |

*Figure S142: STED images of LA-4 in green and $TiO_2$ nanotubes in red channel. Cells are stained with Sir Actin label for F-actin. Cells which have been grown under the same conditions as all experimental cells but haven't been exposed to $TiO_2$ nanotubes.*

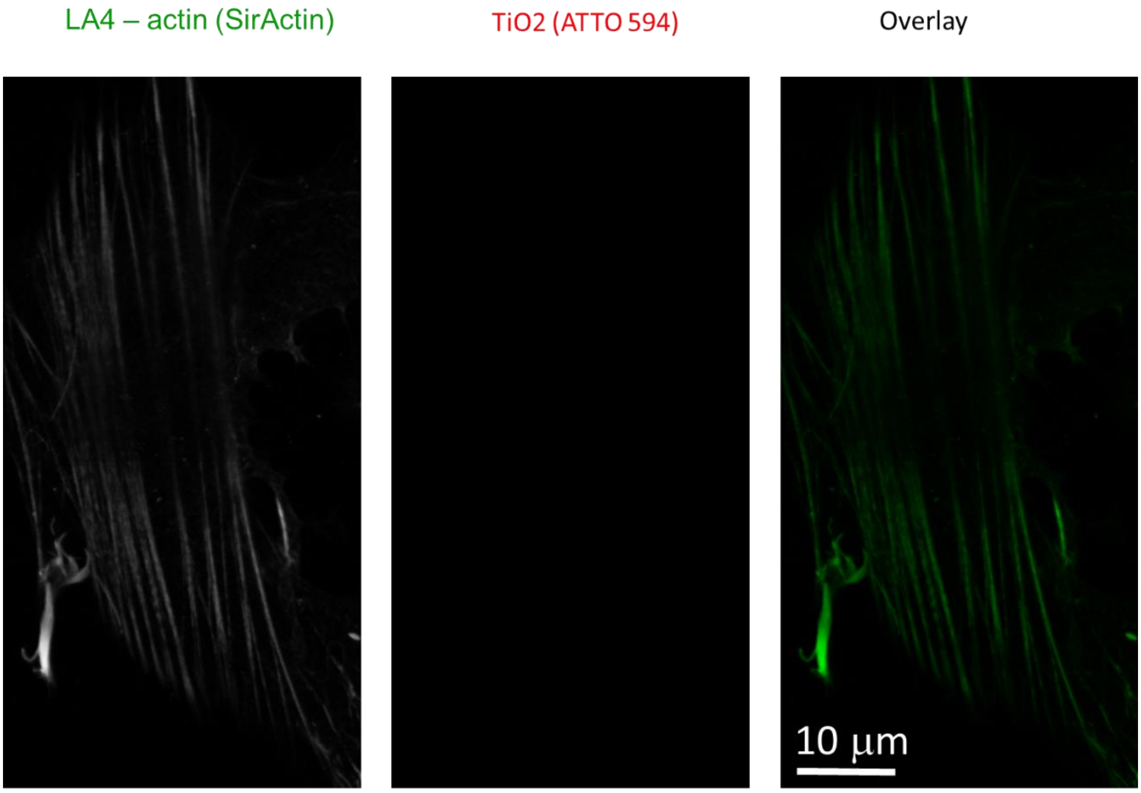

| | | | | | |
|---|---|---|---|---|---|
| Cell line | LA-4 (actin, Sir Actin) | pixelsize (x,y) | 30 nm | 561nm | /% |
| NPs | TiO2 (ATTO 594) | FOV (x,y) | 36 x 72 mm | 640nm | 15% |
| exposure | CONTROLE | pixelsize (z) | / nm | STED | 9% |
| imaging | xy STED | FOV (z) | / mm | filter sets | 605 nm – 625 nm, 650 nm – 720 nm |
| | | imaging time | / min | dwell time | 10 ms |
| | | number of frames | / | objective | wi60x (NA1.2) |

*Figure S143: STED images of LA-4 in green and $TiO_2$ nanotubes in red channel. Cells are stained with Sir Actin label for F-actin. Cells which have been grown under the same conditions as all experimental cells but haven't been exposed to $TiO_2$ nanotubes.*

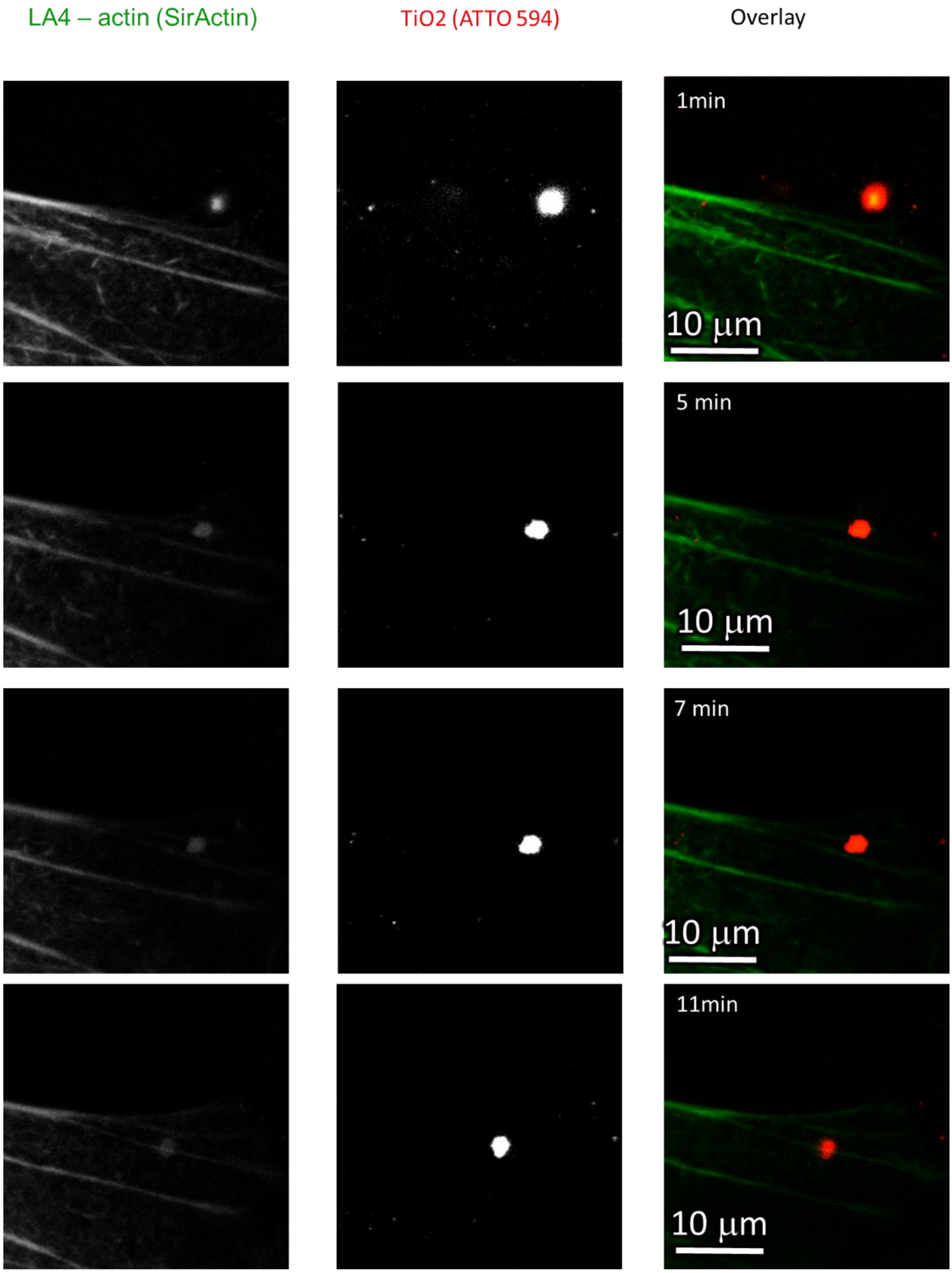

| | |
|---|---|
| Cell line | LA-4 (actin, Sir Actin) |
| NPs | TiO2 (ATTO 594) |
| exposure | 10:1, 0h-1h |
| imaging | xyt STED, 0h-1h |

| | |
|---|---|
| pixelsize (x,y) | 80 nm |
| FOV (x,y) | 33 x 23 mm |
| pixelsize (z) | / nm |
| FOV (z) | / mm |
| imaging time | 25 min |
| number of frames | 221 |

| | |
|---|---|
| 561nm | 20% |
| 640nm | 25% |
| STED | 12% |
| filter sets | 605 nm – 625 nm, 650 nm – 720 nm |
| dwell time | 10 ms |
| objective | wi60x (NA1.2) |

*Figure S144: STED images of LA-4 in green and $TiO_2$ nanotubes in red channel. Cells are stained with Sir Actin label for F-actin. Images are four different time points with 5, 2 and 4 min increments, respectively. Actin actively orients towards the agregate of $TiO_2$ nanotubes, which is then pulled into the cell interior.*

#### Link to time-lapse

- Main experiment
    - [http://lbfnanobiodatabase.ijs.si/file/data/cauliflowerpaper/e01_m02_LA-4_SA_uptake_of_TiO2_ATTO_594_25us.gif](http://lbfnanobiodatabase.ijs.si/file/data/cauliflowerpaper/e01_m02_LA-4_SA_uptake_of_TiO2_ATTO_594_25us.gif)

*Movie S20: Actin filaments (green) folding and uptaking large TiO2 aggregate (red)*

- Supplementary experiment
    - [http://lbfnanobiodatabase.ijs.si/file/data/cauliflowerpaper/20190127_e01_m02_LA-4_SA_uptake_TiO2_ATTO_594_50us.gif](http://lbfnanobiodatabase.ijs.si/file/data/cauliflowerpaper/20190127_e01_m02_LA-4_SA_uptake_TiO2_ATTO_594_50us.gif)

*Movie S21: Uptake of a single TiO2 aggregate (red). Observe dynamics of small actin filaments (green) branching from the main fibres.*

#### S3e – Actin branching

##### Main message

Branching of actin filaments (usually parallel in cells which are not engaged in any activity) after incubation with $TiO_2$ for more than 24h.

Supporting raw and analysed data:

Figure S145-Figure S147

##### Materials and methods

- Main experiment:
  - LA-4 cells were seeded @60% confluence in an Ibidi #1.5H µ-Dish and incubated at 37°C and 5% CO2 for 24 hours
  - After incubation 100 nM of SirActin and 35 µl freshly filtered 1 mg/ml $TiO_2$-40-ATTO 594 in 100x dcb was added directly to the cells (in 400 µL medium) and mixed to achieve 10:1 surface dose and returned into the incubator
  - Cells were observed 12 hours later in 400 µL medium with label at the microscope stage heated at 37°C for 2h

- Supplement experiments
  - Samples were prepared the same way and images were processed the same way

- Analysis (Image J):
  - For STED images in both channels sharpening algorithm ($\sigma = 2$, $c = 0,5$) was used to gain For confocal images Gaussian Blur (0,5 pixel) was applied to reduce grains

##### Experiment names

- Main experiment name:
  - 20190118_e01_m03_LA-4_0.1 um SA and 10 to 1TiO2_ATTO 594_overnight incubation_Actin branching_STED and confocal
  - Supplement Experiment names:
  - 20190118_e01_m01_LA-4_0.1 um SA and 10 to 1TiO2_ATTO 594_overnight incubation_Actin branching_STED and confocal
  - 20190118_e01_m07_LA-4_0.1 um SA and 10 to 1TiO2_ATTO 594_overnight incubation_Actin branching_STED and confocal

##### Controls and statistics

Branching of actin cytoskeleton after the exposure and incubation with $TiO_2$ nanotubes:

Figure S145-Figure S147

LA-4 membrane (CellMaskOrange) | TiO2 (Alexa647) | overlay

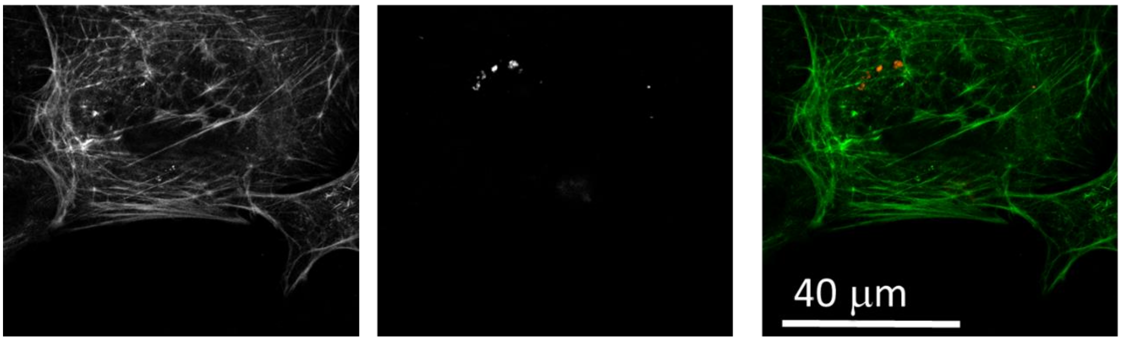

| | |
|---|---|
| Cell line | LA-4 (actin, SirActin) |
| NPs | TiO2 (ATTO 594) |
| exposure | 10:1, 24h-36h |
| imaging | xy confocal, 36h |

| | |
|---|---|
| pixelsize (x,y) (confocal) | 200 nm |
| FOV (x,y) | 80 µm |
| pixelsize (z) | / nm |
| FOV (z) | / µm |
| imaging time | / |
| number of frames | / |

| | |
|---|---|
| 561nm | 5% |
| 640nm | 10% |
| STED | / |
| filter sets | 605 nm – 625 nm, 650 nm – 720 nm |
| dwell time | 10 µs |
| objective | wi60x (NA 1.2) |

LA-4 membrane (CellMaskOrange) | TiO2 (Alexa647) | overlay

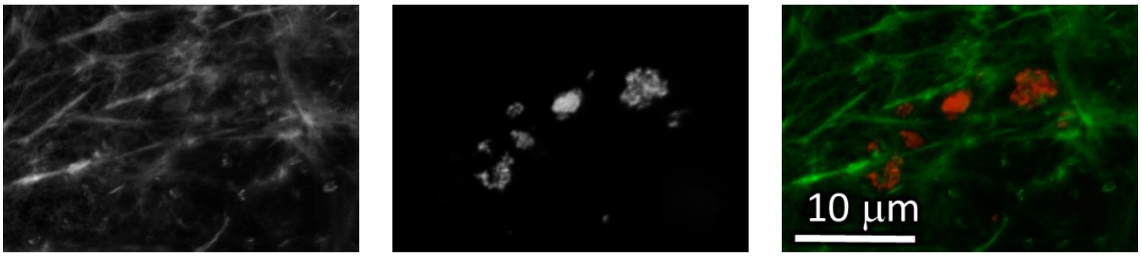

| | |
|---|---|
| Cell line | LA-4 (actin, SirActin) |
| NPs | TiO2 (ATTO 594) |
| exposure | 10:1, 24h-36h |
| imaging | xy STED, 36h |

| | |
|---|---|
| pixelsize (x,y) (confocal) | 25 nm |
| FOV (x,y) | 29 * 22 µm |
| pixelsize (z) | / nm |
| FOV (z) | / µm |
| imaging time | / |
| number of frames | / |

| | |
|---|---|
| 561nm | 10% |
| 640nm | 15% |
| STED | 15% |
| filter sets | 605 nm – 625 nm, 650 nm – 720 nm |
| dwell time | 10 µs |
| objective | wi60x (NA 1.2) |

*Figure S145: Confocal and STED images of LA-4 in green and $TiO_2$ nanotubes in red channel. Cells are stained with Sir Actin label for F-actin. Confocal image of larger FoV in the upper window shows branching of actin filaments after overnight incubation with $TiO_2$ nanotubes. In the lower window is the STED zoom-in of the former confocal image, showing more intricately actine structure and interaction with $TiO_2$ aggregates.*

LA-4 membrane (CellMaskOrange) | TiO2 (Alexa647) | overlay

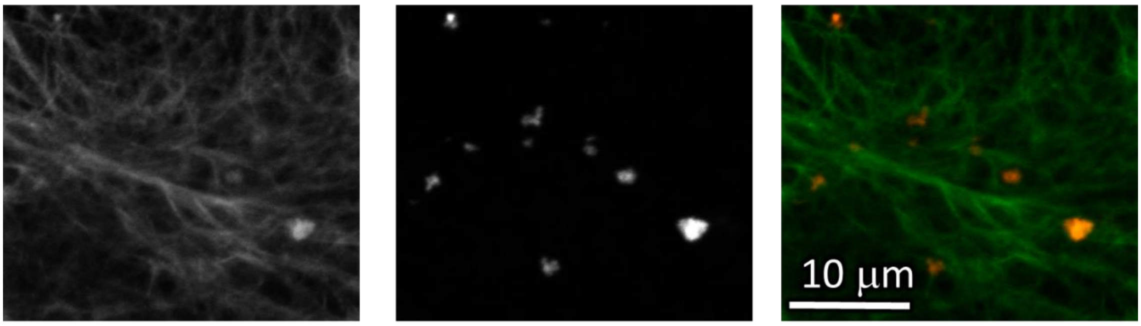

| | |
|---|---|
| Cell line | LA-4 (actin, SirActin) |
| NPs | TiO2 (ATTO 594) |
| exposure | 10:1, 24h-36h |
| imaging | xy STED, 36h |

| | |
|---|---|
| pixelsize (x,y) (confocal) | 25 nm |
| FOV (x,y) | 29 * 22 µm |
| pixelsize (z) | / nm |
| FOV (z) | / µm |
| imaging time | / |
| number of frames | / |

| | |
|---|---|
| 561nm | 10% |
| 640nm | 15% |
| STED | 15% |
| filter sets | 605 nm – 625 nm, 650 nm – 720 nm |
| dwell time | 10 µs |
| objective | wi60x (NA 1.2) |

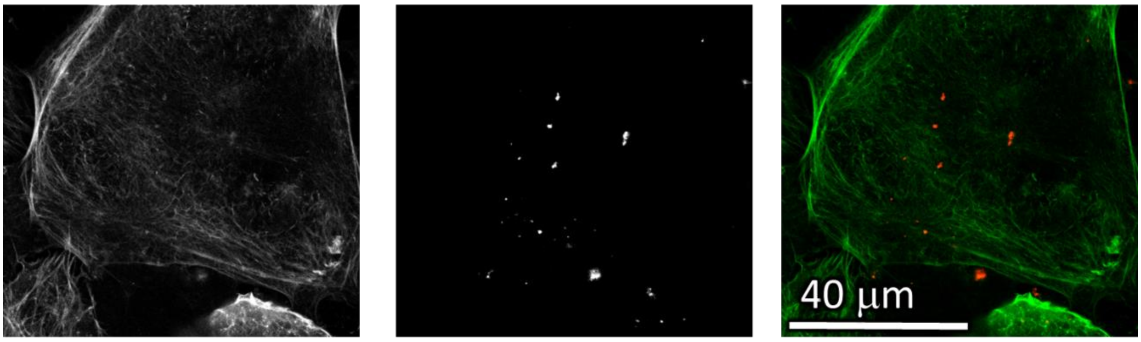

| | |
|---|---|
| Cell line | LA-4 (actin, SirActin) |
| NPs | TiO2 (ATTO 594) |
| exposure | 10:1, 24h-36h |
| imaging | xy confocal, 36h |

| | |
|---|---|
| pixelsize (x,y) (confocal) | 200 nm |
| FOV (x,y) | 80 µm |
| pixelsize (z) | / nm |
| FOV (z) | / µm |
| imaging time | / |
| number of frames | / |

| | |
|---|---|
| 561nm | 5% |
| 640nm | 10% |
| STED | / |
| filter sets | 605 nm – 625 nm, 650 nm – 720 nm |
| dwell time | 10 µs |
| objective | wi60x (NA 1.2) |

*Figure S146: Confocal and STED images of LA-4 in green and $TiO_2$ nanotubes in red channel. Cells are stained with Sir Actin label for F-actin. Confocal image of larger FoV in the lower window shows branching of actin filaments after overnight incubation with $TiO_2$ nanotubes. In the upper window is the STED zoom-in of the former confocal image, showing more intricately actine structure and interaction with $TiO_2$ aggregates.*

LA-4 membrane (CellMaskOrange) | TiO2 (Alexa647) | overlay

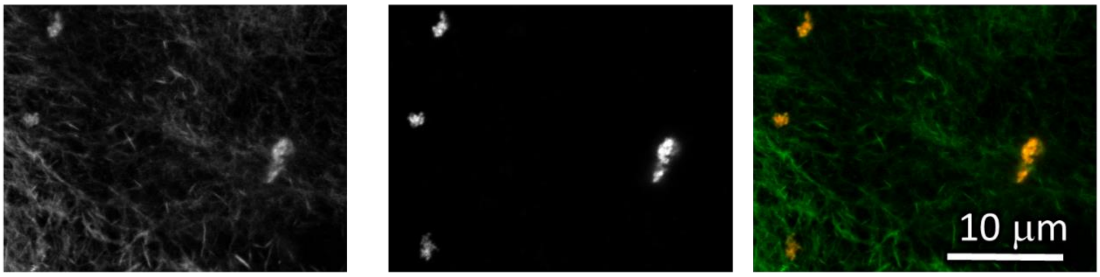

| | |
|---|---|
| Cell line | LA-4 (actin, SirActin) |
| NPs | TiO2 (ATTO 594) |
| exposure | 10:1, 24h-36h |
| imaging | xy STED, 36h |

| | |
|---|---|
| pixelsize (x,y) (confocal) | 25 nm |
| FOV (x,y) | 29 * 22 μm |
| pixelsize (z) | / nm |
| FOV (z) | / μm |
| imaging time | / |
| number of frames | / |

| | |
|---|---|
| 561nm | 10% |
| 640nm | 15% |
| STED | 15% |
| filter sets | 605 nm – 625 nm, 650 nm – 720 nm |
| dwell time | 10 μs |
| objective | wi60x (NA 1.2) |

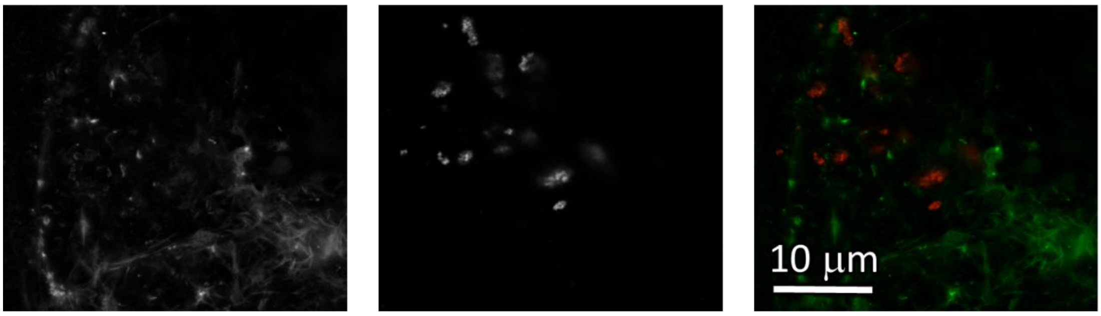

| | |
|---|---|
| Cell line | LA-4 (actin, SirActin) |
| NPs | TiO2 (ATTO 594) |
| exposure | 10:1, 24h-36h |
| imaging | xy STED, 36h |

| | |
|---|---|
| pixelsize (x,y) (confocal) | 25 nm |
| FOV (x,y) | 35 * 31 μm |
| pixelsize (z) | / nm |
| FOV (z) | / μm |
| imaging time | / |
| number of frames | / |

| | |
|---|---|
| 561nm | 10% |
| 640nm | 15% |
| STED | 15% |
| filter sets | 605 nm – 625 nm, 650 nm – 720 nm |
| dwell time | 10 μs |
| objective | wi60x (NA 1.2) |

*Figure S147: STED images of LA-4 in green and $TiO_2$ nanotubes in red channel. Cells are stained with Sir Actin label for F-actin. In both windows are STED images showing branched actine filaments and interaction with $TiO_2$ aggregates after the exposure and incubation with $TiO_2$ nanotubes. For non-branched, control, experiment see S3d – Nanoparticle actin interaction.*

#### S3f – Actin in cauliflowers

##### Main message

After exocytosis of nanomaterial, some remnants of actin can be seen in cauliflower-like formations.

Supporting raw and analysed data:

Figure S148-Figure S151

##### Materials and methods

- Main experiment:
  - LA-4 cells were seeded @60% confluence in an Ibidi #1.5H μ-Dish in the cell medium
  - after 24 hours 35 ml of freshly filtered 1 mg/ml $TiO_2$-17-Alexa 647 in 100x diluted bicarbonate buffer was added directly to the cells (in 400 μL medium) and mixed to achieve 10:1 surface dose.
  - After 48 hours cells were incubated with 100 nM Sir Actin overnight in the incubator at 37°C, 5% CO2. According to manufacturer of the label, it is not necessary to wash it if there is no background, so we imaged the cells without washing them. Cells were imaged on the microscope stage heated on 37°C.

- Images analysed with ImageJ
  - Signal was multiplied by 1.8x
  - Sharpening algorithm ($\sigma = 3$, $c = 0,3$) was used

##### Experiment names

- Main experiment name:
  - 20190627_e05 m02 s02_1um SA & TiO2 S520_Remnants of actin in cauliflower_ STED and confocal

- Supplement Experiment names:
  - 20190619_e01 m01 s01 NEG CONTROLE for10 mM MBCD LA-4 Actin and 10 - 1 S520 TiO2_actin in cauliflowers_STED
  - 20190619_e01 m02 s01 NEG CONTROLE for10 mM MBCD LA-4 Actin and 10 - 1 S520 TiO2_actin in cauliflowers_confocal
  - 20190619_e07 m01 s01_LA-4 SA and 10 - 1 S520 TiO2_actin in cauliflowers_confocal and STED
  - 20190619_e04 m01 s01_100 μM Resveratrol LA-4_SA and 10 - 1 S520 TiO_actin in caulflowers_STED and confocal
  - 20190627_e05 m03 s03 LA-4 0.1um SA & 10 - 1 TiOs S520 overnightincubation 3D and STED

##### Controls and statistics

Actin filaments are exocytosed together with the nanomaterial:

Figure S148-Figure S150

Actin filaments are exocytosed together with the nanomaterial even in cells treated with Resveratrol

Figure S151

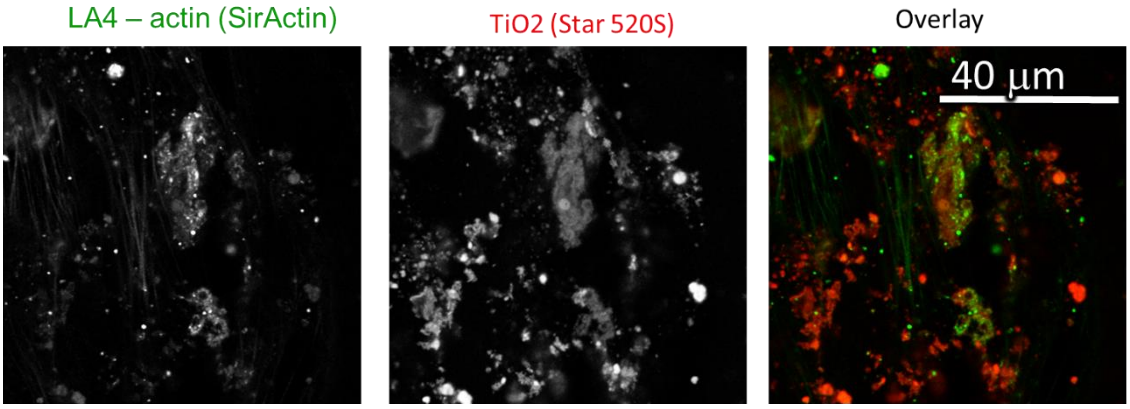

| | |
|---|---|
| Cell line | LA-4 (actin, Sir Actin) |
| NPs | TiO2 (Star 520) |
| exposure | 10:1, 24h-48h |
| imaging | xy confocal, 48h |

| | |
|---|---|
| pixelsize (x,y) | 200 nm |
| FOV (x,y) | 80 um |
| pixelsize (z) | /nm |
| FOV (z) | /um |
| imaging time | /min |
| number z-stacks | |

| | |
|---|---|
| 561nm | 30% |
| 640nm | 50% |
| STED | 0% |
| filter sets | 605 nm – 625 nm, 650 nm – 720 nm |
| dwell time | 10 ms |
| objective | wi60x (NA1.2) |

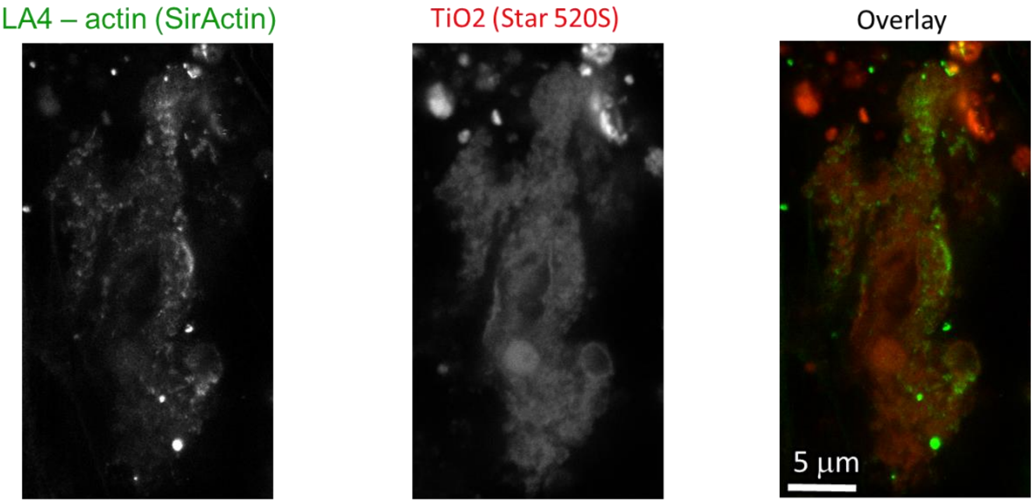

| | |
|---|---|
| Cell line | LA-4 (actin, Sir Actin) |
| NPs | TiO2 (Star 520) |
| exposure | 10:1, 24h-48h |
| imaging | xy STED, 48h |

| | |
|---|---|
| pixelsize (x,y) | 30x30 nm |
| FOV (x,y) | 18x32 um |
| pixelsize (z) | /nm |
| FOV (z) | /um |
| imaging time | / min |
| number z-stacks | / |

| | |
|---|---|
| 561nm | 30% |
| 640nm | 60% |
| STED | 30% |
| filter sets | 605 nm – 625 nm, 650 nm – 720 nm |
| dwell time | 10 ms |
| objective | wi60x (NA1.2) |

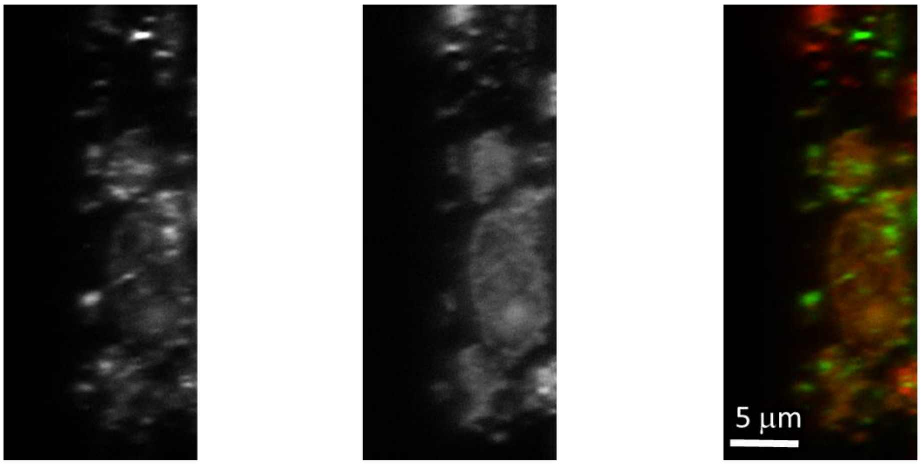

| | |
|---|---|
| Cell line | LA-4 (actin, Sir Actin) |
| NPs | TiO2 (Star 520) |
| exposure | 10:1, 24h-48h |
| imaging | yz STED, 48h |

| | |
|---|---|
| pixelsize (x,y) | 50 nm |
| FOV (x,y) | 33 um |
| pixelsize (z) | 54 nm |
| FOV (z) | 15 um |
| imaging time | / min |
| number z-stacks | / |

| | |
|---|---|
| 561nm | 30% |
| 640nm | 60% |
| STED | 15% |
| filter sets | 605 nm – 625 nm, 650 nm – 720 nm |
| dwell time | 10 ms |
| objective | wi60x (NA1.2) |

*Figure S148: Confocal and STED images of LA-4 in green and $TiO_2$ nanotubes in red channel. Cells are stained with Sir Actin label for F-actin. Confocal image of larger FoV in the upper window shows actin cytoskeleton of few cells with a lot of nanomaterial on the surface and one large cauliflower-like structure. In the middle window is STED zoom-in (XY plane) of the cauliflower-like structure form the upper window where we can clearly see actin filaments (green) intertwined with nanomaterial (red). In the lower window we see the STED zoom-in (YZ plane) of the same cauliflower-like structure shown in the middle window. Again, actin filaments are noticable in the cauliflower-like structure bulk.*

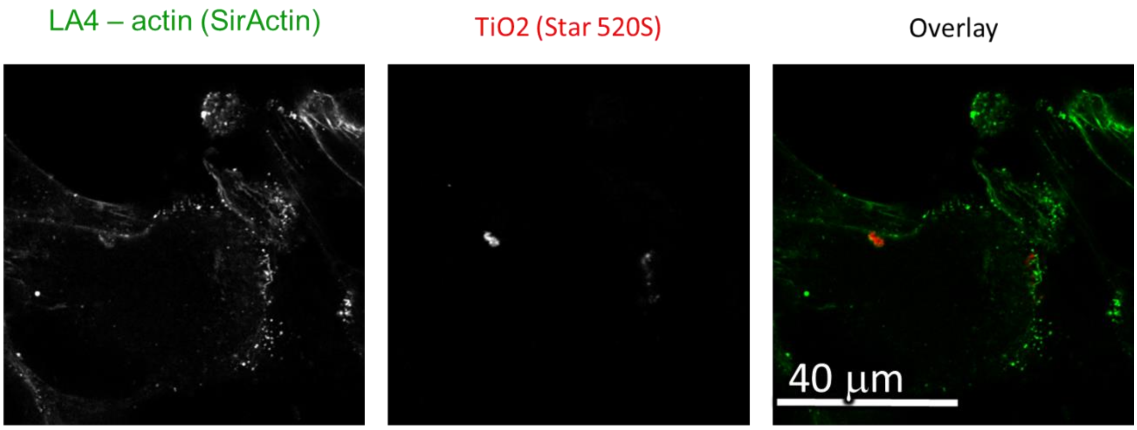

| | |
|---|---|
| Cell line | LA-4 (actin, 0,1 µM SirActin 12h) |
| NPs | TiO2 (Star 520) |
| exposure | 10:1, 24h-48h |
| imaging | xy confocal, 48h |

| | |
|---|---|
| pixelsize (x,y) | 200 nm |
| FOV (x,y) | 80 um |
| pixelsize (z) | /nm |
| FOV (z) | /um |
| imaging time | /min |
| number z-stacks | / |

| | |
|---|---|
| 561nm | 20% |
| 640nm | 20% |
| STED | 0% |
| filter sets | 605 nm – 625 nm, 650 nm – 720 nm |
| dwell time | 10 ms |
| objective | wi60x (NA1.2) |

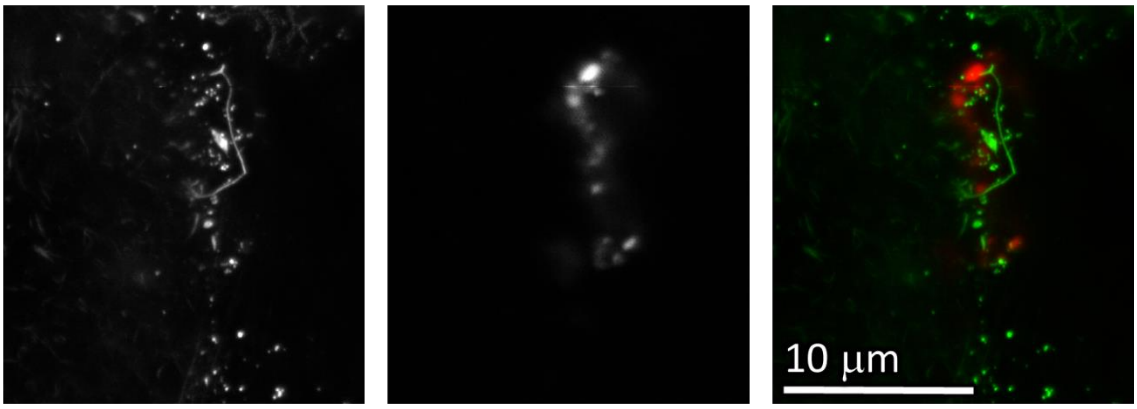

| | |
|---|---|
| Cell line | LA-4 (actin, Sir Actin) |
| NPs | TiO2 (Star 520) |
| exposure | 10:1, 48h-24h |
| imaging | xy STED, 48h |

| | |
|---|---|
| pixelsize (x,y) | 15 nm |
| FOV (x,y) | 19x21 um |
| pixelsize (z) | /nm |
| FOV (z) | /um |
| imaging time | /min |
| number z-stacks | / |

| | |
|---|---|
| 561nm | 30% |
| 640nm | 40% |
| STED | 15% |
| filter sets | 605 nm – 625 nm, 650 nm – 720 nm |
| dwell time | 10 ms |
| objective | wi60x (NA1.2) |

*Figure S149: Confocal and STED images of LA-4 in green and $TiO_2$ nanotubes in red channel. Cells are stained with Sir Actin label for F-actin. Confocal image of larger FoV in the upper window shows actin cytoskeleton of two cells with some nanomaterial on the surface and one small cauliflower-like structure (red aggregate). In the lower window is STED zoom-in (XY plane) of the cauliflower-like structure form the upper window where we can clearly see actin filaments (green) sitting on the top of nanomaterial (red).*

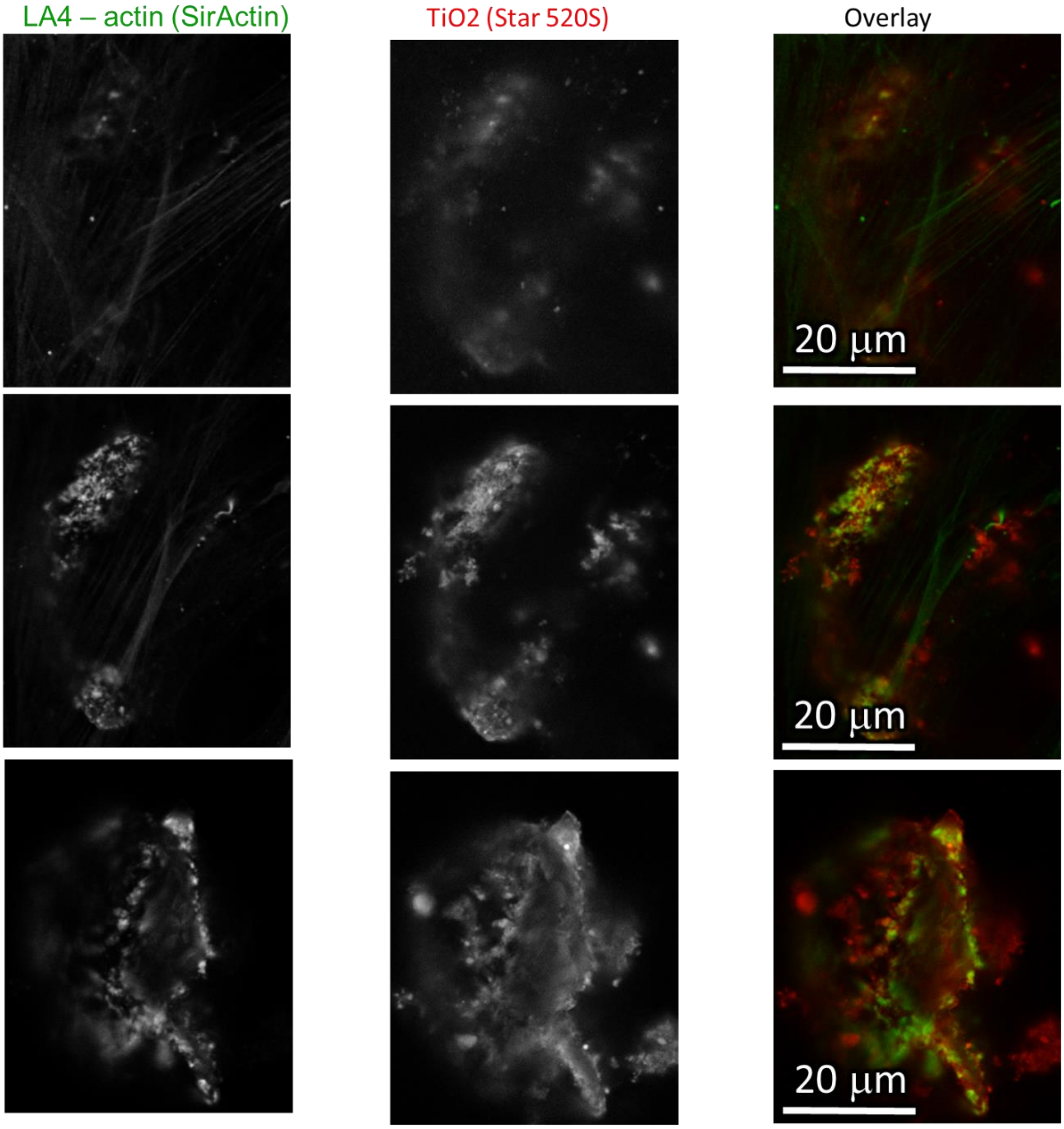

| | |
|---|---|
| Cell line | LA-4 (actin, Sir Actin) |
| NPs | TiO2 (Star 520) |
| exposure | 10:1, 24h-48h |
| imaging | xyz STED, 48h |

| | |
|---|---|
| pixelsize (x,y) | 100 nm |
| FOV (x,y) | 43x53 um |
| pixelsize (z) | 100 nm |
| FOV (z) | 15 um |
| imaging time | /min |
| number z-stacks | 149 |

| | |
|---|---|
| 561nm | 30% |
| 640nm | 40% |
| STED | 15% |
| filter sets | 605 nm – 625 nm, 650 nm – 720 nm |
| dwell time | 10 ms |
| objective | wi60x (NA1.2) |

*Figure S150: STED images of LA-4 in green and $TiO_2$ nanotubes in red channel. Cells are stained with Sir Actin label for F-actin. Three different XY plane slices from a whole Z-stack (3D) of a huge cauliflower-like structure sitting on the surface of the LA-4 cell (green cytoskeleton).*

Link to 3D

- http://lbfnanobiodatabase.ijs.si/file/data/cauliflowerpaper/20190627_e05m03s03_LA-4_0.1_uM_SA_and_10_to_1_TiO2_StarS520_remnants_of_actin_in_cauliflower_from_3D.mp4

*Movie 22: 3D rendering of the actin remains (green) in a large cauliflower on top of LA-4 epithelial cell:*

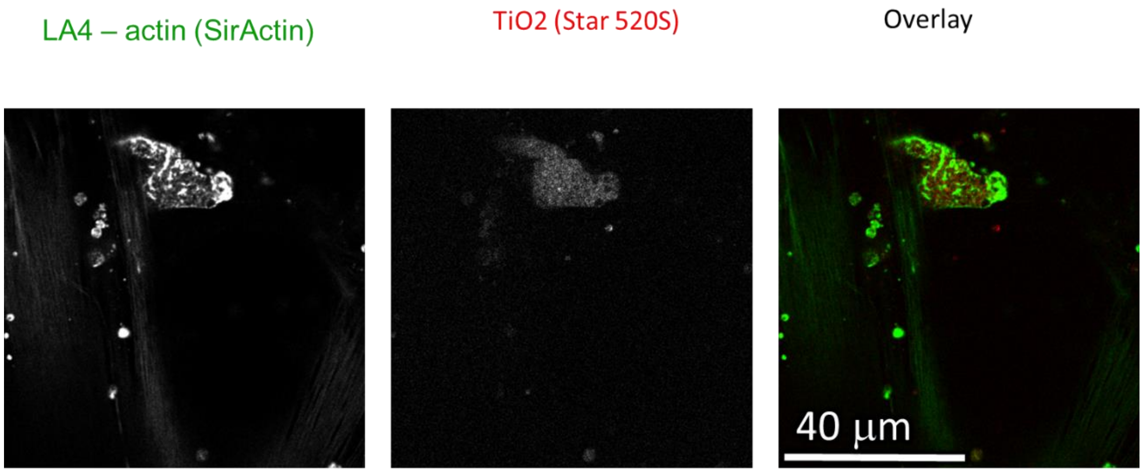

| | |
|---|---|
| Cell line | LA-4 (actin, Sir Actin) |
| NPs | TiO2 (Star 520) |
| exposure | 10:1, 24h-48h & 100 mM Resveratrol |
| imaging | xy confocal, 48h |

| | |
|---|---|
| pixelsize (x,y) | 200 nm |
| FOV (x,y) | 80x80um |
| pixelsize (z) | /nm |
| FOV (z) | /um |
| imaging time | /min |
| number z-stacks | / |

| | |
|---|---|
| 561nm | 20% |
| 640nm | 20% |
| STED | 0% |
| filter sets | 605 nm – 625 nm, 650 nm – 720 nm |
| dwell time | 10 ms |
| objective | wi60x (NA1.2) |

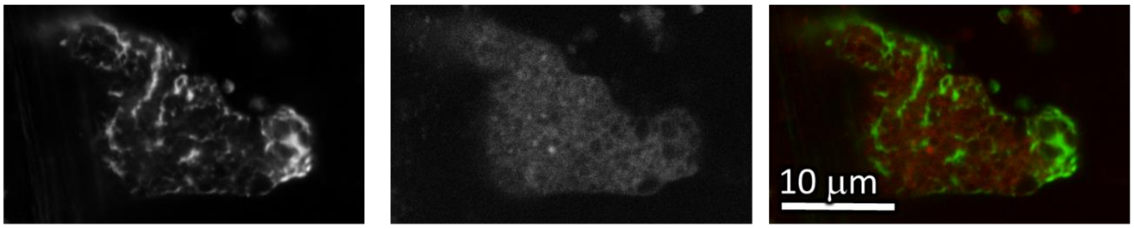

| | |
|---|---|
| Cell line | LA-4 (actin, Sir Actin) |
| NPs | TiO2 (Star 520) |
| exposure | 10:1, 24h-48h & 100 mM Resveratrol |
| imaging | xy STED, 48h |

| | |
|---|---|
| pixelsize (x,y) | 30 nm |
| FOV (x,y) | 32x20 um |
| pixelsize (z) | /nm |
| FOV (z) | /um |
| imaging time | /min |
| number z-stacks | |

| | |
|---|---|
| 561nm | 50% |
| 640nm | 50% |
| STED | 20% |
| filter sets | 605 nm – 625 nm, 650 nm – 720 nm |
| dwell time | 10 ms |
| objective | wi60x (NA1.2) |

*Figure S151: Confocal images of LA-4 in green and $TiO_2$ nanotubes in red channel. Cells are stained with Sir Actin label for F-actin. Confocal image of larger FoV in the upper window shows actin cytoskeleton of few cells with a lot of nanomaterial on the surface and one large cauliflower-like structure. In the lower window is STED zoom-in (XY plane) of the cauliflower-like structure form the upper window where we can clearly see actin filaments (green) with nanomaterial (red). Cells in this experiment have been treated with 100 μM Resveratrol in order to inhibit the activity of a FAS enzyme complex. During the*

*FAS inhibition we don't see big cauliflowers when we stain for plasma membrane (there is no colocalisation of plasma membrane and nanomaterial), but here we do observe cauliflowers and actin fibers excreted together with nanomaterial.*

#### S3g – Genomics – actin related expressions

See supplement S2d – Transcriptomics *in vitro* and *in vivo* after exposure to $TiO_2$.

### S4 – Macrophage action against epithelial defence

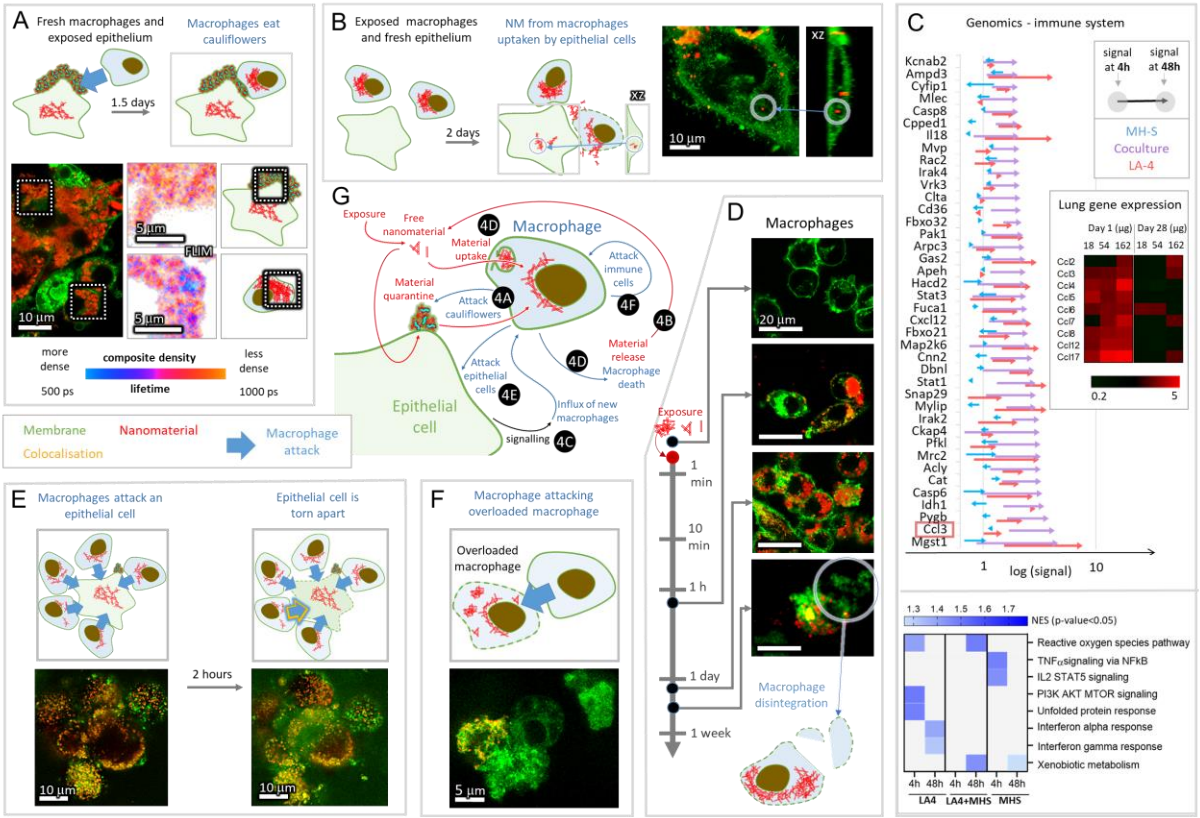

#### S4a – MH-S eat cauliflowers

##### Main message

2 days after previously non-exposed macrophages are added to a LA-4 culture with cauliflowers, they are seen filled up with nanomaterial. Since the LA-4 culture was washed prior to adding the macrophages to remove all free-floating nanomaterial, the only available source of such a huge amount of nanomaterial is from the cauliflowers. The fluorescence lifetime of the nanomaterial in macrophages lower than the fluorescence lifetime of nanomaterial in cauliflowers. Since lower lifetime corresponding to a more aggregated form of nanomaterial, the nanomaterial in macrophages is more tightly-packed than in the cauliflowers.

Supporting raw and analysed data:

Figure S152-Figure S155

##### Materials and methods

- experiment 20190621_e03.100_s01_FLIM:
    - LA-4 cells were seeded @50% confluence in an Ibidi #1.5H µ-Dish with a 4-Well Culture-Insert (Ibidi)
    - after 24 hours, 35 µl freshly filtered 1 mg/ml $TiO_2$-17-Alexa647 in 100x dcb was mixed into 130 µl fresh F12K medium and added to cells in one of the inserts to achieve 100:1 surface dose
    - 48 hours later, medium was removed from the cells and cells were washed with 2x 100 µl PBS to remove freely floating nanomaterial.
    - fresh (nonexposed) macrophages MH-S were added at 30% confluency to the exposed and washed LA-4, and were kept in a 1:1 mixture of media (total volume 140 µl) until observation
    - 34 hours later, the cells in the well were first incubated with 1.6 µg/ml CellMaskOrange in LCIS for 5 minutes, then incubated 15 minutes with freshly diluted 5 mM SAG-38 in LCIS at room temperature, and finally flushed with LCIS and observed in 150 µl LCIS in the home-made incubator at 37C

| | |
|---|---|
| Cell line | LA-4 and MH-S (membrane, CellMaskOrange and lipid bodies, SAG-38) |
| NPs | TiO2 (Alexa647) |
| exposure | 100:1, 48h LA-4, observed after 34h with MH-S |
| imaging | xy confocal |

| | |
|---|---|
| pixelsize (x,y) | 100 nm |
| FOV (x,y) | 70.0 µm |
| pixelsize (z) | |
| FOV (z) | |
| imaging time | |
| number of frames | |

| | |
|---|---|
| 561nm | 10% |
| 640nm | 10% |
| STED | |
| filter sets | 605 nm – 625 nm, 650 nm – 720 nm |
| dwell time | 10 µs |
| objective | 60x wi (NA 1.2) |

| | |
|---|---|
| Cell line | LA-4 and MH-S (membrane, CellMaskOrange and lipid bodies, SAG-38) |
| NPs | TiO2 (Alexa647) |
| exposure | 100:1, 48h LA-4, observed after 34h with MH-S |
| imaging | xy confocal FLIM |

| | |
|---|---|
| pixelsize (x,y) | 100 nm |
| FOV (x,y) | 51.2 µm |

| | |
|---|---|
| TCSPC pixelsize | 122 ps |
| TCSPC FOV | 19.5 ns |

| | |
|---|---|
| 561nm | (10%) |
| 640nm | 10% |
| STED | |
| diffraction grating | 523 – 722 nm |
| dwell time | 100 µs |
| objective | 60x wi (NA1.2) |

- fluorescence lifetime measurement and analysis:
    - see S2c – FLIM of cauliflowers

#### Experiment names

- Main experiment name:
    - 20190621_e03.100_s01_LA-4 MH-S CellMask TiO2-17-Alexa647 100.1_FLIM.msr

- Supplement Experiment names:
    - 20190621_e03.100_s01_LA-4 MH-S CellMask TiO2-17-Alexa647 100.1_FLIM.msr
    - 20190621_e03.100_s0X_cellsB.msr

#### Controls and statistics

Cross-sections of the cocultures of MH-S and LA-4 after the previously nonexposed MH-S have been in a coculture with LA-4 with already formed cauliflowers:

Figure S152-Figure S154

Fluorescence lifetime mapping of Figure S154:

Figure S155

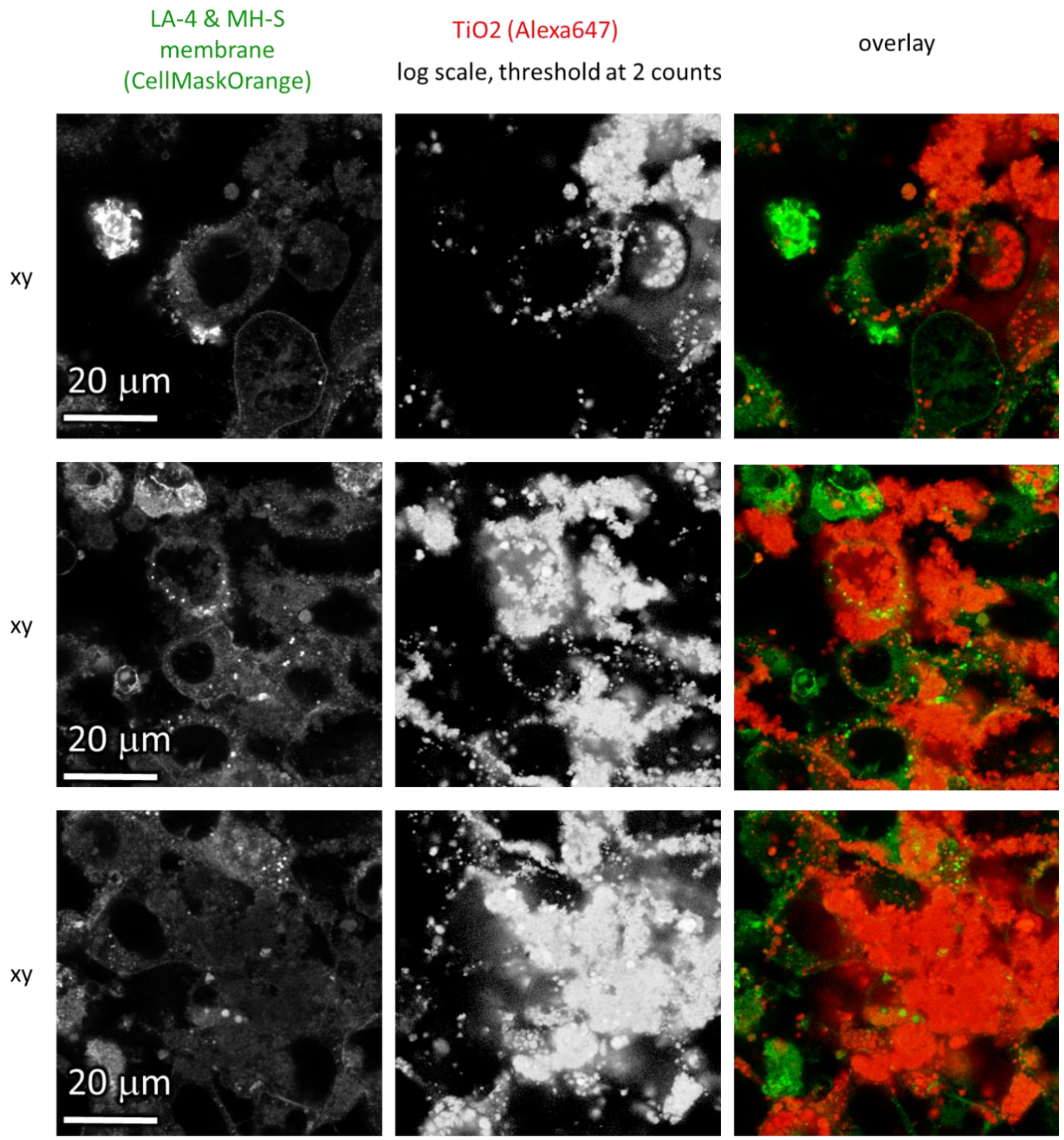

*Figure S152: Statistics for xy cross-sections of the cocultures of MH-S and LA-4 after the previously nonexposed MH-S have been in a coculture with LA-4 with already formed cauliflowers for 34 hours.*

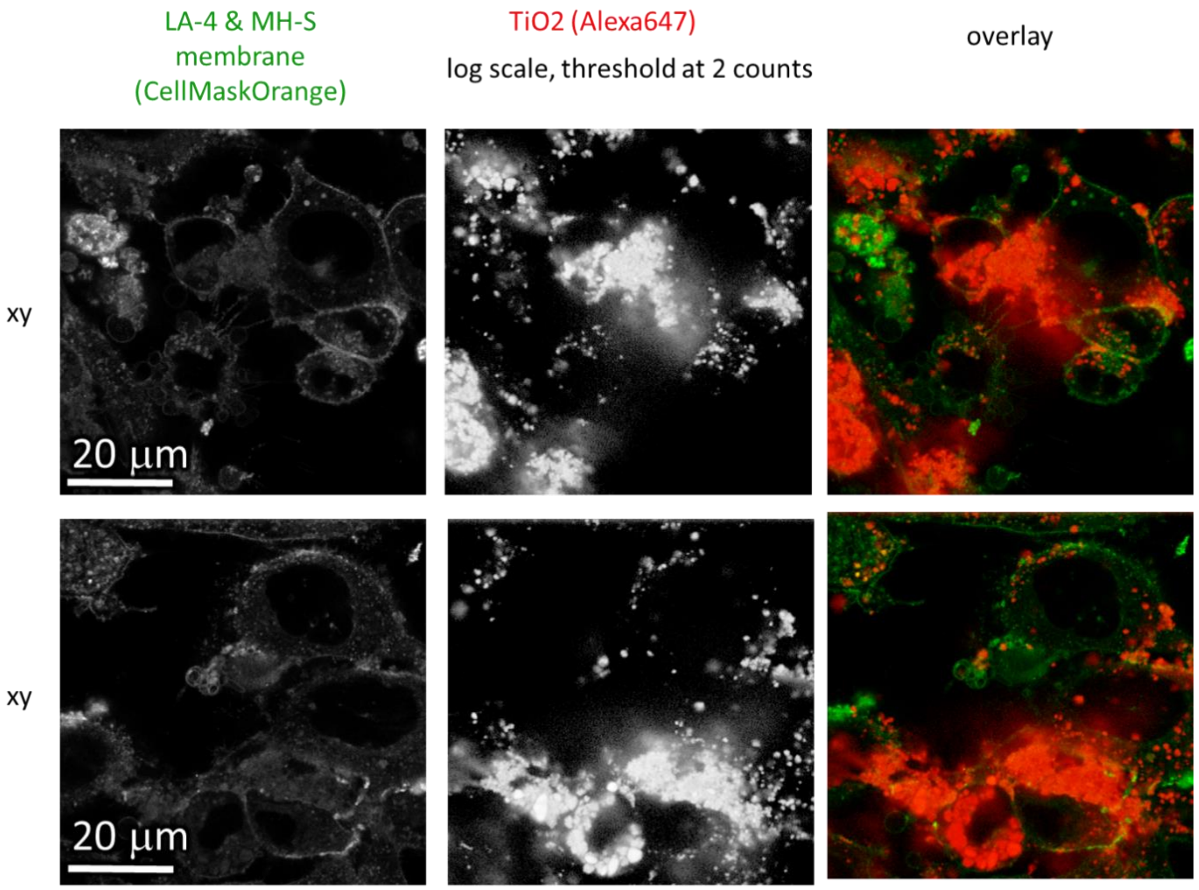

| | |
|---|---|
| Cell line | LA-4 and MH-S (membrane, CellMaskOrange and lipid bodies, SAG-38) |
| NPs | $TiO_2$ (Alexa647) |
| exposure | 100:1, 48h LA-4, observed after 34h with fresh MH-S |
| imaging | xy confocal |

| | |
|---|---|
| pixelsize (x,y) | 150 nm |
| FOV (x,y) | 70.0 µm |
| pixelsize (z) | |
| FOV (z) | |
| imaging time | |
| number of frames | |

| | |
|---|---|
| 561nm | 10% |
| 640nm | 10% |
| STED | |
| filter sets | 605 nm – 625 nm, 650 nm – 720 nm |
| dwell time | 10 µs |
| objective | 60x wi (NA 1.2) |

*Figure S153: Statistics for xy cross-sections of the cocultures of MH-S and LA-4 after the previously non-exposed MH-S have been in a coculture with LA-4 with already formed cauliflowers for 34 hours.*

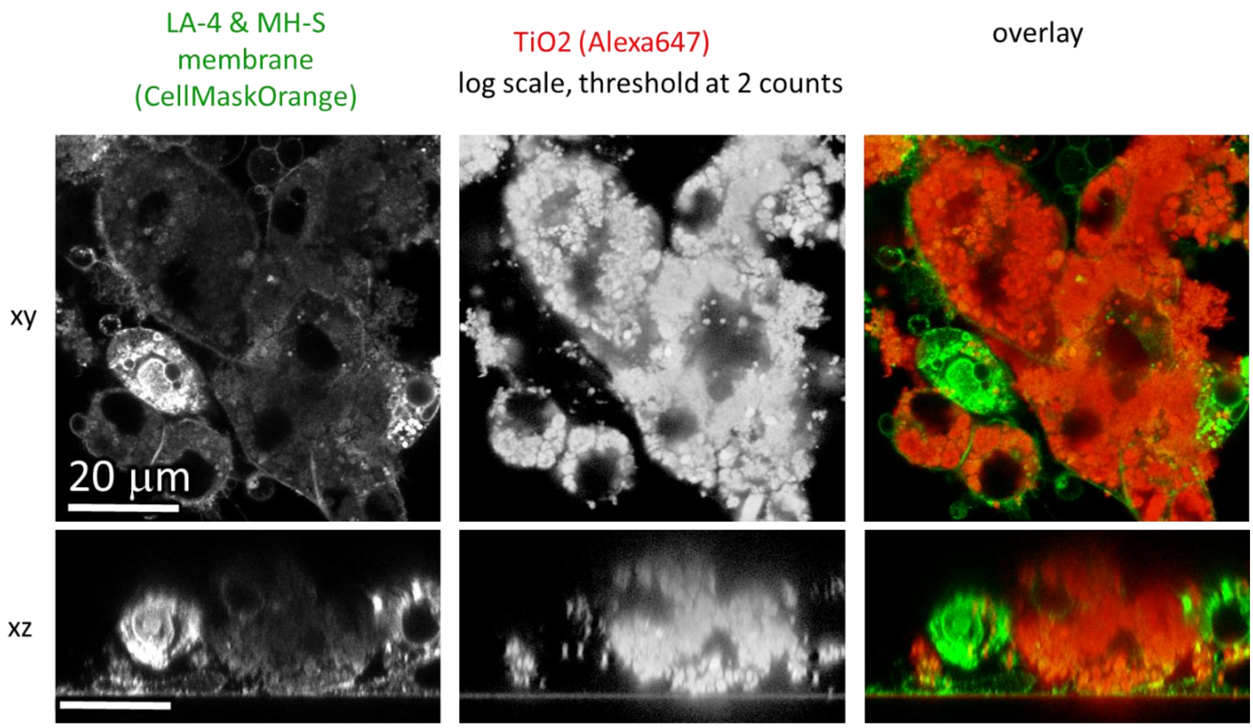

xy

| | |
|---|---|
| Cell line | LA-4 and MH-S (membrane, CellMaskOrange and lipid bodies, SAG-38) |
| NPs | TiO2 (Alexa647) |
| exposure | 100:1, 48h LA-4, observed after 34h with fresh MH-S |
| imaging | xy confocal |

| | |
|---|---|
| pixelsize (x,y) | 100 nm |
| FOV (x,y) | 70.0 µm |
| pixelsize (z) | |
| FOV (z) | |
| imaging time | |
| number of frames | |

| | |
|---|---|
| 561nm | 10% |
| 640nm | 10% |
| STED | |
| filter sets | 605 nm – 625 nm, 650 nm – 720 nm |
| dwell time | 10 µs |
| objective | 60x wi (NA 1.2) |

xz

| | |
|---|---|
| Cell line | LA-4 and MH-S (membrane, CellMaskOrange and lipid bodies, SAG-38) |
| NPs | TiO2 (Alexa647) |
| exposure | 100:1, 48h LA-4, observed after 34h with fresh MH-S |
| imaging | xz confocal |

| | |
|---|---|
| pixelsize (x,y) | 100 nm |
| FOV (x,y) | 70.0 µm |
| pixelsize (z) | 131 nm |
| FOV (z) | 35.0 µm |
| imaging time | |
| number of frames | |

| | |
|---|---|
| 561nm | 10% |
| 640nm | 10% |
| STED | |
| filter sets | 605 nm – 625 nm, 650 nm – 720 nm |
| dwell time | 10 µs |
| objective | 60x wi (NA 1.2) |

*Figure S154: Xy and xz cross-sections of the cocultures of MH-S and LA-4 after the previously non-exposed MH-S have been in a coculture with LA-4 with already formed cauliflowers for 34 h.*

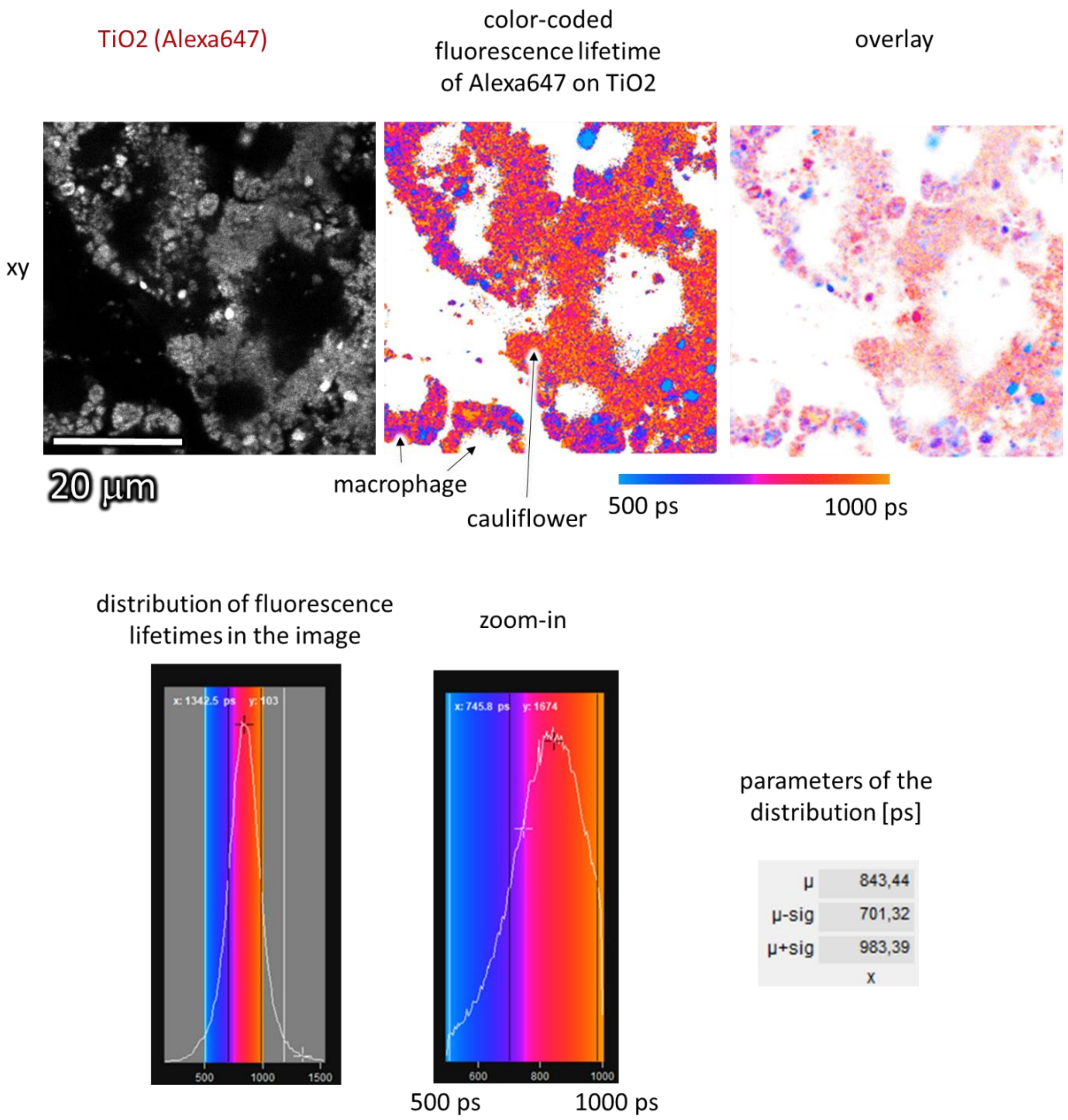

| | |
|---|---|
| Cell line | LA-4 and MH-S (membrane, CellMaskOrange and lipid bodies, SAG-38) |
| NPs | TiO2 (Alexa647) |
| exposure | 100:1, 48h LA-4, observed after 34h with fresh MH-S |
| imaging | xy confocal FLIM |

| | |
|---|---|
| pixelsize (x,y) | 100 nm |
| FOV (x,y) | 51.2 µm |
| TCSPC pixelsize | 122 ps |
| TCSPC FOV | 19.5 ns |

| | |
|---|---|
| 561nm | |
| 640nm | 10% |
| STED | |
| diffraction grating | 523 – 722 nm |
| dwell time | 100 µs |
| objective | 60x wi (NA1.2) |

*Figure S155: Analysis of fluorescence lifetimes of the coculture of MH-S and LA-4 after the previously nonexposed MH-S have been in a coculture with LA-4 with already formed cauliflowers. The fluorescence image, fluorescence-lifetime-color-coded image and distribution of fluorescence lifetimes in the image are shown.*

#### S4b – Re-uptake of nanomaterial into LA-4

##### Main message

Enter text please.

Supporting raw and analysed data:

Figure S156-Figure S158

##### Materials and methods

- experiment setup:
  - LA-4 and MH-S cells were seeded separately @15% confluence in two wells on an Ibidi #1.5H μ-Slide 8-well
  - after 8 hours, 10 μl 1 mg/ml $TiO_2$-17-Alexa647 in 100x dcb was added to MH-S and mixed to achieve 10:1 surface dose
  - 47 hours later, the medium was removed from MH-S cells and they were flushed with 200 μl PBS to remove mobile nanomaterial from the sample. After adding 100 μl fresh MH-S medium, they were scraped and added to LA-4 with 100 μl freshly changed LA-4 medium.
  - after 45 hours, the cells were incubated with 1.6 μg/ml CellMaskOrange for 7 minutes at 37C, afterwards they were observed in 200 μL LCIS in the home-made stage-top incubator at 37C

- analysis:
  - logarithmic scale in red and green channel, threshold at 3 counts, saturation at 200 counts (green channel) or 10 counts (red channel)

| | |
|---|---|
| Cell line | LA-4 and MH-S (membrane, CellMaskOrange) |
| NPs | TiO2 (Alexa647) |
| exposure | 10:1, 47h MH-S, observed after 45h with LA-4 |
| imaging | xy STED |

| | |
|---|---|
| pixelsize (x,y) | 50 nm |
| FOV (x,y) | 70.2 μm |
| pixelsize (z) | |
| FOV (z) | |
| imaging time | |
| number of frames | |

| | |
|---|---|
| 561nm | 5% |
| 640nm | 5% |
| STED | 5%, 71% 3D STED |
| filter sets | 605 nm – 625 nm, 650 nm – 720 nm |
| dwell time | 10 μs |
| objective | 60x wi (NA1.2) |

| | |
|---|---|
| Cell line | LA-4 and MH-S (membrane, CellMaskOrange) |
| NPs | TiO2 (Alexa647) |
| exposure | 10:1, 47h MH-S, observed after 45h with LA-4 |
| imaging | xz STED |

| | |
|---|---|
| pixelsize (x,y) | 50 nm |
| FOV (x,y) | 69.6 μm |
| pixelsize (z) | 50 nm |
| FOV (z) | 30.0 μm |
| imaging time | |
| number of frames | |

| | |
|---|---|
| 561nm | 5% |
| 640nm | 5% |
| STED | 5%, 71% 3D STED |
| filter sets | 605 nm – 625 nm, 650 nm – 720 nm |
| dwell time | 10 μs |
| objective | 60x wi (NA1.2) |

##### Experiment names

- Main experiment name:
  - 20190802_e10_s03_LA-4 MH-S CMO TiO2-17-Alexa647_0000 NN00 10.1_2 days nothing 2 days with full MH-S.msr

- Supplement experiment name:
  - 20190802/e10_s01_LA-4 MH-S CMO TiO2-17-Alexa647_0000 NN00 10.1_2 days nothing 2 days with full MH-S.msr
  - 20190802/e10_s02_LA-4 MH-S CMO TiO2-17-Alexa647_0000 NN00 10.1_2 days nothing 2 days with full MH-S.msr
  - 20190802/e10_s03_LA-4 MH-S CMO TiO2-17-Alexa647_0000 NN00 10.1_2 days nothing 2 days with full MH-S.msr
  - 20190802/e10_s04_LA-4 MH-S CMO TiO2-17-Alexa647_0000 NN00 10.1_2 days nothing 2 days with full MH-S.msr
  - 20190802/e10_s06_LA-4 MH-S CMO TiO2-17-Alexa647_0000 NN00 10.1_2 days nothing 2 days with full MH-S.msr

##### Controls and statistics

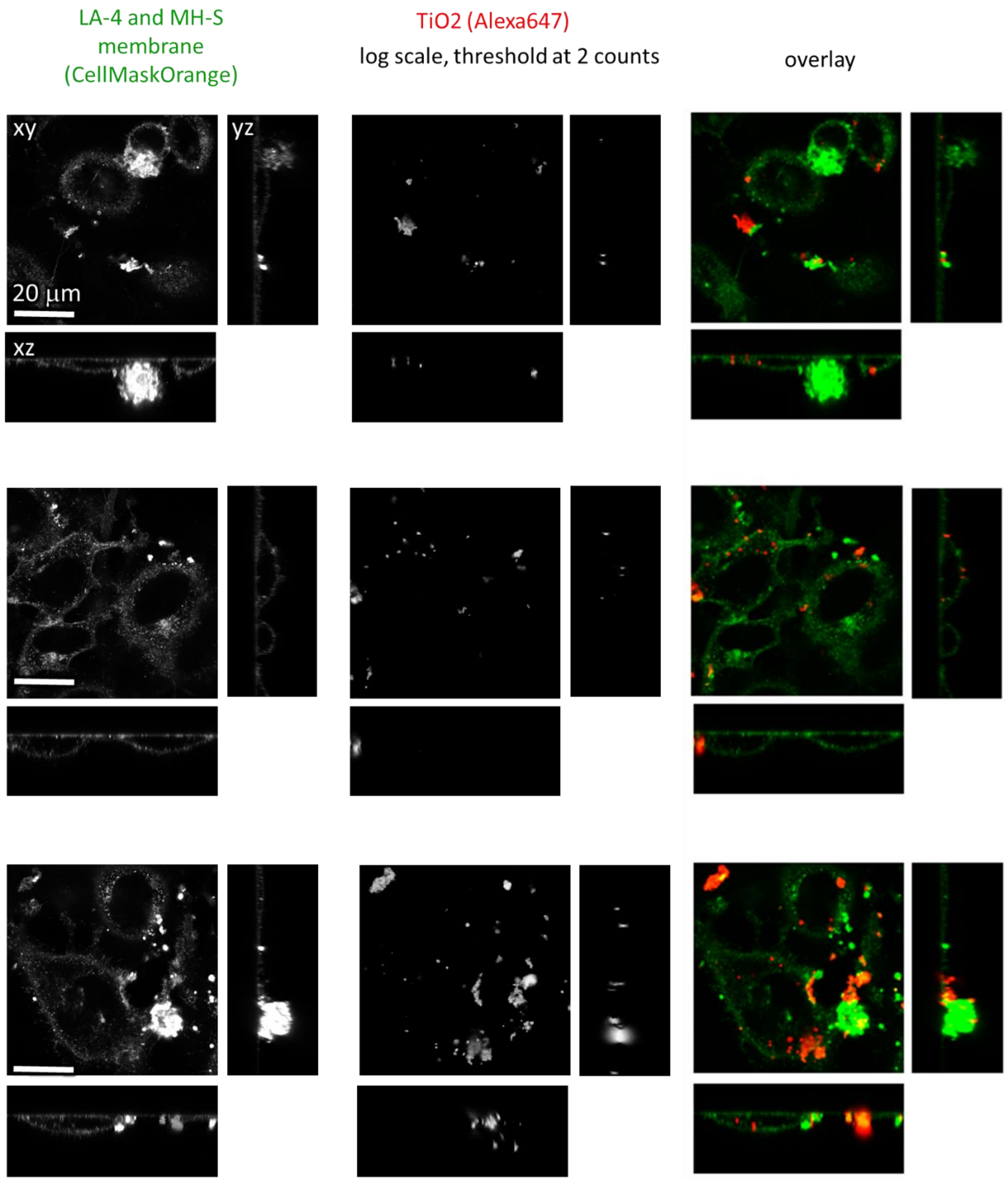

*Figure S156: Localisation of nanomaterial inside LA-4, which were exposed to nanomaterial solely by exposure to nanomaterial-laden macrophages (xy, xz and yz cross-sections)*

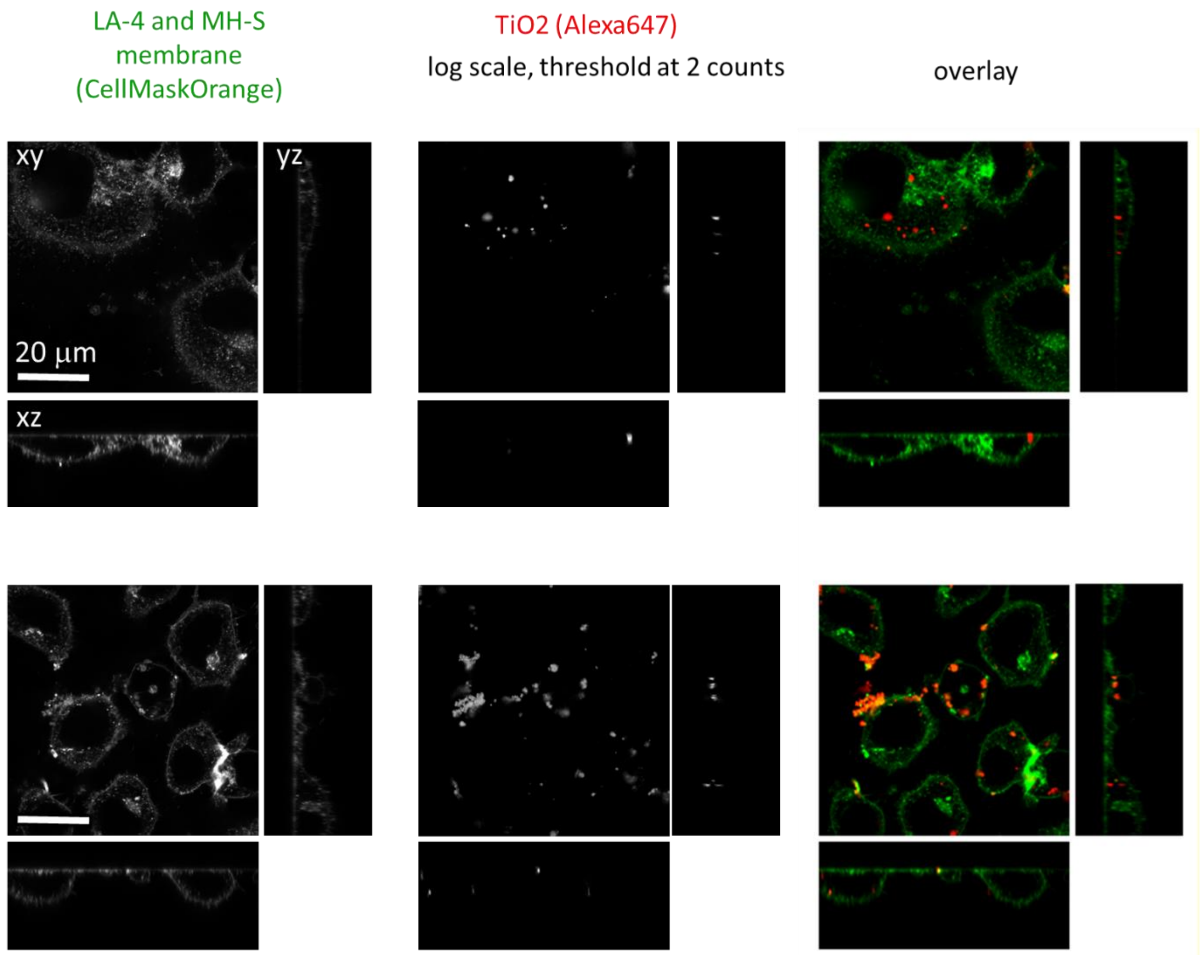

*Figure S157: Localisation of nanomaterial inside LA-4, which were exposed to nanomaterial solely by exposure to nanomaterial-laden macrophages (xy, xz and yz cross-sections)*

| | |
|---|---|
| Cell line | LA-4 and MH-S (membrane, CellMaskOrange) |
| NPs | TiO2 (Alexa647) |
| exposure | 10:1, 47h MH-S, observed after 45h with LA-4 |
| imaging | xy STED |

| | |
|---|---|
| pixelsize (x,y) | 50 nm |
| FOV (x,y) | 70.2 µm |
| pixelsize (z) | |
| FOV (z) | |
| imaging time | |
| number of frames | |

| | |
|---|---|
| 561nm | 5% |
| 640nm | 5% |
| STED | 5%, 71% 3D STED |
| filter sets | 605 nm – 625 nm, 650 nm – 720 nm |
| dwell time | 10 µs |
| objective | 60x wi (NA1.2) |

| | |
|---|---|
| Cell line | LA-4 and MH-S (membrane, CellMaskOrange) |
| NPs | TiO2 (Alexa647) |
| exposure | 10:1, 47h MH-S, observed after 45h with LA-4 |
| imaging | xz STED |

| | |
|---|---|
| pixelsize (x,y) | 50 nm |
| FOV (x,y) | 69.6 µm |
| pixelsize (z) | 50 nm |
| FOV (z) | 30.0 µm |
| imaging time | |
| number of frames | |

| | |
|---|---|
| 561nm | 5% |
| 640nm | 5% |
| STED | 5%, 71% 3D STED |
| filter sets | 605 nm – 625 nm, 650 nm – 720 nm |
| dwell time | 10 µs |
| objective | 60x wi (NA1.2) |

| | |
|---|---|
| Cell line | LA-4 and MH-S (membrane, CellMaskOrange) |
| NPs | TiO2 (Alexa647) |
| exposure | 10:1, 47h MH-S, observed after 45h with LA-4 |
| imaging | yz STED |

| | |
|---|---|
| pixelsize (x,y) | 50 nm |
| FOV (x,y) | 69.6 µm |
| pixelsize (z) | 50 nm |
| FOV (z) | 30.0 µm |
| imaging time | |
| number of frames | |

| | |
|---|---|
| 561nm | 5% |
| 640nm | 5% |
| STED | 5%, 71% 3D STED |
| filter sets | 605 nm – 625 nm, 650 nm – 720 nm |
| dwell time | 10 µs |
| objective | 60x wi (NA1.2) |

*Figure S158: Imaging parameters for the upper two Figures.*

#### S4c – Genomics – immune system

See supplement S2d – Transcriptomics *in vitro* and *in vivo* after exposure to $TiO_2$.

#### S4d – MH-S eat nanomaterial

##### Main message

Macrophages quickly internalise nanomaterial and are full of nanomaterial after a few days. However, they are also seen to die after a few days of exposure to nanomaterial.

Supporting raw and analysed data:

Figure S159-Figure S169

##### Materials and methods

*Experiment at time 0h:*

- experiment: 20190412_e05_t01
- protocol:
    - MH-S cells were seeded @20% confluence in an Ibidi #1.5H µ-Dish
    - after 4 days 100 µl 5 µg/ml CellMaskOrange was added to 400 µl old cell medium on the cells. After 7 minutes at room temperature, the medium was removed and cells were observed in 400 µl LICS in the home-made stage-top incubator at 37C.
- analysis:
    - rescaling Green channel to 65 counts

##### Link to time-lapse

- http://lbfnanobiodatabase.ijs.si/file/data/cauliflowerpaper/20190412_e05_t01b_MH-S_CellMask_xyt_10_minutes_1s_is_2minutes.gif

*Movie S23: Time-lapse of non-exposed macrophages (CellMask Orange, green), 1 s in movie corresponds to 2 minutes in real time.*

- http://lbfnanobiodatabase.ijs.si/file/data/cauliflowerpaper/20190412_e05_t04_xyt_MH-S_CellMask_MH-S_dividing_10minutes_1s_is_2minutes.gif

*Movie S24: Time-lapse of non-exposed macrophages and their division (CellMask Orange, green), 1 s in movie corresponds to 2 minutes in real time.*

##### Link to 3D

- http://lbfnanobiodatabase.ijs.si/file/data/cauliflowerpaper/20190412_e05_t01_MH-S_CellMask_xyz_FOV_70um_x_70um_x_30um.mp4

*Movie S25: 3D of non-exposed macrophages (CellMask Orange, green), field-of-view is 70 x 70 x 30 µm.*

| | | | | | |
|---|---|---|---|---|---|
| Cell line | MH-S (membrane, CellMaskOrange) | pixelsize (x,y) | 100 nm | 561nm | 5% |
| NPs | | FOV (x,y) | 70.2 µm | 640nm | 5% |
| exposure | | pixelsize (z) | | STED | |
| imaging | xy confocal | FOV (z) | | filter sets | 605 nm – 625 nm |
| | | imaging time | | dwell time | 10 µs |
| | | number of frames | | objective | 60x wi (NA1.2) |

| | | | | | |
|---|---|---|---|---|---|
| Cell line | MH-S (membrane, CellMaskOrange) | pixelsize (x,y) | 200 nm | 561nm | 5% |
| NPs | | FOV (x,y) | 70.2 µm | 640nm | 5% |
| exposure | | pixelsize (z) | 1 µm | STED | |
| imaging | xyz confocal | FOV (z) | 30 µm | filter sets | 605 nm – 625 nm |
| | | imaging time | | dwell time | 10 µs |
| | | number of frames | | objective | 60x wi (NA1.2) |

| | | | | | |
|---|---|---|---|---|---|
| Cell line | MH-S (membrane, CellMaskOrange) | pixelsize (x,y) | 200 nm | 561nm | 5% |
| NPs | | FOV (x,y) | 70.2 µm | 640nm | 5% |
| exposure | | pixelsize (z) | | STED | |
| imaging | xyt confocal | FOV (z) | | filter sets | 605 nm – 625 nm |
| | | imaging time | 10 minutes | dwell time | 10 µs |
| | | number of frames | 60 | objective | 60x wi (NA1.2) |

*Experiment at time 2h:*

- experiment: 20190412/e06_t03…t13
- protocol:
  - MH-S cells were seeded @20% confluence in an Ibidi #1.5H µ-Dish
  - after 4 days 100 µl 5 µg/ml CellMaskOrange was added to 400 µl old cell medium on the cells. After 7 minutes at room temperature, the medium was removed and cells were observed in 400 µl LICS in the home-made stage-top incubator at 37C.
  - just prior to filming, freshly filtered 5 ml 1 mg/ml $TiO_2$-40-Alexa647 in 100x dcb was added directly to the cells (in 400 µl LCIS) and mixed to achieve 2:1 surface dose
- analysis:
  - logarithmic scale in red channel, cut-off at 2 counts, saturation at 256 counts, linear scale on green channel, saturation at 150 counts

Link to time-lapse

- http://lbfnanobiodatabase.ijs.si/file/data/cauliflowerpaper/20190412_e06_MH-S_CellMask_TiO2Alexa647_1.1_live_xyt_total_2_h_with_breaks_for_xyz_1s_is_2min.gif

*Movie S26: Live movie of macrophages (CellMask, green) exposed to 1:1 surface dose of $TiO_2$, from 0h – 2 hours (1 second corresponds to 2 minutes real time).*

#### Link to 3D

- http://lbfnanobiodatabase.ijs.si/file/data/cauliflowerpaper/20190412_e06_MH-S_CellMask_TiO2Alexa647_1.1_xyz_FOV_70_x_70_x_30_um_1.mp4

*Movie S27: Comparison of 3D renders of macrophages (CellMask, green) exposed to 1:1 surface dose of $TiO_2$ nanotubes (Alexa 647, red) at different time-points from 0h – 2 hours, field-of-view is 70 x 70 x 30 µm.*

http://lbfnanobiodatabase.ijs.si/file/data/cauliflowerpaper/20190412_e06_MHS_CellMask_TiO2Alexa647_1.1_xyz_FOV_70_x_70_x_30_um.mp4

*Movie S28: 3D movie of macrophages (CellMask, green) exposed to 1:1 surface dose of $TiO_2$ nanotubes (Alexa 647, red) at different time-points from 0h – 2 hours, field-of-view is 70 x 70 x 30 µm.*

- separate timepoints:
  - http://lbfnanobiodatabase.ijs.si/file/data/cauliflowerpaper/20190412_e06_t04_MH-S_CellMask_TiO2Alexa647_1.1_live_xyz_FOV_70_x_70_x_30_um_25min.gif

*Movie S29: Macrophages (CellMask, green) exposed to 1:1 surface dose of $TiO_2$ nanotubes (Alexa 647, red) after 25 minutes*

  - http://lbfnanobiodatabase.ijs.si/file/data/cauliflowerpaper/20190412_e06_t06_MH-S_CellMask_TiO2Alexa647_1.1_live_xyz_FOV_70_x_70_x_30_um_45min.gif

*Movie S30: Macrophages (CellMask, green) exposed to 1:1 surface dose of $TiO_2$ nanotubes (Alexa 647, red) after 45 minutes*

  - http://lbfnanobiodatabase.ijs.si/file/data/cauliflowerpaper/20190412_e06_t08_MH-S_CellMask_TiO2Alexa647_1.1_live_xyz_FOV_70_x_70_x_30_um_1h_25min.gif

*Movie S31: Macrophages (CellMask, green) exposed to 1:1 surface dose of $TiO_2$ nanotubes (Alexa 647, red) after 1h 25 minutes*

  - http://lbfnanobiodatabase.ijs.si/file/data/cauliflowerpaper/20190412_e06_t09_MH-S_CellMask_TiO2Alexa647_1.1_live_xyz_FOV_70_x_70_x_30_um_1h_45min.gif

*Movie S32: Macrophages (CellMask, green) exposed to 1:1 surface dose of $TiO_2$ nanotubes (Alexa 647, red) after 1h 45 minutes*

  - http://lbfnanobiodatabase.ijs.si/file/data/cauliflowerpaper/20190412_e06_t12_MH-S_CellMask_TiO2Alexa647_1.1_live_xyz_FOV_70_x_70_x_30_um_2h.gif

*Movie S33: Macrophages (CellMask, green) exposed to 1:1 surface dose of $TiO_2$ nanotubes (Alexa 647, red) after 2 hours*

  - http://lbfnanobiodatabase.ijs.si/file/data/cauliflowerpaper/20190412_e06_t13_MH-

[S_CellMask_TiO2Alexa647_1.1_live_xyz_FOV_70_x_70_x_30_um_2h_10min.gif]

*Movie S34: Macrophages (CellMask, green) exposed to 1:1 surface dose of $TiO_2$ nanotubes (Alexa 647, red) after 2 h 10 minutes*

| | | | | | |
|---|---|---|---|---|---|
| Cell line | MH-S (membrane, CellMaskOrange) | pixelsize (x,y) | 200 nm | 561nm | 5% |
| NPs | TiO2 (Alexa647) | FOV (x,y) | 70.2 µm | 640nm | 5% |
| exposure | 2:1, 25min – 2h | pixelsize (z) | | STED | |
| imaging | xyt confocal | FOV (z) | | filter sets | 605 nm – 625 nm, 650 nm – 720 nm |
| | | imaging time | 95 min | dwell time | 10 µs |
| | | number of frames | 491 | objective | 60x wi (NA 1.2) |

| | | | | | |
|---|---|---|---|---|---|
| Cell line | MH-S (membrane, CellMaskOrange) | pixelsize (x,y) | 200 nm | 561nm | 5% |
| NPs | TiO2 (Alexa647) | FOV (x,y) | 70.2 µm | 640nm | 5% |
| exposure | 2:1, 0min – 2h | pixelsize (z) | 500 nm | STED | |
| imaging | xyz confocal | FOV (z) | 30 µm | filter sets | 605 nm – 625 nm, 650 nm – 720 nm |
| | | imaging time | | dwell time | 10 µs |
| | | number of frames | | objective | 60x wi (NA 1.2) |

##### *Experiment at time-point 2 days:*

- experiment: 20190412/e07_t04
- protocol:
  - MH-S cells were seeded @20% confluence in an Ibidi #1.5H µ-Dish
  - after 1 day 30 µl freshly filtered 1 mg/ml $TiO_2$-40-Alexa647 in 100x dcb was added to 400µL cell medium already on the cells and mixed to achieve 10:1 surface dose
  - 53 hours later, 100 µl 5 µg/ml CellMaskOrange was added to 400 µl old cell medium on the cells. After 9 minutes at 37C, the medium was removed and cells were observed in 400 µl LICS in the home-made stage-top incubator at 37C.
- analysis:
  - logarithmic scale in red channel, cut-off at 2 counts, linear scale on green channel, saturation at 40 counts

##### Link to 3D

- *http://lbfnanobiodatabase.ijs.si/file/data/cauliflowerpaper/20190412_e07_t04_MH-S_CellMask_TiO2-Alexa647_10.1_2_days_xyz_FOV_70_x_70_x_30_um.mp4*

*Movie S35: Macrophages (CellMask, green) after 2 days of exposure to 10:1 surface dose of $TiO_2$ nanotubes (Alexa 647, red), field-of-view is 70 x 70 x 30 µm.*

##### Link to time-lapse

- http://lbfnanobiodatabase.ijs.si/file/data/cauliflowerpaper/20190412_e07_t06_MH-S_CellMask_TiO2-Alexa647_10.1_2_days_xyt_11min_1s_is_1min.gif

*Movie S36: Dynamics of macrophages (CellMask, green) after 2 days of exposure to 10:1 surface dose of $TiO_2$ nanotubes (Alexa 647, red), 1 second in movie corresponds to 1 minute real time.*

| | |
|---|---|
| Cell line | MH-S (membrane, CellMaskOrange) |
| NPs | TiO2 (Alexa647) |
| exposure | 10:1, 2 days |
| imaging | xyz confocal |

| | |
|---|---|
| pixelsize (x,y) | 150 nm |
| FOV (x,y) | 70.1 µm |
| pixelsize (z) | 500 nm |
| FOV (z) | 30 µm |
| imaging time | |
| number of frames | |

| | |
|---|---|
| 561nm | 5% |
| 640nm | 5% |
| STED | |
| filter sets | 605 nm – 625 nm, 650 nm – 720 nm |
| dwell time | 10 µs |
| objective | 60x wi (NA 1.2) |

| | |
|---|---|
| Cell line | MH-S (membrane, CellMaskOrange) |
| NPs | TiO2 (Alexa647) |
| exposure | 10:1, 2 days |
| imaging | xyt confocal |

| | |
|---|---|
| pixelsize (x,y) | 150 nm |
| FOV (x,y) | 70.1 µm |
| pixelsize (z) | |
| FOV (z) | |
| imaging time | 10 min |
| number of frames | 65 |

| | |
|---|---|
| 561nm | 5% |
| 640nm | 5% |
| STED | |
| filter sets | 605 nm – 625 nm, 650 nm – 720 nm |
| dwell time | 10 µs |
| objective | 60x wi (NA 1.2) |

##### *Experiment at time 4 days:*

- 20190225_e02 m03 s03_dying MH-S_SR DPPE and TiO2_ATTO 594_ over 48h_1 to 10

- protocol:
  - MH-S cells were seeded @30% confluence in an Ibidi #1.5H µ-Dish
  - after 24 hours 35 ml freshly filtered 1 mg/ml $TiO_2$-40-ATTO594 in 100x dcb was added directly to the cells (in 400 µl medium) and mixed to achieve 1:1, 10:1 and 100:1 surface doses
  - 72 hours later, the cells were incubated with 1 um StarRed-DPPE for 5 minutes at the room temperature, afterwards they were flushed with 1x400 ml LCIS (in order not to lose too many macrophages which are not completely adherent) and observed in 400 mL LCIS on the microscope stage heated at 37°C

- analysis:
  - Green channel signal was multiplied by the factor of 1,5 and Gaussian Blur (0,5 pixel) was applied
  - Red channel was multiplied as much as the green in order to conserve colocalization signal and Unshrap Mask (3 pixels with weight 0,6) was applied to better visualise nanoparticle structure

##### Link to 3D

- http://lbfnanobiodatabase.ijs.si/file/data/cauliflowerpaper/20190225_e02_m03_s03_dying_MH-S_SR_DPPE_and_TiO2_ATTO_594__over_48h_1_to_10_b.mp4

*Movie S37:Macrophage (StarRed DPPE, green) after 72 h of exposure to 10:1 dose of $TiO_2$ nanotubes (ATTO594, red).*

- http://lbfnanobiodatabase.ijs.si/file/data/cauliflowerpaper/20190225_dying_MHS_SR_DPPE_and_TiO2_ATTO_594__over_48h_1_to_1.gif

*Movie S38: Macrophage (StarRed DPPE, green) after 72 h of exposure to 1:1 dose of $TiO_2$ nanotubes (ATTO594, red).*

- http://lbfnanobiodatabase.ijs.si/file/data/cauliflowerpaper/20190225_e02_m02_s02_dying_MHS_SR_DPPE_and_TiO2_ATTO_594__over_48h_1_to_10.gif

*Movie S39: Macrophage (StarRed DPPE, green) after 72 h of exposure to 10:1 dose of $TiO_2$ nanotubes (ATTO594, red).*

- http://lbfnanobiodatabase.ijs.si/file/data/cauliflowerpaper/20190225_e02_m03_s03_dying_MH-S_SR_DPPE_and_TiO2_ATTO_594__over_48h_1_to_10_b.mp4

*Movie S40: Macrophage (StarRed DPPE, green) after 72 h of exposure to 10:1 dose of $TiO_2$ nanotubes (ATTO594, red).*

- http://lbfnanobiodatabase.ijs.si/file/data/cauliflowerpaper/20190225_e03_m01_s01_dying_MH-S_SR_DPPE_and_TiO2_ATTO_594__over_48h_1_to_100_agglomerats_of_NP_and_3D.gif

*Movie S41: Macrophage (StarRed DPPE, green) after 72 h of exposure to 100:1 dose of $TiO_2$ nanotubes (ATTO594, red).*

- http://lbfnanobiodatabase.ijs.si/file/data/cauliflowerpaper/3D_72h_MH-S_SR_DPPE_&_TiO2_ATTO_594_dose_dependend_desintegration.mp4

*Movie S42: Comparison of macrophages (StarRed DPPE, green) after 72 h of exposure to 1:1, 10:1, 10:1 and 100:1 dose of $TiO_2$ nanotubes (ATTO594, red).*

| | |
|---|---|
| Cell line | LA-4 (membrane, SR-DPPE) |
| NPs | TiO2_ATTO 594 |
| exposure | 1:10 |
| imaging | xyz STED, @72h |

| | |
|---|---|
| pixelsize (x,y) | 150 nm |
| FOV (x,y) | 32x28 µm |
| pixelsize (z) | 150 nm |
| FOV (z) | 20 µm |
| imaging time | / |
| number of stacks | 106 |

| | |
|---|---|
| 561nm | 25% |
| 640nm | 40% |
| STED | 23% |
| filter sets | 605 nm – 625 nm, 400nm-780nm |
| dwell time | 10 µs |
| objective | wi60x (NA1.2) |

#### Experiment names

- Main experiment name:
  - **experiment at time 0h**
    - 2019_04_12/e05_t01_MH-S CellMask_xyt.msr
  - **experiment at time 2h**
    - 2019_04_12/e06_t03_MH-S CellMask TiO2-Alexa647_1.1 live_xyz xyt.msr
    - 2019_04_12/e06_t04_MH-S CellMask TiO2-Alexa647_1.1 live_xyz xyt.msr
    - 2019_04_12/e06_t06_MH-S CellMask TiO2-Alexa647_1.1 live_xyz xyt.msr

    - 2019_04_12/e06_t07_MH-S CellMask TiO2-Alexa647_1.1 live_xyz xyt.msr
    - 2019_04_12/e06_t08_MH-S CellMask TiO2-Alexa647_1.1 live_xyz xyt.msr
    - 2019_04_12/e06_t09_MH-S CellMask TiO2-Alexa647_1.1 live_xyz xyt.msr
    - 2019_04_12/e06_t10_MH-S CellMask TiO2-Alexa647_1.1 live_xyz xyt.msr
    - 2019_04_12/e06_t12_MH-S CellMask TiO2-Alexa647_1.1 live_xyz xyt.msr
    - 2019_04_12/e06_t13_MH-S CellMask TiO2-Alexa647_1.1 live_xyz xyt.msr
    - 2019_04_12/e06_t14_MH-S CellMask TiO2-Alexa647_1.1 live_xyz xyt.msr

  - **experiment at time 2 days**
    - 2019_04_12/e07_t04_MH-S CellMask TiO2-Alexa647_10.1 2 days_xyz.msr

  - **experiment at time 4 days**
    - 20190225_e02 m03 s03_dying MH-S_SR DPPE and TiO2_ATTO 594_ over 48h_1 to 10

- Supplement experiment name:

  - **experiment at time 0h**
    - 20190412/ e05_t01_MH-S CellMask_xyt.msr
    - 20190412/ e05_t04_MH-S CellMask_xyt.msr
    - 20190802/e03_s01_MH-S CMO 4 days nothing_happy MH-S.msr
    - 20190802/e03_s02_MH-S CMO 4 days nothing_happy MH-S.msr
    - 20190802/e03_s03_MH-S CMO 4 days nothing_happy MH-S.msr

  - **experiment at time 2h**
    - 2019_04_12/e06_t04_MH-S CellMask TiO2-Alexa647_1.1 live_xyz xyt.msr
    - 2019_04_12/e06_t06_MH-S CellMask TiO2-Alexa647_1.1 live_xyz xyt.msr
    - 2019_04_12/e06_t09_MH-S CellMask TiO2-Alexa647_1.1 live_xyz xyt.msr
    - 2019_04_12/e06_t13_MH-S CellMask TiO2-Alexa647_1.1 live_xyz xyt.msr

  - **experiment at time 2 days**
    - 2019_04_12/e07_t04_MH-S CellMask TiO2-Alexa647_10.1 2 days_xyz.msr
    - 20190712_MH-S NP_s03_t04_MH-S eating.msr
    - 20190712_MH-S NP_s03_t06_STED MH-S eating.msr
    - 20190712_MH-S NP_s04_t08_STED.msr
    - 20190712_MH-S NP_s07_t12_STED no cauliflowers.msr
    - 20190712_MH-S NP_s11_t16_glued MH-S.msr

  - **experiment at time 4 days**
    - 20190225_e01 m04 s05_dying MH-S_SR DPPE and TiO2_ATTO 594_ over 48h_1 to 1
    - 20190225_e02 m02 s02_dying MH-S_SR DPPE and TiO2_ATTO 594_ over 48h_1 to 10
    - 20190225_e03 m01 s01_dying MH-S_SR DPPE and TiO2_ATTO 594_ over 48h_1 to 100 agglomerates of NP and 3D

##### Controls and statistics

Non-exposed macrophages at time 0h:

Figure S159-Figure S161

Initial internalisation of nanomaterial in macrophages at time 0-2h:

Figure S162

Macrophages are full on nanomaterial at 2 days of exposure:

Figure S163-Figure S165

*Experiment at time 0h*

- 20190412_e05_t01

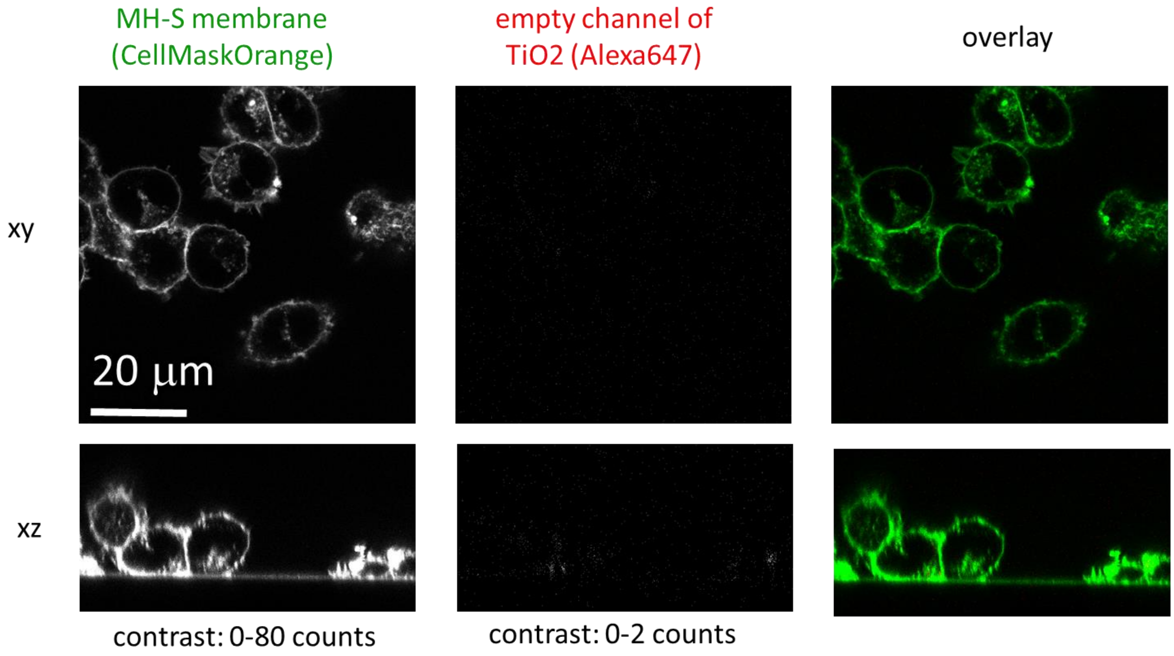

xy

| | |
|---|---|
| Cell line | MH-S (membrane, CellMaskOrange) |
| NPs | |
| exposure | |
| imaging | xy confocal |

| | |
|---|---|
| pixelsize (x,y) | 100 nm |
| FOV (x,y) | 70.2 µm |
| pixelsize (z) | |
| FOV (z) | |
| imaging time | |
| number of frames | |

| | |
|---|---|
| 561nm | 5% |
| 640nm | 5% |
| STED | |
| filter sets | 605 nm – 625 nm |
| dwell time | 10 µs |
| objective | 60x wi (NA1.2) |

xz

| | |
|---|---|
| Cell line | MH-S (membrane, CellMaskOrange) |
| NPs | |
| exposure | |
| imaging | xz confocal |

| | |
|---|---|
| pixelsize (x,y) | 100 nm |
| FOV (x,y) | 70.0 µm |
| pixelsize (z) | 131 nm |
| FOV (z) | 34.9 µm |
| imaging time | |
| number of frames | |

| | |
|---|---|
| 561nm | 10% |
| 640nm | 10% |
| STED | |
| filter sets | 605 nm – 625 nm |
| dwell time | 10 µs |
| objective | 60x wi (NA1.2) |

*Figure S159: Non-exposed macrophages at time 0h*

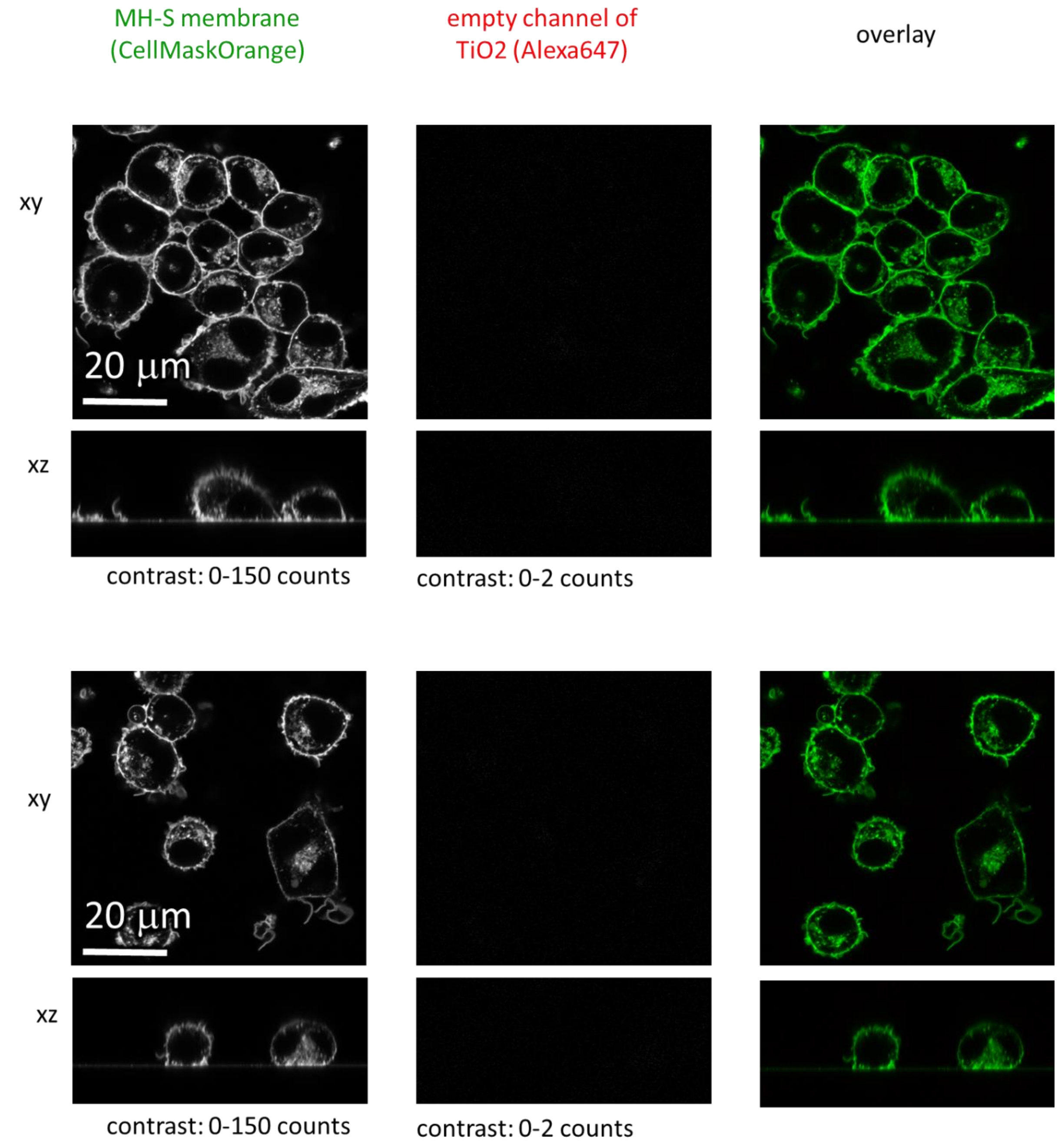

*Figure S160: Non-exposed macrophages at time 0h*

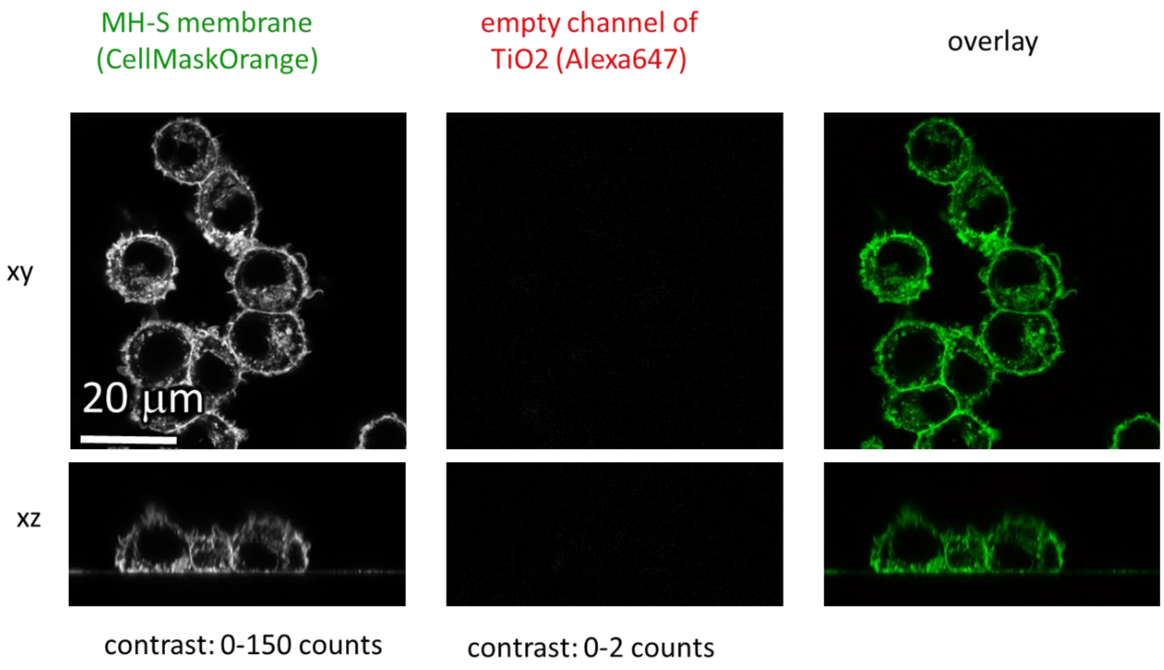

xy

| | | | | | | |
|---|---|---|---|---|---|---|
| Cell line | MH-S (membrane, CellMaskOrange) | pixelsize (x,y) | 50 nm | | 561nm | 5% |
| NPs | | FOV (x,y) | 70.2 µm | | 640nm | 5% |
| exposure | | pixelsize (z) | | | STED | 5% |
| imaging | xy confocal | FOV (z) | | | filter sets | 605 nm – 625 nm |
| | | imaging time | | | dwell time | 10 µs |
| | | number of frames | | | objective | 60x wi (NA1.2) |

xz

| | | | | | | |
|---|---|---|---|---|---|---|
| Cell line | MH-S (membrane, CellMaskOrange) | pixelsize (x,y) | 50 nm | | 561nm | 5% |
| NPs | | FOV (x,y) | 70.0 µm | | 640nm | 5% |
| exposure | | pixelsize (z) | 50 nm | | STED | 5% |
| imaging | xz confocal | FOV (z) | 30 µm | | filter sets | 605 nm – 625 nm |
| | | imaging time | | | dwell time | 10 µs |
| | | number of frames | | | objective | 60x wi (NA1.2) |

*Figure S161: Non-exposed macrophages at time 0h*

#### *Experiment at time 0-2h*

- 20190412/e06_t03…t13

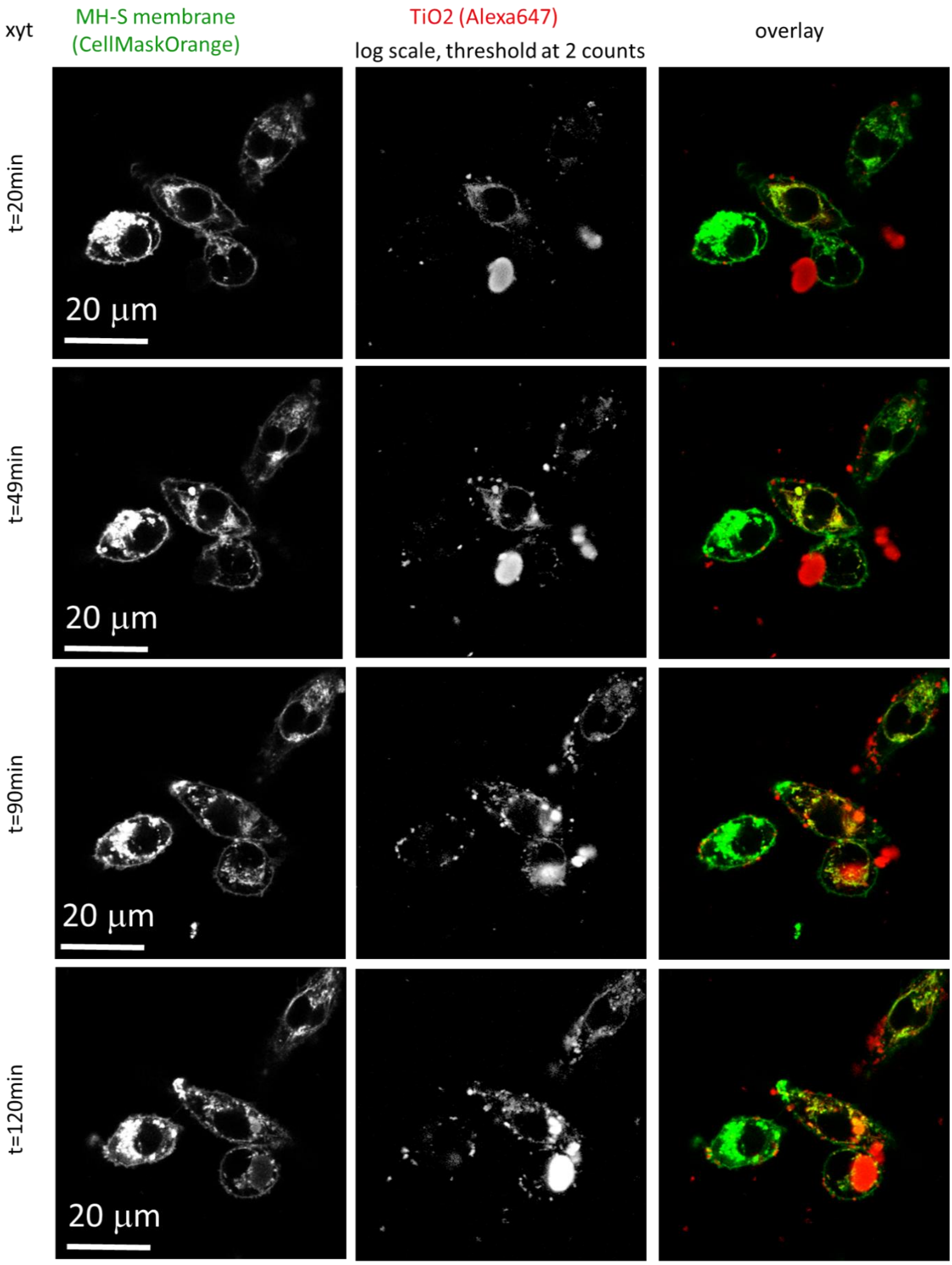

xyt
MH-S membrane
(CellMaskOrange)
TiO2 (Alexa647)
log scale, threshold at 2 counts
overlay
t=20min
t=49min
t=90min
t=120min
20 μm

| | | | | | |
|---|---|---|---|---|---|
| Cell line | MH-S (membrane, CellMaskOrange) | pixelsize (x,y) | 200 nm | 561nm | 5% |
| NPs | TiO2 (Alexa647) | FOV (x,y) | 70.2 µm | 640nm | 5% |
| exposure | 2:1, 0min – 2h | pixelsize (z) | 500 nm | STED | |
| imaging | xyz confocal | FOV (z) | 30 µm | filter sets | 605 nm – 625 nm, 650 nm – 720 nm |
| | | imaging time | | dwell time | 10 µs |
| | | number of frames | | objective | 60x wi (NA 1.2) |

*Figure S162: Initial internalisation of nanomaterial in macrophages at time 0-2h*

##### *Experiment at time 2 days*

- 20190412/e07_t04

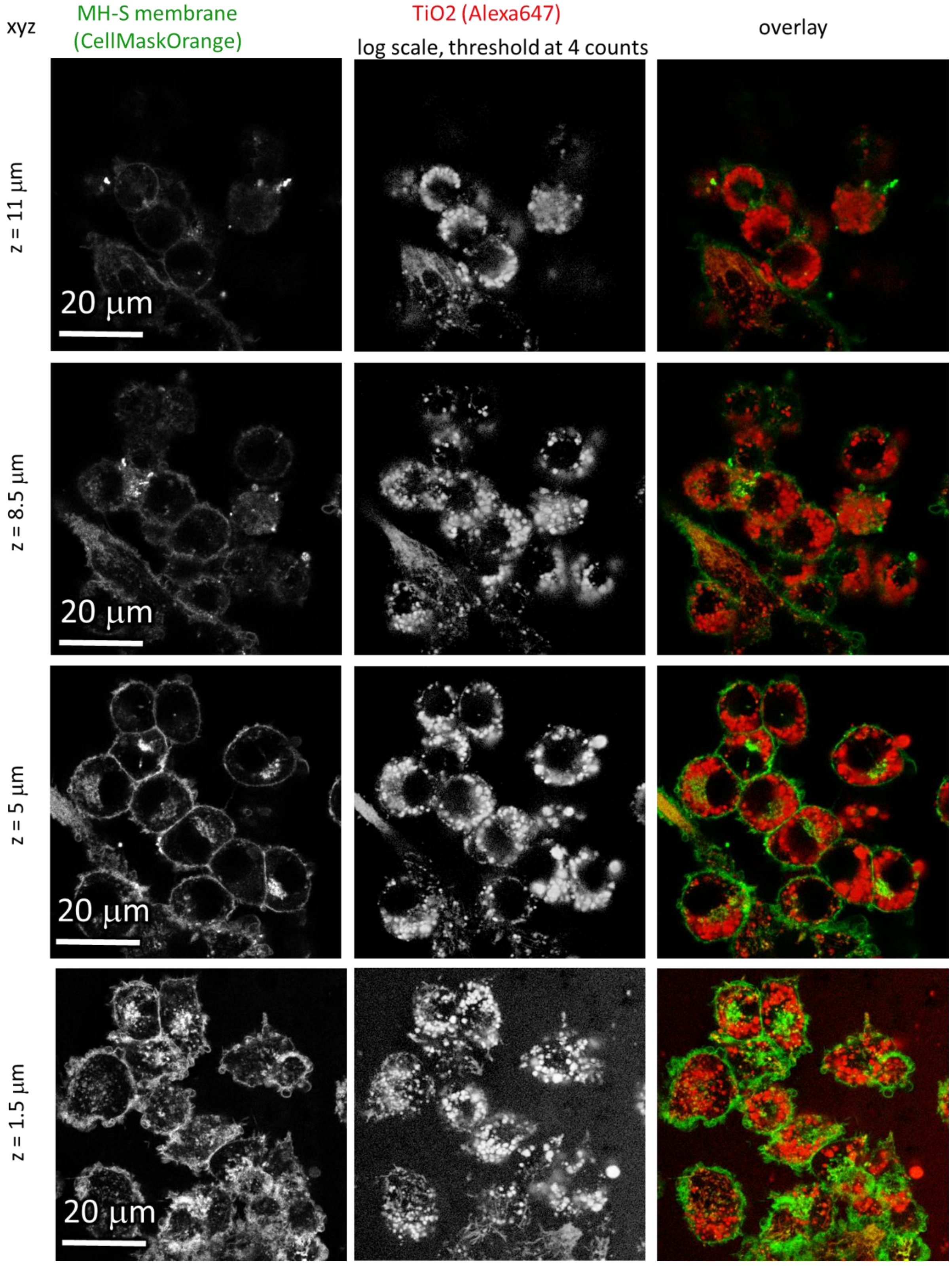
xyz
MH-S membrane
(CellMaskOrange)
TiO2 (Alexa647)
log scale, threshold at 4 counts
overlay
z = 11 µm
z = 8.5 µm
z = 5 µm
z = 1.5 µm
20 µm
20 µm
20 µm
20 µm

| | |
|---|---|
| Cell line | MH-S (membrane, CellMaskOrange) |
| NPs | TiO2 (Alexa647) |
| exposure | 10:1, 2 days |
| imaging | xyz confocal |

| | |
|---|---|
| pixelsize (x,y) | 150 nm |
| FOV (x,y) | 70.1 µm |
| pixelsize (z) | 500 nm |
| FOV (z) | 30 µm |
| imaging time | |
| number of frames | |

| | |
|---|---|
| 561nm | 5% |
| 640nm | 5% |
| STED | |
| filter sets | 605 nm – 625 nm, 650 nm – 720 nm |
| dwell time | 10 µs |
| objective | 60x wi (NA 1.2) |

| | |
|---|---|
| Cell line | MH-S (membrane, CellMaskOrange) |
| NPs | TiO2 (Alexa647) |
| exposure | 10:1, 2 days |
| imaging | xyt confocal |

| | |
|---|---|
| pixelsize (x,y) | 150 nm |
| FOV (x,y) | 70.1 µm |
| pixelsize (z) | |
| FOV (z) | |
| imaging time | 10 min |
| number of frames | 65 |

| | |
|---|---|
| 561nm | 5% |
| 640nm | 5% |
| STED | |
| filter sets | 605 nm – 625 nm, 650 nm – 720 nm |
| dwell time | 10 µs |
| objective | 60x wi (NA 1.2) |

*Figure S163: Macrophages are full on nanomaterial at 2 days of exposure*

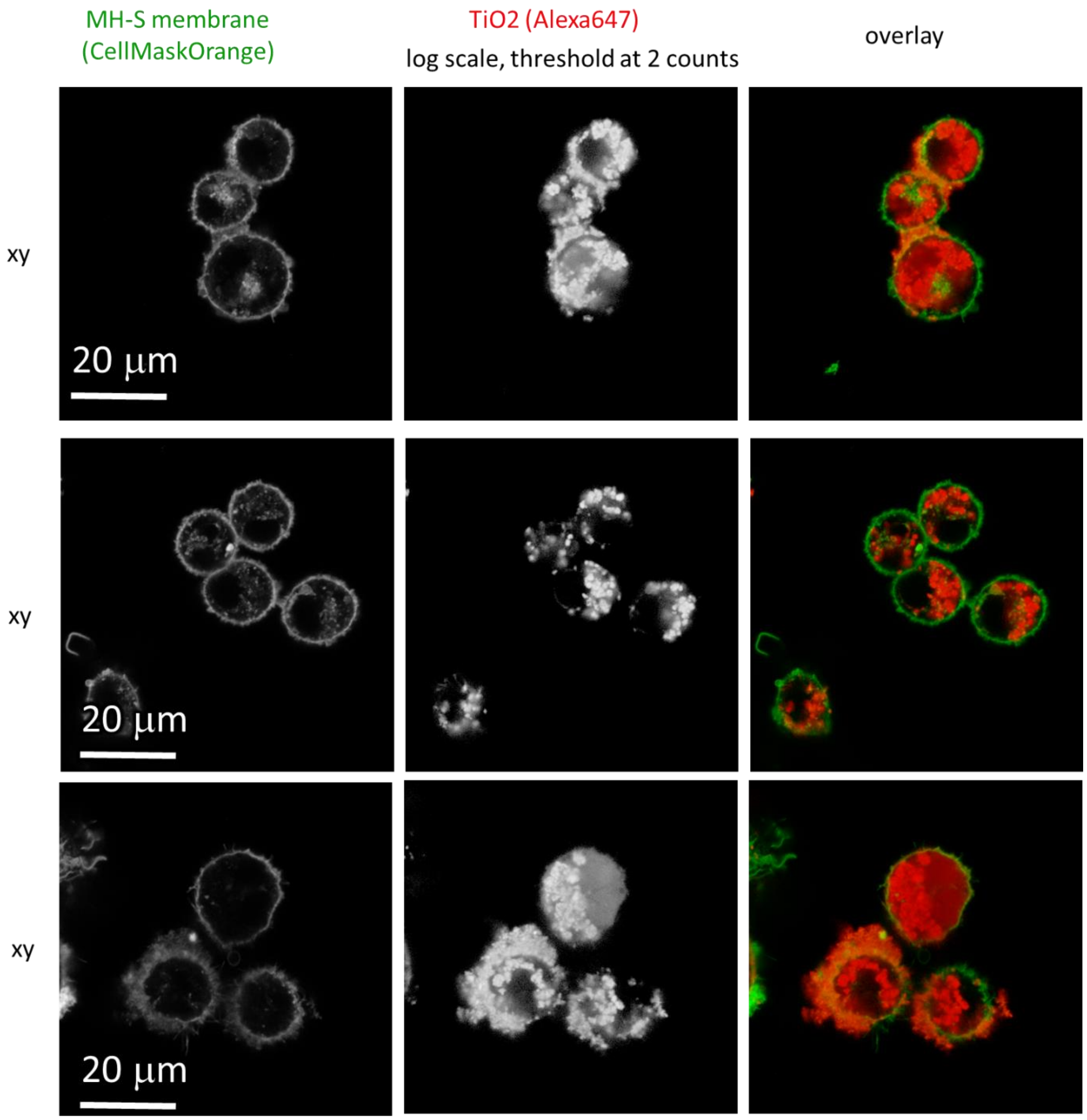

| | |
|---|---|
| Cell line | MH-S (membrane, CellMaskOrange) |
| NPs | TiO2 (Alexa647) |
| exposure | 10:1, 2 days |
| imaging | xy confocal |

| | |
|---|---|
| pixelsize (x,y) | 100 nm |
| FOV (x,y) | 70.0 µm |
| pixelsize (z) | |
| FOV (z) | |
| imaging time | |
| number of frames | |

| | |
|---|---|
| 561nm | 10% |
| 640nm | 10% |
| STED | |
| filter sets | 605 nm – 625 nm, 650 nm – 720 nm |
| dwell time | 10 µs |
| objective | 60x wi (NA 1.2) |

*Figure S164: Macrophages are full on nanomaterial at 2 days of exposure*

| MH-S membrane (CellMaskOrange) | TiO2 (Alexa647) log scale, threshold at 2 counts | overlay |
|---|---|---|

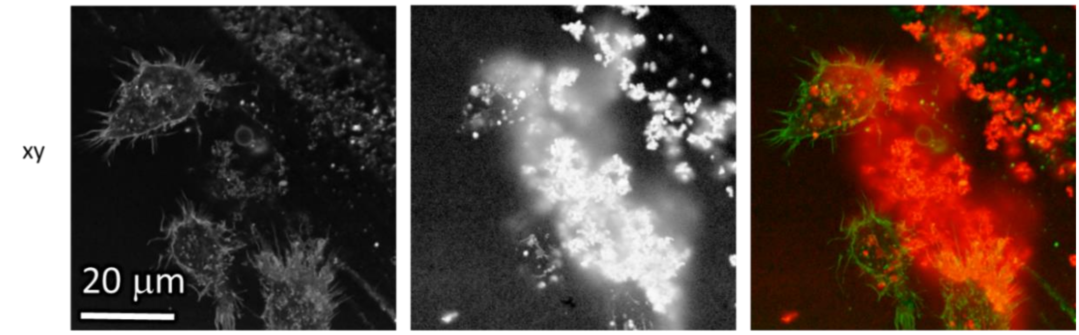

| Cell line | MH-S (membrane, CellMaskOrange) |
|---|---|
| NPs | TiO2 (Alexa647) |
| exposure | 10:1, 2 days |
| imaging | xy confocal |

| pixelsize (x,y) | 100 nm |
|---|---|
| FOV (x,y) | 70.0 µm |
| pixelsize (z) | |
| FOV (z) | |
| imaging time | |
| number of frames | |

| 561nm | 10% |
|---|---|
| 640nm | 100% |
| STED | |
| filter sets | 605 nm – 625 nm, 650 nm – 720 nm |
| dwell time | 10 µs |
| objective | 60x wi (NA 1.2) |

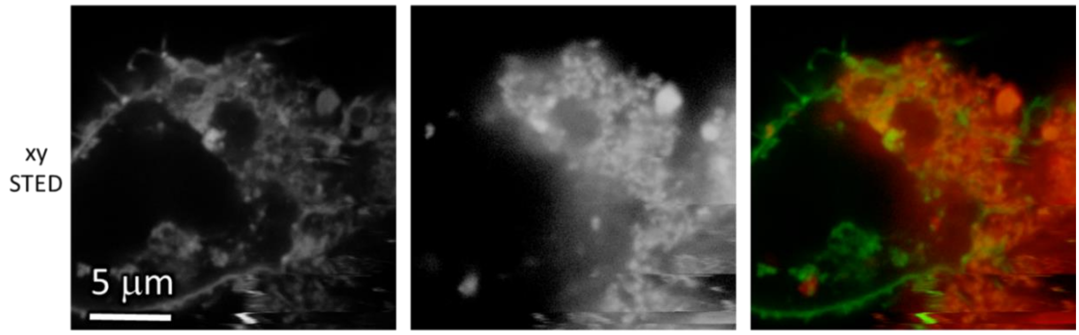

| Cell line | MH-S (membrane, CellMaskOrange) |
|---|---|
| NPs | TiO2 (Alexa647) |
| exposure | 10:1, 2 days |
| imaging | xy confocal |

| pixelsize (x,y) | 30 nm |
|---|---|
| FOV (x,y) | 20.0 µm |
| pixelsize (z) | |
| FOV (z) | |
| imaging time | |
| number of frames | |

| 561nm | 10% |
|---|---|
| 640nm | 10% |
| STED | 25%, 71% 3D STED |
| filter sets | 605 nm – 625 nm, 650 nm – 720 nm |
| dwell time | 10 µs |
| objective | 60x wi (NA 1.2) |

*Figure S165: Macrophages are full on nanomaterial at 2 days of exposure*

##### *Experiment at time 4 days*

- 20190225_e02 m03 s03_dying MH-S_SR DPPE and TiO2_ATTO 594_ over 48h_1 to 10

Macrophages exposed to different surface doses of $TiO_2$NTs.

Figure S166 1:1,

Figure S167-168 10:1 and

Figure S169 100:1 surface dose of $TiO_2$ nanotubes

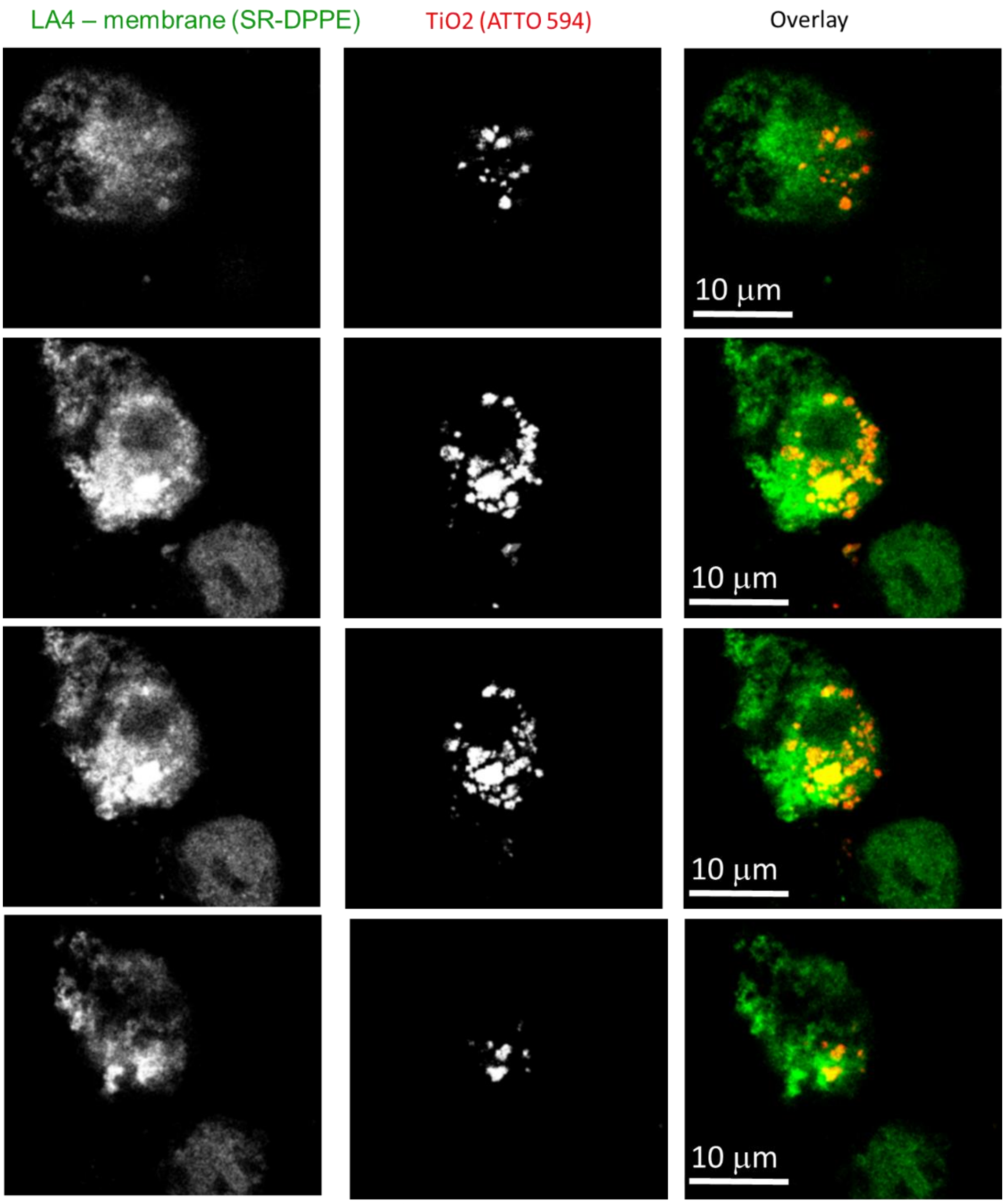

| | | | | | |
|---|---|---|---|---|---|
| Cell line | LA-4 (membrane, SR-DPPE) | pixelsize (x,y) | 150 nm | 561nm | 25% |
| NPs | TiO2_ATTO 594 | FOV (x,y) | 32x28 µm | 640nm | 40% |
| exposure | 1:10 | pixelsize (z) | 150 nm | STED | 23% |
| imaging | xyz STED, @72h | FOV (z) | 20 µm | filter sets | 605 nm – 625 nm, 400nm-780nm |
| | | imaging time | / | dwell time | 10 µs |
| | | number of stacks | 106 | objective | wi60x (NA1.2) |

*Figure S166: STED images of LA-4 in green and $TiO_2$ nanotubes in red channel. MH-S cells are labelled with Star Red – DPPE label for membrane. On figure are four representative XY plane slices from a whole Z-stack (3D). Macrophage (green) are devouring nanomaterial (red) which is seen as a colocalization of both signals in yellow color. When macrophages are full of nanomaterial we observe that they start to shring in size, dissintegrate and finally die. In this experiment surface dose of $TiO_2$ nanotubes is 10:1.*

- 20190225_e01 m04 s05_dying MH-S_SR DPPE and TiO2_ATTO 594_ over 48h_1 to 1

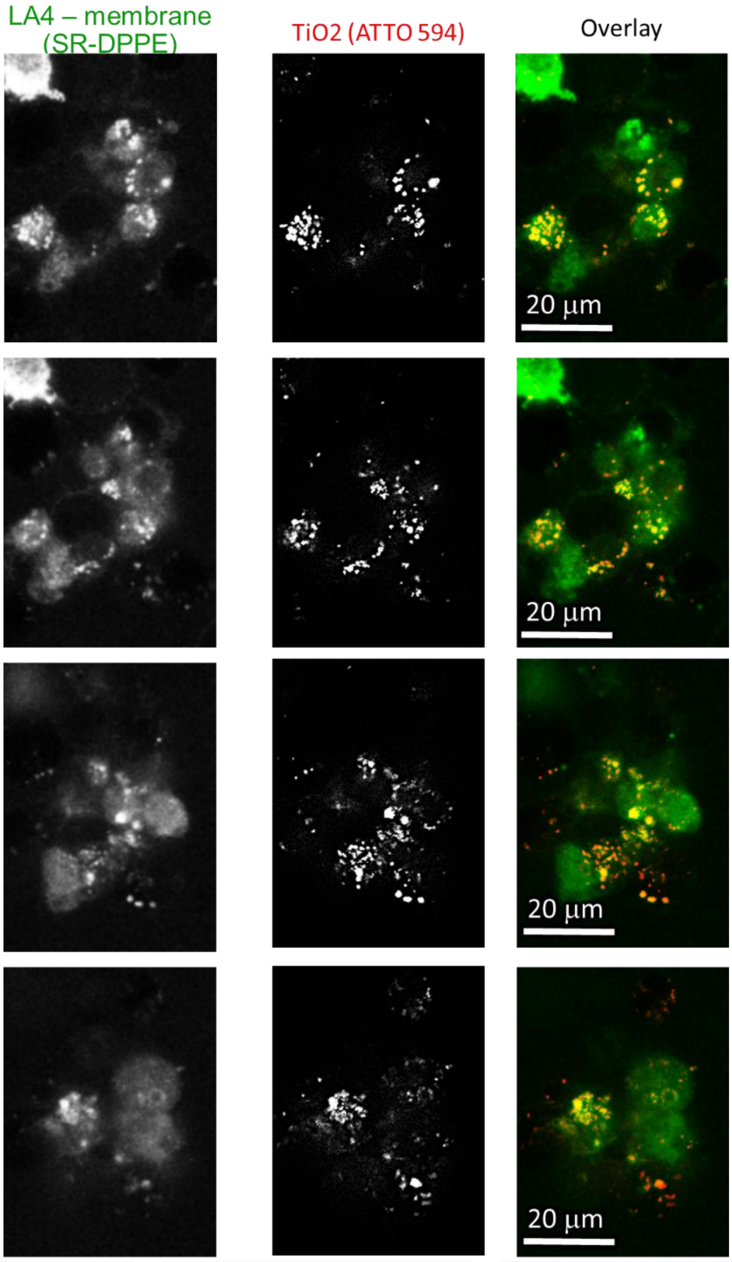

| | |
|---|---|
| Cell line | LA-4 (membrane, SR-DPPE) |
| NPs | TiO2_ATTO 594 |
| exposure | 1:1 |
| imaging | xzz STED, @72h |

| | |
|---|---|
| pixelsize (x,y) | 200 nm |
| FOV (x,y) | 47x65 μm |
| pixelsize (z) | 150 nm |
| FOV (z) | 21 μm |
| imaging time | / |
| number of stacks | 140 |

| | |
|---|---|
| 561nm | 25% |
| 640nm | 40% |
| STED | 23% |
| filter sets | 605 nm – 625 nm, 400nm-780nm |
| dwell time | 10 μs |
| objective | wi60x (NA1.2) |

*Figure S167: STED images of LA-4 in green and $TiO_2$ nanotubes in red channel. MH-S cells are labelled with Star Red – DPPE label for membrane. On figure are four representative XY plane slices from a whole Z-stack (3D). Macrophage (green) are devouring nanomaterial (red) which is seen as a colocalization of both signals in yellow color. When macrophages are full of nanomaterial we observe that they start to shring in size, dissintegrate and finally die. In this experiment surface dose of $TiO_2$ nanotubes is 1:1.*

- 20190225_e02 m02 s02_dying MH-S_SR DPPE and TiO2_ATTO 594_ over 48h_1 to 10

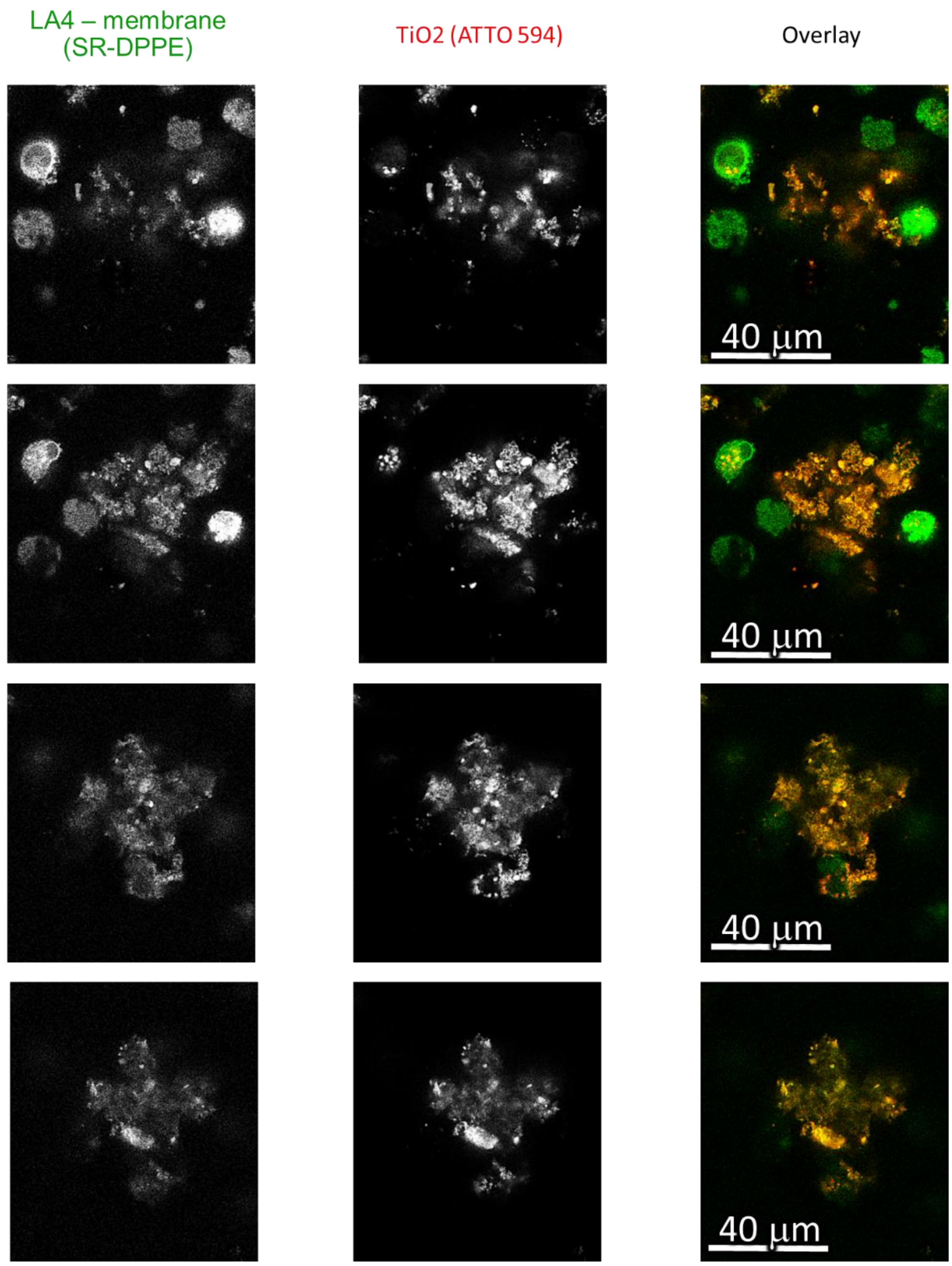

| | |
|---|---|
| Cell line | LA-4 (membrane, SR-DPPE) |
| NPs | TiO2_ATTO 594 |
| exposure | 1:10 |
| imaging | xzz STED, @72h |

| | |
|---|---|
| pixelsize (x,y) | 260 nm |
| FOV (x,y) | 80x90 µm |
| pixelsize (z) | 150 nm |
| FOV (z) | 25 µm |
| imaging time | / |
| number of stacks | 87 |

| | |
|---|---|
| 561nm | 25% |
| 640nm | 40% |
| STED | 23% |
| filter sets | 605 nm – 625 nm, 400nm-780nm |
| dwell time | 10 µs |
| objective | wi60x (NA1.2) |

*Figure S168: STED images of LA-4 in green and $TiO_2$ nanotubes in red channel. MH-S cells are labelled with Star Red – DPPE label for membrane. On figure are four representative XY plane slices from a whole Z-stack (3D). Macrophage (green) are devouring nanomaterial (red) which is seen as a colocalization of both signals in yellow color. When macrophages are full of nanomaterial we observe that they start to shring in size, dissintegrate and finally die. In this experiment surface dose of $TiO_2$ nanotubes is 10:1.*

- 20190225_e03 m01 s01_dying MH-S_SR DPPE and TiO2_ATTO 594_ over 48h_1 to 100 agglomerates of NP and 3D

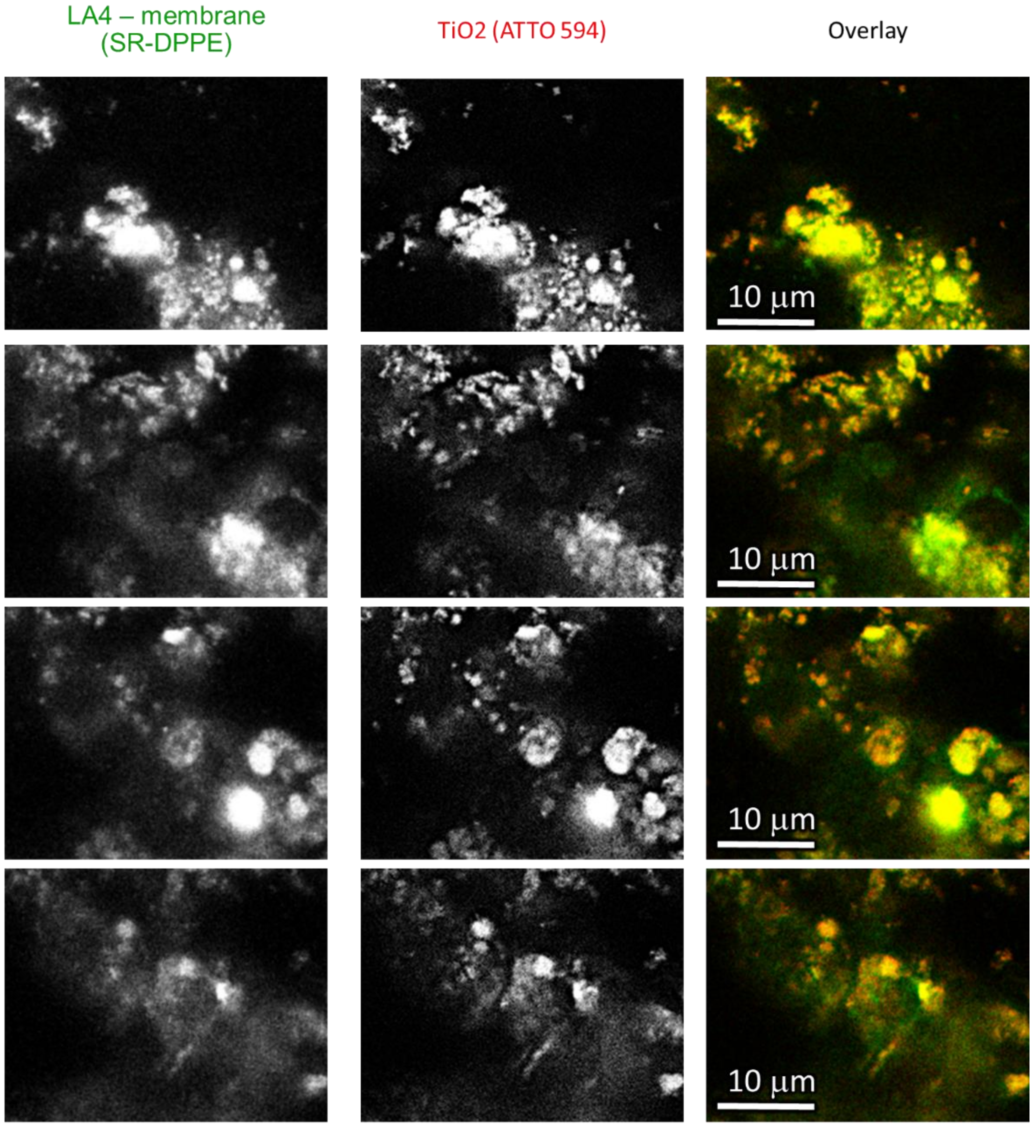

| | |
|---|---|
| Cell line | LA-4 (membrane, SR-DPPE) |
| NPs | TiO2_ATTO 594 |
| exposure | 1:100 |
| imaging | xzz STED, @72h |

| | |
|---|---|
| pixelsize (x,y) | 150 nm |
| FOV (x,y) | 33x25 µm |
| pixelsize (z) | 150 nm |
| FOV (z) | 25 µm |
| imaging time | / |
| number of stacks | 116 |

| | |
|---|---|
| 561nm | 25% |
| 640nm | 40% |
| STED | 23% |
| filter sets | 605 nm – 625 nm, 400nm-780nm |
| dwell time | 10 µs |
| objective | wi60x (NA1.2) |

*Figure S169: TED images of LA-4 in green and $TiO_2$ nanotubes in red channel. MH-S cells are labelled with Star Red – DPPE label for membrane. On figure are four representative XY plane slices from a whole Z-stack (3D). Macrophage (green) are devouring nanomaterial (red) which is seen as a colocalization of both signals in yellow color. When macrophages are full of nanomaterial we observe that they start to shring in size, dissintegrate and finally die. In this experiment surface dose of $TiO_2$ nanotubes is 1:100. Here macrophages are completely seald in a dense detritus of nanomaterial, live and dead macrophages resembling structures we've seen on the citoviva dark-field images (Fig.1a)..*

#### Link to 3D

- [http://lbfnanobiodatabase.ijs.si/file/data/cauliflowerpaper/20190225_e02_m03_s03_dying_MH-S_SR_DPPE_and_TiO2_ATTO_594__over_48h_1_to_10_b.mp4](http://lbfnanobiodatabase.ijs.si/file/data/cauliflowerpaper/20190225_e02_m03_s03_dying_MH-S_SR_DPPE_and_TiO2_ATTO_594__over_48h_1_to_10_b.mp4)

*Movie S43: Macrophage (green) are devouring $TiO_2$ nanotubes (red), surface dose 10:1.*

- http://lbfnanobiodatabase.ijs.si/file/data/cauliflowerpaper/20190225_dying_MHS_SR_DPPE_and_TiO2_ATTO_594__over_48h_1_to_1.gif

*Movie S44: Macrophage (green) are devouring $TiO_2$ nanotubes (red), surface dose 1:1.*

- http://lbfnanobiodatabase.ijs.si/file/data/cauliflowerpaper/20190225_e02_m02_s02_dying_MH-S_SR_DPPE_and_TiO2_ATTO_594__over_48h_1_to_10.gif

*Movie S45: Macrophage (green) are devouring $TiO_2$ nanotubes (red), surface dose 10:1.*

- http://lbfnanobiodatabase.ijs.si/file/data/cauliflowerpaper/20190225_e03_m01_s01_dying_MH-S_SR_DPPE_and_TiO2_ATTO_594__over_48h_1_to_100_agglomerats_of_NP_and_3D.gif

*Movie S46: Macrophage (green) are devouring $TiO_2$ nanotubes (red), surface dose 100:1.*

#### S4e – Macrophages attacking epithelial cells

##### Main message

Several macrophages surround and attacked an epithelial cell in the middle (discernible by its bigger size) which is full of nanomaterial.

Supporting raw and analysed data:

Figure S170

##### Materials and methods

- Main Experiment:
    - LA-4 cells were seeded @60% confluence in an Ibidi #1.5H µ-Dish
    - MH-S were seeded at 20% confluence in an Ibidi #1.5H µ-Dish
    - after 24 hours cultures were mixed in a cocultures and left in the incubator for another 24h
    - After 48h 35 µL freshly filtered 1 mg/mL $TiO_2$-40-ATTO 594 in 100x dcb was added directly to the cells (in 400 µL medium) and mixed to achieve 10:1 surface dose
    - Next day, cells were incubated with 1 um Star Red-DPPE for 10-15 minutes 2 hours in an incubator at 37°C, 5% CO2
    - afterwards they were washed with 1x400 µL LCIS and observed in 400 µL LCIS at microscope stage heated on 37°C

- Images analysed with ImageJ
    - Signal was multiplied and 0,5 pix Gaussian blur was applied Gaussian Blur 0.5 pix
    - Red channel has been analysed in the same way, the only difference being is the sharpening algorithm ($\sigma = 3$, $c = 0{,}5$) to more accentuate the nanomaterial

##### Experimental names

- Main Experiment:
    - 20190423_e01 m02 s02_LA-4 and MH-S_SR DPPE_1 and 10 to 1_ TiO_2 ATTO 594_ TL1
    - 20190423_e01 m02 s02_LA-4 and MH-S_SR DPPE_1 and 10 to 1_ TiO_2 ATTO 594_ TL2

##### Controls and statistics

Macrophages are attacking a LA-4 cell:

Figure S170

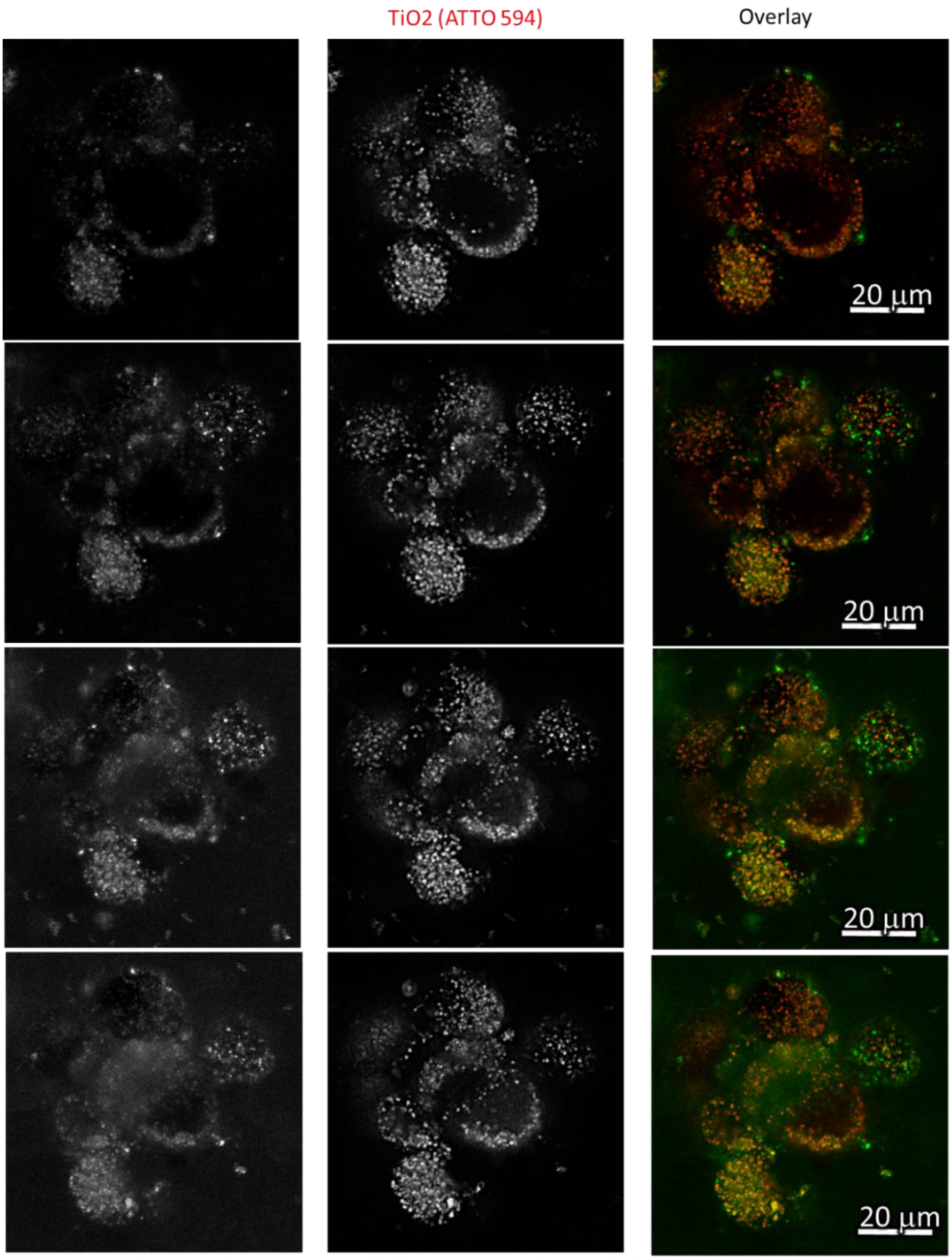

| | |
|---|---|
| Cell line | LA-4 & MHS(membrane, SR-DPPE) |
| NPs | TiO2 (ATTO 594) |
| exposure | 10:1, 48h-72h |
| imaging | xyt STED, 0h-1h |

| | |
|---|---|
| pixelsize (x,y) | 80 nm |
| FOV (x,y) | 55 um |
| pixelsize (z) | / nm |
| FOV (z) | / mm |
| imaging time | 2 hours |
| number of frames | 364 |

| | |
|---|---|
| 561nm | 6% |
| 640nm | 19% |
| STED | 20% |
| filter sets | 605 nm – 625 nm, 650 nm – 720 nm |
| dwell time | 10 ms |
| objective | wi60x (NA1.2) |

*Figure S170: STED images of LA-4 and MH-S in green and $TiO_2$ nanotubes in red channel. Cells are stained with Star Red – DPPE label for membrane. Four time points from a time lapse video. Five macrophages are surrunding and attacking one LA-4 cell, eventually braking the integrity on the LA-4 cell.*

Link to time-lapse

- *http://lbfnanobiodatabase.ijs.si/file/data/cauliflowerpaper/20190423_e01_m02_s02_LA-4_and_MH-S_SR_DPPE_1_and_10_to_1___TiO_2_ATTO_594___TL1_25_us.gif*

*Movie S47: Time-lapse video of five macrophages (green, Star Red DPPE) surrunding and attacking one LA-4 cell, eventually braking the integrity on the LA-4 cell.*

#### S4f – Macrophages attacking another macrophage

##### Main message

Macrophage is approaching and engulfing another macrophage which is completely filled with nanomaterial is already apoptotic and shrinking. Event takes place on the surface of the LA-4 cell.

Supporting raw and analysed data:

Figure S171

##### Materials and methods

- Main experiment:
  - LA-4 cells were seeded @30% confluence in an Ibidi #1.5H µ-Dish
  - MH-S cells were seeded @30% confluence in a separate Ibidi #1.5H µ-Dish
  - After 24 hours of separate incubation, LA-4 were labelled with 0,1 µM SirActin and MH-S were labelled with <1 µM cell mask and mixed together. Growth media for cocultures was mixture of F12K and RPMI-1640 in 1:1 ratio
  - after 48 hours 35 mL freshly filtered 1 mg/mL $TiO_2$-40-ATTO 594 in 100x dcb was added directly to the cells (in 400 µL medium) and mixed to achieve 10:1 surface dose and incubated overnight in the incubator at 37°C and 5% CO2
  - Next day cells were flushed with 1x400 µL of LCIS and imaged on heated microscope stage in LCIS (37°C)

- Analysis was performed in the ImageJ software:
  - Signal was multiplied in order to get adequate visibility for 3D rendering. Signal of stacks of images containing MH-S was multiplied 2x more because signal intensity was not the same. MH-S and LA-4 have been labeled with different labels and they did not yield the same number of counts on our detectors.
  - Additionaly 1 pixel Gaussian Blur was applied in a green channel and sharpening algorithm ($\sigma$ = 2.5, c = 0.6) in the red channel

##### Experimental names

- Main experiment:
  - 20190204_LA-4_SA and MH-S_CM coculture_10 to 1 TiO2_ATTO 594_MH-S attacking another MH-S_3D

##### Controls and statistics

Macrophage devouring another macrophage, completely full of nanomaterial, on the top pf the LA-4 cell:

Figure S171

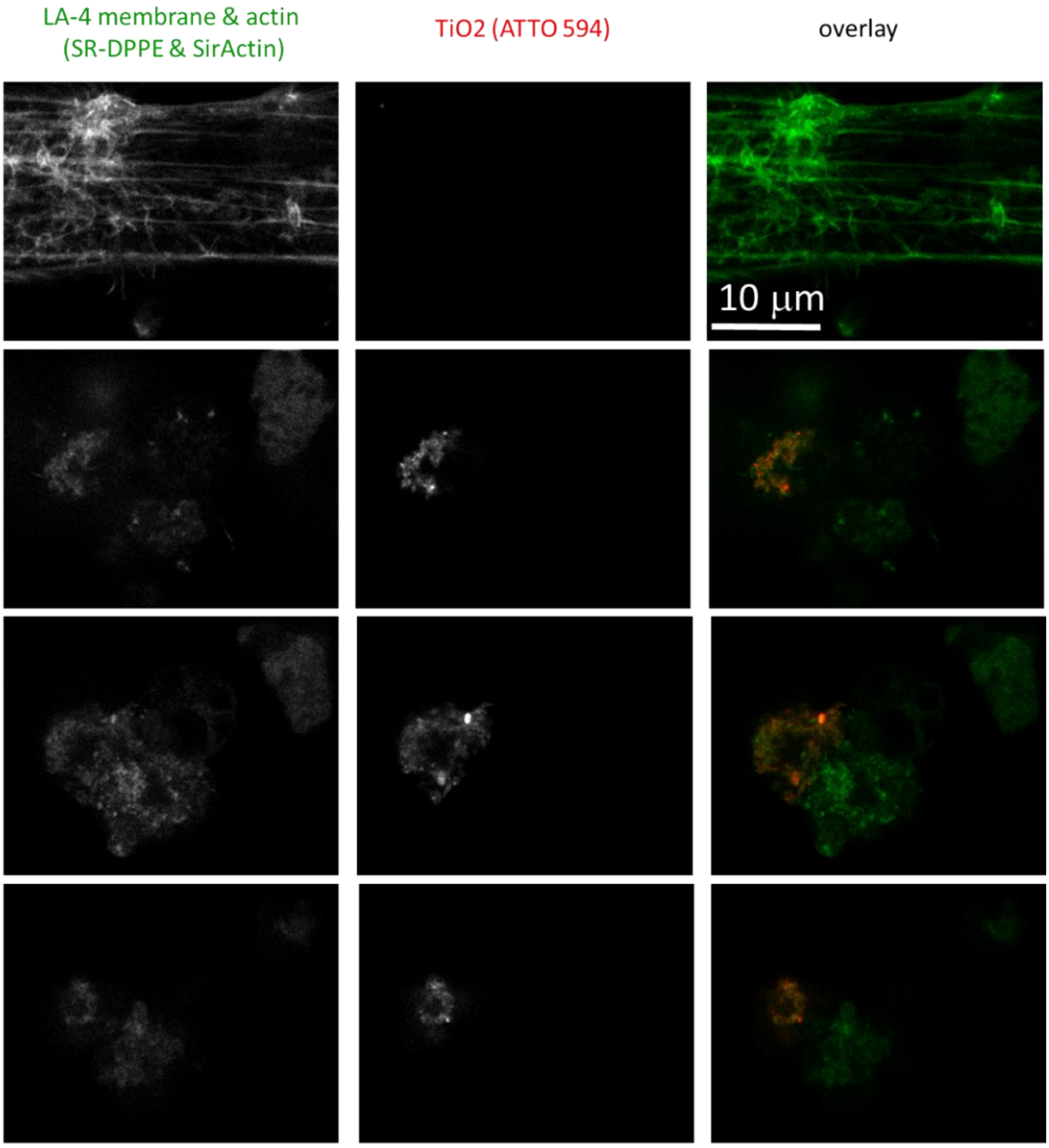

| | |
|---|---|
| Cell line | LA-4 (actin, Sir Actin) & MHS (membrane, CellMaskOrange) |
| NPs | TiO2 (ATTO 594) |
| exposure | 10:1, 48h-55h |
| imaging | xyt STED, 55h |

| | |
|---|---|
| pixelsize (x,y) | 100 nm |
| FOV (x,y) | 31 x 24 µm |
| pixelsize (z) | 120 nm |
| FOV (z) | 12 µm |
| imaging time | |
| number of z-sections | 99 |

| | |
|---|---|
| 561nm | 15% |
| 640nm | 11% |
| STED | 9% |
| filter sets | 605 nm – 625 nm, 650 nm – 720 nm |
| dwell time | 10 µs |
| objective | 60wix (NA1.2) |

*Figure S171: STED images of LA-4 and MH-S in green and $TiO_2$ nanotubes in red channel. LA-4 cells are stained with Sir-Actin (green cytoskeleton filaments) and MH-S are labelled with Star Red – DPPE label for membrane. On figure are four representative XY plane slices from a whole Z-stack (3D). One macrophage (green) on top of the LA-4 cell (green filaments) is devouring another macrophage (green, red, yellow colocalization) which is full of nanomaterial and possibly dying.*

Link to 3D

- http://lbfnanobiodatabase.ijs.si/file/data/cauliflowerpaper/20190204_LA-4_SA_and_MH-S_CM_coculture_10_to_1_TiO2_ATTO_594_MH-S_attacking_another_MH-S_3D.mp4

*Movie S48: Macrophage (CellMask Orange, green) attacking another macrophage full of $TiO_2$ nanotubes (Atto 594, red) on top of epithelial cell.*

### S5 – Towards predictive toxicology

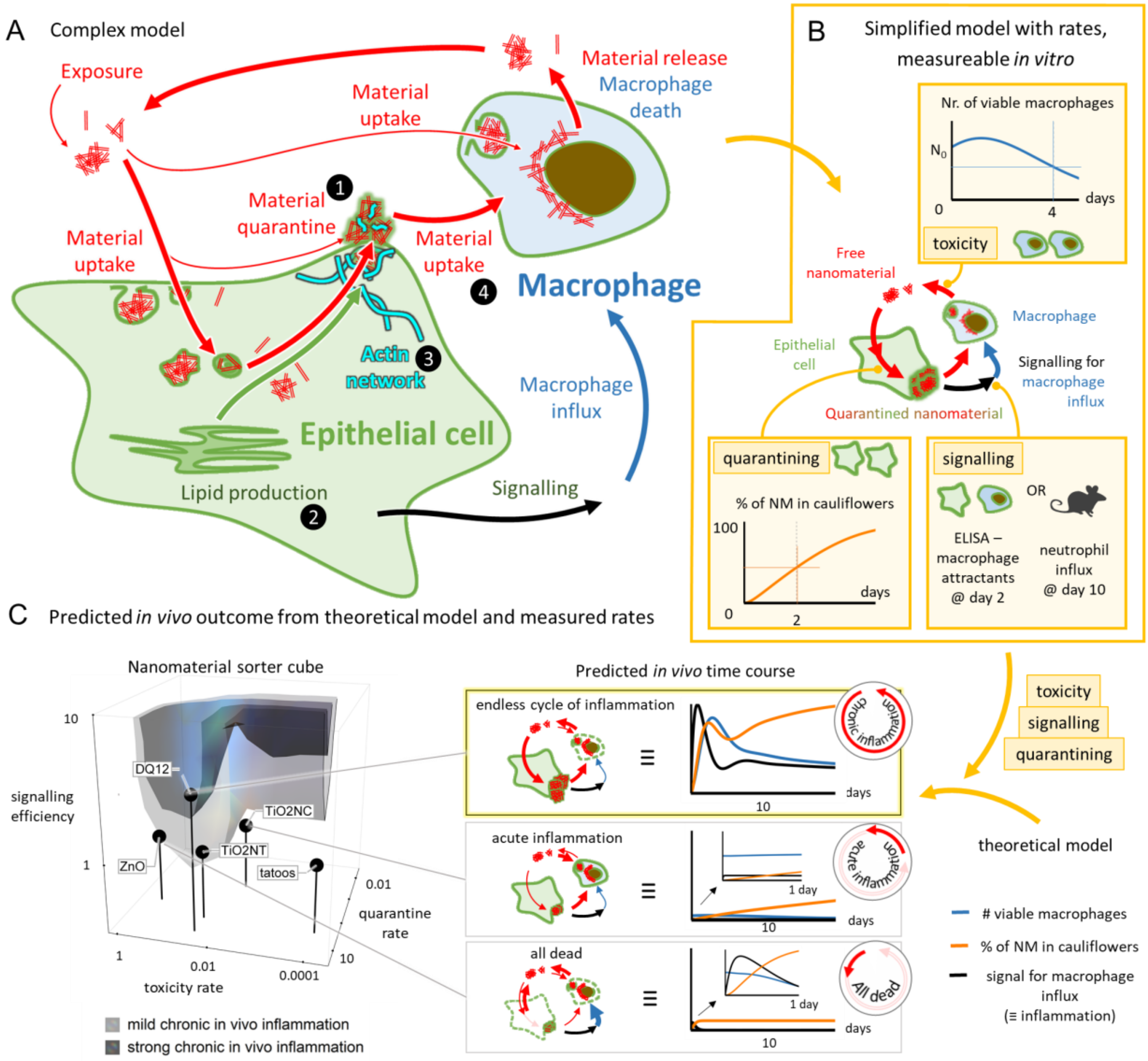

#### Main message

Based on the complex scheme in Figure5a, we designed a theoretical model of the events following exposure to nanomaterial with the least complexity possible (S5b). The rates and parameters in the model can be measured *in vivo* or in *in vitro* systems for any desired nanomaterial (S5b). Based on the time evolution of the theoretical model, outcomes such as chronic and acute inflammation may be predicted for a wide variety of nanomaterials (S5c).

#### S5b – Model of chronic inflammation following nanomaterial exposure and determination of its parameters

##### Model of chronic inflammation following nanomaterial exposure

###### Main message

The theoretical model of chronic inflammation following nanomaterial exposure is described by a series of differential equations, describing the events observed in in vitro and in vivo experiments in this work. This minimal-complexity in vivo model consists of 6 variables (surface of nanomaterial in epithelial cells *npLA4*, in cauliflowers *npCF*, in macrophages *npMHS* and freely-floating nanomaterial *npFree*, surface of macrophages *sMHS* and surface of epithelial cells *sLA4*), 4 fixed parameters which are calibrated for each model system and later locked (endocytosis rate *endo*, rate of cauliflower endocytosis *endo*cfuEff*, delay between cauliflower production and signalling for macrophage influx *delay*, and epithelial cell replication rate *LA4Rep*) and 3 nanomaterial-associated parameters (cauliflower formation rate *cff*, signalling efficiency *signEff*, and toxicity *tox*). Separate *in vitro* models were obtained from the *in vivo* model by swapping the macrophage influx with macrophage replication and leaving out non-existent cells for monocultures.
The system of equations was solved numerically using Wolfram Mathematica 12.0, licence L5063-5112 to obtain the time evolution of the model. The same software was also used for visualization of the results.

Supporting material:

Figure S172 - Figure S175

Rate equations and dynamic evolution of *in vivo* system and *in vitro* monocultures and cocultures:

###### *In vivo model*

rate equations:

```
D[npFree[t], t] == -endo sLa4[t] * npFree[t] - endo sMhs[t] * npFree[t] + tox npMhs[t] * npMhs[t] / sMhs[t] + tox npLa4[t] * npLa4[t] / sLa4[t]
```

| change of free NM | free NM endocytosed by epithelial cells | free NM endocytosed by macrophages | released from dead macrophages | released from dead epithelial cells |
|---|---|---|---|---|

```
D[npLa4[t], t] == endo sLa4[t] * npFree[t] - cff sLa4[t] Tanh[npLa4[t] / (sLa4[t] * cff * timecf)] - tox npLa4[t] * npLa4[t] / sLa4[t]
```

| change of NM in epithelial cells | free NM endocytosed by epithelial cells | excreted into cauliflowers by epithelial cells | released from dead epithelial cells |
|---|---|---|---|

```
D[npCF[t], t] == cff sLa4[t] Tanh[npLa4[t] / (sLa4[t] * cff * timecf)] - endo * cfuEff sMhs[t] * npCF[t]
```

| change of NM in cauliflowers | excreted into cauliflowers by epithelial cells | cauliflowers endocytosed by macrophages |
|---|---|---|

```
D[npMhs[t], t] == endo sMhs[t] * npFree[t] + endo * cfuEff sMhs[t] * npCF[t] - tox npMhs[t] * npMhs[t] / sMhs[t]
```

| change of NM in macrophages | free NM endocytosed by macrophages | cauliflowers endocytosed by macrophages | released from dead macrophages |
|---|---|---|---|

```
D[sMhs[t], t] == -tox npMhs [t] + signalEff cff sLa4[t - delay] Tanh[npLa4[t - delay] / (sLa4[t - delay] * cff * timecf)] * macrophageSurface
```

| change of number of macrophages | dying macrophages | influx of new macrophages, signalled by cauliflower production (increased lipid production) in epithelial cells at an earlier time |
|---|---|---|

```
D[sLa4[t], t] == -tox npLa4[t] + La4Rep sLa4[t] (1 - sLa4[t] / la4Max)
```

| change of number of epithelial cells | dying epithelial cells | replication of epithelial cells |
|---|---|---|

initial conditions:

```
npFree[0] = exposureSurfaceDose = 10;

npLa4[0] == 0,  npLa4[t /; t < 0] == 0,

npCF[0] == 0,

npMhs[0] == 0,

sMhs[0] == macrophageSurface = N[1 / 40];

sLa4[0] == 1    sLa4[t /; t < 0] == 1,

la4Max = 1;
```

At t = 0, we add nanomaterial to fully confluent epithelial cells and macrophages. The ratio

surface of nanomaterial : surface of epithelial cells : surface of macrophages = 10 : 1 : 1/40

variables:

npFree[t] : surface of free-floating nanomaterial, normed to the surface of epithelial cells at t = 0
npLa4[t] : total surface of nanomaterial inside epithelial cells, normed to the surface of epithelial cells at t = 0
npCF[t] : surface of nanomaterial in cauliflowers, normed to the surface of epithelial cells at t = 0
npMhs[t] : total surface of nanomaterial inside macrophages, normed to the surface of epithelial cells at t = 0
sMhs[t] : surface of macrophages, normed to the surface of epithelial cells at t = 0
sLa4[t] : surface of epithelial cells, normed to the surface of epithelial cells at t = 0
t : time, measured in days

*parameters – rate coefficients*

*endo* : rate of endocytosis = 1
***tox*** : toxicity of nanomaterial
***cff*** : rate of cauliflower formation
*timecf* : norming factor for exocytosis saturation = 1
*cfuEff* : rate of cauliflower endocytosis compared to endocytosis of free NM = 1
***signalEff*** : signalling efficiency – how many macrophages are infiltrated by the released signal
*delay* : delay between signal excretion and macrophage infiltration = 0.1
*La4Rep* : replication rate of epithelial cells = 0.3

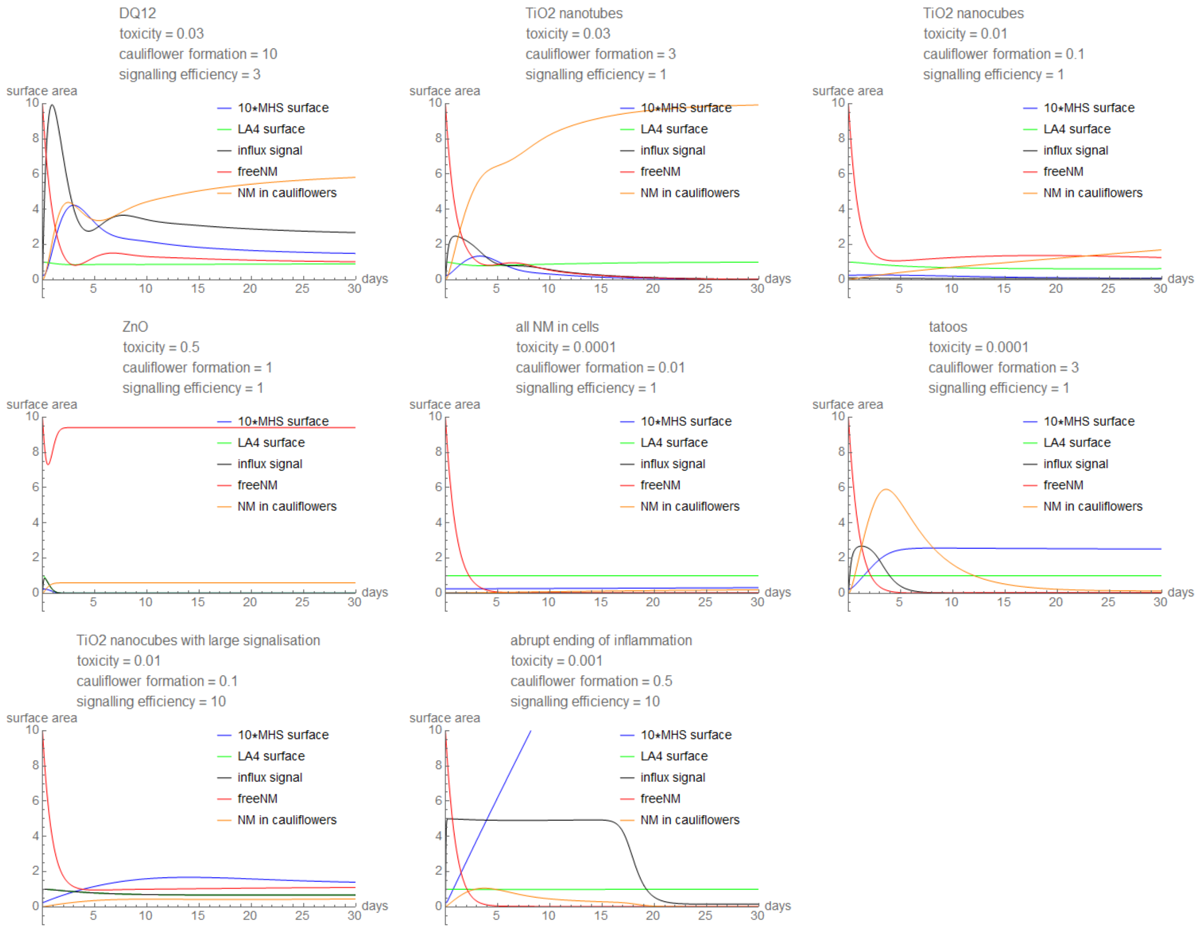

*Figure S172: Time evolution of the in vivo theoretical model for 8 different nanomaterials.*

##### *In vitro coculture model*

For the descriptions of the terms see the *in vivo* model - the rate equations are similar to the *in vivo* model, with some terms being left out and MH-S influx is swapped with MH-S replication.

rate equations:

```
enacbe[{endo_, tox_, cff_, timecf_, cfuEff_, delay_, signalEff_(*signal efficiancy*), La4Rep_(*replication rate of LA4*)}] :=
 {D[npFree[t], t] == -endo sLa4[t] * npFree[t] - endo sMhs[t] * npFree[t] + tox npMhs[t] * npMhs[t] / sMhs[t] + tox npLa4[t] * npLa4[t] / sLa4[t],

  D[npLa4[t], t] == endo sLa4[t] * npFree[t] - cff sLa4[t] Tanh[npLa4[t] / (sLa4[t] * cff * timecf)] - tox npLa4[t] * npLa4[t] / sLa4[t],

  D[npCF[t], t] == cff sLa4[t] Tanh[npLa4[t] / (sLa4[t] * cff * timecf)] - endo * cfuEff sMhs[t] * npCF[t] ,

  D[npMhs[t], t] == endo sMhs[t] * npFree[t] + endo * cfuEff sMhs[t] * npCF[t] - tox npMhs[t] * npMhs[t] / sMhs[t],

  D[sMhs[t], t] == -tox npMhs [t] + La4Rep sMhs[t] (1 - sMhs[t] / la4Max),

  D[sLa4[t], t] == -tox npLa4[t] + La4Rep sLa4[t] (1 - sLa4[t] / la4Max),

  sLa4[t /; t < 0] == 1,
  npLa4[t /; t < 0] == 0,
```

initial conditions:

```
npFree[0] = exposureSurfaceDose = 10;

npLa4[0] == 0,  npLa4[t /; t < 0] == 0,

npCF[0] == 0,

npMhs[0] == 0,

sMhs[0] == macrophageSurface = N[1 / 40];

sLa4[0] == 1    sLa4[t /; t < 0] == 1,

la4Max = 1;
```

At t = 0, we add nanomaterial to fully confluent epithelial cells and macrophages. The ratio

surface of nanomaterial : surface of epithelial cells : surface of macrophages = 10 : 1 : 1/40

variables:

npFree[t] : surface of free-floating nanomaterial, normed to the surface of epithelial cells at t = 0
npLa4[t] : total surface of nanomaterial inside epithelial cells, normed to the surface of epithelial cells at t = 0
npCF[t] : surface of nanomaterial in cauliflowers, normed to the surface of epithelial cells at t = 0
npMhs[t] : total surface of nanomaterial inside macrophages, normed to the surface of epithelial cells at t = 0
sMhs[t] : surface of macrophages, normed to the surface of epithelial cells at t = 0
sLa4[t] : surface of epithelial cells, normed to the surface of epithelial cells at t = 0
t : time, measured in days

*parameters – rate coefficients*

*endo* : rate of endocytosis = 1
***tox*** : toxicity of nanomaterial
***cff*** : rate of cauliflower formation
*timecf* : norming factor for exocytosis saturation = 1
*cfuEff* : rate of cauliflower endocytosis compared to endocytosis of free NM = 1
***signalEff*** : signalling efficiency – how many macrophages are infiltrated by the released signal
*delay* : delay between signal excretion and macrophage infiltration = 0.1
*La4Rep* : replication rate of epithelial cells = 0.3

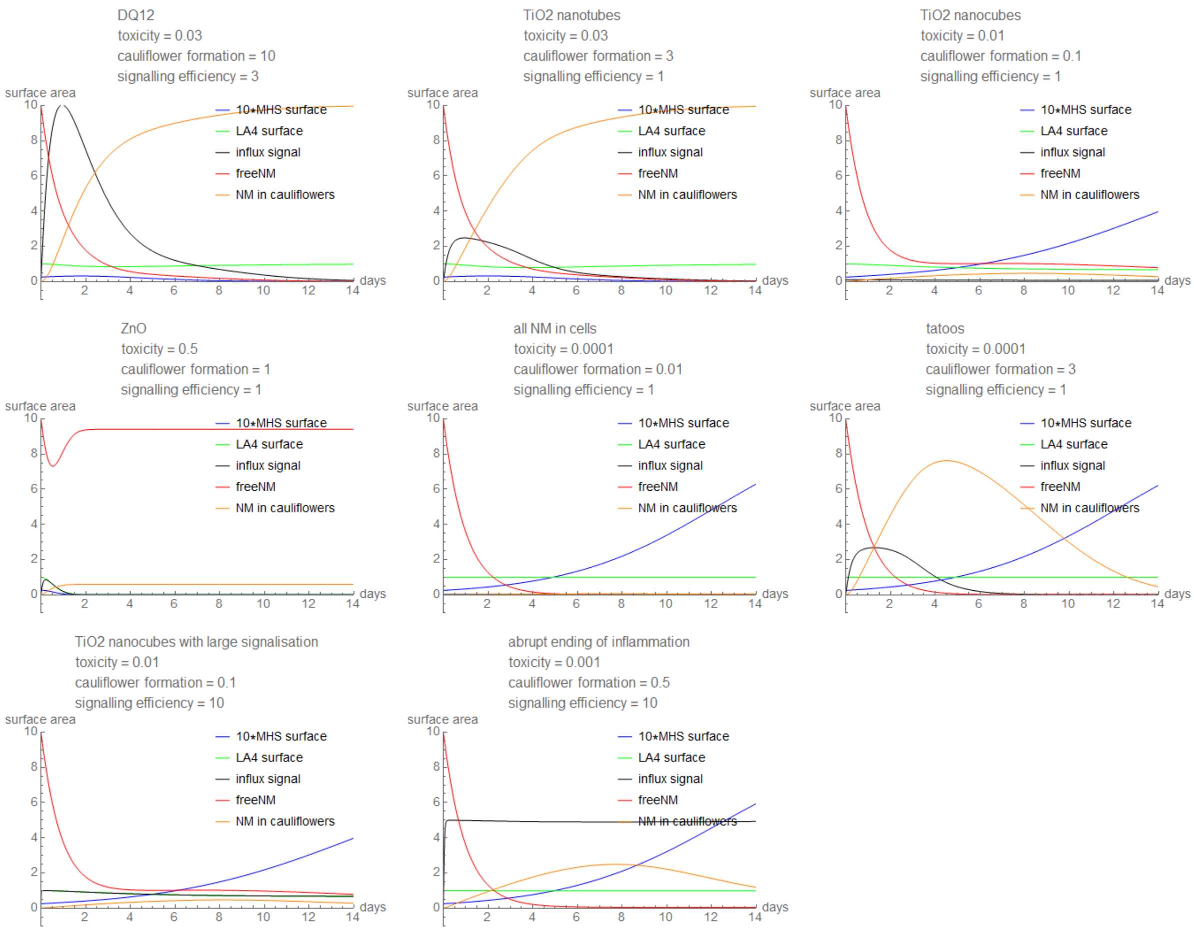

*Figure S173: Time evolution of the in vitro coculture theoretical model for 8 different nanomaterials.*

#### *In vitro LA-4 monoculture model*

For the descriptions of the terms see the *in vivo* model - the rate equations for LA-4 monoculture are similar to the *in vivo* model, with some terms being left out.

rate equations:

```
D[npFree[t], t] == -endo sLa4[t] * npFree[t] + tox npLa4[t] * npLa4[t] / sLa4[t],

D[npLa4[t], t] == endo sLa4[t] * npFree[t] - cff sLa4[t] Tanh[npLa4[t] / (sLa4[t] * cff * timecf)] - tox npLa4[t] * npLa4[t] / sLa4[t],

D[npCF[t], t] == cff sLa4[t] Tanh[npLa4[t] / (sLa4[t] * cff * timecf)] ,

D[sLa4[t], t] == -tox npLa4[t] + la4Rep sLa4[t] (1 - sLa4[t] / la4Max),
```

initial conditions:

```
npFree[0] == exposureSurfaceDose = 10;

npLa4[0] == 0,

npCF[0] == 0,

sLa4[0] == 1

la4Max = 1;
```

At t = 0, we add nanomaterial to fully confluent epithelial cells. The ratio

surface of nanomaterial : surface of epithelial cells = 10 : 1 : 1/40 variables:

npFree[t] : surface of free-floating nanomaterial, normed to the surface of epithelial cells at t = 0
npLa4[t] : total surface of nanomaterial inside epithelial cells, normed to the surface of epithelial cells at t = 0
npCF[t] : surface of nanomaterial in cauliflowers, normed to the surface of epithelial cells at t = 0
sLa4[t] : surface of epithelial cells, normed to the surface of epithelial cells at t = 0
t : time, measured in days

*parameters – rate coefficients*

*endo* : rate of endocytosis = 1
*tox* : toxicity of nanomaterial
*cff* : rate of cauliflower formation
*timecf* : norming factor for exocytosis saturation = 1
*La4Rep* : replication rate of epithelial cells = 0.3

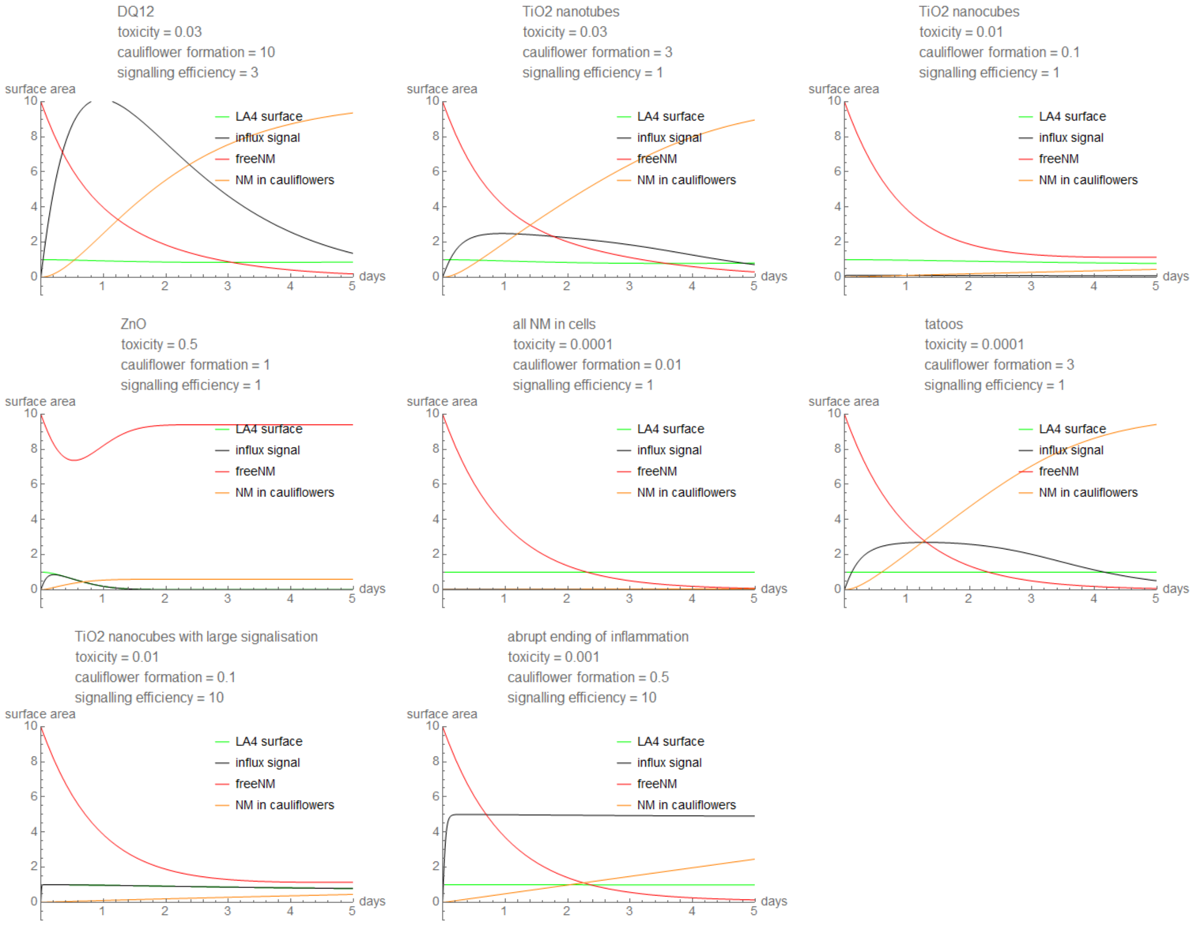

*Figure S174: Time evolution of the in vitro LA-4 monoculture theoretical model for 8 different nanomaterials.*

#### *In vitro MH-S monoculture model*

For the descriptions of the terms see the *in vivo* model - the rate equations for MH-S monoculture are similar to the *in vivo* model, with some terms being left out.

rate equations:

```
D[npFree[t], t] == -endo sMhs[t] * npFree[t] + tox npMhs[t] * npMhs[t] / sMhs[t],

D[npMhs[t], t] == endo sMhs[t] * npFree[t] - tox npMhs[t] * npMhs[t] / sMhs[t],

D[sMhs[t], t] == -tox npMhs [t] + La4Rep sMhs[t] (1 - sMhs[t] / la4Max),
```

initial conditions:

```
npFree[0] == exposureSurfaceDose = 10;

npMhs[0] == 0,

sMhs[0] == macrophageSurface = N[1/40];

la4Max = 1;
```

At t = 0, we add nanomaterial to macrophages. The ratio

surface of nanomaterial : surface of the dish: surface of macrophages = 10 : 1 : 1/40

variables:

npFree[t] : surface of free-floating nanomaterial, normed to the surface of epithelial cells at t = 0
npMhs[t] : total surface of nanomaterial inside macrophages, normed to the surface of epithelial cells at t = 0
sMhs[t] : surface of macrophages, normed to the surface of epithelial cells at t = 0
t : time, measured in days

*parameters – rate coefficients*

*endo* : rate of endocytosis = 1
*tox* : toxicity of nanomaterial
*La4Rep* : replication rate of epithelial cells = 0.3

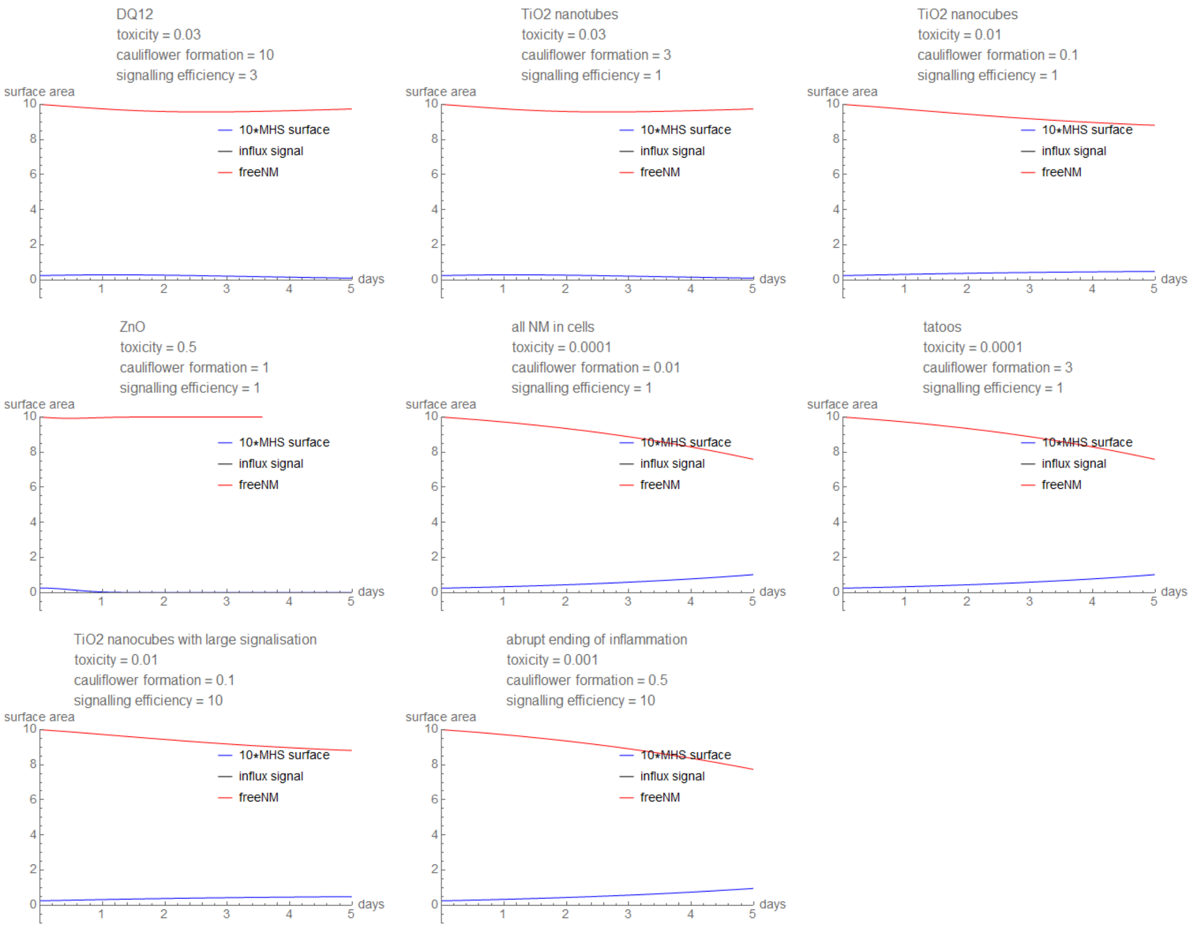

*Figure S175: Time evolution of the in vitro MH-S monoculture theoretical model for 8 different nanomaterials.*

#### Determination of rates for the theoretical model from *in vitro* assay results

##### Main message

In order to determine the nanomaterial-specific model parameters toxicity rate (*tox*), quarantine rate (*cff*) and signalling efficiency (*signalEff*) from the model in S5b for a desired nanomaterial, one must first determine the toxicity of the nanomaterial (*tox*) from real-life *in vitro* MH-S monoculture viability as shown below. By combining this with the observed magnitude of cauliflowers in a real-life *in vitro* LA-4 monoculture, the cauliflower formation rate (*cff*) may be determined as shown below. Since the amount of cauliflowers depends both on the toxicity of the nanomaterial as well as the exocytosis process itself, the *cff* cannot be measured directly. From there on, efficiency of signaling and monocyte influx replacing the dying macrophages *(signalEff)* is calculated from the rates *tox*, *cff* and either via measured polymorphonuclear cell influx *in vivo* after 10 days or measured macrophage attractants in *in vitro* co-culture of LA-4 and MH-S after 2 days (pre-calibrated with *in vivo*).
All analysis and visualisation in this section were done in Wolfram Mathematica 12.0, licence L5063-5112.

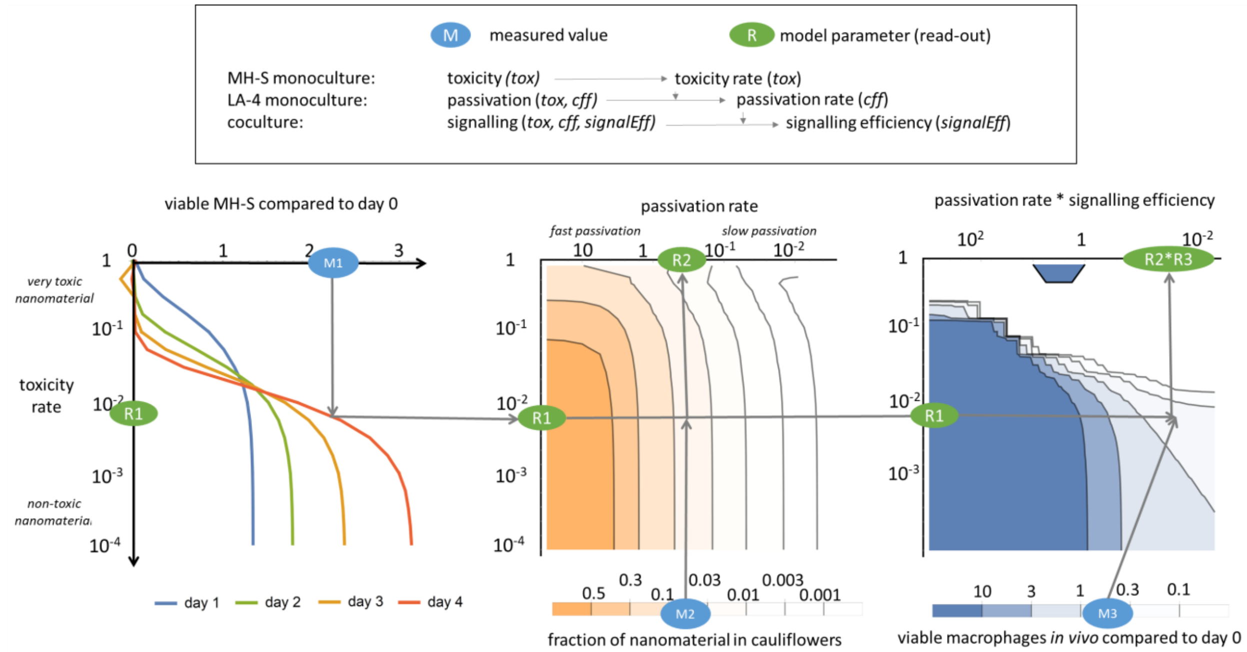

*Figure S176: Schematic of workflow and determination of parameters of the model.*

#### S5c – Phase space of chronic inflammation

##### Main message

The phase space was scanned by calculating the time evolution of the appropriate system of equations from section S5b for a set of nanomaterials with appropriately interspaced parameters: toxicity rate (*tox*), quarantine rate (*cff*) and signalling efficiency (*signalEff*). For each parameter, 30 logarithmically-equally-spaced values in a sensible range were chosen – the total amount of values in the grid was thus 30 x 30 x 30 = 27.000. The parameter values for plotted nanomaterials were approximated from the known behaviour of the *in vivo* and *in vitro* systems following exposure to these nanomaterials.
For three chosen nanomaterials, the time-courses were simulated.
All the analysis in this section was done in Wolfram Mathematica 12.0, licence L5063-5112.
More information can be found in S5b.

Supporting data:

Figure S177

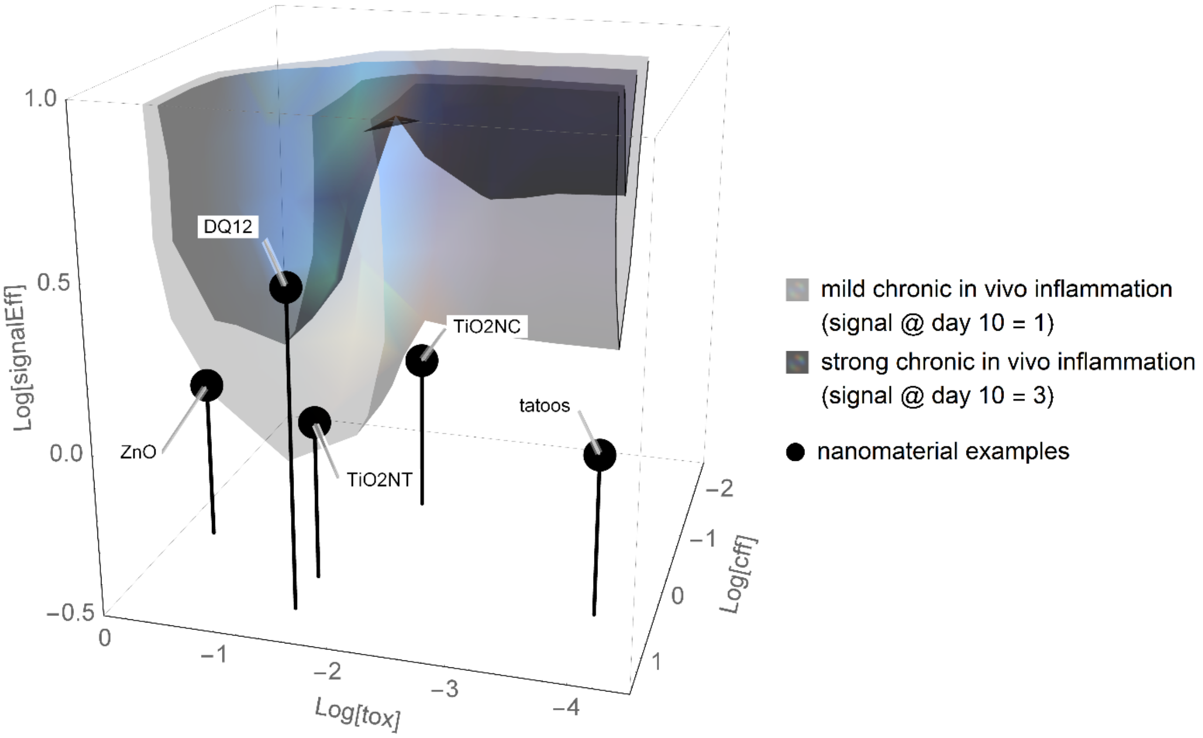

*Figure S177: The black contour in the 3D parameter phase space plot represents the set of parameters with the predicted influx signal in vivo at day 10 equal to 3 (strong chronic inflammation) and the grey contour with the predicted influx signal in vivo at day 10 equal to 1 (weak chronic inflammation) (black line on Figure S172). Black spheres depict the estimated location of the nanomaterials in the cube.*
